# Distinguishable topological properties of functional genome networks in HIV-1 reservoirs

**DOI:** 10.1101/2024.02.05.578936

**Authors:** Janusz Wiśniewski, Kamil Więcek, Haider Ali, Krzysztof Pyrc, Anna Kula-Păcurar, Marek Wagner, Heng-Chang Chen

## Abstract

HIV-1 reservoirs display heterogeneous nature, lodging both intact and defective proviruses. Recent evidence has shed light on their difference, particularly in the context of immune-mediated selection. To deepen our understanding of such heterogeneous HIV-1 reservoirs and their functional implications, we pioneered the integration of basic concepts of graph theory to characterize the composition of HIV-1 reservoirs. Our analysis revealed noticeable topological properties in networks, featuring immunologic signatures enriched by genes harboring intact and defective proviruses, when comparing antiretroviral therapy (ART)-treated HIV-1-infected individuals and elite controllers. The key variable, the rich factor, played a pivotal role in classifying distinct topological properties in networks. The host gene expression strengthened the accuracy of classification between elite controllers and ART-treated patients. Overall, our work provides a prime example of leveraging genomic approaches alongside mathematical tools to unravel the complexities of HIV-1 reservoirs.

## Introduction

The establishment of latent HIV-1 reservoirs is a complex disease progression mechanism. It involves various types of immune cells and responses converging at the site of HIV-1 infection, aiming at restricting viral propagation. The presence of latent proviruses, causing viral rebound upon interruption of ART, impedes treatment efficacy. A more comprehensive understanding of the establishment of latent HIV-1 reservoirs is essential for designing a potential functional cure against HIV-1 infection. Recent studies have significantly advance our knowledge of latent HIV-1 reservoirs, emphasizing the diverse strengths of immune-mediated selection forces acting on reservoir cells harboring intact and defective proviruses. This diversity results in distinct configurations of reservoirs between each other (Pinzone et al, 2019; Einkauf et al, 2019; Antar et al, 2020; Gandhi et al, 2021; Rozera et al, 2022; Einkauf et al, 2022; Duette et al, 2022; Cho et al, 2022; Lian et al, 2023; Sun et al, 2023; Dufour et al, 2023). The impact of immune selection pressure in elite controllers appears more pronounced than in post-treatment controllers (Jiang et al, 2020; Lian et al, 2021). Furthermore, unique phenotypic signatures associated with reservoir cells harboring intact proviruses (Dufour et al, 2023; Sun et al, 2023) and distinct transcriptomic signatures in HIV-1-infected memory CD4 T cells under ART (Clark et al, 2023) have been reported. These findings underscore the specific microenvironment of latent HIV-1 reservoirs, providing a fertile ground for further investigations into their configurations.

We previously hypothesized that the frequency of HIV-1 integration could serve as a proxy for enriched immunologic signatures during HIV-1 infection (Chen, 2023). Building on this hypothesis, in our current study we employ various theorems and principles involved in graph theory (Diestel, 2017) to obtain a fresh perspective on latent HIV-1 reservoirs. We characterized the network property structured by enriched signatures in ART-treated patients and elite controllers, distinguished by reservoirs harboring intact and defective proviruses, respectively. 309 unique vertices representing enriched signatures were linked by edges indicating correlation coefficients measured between two adjacent vertices. We incorporated variables from enriched signatures, transcriptome, and the spatial genome to illustrate their topological structure. Our observations revealed that variables among enriched signatures were sufficient to maintain an assortative structure of the network across all models tested in this work. Notably, the rich factor variable played a pivotal role in classifying topological properties in ART-treated patients and elite controllers, separated by reservoirs harboring intact and defective proviruses. Interestingly, host gene expression signified in enhancing the accuracy of classification. Altogether, this work introduced a novel perspective on latent HIV-1 reservoirs through topological graphs, providing a method to characterize the architecture of a network associated with its function.

## Results

### Different immunologic signatures enriched in ART-treated patients and elite controllers

A total of 958 and 275 provirus-targeted host genes were collected respectively from HIV-1-infected individuals subjected to ART (Einkauf et al, 2019; Patro et al, 2019; Brandt et al, 2021; Huang et al, 2021; Simonetti et al, 2021; Einkauf et al, 2022; Joseph et al, 2022; Lian et al, 2023) and elite controllers ((Jiang et al, 2020; Lian et al, 2021) (**Extended data Fig. 1a**) in this study. These genes were assigned to four groups (ART-intact, ART-defective, EC-intact, and EC-defective). We first performed the over-representation analysis using MSigDb C7 immunologic signature gene sets on these 1233 genes and revealed that the majority of enriched immunologic signatures differed between reservoirs harboring intact or defective proviruses in both ART-treated patients and elite controllers (**Fig. 1a**, **Extended data Fig. 1b**, **Supplementary Tables 1-4**). The overlays of enriched signatures between different comparisons were demonstrated in **Extended data Fig. 1b**. Significantly, signatures harboring intact proviruses displayed a better magnitude of enrichment, and the overall enrichment was more intense in elite controllers than in ART-treated patients (**Fig. 1b**). While an abundant number (n = 773) of signatures were enriched by genes harboring defective proviruses in ART-treated patients, less than half (n = 239) exceeded the mean of the rich factor calculated using all enriched signatures in ART-treated patients (mean: 2.799). Altogether, these findings indicate that enriched signatures were influenced by different HIV-1 reservoirs (intact *versus* defective) in ART-treated patients and elite controllers.

**Fig. 1.**
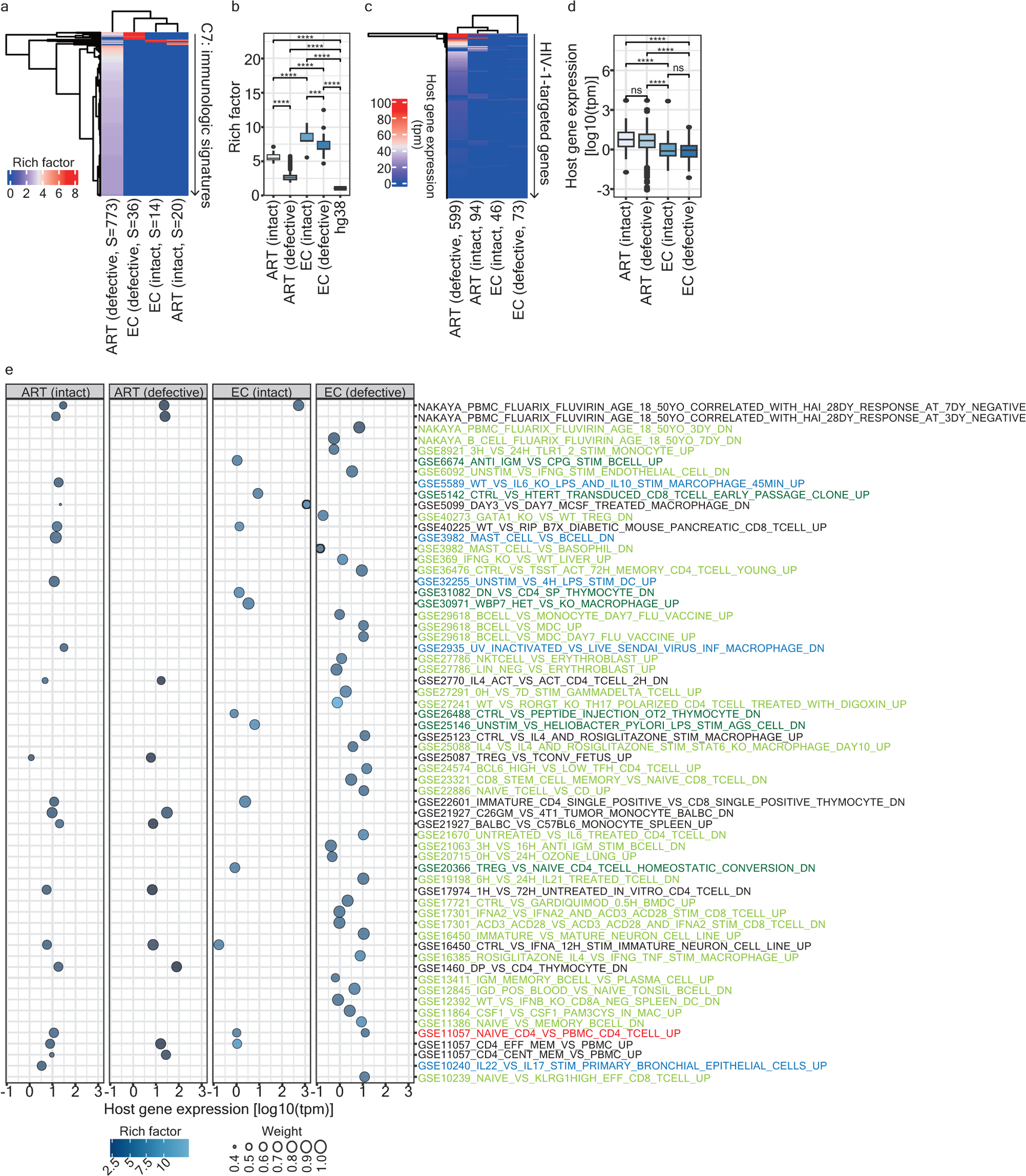
Distinct enriched signatures between HIV-1-infected individuals and elite controllers. (a) A clustering heatmap illustrating immunologic signatures enriched by host genes harboring intact and defective proviruses in both HIV-1-infected patients and elite controllers. Parentheses beneath the column denote the integrity of the provirus genome (intact *versus* defective) along with the number of enriched signatures. The color scale depicts the magnitude of enrichment as represented by rich factors (S: signatures). (b) A box plot displaying the enrichment of signatures represented by rich factors. Details on the commands used for rich factors calculation and the inclusion of a random control “hg38” are described in Chen (2023)^15^ and in **Online Methods**. Statistical significance was determined using the Wilcoxon test in R with default options. The mean and median values for different groups are provided: ART-intact: Mean, 5.655; Median, 5.444. ART-defective: Mean, 2.725; Median, 2.621. EC-intact: Mean, 8.365; Median, 8.030. EC-defective: Mean, 7.304; Median, 6.829. hg38 (control): Mean, 1.039; Median, 1.039. (c) A clustering heatmap showcasing host gene expression (tpm) of genes retrieved from enriched signatures. Parentheses beneath the column indicate the integrity of the provirus genome (intact *versus* defective) and the total number of retrieved genes. (d) A box plot, represented on a logarithmic scale, representing host gene expression (tpm) of the genes retrieved from enriched signatures. The mean and median values for different groups are provided: ART-intact: Mean, 1.87; Median, 0.754. ART-defective: Mean, 1.421; Median, 0.672. EC-intact: Mean, 2.016; Median, −0.108. EC-defective: Mean, 0.432; Median, −0.049 (values on a logarithmic scale). (e) A bubble plot offering insights into selected enriched signatures in HIV-1-infected patients and the complete list in elite controllers. The color scale represents the enrichment magnitude as indicated by rich factors. Weight, calculated based on transcribed genes, highlights difference between genes harboring intact *versus* defective proviruses in both ART-treated patients and elite controllers. Signatures descriptions in different colors signify unique enrichments in reservoirs: Blue (n = 5): intact proviruses in ART-treated patients; Dark green (n = 7): intact proviruses in elite controllers; Light green (n = 34): defective proviruses in elite controllers. A Red (n = 1): Shared in reservoirs with intact and defective proviruses in elite controllers.

### Distinct transcriptome patterns of enriched signatures between ART-treated patients and elite controllers

The involvement of a gene’s transcriptional status in the interaction between HIV-1 integration and enriched signatures remains unclear. To tackle this question, we collected transcriptome data from three studies (Jiang et al, 2020; Einkauf et al, 2022; Clark et al, 2023) and overlaid it with genes retrieved from signatures enriched in four groups, respectively. We observed inconsistencies of the genes targeted by intact or defective proviruses in ART-treated patients with a wide range of gene expression profiles (**Fig. 1c**). However, in a few cases, gene expression was detectable regardless of the integrity of a provirus genome (**Fig. 1c**). The same pattern was observed in elite controllers (**Fig. 1c**). It is noteworthy that our observation aligns with the previous finding that host gene expression patterns in HIV-1 reservoirs (CD4 T cells) are distinct(Clark et al, 2023). Compared to ART-treated patients, the overall host gene expression of genes targeted by proviruses was moderate in elite controllers (**Fig. 1c** and **1d**). The overlays of genes retrieved from enriched immunologic signatures are shown in **Extended data Fig. 1c**.

We continued by calculating the mean of Transcript Per Million (tpm) and the rich factor, as well as the weight across enriched signatures in four groups. Initially, we conducted a cross-comparison based on the 20 enriched signatures harboring intact proviruses against the entire list of enriched signatures harboring defective proviruses in ART-treated patients (**Fig. 1e**) and all enriched signatures in elite controllers (**Fig. 1e**). Subsequently, we retrieved 43 genes present in nine enriched signatures found only in reservoirs harboring intact proviruses in ART-treated patients (**Extended data Fig. 1b**) and performed a KEGG pathway over-representation analysis (**Extended data Fig. 1d**). Nine enriched pathways covering Immune system, Infectious disease: viral, and Cancer: overview were revealed (**Extended data Fig. 1d**). No pathways were enriched when conducting the same analysis with 53 genes present in the enriched signatures harboring both intact and defective proviruses. Further, we selected four genes from these nine enriched pathways and performed RT-qPCR. A positive correlation of host gene expression measured by RT-qPCR and RNA sequencing (Jiang et al, 2020; Einkauf et al, 2022; Clark et al, 2023) (tpm) was observed (*R^2^* = 0.427) (**Extended data Fig. 1e**). At the signature level, we observed a moderately positive correlation (*R^2^*= 0.134) between the mean of tpm from the genes appearing in enriched signatures harboring both intact and defective proviruses in ART-treated patients (**Extended data Fig. 1f**). In the case of elite controllers, only one enriched signature (signature description highlighted in red) was shared between the genes targeted by intact and defective proviruses (**Fig. 1e**). We repeated the KEGG pathway over-representation analysis on genes retrieved from unique signatures enriched in reservoirs harboring either intact (n = 40) or defective (n = 67) proviruses in elite controllers. We observed that only one pathway, Lysine degradation (hsa00310), was enriched in the former case and none of the pathways was enriched in the latter case. These findings suggest that host gene expression may play a role in the identification of enriched signatures in reservoirs harboring intact *versus* defective proviruses in both ART-treated patients and elite controllers.

### Distinct assortativity of network properties between ART-treated patients and elite controllers

To explore how host gene expression and whether the spatial genome also influences the topological property comprising enriched signatures in different groups, we characterized the network architecture (**Extended data Fig. 2a**) using various combinations of category attributes related to enriched signatures (Cat 1), host gene expression (Cat 2) and the spacial genome (Cat 3). Of note, for the ART-defective group we selected the signatures with the enrichment of rich factor exceeding the mean of the enrichment scale (n = 239). With the exception of networks constructed using two and three categories of attributes in elite controllers associated with defective proviruses (**Extended data Fig. 2a**), the majority of the network architectures were represented as disconnected graphs. This indicates that, depending on the utilization of attributes, some networks can consist of two or more subsets of enriched signatures with either low or non-correlation (**Extended data Fig. 2a**). Relative to elite controllers, the network architecture was more assortative in ART-treated patients, especially with reservoirs harboring intact proviruses (**Fig. 2a** and **Extended data Fig. 2b**). Intriguingly, the Cat 2 attribute strengthened the topological property, while Cat 3 attributes failed to reinforce the network structure (**Fig. 2a**). This suggests that Cat 1 and 2 attributes were pivotal in determining the topological property. Consistent with this finding, we observed a higher level of degree connectivity between two adjacent enriched signatures in ART-treated patients compared to elite controllers (**Fig. 2b**). A ranking of the top 10 enriched signatures with the most frequent contact with adjacent signatures demonstrated that signatures such as vertices #41 and #49 were the most essential hubs of connectedness (**Fig 2c**). The coordinates of the series numbers of vertices and incident enriched signatures are listed in **Supplementary Table 5**. Overall, these findings suggest that the network architecture could be more connective in ART-treated patients than in elite controllers.

**Fig. 2.**
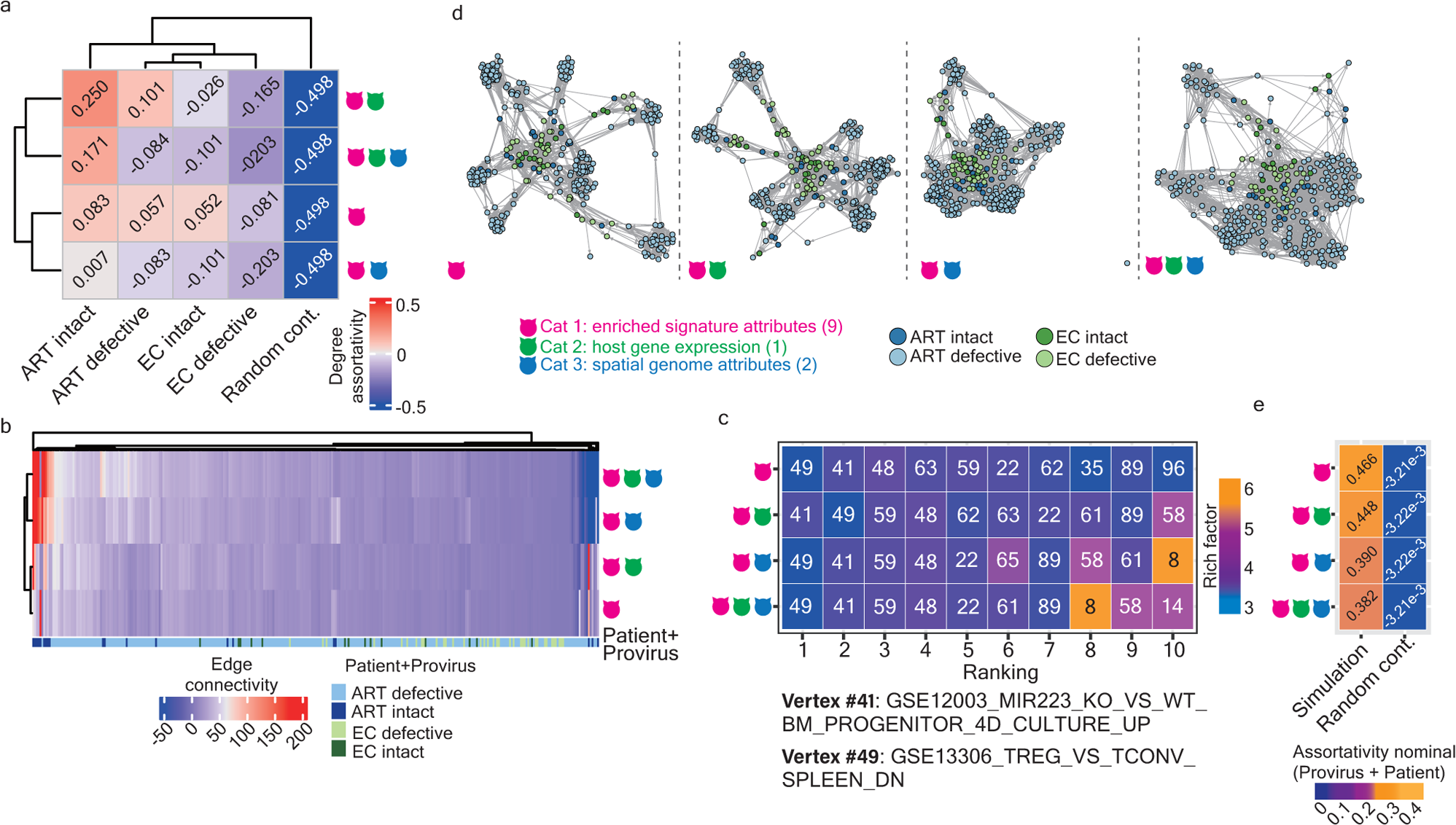
Characteristics of the topological property of the network structured by enriched signatures. (a) A clustering heatmap representing degree assortativity coefficients calculated based on individual networks, using Cat 1, Cat 1 plus Cat 2, Cat1 plus Cat 3, and Cat 1, 2, and 3 attributes. The color scale illustrates the degree of assortativity. (b) A clustering heatmap displaying the edge connectivity of all enriched signatures across four networks (ART-intact, ART-defective, EC-intact, and EC-defective). Edges were calculated using different combinations of category attributes. The color scale denotes the magnitude of degree connectivity. The color code annotation beneath the heatmap denotes the networks where an enriched signature in the “From” ID column was retrieved. (c) A heatmap representing the top 10 enriched signatures with the highest connectedness across four networks. Color codes represent the enrichment magnitude of the rich factor corresponding to each enriched signature. (d) Tetrapartitie graphs illustrating interactions among enriched signatures across four networks, using Cat 1, Cat 1 plus Cat 2, Cat1 plus Cat 3, and Cat 1, 2, and 3 attributes. Signatures marked in dark blue, light blue, dark green, and light green were enriched in reservoirs harboring intact and defective proviruses in ART-treated patients and elite controllers, respectively. Enriched signatures (vertices in tetrapartitie graphs) were linked by edges representing their correlation coefficients. (e) A heatmap representing nominal assortativity coefficients calculated based on the integrity of the provirus genome (intact *versus* defective) and patients (ART-treated *versus* elite controllers) in each network shown in panel (d).

We further examined the topological interaction between two adjacent signatures located in different networks. Initially, we observed a significant number of enriched signatures with a lack of correlation, particularly in the cases involving signatures harboring intact *versus* defective proviruses in ART-treated patients (**Extended data Fig. 3a**) and signatures harboring defective proviruses in ART-treated patients *versus* elite controllers (**Extended data Fig. 3a**). This observation is reflected in the larger interquartile range shown in **Extended data Fig 3b** and is supported by the larger average Euclidean distance (**Extended data Fig. 3c**). Finally, we represented tetrapartite graphs illustrating the interaction of four network architectures (**Fig. 2d**) and observed that all four networks were more structured and distinguishable, particularly when Cat 1 and Cat 1 plus Cat 2 attributes were applied (**Fig. 2d** and **2e**). In summary, these findings suggest that the structural composition of the networks differs from one another and can be influenced by attributes associated with enriched signatures and host gene expression, with the impact of the spatial genome being less influential.

### Rich factor holds paramount significance for classifying network properties

To identify which attributes among the three categories could better classify different properties, we assessed the area under the curve (AUC) of receiver operating characteristic (ROC) curves using logistic regression classifiers constructed with randomly selected predictor variables (**Fig. 3a-3d**). All classifiers effectively distinguished intact *versus* defective proviruses (**Fig. 3a-3d**, top panels namely “Provirus”) and properties of enriched signatures between ART-treated patients *versus* elite controllers (**Fig. 3a-3d** middle panels namely “Patient”). AUC values increased as the number of the attributes in Cat 1 was added (**Fig. 3a-3d**). However, classifiers were less effective in discriminating when considering the combined scenario of “Provirus” plus “Patient” (**Fig. 3a-3d**, bottom panels namely “Provirus+Patient”), especially when Cat 3 attributes were included. It is important to stress that although all classifiers displayed acceptable prediction power, AUC values occasionally varied, indicating that each predictor variable possessed different propensities that could influence the topology of the network. We also constructed random-forest classifiers and used them to rank the importance of predictor variables (**Fig. 3e**). The rich factor variable among all Cat 1 attributes, was of paramount importance in underpinning the prediction power of the models (**Fig. 3e**), and using Cat 1 attributes alone was sufficient for accurate models (**Fig. 3f**). In addition, we observed that cooperation with host gene expression enhanced the robustness of classifiers (**Fig. 3f**). Overall, these findings suggest that Cat 1 attributes, especially rich factor was crucial for classifying networks harboring intact *versus* defective proviruses in ART-treated patients and elite controllers.

**Fig. 3.**
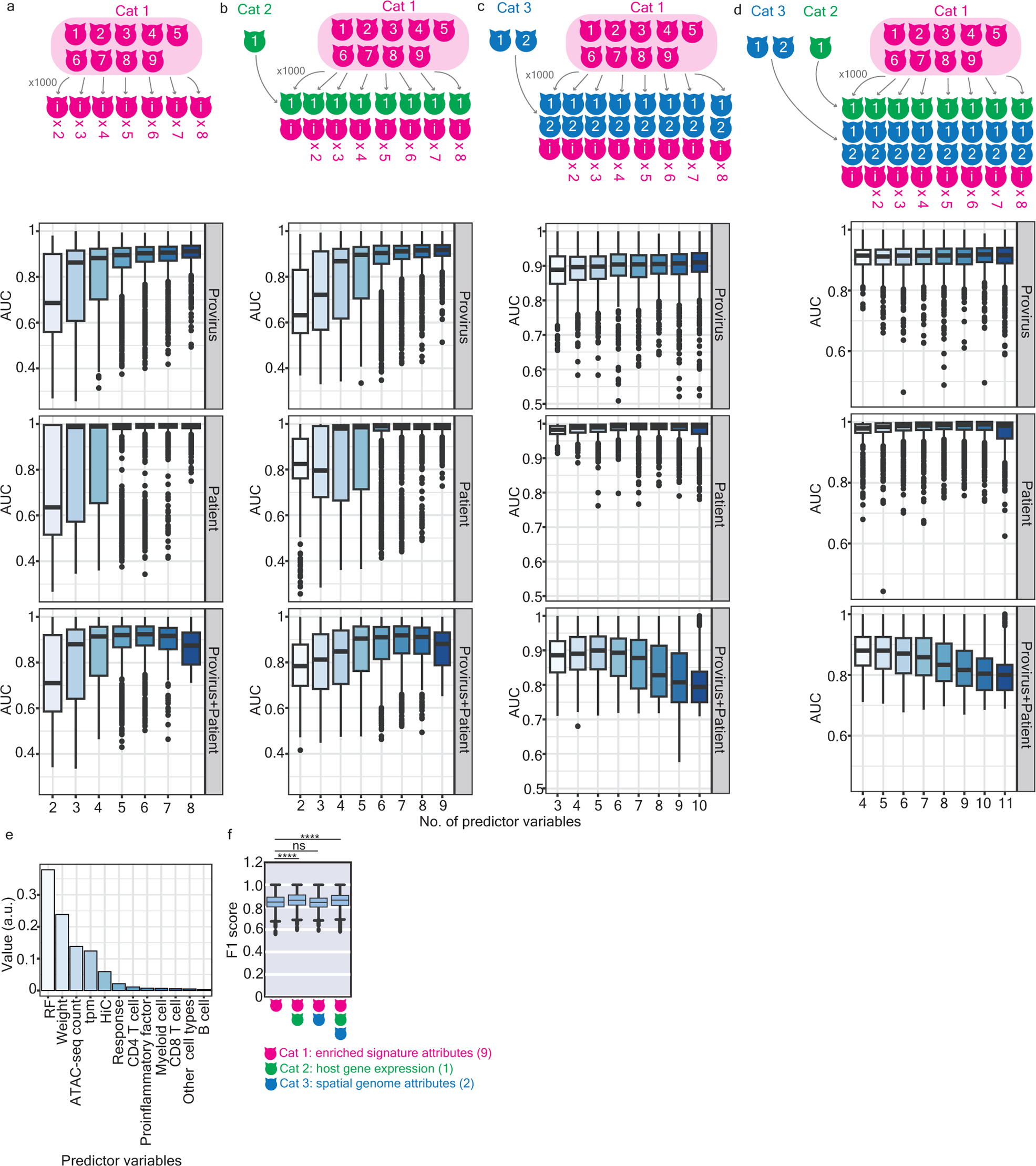
Classification of networks between ART-treated patients and elite controllers. (a, b, c, d) The are under the curve (AUC) of logistic regression models, constructed by bootstrapping selected numbers of predictor variables using Cat 1 (a), Cat 1 plus Cat 2 (b), Cat 1 plus Cat 3 (c), and Cat 1, 2, and 3 attributes (d), calculated to classify the networks associated with intact *versus* defective proviruses (top panel), networks in ART-treated patients *versus* elite controllers (middle panel), and networks associated with intact *versus* defective proviruses, separated by ART-treated patients *versus* elite controllers (bottom panel). The *x*-axis labels representing the number of bootstrapped predictor variables used to construct models. Each classification iteration was repeated 1,000 times for statistical significance. (e) A bar plot representing the ranking of predictor variables importance using a random forest classifier (detailed in **Online Methods**). (f) A box plot illustrating the prediction power for classifying networks harboring intact *versu*s defective proviruses, separated by ART-treated patients *versus* elite controllers. The F1 score was calculated based on 1,000 times of individual train-test splits in models. Significance levels are denoted as follows: * *p* 0.05, ** *p* 0.01, *** *p* 0.001, **** *p* 0.0001.

### Dynamics of the network architecture in a longitudinal order in ART-treated patients

We previously hypothesized that alterations in enriched immunologic signatures are significant, and display a high degree of specificity concerning different immune cell types and proinflammatory soluble factors required alongside HIV-1 infections (Chen, 2023)^15^. In line with this, we applied the same rationale, as described above, to investigate whether the network architecture of enriched signatures changes alongside HIV-1 infections associated with ART, and comparing them to that depicted in elite controllers (**Extended data Fig. 5a**). Similar to the topology characterized in **Extended data Fig. 2a**, the network architecture in a longitudinal order was represented as disconnected graphs (**Extended data Fig. 5a**). The network in pretreatment HIV-1-infected individuals strongly resembled a null graph with zero degrees of assortativity (**Fig. 4a** and **Extended data Fig. 5a**). We assume that this observation is likely due to the low number of enriched signatures (n = 8). Although the degree of assortativity varied among HIV-1-infected individuals subjected to short and long period of ART and elite controllers when different category attributes were applied, it appeared that Cat 1 attributes were pivotal in offering a better structural property of the network (**Fig. 4a**). A higher level of degree connectivity was observed in signatures enriched in HIV-1-infected individuals subjected to a long period of ART (**Fig. 4b**). Vertices #14, #47, and #55 were identified as the most essential hubs of connectedness (**Fig 4b** and **4c**). The coordinates of the series numbers of vertices and incident enriched signatures are listed in **Supplementary Table 6**.

**Fig. 4.**
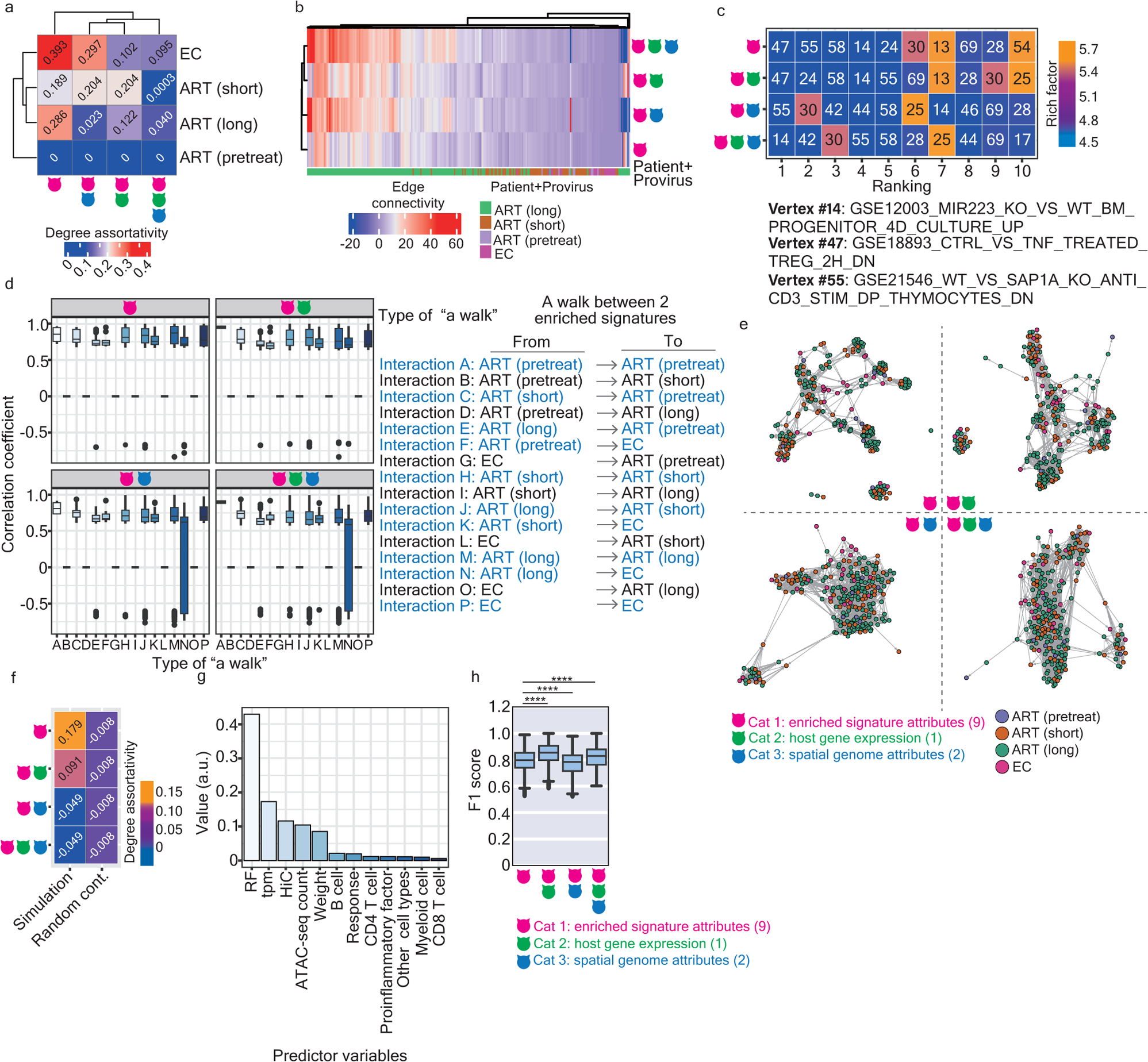
Dynamics of the networks alongside HIV-1 infections in patients subjected to ART and in elite controllers. (a) A clustering heatmap representing degree assortativity coefficients based on individual networks, considering various attibites (Cat 1, Cat 1 plus Cat 2, Cat1 plus Cat 3, and Cat 1, 2, and 3 attributes) for pretreatment HIV-1-infected individuals, patients subjected to a short and a long period of ART, and elite controllers. (b) A clustering heatmap illustrating the edge connectivity of all enriched signatures capable of bridging possible walks in HIV-1-infected individuals in a longitudinal order and in elite controllers. Edges between adjacent enriched signatures were calculated using various combinations of category attributes. The color scale denotes the magnitude of degree connectivity. The color code annotation beneath the heatmap denotes the networks where an enriched signature in the “From” ID column was retrieved. (c) A heatmap representing the top 10 enriched signatures with the highest connectedness in HIV-1-infected individuals in a longitudinal order and in elite controllers. Color codes represent the enrichment magnitude of the rich factor corresponding to each enriched signature. (d) A box plot illustrating the correlation coefficient between two adjacent enriched signatures. All possible pairs of walks are provided in **Supplementary Table 7**. Statistical significance was determined using the Wilcoxon test in R with default options. The uni-direction of a walk is defined in **Online Methods**. Types of a walk written in black indicate the absence of such a connectedness in the complete edge list. (e) The tetrapartitie graphs illustreating the interactions among enriched signatures across four networks using different attribute combinations, namely Cat 1, Cat 1 plus Cat 2, Cat1 plus Cat 3, and Cat 1, 2, and 3. Signatures marked in violet, marron, green, and deep pink were enriched in reservoirs from pretreatment HIV-1-infected individuals, patients subjected to a short and a long period of ART, and elite controllers, respectively. Enriched signatures (vertices in tetrapartitie graphs) were linked by edges representing their correlation coefficients. (f) A heatmap representing the degree assortativity coefficient based on tetraparitie graphs shown in panel (e). (g) A bar plot ranking the importance of predictor variables using a random forest classifier, detailed in **Online Methods**. (h) A box plot representing the prediction power for classifying networks in HIV-1-infected individuals in a longitudinal order and in elite controllers. The F1 score was calculated based on 1,000 times of individual train-test splits in models. Significance levels are denoted as follows: * *p* 0.05, ** *p* 0.01, *** *p* 0.001, **** *p* 0.0001.

We further compared the correlation coefficient between two adjacent enriched signatures bridged by a walk with direction (**Fig. 4d**). According to our definition of the direction of a walk described in **Online Methods**, we observed that a walk was only realized in the incident that an enriched signature connects to another, either present in the same network or in the networks that already exist (**Fig. 4d**, walks highlighted in blue). Interestingly, no direct walk from signatures enriched in elite controllers to those enriched in non-elite controllers was observed (**Fig. 4d**, walks written in black). This observation may imply a difference in the intrinsic property between elite controllers and ART-treated patients in the context of enriched signatures, host gene expression, and the spatial genome. We also observed that although the correlation coefficient between two adjacent enriched signatures varied in most types of walks (**Fig. 4d**), the proportion of types of walks associated with high (> −0.5) and low (< −0.5) correlation coefficients showed no significant difference (**Extended data Fig. 5f**).

In the final step, we illustrated tetrapartite graphs depicting the interaction among three networks in a longitudinal order and the network in elite controllers (**Fig. 4e**). Once again, we observed that Cat 1 attributes were sufficient to sustain its topology (**Fig. 4e** and **Fig. 4f**). Subsequently, we computed the importance of each variable and found, once again, that the rich factor variable was of paramount importance for ensuring robust prediction power, followed by the host gene expression variable (**Fig. 4g**). Of note, the host gene expression variable played an important role in distinguishing the networks between ART-treated patients and elite controllers (**Fig. 4h** and **Extended data Fig. 5g**) and exhibited moderate effectiveness in classifying dynamic networks between ART-treated patients at different stages of treatment and elite controllers (**Fig. 4h** and **Extended data Fig. 5h**).

## Discussion

While HIV-1 latency has been a subject of extensive research for many years, our current understanding remains limited in precisely visualizing latent HIV-1 reservoirs responsible for viral rebound. In this work, we proactively employed graph-theoretical tools to define the topological properties of a network formed by enriched immunologic signatures. Despite observing a substantial number of enriched signatures in reservoirs harboring defective proviruses in ART-treated patients (n=773), Only 30.9% (n = 239) and 0.91% (n =7) of the signatures with the enrichment of rich factor exceeded the mean (2.799) and the median (2.621) of the enrichment scale. This suggests that these signatures might be either temporarily enriched or represent background noise. We hypothesize that proviruses associated with these signatures could potentially be eliminated during the selection process.

In this work, we defined the network architecture based on correlation coefficients between two adjacent enriched signatures. The utilization of different attributes could influence its topology. Of note, we observed that Cat 1 attributes could be deemed as pillars supporting property’s topology. The rich factor variable within Cat 1 alone already demonstrated sustained predictive strength. However, we also observed that, relative to ART-treated patients, models applied to elite controllers faltered (**Extended data Fig. 4a**), suggesting that, in elite controllers, either the sample size was small or additional factors that governing such HIV-1 reservoirs have not yet been identified. Although the contribution of Cat 3 attributes was not emphasized in this work, a more detailed examination of the influence of different hierarchical 3D genome organizations on the property’s topology will be required. Overall, in contrast to elite controllers, the characteristics of the network architecture in ART-treated patients include: (1) a less intense magnitude of enrichment of signatures, (2) high degree of assortativity, and (3) high connectedness between two adjacent vertices (**Fig. 5**). This signifies that the network architecture was more connective and structural in ART-treated patients (**Fig. 5**).

**Fig. 5.**
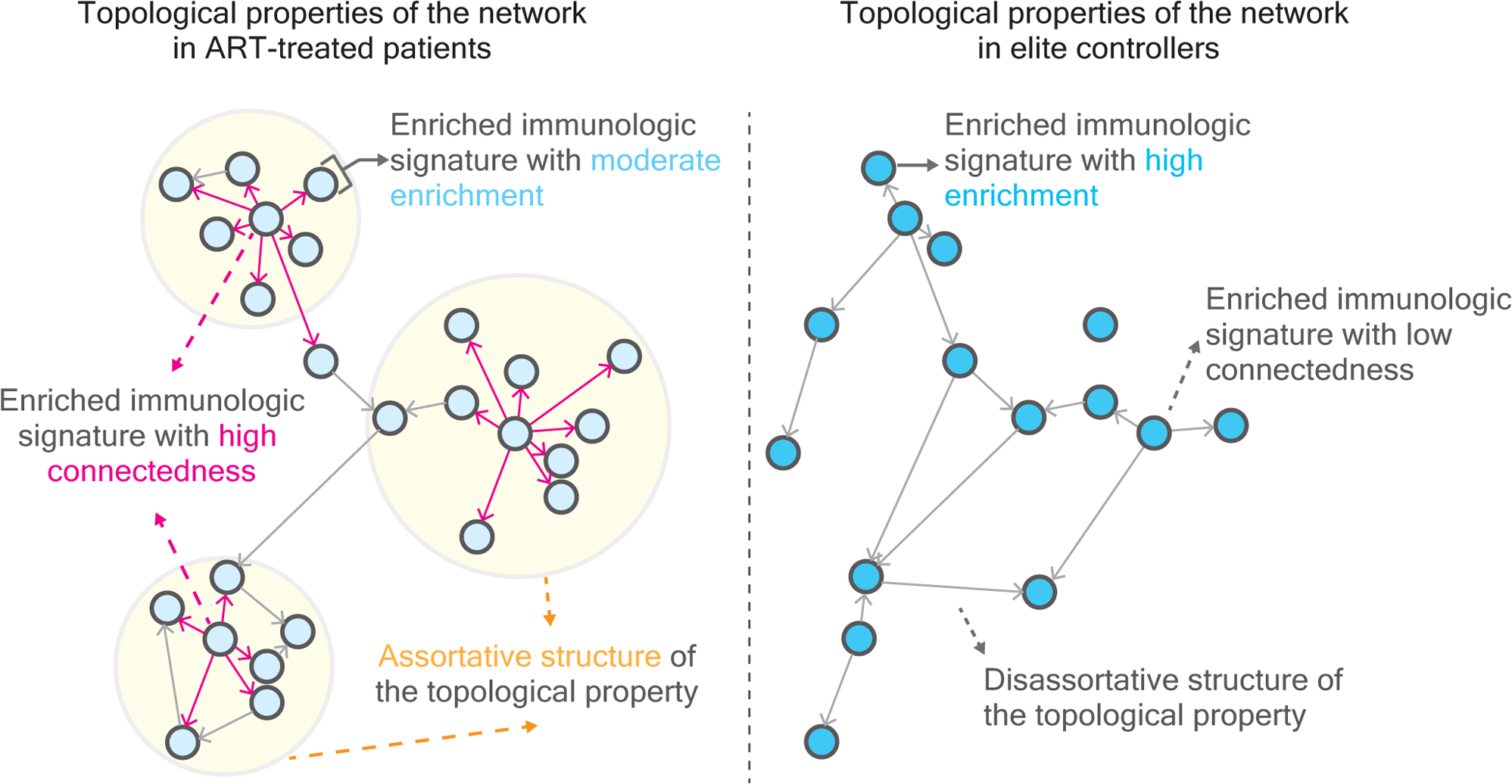
Proposed model of distinct topological properties of functional genome networks between ART-treated patients and elite controllers. We hypothesize that, in comparison to elite controllers, the network structured formed by enriched immunologic signatures in ART-treated patients is moderately enriched and exhibits the higher level of connectedness, resulting in a more assortative architecture. This assortative structure is particularly in reservoirs harboring intact proviruses in ART-treated patients.

We also explored the evolutionary dynamics of the network using longitudinal HIV-1 integration. Despite grappling with the challenges of a limited number of integration sites retrieved from pretreatment HIV-1 individuals, we extrapolated our findings by examining uni-directional walks from signatures enriched in ART-treated patients to those enriched in elite controllers (**Fig. 4d**). This led us to postulate the existence of an intrinsic property barrier between these two groups of HIV-1-infected individuals. Our proposition suggests that connectedness from signatures enriched in ART-treated patients to elite controllers reveals the potential of proviruses in ART-treated patients to reach a state of deep latency, resembling the transcriptional state of proviruses in elite controllers. Conversely, a lack of connectedness in the reverse direction indicates that the awakening of proviruses from deep latency was less feasible. Latent HIV-1 reservoirs are known to be stable (Chun et al, 1997, 1999; Finzi et al, 1999), and *ex vivo*, a subset of the intact and replication-competent proviruses fail to be reactivated even after multiple rounds of stimulation (Ho et al, 2013), suggesting that in some cases, deep latency can be refractory to reactivation. Overall, this work represents an inaugural step in utilizing genomic approaches with graph-theoretical tools to enhance our understanding of latent HIV-1 reservoirs. The subsequent step involves discerning the biological functions of individual enriched signatures and investigating further ramifications for the microeviroment of deeply latent HIV-1 reservoirs.

## Materials and Methods

### Acquisition and procession of public datasets

A through literature search was conducted on PubMed, using the keywords (((intact) OR (intact provirus) OR (intact proviruses)) AND ((HIV) OR (HIV-1))), accessed on March 27, 2023, as previously described in Więcek and Chen (2023) (Więcek & Chen, 2023). Research articles were selected from the first 1,000 cited papers in PubMed between 2005 to March 2023. For this study, we analyzed eight (Einkauf et al, 2019; Patro et al, 2019; Brandt et al, 2021; Huang et al, 2021; Simonetti et al, 2021; Einkauf et al, 2022; Joseph et al, 2022; Lian et al, 2023) studies related to HIV-1 integtation in ART-treated patients and two (Jiang et al, 2020; Lian et al, 2021) studies related to elite controllers (**Extended data Fig. 1a**). Host genes targeted by HIV-1 integration, as reported by Jiang et al. (2020) (Jiang et al, 2020), were previously analyzed and documented in Chen (2023) (Chen, 2023). Transcriptome sequencing data for ART-treated patients were retrieved from Einkauf et al. (2022) (Einkauf et al, 2022) (GEO: GSE144334) and Clark et al. (2023) (Clark et al, 2023). Transcriptome sequencing data for elite controllers were retrieved from Jiang et al. (2020) (Jiang et al, 2020) (GEO: GSE144332). ATAC-seq (GEO: GSE144329) and HiC datasets (GEO: GSE168337) were retrieved from Jiang et al. (2020) (Jiang et al, 2020) and Einkauf et al. (2022) (Einkauf et al, 2022), respectively.

### MSigDb over-representation analysis

A total of 958 provirus-targeted host genes (200 genes harboring intact *versus* 758 genes harboring defective proviruses) and 275 provirus-targeted host genes (111 genes harboring intact *versus* 164 genes harboring defective proviruses) were collected respectively from HIV-1-infected individuals subjected to ART (eight studies) (Einkauf et al, 2019; Patro et al, 2019; Brandt et al, 2021; Huang et al, 2021; Simonetti et al, 2021; Einkauf et al, 2022; Joseph et al, 2022; Lian et al, 2023) and elite controllers (two studies) (Jiang et al, 2020; Lian et al, 2021) to initiate the analysis. The R package clusterProfiler (Version 4.4.1) (Yu et al, 2012; Wu et al, 2021) was used to compute enriched immunologic signatures with the function `enricher()` and default options. Over-representation analysis (Boyle et al, 2004) was performed using C7 immunologic signature gene sets from the Molecular Signatures Database (MSigDb) (Subramanian et al, 2005; Liberzon et al, 2011, 2015) as the background. Enriched signatures with *p*-values (adjusted by the Benjamini-Hochberg method) below 0.05 were selected. The command lines for calculating rich factors and generating a random control labeled “hg38” in **Fig. 1b** were detailed in Chen (2023) (Chen, 2023). In brief, the rich factor, representing the enrichment score for each enriched immunologic signature, was calculated by dividing GeneRatio by BgRatio. In **Fig. 1b**, “hg38” denotes the rich factor calculated using all protein-coding genes (rich factor: median, 1.162; mean, 1.154). Weight was demonstrated by dividing the total number of the transcribed genes retrieved from individual enriched signatures but not present in counterparts, such as genes targeted by intact proviruses rather than defective ones in either ART-treated patients or elite controllers. The outputs of over-representation analysis for genes harboring intact and defective proviruses in ART-treated patients and elite controllers are presented in this study and can be found in **Supplementary Tables 1-4**. Outputs related to longitudinal HIV-1 integration were downloaded from Chen (2023) (Chen, 2023).

### Assignment of predictor variables in category 1 (Cat 1)-, category 2 (Cat 2)-, and category 3 (Cat 3)-attributes

We utilized all enriched immunologic signatures from ART-intact (n = 20), EC-intact (n = 14), EC-defective (n = 36), and selected the enriched signatures in ART-defective with rich factors over than the mean (2.799) of the enrichment scale (n = 239) as the input signatures (n = 309) to illustrate topological properties of the network. Nine attributes: (1) rich factor, (2) weight, (3) involvement of CD4 T cells, (4) involvement of CD8 T cells, (5) involvement of B cells, (6) involvement of myeloid cells, (7) involvement of other cell types, (8) involvement of proinflammatory factors and (9) immune response labeled in immunologic signatures were used in Cat 1 attributes. Attributes (3) to (7) were constructed based on the presence of the indicated cell type in the signature description, with a character “1” denoting its presence, and a character “0” indicated their absence. A character “1” was denoted if proinflammatory factor is present in the signature description; otherwise, a character “0” was given. For immune response, a character “0” indicated no description in the signature description, a character “1” indicated down-regulation, and a character “2” indicated up-regulation. Attributes (1) and (2) were numeric attributes. Cat 2 attribute referred to Transcript Per Million (tpm), calculated by dividing RNA sequencing raw reads (Jiang et al, 2020; Einkauf et al, 2022; Clark et al, 2023) by the length of a gene in kilobases (reads per kilobase, RPK), followed by dividing by the sum of all RPK values divided by 1,000,000. Cat 3 attributes included data from ATAC-seq and intrachromosome HiC followed by high throughput sequencing. The HiC outputs (GSM5136368, GSM5136369, and HiC_GSM5136370) (Einkauf et al, 2022) with 10-kilobase resolution were combined and overlaid on genes retrieved from enriched signatures in order to determine their topological distribution using the command *intersect* with default options in bedtools (Quinlan & Hall, 2010). The same pool of the gene was also overlaid on the ATAC-seq readout analyzed by Einkauf et al. (2022) (Einkauf et al, 2022) to identify genes within ATAC-seq coverage regions. A comprehensive list of all attributes associated with enriched signatures harboring intact and defective proviruses in ART-treated patients and elite controllers, as well as enriched signatures obtained from longitudinal HIV-1 integrations, is provided in **Supplementary Tables 5** and **6**, respectively.

### Measurement of correlation coefficients of enriched immunologic signatures

The R package “Hmisc” (https://CRAN.R-project.org/package=Hmisc) was employed to calculate correlation coefficients among enriched signatures using the function `rocrr()`. Subsequently, correlation matrices were transformed into data frames containing four columns. The first two columns served as ID columns, designating enriched signatures as “From” and “To” within a walk, respectively. The remaining columns included the correlation coefficient and the associated *p-*value. To filter out weak or spurious connections, correlation coefficients with a *p-*value > 0.05 was excluded. **Supplementary Tables 5** and **6** provide the correspondence between ID numbers and the associated enriched signatures.

### Visualization of the network architecture

For each individual network, *w*e utilized the previously mentioned correlation matrix as the edge list. In this list, the columns of the row and the column name were designated as the “|From” and “To” vertices, respectively, with the correlation coefficient representing the edge. The node list consisted of a series ID for all enriched signatures, information about the integrity of a proviral genome, and the classification of HIV-1-infected individuals (ART-treated patients *versus* elite controllers). “From” and “To” signatures were selected to remain in the same network. It is important to note that we generated separate node lists for each property. The network structure was then established using the function `graph_from_data_frame()` from the R package “igraph” (https://igraph.org) (Csárdi et al, 2023) with the following arguments: d for the the edge list, vertices for the node list, and directed = TRUE to account for directed edges in the network.

For the visualization of two networks (bipartite graphs) we selected two adjacent signatures enriched in distinct topological properties to illustrate their topological interactions. It is important to emphasize that only a single and uni-directional (from the row denoted as “From” to the column denoted as “To”) bridged by two adjacent enriched signatures was considered in this study. The same function and arguments from the R package “igraph” (https://igraph.org) (Csárdi et al, 2023) were employed for visualizing these networks.

Regarding the four networks (tetrapartite graphs), *t*he complete edge and the node list, which includes all pairs of two adjacent enriched signatures and the corresponding correlation coefficients, were used as arguments, d and vertices, respectively, in the function `graph_from_data_frame()` to depict these tetrapartite graphs. Edge connectivity (**Fig. 2b** and **4b**) was calculated based on the correlation matrix resulted from the tetrapartite graph. Connectedness was determined by dividing the number of edges of each signature, shown in the first ID column (namely “From”), by the mean of correlation coefficients of edges radiating from the incident signature. The complete edge list for plotting **Fig. 4d** is provided in **Supplementary Table 7**. The percentage and absolute number of walks between two adjacent signatures are shown in **Extended data Fig. 5b-5e**. It is important to note that in **Fig. 4d**, types of a walk written in black indicate the absence of such connectedness in the complete edge list.

### Clustering heatmap

The cluster heatmaps representing the magnitude of the enrichment of immunologic signatures (**Fig. 1a**), host gene expression of the genes retrieved from enriched signatures (**Fig. 1c**), assortativity analysis (**Fig. 2a** and **Fig. 4a**), edge connectivity of enriched immunologic signatures (**Fig. 2b** and **Fig. 4b**), and the number of walks between two adjacent enriched signatures (**Extended data Fig. 5b-5e**) were created using the R package ComplexHeatmap (Gu et al, 2016) with default options. All plots presented in this manuscript were generated in R with default options.

### Measurement of degree and nominal assortativity coefficient and Euclidean distance of the network architecture

Degree and nominal assortativity coefficients were computed using the function s`assortativity_degree()` and `assortativity_nominal()` in the R package “igraph” (https://igraph.org) (Csárdi et al, 2023). For **Extended data Fig. 2b**, 10 enriched signatures in each scenario were randomly sampled with replacement to obtain degree assortativity coefficients using the mentioned function, and this process was repeated 1,000 times. Statistical tests were performed with R with default options. Additionally, Euclidean distance (**Extended data Fig. 3c**) was calculated using the function `dist()`with the argument method for “euclidean” in the R package “stats”, which is a part of R (https://www.r-project.org/).

### Classification of the networks

Logistic regression-based classification: we divided the complete dataset. containing 309 enriched signatures associated with attributes into a training set (80% of the dataset) and a testing set (20% of the dataset) for logistic regression running on R. The logistic regression model was fitted using the function `glm()`with the argument family specified as “binomial” in the R package “stats” (https://www.r-project.org/). Three types of responses were considered: provirus (intact *versus* defective provirus), patient (ART-treated patients *versus* elite controllers), and provirus + patient. These responses were included in an object of class “formula”. Different numbers and combinations of the “term” referred to category attributes in an object of class “formula” were bootstrapped with replacement and this process was repeated 1,000 times. Receiver operating characteristic (ROC) and the area under the curve (AUC) were calculated using the functions `multiclass.roc()` and `auc()` in the R package “pROC” (Robin et al, 2011), respectively. The whole procedure was repeated 1,000 times for statistical robustness.

Random forest classification: Separate models were made for six classification tasks, all following the same pipeline: M1 – multiclass classification of enriched signatures harboring intact *versus* defective proviruses in ART-treated patients and elite controllers, M2 – binary classification of enriched signatures harboring intact *versus* defective proviruses in ART-treated patients, M3 – binary classification of enriched signatures harboring intact *versus* defective proviruses in elite controllers, M4 – multiclass classification of immunologic signatures enriched in pretreatment HIV-1-infected individuals, patients subjected to a short and long period of ART and elite controllers, M5 – as in M4, but excluding elite controllers, M6 – as in M4, but excluding pretreatment HIV-1-infected individuals. The pipeline was implemented in Python 3.11.6 using pandas 2.1.3 (https://pandas.pydata.org/docs/index.html) and scikit-learn 1.3.2 packages (https://scikit-learn.org/stable/whats_new/v1.3.html). Plots were generated using seaborn 0.13.0 package (Waskom, 2021).

First, 30% of the data was reserved for testing, ensuring class balance. Subsequently, random forest classifiers from the Python scikit-learn package (https://jmlr.csail.mit.edu/papers/v12/pedregosa11a.html) were trained on the training set with default parameters to extract impurity-based feature importance. In all cases, all Cat 1 attributes except rich factor and weight exhibited much lower importance than other attributes and were thus removed from further classification.

Next, grid search cross-validation was conducted to select hyperparameters for the final random forest classifiers in each task. The tested hyperparameter values were included: n_estimators (20, 100), criterion (gini, log_loss), max_features (sqrt, log2), min_samples_split (3, 5, 10), min_samples_leaf (1, 4), and class_weight (None, balanced). The remaining parameters were set to default values. Given the small size of the data sets, especially minority classes, repeated stratified k-fold validation was performed with 10 iterations of 5 randomly selected validation splits. The macro-averaged F1 score guided model selection in multiclass tasks (M1, M4-M6), while the positive class F1 score was used in binary classification tasks (M2 and M3).

Finally, model evaluation was carried out. Due to the limited size of the datasets and the presence of minority classes, which could lead to strong dependence of model performance on a specific selection of samples for the test set in a single train-test split, each model was independently re-trained and evaluated on 1000 randomly generated stratified 70%: 30% splits to mitigate this bias and generate robust statistics.

To compare the impact of Cat 1, 2, and 3 attributes on classification, the following approaches were employed for each task: classification using Cat 1 attributes, Cat 1 and Cat 2 attributes, Cat1 and Cat 3 attributes, and all category attributes. Resulting distributions of F1 scores, macro-averaged for multiclass tasks and positive class for binary tasks, were presented using kernel density estimation (KDE) and box plots. Median F1 scores were compared, and statistical significance was assessed using the Wilcoxon test from the Python SciPy 1.11.3 package (Virtanen et al, 2020). In some cases, KDE plots displayed multiple maxima, particularly in tasks M2 and M3 (**Extended data Fig. 4c** and **4d**), indicating binary classification tasks with small positive classes. This phenomenon is attributed to the discrete difference in F1 score resulting from even a single positive class sample having a different prediction. This underscores the importance of evaluating models across multiple independent data splits. Additionally, it is noteworthy that the accuracy of classifying the networks in longitudinal ART-treated patients *versus* elite controllers improved when a small sample size from pretreatment HIV-1-infected individuals was removed (**Extended data Fig. 5h**).

### Cell culture

The human T lymphoblast Sup T1 cell line was cultured at 37°C under a 95% air and 5% CO_2_ atmosphere in Roswell Park Memorial Institute medium (RPMI 1640 medium; Sartorius #01-100-1A) supplemented with 10% fetal bovine serum (FBS; Gibco #A5256801) and 1% penicillin-streptomycin (PAN Biotech #P06-07100). HEK 293T cells were cultured under the same conditions in Dulbecco’s modified Eagle’s medium (DMEM; Gibco #11965-092) supplemented with 5% FBS and 1% penicillin-streptomycin. Routine testing for mycoplasma contamination was conducted every two weeks.

### Transfection and viral infection

For the preparation of viral stocks, 2.5 × 10^6^ HEK 293T cells in 10-cm dishes were transfected with 6.66 µg of pVSV-G and 13.33 µg of HIV-1 plasmid (NL4.3_ΔEnv_EGFP). After 16 hours, the DMEM medium was replaced with the RPMI medium and infectious viral particles were collected 48 and 72 h after transfection. Infection of SupT1 cells was performed by spinoculation; 1×10^6^ SupT1 cells in 100 µl of RPMI supplemented with 8 µg/ml of polybrene were infected with NL4.3_ΔEnv_EGFP by spinoculation (1000×g, 2h, 32°C) at an MOI of approximately 0.5. The medium was replaced with 3 mL fresh RPMI 24 h after infection. The efficiency of infection was monitored four days after infection by flow cytometry analysis.

### RNA extraction and reverse transcription (RT)

Total RNA extraction was carried out using the AllPrep DNA/RNA Mini Kit(50) (Qiagen Cat. No. 80204) following the manufacturer’s instructions. Subsequently, mRNA purification was performed with RNeasy Pure mRNA Bead Kit (Qiagen Cat. No. 180244). 28 ng mRNA was used to synthesize cDNA using 50 μM Oligo(dT)20 primers, and NG dART RT-PCR Kit (EURx Cat. No. E0802-02) were used at 50°C for 1 h.

### Quantitative PCR (qPCR)

Four genes, namely, *chuk*, *hif1a*, *jak1*, and *oas2* were selected for qPCR. Each qPCR reaction was assembled with FastStart Universal SYBR Green Master (Rox) (Roche Cat. No. 4913850001) and 2 µl of undiluted cDNA sample in 10 µl total volume. The qPCR reaction conditions were set as follows: 95°C for 10 min and 95°C for 10 s, 58°C for 30 s, 72°C for 30 s for 50 cycles. The reaction was terminated by performing a melting curve analysis at the end of amplification. The obtained qRT-PCR data were analyzed using the ΔΔCt method, with normalization to the expression of β-actin (ACTB). Primer sequences used for the qPCR reaction are provided in **Supplementary Table 8**.

### Statistics

All statistical tests were performed using R with default options and specific details are provided in the main text and figure legends where applicable.

### Data availability

Publicly available datasets were analyzed in this study and their origins are detailed in the **Acquisition and procession of public datasets** section. The analyzed data, comprising lists of enriched immunologic signatures, coordinates between ID numbers and incident enriched signatures associated with predictor variables, and lists of all possible walks are provided in **Supplementary Tables**.

### Code availability

All code and scripts provided in this work are available on GitHub (https://github.com/HCAngelC/Network_structure_of_HIV_IS). The open-source packages used in this study, which have not been assigned DOIs, are listed as follows: The R package “Hmisc” was used to calculate correlation coefficients (Harrell Jr., F., & Dupont, Ch. (2019). Hmisc: Harrell Miscellaneous. R Package Version 4.2-0. https://CRAN.R-project.org/package=Hmisc). The R package “igraph” was used for the analysis of the network structure (Csardi G, & Nepusz T (2006). The igraph software package for complex network research. InterJournal, Complex Systems, 1695. https://igraph.org). Python 3.11.6 with pandas 2.3.1 was used to construct random forest-based classifiers (https://pandas.pydata.org/docs/). Python scikit-learn 1.3.2 package was used to construct random forest-based classifiers (Scikit-learn: Machine Learning in Python, Pedregosa et al., JMLR 12, pp. 2825-2830, 201).

## Acknowledgments

HCC acknowledges funding from the Narodowe Centrum Nauki (Sonata Bis Grant UMO-2022/46/E/NZ6/00022). AKP and HA acknowledge funding from the National Science Centre, Poland (Sonata Bis Grant UMO2018/30/E/NZ1/00874). AKP acknowledges funding from the National Science Centre, Poland (OPUS Grant UMO-2022/45/B/NZ3/03890). MW acknowledges funding from the National Science Centre, Poland (Sonata Bis Grant UMO-2022/46/E/NZ6/00131).

## Author contributions

Conceptualization, H.-C.C.; methodology, H.-C.C., and J.W.; software, H.-C.C., and J.W., formal analysis, H.-C.C., and J.W., investigation, H.-C.C., J.W., A.K.-P., K.W., and H.A.; resources, H.-C.C.; data curation, H.-C.C., and J.W.; writing of original draft manuscript, H.-C.C., M.W., J.W., and K.W.; writing, manuscript review and editing, H.-C.C., M.W., A.K.-P., and J.W.; visualization, H.-C.C., and J.W.; supervision, H.-C.C., project administration, H.-C.C.; funding acquisition, H.-C.C.

---

**Extended data Fig. 1.**
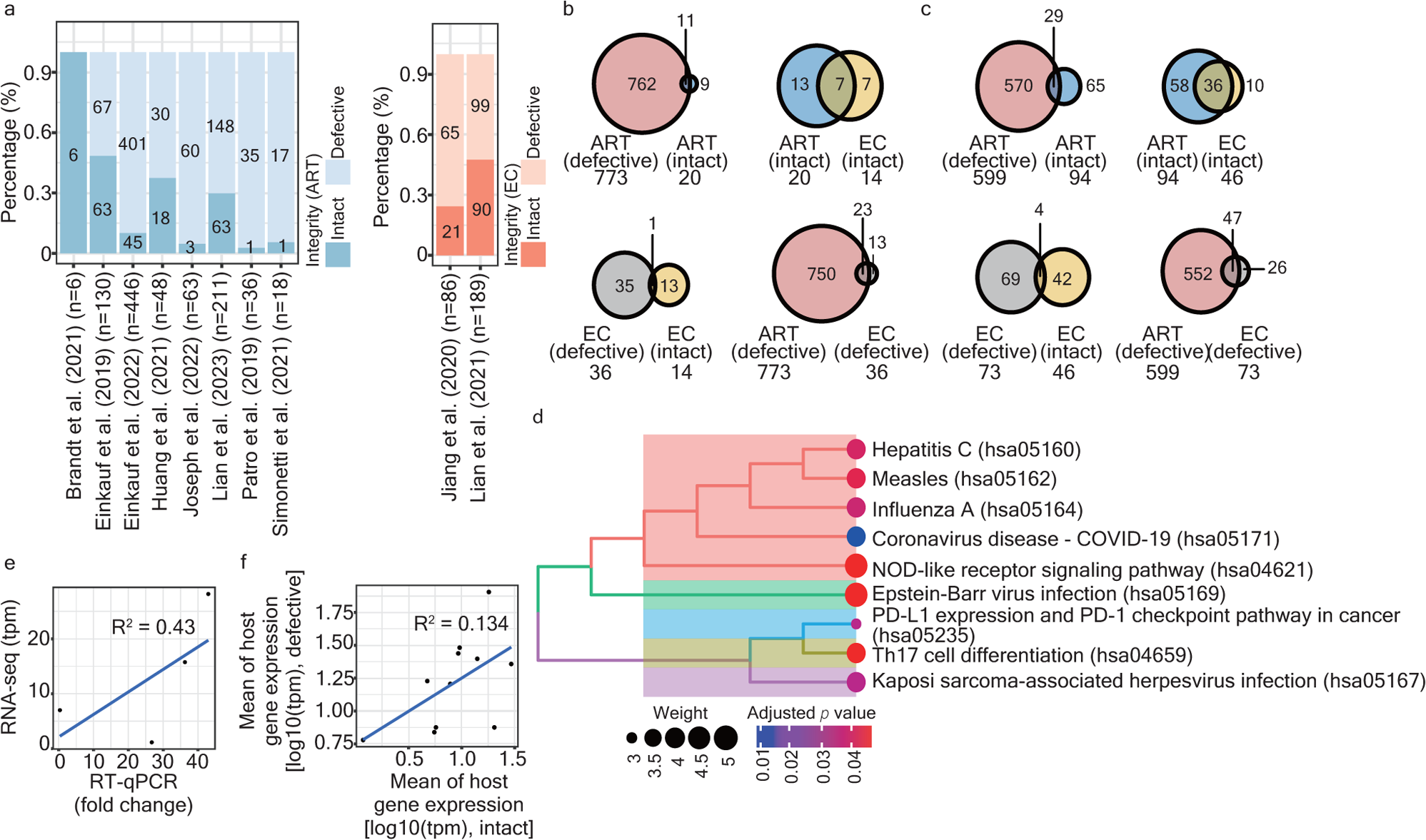
Characteristics of enriched signatures in ART-treated patients and elite controllers. (a) A stacked bar plot representing the proportion of genes harboring intact *versus* defective proviruses in research articles. The left panel lists articles in which provirus-targeted genes were retrieved from ART-treated patients, while the right panel lists articles where provirus-targeted genes were retrieved from elite controllers. (b) Venn diagram representing the overlap of enriched signatures between different comparisons, with the total number of enriched signatures indicated under the name of respective groups. (c) Venn diagram representing the overlap of provirus-targeted genes retrieved from enriched signatures between different comparisons, with the total number of provirus-targeted genes indicated under the name of respective groups. (d) A clustering tree plot representing KEGG pathways enriched by 43 genes retrieved from signatures uniquely enriched in reservoirs harboring intact proviruses in ART-treated patients. (e) A scatter plot representing the correlation of expression of four selected genes in signatures uniquely enriched in reservoirs harboring intact proviruses in ART-treated patients. The *x*-axis indicates the measurement of host gene expression from RT-qPCR, with β-actin (ACTB) as a reference. Data are the mean of two independent measurements. The *y*-axis indicates the measurement retrieved from RNA sequencing published in Einkauf et al. (2022)^6^ and Clark et al. (2023)^14^ (detaileds provided in **Online Methods**). (f) A scatter plot representing the mean transcriptional state of eleven common signatures enriched in both reservoirs harboring intact and defective proviruses in ART-treated patients. Each dot represents a unique enriched immunologic signature.

**Extended data Fig. 2.**
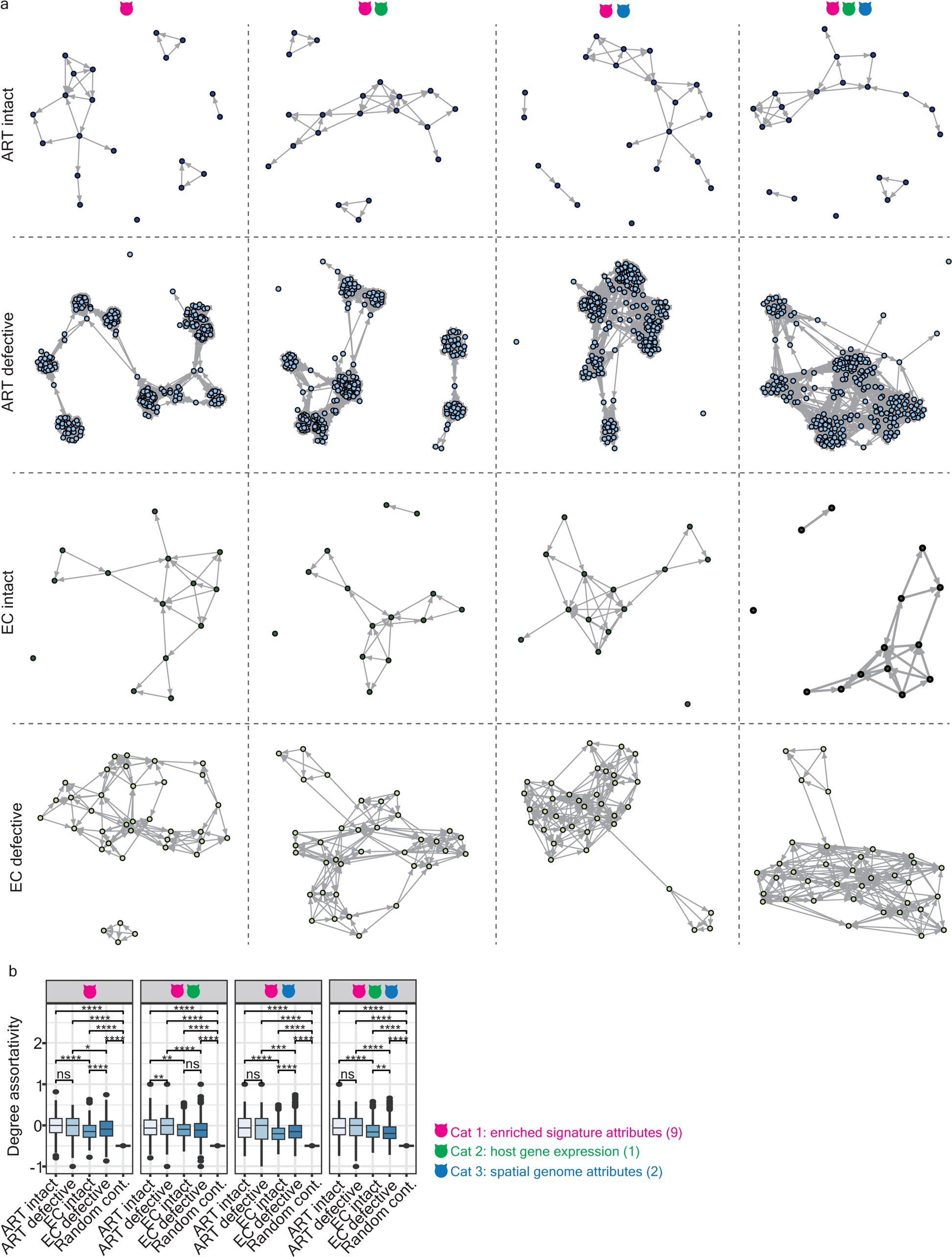
Topological properties of individual networks. (a) Graphs illustrating the topological properties of the network structured by enriched signatures (vertices) linked by edges, representing correlation coefficients using Cat 1, Cat 1 plus Cat 2, Cat1 plus Cat 3, and Cat 1, 2, and 3 attributes (ordered from left to right). The graphs, from top to bottom, illustrate the networks associated with intact and defective proviruses in ART-treated patients and elite controllers in a sequential order. Enriched signatures (vertices in the graphs) in reservoirs harboring intact and defective proviruses in ART-treated patients and elite controllers are labeled in dark blue, light blue, dark green, and light green, respectively. Predictor variables among different category attributes are described in detail in **Online Methods**. Vertices represent individual enriched signatures, while edges represent the correlation coefficient between two adjacent enriched signatures. (b) A box plot representing the degree assortativity coefficient of individual networks constructed based on different combinations of category attributes using the bootstrapping method detailed in **Online Methods**. Statistical significance was determined using the Wilcoxon test in R with default options. Significance levels are denoted as follows: * *p* 0.05, ** *p* 0.01, *** *p* 0.001, **** *p* 0.0001.

**Extended data Fig. 3.**
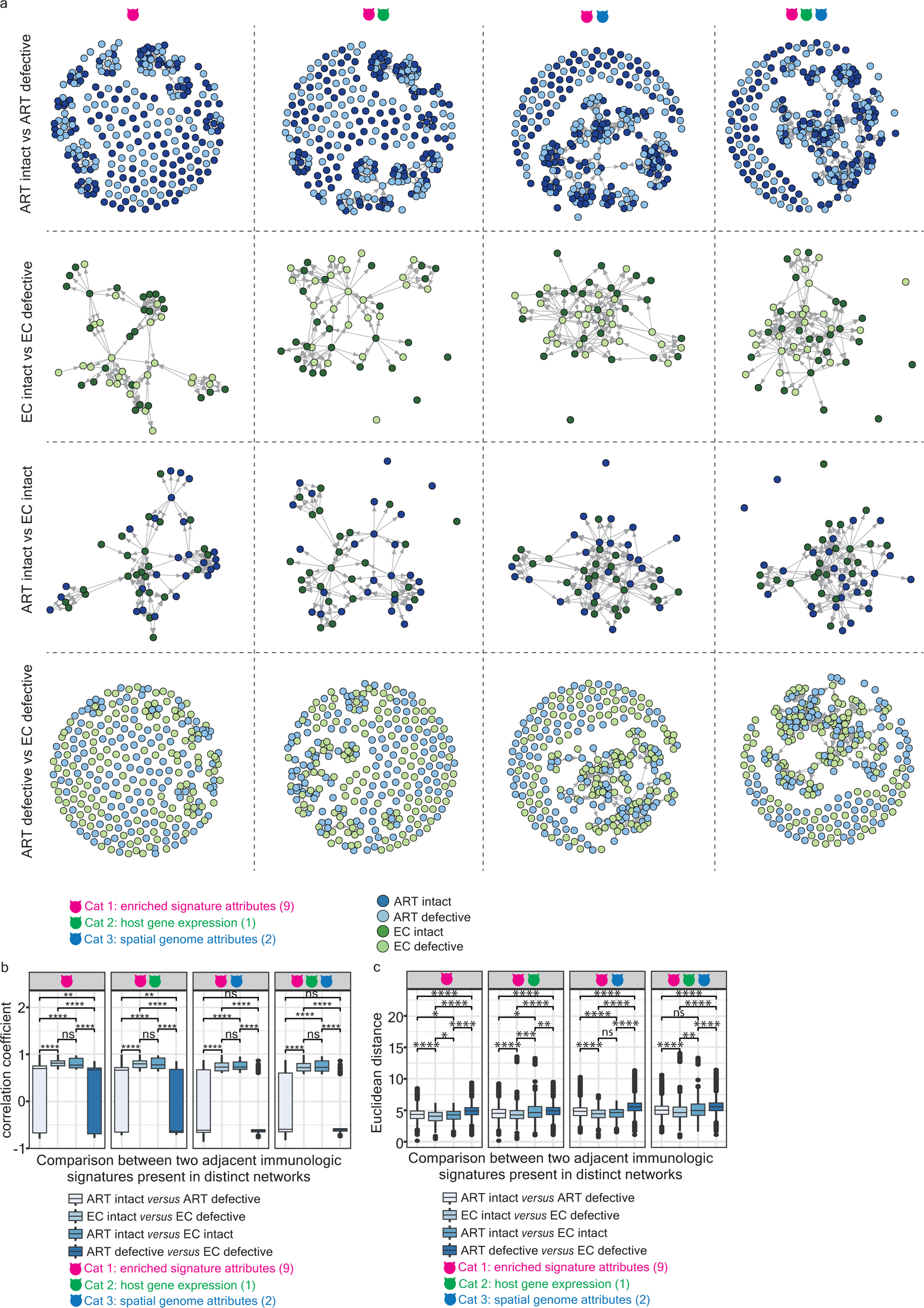
Topological interactions of two distinct networks. (a) Bipartite graphs illustrating the interaction between two adjacent enriched signatures present in distinct networks using Cat 1, Cat 1 plus Cat 2, Cat1 plus Cat 3, and Cat 1, 2, and 3 attributes (ordered from left to right). The graphs, from top to bottom, illustrate the topological interactions of the networks harboring intact (dark blue) *versus* defective (light blue) proviruses in ART-treated patients, the networks harboring intact (dark green) *versus* defective (light green) proviruses in elite controllers, the networks harboring intact proviruses in ART-treated patients (dark blue) *versus* elite controllers (dark green), and the networks harboring defective proviruses in ART-treated patients (light blue) *versus* elite controllers (light green). Predictor variables among different category attributes are described in detail in **Online Methods**. Vertices represent individual enriched signatures, and edges represent the correlation coefficient between two adjacent enriched signatures. (b) A box plot representing the correlation coefficients between two adjacent and distinct enriched signatures present in different networks using Cat 1, Cat 1 plus Cat 2, Cat1 plus Cat 3, and Cat 1, 2, and 3 attributes (ordered from left to right). Statistical significance was determined using the Wilcoxon test in R with default options. (c) A box plot representing the Euclidean distance between two adjacent and distinct enriched signatures present in different networks using Cat 1, Cat 1 plus Cat 2, Cat1 plus Cat 3, and Cat 1, 2, and 3 attributes (ordered from left to right). Statistical significance was determined using the Wilcoxon test in R with default options. Significance levels are denoted as follows: * *p* 0.05, ** *p* 0.01, *** *p* 0.001, **** *p* 0.0001.

**Extended data Fig. 4.**
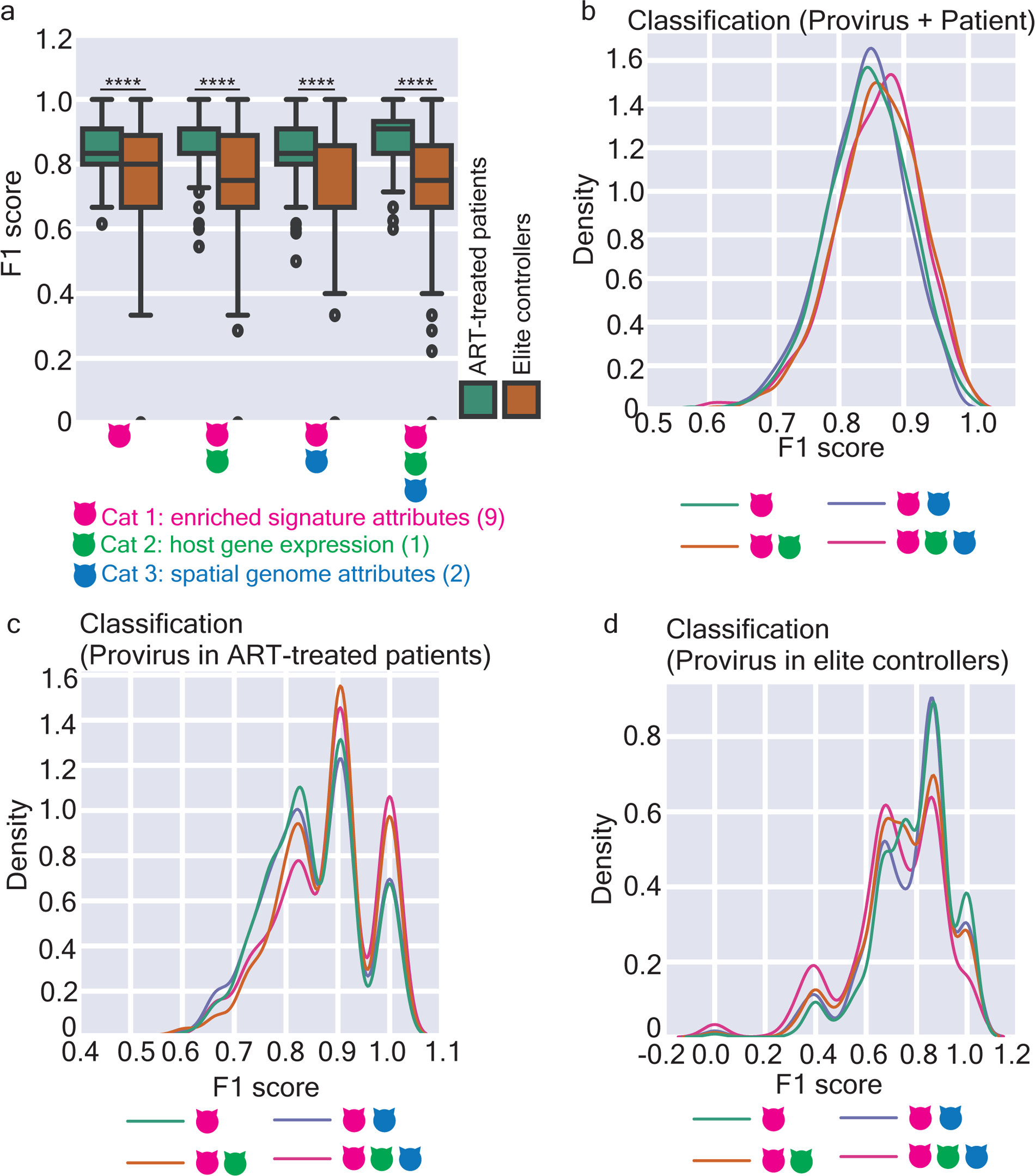
Random forest-based classifiers for predicting networks harboring intact *versus* defective proviruses. (a) A dodged box plot representing the accuracy of classification for networks associated with intact *versus* defective proviruses using random forest algorithms based on Cat 1, Cat 1 plus Cat 2, Cat1 plus Cat 3, and Cat 1, 2, and 3 attributes (ordered from left to right). Significance levels are denoted as follows: * *p* 0.05, ** *p* 0.01, *** *p* 0.001, **** *p* 0.0001. (b, c, d) A density plot representing the prediction quality of individual models constructed using different combinations of category attributes to classify networks associated with intact *versus* defective proviruses in ART-treated patients and elite controllers (b), networks associated with intact *versus* defective proviruses in ART-treated patients (c), and networks associated with intact *versus* defective proviruses in elite controllers (d). Interpretation of panels (c) and (d) is provided in **Online Methods**.

**Extended data Fig. 5.**
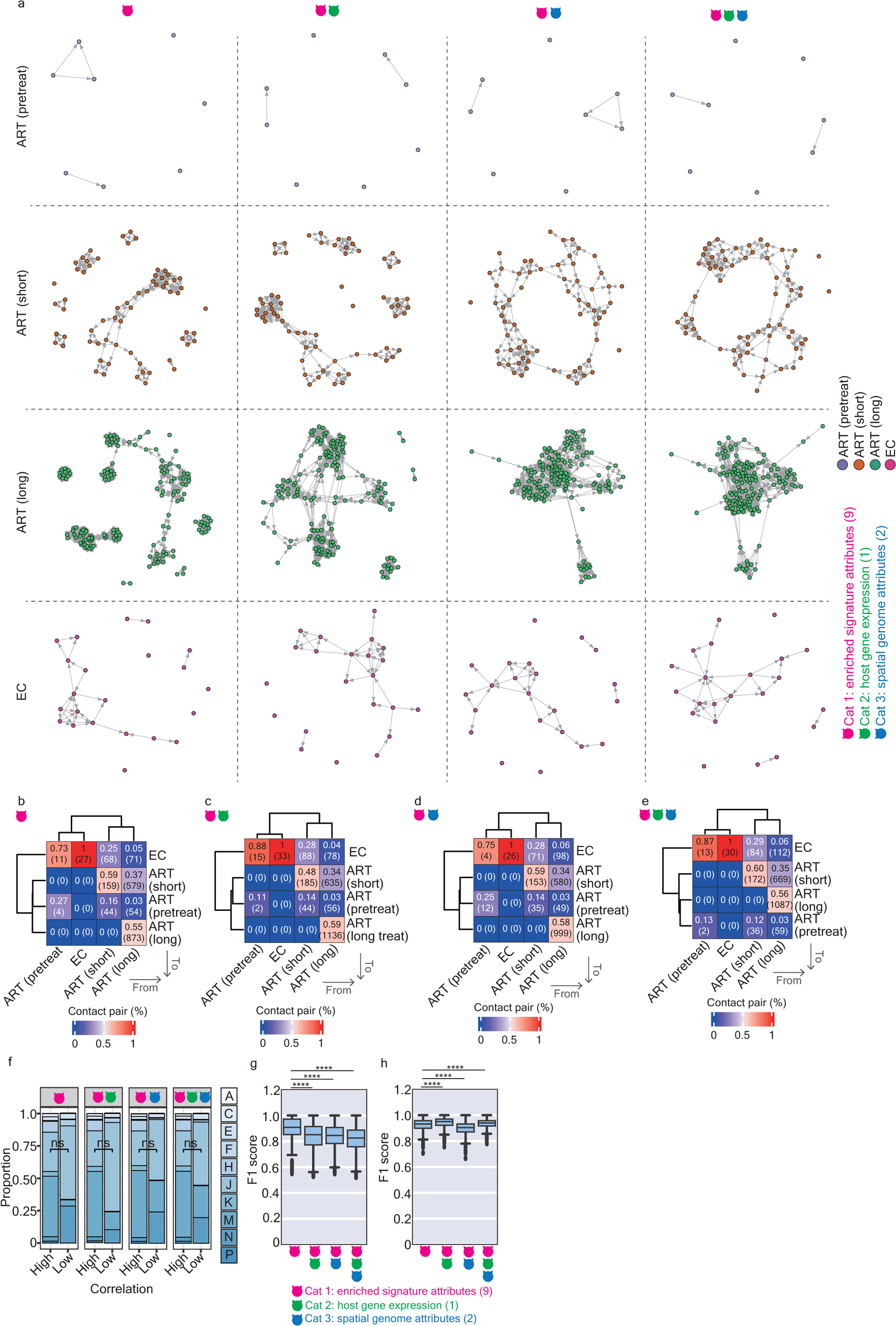
Longitudinal dynamics of the networks alongside HIV-1 infections associated with ART and in elite controllers. (a) Graphs illustrate the topological properties of the network structured by enriched signatures (vertices) linked by edges, representing correlation coefficients using Cat 1, Cat 1 plus Cat 2, Cat1 plus Cat 3, and Cat 1, 2, and 3 attributes (ordered from left to right). The graphs, from top to bottom, illustrate the networks structured by signatures enriched in pretreatment HIV-1-infected individuals, patients subjected to a short and long period of ART, and elite controllers. Enriched signatures (vertices in graphs) in pretreatment HIV-1-infected individuals, patients subjected to a short and a long period of ART, and elite controllers are labeled in violet, marron, green, and deep pink, respectively. Predictor variables among different category attributes are described in **Online Methods**. Vertices represent individual enriched signatures, while edges represent the correlation coefficient between two adjacent enriched signatures. (b, c, d, e) A clustering heatmap representing the percentage of two adjacent enriched signatures that were present in either the same or distinct networks using Cat 1, Cat 1 plus Cat 2, Cat 1 plus Cat 3, and Cat 1, 2, and 3 attributes (ordered from left to right). Parentheses placed in squares indicate the absolute number of walks. The color scale represents the percentage calculated by using the number of enriched signatures retrieved in rows, namely “To”, divided by the number of enriched signatures retrieved in columns, namely “From”. (f) A stacked bar plot representing the proportion of connected vertices in pairs associated with a high correlation coefficient (> −0.5) and those associated with a low correlation coefficient (< −0.5). (g) A box plot representing the prediction power for classifying networks in pretreatment HIV-1-infected individuals, patients subjected to a short and a long period of ART. The F1 score was calculated based on 1,000 times of individual train-test splits in models. (h) A box plot representing the prediction power for classifying networks in HIV-1-infected individuals subjected to a short and a long period of ART, and elite controllers. The F1 score was calculated based on 1,000 times of individual train-test splits in models. Significance levels are denoted as follows: * *p* 0.05, ** *p* 0.01, *** *p* 0.001, **** *p* 0.0001.

