## Appendix Table S1 for "Distinguishable topological properties of functional genome networks in HIV-1 reservoirs"

**Supplementary Table 1. List of enriched immunologic signatures harboring intact proviruses in ART-treated patients**

| ID | Description | GeneRatio | BgRatio | pvalue | p.adjust | qvalue | geneID | Count |
|---|---|---|---|---|---|---|---|---|
| NAKAYA_PBMC_FLUARIX_FLUVIRIN_AGE_18_50YO_CORRELATED_WITH_HAI_28DY_RESPONSE_AT_3DY_NEGATIVE | NAKAYA_PBMC_FLUARIX_FLUVIRIN_AGE_18_50YO_CORRELATED_WITH_HAI_28DY_RESPONSE_AT_3DY_NEGATIVE | 16/164 | 364/21355 | 2.3E-08 | 9.86E-05 | 9E-05 | 288/7109/54664/378938/51735/10782/3738/246175/3716/54856/51176/79066/9692/54665/7574/2521 | 16 |
| NAKAYA_PBMC_FLUARIX_FLUVIRIN_AGE_18_50YO_CORRELATED_WITH_HAI_28DY_RESPONSE_AT_7DY_NEGATIVE | NAKAYA_PBMC_FLUARIX_FLUVIRIN_AGE_18_50YO_CORRELATED_WITH_HAI_28DY_RESPONSE_AT_7DY_NEGATIVE | 17/164 | 475/21355 | 1.6E-07 | 0.000345 | 0.0003 | 288/6777/7109/378938/6774/10782/1997/84196/23042/11273/246175/9882/54856/51176/196528/23049/2521 | 17 |
| GSE5099_DAY3_VS_DAY7_MCSF_TREATED_MACROPHAGE_DN | GSE5099_DAY3_VS_DAY7_MCSF_TREATED_MACROPHAGE_DN | 10/164 | 183/21355 | 1.6E-06 | 0.002257 | 0.0021 | 1416/57690/378938/8418/23397/57459/246175/8832/51176/3183 | 10 |
| GSE40225_WT_VS_RIP_B7X_DIABETIC_MOUSE_PANCREATIC_CD8_TCELL_UP | GSE40225_WT_VS_RIP_B7X_DIABETIC_MOUSE_PANCREATIC_CD8_TCELL_UP | 10/164 | 199/21355 | 3.3E-06 | 0.00301 | 0.0028 | 64853/51466/63892/55770/9873/79230/54476/4306/28968/23049 | 10 |
| GSE2770_IL4_ACT_VS_ACT_CD4_TCELL_2H_DN | GSE2770_IL4_ACT_VS_ACT_CD4_TCELL_2H_DN | 10/164 | 200/21355 | 3.5E-06 | 0.00301 | 0.0028 | 116984/51735/23214/9135/64375/84166/10425/8178/4939/10613 | 10 |
| GSE21927_BALBC_VS_C57BL6_MONOCYTE_SPLEEN_UP | GSE21927_BALBC_VS_C57BL6_MONOCYTE_SPLEEN_UP | 9/164 | 189/21355 | 1.6E-05 | 0.010695 | 0.01 | 64853/10521/83852/1147/104/11273/51176/54811/28968 | 9 |
| GSE11057_CD4_CENT_MEM_VS_PBMC_UP | GSE11057_CD4_CENT_MEM_VS_PBMC_UP | 9/164 | 197/21355 | 2.2E-05 | 0.010695 | 0.01 | 89894/6777/55870/23048/2035/55690/8832/9882/90987 | 9 |
| GSE11057_NAIVE_CD4_VS_PBMC_CD4_TCELL_UP | GSE11057_NAIVE_CD4_VS_PBMC_CD4_TCELL_UP | 9/164 | 197/21355 | 2.2E-05 | 0.010695 | 0.01 | 6777/51466/9252/23214/246175/54856/79066/90987/54665 | 9 |
| GSE3982_MAST_CELL_VS_BCELL_DN | GSE3982_MAST_CELL_VS_BCELL_DN | 9/164 | 197/21355 | 2.2E-05 | 0.010695 | 0.01 | 51466/63892/953/8879/3738/10664/3716/9692/4939 | 9 |
| GSE2935_UV_INACTIVATED_VS_LIVE_SENDAI_VIRUS_INF_MACROPHAGE_DN | GSE2935_UV_INACTIVATED_VS_LIVE_SENDAI_VIRUS_INF_MACROPHAGE_DN | 8/164 | 179/21355 | 7.5E-05 | 0.032352 | 0.0303 | 953/493/10111/54476/11273/246175/3683/10613 | 8 |
| GSE32255_UNSTIM_VS_4H_LPS_STIM_DC_UP | GSE32255_UNSTIM_VS_4H_LPS_STIM_DC_UP | 8/164 | 186/21355 | 9.8E-05 | 0.034891 | 0.0326 | 953/23048/3738/140609/54476/8832/9882/4306 | 8 |
| GSE16450_CTRL_VS_IFNA_12H_STIM_IMMATURE_NEURON_CELL_LINE_UP | GSE16450_CTRL_VS_IFNA_12H_STIM_IMMATURE_NEURON_CELL_LINE_UP | 8/164 | 197/21355 | 0.00015 | 0.034891 | 0.0326 | 63892/9873/64754/23315/283450/2530/55809/2521 | 8 |
| GSE25087_TREG_VS_TCONV_FETUS_UP | GSE25087_TREG_VS_TCONV_FETUS_UP | 8/164 | 198/21355 | 0.00015 | 0.034891 | 0.0326 | 953/23048/81671/672/196/29110/64375/3683 | 8 |
| GSE11057_CD4_EFF_MEM_VS_PBMC_UP | GSE11057_CD4_EFF_MEM_VS_PBMC_UP | 8/164 | 199/21355 | 0.00016 | 0.034891 | 0.0326 | 288/89894/6777/51466/9252/3835/2035/54856 | 8 |
| GSE17974_1H_VS_72H_UNTREATED_IN_VITRO_CD4_TCELL_DN | GSE17974_1H_VS_72H_UNTREATED_IN_VITRO_CD4_TCELL_DN | 8/164 | 199/21355 | 0.00016 | 0.034891 | 0.0326 | 23048/90861/26263/10664/2108/2175/9978/10694 | 8 |
| GSE21927_C26GM_VS_4T1_TUMOR_MONOCYTE_BALBC_DN | GSE21927_C26GM_VS_4T1_TUMOR_MONOCYTE_BALBC_DN | 8/164 | 199/21355 | 0.00016 | 0.034891 | 0.0326 | 51466/7328/23315/1997/2035/6872/283450/81856 | 8 |
| GSE10240_IL22_VS_IL17_STIM_PRIMARY_BRONCHIAL_EPITHELIAL_CELLS_UP | GSE10240_IL22_VS_IL17_STIM_PRIMARY_BRONCHIAL_EPITHELIAL_CELLS_UP | 8/164 | 200/21355 | 0.00016 | 0.034891 | 0.0326 | 51735/1147/26263/84196/9978/29110/7700/54665 | 8 |
| GSE1460_DP_VS_CD4_THYMOCYTE_DN | GSE1460_DP_VS_CD4_THYMOCYTE_DN | 8/164 | 200/21355 | 0.00016 | 0.034891 | 0.0326 | 7699/51466/8879/104/9882/196/92822/4939 | 8 |
| GSE22601_IMMATURE_CD4_SINGLE_POSITIVE_VS_CD8_SINGLE_POSITIVE_THYMOCYTE_DN | GSE22601_IMMATURE_CD4_SINGLE_POSITIVE_VS_CD8_SINGLE_POSITIVE_THYMOCYTE_DN | 8/164 | 200/21355 | 0.00016 | 0.034891 | 0.0326 | 116984/8897/6774/51735/3091/54739/196/64375 | 8 |
| GSE5589_WT_VS_IL6_KO_LPS_AND_IL10_STIM_MACROPHAGE_45MIN_UP | GSE5589_WT_VS_IL6_KO_LPS_AND_IL10_STIM_MACROPHAGE_45MIN_UP | 8/164 | 200/21355 | 0.00016 | 0.034891 | 0.0326 | 7109/9738/3835/23397/90861/672/57153/10613 | 8 |
