## Appendix Table S2 for "Distinguishable topological properties of functional genome networks in HIV-1 reservoirs"

**Supplementary Table 2. List of enriched immunologic signatures harboring defective proviruses in ART-treated patients**

| ID | Description | GeneRatio | BgRatio | pvalue | p.adjust | qvalue | geneID | Count |
|---|---|---|---|---|---|---|---|---|
| GSE27241_WT_VS_RORGT_KO_TH17_POLARIZED_CD4_TCELL_TREATED_WITH_DIGOXIN_UP | GSE27241_WT_VS_RORGT_KO_TH17_POLARIZED_CD4_TCELL_TREATED_WITH_DIGOXIN_UP | 28/611 | 180/21355 | 3E-13 | 1.1E-09 | 8.2E-10 | 65125/171023/23085/1387/4763/9873/23731/9735/672/22992/2081/29028/8729/23139/26207/1060/9611/23499/5925/9918/8289/9520/5578/10788/65979/284058/5514/6654 | 28 |
| NAKAYA_PBMC_FLUARIX_FLUVIRIN_AGE_18_50YO_CORRELATED_WITH_HAI_28DY_RESPONSE_AT_7DY_NEGATIVE | NAKAYA_PBMC_FLUARIX_FLUVIRIN_AGE_18_50YO_CORRELATED_WITH_HAI_28DY_RESPONSE_AT_7DY_NEGATIVE | 46/611 | 475/21355 | 6E-13 | 1.1E-09 | 8.2E-10 | 6777/171023/60468/80264/4670/1540/1105/150864/6774/64848/4820/9840/2272/51696/84196/3480/5257/89846/79828/79813/23774/84458/3707/25981/255231/23141/4850/5295/1606/4026/57494/378938/26036/2521/23198/50852/5094/51176/7798/7109/163702/10198/9263/9129/7403/26043 | 46 |
| GSE37532_WT_VS_PPARG_KO_LN_TCONV_UP | GSE37532_WT_VS_PPARG_KO_LN_TCONV_UP | 29/611 | 200/21355 | 7E-13 | 1.1E-09 | 8.2E-10 | 60468/9736/55187/7173/57690/51696/91775/55619/22848/7182/1520/84458/65117/7745/5305/9466/9807/135112/23019/51735/80728/64750/114836/55758/51719/54842/163486/27244/7267 | 29 |
| GSE21927_C26GM_VS_4T1_TUMOR_MONOCYTE_BALBC_DN | GSE21927_C26GM_VS_4T1_TUMOR_MONOCYTE_BALBC_DN | 28/611 | 199/21355 | 4E-12 | 4.4E-09 | 3.3E-09 | 64766/1387/1540/1265/4820/81669/5048/22861/546/10006/51108/22848/54838/26207/23499/10163/1432/120/54878/10905/1739/2035/1911/1871/60685/9263/9459/7267 | 28 |
| OCONNOR_PBMC_MENVEO_ACWYVAX_AGE_30_70YO_7DY_AFTER_SECOND_DOSE_VS_7DY_AFTER_FIRST_DOSE_UP | OCONNOR_PBMC_MENVEO_ACWYVAX_AGE_30_70YO_7DY_AFTER_SECOND_DOSE_VS_7DY_AFTER_FIRST_DOSE_UP | 23/611 | 145/21355 | 3E-11 | 2.5E-08 | 1.9E-08 | 23085/55095/23633/22992/7322/84316/1729/5339/4967/10163/1778/51586/23248/6645/7109/284058/4134/124446/4791/23534/493/5514/10963 | 23 |
| NAKAYA_PBMC_FLUARIX_FLUVIRIN_AGE_18_50YO_CORRELATED_WITH_HAI_28DY_RESPONSE_AT_3DY_NEGATIVE | NAKAYA_PBMC_FLUARIX_FLUVIRIN_AGE_18_50YO_CORRELATED_WITH_HAI_28DY_RESPONSE_AT_3DY_NEGATIVE | 36/611 | 364/21355 | 1E-10 | 9.2E-08 | 6.8E-08 | 60468/10390/1105/253461/4820/9782/11320/546/84830/9728/79828/23774/83478/84458/3707/5775/3655/5295/10163/5588/65117/378938/2521/552900/50852/5094/114799/51176/51735/55818/7109/9263/79982/7403/84441/7267 | 36 |
| GSE25677_MPL_VS_R848_STIM_BCELL_DN | GSE25677_MPL_VS_R848_STIM_BCELL_DN | 24/611 | 181/21355 | 4E-10 | 3.1E-07 | 2.3E-07 | 10672/85464/60468/9873/23370/56913/1265/8826/1786/10163/1606/3660/4627/120/50650/3454/10403/7170/30844/10129/6197/9402/25853/84636 | 24 |
| GSE10325_LUPUS_CD4_TCELL_VS_LUPUS_BCELL_UP | GSE10325_LUPUS_CD4_TCELL_VS_LUPUS_BCELL_UP | 25/611 | 199/21355 | 6E-10 | 3.7E-07 | 2.7E-07 | 6777/1540/944/11320/9840/2272/51696/84830/3707/5775/23059/9797/3655/1786/51271/5295/8631/5588/1606/23102/5578/50852/51176/51735/493 | 25 |
| GSE13411_SWITCHED_MEMORY_BCELL_VS_PLASMA_CELL_UP | GSE13411_SWITCHED_MEMORY_BCELL_VS_PLASMA_CELL_UP | 25/611 | 200/21355 | 7E-10 | 3.7E-07 | 2.7E-07 | 51742/64421/1105/26065/55187/64848/55666/8500/1455/79828/23515/83478/10464/6894/11165/5295/1606/9712/259230/6197/57711/51735/55818/9129/79982 | 25 |
| GSE15624_CTRL_VS_3H_HALOFUGINONE_TREATED_CD4_TCELL_UP | GSE15624_CTRL_VS_3H_HALOFUGINONE_TREATED_CD4_TCELL_UP | 22/611 | 160/21355 | 1E-09 | 5.8E-07 | 4.3E-07 | 9873/23304/7046/23130/56913/4775/8826/91775/55619/1455/23186/3655/7920/1432/10425/55193/283209/117584/50852/10906/10198/10657 | 22 |
| GSE16450_IMMATURE_VS_MATURE_NEURON_CELL_LINE_UP | GSE16450_IMMATURE_VS_MATURE_NEURON_CELL_LINE_UP | 24/611 | 195/21355 | 2E-09 | 9.1E-07 | 6.8E-07 | 85464/23112/4763/23370/9267/4820/22861/25777/84166/89846/1455/84961/8705/3655/9960/5295/5588/120/378938/5165/284058/57337/388685/51317 | 24 |
| GSE14350_TREG_VS_TEFF_UP | GSE14350_TREG_VS_TEFF_UP | 24/611 | 199/21355 | 3E-09 | 1.1E-06 | 8.3E-07 | 56261/85464/51552/11329/9698/22828/55252/9728/1729/1203/84458/55870/9466/26036/29123/50852/51735/1399/63977/64750/55023/10198/4354/6654 | 24 |
| GSE2770_UNTREATED_VS_IL12_TREATED_ACT_CD4_TCELL_2H_UP | GSE2770_UNTREATED_VS_IL12_TREATED_ACT_CD4_TCELL_2H_UP | 24/611 | 200/21355 | 3E-09 | 1.1E-06 | 8.3E-07 | 6777/3936/7414/23214/6774/8826/3784/9797/1431/7009/4215/9693/4627/120/9918/157680/10788/10129/23604/6873/8031/8428/9459/2289 | 24 |

| | | | | | | | | |
|---|---|---|---|---|---|---|---|---|
| GSE45881_CXCR6HI_VS_CXCR1LO_COLONIC_LAMINA_PROPRIA_UP | GSE45881_CXCR6HI_VS_CXCR1LO_COLONIC_LAMINA_PROPRIA_UP | 24/611 | 200/21355 | 3E-09 | 1.1E-06 | 8.3E-07 | 4204/64848/4820/5048/8847/22861/6526/22887/4928/9967/22848/23499/1606/51586/26036/79745/23198/54870/23355/51735/6873/54521/10198/55758 | 24 |
| GSE9988_ANTI_TREM1_VS_CTRL_TREATED_MONOCYTES_DN | GSE9988_ANTI_TREM1_VS_CTRL_TREATED_MONOCYTES_DN | 24/611 | 200/21355 | 3E-09 | 1.1E-06 | 8.3E-07 | 377/11329/23369/51696/4297/89846/54838/9960/9611/1432/2885/8289/23102/54878/51/3454/155038/8031/25853/9020/55023/55758/51719/10521 | 24 |
| GSE11057_EFF_MEM_VS_CENT_MEM_CD4_TCELL_DN | GSE11057_EFF_MEM_VS_CENT_MEM_CD4_TCELL_DN | 23/611 | 188/21355 | 5E-09 | 1.5E-06 | 1.1E-06 | 85464/150864/9736/1119/55095/23077/200424/4297/5533/57634/50807/55870/4363/10144/753/23499/10055/9918/26036/51176/22866/55023/84441 | 23 |
| GSE32901_TH1_VS_TH17_ENRICHED_CD4_TCELL_UP | GSE32901_TH1_VS_TH17_ENRICHED_CD4_TCELL_UP | 22/611 | 174/21355 | 6E-09 | 1.7E-06 | 1.2E-06 | 85464/11329/23370/1105/117583/10006/1455/10048/1432/5588/1606/4627/120/3560/64333/55291/84937/51176/9402/55023/57337/60685 | 22 |
| NAKAYA_PBMC_FLUARIX_FLUVIRIN_AGE_18_50YO_3DY_DN | NAKAYA_PBMC_FLUARIX_FLUVIRIN_AGE_18_50YO_3DY_DN | 36/611 | 433/21355 | 1E-08 | 3.2E-06 | 2.4E-06 | 171023/80264/7799/54934/4670/23130/1540/4820/81669/22861/22887/155435/22992/51696/3480/378805/860/22848/3707/9960/9693/1606/65117/29123/5515/283209/2521/5770/54870/23060/5094/57711/283989/55818/7109/11184 | 36 |
| GSE32533_WT_VS_MIR17_KO_ACT_CD4_TCELL_DN | GSE32533_WT_VS_MIR17_KO_ACT_CD4_TCELL_DN | 23/611 | 197/21355 | 1E-08 | 3.2E-06 | 2.4E-06 | 85464/377/3936/23130/9267/4820/80196/23476/9851/7409/5925/3454/5090/7170/23527/10905/155038/84181/1315/284058/64750/10521/9459 | 23 |
| GSE10325_CD4_TCELL_VS_BCELL_UP | GSE10325_CD4_TCELL_VS_BCELL_UP | 23/611 | 198/21355 | 1E-08 | 3.3E-06 | 2.5E-06 | 6777/1540/944/11320/22848/3707/5775/23059/3655/1786/5295/8631/138151/5588/3560/23102/5305/5578/6197/50852/51176/493/2289 | 23 |
| GSE21063_CTRL_VS_ANTI_IGM_STIM_BCELL_NFATC1_KO_3H_DN | GSE21063_CTRL_VS_ANTI_IGM_STIM_BCELL_NFATC1_KO_3H_DN | 22/611 | 188/21355 | 2E-08 | 5.7E-06 | 4.2E-06 | 7799/65059/7456/55187/84301/81669/11320/55852/7182/5295/57494/148867/79745/7644/23198/9727/84181/7798/7109/64750/388685/23032 | 22 |
| GSE11057_CD4_CENT_MEM_VS_PBMC_UP | GSE11057_CD4_CENT_MEM_VS_PBMC_UP | 22/611 | 197/21355 | 6E-08 | 1.2E-05 | 8.7E-06 | 6777/171023/55690/51696/89845/4297/89846/257160/3707/55870/4363/23141/5588/1606/3710/23048/2035/1911/9020/9949/4012/11184 | 22 |
| GSE12366_GC_BCELL_VS_PLASMA_CELL_UP | GSE12366_GC_BCELL_VS_PLASMA_CELL_UP | 22/611 | 197/21355 | 6E-08 | 1.2E-05 | 8.7E-06 | 51742/65125/6777/60468/9736/8847/23369/701/84316/26528/55421/84458/10055/120/817/5090/5888/259230/7443/1399/388685/80025 | 22 |
| GSE16450_CTRL_VS_IFNA_12H_STIM_IMMATURE_NEURON_CELL_LINE_UP | GSE16450_CTRL_VS_IFNA_12H_STIM_IMMATURE_NEURON_CELL_LINE_UP | 22/611 | 197/21355 | 6E-08 | 1.2E-05 | 8.7E-06 | 171023/4763/9873/987/55187/641/3655/8631/5588/4026/2521/157680/5090/5578/1739/79613/9577/197135/5073/31/84441/5514 | 22 |
| GSE4984_LPS_VS_VEHICLE_CTRL_TREATED_DC_DN | GSE4984_LPS_VS_VEHICLE_CTRL_TREATED_DC_DN | 22/611 | 198/21355 | 6E-08 | 1.2E-05 | 8.8E-06 | 65125/4763/23304/4204/8826/2081/8073/965/2909/158358/50807/4967/23406/7009/5925/23102/148867/3594/23019/10788/57534/493 | 22 |
| GSE12366_PLASMA_CELL_VS_NAIVE_BCELL_DN | GSE12366_PLASMA_CELL_VS_NAIVE_BCELL_DN | 22/611 | 199/21355 | 7E-08 | 1.2E-05 | 8.8E-06 | 51742/51552/64421/10390/29028/84376/79828/10163/4627/120/8289/405/7170/10096/6197/1399/5205/22834/114836/9459/388685/4791 | 22 |
| GSE5542_UNTREATED_VS_IFNA_AND_IFNG_TREATED_EPITHELIAL_CELLS_6H_DN | GSE5542_UNTREATED_VS_IFNA_AND_IFNG_TREATED_EPITHELIAL_CELLS_6H_DN | 22/611 | 199/21355 | 7E-08 | 1.2E-05 | 8.8E-06 | 23132/4763/23130/6938/672/965/2909/84316/55421/7182/8705/79956/4627/9962/23102/405/5305/5770/115/4354/25938/163486 | 22 |
| GSE30083_SP1_VS_SP4_THYMOCYTE_DN | GSE30083_SP1_VS_SP4_THYMOCYTE_DN | 22/611 | 200/21355 | 7E-08 | 1.2E-05 | 8.8E-06 | 727/6774/944/91775/84166/4297/5533/1520/3655/22806/120/3560/976/148867/23248/6645/64750/10906/5073/55784/9953/27244 | 22 |
| GSE37533_PPARG1_FOXP3_VS_PPARG2_FOXP3_TRANSDUCED_CD4_TCELL_DN | GSE37533_PPARG1_FOXP3_VS_PPARG2_FOXP3_TRANSDUCED_CD4_TCELL_DN | 22/611 | 200/21355 | 7E-08 | 1.2E-05 | 8.8E-06 | 9202/23731/10390/7456/9267/81669/25777/57690/155435/51696/7322/23059/1432/10425/54165/11064/6197/27296/55683/23607/55023/114836 | 22 |

| | | | | | | | | |
|---|---|---|---|---|---|---|---|---|
| GSE411_WT_VS_SOCS3_KO_MACROPHAGE_DN | GSE411_WT_VS_SOCS3_KO_MACROPHAGE_DN | 22/611 | 200/21355 | 7E-08 | 1.2E-05 | 8.8E-06 | 9135/54934/4670/4204/7456/150864/25777/155435/23077/2909/6249/83478/50807/6767/3710/8289/92170/84181/23607/284058/10198/10657 | 22 |
| GSE9988_LOW_LPS_VS_CTRL_TREATED_MONOCYTE_DN | GSE9988_LOW_LPS_VS_CTRL_TREATED_MONOCYTE_DN | 22/611 | 200/21355 | 7E-08 | 1.2E-05 | 8.8E-06 | 23369/51696/8073/6780/80196/54838/9960/10163/54878/7769/155038/54870/10992/92912/55818/25853/9020/284058/55758/51719/10521/80331 | 22 |
| ANDERSON_BLOOD_CN54GP140_ADJUVANTED_WITH_GLA_AF_AGE_18_45YO_1DY_DN | ANDERSON_BLOOD_CN54GP140_ADJUVANTED_WITH_GLA_AF_AGE_18_45YO_1DY_DN | 14/611 | 85/21355 | 1E-07 | 1.9E-05 | 1.4E-05 | 7534/377/1387/22887/5976/7455/5977/4627/54878/10992/23355/23013/9577/8428 | 14 |
| GSE13411_IGM_MEMORY_BCELL_VS_PLASMA_CELL_UP | GSE13411_IGM_MEMORY_BCELL_VS_PLASMA_CELL_UP | 21/611 | 198/21355 | 3E-07 | 3.6E-05 | 2.6E-05 | 6777/60468/80264/7414/1105/22861/1729/83478/10464/3707/6894/11165/5976/10163/1606/5090/11064/23355/9844/9459/7403 | 21 |
| GSE16522_ANTI_CD3CD28_STIM_VS_UNSTIM_NAIVE_CD8_TCELL_DN | GSE16522_ANTI_CD3CD28_STIM_VS_UNSTIM_NAIVE_CD8_TCELL_DN | 21/611 | 198/21355 | 3E-07 | 3.6E-05 | 2.6E-05 | 55690/9267/1265/57690/29028/5257/1455/342945/55852/4967/6929/4627/5933/9918/549/2521/5770/7443/6891/57198/2590 | 21 |
| GSE36476_CTRL_VS_TSST_ACT_16H_MEMORY_CD4_TCELL_YOUNG_UP | GSE36476_CTRL_VS_TSST_ACT_16H_MEMORY_CD4_TCELL_YOUNG_UP | 21/611 | 198/21355 | 3E-07 | 3.6E-05 | 2.6E-05 | 54934/23370/23130/1105/1265/9736/25777/11320/9976/51696/5775/26207/9960/5295/8631/5588/120/7769/1911/51176/9459 | 21 |
| GSE11924_TFH_VS_TH1_CD4_TCELL_UP | GSE11924_TFH_VS_TH1_CD4_TCELL_UP | 21/611 | 199/21355 | 3E-07 | 3.6E-05 | 2.6E-05 | 1387/22828/1105/8826/25777/10006/2272/3480/9728/8705/23303/9960/4850/4627/405/23527/10096/51735/9020/23287/2590 | 21 |
| GSE25087_TREG_VS_TCONV_ADULT_UP | GSE25087_TREG_VS_TCONV_ADULT_UP | 21/611 | 199/21355 | 3E-07 | 3.6E-05 | 2.6E-05 | 8826/23347/84196/29028/5257/965/23150/701/55010/202052/4363/8631/5933/3560/3594/6645/57198/57534/493/84636/5906 | 21 |
| GSE37301_COMMON_LYMPHOID_PROGENITOR_VS_PRO_BCELL_DN | GSE37301_COMMON_LYMPHOID_PROGENITOR_VS_PRO_BCELL_DN | 21/611 | 199/21355 | 3E-07 | 3.6E-05 | 2.6E-05 | 4775/1265/64848/253461/8826/4820/9976/10036/1786/8904/120/6613/9807/5888/10788/4931/22834/55758/9844/9459/163486 | 21 |
| GSE39820_CTRL_VS_TGFBETA1_IL6_IL23A_CD4_TCELL_UP | GSE39820_CTRL_VS_TGFBETA1_IL6_IL23A_CD4_TCELL_UP | 21/611 | 199/21355 | 3E-07 | 3.6E-05 | 2.6E-05 | 51742/54934/23731/7456/26065/944/6526/23515/55870/22908/50650/283209/375748/259230/64783/54521/9844/23524/163486/7403/26043 | 21 |
| GSE45365_WT_VS_IFNAR_KO_CD8A_DC_MCMV_INFECTION_UP | GSE45365_WT_VS_IFNAR_KO_CD8A_DC_MCMV_INFECTION_UP | 21/611 | 199/21355 | 3E-07 | 3.6E-05 | 2.6E-05 | 9887/7414/823/4297/8729/23476/22848/1203/50807/23499/138151/51586/8289/23102/54815/4277/10956/23534/56942/473/26043 | 21 |
| GSE26488_CTRL_VS_PEPTIDE_INJECTION_OT2_THYMOCYTE_DN | GSE26488_CTRL_VS_PEPTIDE_INJECTION_OT2_THYMOCYTE_DN | 20/611 | 183/21355 | 3E-07 | 3.6E-05 | 2.6E-05 | 60468/9873/23370/3936/4204/56913/8826/89846/54899/10163/1606/3660/4627/120/8289/50650/7040/10403/10129/51735 | 20 |
| GSE21670_UNTREATED_VS_IL6_TREATED_CD4_TCELL_DN | GSE21670_UNTREATED_VS_IL6_TREATED_CD4_TCELL_DN | 21/611 | 200/21355 | 3E-07 | 3.6E-05 | 2.6E-05 | 23112/11329/55690/1265/55187/22992/22848/3707/50807/26207/3710/8289/54815/5305/3454/64333/10956/54617/64750/2590/9953 | 21 |
| GSE24210_CTRL_VS_IL35_TREATED_TCONV_CD4_TCELL_UP | GSE24210_CTRL_VS_IL35_TREATED_TCONV_CD4_TCELL_UP | 21/611 | 200/21355 | 3E-07 | 3.6E-05 | 2.6E-05 | 9267/81669/401409/23515/84458/7187/3655/4215/54899/5880/3660/3710/4026/9712/10129/55818/2175/54521/55784/51317/473 | 21 |
| GSE35685_CD34POS_CD38NEG_VS_CD34POS_CD10NEG_CD62LPOS_BONE_MARROW_UP | GSE35685_CD34POS_CD38NEG_VS_CD34POS_CD10NEG_CD62LPOS_BONE_MARROW_UP | 21/611 | 200/21355 | 3E-07 | 3.6E-05 | 2.6E-05 | 65125/28977/7414/987/26065/8826/9782/23077/4733/23476/721/5339/4215/10425/4700/7251/115/51719/2643/473/7267 | 21 |
| GSE45739_UNSTIM_VS_ACD3_ACD28_STIM_NRAS_KO_CD4_TCELL_UP | GSE45739_UNSTIM_VS_ACD3_ACD28_STIM_NRAS_KO_CD4_TCELL_UP | 21/611 | 200/21355 | 3E-07 | 3.6E-05 | 2.6E-05 | 6777/171023/1540/81669/9840/2272/2926/10892/3655/10144/6767/1606/5578/10129/23355/50852/1911/51176/9949/10800/473 | 21 |
| GSE38696_LIGHT_ZONE_VS_DARK_ZONE_BCELL_UP | GSE38696_LIGHT_ZONE_VS_DARK_ZONE_BCELL_UP | 20/611 | 184/21355 | 3E-07 | 3.8E-05 | 2.8E-05 | 2869/11329/23370/55187/23077/79813/84961/26207/7409/23499/5588/549/5305/283209/23198/23365/8065/51735/84186/10521 | 20 |
| GSE15330_HSC_VS_MEGAKARYOCYTE_ERYTHROID_PROGENITOR_DN | GSE15330_HSC_VS_MEGAKARYOCYTE_ERYTHROID_PROGENITOR_DN | 19/611 | 168/21355 | 4E-07 | 3.9E-05 | 2.9E-05 | 60468/9873/944/23077/3480/51108/79828/79663/3157/4215/22955/146691/1606/50852/84937/51735/23607/27244/7267 | 19 |

| | | | | | | | | |
|---|---|---|---|---|---|---|---|---|
| QI_PBMC_ZOSTAVAX_AGE_50_75YO_CORRELATED_WITH_CONTRACTION_OF_VZV_SPECIFIC_T_CELLS_PEAK_TO_28DY_AT_1DYPOSITIVE | QI_PBMC_ZOSTAVAX_AGE_50_75YO_CORRELATED_WITH_CONTRACTION_OF_VZV_SPECIFIC_T_CELLS_PEAK_TO_28DY_AT_1DYPOSITIVE | 26/611 | 294/21355 | 4E-07 | 4.2E-05 | 3.1E-05 | 1387/3936/1105/6774/4820/3106/155435/80196/51271/5880/4627/8289/23048/976/79801/23198/23527/155038/5094/23013/8031/25853/55023/51719/10800/473 | 26 |
| GSE1925_CTRL_VS_IFNG_PRIMED_MACROPHAGE_3H_IFNG_STIM_UP | GSE1925_CTRL_VS_IFNG_PRIMED_MACROPHAGE_3H_IFNG_STIM_UP | 19/611 | 174/21355 | 6E-07 | 6.4E-05 | 4.7E-05 | 27340/944/23047/6249/7182/1520/84961/4627/57649/2961/55291/56852/6645/51735/55500/54617/114836/54842/7403 | 19 |
| GSE9601_UNTREATED_VS_NFKB_INHIBITOR_TREATED_HCMV_INF_MONOCYTE_DN | GSE9601_UNTREATED_VS_NFKB_INHIBITOR_TREATED_HCMV_INF_MONOCYTE_DN | 19/611 | 174/21355 | 6E-07 | 6.4E-05 | 4.7E-05 | 51742/9135/4763/9202/253461/5257/6249/22848/84961/9693/22806/50650/51/64333/8672/6873/55275/7267/23032 | 19 |
| GSE7219_UNSTIM_VS_LPS_AND_ANTI_CD40_STIM_NIK_NFKB2_KO_DC_UP | GSE7219_UNSTIM_VS_LPS_AND_ANTI_CD40_STIM_NIK_NFKB2_KO_DC_UP | 19/611 | 178/21355 | 9E-07 | 8.9E-05 | 6.6E-05 | 60468/9267/4775/944/58513/5295/5588/1606/120/9466/259230/10788/10129/51176/23607/114836/9844/51317/9953 | 19 |
| GSE13411_NAIVE_BCELL_VS_PLASMA_CELL_UP | GSE13411_NAIVE_BCELL_VS_PLASMA_CELL_UP | 20/611 | 196/21355 | 1E-06 | 9E-05 | 6.7E-05 | 2869/60468/9202/64421/4775/23064/6894/11165/5976/23499/1606/9466/976/11064/23355/1911/9263/9459/31/7403 | 20 |
| GSE23321_CD8_STEM_CELL_MEMORY_VS_NAIVE_CD8_TCELL_DN | GSE23321_CD8_STEM_CELL_MEMORY_VS_NAIVE_CD8_TCELL_DN | 20/611 | 196/21355 | 1E-06 | 9E-05 | 6.7E-05 | 85464/23112/4763/23130/11320/55252/29028/5257/257160/8705/79956/5880/5933/196074/10403/10905/2035/55758/124446/25938 | 20 |
| GSE15330_HSC_VS_MEGAKARYOCYTE_ERYTHROID_PROGENITOR_UP | GSE15330_HSC_VS_MEGAKARYOCYTE_ERYTHROID_PROGENITOR_UP | 18/611 | 162/21355 | 1E-06 | 9E-05 | 6.7E-05 | 85464/60468/1540/3480/2909/79828/79663/4215/22955/146691/9466/50852/84937/51176/51735/9402/27244/7267 | 18 |
| GSE25088_CTRL_VS_IL4_AND_ROSIGLITAZONE_STIM_STAT6_KO_MACROPHAGE_UP | GSE25088_CTRL_VS_IL4_AND_ROSIGLITAZONE_STIM_STAT6_KO_MACROPHAGE_UP | 20/611 | 197/21355 | 1E-06 | 9.3E-05 | 6.9E-05 | 23201/23731/54834/3069/29028/3480/91775/23150/2909/7009/5880/9918/405/54878/6599/1739/10198/10521/2590/7267 | 20 |
| GSE9988_ANTI_TREM1_VS_VEHICLE_TREATED_MONOCYTES_DN | GSE9988_ANTI_TREM1_VS_VEHICLE_TREATED_MONOCYTES_DN | 20/611 | 197/21355 | 1E-06 | 9.3E-05 | 6.9E-05 | 11329/25777/11320/51696/8073/6794/54838/23406/4215/1432/2885/6613/9466/155038/8031/25853/22834/8428/54521/10521 | 20 |
| GSE10325_CD4_TCELL_VS_MYELOID_UP | GSE10325_CD4_TCELL_VS_MYELOID_UP | 20/611 | 199/21355 | 1E-06 | 9.9E-05 | 7.3E-05 | 1540/11320/9840/2272/3707/5775/23059/3655/1786/10055/5295/8631/6767/5588/1606/3560/5900/5578/50852/51176 | 20 |
| GSE12845_IGD_POS_BLOOD_VS_DARKZONE_GC_TONSIL_BCELL_DN | GSE12845_IGD_POS_BLOOD_VS_DARKZONE_GC_TONSIL_BCELL_DN | 20/611 | 199/21355 | 1E-06 | 9.9E-05 | 7.3E-05 | 6777/171023/60468/3936/9736/6780/3707/10055/3183/65117/5165/30844/2961/5578/54870/259230/7251/23013/55758/4012 | 20 |
| GSE21927_SPLEEN_C57BL6_VS_EL4_TUMOR_BALBC_MONOCYTES_UP | GSE21927_SPLEEN_C57BL6_VS_EL4_TUMOR_BALBC_MONOCYTES_UP | 20/611 | 199/21355 | 1E-06 | 9.9E-05 | 7.3E-05 | 1387/9736/81669/5048/9976/23077/55252/10006/4297/23774/22848/5533/9960/23499/1432/120/65117/54878/10905/2035 | 20 |
| GSE24574_BCL6_HIGH_VS_LOW_TFH_CD4_TCELL_UP | GSE24574_BCL6_HIGH_VS_LOW_TFH_CD4_TCELL_UP | 20/611 | 199/21355 | 1E-06 | 9.9E-05 | 7.3E-05 | 7534/51742/9873/7799/23370/9267/9736/1119/9782/25777/7554/22992/2272/3707/8705/9693/1606/120/9712/9263 | 20 |
| GSE27670_BLIMP1_VS_LMP1_TRANSDUCED_GC_BCELL_DN | GSE27670_BLIMP1_VS_LMP1_TRANSDUCED_GC_BCELL_DN | 20/611 | 199/21355 | 1E-06 | 9.9E-05 | 7.3E-05 | 377/64324/7046/1265/23633/3069/23369/10006/22955/79745/5165/5090/64333/5888/9274/1399/55500/10198/4354/9694 | 20 |
| GSE16385_ROSIGLITAZONE_IL4_VS_IFNG_TNF_STIM_MACROPHAGE_UP | GSE16385_ROSIGLITAZONE_IL4_VS_IFNG_TNF_STIM_MACROPHAGE_UP | 20/611 | 200/21355 | 1E-06 | 9.9E-05 | 7.3E-05 | 51742/171023/60468/1387/22992/89845/55619/55253/5775/26207/8631/10479/120/3560/23102/157680/51719/2643/9129/79982 | 20 |
| GSE3039_CD4_TCELL_VS_NKT_CELL_UP | GSE3039_CD4_TCELL_VS_NKT_CELL_UP | 20/611 | 200/21355 | 1E-06 | 9.9E-05 | 7.3E-05 | 85464/6777/7414/8826/401409/337867/51696/84306/5257/84376/7182/6337/79956/23141/8289/155038/11064/51176/57337/163486 | 20 |
| GSE35825_IFNA_VS_IFNG_STIM_MACROPHAGE_DN | GSE35825_IFNA_VS_IFNG_STIM_MACROPHAGE_DN | 20/611 | 200/21355 | 1E-06 | 9.9E-05 | 7.3E-05 | 85464/23112/90273/54934/4775/9889/23064/6938/23369/546/11165/1786/23141/283209/64333/10788/10129/8672/57198/163486 | 20 |
| GSE6674_ANTI_IGM_VS_CPG_STIM_BCELL_UP | GSE6674_ANTI_IGM_VS_CPG_STIM_BCELL_UP | 20/611 | 200/21355 | 1E-06 | 9.9E-05 | 7.3E-05 | 11329/23370/23131/4775/150864/51696/50807/84961/4215/120/5305/283209/9807/10403/23248/259230/23604/57337/60685/27244 | 20 |

| | | | | | | | | |
|---|---|---|---|---|---|---|---|---|
| GSE7460_FOXP3_MUT_VS_WT_ACT_WITH_TGFB_TCONV_DN | GSE7460_FOXP3_MUT_VS_WT_ACT_WITH_TGFB_TCONV_DN | 20/611 | 200/21355 | 1E-06 | 9.9E-05 | 7.3E-05 | 85464/6777/11329/150864/155435/9967/26207/23141/50650/9466/976/5578/50852/51176/115/57337/124446/7403/473/27244 | 20 |
| KAECH_NAIVE_VS_DAY8_EFF_CD8_TCELL_UP | KAECH_NAIVE_VS_DAY8_EFF_CD8_TCELL_UP | 20/611 | 200/21355 | 1E-06 | 9.9E-05 | 7.3E-05 | 23731/57585/23077/3480/4297/51108/6249/1520/6223/4850/1606/1289/7813/5305/84937/51176/23450/473/27244/7267 | 20 |
| GSE25146_UNSTIM_VS_HELIOBACTER_PYLORI_LPS_STIM_AGS_CELL_DN | GSE25146_UNSTIM_VS_HELIOBACTER_PYLORI_LPS_STIM_AGS_CELL_DN | 18/611 | 166/21355 | 1E-06 | 0.0001 | 7.6E-05 | 51742/4820/81669/23077/55619/84166/10892/84961/55870/23102/64333/11064/84937/9402/10521/4791/51317/23032 | 18 |
| GSE37301_PRO_BCELL_VS_CD4_TCELL_UP | GSE37301_PRO_BCELL_VS_CD4_TCELL_UP | 18/611 | 170/21355 | 2E-06 | 0.00014 | 0.00011 | 4763/7799/944/51072/59269/91775/6894/157378/55972/5588/5933/50650/5770/6599/64333/80728/64750/6641 | 18 |
| GSE27896_HDAC6_KO_VS_WT_TREG_UP | GSE27896_HDAC6_KO_VS_WT_TREG_UP | 18/611 | 176/21355 | 3E-06 | 0.00023 | 0.00017 | 23370/9698/22828/23064/25777/10006/7182/3707/5339/23303/7409/6767/22806/9466/3594/83891/22870/63977 | 18 |
| GSE11386_NAIVE_VS_MEMORY_BCELL_DN | GSE11386_NAIVE_VS_MEMORY_BCELL_DN | 17/611 | 159/21355 | 3E-06 | 0.00023 | 0.00017 | 64766/60468/64324/6938/23077/84166/378805/23293/50807/55870/23384/5578/8672/51176/7982/84186/55500 | 17 |
| GSE10273_HIGH_VS_LOW_IL7_TREATED_IRF4_8_NULL_PRE_BCELL_DN | GSE10273_HIGH_VS_LOW_IL7_TREATED_IRF4_8_NULL_PRE_BCELL_DN | 19/611 | 198/21355 | 4E-06 | 0.00029 | 0.00021 | 132789/6777/11329/23130/150864/55187/23064/57690/727957/7187/7409/1778/3710/10956/4700/64750/114836/84433/7267 | 19 |
| GSE21927_SPLENIC_VS_TUMOR_MONOCYTES_FROM_C26GM_TUMOROUS_MICE_BALBC_DN | GSE21927_SPLENIC_VS_TUMOR_MONOCYTES_FROM_C26GM_TUMOROUS_MICE_BALBC_DN | 19/611 | 198/21355 | 4E-06 | 0.00029 | 0.00021 | 64766/1540/4820/81669/5048/22887/23077/546/10006/51108/23774/79663/54838/9611/23499/120/10905/7109/11184 | 19 |
| GSE10325_LUPUS_CD4_TCELL_VS_LUPUS_MYELOID_UP | GSE10325_LUPUS_CD4_TCELL_VS_LUPUS_MYELOID_UP | 19/611 | 199/21355 | 5E-06 | 0.00029 | 0.00021 | 6777/60468/9840/2272/7322/5533/5775/9797/3655/8631/6767/5588/3710/3560/5578/1911/51176/51735/9402 | 19 |
| GSE11057_NAIVE_VS_CENT_MEMORY_CD4_TCELL_DN | GSE11057_NAIVE_VS_CENT_MEMORY_CD4_TCELL_DN | 19/611 | 199/21355 | 5E-06 | 0.00029 | 0.00021 | 377/5496/7414/8826/965/2909/257160/4967/4363/23406/9693/51586/4026/135112/10403/30844/10788/493/163486 | 19 |
| GSE13306_LAMINA_PROPRIA_VS_SPLEEN_TREG_UP | GSE13306_LAMINA_PROPRIA_VS_SPLEEN_TREG_UP | 19/611 | 199/21355 | 5E-06 | 0.00029 | 0.00021 | 171023/60468/1387/64324/3683/4245/23150/7009/5976/51586/5933/5770/5090/7170/155038/259230/84636/5514/27244 | 19 |
| GSE21670_UNTREATED_VS_TGFB_TREATED_STAT3_KO_CD4_TCELL_DN | GSE21670_UNTREATED_VS_TGFB_TREATED_STAT3_KO_CD4_TCELL_DN | 19/611 | 199/21355 | 5E-06 | 0.00029 | 0.00021 | 56261/9873/9267/1540/944/23633/401409/9840/7421/50807/165918/5588/817/5770/22866/55500/10906/11184/5906 | 19 |
| GSE22886_NAIVE_CD8_TCELL_VS_MONOCYTE_UP | GSE22886_NAIVE_CD8_TCELL_VS_MONOCYTE_UP | 19/611 | 199/21355 | 5E-06 | 0.00029 | 0.00021 | 60468/4204/987/11320/8287/5775/3655/23499/8631/5588/3710/3560/5578/57711/51176/51735/9402/9949/104 | 19 |
| GSE22886_NAIVE_TCELL_VS_DC_UP | GSE22886_NAIVE_TCELL_VS_DC_UP | 19/611 | 199/21355 | 5E-06 | 0.00029 | 0.00021 | 6777/171023/80264/11329/987/26065/22861/22887/23347/23077/4297/5533/8631/5588/1606/120/3710/7644/51176 | 19 |
| GSE26495_NAIVE_VS_PD1LOW_CD8_TCELL_DN | GSE26495_NAIVE_VS_PD1LOW_CD8_TCELL_DN | 19/611 | 199/21355 | 5E-06 | 0.00029 | 0.00021 | 3683/9976/8073/965/2909/5775/255231/4026/3560/4277/8742/5900/8672/6645/115/4134/114836/55784/493 | 19 |
| GSE2770_IL12_VS_TGFB_AND_IL12_TREATED_ACT_CD4_TCELL_6H_DN | GSE2770_IL12_VS_TGFB_AND_IL12_TREATED_ACT_CD4_TCELL_6H_DN | 19/611 | 199/21355 | 5E-06 | 0.00029 | 0.00021 | 26574/944/22861/4733/701/23059/1060/7409/9611/1432/5977/2885/65117/8289/8672/64750/55023/56942/163486 | 19 |
| GSE12392_CD8A_POS_VS_NEG_SPLEEN_DC_DN | GSE12392_CD8A_POS_VS_NEG_SPLEEN_DC_DN | 19/611 | 200/21355 | 5E-06 | 0.00029 | 0.00021 | 171023/23370/9735/7414/672/25777/3106/23150/701/84166/23139/5339/10615/10055/1778/9918/79801/10788/10800 | 19 |
| GSE17301_CTRL_VS_48H_ACD3_ACD28_IFNA5_STIM_CD8_TCELL_UP | GSE17301_CTRL_VS_48H_ACD3_ACD28_IFNA5_STIM_CD8_TCELL_UP | 19/611 | 200/21355 | 5E-06 | 0.00029 | 0.00021 | 56261/23112/65059/4204/2081/546/10048/8289/1954/50650/5305/259230/51735/79157/4354/25938/51317/473/27244 | 19 |
| GSE21546_WT_VS_SAP1A_KO_AND_ELK1_KO_DP_THYMOCYTES_DN | GSE21546_WT_VS_SAP1A_KO_AND_ELK1_KO_DP_THYMOCYTES_DN | 19/611 | 200/21355 | 5E-06 | 0.00029 | 0.00021 | 60468/9873/26065/117583/155435/26207/23048/54165/51/155038/10129/1911/7798/65979/57198/4134/2643/27244/7267 | 19 |

| | | | | | | | | |
|---|---|---|---|---|---|---|---|---|
| GSE22601_IMMATURE_CD4_SINGLE_POSITIVE_VS_DOUBLE_POSITIVE_THYMOCYTE_DN | GSE22601_IMMATURE_CD4_SINGLE_POSITIVE_VS_DOUBLE_POSITIVE_THYMOCYTE_DN | 19/611 | 200/21355 | 5E-06 | 0.00029 | 0.00021 | 56261/85464/7414/150864/59269/11320/51696/79718/55619/23150/22848/50807/6767/22806/5305/7040/6197/5514/7267 | 19 |
| GSE30962_ACUTE_VS_CHRONIC_LCMV_SECONDARY_INF_CD8_TCELL_UP | GSE30962_ACUTE_VS_CHRONIC_LCMV_SECONDARY_INF_CD8_TCELL_UP | 19/611 | 200/21355 | 5E-06 | 0.00029 | 0.00021 | 85464/11329/55690/57585/57690/51696/26207/753/9960/23048/283209/9807/10788/10129/51176/22834/51317/84441/473 | 19 |
| GSE31082_DN_VS_CD8_SP_THYMOCYTE_DN | GSE31082_DN_VS_CD8_SP_THYMOCYTE_DN | 19/611 | 200/21355 | 5E-06 | 0.00029 | 0.00021 | 65125/85464/1387/7456/9267/150864/10892/23515/7187/9611/5880/50650/83891/9402/283131/4012/55784/80331/473 | 19 |
| GSE39820_CTRL_VS_TGFBETA3_IL6_IL23A_CD4_TCELL_UP | GSE39820_CTRL_VS_TGFBETA3_IL6_IL23A_CD4_TCELL_UP | 19/611 | 200/21355 | 5E-06 | 0.00029 | 0.00021 | 51742/56261/26065/23369/51696/23515/9851/50807/22908/51455/50650/259230/10992/92181/1871/54521/79692/9844/27244 | 19 |
| GSE5542_UNTREATED_VS_IFNG_TREATED_EPITHELIAL_CELLS_6H_DN | GSE5542_UNTREATED_VS_IFNG_TREATED_EPITHELIAL_CELLS_6H_DN | 19/611 | 200/21355 | 5E-06 | 0.00029 | 0.00021 | 377/4763/3936/965/2909/84316/55421/8705/79956/4627/9962/23102/4277/5305/7040/1739/115/25938/163486 | 19 |
| KAECH_NAIVE_VS_MEMORY_CD8_TCELL_UP | KAECH_NAIVE_VS_MEMORY_CD8_TCELL_UP | 19/611 | 200/21355 | 5E-06 | 0.00029 | 0.00021 | 65125/64766/54934/23731/9736/57585/3069/3480/4297/51108/6767/1289/5305/84937/10198/4354/473/27244/7267 | 19 |
| GSE15330_LYMPHOID_MULTIPOTENT_VS_MEGAKARYOCYTE_ERYTHROID_PROGENITOR_IKAROS_KO_UP | GSE15330_LYMPHOID_MULTIPOTENT_VS_MEGAKARYOCYTE_ERYTHROID_PROGENITOR_IKAROS_KO_UP | 17/611 | 166/21355 | 6E-06 | 0.00032 | 0.00024 | 60468/57690/3480/51108/79828/79663/4215/22955/146691/9466/50852/84937/51176/51735/23607/27244/7267 | 17 |
| GSE14699_NAIVE_VS_DELETIONAL_TOLERANCE_CD8_TCELL_UP | GSE14699_NAIVE_VS_DELETIONAL_TOLERANCE_CD8_TCELL_UP | 16/611 | 152/21355 | 8E-06 | 0.00042 | 0.00031 | 23201/944/57690/84166/2909/79828/79663/84961/55870/9466/324/50852/51176/51317/27244/7267 | 16 |
| NAKAYA_MONOCYTE_FLUMIST_AGE_18_50YO_7DY_UP | NAKAYA_MONOCYTE_FLUMIST_AGE_18_50YO_7DY_UP | 32/611 | 478/21355 | 9E-06 | 0.00047 | 0.00035 | 9135/545/10390/7456/146057/23064/5048/22992/6249/1203/10464/54838/3655/55972/6929/10144/79956/23141/9611/9466/54878/7769/23019/4700/1739/23163/7109/55500/2289/2590/5514/7267 | 32 |
| GSE21063_CTRL_VS_ANTI_IGM_STIM_BCELL_NFATC1_KO_8H_UP | GSE21063_CTRL_VS_ANTI_IGM_STIM_BCELL_NFATC1_KO_8H_UP | 18/611 | 191/21355 | 1E-05 | 0.00054 | 0.0004 | 60468/23131/23130/1105/1119/84301/81669/23774/3707/84961/9693/5295/7040/23198/23163/23013/7798/64750 | 18 |
| GSE7348_LPS_VS_TOLERIZED_AND_LPS_STIM_MACROPHAGE_DN | GSE7348_LPS_VS_TOLERIZED_AND_LPS_STIM_MACROPHAGE_DN | 17/611 | 174/21355 | 1E-05 | 0.00058 | 0.00043 | 1540/944/23077/80196/257160/7920/5588/1778/3594/5165/30844/65979/23607/54521/51317/6641/6654 | 17 |
| GSE32255_WT_UNSTIM_VS_JMJD2D_KNOCKDOWN_4H_LPS_STIM_DC_UP | GSE32255_WT_UNSTIM_VS_JMJD2D_KNOCKDOWN_4H_LPS_STIM_DC_UP | 17/611 | 175/21355 | 1E-05 | 0.00062 | 0.00046 | 60468/1387/4763/7799/56913/944/11320/3480/22848/6894/5588/57649/9712/4700/55818/120425/51317 | 17 |
| GSE7568_IL4_TGFB_DEXAMETHASONE_VS_IL4_TGFB_TREATED_MACROPHAGE_DN | GSE7568_IL4_TGFB_DEXAMETHASONE_VS_IL4_TGFB_TREATED_MACROPHAGE_DN | 17/611 | 175/21355 | 1E-05 | 0.00062 | 0.00046 | 132789/85464/23304/4775/823/9728/257160/3707/79956/51271/54878/283209/23527/6645/51317/9953/51720 | 17 |
| GSE16385_IL4_VS_ROSIGLITAZONE_STIM_MACROPHAGE_UP | GSE16385_IL4_VS_ROSIGLITAZONE_STIM_MACROPHAGE_UP | 18/611 | 194/21355 | 1E-05 | 0.00064 | 0.00048 | 7414/8826/23369/29028/79718/3480/23150/2909/7009/79956/23499/9918/196074/157680/6599/11163/2590/7267 | 18 |
| GSE16450_CTRL_VS_IFNA_6H_STIM_IMMATURE_NEURON_CELL_LINE_DN | GSE16450_CTRL_VS_IFNA_6H_STIM_IMMATURE_NEURON_CELL_LINE_DN | 18/611 | 195/21355 | 1E-05 | 0.00068 | 0.00051 | 64766/11329/26157/25777/9976/84166/84961/8705/26207/5295/57494/50650/5165/1911/388685/493/10800/27244 | 18 |
| GSE11057_PBMC_VS_MEM_CD4_TCELL_DN | GSE11057_PBMC_VS_MEM_CD4_TCELL_DN | 18/611 | 197/21355 | 2E-05 | 0.00071 | 0.00053 | 51742/6777/23370/9267/944/5820/51696/4297/257160/5533/5588/3710/50852/2035/1911/9020/57198/9949 | 18 |
| GSE22886_NAIVE_CD4_TCELL_VS_MONOCYTE_UP | GSE22886_NAIVE_CD4_TCELL_VS_MONOCYTE_UP | 18/611 | 197/21355 | 2E-05 | 0.00071 | 0.00053 | 23112/60468/987/11320/9840/5775/3655/5295/8631/5588/1606/3710/5578/50852/57711/51176/9949/27244 | 18 |
| GSE22886_CD4_TCELL_VS_BCELL_NAIVE_UP | GSE22886_CD4_TCELL_VS_BCELL_NAIVE_UP | 18/611 | 198/21355 | 2E-05 | 0.00071 | 0.00053 | 65125/253461/3683/2926/3707/5775/26207/10055/5295/51586/9466/976/5900/5578/10788/50852/51176/2289 | 18 |
| GSE22886_CD8_TCELL_VS_BCELL_NAIVE_UP | GSE22886_CD8_TCELL_VS_BCELL_NAIVE_UP | 18/611 | 198/21355 | 2E-05 | 0.00071 | 0.00053 | 253461/3683/6780/5775/26207/138151/5588/51586/3560/9466/5578/51176/9402/10198/493/9673/163486/2289 | 18 |

| | | | | | | | | |
|---|---|---|---|---|---|---|---|---|
| GSE22886_NAIVE_TCELL_VS_MONOCYTE_UP | GSE22886_NAIVE_TCELL_VS_MONOCYTE_UP | 18/611 | 198/21355 | 2E-05 | 0.00071 | 0.00053 | 171023/60468/987/11320/9797/3655/8631/5588/1606/3710/5578/57711/51176/55818/9402/9949/10198/27244 | 18 |
| GSE29618_PRE_VS_DAY7_FLU_VACCINE_MDC_DN | GSE29618_PRE_VS_DAY7_FLU_VACCINE_MDC_DN | 18/611 | 198/21355 | 2E-05 | 0.00071 | 0.00053 | 55252/3784/79718/91775/79230/7322/79663/6894/23499/22908/138151/1606/92170/54870/8672/7982/79632/5514 | 18 |
| GSE7348_UNSTIM_VS_TOLERIZED_AND_LPS_STIM_MACROPHAGE_UP | GSE7348_UNSTIM_VS_TOLERIZED_AND_LPS_STIM_MACROPHAGE_UP | 16/611 | 162/21355 | 2E-05 | 0.00071 | 0.00053 | 64324/22828/585/5820/6938/57585/57690/79813/23303/23048/51460/79613/64783/7109/115/27244 | 16 |
| GSE10094_LCMV_VS_LISTERIA_IND_EFF_CD4_TCELL_UP | GSE10094_LCMV_VS_LISTERIA_IND_EFF_CD4_TCELL_UP | 18/611 | 199/21355 | 2E-05 | 0.00071 | 0.00053 | 65125/51552/11329/10390/150864/3683/5048/155435/55619/197322/84316/23515/9693/23443/148867/7109/8428/51720 | 18 |
| GSE1460_INTRATHYMIC_T_PROGENITOR_VS_CD4_THYMOCYTE_DN | GSE1460_INTRATHYMIC_T_PROGENITOR_VS_CD4_THYMOCYTE_DN | 18/611 | 199/21355 | 2E-05 | 0.00071 | 0.00053 | 60468/7414/22861/3106/3707/9797/23406/79956/7409/8631/5880/3560/9466/30844/2643/9459/79632/104 | 18 |
| GSE17721_CTRL_VS_CPG_0.5H_BMDC_UP | GSE17721_CTRL_VS_CPG_0.5H_BMDC_UP | 18/611 | 199/21355 | 2E-05 | 0.00071 | 0.00053 | 377/171023/90273/23370/23167/4775/7187/3655/26207/8904/1432/5977/9918/976/128866/9949/55758/79657 | 18 |
| GSE22886_NAIVE_CD8_TCELL_VS_DC_UP | GSE22886_NAIVE_CD8_TCELL_VS_DC_UP | 18/611 | 199/21355 | 2E-05 | 0.00071 | 0.00053 | 6777/60468/80264/11329/26574/9267/987/9782/22887/4297/5775/5588/3710/3560/5305/57711/9402/104 | 18 |
| GSE22886_TCELL_VS_BCELL_NAIVE_UP | GSE22886_TCELL_VS_BCELL_NAIVE_UP | 18/611 | 199/21355 | 2E-05 | 0.00071 | 0.00053 | 6777/5496/3683/2926/3707/3655/26207/10055/5588/1606/51586/9466/976/5578/9274/51176/22870/2289 | 18 |
| GSE2770_IL12_VS_IL4_TREATED_ACT_CD4_TCELL_48H_DN | GSE2770_IL12_VS_IL4_TREATED_ACT_CD4_TCELL_48H_DN | 18/611 | 199/21355 | 2E-05 | 0.00071 | 0.00053 | 4204/22828/4775/23064/5820/200424/4297/2909/9851/50807/5339/7187/54899/3710/3560/84181/6197/23607 | 18 |
| GSE32986_GMCSF_AND_CURDLAN_LOWDOSE_VS_GMCSF_AND_CURDLAN_HIGHDOSE_STIM_DC_UP | GSE32986_GMCSF_AND_CURDLAN_LOWDOSE_VS_GMCSF_AND_CURDLAN_HIGHDOSE_STIM_DC_UP | 18/611 | 199/21355 | 2E-05 | 0.00071 | 0.00053 | 85464/11329/23731/64324/65059/64848/55252/727957/79813/23303/138151/5933/3077/324/10906/10198/51317/124565 | 18 |
| GSE35435_RESTING_VS_IL4_TREATED_MACROPHAGE_UP | GSE35435_RESTING_VS_IL4_TREATED_MACROPHAGE_UP | 18/611 | 199/21355 | 2E-05 | 0.00071 | 0.00053 | 65125/23370/146057/4820/89845/8073/4297/79663/53944/7187/7813/4026/23443/7769/7644/79982/27244/6654 | 18 |
| GSE37301_COMMON_LYMPHOID_PROGENITOR_VS_CD4_TCELL_DN | GSE37301_COMMON_LYMPHOID_PROGENITOR_VS_CD4_TCELL_DN | 18/611 | 199/21355 | 2E-05 | 0.00071 | 0.00053 | 4775/64848/253461/9976/7322/1455/10036/5977/4627/9807/5090/5888/10788/4931/5094/9844/163486/6654 | 18 |
| GSE40273_GATA1_KO_VS_WT_TREG_DN | GSE40273_GATA1_KO_VS_WT_TREG_DN | 18/611 | 199/21355 | 2E-05 | 0.00071 | 0.00053 | 60468/9202/55690/23370/9267/25777/10892/5339/120/4026/9466/54165/29123/5770/23527/155038/29761/84433 | 18 |
| GSE5542_UNTREATED_VS_IFNA_TREATED_EPITHELIAL_CELLS_6H_UP | GSE5542_UNTREATED_VS_IFNA_TREATED_EPITHELIAL_CELLS_6H_UP | 18/611 | 199/21355 | 2E-05 | 0.00071 | 0.00053 | 6777/377/8826/823/2081/965/23139/4967/23406/3560/57649/7040/135112/7798/283989/1822/493/10800 | 18 |
| GSE6092_UNSTIM_VS_IFNG_STIM_ENDOTHELIAL_CELL_DN | GSE6092_UNSTIM_VS_IFNG_STIM_ENDOTHELIAL_CELL_DN | 18/611 | 199/21355 | 2E-05 | 0.00071 | 0.00053 | 23112/60468/1265/25777/29980/337867/84196/5339/26207/1060/8631/1606/50650/26036/157680/2035/54521/4012 | 18 |
| GSE7460_CD8_TCELL_VS_TREG_ACT_DN | GSE7460_CD8_TCELL_VS_TREG_ACT_DN | 18/611 | 199/21355 | 2E-05 | 0.00071 | 0.00053 | 23181/7799/4670/23047/701/10615/1060/10055/5925/5770/5165/10403/7443/7251/50852/23607/8428/473 | 18 |
| GSE10239_MEMORY_VS_KLRG1HIGH_EFF_CD8_TCELL_DN | GSE10239_MEMORY_VS_KLRG1HIGH_EFF_CD8_TCELL_DN | 18/611 | 200/21355 | 2E-05 | 0.00071 | 0.00053 | 7534/171023/51552/23047/58513/23150/701/51108/2926/753/9693/10479/8904/6599/10096/11163/128866/9694 | 18 |
| GSE12003_MIR223_KO_VS_WT_BM_PROGENITOR_4D_CULTURE_UP | GSE12003_MIR223_KO_VS_WT_BM_PROGENITOR_4D_CULTURE_UP | 18/611 | 200/21355 | 2E-05 | 0.00071 | 0.00053 | 60468/54934/7046/9736/6938/23077/4297/22848/4215/51455/51176/51735/9577/115/55500/8428/23287/7267 | 18 |
| GSE17721_0.5H_VS_12H_GARDIQUIMOD_BMDC_DN | GSE17721_0.5H_VS_12H_GARDIQUIMOD_BMDC_DN | 18/611 | 200/21355 | 2E-05 | 0.00071 | 0.00053 | 9698/9889/81669/80196/23515/3707/23293/54899/7409/1606/57410/5933/10905/8672/117584/8726/51317/51720 | 18 |
| GSE17721_PAM3CSK4_VS_GADIQUIMOD_8H_BMDC_DN | GSE17721_PAM3CSK4_VS_GADIQUIMOD_8H_BMDC_DN | 18/611 | 200/21355 | 2E-05 | 0.00071 | 0.00053 | 51742/23214/6774/672/4297/7322/83478/10048/4795/5295/5977/1606/5933/54815/23198/117584/128866/55500 | 18 |

| | | | | | | | | |
|---|---|---|---|---|---|---|---|---|
| GSE22601_CD4_SINGLE_POSITIVE_VS_CD8_SINGLE_POSITIVE_THYMOCYTE_DN | GSE22601_CD4_SINGLE_POSITIVE_VS_CD8_SINGLE_POSITIVE_THYMOCYTE_DN | 18/611 | 200/21355 | 2E-05 | 0.00071 | 0.00053 | 7534/57102/54834/401409/155435/337867/3707/3655/1606/3660/2885/65117/29123/155038/4700/57337/114836/55784 | 18 |
| GSE24671_CTRL_VS_BAKIMULC_INFECTED_MOUSE_SPLENOCYTES_UP | GSE24671_CTRL_VS_BAKIMULC_INFECTED_MOUSE_SPLENOCYTES_UP | 18/611 | 200/21355 | 2E-05 | 0.00071 | 0.00053 | 85464/90273/10390/9267/83478/84458/26207/8631/4627/7040/23248/84181/259230/56852/9402/54521/51719/51317 | 18 |
| GSE31082_CD4_VS_CD8_SP_THYMOCYTE_UP | GSE31082_CD4_VS_CD8_SP_THYMOCYTE_UP | 18/611 | 200/21355 | 2E-05 | 0.00071 | 0.00053 | 23085/23214/26065/9736/51696/91775/9728/22848/54838/9693/5305/50852/23607/55758/10521/9459/54842/5906 | 18 |
| GSE31082_DN_VS_CD4_SP_THYMOCYTE_UP | GSE31082_DN_VS_CD4_SP_THYMOCYTE_UP | 18/611 | 200/21355 | 2E-05 | 0.00071 | 0.00053 | 11052/56005/4670/586/79718/10111/84316/26528/55421/51068/165918/729852/549/10767/2961/7443/7251/114836 | 18 |
| GSE33162_HDAC3_KO_VS_HDAC3_KO_4H_LPS_STIM_MACROPHAGE_UP | GSE33162_HDAC3_KO_VS_HDAC3_KO_4H_LPS_STIM_MACROPHAGE_UP | 18/611 | 200/21355 | 2E-05 | 0.00071 | 0.00053 | 150864/22887/23369/23077/79718/84316/26528/6794/79036/3183/65117/5305/54870/1399/64783/128866/51719/7403 | 18 |
| GSE39820_TGFBETA1_IL6_VS_TGFBETA1_IL6_IL23A_TREATED_CD4_TCELL_UP | GSE39820_TGFBETA1_IL6_VS_TGFBETA1_IL6_IL23A_TREATED_CD4_TCELL_UP | 18/611 | 200/21355 | 2E-05 | 0.00071 | 0.00053 | 51742/10672/23131/7046/3064/8826/5048/89970/11320/10111/5295/22806/26036/5090/1911/55683/80728/10800 | 18 |
| GSE41867_DAY6_EFFECTOR_VS_DAY30_MEMORY_CD8_TCELL_LCMV_ARMSTRONG_DN | GSE41867_DAY6_EFFECTOR_VS_DAY30_MEMORY_CD8_TCELL_LCMV_ARMSTRONG_DN | 18/611 | 200/21355 | 2E-05 | 0.00071 | 0.00053 | 23112/7799/10390/64324/9267/8826/3683/860/84458/23293/9611/4627/324/23248/84181/54521/51719/51317 | 18 |
| GSE5589_LPS_AND_IL10_VS_LPS_AND_IL6_STIM_MACROPHAGE_45MIN_DN | GSE5589_LPS_AND_IL10_VS_LPS_AND_IL6_STIM_MACROPHAGE_45MIN_DN | 18/611 | 200/21355 | 2E-05 | 0.00071 | 0.00053 | 28977/7414/337867/641/55619/55010/80196/11165/2521/3594/10096/10906/10198/2643/83990/11184/163486/84636 | 18 |
| GSE7460_CTRL_VS_TGFB_TREATED_ACT_FOXP3_HET_TCONV_UP | GSE7460_CTRL_VS_TGFB_TREATED_ACT_FOXP3_HET_TCONV_UP | 18/611 | 200/21355 | 2E-05 | 0.00071 | 0.00053 | 8844/11052/23132/23214/150864/5048/8073/10111/4363/51455/3607/3560/10767/83891/2961/5094/8726/31 | 18 |
| GSE9006_HEALTHY_VS_TYPE_1_DIABETES_PBMC_1MONTH_POST_DX_UP | GSE9006_HEALTHY_VS_TYPE_1_DIABETES_PBMC_1MONTH_POST_DX_UP | 18/611 | 200/21355 | 2E-05 | 0.00071 | 0.00053 | 7534/11052/4670/26574/26065/9782/8073/6780/55421/10144/5295/3607/23443/8672/6891/6645/5094/79982 | 18 |
| GSE9650_NAIVE_VS_MEMORY_CD8_TCELL_UP | GSE9650_NAIVE_VS_MEMORY_CD8_TCELL_UP | 18/611 | 200/21355 | 2E-05 | 0.00071 | 0.00053 | 64766/54934/23731/9736/57585/3069/3480/4297/51108/1520/5305/10767/54870/84937/10198/4354/27244/7267 | 18 |
| GSE10147_IL3_VS_IL3_AND_HIVP17_STIM_PDC_UP | GSE10147_IL3_VS_IL3_AND_HIVP17_STIM_PDC_UP | 16/611 | 166/21355 | 2E-05 | 0.00088 | 0.00065 | 57521/7414/146057/641/53944/84961/23059/5976/5925/23248/114804/23355/23163/22870/2643/23032 | 16 |
| NAKAYA_MYELOID_DENDRITIC_CELL_FLUARIX_FLUVIRIN_AGE_18_50YO_7DY_DN | NAKAYA_MYELOID_DENDRITIC_CELL_FLUARIX_FLUVIRIN_AGE_18_50YO_7DY_DN | 21/611 | 269/21355 | 3E-05 | 0.00126 | 0.00094 | 10672/7799/4255/987/7326/5257/23515/3183/23048/405/10905/23060/10454/1739/1911/9949/10198/55758/493/54842/7267 | 21 |
| GSE27291_0H_VS_7D_STIM_GAMMADELTA_TCELL_DN | GSE27291_0H_VS_7D_STIM_GAMMADELTA_TCELL_DN | 17/611 | 190/21355 | 3E-05 | 0.00127 | 0.00094 | 1387/4763/3683/23347/8287/10048/1606/3710/405/5305/51460/79745/29123/64333/55023/54521/163486 | 17 |
| GSE14415_INDUCED_TREG_VS_FOXP3_KO_INDUCED_TREG_UP | GSE14415_INDUCED_TREG_VS_FOXP3_KO_INDUCED_TREG_UP | 16/611 | 172/21355 | 4E-05 | 0.00133 | 0.00098 | 944/23064/155435/2272/51108/79828/6249/8289/9466/50852/84937/51176/51735/23607/27244/7267 | 16 |
| GSE14415_FOXP3_KO_NATURAL_TREG_VS_TCONV_DN | GSE14415_FOXP3_KO_NATURAL_TREG_VS_TCONV_DN | 16/611 | 173/21355 | 4E-05 | 0.00141 | 0.00105 | 85464/944/57690/23369/10111/51108/79828/22955/50852/84937/51176/51735/23607/114836/27244/7267 | 16 |
| GSE21670_IL6_VS_TGFB_AND_IL6_TREATED_CD4_TCELL_UP | GSE21670_IL6_VS_TGFB_AND_IL6_TREATED_CD4_TCELL_UP | 17/611 | 193/21355 | 4E-05 | 0.00152 | 0.00113 | 11329/55690/27340/26157/11320/2272/22848/3655/26207/9960/1606/259283/155038/5578/51176/51735/84441 | 17 |
| GSE18893_TCONV_VS_TREG_2H_TNF_STIM_DN | GSE18893_TCONV_VS_TREG_2H_TNF_STIM_DN | 17/611 | 194/21355 | 5E-05 | 0.00161 | 0.00119 | 1387/23167/3106/11320/57690/23077/51696/8500/84458/57649/7769/7644/155038/10129/8065/10521/388685 | 17 |
| GSE17974_IL4_AND_ANTI_IL12_VS_UNTREATED_72H_ACT_CD4_TCELL_DN | GSE17974_IL4_AND_ANTI_IL12_VS_UNTREATED_72H_ACT_CD4_TCELL_DN | 17/611 | 195/21355 | 5E-05 | 0.0017 | 0.00126 | 8844/7456/9976/91775/965/4297/26207/9693/83891/23198/155038/5888/10788/50852/84766/2643/84636 | 17 |
| GSE13485_CTRL_VS_DAY1_YF17D_VACCINE_PBMC_UP | GSE13485_CTRL_VS_DAY1_YF17D_VACCINE_PBMC_UP | 17/611 | 196/21355 | 5E-05 | 0.00178 | 0.00132 | 1073/55526/23167/9567/55010/6780/2926/165918/23186/10479/54165/5515/117584/1399/64783/9377/9673 | 17 |

| | | | | | | | | |
|---|---|---|---|---|---|---|---|---|
| GSE17974_0.5H_VS_72H_IL4_AND_ANTI_IL12_ACT_CD4_TCELL_DN | GSE17974_0.5H_VS_72H_IL4_AND_ANTI_IL12_ACT_CD4_TCELL_DN | 17/611 | 196/21355 | 5E-05 | 0.00178 | 0.00132 | 586/79602/4245/965/84316/51108/342945/3183/817/5690/23019/9577/2175/10906/83990/11184/10963 | 17 |
| GSE9960_HEALTHY_VS_GRAM_NEG_AND_POS_SEPSIS_PBMC_DN | GSE9960_HEALTHY_VS_GRAM_NEG_AND_POS_SEPSIS_PBMC_DN | 17/611 | 196/21355 | 5E-05 | 0.00178 | 0.00132 | 8844/9873/64324/987/9889/22992/546/84376/965/80196/5295/26036/54165/157680/11064/117584/9263 | 17 |
| GSE26030_TH1_VS_TH17_DAY5_POST_POLARIZATION_DN | GSE26030_TH1_VS_TH17_DAY5_POST_POLARIZATION_DN | 17/611 | 197/21355 | 6E-05 | 0.00185 | 0.00137 | 3936/1119/5820/3683/59269/83478/7187/4363/23406/9962/5305/976/3594/92181/2175/31/2289 | 17 |
| GSE22342_CD11C_HIGH_VS_LOW_DECIDUAL_MACROPHAGES_UP | GSE22342_CD11C_HIGH_VS_LOW_DECIDUAL_MACROPHAGES_UP | 17/611 | 198/21355 | 6E-05 | 0.00185 | 0.00137 | 9267/3683/29028/965/202052/11165/9611/1432/4627/3077/23102/5690/55291/8672/7109/55023/84636 | 17 |
| GSE22886_NAIVE_CD8_TCELL_VS_NKCELL_UP | GSE22886_NAIVE_CD8_TCELL_VS_NKCELL_UP | 17/611 | 198/21355 | 6E-05 | 0.00185 | 0.00137 | 1265/944/50514/7326/4733/2926/10892/4363/22955/6767/1606/5578/11064/51176/51719/104/2289 | 17 |
| GSE25087_TREG_VS_TCONV_FETUS_UP | GSE25087_TREG_VS_TCONV_FETUS_UP | 17/611 | 198/21355 | 6E-05 | 0.00185 | 0.00137 | 8844/4763/727/3683/672/55010/10144/10055/7813/3560/23048/283209/57534/51501/9953/6654/81671 | 17 |
| GSE13484_UNSTIM_VS_3H_YF17D_VACCINE_STIM_PBMC_UP | GSE13484_UNSTIM_VS_3H_YF17D_VACCINE_STIM_PBMC_UP | 17/611 | 199/21355 | 6E-05 | 0.00185 | 0.00137 | 11052/23370/146057/987/6938/9782/55852/1729/6929/10163/54878/1739/23013/22870/60685/9459/79982 | 17 |
| GSE17721_0.5H_VS_8H_GARDIQUIMOD_BMDC_DN | GSE17721_0.5H_VS_8H_GARDIQUIMOD_BMDC_DN | 17/611 | 199/21355 | 6E-05 | 0.00185 | 0.00137 | 1073/23167/6774/9567/81669/3106/4297/23515/3707/5933/4026/50650/148867/5515/29761/54870/283131 | 17 |
| GSE22886_NAIVE_CD4_TCELL_VS_12H_ACT_TH2_UP | GSE22886_NAIVE_CD4_TCELL_VS_12H_ACT_TH2_UP | 17/611 | 199/21355 | 6E-05 | 0.00185 | 0.00137 | 6777/1387/9736/55187/4820/22861/23077/3707/3710/7644/1739/27296/5094/1911/7109/55758/7267 | 17 |
| GSE22886_NAIVE_CD8_TCELL_VS_MEMORY_TCELL_DN | GSE22886_NAIVE_CD8_TCELL_VS_MEMORY_TCELL_DN | 17/611 | 199/21355 | 6E-05 | 0.00185 | 0.00137 | 51552/1540/51696/26528/7009/9960/120/5933/54815/51460/23527/1399/57198/8428/51719/5906/27244 | 17 |
| GSE22886_NAIVE_VS_MEMORY_TCELL_DN | GSE22886_NAIVE_VS_MEMORY_TCELL_DN | 17/611 | 199/21355 | 6E-05 | 0.00185 | 0.00137 | 7534/6777/9782/2081/51696/23150/26528/5933/6613/54815/23527/30844/50852/57198/51719/5906/27244 | 17 |
| GSE43863_LY6C_INT_CXCR5POS_VS_LY6C_LOW_CXCR5NEG_EFFECTOR_CD4_TCELL_UP | GSE43863_LY6C_INT_CXCR5POS_VS_LY6C_LOW_CXCR5NEG_EFFECTOR_CD4_TCELL_UP | 17/611 | 199/21355 | 6E-05 | 0.00185 | 0.00137 | 9873/27031/9267/57690/22992/51696/3784/8287/3707/6929/10163/8289/9577/163702/55023/54521/4012 | 17 |
| GSE10273_LOW_IL7_VS_HIGH_IL7_AND_IRF4_IN_IRF4_8_NULL_PRE_BCELL_DN | GSE10273_LOW_IL7_VS_HIGH_IL7_AND_IRF4_IN_IRF4_8_NULL_PRE_BCELL_DN | 17/611 | 200/21355 | 7E-05 | 0.00185 | 0.00137 | 64766/1387/11329/10390/55187/727957/7187/23141/7409/5977/3710/25862/55683/22834/64750/79157/2590 | 17 |
| GSE16450_CTRL_VS_IFNA_6H_STIM_MATURE_NEURON_CELL_LINE_DN | GSE16450_CTRL_VS_IFNA_6H_STIM_MATURE_NEURON_CELL_LINE_DN | 17/611 | 200/21355 | 7E-05 | 0.00185 | 0.00137 | 7534/8844/23112/3683/337867/8073/55666/3707/6794/4627/8289/378938/54815/5690/3454/10956/54870 | 17 |
| GSE17186_BLOOD_VS_CORD_BLOOD_CD21LOW_TRANSITIONAL_BCELL_UP | GSE17186_BLOOD_VS_CORD_BLOOD_CD21LOW_TRANSITIONAL_BCELL_UP | 17/611 | 200/21355 | 7E-05 | 0.00185 | 0.00137 | 85464/6777/9267/4775/23064/25777/51696/9960/4627/120/5770/23248/84937/51735/4012/2590/27244 | 17 |
| GSE17301_ACD3_ACD28_VS_ACD3_ACD28_AND_IFNA2_STIM_CD8_TCELL_DN | GSE17301_ACD3_ACD28_VS_ACD3_ACD28_AND_IFNA2_STIM_CD8_TCELL_DN | 17/611 | 200/21355 | 7E-05 | 0.00185 | 0.00137 | 1387/9267/56913/987/25777/11320/9840/6249/1729/1203/7187/26207/4850/7699/3454/57337/60685 | 17 |
| GSE17721_CTRL_VS_GARDIQUIMOD_0.5H_BMDC_UP | GSE17721_CTRL_VS_GARDIQUIMOD_0.5H_BMDC_UP | 17/611 | 200/21355 | 7E-05 | 0.00185 | 0.00137 | 10672/56261/2869/28977/4670/1119/9567/8826/23077/84830/23406/51271/51455/5770/25853/54617/10906 | 17 |
| GSE18893_TCONV_VS_TREG_2H_CULTURE_DN | GSE18893_TCONV_VS_TREG_2H_CULTURE_DN | 17/611 | 200/21355 | 7E-05 | 0.00185 | 0.00137 | 56261/1073/51552/11320/2081/6780/79956/7920/4026/23102/976/5770/22834/55500/64750/10906/79657 | 17 |
| GSE24026_PD1_LIGATION_VS_CTRL_IN_ACT_TCELL_LINE_DN | GSE24026_PD1_LIGATION_VS_CTRL_IN_ACT_TCELL_LINE_DN | 17/611 | 200/21355 | 7E-05 | 0.00185 | 0.00137 | 23214/4775/58513/64848/7187/9797/6929/65117/54815/54878/9807/55683/22870/55818/55500/10198/60685 | 17 |
| GSE24726_WT_VS_E2_2_KO_PDC_DAY6_POST_DELETION_DN | GSE24726_WT_VS_E2_2_KO_PDC_DAY6_POST_DELETION_DN | 17/611 | 200/21355 | 7E-05 | 0.00185 | 0.00137 | 65125/10672/85464/9698/65059/7456/1265/6938/546/3655/23303/378938/11163/80728/9459/4354/473 | 17 |
| GSE26030_UNSTIM_VS_RESTIM_TH1_DAY15_POST_POLARIZATION_UP | GSE26030_UNSTIM_VS_RESTIM_TH1_DAY15_POST_POLARIZATION_UP | 17/611 | 200/21355 | 7E-05 | 0.00185 | 0.00137 | 7534/90273/6774/3683/401409/29980/3106/860/6337/1060/5305/259230/1739/80728/9020/8428/60685 | 17 |
| GSE2770_IL4_ACT_VS_ACT_CD4_TCELL_2H_DN | GSE2770_IL4_ACT_VS_ACT_CD4_TCELL_2H_DN | 17/611 | 200/21355 | 7E-05 | 0.00185 | 0.00137 | 11052/9135/23214/5820/4245/84166/51068/7187/10425/4026/9962/50650/5770/155038/6891/51735/9020 | 17 |
| GSE2770_TGFB_AND_IL4_VS_IL12_TREATED_ACT_CD4_TCELL_6H_UP | GSE2770_TGFB_AND_IL4_VS_IL12_TREATED_ACT_CD4_TCELL_6H_UP | 17/611 | 200/21355 | 7E-05 | 0.00185 | 0.00137 | 7456/55187/8826/25777/11320/23077/55252/23150/22848/1520/9797/5588/4627/23198/11064/1911/84937 | 17 |

| | | | | | | | | |
|---|---|---|---|---|---|---|---|---|
| GSE2770_UNTREATED_VS_IL4_TREATED_ACT_CD4_TCELL_6H_DN | GSE2770_UNTREATED_VS_IL4_TREATED_ACT_CD4_TCELL_6H_DN | 17/611 | 200/21355 | 7E-05 | 0.00185 | 0.00137 | 10672/9135/7456/4775/79718/4215/54899/5880/120/2885/29123/10992/6197/22870/55023/163486/80025 | 17 |
| GSE27786_LIN_NEG_VS_CD8_TCELL_DN | GSE27786_LIN_NEG_VS_CD8_TCELL_DN | 17/611 | 200/21355 | 7E-05 | 0.00185 | 0.00137 | 7799/23167/10390/55187/3683/81669/84166/89846/1455/5339/10048/378938/9807/23248/10992/6197/114799 | 17 |
| GSE31082_DN_VS_CD4_SP_THYMOCYTE_DN | GSE31082_DN_VS_CD4_SP_THYMOCYTE_DN | 17/611 | 200/21355 | 7E-05 | 0.00185 | 0.00137 | 60468/1387/65059/23214/9267/150864/944/9728/7187/54838/55972/3454/283131/9020/8428/9459/473 | 17 |
| GSE33425_CD161_HIGH_VS_INT_CD8_TCELL_UP | GSE33425_CD161_HIGH_VS_INT_CD8_TCELL_UP | 17/611 | 200/21355 | 7E-05 | 0.00185 | 0.00137 | 51552/90273/7414/1105/8826/7326/4245/1520/221477/55870/3655/255231/5925/5977/5090/23607/51720 | 17 |
| GSE339_CD4POS_VS_CD4CD8DN_DC_UP | GSE339_CD4POS_VS_CD4CD8DN_DC_UP | 17/611 | 200/21355 | 7E-05 | 0.00185 | 0.00137 | 11329/10111/2909/10615/11165/22955/10163/1432/3607/3077/57649/5305/10956/1739/5094/10906/27244 | 17 |
| GSE369_PRE_VS_POST_IL6_INJECTION_IFNG_WT_LIVER_DN | GSE369_PRE_VS_POST_IL6_INJECTION_IFNG_WT_LIVER_DN | 17/611 | 200/21355 | 7E-05 | 0.00185 | 0.00137 | 85464/4763/9873/23214/150864/823/23369/23347/3655/1606/120/50650/324/5900/30844/25938/5906 | 17 |
| GSE411_UNSTIM_VS_100MIN_IL6_STIM_SOCS3_KO_MACROPHAGE_DN | GSE411_UNSTIM_VS_100MIN_IL6_STIM_SOCS3_KO_MACROPHAGE_DN | 17/611 | 200/21355 | 7E-05 | 0.00185 | 0.00137 | 10672/56261/5496/150864/23071/59269/7326/10006/3157/50807/9797/3655/3560/5165/83891/4134/7403 | 17 |
| GSE41867_DAY8_VS_DAY15_LCMV_CLONE13_EFFECTOR_CD8_TCELL_UP | GSE41867_DAY8_VS_DAY15_LCMV_CLONE13_EFFECTOR_CD8_TCELL_UP | 17/611 | 200/21355 | 7E-05 | 0.00185 | 0.00137 | 1387/7799/9698/10390/4820/57585/337867/8073/23293/51586/23384/9712/259230/56852/1911/9402/51317 | 17 |
| GSE45739_UNSTIM_VS_ACD3_ACD28_STIM_WT_CD4_TCELL_UP | GSE45739_UNSTIM_VS_ACD3_ACD28_STIM_WT_CD4_TCELL_UP | 17/611 | 200/21355 | 7E-05 | 0.00185 | 0.00137 | 1540/5820/9840/2272/84830/1203/3655/10144/4215/1606/120/3077/9466/50852/51176/10800/473 | 17 |
| GSE5503_PLN_DC_VS_SPLEEN_DC_ACTIVATED_ALLOGENIC_TCELL_DN | GSE5503_PLN_DC_VS_SPLEEN_DC_ACTIVATED_ALLOGENIC_TCELL_DN | 17/611 | 200/21355 | 7E-05 | 0.00185 | 0.00137 | 23167/23064/54834/823/10006/53944/9491/7187/22908/79036/549/9466/10905/11163/57198/64750/2643 | 17 |
| GSE5679_CTRL_VS_PPARG_LIGAND_ROSIGLITAZONE_TREATED_DC_DN | GSE5679_CTRL_VS_PPARG_LIGAND_ROSIGLITAZONE_TREATED_DC_DN | 17/611 | 200/21355 | 7E-05 | 0.00185 | 0.00137 | 23112/6777/22828/81669/3106/546/753/5295/10425/976/9807/5578/8672/23365/1399/197135/8428 | 17 |
| GSE5679_PPARG_LIGAND_ROSIGLITAZONE_VS_ROSIGLITAZONE_AND_RARA_AGONIST_AM580_TREATED_DC_UP | GSE5679_PPARG_LIGAND_ROSIGLITAZONE_VS_ROSIGLITAZONE_AND_RARA_AGONIST_AM580_TREATED_DC_UP | 17/611 | 200/21355 | 7E-05 | 0.00185 | 0.00137 | 8844/56005/65059/944/401409/6526/50807/3594/5090/64968/114804/7443/7798/114836/4354/51501/81671 | 17 |
| GSE7831_UNSTIM_VS_CPG_STIM_PDC_4H_UP | GSE7831_UNSTIM_VS_CPG_STIM_PDC_4H_UP | 17/611 | 200/21355 | 7E-05 | 0.00185 | 0.00137 | 7534/3936/22828/1265/51072/5048/25777/23406/10048/26207/1606/5090/64333/11163/51176/51719/2590 | 17 |
| GSE9650_NAIVE_VS_EFF_CD8_TCELL_UP | GSE9650_NAIVE_VS_EFF_CD8_TCELL_UP | 17/611 | 200/21355 | 7E-05 | 0.00185 | 0.00137 | 54934/9736/57585/23077/3480/4297/51108/1520/3707/6223/1606/84937/23607/4354/473/27244/7267 | 17 |
| GSE7568_IL4_VS_IL4_AND_TGFB_TREATED_MACROPHAGE_24H_UP | GSE7568_IL4_VS_IL4_AND_TGFB_TREATED_MACROPHAGE_24H_UP | 15/611 | 167/21355 | 9E-05 | 0.00262 | 0.00194 | 132789/9873/23304/9267/79956/9918/29761/51176/51735/84186/55023/60685/51317/84636/26043 | 15 |
| THAKAR_PBMC_INACTIVATED_INFLUENZA_AGE_21_30YO_NONRESPONDER_28DY_UP | THAKAR_PBMC_INACTIVATED_INFLUENZA_AGE_21_30YO_NONRESPONDER_28DY_UP | 15/611 | 174/21355 | 0.0002 | 0.00413 | 0.00306 | 23167/10390/150864/6938/23369/23347/10006/79718/1203/83478/3157/26036/64783/79692/147657 | 15 |
| GSE5099_CLASSICAL_M1_VS_ALTERNATIVE_M2_MACROPHAGE_UP | GSE5099_CLASSICAL_M1_VS_ALTERNATIVE_M2_MACROPHAGE_UP | 16/611 | 194/21355 | 0.0002 | 0.00418 | 0.0031 | 10390/26157/64848/4245/3106/23406/6223/120/9466/3594/155038/6197/2035/57198/64750/9844 | 16 |
| GSE17974_CTRL_VS_ACT_IL4_AND_ANTI_IL12_24H_CD4_TCELL_UP | GSE17974_CTRL_VS_ACT_IL4_AND_ANTI_IL12_24H_CD4_TCELL_UP | 16/611 | 195/21355 | 0.0002 | 0.00418 | 0.0031 | 60468/944/256435/22861/11320/155435/7554/2272/9728/5533/84458/3707/3655/57337/60685/84441 | 16 |
| GSE9601_UNTREATED_VS_PI3K_INHIBITOR_TREATED_HCMV_INF_MONOCYTE_UP | GSE9601_UNTREATED_VS_PI3K_INHIBITOR_TREATED_HCMV_INF_MONOCYTE_UP | 15/611 | 176/21355 | 0.0002 | 0.00418 | 0.0031 | 3936/672/1455/1786/5588/4627/51586/29123/30844/10956/10992/128866/120425/1010/84766 | 15 |
| GSE17974_0.5H_VS_72H_IL4_AND_ANTI_IL12_ACT_CD4_TCELL_UP | GSE17974_0.5H_VS_72H_IL4_AND_ANTI_IL12_ACT_CD4_TCELL_UP | 16/611 | 196/21355 | 0.0002 | 0.00418 | 0.0031 | 11329/64324/1105/26065/55095/51696/84458/3655/9960/9693/5295/9712/51735/7109/9263/7403 | 16 |
| GSE13411_PLASMA_CELL_VS_MEMORY_BCELL_DN | GSE13411_PLASMA_CELL_VS_MEMORY_BCELL_DN | 16/611 | 197/21355 | 0.0002 | 0.00418 | 0.0031 | 23370/3936/5257/55619/1455/83478/6894/54838/11165/9611/10163/23048/5305/11064/23355/9459 | 16 |
| GSE21546_ELK1_KO_VS_SAP1A_KO_AND_ELK1_KO_DP_THYMOCYTES_DN | GSE21546_ELK1_KO_VS_SAP1A_KO_AND_ELK1_KO_DP_THYMOCYTES_DN | 16/611 | 197/21355 | 0.0002 | 0.00418 | 0.0031 | 23649/9735/4670/8847/672/155435/641/84316/10036/10615/9918/23102/2521/219988/9129/23450 | 16 |
| GSE36476_YOUNG_VS_OLD_DONOR_MEMORY_CD4_TCELL_40H_TSST_ACT_DN | GSE36476_YOUNG_VS_OLD_DONOR_MEMORY_CD4_TCELL_40H_TSST_ACT_DN | 16/611 | 197/21355 | 0.0002 | 0.00418 | 0.0031 | 2869/11329/23304/2272/84830/51108/7187/10144/23141/10163/5090/5900/5578/9020/4791/79982 | 16 |

| | | | | | | | | |
|---|---|---|---|---|---|---|---|---|
| GSE37416_0H_VS_48H_F_TULARENSIS_LVS_NEUTROPHIL_UP | GSE37416_0H_VS_48H_F_TULARENSIS_LVS_NEUTROPHIL_UP | 16/611 | 197/21355 | 0.0002 | 0.00418 | 0.0031 | 51742/11329/23304/3936/7456/150864/26157/1265/253461/860/157378/10425/7813/10403/64333/51719 | 16 |
| GSE9988_ANTI_TREM1_AND_LPS_VS_CTRL_TREATED_MONOCYTES_DN | GSE9988_ANTI_TREM1_AND_LPS_VS_CTRL_TREATED_MONOCYTES_DN | 16/611 | 197/21355 | 0.0002 | 0.00418 | 0.0031 | 22992/51696/4297/6780/54838/9960/1432/8289/54878/9520/155038/10992/25853/9020/8428/10521 | 16 |
| GSE11057_NAIVE_VS_MEMORY_CD4_TCELL_DN | GSE11057_NAIVE_VS_MEMORY_CD4_TCELL_DN | 16/611 | 198/21355 | 0.0002 | 0.00418 | 0.0031 | 253461/8826/965/2909/257160/4967/23406/255231/4026/3560/135112/30844/10788/9949/57534/493 | 16 |
| GSE11864_UNTREATED_VS_CSF1_PAM3CYS_IN_MAC_UP | GSE11864_UNTREATED_VS_CSF1_PAM3CYS_IN_MAC_UP | 16/611 | 198/21355 | 0.0002 | 0.00418 | 0.0031 | 85464/1073/54934/7414/55095/91775/79663/55852/5775/165918/23406/26036/79745/4354/51317/5906 | 16 |
| GSE13738_TCR_VS_BYSTANDER_ACTIVATED_CD4_TCELL_UP | GSE13738_TCR_VS_BYSTANDER_ACTIVATED_CD4_TCELL_UP | 16/611 | 198/21355 | 0.0002 | 0.00418 | 0.0031 | 11052/55526/4775/146712/7326/6526/6894/22978/64968/23248/23355/9577/9459/4354/55784/9673 | 16 |
| GSE21360_PRIMARY_VS_QUATERNARY_MEMORY_CD8_TCELL_DN | GSE21360_PRIMARY_VS_QUATERNARY_MEMORY_CD8_TCELL_DN | 16/611 | 198/21355 | 0.0002 | 0.00418 | 0.0031 | 9976/10006/23774/9611/23499/8631/10163/5588/5977/120/157680/55291/22834/64750/54521/23287 | 16 |
| GSE22886_NAIVE_TCELL_VS_NKCELL_UP | GSE22886_NAIVE_TCELL_VS_NKCELL_UP | 16/611 | 198/21355 | 0.0002 | 0.00418 | 0.0031 | 6777/171023/23370/1265/944/2926/10892/4363/22955/6767/1606/10767/5578/51176/104/2289 | 16 |
| GSE22886_UNSTIM_VS_STIM_MEMORY_TCELL_UP | GSE22886_UNSTIM_VS_STIM_MEMORY_TCELL_UP | 16/611 | 198/21355 | 0.0002 | 0.00418 | 0.0031 | 11329/1265/26065/25777/5775/3655/26207/5295/8631/3077/5305/976/8742/23527/55291/27244 | 16 |
| GSE25088_IL4_VS_IL4_AND_ROSIGLITAZONE_STIM_STAT6_KO_MACROPHAGE_DAY10_UP | GSE25088_IL4_VS_IL4_AND_ROSIGLITAZONE_STIM_STAT6_KO_MACROPHAGE_DAY10_UP | 16/611 | 198/21355 | 0.0002 | 0.00418 | 0.0031 | 85464/23112/55526/55690/4775/860/157378/23406/120/54878/64333/64750/493/51317/23287/473 | 16 |
| GSE2770_IL12_AND_TGFB_ACT_VS_ACT_CD4_TCELL_6H_DN | GSE2770_IL12_AND_TGFB_ACT_VS_ACT_CD4_TCELL_6H_DN | 16/611 | 198/21355 | 0.0002 | 0.00418 | 0.0031 | 7414/944/23150/6780/80196/138151/2521/5770/9712/8726/10906/2643/493/9673/31/10963 | 16 |
| GSE2770_IL4_ACT_VS_ACT_CD4_TCELL_6H_UP | GSE2770_IL4_ACT_VS_ACT_CD4_TCELL_6H_UP | 16/611 | 198/21355 | 0.0002 | 0.00418 | 0.0031 | 54934/7414/26157/2081/256236/84166/84316/51068/1431/6929/120/3594/7644/11184/124565/473 | 16 |
| GSE32986_GMCSF_VS_GMCSF_AND_CURDLAN_HIGHDOSE_STIM_DC_DN | GSE32986_GMCSF_VS_GMCSF_AND_CURDLAN_HIGHDOSE_STIM_DC_DN | 16/611 | 198/21355 | 0.0002 | 0.00418 | 0.0031 | 4775/1119/23633/55252/55619/6780/55852/3707/25981/51586/7813/51460/23019/64750/10198/163486 | 16 |
| GSE36476_CTRL_VS_TSST_ACT_16H_MEMORY_CD4_TCELL_OLD_UP | GSE36476_CTRL_VS_TSST_ACT_16H_MEMORY_CD4_TCELL_OLD_UP | 16/611 | 198/21355 | 0.0002 | 0.00418 | 0.0031 | 4763/54934/23370/1105/25777/11320/23077/51696/26207/9960/5588/120/7769/57711/51176/9459 | 16 |
| GSE36476_CTRL_VS_TSST_ACT_40H_MEMORY_CD4_TCELL_YOUNG_UP | GSE36476_CTRL_VS_TSST_ACT_40H_MEMORY_CD4_TCELL_YOUNG_UP | 16/611 | 198/21355 | 0.0002 | 0.00418 | 0.0031 | 54934/23370/9267/1105/9736/25777/11320/51696/5775/10144/26207/9960/5295/120/51176/7109 | 16 |
| GSE3982_EOSINOPHIL_VS_BASOPHIL_DN | GSE3982_EOSINOPHIL_VS_BASOPHIL_DN | 16/611 | 198/21355 | 0.0002 | 0.00418 | 0.0031 | 4763/11329/5148/9840/84316/10464/9611/8631/5588/120/157680/51/6599/7443/10330/9694 | 16 |
| GSE3982_MAST_CELL_VS_TH1_DN | GSE3982_MAST_CELL_VS_TH1_DN | 16/611 | 198/21355 | 0.0002 | 0.00418 | 0.0031 | 5496/4763/23370/9735/1540/57102/59269/641/5533/10615/5933/29123/5165/6891/2643/2289 | 16 |
| GSE5542_UNTREATED_VS_IFNG_TREATED_EPITHELIAL_CELLS_24H_UP | GSE5542_UNTREATED_VS_IFNG_TREATED_EPITHELIAL_CELLS_24H_UP | 16/611 | 198/21355 | 0.0002 | 0.00418 | 0.0031 | 8844/6777/377/8826/2909/55605/8729/4967/5925/3710/3560/3594/6645/23365/7798/283989 | 16 |
| GSE4984_GALECTIN1_VS_LPS_STIM_DC_UP | GSE4984_GALECTIN1_VS_LPS_STIM_DC_UP | 14/611 | 159/21355 | 0.0002 | 0.00418 | 0.0031 | 51742/9267/585/25777/7182/4627/8289/157680/29761/11064/23163/23607/25853/27244 | 14 |
| GSE11057_CD4_EFF_MEM_VS_PBMC_UP | GSE11057_CD4_EFF_MEM_VS_PBMC_UP | 16/611 | 199/21355 | 0.0002 | 0.00418 | 0.0031 | 51742/6777/284001/51696/79718/10892/3707/3710/51460/114804/50852/2035/9020/57198/9949/51719 | 16 |
| GSE11961_MARGINAL_ZONE_BCELL_VS_GERMINAL_CENTER_BCELL_DAY40_UP | GSE11961_MARGINAL_ZONE_BCELL_VS_GERMINAL_CENTER_BCELL_DAY40_UP | 16/611 | 199/21355 | 0.0002 | 0.00418 | 0.0031 | 51072/2272/84166/8500/22848/7182/9851/23186/5295/57410/5933/23443/9466/114804/10129/57198 | 16 |
| GSE12845_IGD_NEG_BLOOD_VS_DARKZONE_GC_TONSIL_BCELL_DN | GSE12845_IGD_NEG_BLOOD_VS_DARKZONE_GC_TONSIL_BCELL_DN | 16/611 | 199/21355 | 0.0002 | 0.00418 | 0.0031 | 171023/5496/4670/3936/1119/4820/7326/3707/50807/26207/65117/259230/10096/4012/11184/51501 | 16 |
| GSE12845_IGD_POS_BLOOD_VS_NAIVE_TONSIL_BCELL_DN | GSE12845_IGD_POS_BLOOD_VS_NAIVE_TONSIL_BCELL_DN | 16/611 | 199/21355 | 0.0002 | 0.00418 | 0.0031 | 2869/23112/54934/9698/1105/22992/2885/65117/4026/9962/976/8672/23013/4012/163486/5514 | 16 |
| GSE13306_TREG_VS_TCONV_SPLEEN_DN | GSE13306_TREG_VS_TCONV_SPLEEN_DN | 16/611 | 199/21355 | 0.0002 | 0.00418 | 0.0031 | 23370/7456/701/22848/1786/255231/283209/259230/5094/80728/4012/11184/10800/54842/2289/9953 | 16 |

| | | | | | | | | |
|---|---|---|---|---|---|---|---|---|
| GSE15733_BM_VS_SPLEEN_MEMORY_CD4_TCELL_DN | GSE15733_BM_VS_SPLEEN_MEMORY_CD4_TCELL_DN | 16/611 | 199/21355 | 0.0002 | 0.00418 | 0.0031 | 7534/54934/23370/26574/9267/91775/7322/6249/10163/5305/9466/64968/84181/5578/114836/31 | 16 |
| GSE21546_UNSTIM_VS_ANTI_CD3_STIM_DP_THYMOCYTES_DN | GSE21546_UNSTIM_VS_ANTI_CD3_STIM_DP_THYMOCYTES_DN | 16/611 | 199/21355 | 0.0002 | 0.00418 | 0.0031 | 9735/4670/672/641/10036/6894/10615/1786/4215/23141/23102/2521/5578/4931/6645/9129 | 16 |
| GSE22432_CONVENTIONAL_CDC_VS_PLASMACYTOID_PDC_UP | GSE22432_CONVENTIONAL_CDC_VS_PLASMACYTOID_PDC_UP | 16/611 | 199/21355 | 0.0002 | 0.00418 | 0.0031 | 9735/57102/23511/3480/91942/3157/23406/6929/1289/3607/23102/2521/7443/92181/283989/5906 | 16 |
| GSE24671_CTRL_VS_SENDAI_VIRUS_INFECTED_MOUSE_SPLENOCYTES_DN | GSE24671_CTRL_VS_SENDAI_VIRUS_INFECTED_MOUSE_SPLENOCYTES_DN | 16/611 | 199/21355 | 0.0002 | 0.00418 | 0.0031 | 55690/4775/55187/3683/25777/57690/84196/6249/124245/7187/26207/9693/9611/10788/8672/2035 | 16 |
| GSE28726_ACT_CD4_TCELL_VS_ACT_NKTCELL_DN | GSE28726_ACT_CD4_TCELL_VS_ACT_NKTCELL_DN | 16/611 | 199/21355 | 0.0002 | 0.00418 | 0.0031 | 9873/23370/9267/1540/26065/22992/3707/8705/3655/5588/1606/120/5515/23355/55818/9263 | 16 |
| GSE39820_TGFBETA3_IL6_VS_TGFBETA3_IL6_IL23A_TREATED_CD4_TCELL_UP | GSE39820_TGFBETA3_IL6_VS_TGFBETA3_IL6_IL23A_TREATED_CD4_TCELL_UP | 16/611 | 199/21355 | 0.0002 | 0.00418 | 0.0031 | 9873/22828/26065/3064/8826/89970/2081/29028/3480/51807/157378/55972/26207/56852/79157/5906 | 16 |
| GSE40068_BCL6_POS_VS_NEG_CXCR5_POS_TFH_DN | GSE40068_BCL6_POS_VS_NEG_CXCR5_POS_TFH_DN | 16/611 | 199/21355 | 0.0002 | 0.00418 | 0.0031 | 1073/9202/6938/2909/6249/1520/753/9693/9466/84937/114799/51735/1822/23524/473/7267 | 16 |
| GSE40273_EOS_KO_VS_WT_TREG_DN | GSE40273_EOS_KO_VS_WT_TREG_DN | 16/611 | 199/21355 | 0.0002 | 0.00418 | 0.0031 | 60468/585/1265/57690/55972/26207/22806/10425/54165/114804/29761/23163/55784/84433/2289/80025 | 16 |
| GSE42724_MEMORY_BCELL_VS_PLASMABLAST_UP | GSE42724_MEMORY_BCELL_VS_PLASMABLAST_UP | 16/611 | 199/21355 | 0.0002 | 0.00418 | 0.0031 | 2869/23214/9267/9889/9567/4297/7421/257160/64968/6891/2035/22870/219988/55500/60685/2643 | 16 |
| GSE21033_1H_VS_12H_POLYIC_STIM_DC_DN | GSE21033_1H_VS_12H_POLYIC_STIM_DC_DN | 13/611 | 141/21355 | 0.0002 | 0.00418 | 0.0031 | 132789/65125/23112/1540/4820/3183/120/3077/23443/29761/56852/50852/10657 | 13 |
| GSE25677_MPL_VS_MPL_AND_R848_STIM_BCELL_UP | GSE25677_MPL_VS_MPL_AND_R848_STIM_BCELL_UP | 14/611 | 160/21355 | 0.0002 | 0.00418 | 0.0031 | 10672/9873/23347/1520/157378/23186/51586/55236/324/51460/10403/84636/80025/473 | 14 |
| GSE13484_UNSTIM_VS_YF17D_VACCINE_STIM_PBMC_UP | GSE13484_UNSTIM_VS_YF17D_VACCINE_STIM_PBMC_UP | 16/611 | 200/21355 | 0.0002 | 0.00418 | 0.0031 | 11052/54934/3936/23369/51696/8073/54838/4215/6767/138151/79801/5090/54870/55818/60685/7267 | 16 |
| GSE13738_RESTING_VS_TCR_ACTIVATED_CD4_TCELL_DN | GSE13738_RESTING_VS_TCR_ACTIVATED_CD4_TCELL_DN | 16/611 | 200/21355 | 0.0002 | 0.00418 | 0.0031 | 28977/586/146712/23071/7326/3069/753/5880/7813/4026/817/23355/84937/55784/84324/9377 | 16 |
| GSE14350_TREG_VS_TEFF_IN_IL2RB_KO_UP | GSE14350_TREG_VS_TEFF_IN_IL2RB_KO_UP | 16/611 | 200/21355 | 0.0002 | 0.00418 | 0.0031 | 9202/23214/26574/55187/81669/23369/55252/84196/1606/120/51586/9466/324/117584/7109/54521 | 16 |
| GSE15330_WT_VS_IKAROS_KO_MEGAKARYOCYTE_ERYTHROID_PROGENITOR_DN | GSE15330_WT_VS_IKAROS_KO_MEGAKARYOCYTE_ERYTHROID_PROGENITOR_DN | 16/611 | 200/21355 | 0.0002 | 0.00418 | 0.0031 | 65125/6777/171023/54834/256435/57585/23186/6223/5880/378938/51/10905/54870/4700/2035/55023 | 16 |
| GSE16385_IFNG_TNF_VS_UNSTIM_MACROPHAGE_ROSIGLITAZONE_TREATED_DN | GSE16385_IFNG_TNF_VS_UNSTIM_MACROPHAGE_ROSIGLITAZONE_TREATED_DN | 16/611 | 200/21355 | 0.0002 | 0.00418 | 0.0031 | 51742/1387/987/55253/3707/5775/120/3560/23102/9466/157680/5205/54521/51719/9129/79982 | 16 |
| GSE16450_IMMATURE_VS_MATURE_NEURON_CELL_LINE_12H_IFNA_STIM_DN | GSE16450_IMMATURE_VS_MATURE_NEURON_CELL_LINE_12H_IFNA_STIM_DN | 16/611 | 200/21355 | 0.0002 | 0.00418 | 0.0031 | 56261/27340/22828/1119/944/58513/57585/8729/4795/23248/54870/375748/54617/79692/60685/54842 | 16 |
| GSE16450_IMMATURE_VS_MATURE_NEURON_CELL_LINE_12H_IFNA_STIM_UP | GSE16450_IMMATURE_VS_MATURE_NEURON_CELL_LINE_12H_IFNA_STIM_UP | 16/611 | 200/21355 | 0.0002 | 0.00418 | 0.0031 | 51742/1387/23370/9267/81669/337867/50807/55870/4215/9693/976/64333/23248/10956/51735/114836 | 16 |
| GSE17186_NAIVE_VS_CD21LOW_TRANSITIONAL_BCELL_DN | GSE17186_NAIVE_VS_CD21LOW_TRANSITIONAL_BCELL_DN | 16/611 | 200/21355 | 0.0002 | 0.00418 | 0.0031 | 6777/27340/9267/4775/23064/6938/1729/55972/6223/5880/3560/283209/9807/728621/284058/27244 | 16 |
| GSE17721_0.5H_VS_4H_POLYIC_BMDC_DN | GSE17721_0.5H_VS_4H_POLYIC_BMDC_DN | 16/611 | 200/21355 | 0.0002 | 0.00418 | 0.0031 | 51742/55690/23167/6780/80196/51271/3660/54815/5165/5900/10905/29761/4642/2643/9129/6654 | 16 |
| GSE17721_PAM3CSK4_VS_CPG_0.5H_BMDC_UP | GSE17721_PAM3CSK4_VS_CPG_0.5H_BMDC_UP | 16/611 | 200/21355 | 0.0002 | 0.00418 | 0.0031 | 377/90273/4204/987/57585/2909/7187/9611/10163/5977/3183/57494/976/7251/128866/115 | 16 |
| GSE20198_IL12_VS_IL12_IL18_TREATED_ACT_CD4_TCELL_DN | GSE20198_IL12_VS_IL12_IL18_TREATED_ACT_CD4_TCELL_DN | 16/611 | 200/21355 | 0.0002 | 0.00418 | 0.0031 | 5496/23131/1105/26065/51072/10006/23150/5339/51455/5977/1606/10425/54165/8726/7109/31 | 16 |
| GSE22601_DOUBLE_NEGATIVE_VS_DOUBLE_POSITIVE_THYMOCYTE_DN | GSE22601_DOUBLE_NEGATIVE_VS_DOUBLE_POSITIVE_THYMOCYTE_DN | 16/611 | 200/21355 | 0.0002 | 0.00418 | 0.0031 | 132789/7534/6774/54834/155435/2081/55252/337867/51068/3707/1606/155038/26133/57337/55784/51720 | 16 |
| GSE22886_NAIVE_CD4_TCELL_VS_48H_ACT_TH1_UP | GSE22886_NAIVE_CD4_TCELL_VS_48H_ACT_TH1_UP | 16/611 | 200/21355 | 0.0002 | 0.00418 | 0.0031 | 6777/9736/23064/4820/81669/22861/22887/9840/7644/23527/5578/23060/1739/51176/22834/23032 | 16 |

| | | | | | | | | |
|---|---|---|---|---|---|---|---|---|
| GSE22919_RESTING_VS_IL2_IL12_IL15_STIM_NK_CELL_DN | GSE22919_RESTING_VS_IL2_IL12_IL15_STIM_NK_CELL_DN | 16/611 | 200/21355 | 0.0002 | 0.00418 | 0.0031 | 9267/1540/585/823/5257/84316/79828/6929/4215/3660/157680/64333/11064/22834/51317/163486 | 16 |
| GSE23984_CTRL_VS_HYPOCALEMIC_VITAMIND_ANALOG_TCELL_DN | GSE23984_CTRL_VS_HYPOCALEMIC_VITAMIND_ANALOG_TCELL_DN | 16/611 | 200/21355 | 0.0002 | 0.00418 | 0.0031 | 56261/7456/8826/59269/23077/4297/89846/3655/1606/3660/2885/5305/283131/23524/27244/23032 | 16 |
| GSE25088_IL4_VS_IL4_AND_ROSIGLITAZONE_STIM_MACROPHAGE_DAY10_UP | GSE25088_IL4_VS_IL4_AND_ROSIGLITAZONE_STIM_MACROPHAGE_DAY10_UP | 16/611 | 200/21355 | 0.0002 | 0.00418 | 0.0031 | 85464/65059/84301/5048/55252/23150/7322/9728/23059/23141/8904/8289/114804/27296/25853/114836 | 16 |
| GSE26351_UNSTIM_VS_WNT_PATHWAY_STIM_HEMATOPOIETIC_PROGENITORS_DN | GSE26351_UNSTIM_VS_WNT_PATHWAY_STIM_HEMATOPOIETIC_PROGENITORS_DN | 16/611 | 200/21355 | 0.0002 | 0.00418 | 0.0031 | 56005/10390/54834/57585/155435/84306/2909/84961/54838/6767/9962/5515/2961/8726/10198/25938 | 16 |
| GSE27786_BCELL_VS_NKCELL_UP | GSE27786_BCELL_VS_NKCELL_UP | 16/611 | 200/21355 | 0.0002 | 0.00418 | 0.0031 | 64766/9202/56913/9736/1455/342945/1606/3710/57494/2521/83891/84181/92181/25853/51317/84433 | 16 |
| GSE27786_CD4_TCELL_VS_NKCELL_UP | GSE27786_CD4_TCELL_VS_NKCELL_UP | 16/611 | 200/21355 | 0.0002 | 0.00418 | 0.0031 | 1540/9736/6938/3747/25777/57690/4297/9611/3607/3183/6891/2035/57198/55500/11184/10330 | 16 |
| GSE27786_NKCELL_VS_ERYTHROBLAST_UP | GSE27786_NKCELL_VS_ERYTHROBLAST_UP | 16/611 | 200/21355 | 0.0002 | 0.00418 | 0.0031 | 28977/9202/23064/6938/91775/22848/5533/3660/8289/2521/92170/4700/8726/22834/83451/31 | 16 |
| GSE3039_ALPHABETA_CD8_TCELL_VS_B1_BCELL_UP | GSE3039_ALPHABETA_CD8_TCELL_VS_B1_BCELL_UP | 16/611 | 200/21355 | 0.0002 | 0.00418 | 0.0031 | 23112/26574/81669/7326/57690/55619/6249/7182/4215/79036/3077/976/157680/51735/63977/55023 | 16 |
| GSE3039_NKT_CELL_VS_ALPHAALPHA_CD8_TCELL_DN | GSE3039_NKT_CELL_VS_ALPHAALPHA_CD8_TCELL_DN | 16/611 | 200/21355 | 0.0002 | 0.00418 | 0.0031 | 3106/8073/91775/84166/197322/1786/8631/5588/22806/3560/3594/155038/6891/79613/80728/79157 | 16 |
| GSE30962_PRIMARY_VS_SECONDARY_ACUTE_LCMV_INF_CD8_TCELL_UP | GSE30962_PRIMARY_VS_SECONDARY_ACUTE_LCMV_INF_CD8_TCELL_UP | 16/611 | 200/21355 | 0.0002 | 0.00418 | 0.0031 | 51072/672/701/55010/50807/10615/9918/79801/10403/2961/9727/5888/7798/114792/83990/2289 | 16 |
| GSE360_CTRL_VS_B_MALAYI_HIGH_DOSE_MAC_DN | GSE360_CTRL_VS_B_MALAYI_HIGH_DOSE_MAC_DN | 16/611 | 200/21355 | 0.0002 | 0.00418 | 0.0031 | 5496/23214/3683/22887/23774/6249/23476/1203/3707/23186/4967/4850/8904/5933/9520/10906 | 16 |
| GSE37301_LYMPHOID_PRIMED_MPP_VS_COMMON_LYMPHOID_PROGENITOR_DN | GSE37301_LYMPHOID_PRIMED_MPP_VS_COMMON_LYMPHOID_PROGENITOR_DN | 16/611 | 200/21355 | 0.0002 | 0.00418 | 0.0031 | 28977/26065/23511/55095/81669/57585/23077/51108/2521/5165/6599/10788/6197/1399/128866/79657 | 16 |
| GSE38681_WT_VS_LYL1_KO_LYMPHOID_PRIMED_MULTIPOTENT_PROGENITOR_DN | GSE38681_WT_VS_LYL1_KO_LYMPHOID_PRIMED_MULTIPOTENT_PROGENITOR_DN | 16/611 | 200/21355 | 0.0002 | 0.00418 | 0.0031 | 7456/6774/3784/6780/23515/3660/5933/817/29761/11163/55500/55023/10906/2643/51720/81671 | 16 |
| GSE40068_CXCR5POS_BCL6POS_TFH_VS_CXCR5NEG_BCL6NEG_CD4_TCELL_UP | GSE40068_CXCR5POS_BCL6POS_TFH_VS_CXCR5NEG_BCL6NEG_CD4_TCELL_UP | 16/611 | 200/21355 | 0.0002 | 0.00418 | 0.0031 | 51742/23112/1073/60468/2909/22848/84458/6223/120/9466/23019/10788/1822/22834/23524/7267 | 16 |
| GSE40666_UNTREATED_VS_IFNA_STIM_EFFECTOR_CD8_TCELL_90MIN_UP | GSE40666_UNTREATED_VS_IFNA_STIM_EFFECTOR_CD8_TCELL_90MIN_UP | 16/611 | 200/21355 | 0.0002 | 0.00418 | 0.0031 | 586/7046/3683/7326/6526/256236/84166/1431/23406/3077/976/5770/4642/9949/51719/493 | 16 |
| GSE41867_DAY8_EFFECTOR_VS_DAY30_EXHAUSTED_CD8_TCELL_LCMV_CLONE13_DN | GSE41867_DAY8_EFFECTOR_VS_DAY30_EXHAUSTED_CD8_TCELL_LCMV_CLONE13_DN | 16/611 | 200/21355 | 0.0002 | 0.00418 | 0.0031 | 9698/9889/23071/6938/59269/2272/79663/23406/4627/120/8726/54521/11184/31/10657/27244 | 16 |
| GSE43863_TH1_VS_TFH_EFFECTOR_CD4_TCELL_DN | GSE43863_TH1_VS_TFH_EFFECTOR_CD4_TCELL_DN | 16/611 | 200/21355 | 0.0002 | 0.00418 | 0.0031 | 1387/9135/146057/25777/57690/51696/5339/84961/8289/9807/84937/55818/9020/57337/54521/23032 | 16 |
| GSE4535_BM_DERIVED_DC_VS_FOLLICULAR_DC_UP | GSE4535_BM_DERIVED_DC_VS_FOLLICULAR_DC_UP | 16/611 | 200/21355 | 0.0002 | 0.00418 | 0.0031 | 65059/944/401409/6526/10892/50807/3594/5770/5090/83891/64968/114804/7798/114836/4354/81671 | 16 |
| GSE5503_MLN_DC_VS_PLN_DC_ACTIVATED_ALLOGENIC_TCELL_DN | GSE5503_MLN_DC_VS_PLN_DC_ACTIVATED_ALLOGENIC_TCELL_DN | 16/611 | 200/21355 | 0.0002 | 0.00418 | 0.0031 | 65125/171023/60468/1387/7799/9736/23047/79718/3157/23141/55236/5305/114804/5578/56852/23524 | 16 |
| GSE5589_IL6_KO_VS_IL10_KO_LPS_AND_IL6_STIM_MACROPHAGE_45MIN_UP | GSE5589_IL6_KO_VS_IL10_KO_LPS_AND_IL6_STIM_MACROPHAGE_45MIN_UP | 16/611 | 200/21355 | 0.0002 | 0.00418 | 0.0031 | 23181/55690/51696/84376/701/157378/4363/1786/54899/871/54870/11064/57198/9844/10800/124565 | 16 |
| GSE5679_PPARG_LIGAND_ROSIGLITAZONE_VS_RARA_AGONIST_AM580_TREATED_DC_DN | GSE5679_PPARG_LIGAND_ROSIGLITAZONE_VS_RARA_AGONIST_AM580_TREATED_DC_DN | 16/611 | 200/21355 | 0.0002 | 0.00418 | 0.0031 | 56261/23370/91775/55619/89846/3655/26207/5295/1606/4627/8289/259230/27296/22834/114836/9953 | 16 |
| GSE5960_TH1_VS_ANERGIC_TH1_DN | GSE5960_TH1_VS_ANERGIC_TH1_DN | 16/611 | 200/21355 | 0.0002 | 0.00418 | 0.0031 | 23731/4204/51072/823/672/8073/4363/5933/2885/5165/6599/26133/5578/4700/8726/4791 | 16 |
| GSE9037_CTRL_VS_LPS_1H_STIM_BMDM_UP | GSE9037_CTRL_VS_LPS_1H_STIM_BMDM_UP | 16/611 | 200/21355 | 0.0002 | 0.00418 | 0.0031 | 1387/1105/84301/81669/25777/4297/165918/4850/10163/4627/25862/552900/51176/4354/79982/6654 | 16 |

| | | | | | | | | |
|---|---|---|---|---|---|---|---|---|
| GSE40274_IRF4_VS_FOXP3_AND_IRF4_TRANSDUCED_ACTIVATED_CD4_TCELL_UP | GSE40274_IRF4_VS_FOXP3_AND_IRF4_TRANSDUCED_ACTIVATED_CD4_TCELL_UP | 15/611 | 180/21355 | 0.0002 | 0.00418 | 0.0031 | 23370/9567/8826/23150/9491/1606/4627/9807/23198/55683/10906/4791/9219/2289/5514 | 15 |
| GSE4590_LARGE_PRE_BCELL_VS_VPREB_POS_LARGE_PRE_BCELL_DN | GSE4590_LARGE_PRE_BCELL_VS_VPREB_POS_LARGE_PRE_BCELL_DN | 15/611 | 180/21355 | 0.0002 | 0.00418 | 0.0031 | 85464/9202/23150/22848/9693/5588/22806/50650/976/51/51176/51735/114836/7267/23032 | 15 |
| GSE13547_CTRL_VS_ANTI_IGM_STIM_ZFX_KO_BCELL_12H_DN | GSE13547_CTRL_VS_ANTI_IGM_STIM_ZFX_KO_BCELL_12H_DN | 14/611 | 162/21355 | 0.0002 | 0.0046 | 0.00341 | 60468/25777/200424/3480/79828/79663/146691/50852/84937/51176/51317/84441/473/7267 | 14 |
| NAKAYA_MYELOID_DENDRITIC_CELL_FLUMIST_AGE_18_50YO_7DY_UP | NAKAYA_MYELOID_DENDRITIC_CELL_FLUMIST_AGE_18_50YO_7DY_UP | 24/611 | 378/21355 | 0.0003 | 0.00478 | 0.00355 | 7799/64421/26574/9267/23047/22887/79718/10111/23476/753/23141/4850/23499/1606/7813/23443/976/3454/5578/11064/55683/55500/2289/7267 | 24 |
| GSE14415_INDUCED_TREG_VS_TCONV_DN | GSE14415_INDUCED_TREG_VS_TCONV_DN | 14/611 | 164/21355 | 0.0003 | 0.00516 | 0.00383 | 57521/256435/57690/9840/5533/157378/26207/9466/259230/50852/51317/54842/84441/9953 | 14 |
| GSE7768_OVA_ALONE_VS_OVA_WITH_MPL_IMMUNIZED_MOUSE_WHOLE_SPLEEN_6H_DN | GSE7768_OVA_ALONE_VS_OVA_WITH_MPL_IMMUNIZED_MOUSE_WHOLE_SPLEEN_6H_DN | 14/611 | 164/21355 | 0.0003 | 0.00516 | 0.00383 | 9887/64421/6774/84166/23515/3660/23198/9520/30844/29761/23607/54617/124446/2643 | 14 |
| THAKAR_PBMC_INACTIVATED_INFLUENZA_AGE_21_30YO_NONRESPONDER_7DY_UP | THAKAR_PBMC_INACTIVATED_INFLUENZA_AGE_21_30YO_NONRESPONDER_7DY_UP | 12/611 | 127/21355 | 0.0003 | 0.00548 | 0.00407 | 23167/10390/150864/9782/23369/1203/83478/3157/26036/55291/79692/147657 | 12 |
| GSE7509_UNSTIM_VS_FCGRIIB_STIM_MONOCYTE_DN | GSE7509_UNSTIM_VS_FCGRIIB_STIM_MONOCYTE_DN | 14/611 | 166/21355 | 0.0003 | 0.0058 | 0.00431 | 51742/65125/7799/58513/81669/57690/55252/9611/23499/9918/5090/11064/10129/55818 | 14 |
| GSE7568_CTRL_VS_3H_TGFB_TREATED_MACROPHAGES_WITH_IL4_AND_DEXAMETHASONE_DN | GSE7568_CTRL_VS_3H_TGFB_TREATED_MACROPHAGES_WITH_IL4_AND_DEXAMETHASONE_DN | 14/611 | 167/21355 | 0.0003 | 0.00615 | 0.00456 | 6774/64848/155435/6780/79828/4215/23141/23499/7455/23198/23248/114836/5073/80025 | 14 |
| HOFT_CD4_POSITIVE_ALPHA_BETA_MEMORY_T_CELL_BCG_VACCINE_AGE_18_45YO_56D_TOP_100_DEG_AFTER_IN_VITRO_RE_STIMULATION_DN | HOFT_CD4_POSITIVE_ALPHA_BETA_MEMORY_T_CELL_BCG_VACCINE_AGE_18_45YO_56D_TOP_100_DEG_AFTER_IN_VITRO_RE_STIMULATION_DN | 7/611 | 47/21355 | 0.0004 | 0.00656 | 0.00487 | 23085/4775/55095/8729/4363/10163/23248 | 7 |
| ZAK_PBMC_MRKAD5_HIV_1_GAG_POL_NEF_AGE_20_50YO_1DY_ADDNL_EXON_LVL_UP | ZAK_PBMC_MRKAD5_HIV_1_GAG_POL_NEF_AGE_20_50YO_1DY_ADDNL_EXON_LVL_UP | 10/611 | 94/21355 | 0.0004 | 0.00661 | 0.0049 | 56913/23064/84166/50650/135112/26133/2175/163486/80025/473 | 10 |
| GSE21927_BALBC_VS_C57BL6_MONOCYTE_SPLEEN_UP | GSE21927_BALBC_VS_C57BL6_MONOCYTE_SPLEEN_UP | 15/611 | 189/21355 | 0.0004 | 0.00679 | 0.00504 | 60468/27031/9698/8847/84376/22848/1289/6197/57711/51176/1871/84186/80728/10521/104 | 15 |
| GSE45365_HEALTHY_VS_MCMV_INFECTION_CD11B_DC_UP | GSE45365_HEALTHY_VS_MCMV_INFECTION_CD11B_DC_UP | 15/611 | 189/21355 | 0.0004 | 0.00679 | 0.00504 | 23181/80264/55690/23131/22828/9267/25777/9967/57634/7644/6599/9020/5205/57337/27244 | 15 |
| GSE7219_WT_VS_NIK_NFKB2_KO_DC_DN | GSE7219_WT_VS_NIK_NFKB2_KO_DC_DN | 14/611 | 169/21355 | 0.0004 | 0.00681 | 0.00506 | 60468/4775/944/23064/23499/5588/1606/120/259230/10788/114836/9844/51317/23032 | 14 |
| GSE13485_CTRL_VS_DAY21_YF17D_VACCINE_PBMC_UP | GSE13485_CTRL_VS_DAY21_YF17D_VACCINE_PBMC_UP | 15/611 | 190/21355 | 0.0004 | 0.0071 | 0.00527 | 377/1073/9873/64421/9889/9567/823/155435/7554/165918/9960/3607/2885/4026/51317 | 15 |
| GSE19888_ADENOSINE_A3R_ACT_VS_TCELL_MEMBRANES_ACT_IN_MAST_CELL_DN | GSE19888_ADENOSINE_A3R_ACT_VS_TCELL_MEMBRANES_ACT_IN_MAST_CELL_DN | 15/611 | 190/21355 | 0.0004 | 0.0071 | 0.00527 | 51552/27340/22828/4820/54834/9976/5933/378938/2521/9520/10992/117584/219988/4791/6641 | 15 |
| GSE7768_OVA_ALONE_VS_OVA_WITH_LPS_IMMUNIZED_MOUSE_WHOLE_SPLEEN_6H_DN | GSE7768_OVA_ALONE_VS_OVA_WITH_LPS_IMMUNIZED_MOUSE_WHOLE_SPLEEN_6H_DN | 14/611 | 170/21355 | 0.0004 | 0.00716 | 0.00531 | 10672/59269/84376/55619/1203/10892/255231/4795/9693/23102/10767/5900/117584/2643 | 14 |
| GSE14415_FOXP3_KO_NATURAL_TREG_VS_TCONV_UP | GSE14415_FOXP3_KO_NATURAL_TREG_VS_TCONV_UP | 14/611 | 171/21355 | 0.0004 | 0.00757 | 0.00562 | 85464/10111/79828/79813/79663/8289/50852/84937/51176/51735/23607/51317/27244/7267 | 14 |
| GSE13738_RESTING_VS_BYSTANDER_ACTIVATED_CD4_TCELL_UP | GSE13738_RESTING_VS_BYSTANDER_ACTIVATED_CD4_TCELL_UP | 15/611 | 194/21355 | 0.0005 | 0.00851 | 0.00631 | 23112/150864/55095/22887/11320/3480/7182/84458/6894/79745/23248/163050/9263/388685/84441 | 15 |
| GSE29618_BCELL_VS_MDC_UP | GSE29618_BCELL_VS_MDC_UP | 15/611 | 194/21355 | 0.0005 | 0.00851 | 0.00631 | 2869/60468/9873/80264/7799/23347/51696/4297/5533/83478/22806/120/9712/9263/55784 | 15 |
| GSE3400_UNTREATED_VS_IFNB_TREATED_MEF_UP | GSE3400_UNTREATED_VS_IFNB_TREATED_MEF_UP | 15/611 | 194/21355 | 0.0005 | 0.00851 | 0.00631 | 23731/51696/55619/2909/1060/10479/51/5090/7251/23013/25853/9949/10521/51317/5906 | 15 |
| GSE4748_LPS_VS_LPS_AND_CYANOBACTERIUM_LPSLIKE_STIM_DC_3H_UP | GSE4748_LPS_VS_LPS_AND_CYANOBACTERIUM_LPSLIKE_STIM_DC_3H_UP | 15/611 | 194/21355 | 0.0005 | 0.00851 | 0.00631 | 23112/586/113178/23071/54834/55605/10144/283209/6645/64750/2643/55275/493/79982/26043 | 15 |
| GSE32901_NAIVE_VS_TH17_ENRICHED_CD4_TCELL_UP | GSE32901_NAIVE_VS_TH17_ENRICHED_CD4_TCELL_UP | 13/611 | 154/21355 | 0.0005 | 0.00851 | 0.00631 | 11329/91775/89846/4850/23499/5588/120/378938/976/10788/51176/9402/6641 | 13 |
| GSE16755_CTRL_VS_IFNA_TREATED_MAC_UP | GSE16755_CTRL_VS_IFNA_TREATED_MAC_UP | 15/611 | 195/21355 | 0.0005 | 0.00851 | 0.00631 | 23214/4775/6938/79718/89846/1786/10144/138151/64333/23355/2035/114799/55818/10521/51317 | 15 |

| | | | | | | | | |
|---|---|---|---|---|---|---|---|---|
| GSE46606_IRF4_KO_VS_WT_CD40L_IL2_IL5_3DAY_STIMULATED_BCELL_DN | GSE46606_IRF4_KO_VS_WT_CD40L_IL2_IL5_3DAY_STIMULATED_BCELL_DN | 15/611 | 195/21355 | 0.0005 | 0.00851 | 0.00631 | 1387/944/23064/23071/23347/4733/79663/83478/5976/79036/10425/11064/51719/79692/4354 | 15 |
| GSE14699_NAIVE_VS_ACT_CD8_TCELL_DN | GSE14699_NAIVE_VS_ACT_CD8_TCELL_DN | 14/611 | 175/21355 | 0.0005 | 0.00851 | 0.00631 | 85464/1540/944/57690/2909/79828/50852/84937/51176/51735/9402/23607/27244/7267 | 14 |
| GSE8685_IL2_ACT_IL2_STARVED_VS_IL21_ACT_IL2_STARVED_CD4_TCELL_UP | GSE8685_IL2_ACT_IL2_STARVED_VS_IL21_ACT_IL2_STARVED_CD4_TCELL_UP | 14/611 | 175/21355 | 0.0005 | 0.00851 | 0.00631 | 171023/9735/79954/29980/701/157378/79956/5515/5900/23198/5578/10330/79657/27244 | 14 |
| GSE19401_PLN_VS_PEYERS_PATCH_FOLLICULAR_DC_UP | GSE19401_PLN_VS_PEYERS_PATCH_FOLLICULAR_DC_UP | 15/611 | 196/21355 | 0.0005 | 0.00851 | 0.00631 | 6777/171023/11329/9736/546/2272/200424/6929/138151/6599/7170/1399/23524/473/7267 | 15 |
| GSE46606_UNSTIM_VS_CD40L_IL2_IL5_DAY1_STIMULATED_BCELL_UP | GSE46606_UNSTIM_VS_CD40L_IL2_IL5_DAY1_STIMULATED_BCELL_UP | 15/611 | 196/21355 | 0.0005 | 0.00851 | 0.00631 | 22828/150864/9976/79813/79956/9960/3660/120/23102/7769/10129/5094/55758/55275/84441 | 15 |
| NAKAYA_B_CELL_FLUMIST_AGE_18_50YO_7DY_DN | NAKAYA_B_CELL_FLUMIST_AGE_18_50YO_7DY_DN | 24/611 | 399/21355 | 0.0005 | 0.00851 | 0.00631 | 5343/1540/9567/23633/29028/79718/965/4297/9728/51068/23293/5339/26207/5976/4850/5295/9466/5690/51/7251/1911/9577/493/473 | 24 |
| GSE21033_1H_VS_12H_POLYIC_STIM_DC_UP | GSE21033_1H_VS_12H_POLYIC_STIM_DC_UP | 14/611 | 176/21355 | 0.0006 | 0.00851 | 0.00631 | 64421/56913/4775/4245/3480/3707/3655/23102/5900/114804/23013/4354/4791/10657 | 14 |
| GSE3720_UNSTIM_VS_PMA_STIM_VD1_GAMMADELTA_TCELL_UP | GSE3720_UNSTIM_VS_PMA_STIM_VD1_GAMMADELTA_TCELL_UP | 14/611 | 176/21355 | 0.0006 | 0.00851 | 0.00631 | 55690/23214/944/79663/5533/6223/5305/7170/64333/552900/128866/84766/114836/9377 | 14 |
| NAKAYA_MYELOID_DENDRITIC_CELL_FLUARIX_FLUVIRIN_AGE_18_50YO_7DY_UP | NAKAYA_MYELOID_DENDRITIC_CELL_FLUARIX_FLUVIRIN_AGE_18_50YO_7DY_UP | 24/611 | 400/21355 | 0.0006 | 0.00851 | 0.00631 | 545/11329/9202/55690/4204/64848/23071/23369/55252/79230/9728/79663/23139/6894/4363/6929/10048/1432/3660/2521/23198/9020/55023/23524 | 24 |
| GSE12845_IGD_NEG_BLOOD_VS_NAIVE_TONSIL_BCELL_DN | GSE12845_IGD_NEG_BLOOD_VS_NAIVE_TONSIL_BCELL_DN | 15/611 | 197/21355 | 0.0006 | 0.00851 | 0.00631 | 9135/54934/27340/1119/4820/22861/22992/50807/23499/8631/3710/8289/29123/7040/473 | 15 |
| GSE22033_UNTREATED_VS_MRL24_TREATED_MEF_DN | GSE22033_UNTREATED_VS_MRL24_TREATED_MEF_DN | 15/611 | 197/21355 | 0.0006 | 0.00851 | 0.00631 | 6774/8847/200424/53944/9851/7699/4627/11064/51735/1822/55500/64750/9949/56942/5514 | 15 |
| GSE3982_MAC_VS_BASOPHIL_DN | GSE3982_MAC_VS_BASOPHIL_DN | 15/611 | 197/21355 | 0.0006 | 0.00851 | 0.00631 | 9887/5148/7414/585/26065/7322/84316/55421/1432/11064/10129/10992/493/23287/7403 | 15 |
| GSE40493_BCL6_KO_VS_WT_TREG_UP | GSE40493_BCL6_KO_VS_WT_TREG_UP | 14/611 | 177/21355 | 0.0006 | 0.00851 | 0.00631 | 132789/55187/55095/89970/8073/10111/89846/23186/10055/1778/23048/55236/57534/79657 | 14 |
| GSE12366_GC_VS_MEMORY_BCELL_UP | GSE12366_GC_VS_MEMORY_BCELL_UP | 15/611 | 198/21355 | 0.0006 | 0.00851 | 0.00631 | 10672/60468/586/29028/8073/7421/55421/10615/1786/10055/817/30844/5888/7443/1871 | 15 |
| GSE13738_RESTING_VS_BYSTANDER_ACTIVATED_CD4_TCELL_DN | GSE13738_RESTING_VS_BYSTANDER_ACTIVATED_CD4_TCELL_DN | 15/611 | 198/21355 | 0.0006 | 0.00851 | 0.00631 | 28977/7046/23130/23071/23150/10048/753/5880/4026/3560/976/8742/4700/8428/9377 | 15 |
| GSE20500_RETINOIC_ACID_VS_RARA_ANTAGONIST_TREATED_CD4_TCELL_DN | GSE20500_RETINOIC_ACID_VS_RARA_ANTAGONIST_TREATED_CD4_TCELL_DN | 15/611 | 198/21355 | 0.0006 | 0.00851 | 0.00631 | 60468/55619/6780/3157/55870/51271/55236/51460/9712/1315/259230/11064/23604/9949/56942 | 15 |
| GSE21927_C26GM_VS_4T1_TUMOR_MONOCYTE_BALBC_UP | GSE21927_C26GM_VS_4T1_TUMOR_MONOCYTE_BALBC_UP | 15/611 | 198/21355 | 0.0006 | 0.00851 | 0.00631 | 23201/9873/11329/4255/253461/11320/84196/4297/23101/23406/255231/5880/163486/6641/6654 | 15 |
| GSE22611_NOD2_VS_CTRL_TRANSDUCED_HEK293T_CELL_DN | GSE22611_NOD2_VS_CTRL_TRANSDUCED_HEK293T_CELL_DN | 15/611 | 198/21355 | 0.0006 | 0.00851 | 0.00631 | 23112/90273/23304/146712/23347/7421/5339/23406/4215/9960/9712/23604/9263/80025/51720 | 15 |
| GSE22886_NAIVE_CD4_TCELL_VS_NKCELL_DN | GSE22886_NAIVE_CD4_TCELL_VS_NKCELL_DN | 15/611 | 198/21355 | 0.0006 | 0.00851 | 0.00631 | 7456/6938/641/10111/9728/80196/23101/54165/7769/92170/23198/1315/23365/55784/51317 | 15 |
| GSE2770_IL12_ACT_VS_ACT_CD4_TCELL_48H_DN | GSE2770_IL12_ACT_VS_ACT_CD4_TCELL_48H_DN | 15/611 | 198/21355 | 0.0006 | 0.00851 | 0.00631 | 9135/9873/23064/5820/23077/23059/1431/6767/5933/2885/3710/3560/6599/6197/83990 | 15 |
| GSE360_L_MAJOR_VS_B_MALAYI_HIGH_DOSE_DC_DN | GSE360_L_MAJOR_VS_B_MALAYI_HIGH_DOSE_DC_DN | 15/611 | 198/21355 | 0.0006 | 0.00851 | 0.00631 | 171023/545/22828/7414/3069/9851/4967/5925/1778/5933/9962/51460/9807/23198/7443 | 15 |
| GSE37301_LYMPHOID_PRIMED_MPP_VS_GRAN_MONO_PROGENITOR_DN | GSE37301_LYMPHOID_PRIMED_MPP_VS_GRAN_MONO_PROGENITOR_DN | 15/611 | 198/21355 | 0.0006 | 0.00851 | 0.00631 | 23112/79602/253461/89845/1455/10036/1786/6613/54878/54870/79058/1911/1871/10198/6654 | 15 |
| GSE37416_0H_VS_3H_F_TULARENSIS_LVS_NEUTROPHIL_UP | GSE37416_0H_VS_3H_F_TULARENSIS_LVS_NEUTROPHIL_UP | 15/611 | 198/21355 | 0.0006 | 0.00851 | 0.00631 | 26157/823/80196/9491/84961/50650/10403/64333/10129/2035/120425/22834/4012/9673/79657 | 15 |

| | | | | | | | | |
|---|---|---|---|---|---|---|---|---|
| GSE37416_0H_VS_6H_F_TULARENSIS_LVS_NEUTROPHIL_UP | GSE37416_0H_VS_6H_F_TULARENSIS_LVS_NEUTROPHIL_UP | 15/611 | 198/21355 | 0.0006 | 0.00851 | 0.00631 | 4763/11329/7414/3683/823/84196/9491/4215/9611/7813/7170/64333/23527/6197/9673 | 15 |
| GSE6259_BCELL_VS_CD8_TCELL_DN | GSE6259_BCELL_VS_CD8_TCELL_DN | 15/611 | 198/21355 | 0.0006 | 0.00851 | 0.00631 | 23112/150864/51696/4928/158358/84458/84961/9693/1432/549/10403/54870/7443/9949/473 | 15 |
| GSE9988_LPS_VS_VEHICLE_TREATED_MONOCYTE_DN | GSE9988_LPS_VS_VEHICLE_TREATED_MONOCYTE_DN | 15/611 | 198/21355 | 0.0006 | 0.00851 | 0.00631 | 64421/51696/8073/6794/54838/23406/4215/138151/6613/9466/5094/25853/8428/54521/10521 | 15 |
| NAKAYA_B_CELL_FLUMIST_AGE_18_50YO_7DY_UP | NAKAYA_B_CELL_FLUMIST_AGE_18_50YO_7DY_UP | 27/611 | 475/21355 | 0.0006 | 0.00851 | 0.00631 | 8844/11052/4763/64421/4204/253461/23071/23347/2081/5925/8631/6767/653519/405/55236/9466/5770/5900/23384/1739/8065/4134/10198/9263/147657/2289/2590 | 27 |
| GSE10240_IL17_VS_IL17_AND_IL22_STIM_PRIMARY_BRONCHIAL_EPITHELIAL_CELLS_UP | GSE10240_IL17_VS_IL17_AND_IL22_STIM_PRIMARY_BRONCHIAL_EPITHELIAL_CELLS_UP | 15/611 | 199/21355 | 0.0006 | 0.00851 | 0.00631 | 23132/7799/55187/54834/257160/7409/5977/2885/54878/157680/6645/7798/65979/64750/54521 | 15 |
| GSE12845_PRE_GC_VS_DARKZONE_GC_TONSIL_BCELL_DN | GSE12845_PRE_GC_VS_DARKZONE_GC_TONSIL_BCELL_DN | 15/611 | 199/21355 | 0.0006 | 0.00851 | 0.00631 | 6777/60468/4763/23071/7326/641/84316/4363/22955/6767/10906/55758/9263/51501/473 | 15 |
| GSE14769_40MIN_VS_360MIN_LPS_BMDM_DN | GSE14769_40MIN_VS_360MIN_LPS_BMDM_DN | 15/611 | 199/21355 | 0.0006 | 0.00851 | 0.00631 | 10672/377/7799/58513/91775/55619/9967/23141/8289/23384/26133/23013/57337/124446/163486 | 15 |
| GSE15767_MED_VS_SCS_MAC_LN_DN | GSE15767_MED_VS_SCS_MAC_LN_DN | 15/611 | 199/21355 | 0.0006 | 0.00851 | 0.00631 | 64766/23132/9267/23064/401409/91775/6249/3710/9466/9807/9402/1822/22834/4134/51720 | 15 |
| GSE17721_LPS_VS_POLYIC_6H_BMDC_DN | GSE17721_LPS_VS_POLYIC_6H_BMDC_DN | 15/611 | 199/21355 | 0.0006 | 0.00851 | 0.00631 | 11329/7456/9567/25777/5339/3655/23406/22955/3710/3077/10788/6645/55818/7109/55023 | 15 |
| GSE17721_POLYIC_VS_CPG_8H_BMDC_UP | GSE17721_POLYIC_VS_CPG_8H_BMDC_UP | 15/611 | 199/21355 | 0.0006 | 0.00851 | 0.00631 | 10390/8826/672/3106/701/257160/23476/1729/23515/54838/5976/1606/64783/11184/10800 | 15 |
| GSE17721_POLYIC_VS_PAM3CSK4_16H_BMDC_DN | GSE17721_POLYIC_VS_PAM3CSK4_16H_BMDC_DN | 15/611 | 199/21355 | 0.0006 | 0.00851 | 0.00631 | 2869/57102/5257/5533/23406/1432/3607/10767/51/1315/56852/23365/84937/283131/23287 | 15 |
| GSE17812_WT_VS_THPOK_KO_MEMORY_CD8_TCELL_UP | GSE17812_WT_VS_THPOK_KO_MEMORY_CD8_TCELL_UP | 15/611 | 199/21355 | 0.0006 | 0.00851 | 0.00631 | 60468/23370/9267/51696/51108/79828/157378/9960/5880/120/10905/552900/117584/51735/2590 | 15 |
| GSE17974_0H_VS_2H_IN_VITRO_ACT_CD4_TCELL_UP | GSE17974_0H_VS_2H_IN_VITRO_ACT_CD4_TCELL_UP | 15/611 | 199/21355 | 0.0006 | 0.00851 | 0.00631 | 23201/1073/155435/2272/89845/5257/5533/6894/3655/10144/817/259283/8672/60685/51317 | 15 |
| GSE20198_UNTREATED_VS_IL12_IL18_TREATED_ACT_CD4_TCELL_UP | GSE20198_UNTREATED_VS_IL12_IL18_TREATED_ACT_CD4_TCELL_UP | 15/611 | 199/21355 | 0.0006 | 0.00851 | 0.00631 | 26065/3747/4733/10497/5533/55972/146691/3607/114804/9712/10992/9274/5094/55683/7798 | 15 |
| GSE23505_UNTREATED_VS_4DAY_IL6_IL1_TGFB_TREATED_CD4_TCELL_UP | GSE23505_UNTREATED_VS_4DAY_IL6_IL1_TGFB_TREATED_CD4_TCELL_UP | 15/611 | 199/21355 | 0.0006 | 0.00851 | 0.00631 | 586/9736/1119/59269/256435/9797/3655/255231/22908/1954/50650/10767/84937/9377/54842 | 15 |
| GSE24142_DN2_VS_DN3_THYMOCYTE_ADULT_DN | GSE24142_DN2_VS_DN3_THYMOCYTE_ADULT_DN | 15/611 | 199/21355 | 0.0006 | 0.00851 | 0.00631 | 54934/586/7046/4820/23186/1431/51455/1606/26036/8672/51176/51735/55818/114836/80331 | 15 |
| GSE2585_CTEC_VS_THYMIC_MACROPHAGE_DN | GSE2585_CTEC_VS_THYMIC_MACROPHAGE_DN | 15/611 | 199/21355 | 0.0006 | 0.00851 | 0.00631 | 9887/9735/9736/3683/79718/55619/5339/9611/324/5094/7798/64783/10198/4012/7267 | 15 |
| GSE27859_MACROPHAGE_VS_CD11C_INT_F480_INT_DC_DN | GSE27859_MACROPHAGE_VS_CD11C_INT_F480_INT_DC_DN | 15/611 | 199/21355 | 0.0006 | 0.00851 | 0.00631 | 65059/944/3069/89845/10036/9797/11165/255231/2885/51460/79801/7170/155038/4700/80025 | 15 |
| GSE360_CTRL_VS_L_MAJOR_DC_UP | GSE360_CTRL_VS_L_MAJOR_DC_UP | 15/611 | 199/21355 | 0.0006 | 0.00851 | 0.00631 | 9887/23370/79602/23347/8500/23059/23303/5880/79036/3077/9520/23607/9844/473/10963 | 15 |
| GSE36009_WT_VS_NLRP10_KO_DC_DN | GSE36009_WT_VS_NLRP10_KO_DC_DN | 15/611 | 199/21355 | 0.0006 | 0.00851 | 0.00631 | 6938/59269/51696/1203/7187/4363/255231/126298/976/3454/259230/117584/7109/4791/9219 | 15 |
| GSE369_SOCS3_KO_VS_WT_LIVER_DN | GSE369_SOCS3_KO_VS_WT_LIVER_DN | 15/611 | 199/21355 | 0.0006 | 0.00851 | 0.00631 | 4204/9267/1540/23064/54834/10006/84376/89846/9728/6223/9918/1739/51176/55784/163486 | 15 |
| GSE37532_WT_VS_PPARG_KO_VISCERAL_ADIPOSE_TISSUE_TREG_UP | GSE37532_WT_VS_PPARG_KO_VISCERAL_ADIPOSE_TISSUE_TREG_UP | 15/611 | 199/21355 | 0.0006 | 0.00851 | 0.00631 | 7534/11052/60468/4670/8826/29980/8073/5257/10615/6929/10055/22978/10403/23248/84324 | 15 |
| GSE39820_CTRL_VS_IL1B_IL6_CD4_TCELL_UP | GSE39820_CTRL_VS_IL1B_IL6_CD4_TCELL_UP | 15/611 | 199/21355 | 0.0006 | 0.00851 | 0.00631 | 51696/641/55619/79828/3157/50807/3655/6929/5295/9962/283209/135112/51176/163486/7403 | 15 |

| | | | | | | | | |
|---|---|---|---|---|---|---|---|---|
| GSE40274_LEF1_VS_FOXP3_AND_LEF1_TRANSDUCED_ACTIVATED_CD4_TCELL_DN | GSE40274_LEF1_VS_FOXP3_AND_LEF1_TRANSDUCED_ACTIVATED_CD4_TCELL_DN | 15/611 | 199/21355 | 0.0006 | 0.00851 | 0.00631 | 11329/23370/11320/138151/120/5305/50852/51176/80728/64750/10198/54842/9673/84441/9953 | 15 |
| GSE40666_UNTREATED_VS_IFNA_STIM_STAT4_KO_EFFECTOR_CD8_TCELL_90MIN_DN | GSE40666_UNTREATED_VS_IFNA_STIM_STAT4_KO_EFFECTOR_CD8_TCELL_90MIN_DN | 15/611 | 199/21355 | 0.0006 | 0.00851 | 0.00631 | 60468/9698/8847/57690/79230/8631/163050/8672/79613/22866/9402/4012/55784/84441/27244 | 15 |
| GSE41176_WT_VS_TAK1_KO_ANTI_IGM_STIM_BCELL_6H_DN | GSE41176_WT_VS_TAK1_KO_ANTI_IGM_STIM_BCELL_6H_DN | 15/611 | 199/21355 | 0.0006 | 0.00851 | 0.00631 | 377/9873/585/23511/7409/5880/2885/9918/4026/23048/552900/10788/23163/10906/2643 | 15 |
| GSE557_WT_VS_CIITA_KO_DC_DN | GSE557_WT_VS_CIITA_KO_DC_DN | 15/611 | 199/21355 | 0.0006 | 0.00851 | 0.00631 | 10672/5496/23071/7326/6526/23369/50807/4967/1778/65117/3560/50650/30844/2035/163702 | 15 |
| GSE8685_IL2_STARVED_VS_IL2_ACT_IL2_STARVED_CD4_TCELL_UP | GSE8685_IL2_STARVED_VS_IL2_ACT_IL2_STARVED_CD4_TCELL_UP | 15/611 | 199/21355 | 0.0006 | 0.00851 | 0.00631 | 51742/23731/23214/155435/22848/5533/753/3660/50650/157680/6891/1739/114799/51735/11184 | 15 |
| GSE9006_TYPE_1_VS_TYPE_2_DIABETES_PBMC_AT_DX_DN | GSE9006_TYPE_1_VS_TYPE_2_DIABETES_PBMC_AT_DX_DN | 15/611 | 199/21355 | 0.0006 | 0.00851 | 0.00631 | 55526/7414/6774/79602/54834/51108/50807/55870/4026/54878/23019/6197/23365/493/163486 | 15 |
| ZAK_PBMC_MRKAD5_HIV_1_GAG_POL_NEF_AGE_20_50YO_CORRELATED_WITH_CD8_T_CELL_RESPONSE_3DY_POSITIVE | ZAK_PBMC_MRKAD5_HIV_1_GAG_POL_NEF_AGE_20_50YO_CORRELATED_WITH_CD8_T_CELL_RESPONSE_3DY_POSITIVE | 9/611 | 84/21355 | 0.0007 | 0.00851 | 0.00631 | 11320/546/84376/84458/55972/5295/114799/55818/64750 | 9 |
| GSE3565_DUSP1_VS_WT_SPLENOCYTES_DN | GSE3565_DUSP1_VS_WT_SPLENOCYTES_DN | 14/611 | 179/21355 | 0.0007 | 0.00851 | 0.00631 | 9873/944/6938/3707/6223/1606/120/5305/64333/51176/51735/23607/114836/4354 | 14 |
| GSE10239_MEMORY_VS_DAY4.5_EFF_CD8_TCELL_DN | GSE10239_MEMORY_VS_DAY4.5_EFF_CD8_TCELL_DN | 15/611 | 200/21355 | 0.0007 | 0.00851 | 0.00631 | 10672/171023/51552/56913/987/51072/23150/701/2926/3157/9693/6599/10096/1871/8726 | 15 |
| GSE10240_CTRL_VS_IL22_STIM_PRIMARY_BRONCHIAL_EPITHELIAL_CELLS_UP | GSE10240_CTRL_VS_IL22_STIM_PRIMARY_BRONCHIAL_EPITHELIAL_CELLS_UP | 15/611 | 200/21355 | 0.0007 | 0.00851 | 0.00631 | 65125/55690/23370/3683/3106/2081/79718/3480/55605/860/4363/23102/9466/155038/7267 | 15 |
| GSE11961_FOLLICULAR_BCELL_VS_GERMINAL_CENTER_BCELL_DAY40_UP | GSE11961_FOLLICULAR_BCELL_VS_GERMINAL_CENTER_BCELL_DAY40_UP | 15/611 | 200/21355 | 0.0007 | 0.00851 | 0.00631 | 64766/11329/55690/8826/55870/26207/976/79745/51/10129/6645/23163/114836/79692/54842 | 15 |
| GSE11961_GERMINAL_CENTER_BCELL_DAY7_VS_PLASMA_CELL_DAY7_UP | GSE11961_GERMINAL_CENTER_BCELL_DAY7_VS_PLASMA_CELL_DAY7_UP | 15/611 | 200/21355 | 0.0007 | 0.00851 | 0.00631 | 64421/586/117583/6526/23077/8287/3607/5933/30844/5888/7798/64783/1822/2289/81671 | 15 |
| GSE11961_MARGINAL_ZONE_BCELL_VS_GERMINAL_CENTER_BCELL_DAY7_DN | GSE11961_MARGINAL_ZONE_BCELL_VS_GERMINAL_CENTER_BCELL_DAY7_DN | 15/611 | 200/21355 | 0.0007 | 0.00851 | 0.00631 | 2869/1119/55852/54838/22955/9693/57410/5933/5888/7798/57534/79632/2289/80025/81671 | 15 |
| GSE12392_WT_VS_IFNAR_KO_CD8A_NEG_SPLEEN_DC_UP | GSE12392_WT_VS_IFNAR_KO_CD8A_NEG_SPLEEN_DC_UP | 15/611 | 200/21355 | 0.0007 | 0.00851 | 0.00631 | 23064/29980/337867/91775/860/1432/1606/9712/6197/80728/9020/8428/9844/84433/27244 | 15 |
| GSE13484_12H_UNSTIM_VS_YF17D_VACCINE_STIM_PBMC_UP | GSE13484_12H_UNSTIM_VS_YF17D_VACCINE_STIM_PBMC_UP | 15/611 | 200/21355 | 0.0007 | 0.00851 | 0.00631 | 26065/8826/55852/23499/7699/138151/79036/8289/157680/23198/8031/9949/60685/4354/83451 | 15 |
| GSE15330_WT_VS_IKAROS_KO_GRANULOCYTE_MONOCYTE_PROGENITOR_UP | GSE15330_WT_VS_IKAROS_KO_GRANULOCYTE_MONOCYTE_PROGENITOR_UP | 15/611 | 200/21355 | 0.0007 | 0.00851 | 0.00631 | 944/146712/9567/79602/5820/59269/3106/197322/1520/3157/50650/976/84181/8672/114836 | 15 |
| GSE15330_WT_VS_IKAROS_KO_HSC_UP | GSE15330_WT_VS_IKAROS_KO_HSC_UP | 15/611 | 200/21355 | 0.0007 | 0.00851 | 0.00631 | 7414/53944/1520/5339/9611/5880/10767/51/10905/26133/54870/1739/55023/4354/9953 | 15 |
| GSE15930_STIM_VS_STIM_AND_IFNAB_48H_CD8_T_CELL_UP | GSE15930_STIM_VS_STIM_AND_IFNAB_48H_CD8_T_CELL_UP | 15/611 | 200/21355 | 0.0007 | 0.00851 | 0.00631 | 64766/4670/1265/25777/84316/1455/79828/54838/255231/9960/5295/5880/1606/5305/26133 | 15 |
| GSE15930_STIM_VS_STIM_AND_IL12_48H_CD8_T_CELL_UP | GSE15930_STIM_VS_STIM_AND_IL12_48H_CD8_T_CELL_UP | 15/611 | 200/21355 | 0.0007 | 0.00851 | 0.00631 | 4670/23214/84196/84316/1520/23139/26207/1606/976/5770/3454/6891/55275/11184/473 | 15 |
| GSE16450_CTRL_VS_IFNA_6H_STIM_IMMATURE_NEURON_CELL_LINE_UP | GSE16450_CTRL_VS_IFNA_6H_STIM_IMMATURE_NEURON_CELL_LINE_UP | 15/611 | 200/21355 | 0.0007 | 0.00851 | 0.00631 | 10672/64421/58513/11320/26207/9693/5925/3077/324/10352/64968/23527/57534/9459/7267 | 15 |
| GSE17721_CPG_VS_GARDIQUIMOD_0.5H_BMDC_DN | GSE17721_CPG_VS_GARDIQUIMOD_0.5H_BMDC_DN | 15/611 | 200/21355 | 0.0007 | 0.00851 | 0.00631 | 377/171023/23370/987/2909/7187/1786/25981/9611/10425/51586/7531/128866/55758/79657 | 15 |
| GSE17721_CPG_VS_GARDIQUIMOD_6H_BMDC_DN | GSE17721_CPG_VS_GARDIQUIMOD_6H_BMDC_DN | 15/611 | 200/21355 | 0.0007 | 0.00851 | 0.00631 | 64766/5496/11329/27340/9736/79602/23071/89970/2926/53944/124245/10055/57410/9129/23032 | 15 |
| GSE19401_UNSTIM_VS_PAM2CSK4_STIM_FOLLICULAR_DC_DN | GSE19401_UNSTIM_VS_PAM2CSK4_STIM_FOLLICULAR_DC_DN | 15/611 | 200/21355 | 0.0007 | 0.00851 | 0.00631 | 23064/55619/965/2926/10036/6894/5925/3660/4277/22978/259230/4700/23607/64750/4354 | 15 |
| GSE20198_IL12_IL18_VS_IFNA_TREATED_ACT_CD4_TCELL_DN | GSE20198_IL12_IL18_VS_IFNA_TREATED_ACT_CD4_TCELL_DN | 15/611 | 200/21355 | 0.0007 | 0.00851 | 0.00631 | 4670/4775/6774/58513/54834/3747/9782/51696/51455/51/114804/29761/8726/4642/9673 | 15 |

| | | | | | | | | |
|---|---|---|---|---|---|---|---|---|
| GSE21546_WT_VS_SAP1A_KO_ANTI_CD3_STIM_DP_THYMOCYTES_DN | GSE21546_WT_VS_SAP1A_KO_ANTI_CD3_STIM_DP_THYMOCYTES_DN | 15/611 | 200/21355 | 0.0007 | 0.00851 | 0.00631 | 1387/9736/7421/80196/55870/4363/8904/55236/2521/54870/10454/6891/1399/7109/57534 | 15 |
| GSE21670_TGFB_VS_TGFB_AND_IL6_TREATED_CD4_TCELL_UP | GSE21670_TGFB_VS_TGFB_AND_IL6_TREATED_CD4_TCELL_UP | 15/611 | 200/21355 | 0.0007 | 0.00851 | 0.00631 | 23181/29028/641/55421/3157/79801/9712/5578/1739/115/5073/57534/124446/56942/10330 | 15 |
| GSE21774_CD62L_POS_CD56_BRIGHT_VS_CD62L_NEG_CD56_DIM_NK_CELL_DN | GSE21774_CD62L_POS_CD56_BRIGHT_VS_CD62L_NEG_CD56_DIM_NK_CELL_DN | 15/611 | 200/21355 | 0.0007 | 0.00851 | 0.00631 | 64766/26065/155435/1729/3655/26207/155038/84181/29761/2035/1911/7982/57198/2643/79632 | 15 |
| GSE22432_MULTIPOTENT_PROGENITOR_VS_CDC_DN | GSE22432_MULTIPOTENT_PROGENITOR_VS_CDC_DN | 15/611 | 200/21355 | 0.0007 | 0.00851 | 0.00631 | 3936/26574/57102/23511/91942/23406/1289/283209/2521/7443/11163/92181/4642/10198/5906 | 15 |
| GSE22601_DOUBLE_POSITIVE_VS_CD8_SINGLE_POSITIVE_THYMOCYTE_UP | GSE22601_DOUBLE_POSITIVE_VS_CD8_SINGLE_POSITIVE_THYMOCYTE_UP | 15/611 | 200/21355 | 0.0007 | 0.00851 | 0.00631 | 85464/11329/25777/155435/337867/91775/3707/1606/10425/26133/5578/55500/114836/54842/9953 | 15 |
| GSE22886_DC_VS_MONOCYTE_DN | GSE22886_DC_VS_MONOCYTE_DN | 15/611 | 200/21355 | 0.0007 | 0.00851 | 0.00631 | 1387/11329/56913/1105/3683/22861/23077/8073/3480/51108/9960/976/11064/2643/23287 | 15 |
| GSE22886_IGG_IGA_MEMORY_BCELL_VS_BM_PLASMA_CELL_UP | GSE22886_IGG_IGA_MEMORY_BCELL_VS_BM_PLASMA_CELL_UP | 15/611 | 200/21355 | 0.0007 | 0.00851 | 0.00631 | 6777/11329/23370/23369/23347/23515/6894/54878/29761/10129/1911/57711/8428/57337/147657 | 15 |
| GSE22886_NAIVE_CD4_TCELL_VS_NEUTROPHIL_UP | GSE22886_NAIVE_CD4_TCELL_VS_NEUTROPHIL_UP | 15/611 | 200/21355 | 0.0007 | 0.00851 | 0.00631 | 54934/25777/4733/26528/23499/5295/6767/10425/10767/26036/7644/5578/51176/10198/23450 | 15 |
| GSE23568_CTRL_VS_ID3_TRANSDUCED_CD8_TCELL_UP | GSE23568_CTRL_VS_ID3_TRANSDUCED_CD8_TCELL_UP | 15/611 | 200/21355 | 0.0007 | 0.00851 | 0.00631 | 1073/1387/23077/1455/55870/3655/10048/1432/5977/1606/9466/976/9402/23607/10963 | 15 |
| GSE24634_TEFF_VS_TCONV_DAY10_IN_CULTURE_UP | GSE24634_TEFF_VS_TCONV_DAY10_IN_CULTURE_UP | 15/611 | 200/21355 | 0.0007 | 0.00851 | 0.00631 | 2869/672/55252/23150/701/79828/10615/3560/23048/79801/5888/7443/6197/55784/2590 | 15 |
| GSE25123_IL4_VS_IL4_AND_ROSIGLITAZONE_STIM_MACROPHAGE_DAY10_UP | GSE25123_IL4_VS_IL4_AND_ROSIGLITAZONE_STIM_MACROPHAGE_DAY10_UP | 15/611 | 200/21355 | 0.0007 | 0.00851 | 0.00631 | 9735/944/401409/672/701/7421/860/10892/10615/7455/30844/9727/114836/10800/81671 | 15 |
| GSE26343_UNSTIM_VS_LPS_STIM_MACROPHAGE_DN | GSE26343_UNSTIM_VS_LPS_STIM_MACROPHAGE_DN | 15/611 | 200/21355 | 0.0007 | 0.00851 | 0.00631 | 65059/150864/10111/701/89846/6249/10163/1432/2885/3077/64333/4700/10906/5073/4354 | 15 |
| GSE26669_CTRL_VS_COSTIM_BLOCK_MLR_CD4_TCELL_UP | GSE26669_CTRL_VS_COSTIM_BLOCK_MLR_CD4_TCELL_UP | 15/611 | 200/21355 | 0.0007 | 0.00851 | 0.00631 | 59269/7173/11320/84166/79828/55852/1786/5880/3660/9918/6891/128866/1822/55275/10963 | 15 |
| GSE2770_UNTREATED_VS_ACT_CD4_TCELL_6H_DN | GSE2770_UNTREATED_VS_ACT_CD4_TCELL_6H_DN | 15/611 | 200/21355 | 0.0007 | 0.00851 | 0.00631 | 8844/117583/256435/9851/84458/3710/65117/23355/23365/84937/283131/9020/163702/5514/7267 | 15 |
| GSE27786_CD8_TCELL_VS_NKTCELL_DN | GSE27786_CD8_TCELL_VS_NKTCELL_DN | 15/611 | 200/21355 | 0.0007 | 0.00851 | 0.00631 | 51552/65059/3683/55619/23150/55852/1729/5533/23139/4627/23048/54815/976/114804/65979 | 15 |
| GSE27786_ERYTHROBLAST_VS_NEUTROPHIL_DN | GSE27786_ERYTHROBLAST_VS_NEUTROPHIL_DN | 15/611 | 200/21355 | 0.0007 | 0.00851 | 0.00631 | 55187/23071/91775/4297/53944/23059/54838/4795/6767/3660/23443/6197/84937/1822/83451 | 15 |
| GSE28726_ACT_CD4_TCELL_VS_ACT_VA24NEG_NKTCELL_DN | GSE28726_ACT_CD4_TCELL_VS_ACT_VA24NEG_NKTCELL_DN | 15/611 | 200/21355 | 0.0007 | 0.00851 | 0.00631 | 7534/9873/7799/23370/9267/26065/9736/25777/3707/5775/3655/120/157680/51176/23524 | 15 |
| GSE29164_UNTREATED_VS_CD8_TCELL_AND_IL12_TREATED_MELANOMA_DAY7_UP | GSE29164_UNTREATED_VS_CD8_TCELL_AND_IL12_TREATED_MELANOMA_DAY7_UP | 15/611 | 200/21355 | 0.0007 | 0.00851 | 0.00631 | 23112/23731/7414/57585/23077/1520/165918/221477/9693/5933/157680/3454/6197/11163/55275 | 15 |
| GSE30083_SP2_VS_SP4_THYMOCYTE_DN | GSE30083_SP2_VS_SP4_THYMOCYTE_DN | 15/611 | 200/21355 | 0.0007 | 0.00851 | 0.00631 | 727/3106/4297/727957/7421/5533/1520/3655/3560/23443/976/6645/114836/55784/9953 | 15 |
| GSE31082_DN_VS_CD8_SP_THYMOCYTE_UP | GSE31082_DN_VS_CD8_SP_THYMOCYTE_UP | 15/611 | 200/21355 | 0.0007 | 0.00851 | 0.00631 | 11052/4670/586/26574/10111/26528/3157/55236/10767/30844/2961/375748/92912/7531/1871 | 15 |
| GSE32986_CURDLAN_LOWDOSE_VS_CURDLAN_HIGHDOSE_STIM_DC_UP | GSE32986_CURDLAN_LOWDOSE_VS_CURDLAN_HIGHDOSE_STIM_DC_UP | 15/611 | 200/21355 | 0.0007 | 0.00851 | 0.00631 | 85464/65059/6938/157378/3655/1786/23303/8289/324/64333/23527/65979/79157/25938/27244 | 15 |
| GSE34392_ST2_KO_VS_WT_DAY8_LCMV_EFFECTOR_CD8_TCELL_UP | GSE34392_ST2_KO_VS_WT_DAY8_LCMV_EFFECTOR_CD8_TCELL_UP | 15/611 | 200/21355 | 0.0007 | 0.00851 | 0.00631 | 85464/23370/150864/9889/6938/7554/120/64333/30844/4700/8672/80728/84433/473/27244 | 15 |
| GSE39820_TGFBETA1_VS_TGFBETA3_IN_IL6_TREATED_CD4_TCELL_UP | GSE39820_TGFBETA1_VS_TGFBETA3_IN_IL6_TREATED_CD4_TCELL_UP | 15/611 | 200/21355 | 0.0007 | 0.00851 | 0.00631 | 7534/9873/9267/26065/8826/89970/2081/6249/57410/26036/51/5090/7170/10905/23355 | 15 |

| | | | | | | | | |
|---|---|---|---|---|---|---|---|---|
| GSE40274_CTRL_VS_FOXP3_TRANSDUCED_ACTIVATED_CD4_TCELL_DN | GSE40274_CTRL_VS_FOXP3_TRANSDUCED_ACTIVATED_CD4_TCELL_DN | 15/611 | 200/21355 | 0.0007 | 0.00851 | 0.00631 | 23201/56261/23370/3106/11320/23077/1203/83478/9960/50650/378938/9466/976/2521/155038 | 15 |
| GSE40277_EOS_AND_LEF1_TRANSDUCED_VS_CTRL_CD4_TCELL_UP | GSE40277_EOS_AND_LEF1_TRANSDUCED_VS_CTRL_CD4_TCELL_UP | 15/611 | 200/21355 | 0.0007 | 0.00851 | 0.00631 | 60468/11329/29980/79828/3660/9962/3560/976/9807/5900/9712/6891/197135/9694/473 | 15 |
| GSE40666_WT_VS_STAT4_KO_CD8_TCELL_WITH_IFNA_STIM_90MIN_UP | GSE40666_WT_VS_STAT4_KO_CD8_TCELL_WITH_IFNA_STIM_90MIN_UP | 15/611 | 200/21355 | 0.0007 | 0.00851 | 0.00631 | 23112/60468/11329/55690/23214/1265/138151/976/148867/23527/259230/50852/4354/9694/84636 | 15 |
| GSE411_UNSTIM_VS_100MIN_IL6_STIM_SOCS3_KO_MACROPHAGE_UP | GSE411_UNSTIM_VS_100MIN_IL6_STIM_SOCS3_KO_MACROPHAGE_UP | 15/611 | 200/21355 | 0.0007 | 0.00851 | 0.00631 | 4204/256435/155435/22848/7187/55870/8631/5880/7745/7040/84181/55023/9459/83451/11184 | 15 |
| GSE41867_DAY6_EFFECTOR_VS_DAY30_EXHAUSTED_CD8_TCELL_LCMV_CLONE13_UP | GSE41867_DAY6_EFFECTOR_VS_DAY30_EXHAUSTED_CD8_TCELL_LCMV_CLONE13_UP | 15/611 | 200/21355 | 0.0007 | 0.00851 | 0.00631 | 585/401409/3106/23077/7009/23303/138151/51586/3560/9807/3594/10906/124446/2643/473 | 15 |
| GSE41867_MEMORY_VS_EXHAUSTED_CD8_TCELL_DAY30_LCMV_DN | GSE41867_MEMORY_VS_EXHAUSTED_CD8_TCELL_DAY30_LCMV_DN | 15/611 | 200/21355 | 0.0007 | 0.00851 | 0.00631 | 6777/64766/56005/7414/23071/4245/3480/1606/22806/1954/51/25853/57198/2590/5906 | 15 |
| GSE43863_TH1_VS_TFH_MEMORY_CD4_TCELL_DN | GSE43863_TH1_VS_TFH_MEMORY_CD4_TCELL_DN | 15/611 | 200/21355 | 0.0007 | 0.00851 | 0.00631 | 23370/7414/1265/4245/22992/89846/6780/1455/5339/10163/4627/120/3710/50650/473 | 15 |
| GSE44649_NAIVE_VS_ACTIVATED_CD8_TCELL_MIR155_KO_DN | GSE44649_NAIVE_VS_ACTIVATED_CD8_TCELL_MIR155_KO_DN | 15/611 | 200/21355 | 0.0007 | 0.00851 | 0.00631 | 51742/10672/56913/987/64848/3683/2926/55972/55193/196074/6197/4931/6873/64750/114836 | 15 |
| GSE5589_UNSTIM_VS_45MIN_LPS_AND_IL6_STIM_MACROPHAGE_UP | GSE5589_UNSTIM_VS_45MIN_LPS_AND_IL6_STIM_MACROPHAGE_UP | 15/611 | 200/21355 | 0.0007 | 0.00851 | 0.00631 | 9267/55010/22848/4363/2885/55193/79801/10129/79058/23013/57198/10800/2590/473/27244 | 15 |
| GSE6259_FLT3L_INDUCED_33D1_POS_DC_VS_BCELL_DN | GSE6259_FLT3L_INDUCED_33D1_POS_DC_VS_BCELL_DN | 15/611 | 200/21355 | 0.0007 | 0.00851 | 0.00631 | 56261/64766/9873/1729/1203/1786/4215/9918/3454/23019/259230/55023/60685/80331/80025 | 15 |
| GSE6674_PL2_3_VS_ANTI_IGM_AND_CPG_STIM_BCELL_DN | GSE6674_PL2_3_VS_ANTI_IGM_AND_CPG_STIM_BCELL_DN | 15/611 | 200/21355 | 0.0007 | 0.00851 | 0.00631 | 65125/85464/7799/11320/55619/55852/3655/26207/23141/50650/26133/55818/25938/10800/23287 | 15 |
| GSE7460_FOXP3_MUT_VS_WT_ACT_TCONV_UP | GSE7460_FOXP3_MUT_VS_WT_ACT_TCONV_UP | 15/611 | 200/21355 | 0.0007 | 0.00851 | 0.00631 | 9873/5148/6774/4245/23150/860/4363/7813/549/3594/23198/283989/10800/5906/81671 | 15 |
| GSE7509_DC_VS_MONOCYTE_WITH_FCGRIIB_STIM_UP | GSE7509_DC_VS_MONOCYTE_WITH_FCGRIIB_STIM_UP | 15/611 | 200/21355 | 0.0007 | 0.00851 | 0.00631 | 23112/6777/23130/8073/2909/7187/54838/10144/1432/9918/6599/23248/9020/7267/26043 | 15 |
| GSE7831_UNSTIM_VS_CPG_STIM_PDC_1H_UP | GSE7831_UNSTIM_VS_CPG_STIM_PDC_1H_UP | 15/611 | 200/21355 | 0.0007 | 0.00851 | 0.00631 | 56913/3683/6938/59269/401409/9967/9693/7920/4627/5305/3594/259230/6197/57198/54521 | 15 |
| GSE8685_IL2_STARVED_VS_IL21_ACT_IL2_STARVED_CD4_TCELL_DN | GSE8685_IL2_STARVED_VS_IL21_ACT_IL2_STARVED_CD4_TCELL_DN | 15/611 | 200/21355 | 0.0007 | 0.00851 | 0.00631 | 64766/9135/9887/7799/29980/89846/9728/1455/5775/255231/120/7170/6645/1911/163486 | 15 |
| GSE9650_EFFECTOR_VS_MEMORY_CD8_TCELL_DN | GSE9650_EFFECTOR_VS_MEMORY_CD8_TCELL_DN | 15/611 | 200/21355 | 0.0007 | 0.00851 | 0.00631 | 54934/944/23077/1520/3707/6223/255231/8289/57649/7443/6645/84937/23607/473/27244 | 15 |
| GSE9988_LPS_VS_CTRL_TREATED_MONOCYTE_DN | GSE9988_LPS_VS_CTRL_TREATED_MONOCYTE_DN | 15/611 | 200/21355 | 0.0007 | 0.00851 | 0.00631 | 51696/8073/23774/54838/9960/10479/10163/1432/54878/7769/10992/25853/9020/284058/80331 | 15 |
| GSE40274_FOXP3_VS_FOXP3_AND_XBP1_TRANSDUCED_ACTIVATED_CD4_TCELL_DN | GSE40274_FOXP3_VS_FOXP3_AND_XBP1_TRANSDUCED_ACTIVATED_CD4_TCELL_DN | 13/611 | 159/21355 | 0.0007 | 0.00855 | 0.00635 | 7046/146057/987/255231/10767/30844/8672/51176/120425/114836/124446/4354/55784 | 13 |
| GSE37301_CD4_TCELL_VS_RAG2_KO_NK_CELL_DN | GSE37301_CD4_TCELL_VS_RAG2_KO_NK_CELL_DN | 14/611 | 180/21355 | 0.0007 | 0.00886 | 0.00657 | 23181/55690/79602/57690/9728/79813/79663/5339/23186/9797/7009/9611/7443/57534 | 14 |
| GSE7768_OVA_ALONE_VS_OVA_WITH_MPL_IMMUNIZED_MOUSE_WHOLE_SPLEEN_6H_UP | GSE7768_OVA_ALONE_VS_OVA_WITH_MPL_IMMUNIZED_MOUSE_WHOLE_SPLEEN_6H_UP | 14/611 | 181/21355 | 0.0007 | 0.00933 | 0.00692 | 171023/5496/23304/7046/6526/79718/701/1060/9918/23048/22978/64783/63977/27244 | 14 |
| GSE7596_AKT_TRANSD_VS_CTRL_CD4_TCONV_WITH_TGFB_UP | GSE7596_AKT_TRANSD_VS_CTRL_CD4_TCONV_WITH_TGFB_UP | 13/611 | 161/21355 | 0.0008 | 0.00955 | 0.00708 | 1540/4820/641/5925/79801/7644/23198/10096/6645/54521/114836/493/7403 | 13 |
| GSE7348_LPS_VS_TOLERIZED_AND_LPS_STIM_MACROPHAGE_UP | GSE7348_LPS_VS_TOLERIZED_AND_LPS_STIM_MACROPHAGE_UP | 14/611 | 182/21355 | 0.0008 | 0.0098 | 0.00727 | 9135/9698/57585/7326/22887/7421/124245/22908/23048/23102/29123/83891/259230/115 | 14 |
| GSE37533_UNTREATED_VS_PIOGLIZATONE_TREATED_CD4_TCELL_FOXP3_TRASDUCED_CD4_TCELL_DN | GSE37533_UNTREATED_VS_PIOGLIZATONE_TREATED_CD4_TCELL_FOXP3_TRASDUCED_CD4_TCELL_DN | 13/611 | 162/21355 | 0.0008 | 0.01006 | 0.00747 | 10672/28977/146057/9567/6249/10892/6223/5977/2885/3077/5305/2175/124446 | 13 |
| NAKAYA_B_CELL_FLUARIX_FLUVIRIN_AGE_18_50YO_7DY_DN | NAKAYA_B_CELL_FLUARIX_FLUVIRIN_AGE_18_50YO_7DY_DN | 19/611 | 294/21355 | 0.0009 | 0.01087 | 0.00806 | 60468/4763/4255/64421/22828/146057/4775/9736/58513/79718/8289/7813/29123/7040/2961/23013/1822/9844/51281 | 19 |

| | | | | | | | | |
|---|---|---|---|---|---|---|---|---|
| GSE37563_WT_VS_CTLA4_KO_CD4_TCELL_D4_POST_IMMUNIZATION_UP | GSE37563_WT_VS_CTLA4_KO_CD4_TCELL_D4_POST_IMMUNIZATION_UP | 13/611 | 165/21355 | 0.001 | 0.01187 | 0.00881 | 23167/23064/91775/89846/727957/10479/146691/7644/55291/22866/51735/9402/10521 | 13 |
| GSE13946_CTRL_VS_DSS_COLITIS_GD_TCELL_FROM_COLON_DN | GSE13946_CTRL_VS_DSS_COLITIS_GD_TCELL_FROM_COLON_DN | 13/611 | 170/21355 | 0.0013 | 0.01553 | 0.01153 | 944/10111/51108/79828/79663/3157/3655/22955/50852/51176/23607/27244/7267 | 13 |
| ZAK_PBMC_MRKAD5_HIV_1_GAG_POL_NEF_AGE_20_50YO_AD5_NAB_TITERS_GTE_200_VS_LTE_200_1DY_UP | ZAK_PBMC_MRKAD5_HIV_1_GAG_POL_NEF_AGE_20_50YO_AD5_NAB_TITERS_GTE_200_VS_LTE_200_1DY_UP | 13/611 | 170/21355 | 0.0013 | 0.01553 | 0.01153 | 9736/11320/3480/3655/8631/7745/9807/114804/10198/84636/84441/27244/7267 | 13 |
| GSE5099_DAY3_VS_DAY7_MCSF_TREATED_MACROPHAGE_UP | GSE5099_DAY3_VS_DAY7_MCSF_TREATED_MACROPHAGE_UP | 14/611 | 192/21355 | 0.0013 | 0.01618 | 0.01201 | 64766/7414/26157/64848/51696/84166/6894/23406/55972/10479/120/3594/155038/9263 | 14 |
| GSE13547_CTRL_VS_ANTI_IGM_STIM_ZFX_KO_BCELL_2H_DN | GSE13547_CTRL_VS_ANTI_IGM_STIM_ZFX_KO_BCELL_2H_DN | 13/611 | 171/21355 | 0.0013 | 0.01618 | 0.01201 | 132789/2869/28977/5820/672/1520/165918/221477/3183/9962/56852/120425/23607 | 13 |
| GSE15624_3H_VS_6H_HALOFUGINONE_TREATED_CD4_TCELL_UP | GSE15624_3H_VS_6H_HALOFUGINONE_TREATED_CD4_TCELL_UP | 13/611 | 171/21355 | 0.0013 | 0.01618 | 0.01201 | 65125/54834/57585/9967/5339/8631/5880/3077/3560/2521/84937/57198/493 | 13 |
| GSE6269_FLU_VS_STREP_PNEUMO_INF_PBMC_UP | GSE6269_FLU_VS_STREP_PNEUMO_INF_PBMC_UP | 13/611 | 171/21355 | 0.0013 | 0.01618 | 0.01201 | 28977/4763/9887/23131/23064/3106/4297/51586/3594/23060/23607/4134/2643 | 13 |
| GSE7568_IL4_VS_IL4_AND_DEXAMETHASONE_TREATED_MACROPHAGE_UP | GSE7568_IL4_VS_IL4_AND_DEXAMETHASONE_TREATED_MACROPHAGE_UP | 13/611 | 171/21355 | 0.0013 | 0.01618 | 0.01201 | 28977/23649/9735/586/3936/91942/1060/3183/26133/56852/1871/6873/9377 | 13 |
| GSE9509_LPS_VS_LPS_AND_IL10_STIM_IL10_KO_MACROPHAGE_30MIN_DN | GSE9509_LPS_VS_LPS_AND_IL10_STIM_IL10_KO_MACROPHAGE_30MIN_DN | 14/611 | 193/21355 | 0.0014 | 0.01681 | 0.01247 | 85464/1387/23649/944/23047/2272/5533/10048/54899/1606/1289/4277/51176/7403 | 14 |
| GSE17974_IL4_AND_ANTI_IL12_VS_UNTREATED_12H_ACT_CD4_TCELL_UP | GSE17974_IL4_AND_ANTI_IL12_VS_UNTREATED_12H_ACT_CD4_TCELL_UP | 14/611 | 194/21355 | 0.0014 | 0.01716 | 0.01273 | 56261/9202/7414/23633/200424/50807/10144/79956/817/22866/54521/54842/6641/81671 | 14 |
| GSE21927_UNTREATED_VS_GMCSF_IL6_TREATED_BONE_MARROW_UP | GSE21927_UNTREATED_VS_GMCSF_IL6_TREATED_BONE_MARROW_UP | 14/611 | 194/21355 | 0.0014 | 0.01716 | 0.01273 | 7534/5496/197322/84830/55421/23499/284338/50650/55236/5578/54617/83990/55275/2289 | 14 |
| GSE25087_TREG_VS_TCONV_FETUS_DN | GSE25087_TREG_VS_TCONV_FETUS_DN | 14/611 | 194/21355 | 0.0014 | 0.01716 | 0.01273 | 10390/7414/7554/337867/7182/6929/26207/79956/120/3454/5165/5094/9263/51720 | 14 |
| GSE2770_TGFB_AND_IL4_ACT_VS_ACT_CD4_TCELL_2H_DN | GSE2770_TGFB_AND_IL4_ACT_VS_ACT_CD4_TCELL_2H_DN | 14/611 | 194/21355 | 0.0014 | 0.01716 | 0.01273 | 56913/9567/23347/3660/50650/4277/135112/26133/29761/55683/65979/10906/2643/7403 | 14 |
| GSE10211_UV_INACT_SENDAI_VS_LIVE_SENDAI_VIRUS_TRACHEAL_EPITHELIAL_CELLS_UP | GSE10211_UV_INACT_SENDAI_VS_LIVE_SENDAI_VIRUS_TRACHEAL_EPITHELIAL_CELLS_UP | 12/611 | 152/21355 | 0.0014 | 0.01716 | 0.01273 | 56261/9567/4215/23499/29123/259230/10788/51176/23607/54617/25938/2590 | 12 |
| GSE33292_DN3_THYMOCYTE_VS_TCELL_LYMPHOMA_FROM_TCF1_KO_UP | GSE33292_DN3_THYMOCYTE_VS_TCELL_LYMPHOMA_FROM_TCF1_KO_UP | 13/611 | 173/21355 | 0.0015 | 0.01716 | 0.01273 | 545/4775/23077/91775/378805/9491/54165/23198/10788/92912/2590/6641/51720 | 13 |
| GSE7219_UNSTIM_VS_LPS_AND_ANTI_CD40_STIM_DC_UP | GSE7219_UNSTIM_VS_LPS_AND_ANTI_CD40_STIM_DC_UP | 13/611 | 173/21355 | 0.0015 | 0.01716 | 0.01273 | 10672/85464/60468/944/5295/1606/10788/51176/23607/114836/54842/9953/23032 | 13 |
| GSE20484_MCSG_VS_CXCL4_MONOCYTE_DERIVED_MACROPHAGE_DN | GSE20484_MCSG_VS_CXCL4_MONOCYTE_DERIVED_MACROPHAGE_DN | 14/611 | 195/21355 | 0.0015 | 0.01716 | 0.01273 | 6777/23181/64421/23064/23369/3480/22908/5305/26036/54165/157680/117584/23365/57337 | 14 |
| GSE34205_RSV_VS_FLU_INF_INFANT_PBMC_DN | GSE34205_RSV_VS_FLU_INF_INFANT_PBMC_DN | 14/611 | 195/21355 | 0.0015 | 0.01716 | 0.01273 | 51552/4763/585/23064/3683/3106/55605/5775/7009/128710/26133/23060/283989/219988 | 14 |
| GSE3982_MAST_CELL_VS_MAC_DN | GSE3982_MAST_CELL_VS_MAC_DN | 14/611 | 195/21355 | 0.0015 | 0.01716 | 0.01273 | 9873/7046/59269/9976/55852/1520/51271/5933/9466/79745/51/1822/79982/2289 | 14 |
| GSE22886_NAIVE_CD4_TCELL_VS_NKCELL_UP | GSE22886_NAIVE_CD4_TCELL_VS_NKCELL_UP | 14/611 | 196/21355 | 0.0016 | 0.01716 | 0.01273 | 1265/944/3784/84830/2926/10892/22955/1606/10767/5578/51176/4354/104/473 | 14 |
| GSE29617_CTRL_VS_DAY3_TIV_FLU_VACCINE_PBMC_2008_UP | GSE29617_CTRL_VS_DAY3_TIV_FLU_VACCINE_PBMC_2008_UP | 14/611 | 196/21355 | 0.0016 | 0.01716 | 0.01273 | 9873/7799/23130/1540/117583/22887/378805/29123/2035/5094/57711/55818/10521/84441 | 14 |
| GSE37416_0H_VS_24H_F_TULARENSIS_LVS_NEUTROPHIL_UP | GSE37416_0H_VS_24H_F_TULARENSIS_LVS_NEUTROPHIL_UP | 14/611 | 196/21355 | 0.0016 | 0.01716 | 0.01273 | 65125/11329/7414/150864/26157/64848/546/1432/51/7170/64333/6197/51719/23450 | 14 |
| GSE41867_DAY6_VS_DAY15_LCMV_ARMSTRONG_EFFECTOR_CD8_TCELL_DN | GSE41867_DAY6_VS_DAY15_LCMV_ARMSTRONG_EFFECTOR_CD8_TCELL_DN | 14/611 | 196/21355 | 0.0016 | 0.01716 | 0.01273 | 65125/60468/22828/26065/23369/22848/23141/9520/55291/10788/84186/7109/4134/9129 | 14 |
| GSE12366_GC_VS_MEMORY_BCELL_DN | GSE12366_GC_VS_MEMORY_BCELL_DN | 14/611 | 197/21355 | 0.0017 | 0.01716 | 0.01273 | 1105/22861/55619/6249/83478/157378/7009/9960/23499/1606/9466/283209/5770/27244 | 14 |
| GSE14000_UNSTIM_VS_16H_LPS_DC_TRANSLATED_RNA_UP | GSE14000_UNSTIM_VS_16H_LPS_DC_TRANSLATED_RNA_UP | 14/611 | 197/21355 | 0.0017 | 0.01716 | 0.01273 | 7414/4775/84376/22848/6894/50807/5775/7187/4215/79956/138151/64333/9844/7267 | 14 |
| GSE29618_LAIV_VS_TIV_FLU_VACCINE_DAY7_MDC_UP | GSE29618_LAIV_VS_TIV_FLU_VACCINE_DAY7_MDC_UP | 14/611 | 197/21355 | 0.0017 | 0.01716 | 0.01273 | 5496/9698/64324/10497/23515/5295/4627/3077/26036/3454/5578/65979/9949/147657 | 14 |

| | | | | | | | | |
|---|---|---|---|---|---|---|---|---|
| GSE45739_NRAS_KO_VS_WT_UNSTIM_CD4_TCELL_DN | GSE45739_NRAS_KO_VS_WT_UNSTIM_CD4_TCELL_DN | 14/611 | 197/21355 | 0.0017 | 0.01716 | 0.01273 | 27031/89970/729013/6929/120/9962/23102/157680/10129/5094/283131/57337/55758/5073 | 14 |
| GSE9988_ANTI_TREM1_AND_LPS_VS_VEHICLE_TREATED_MONOCYTES_DN | GSE9988_ANTI_TREM1_AND_LPS_VS_VEHICLE_TREATED_MONOCYTES_DN | 14/611 | 197/21355 | 0.0017 | 0.01716 | 0.01273 | 25777/51696/8073/6794/54838/23406/4215/6613/9466/155038/8031/25853/8428/10521 | 14 |
| NAKAYA_PLASMACYTOID_DENDRITIC_CELL_FLUARIX_FLUVIRIN_AGE_18_50YO_7DY_UP | NAKAYA_PLASMACYTOID_DENDRITIC_CELL_FLUARIX_FLUVIRIN_AGE_18_50YO_7DY_UP | 15/611 | 219/21355 | 0.0017 | 0.01716 | 0.01273 | 9889/8826/23633/23059/23406/9611/4850/405/10767/5578/1739/5094/23013/283131/9949 | 15 |
| GSE11057_NAIVE_VS_EFF_MEMORY_CD4_TCELL_UP | GSE11057_NAIVE_VS_EFF_MEMORY_CD4_TCELL_UP | 14/611 | 198/21355 | 0.0017 | 0.01716 | 0.01273 | 60468/23731/150864/200424/3480/6894/3655/146691/5578/51176/22866/51735/4354/84441 | 14 |
| GSE12366_GC_VS_NAIVE_BCELL_UP | GSE12366_GC_VS_NAIVE_BCELL_UP | 14/611 | 198/21355 | 0.0017 | 0.01716 | 0.01273 | 10672/944/7326/701/84316/26207/3183/7443/5094/7531/8726/10198/5073/163486 | 14 |
| GSE15330_LYMPHOID_MULTIPOTENT_VS_GRANULOCYTE_MONOCYTE_PROGENITOR_IKAROS_KO_UP | GSE15330_LYMPHOID_MULTIPOTENT_VS_GRANULOCYTE_MONOCYTE_PROGENITOR_IKAROS_KO_UP | 14/611 | 198/21355 | 0.0017 | 0.01716 | 0.01273 | 54934/944/23511/3480/3157/84961/5305/259230/25853/55023/5073/4791/23450/79657 | 14 |
| GSE1791_CTRL_VS_NEUROMEDINU_IN_T_CELL_LINE_6H_UP | GSE1791_CTRL_VS_NEUROMEDINU_IN_T_CELL_LINE_6H_UP | 14/611 | 198/21355 | 0.0017 | 0.01716 | 0.01273 | 7534/586/23347/6780/51108/79663/1729/9960/50650/55291/375748/7251/25853/8428 | 14 |
| GSE17974_1.5H_VS_72H_IL4_AND_ANTI_IL12_ACT_CD4_TCELL_UP | GSE17974_1.5H_VS_72H_IL4_AND_ANTI_IL12_ACT_CD4_TCELL_UP | 14/611 | 198/21355 | 0.0017 | 0.01716 | 0.01273 | 150864/944/55095/57585/22861/4928/79663/653519/54878/51735/84766/54617/7403/9953 | 14 |
| GSE19198_CTRL_VS_IL21_TREATED_TCELL_6H_DN | GSE19198_CTRL_VS_IL21_TREATED_TCELL_6H_DN | 14/611 | 198/21355 | 0.0017 | 0.01716 | 0.01273 | 85464/23085/6938/823/51696/9967/165918/23303/5900/23198/6645/4354/2289/84441 | 14 |
| GSE22033_UNTREATED_VS_ROSIGLITAZONE_TREATED_MEF_DN | GSE22033_UNTREATED_VS_ROSIGLITAZONE_TREATED_MEF_DN | 14/611 | 198/21355 | 0.0017 | 0.01716 | 0.01273 | 1105/79602/23071/89845/10892/50807/23303/9962/5165/23013/4791/54842/7403/6654 | 14 |
| GSE24142_EARLY_THYMIC_PROGENITOR_VS_DN3_THYMOCYTE_DN | GSE24142_EARLY_THYMIC_PROGENITOR_VS_DN3_THYMOCYTE_DN | 14/611 | 198/21355 | 0.0017 | 0.01716 | 0.01273 | 6938/59269/2272/79828/22848/1606/549/64333/51176/9402/83854/114836/4354/2289 | 14 |
| GSE24634_NAIVE_CD4_TCELL_VS_DAY3_IL4_CONV_TREG_UP | GSE24634_NAIVE_CD4_TCELL_VS_DAY3_IL4_CONV_TREG_UP | 14/611 | 198/21355 | 0.0017 | 0.01716 | 0.01273 | 11329/54934/23370/9267/22861/11320/2272/5533/3655/5295/51460/23355/51176/60685 | 14 |
| GSE2770_TGFB_AND_IL4_ACT_VS_ACT_CD4_TCELL_48H_DN | GSE2770_TGFB_AND_IL4_ACT_VS_ACT_CD4_TCELL_48H_DN | 14/611 | 198/21355 | 0.0017 | 0.01716 | 0.01273 | 7414/256236/8073/7322/84316/51068/11165/120/3594/23248/22834/124446/11184/6641 | 14 |
| GSE2770_TGFB_AND_IL4_ACT_VS_ACT_CD4_TCELL_6H_UP | GSE2770_TGFB_AND_IL4_ACT_VS_ACT_CD4_TCELL_6H_UP | 14/611 | 198/21355 | 0.0017 | 0.01716 | 0.01273 | 65125/25777/9840/51696/10163/8289/50650/5090/1911/57198/22834/284058/57337/10521 | 14 |
| GSE29618_PRE_VS_DAY7_POST_LAIV_FLU_VACCINE_BCELL_UP | GSE29618_PRE_VS_DAY7_POST_LAIV_FLU_VACCINE_BCELL_UP | 14/611 | 198/21355 | 0.0017 | 0.01716 | 0.01273 | 22861/22992/51696/79718/26528/57634/5339/5976/51271/9960/9693/4850/7251/9129 | 14 |
| GSE36476_CTRL_VS_TSST_ACT_72H_MEMORY_CD4_TCELL_YOUNG_UP | GSE36476_CTRL_VS_TSST_ACT_72H_MEMORY_CD4_TCELL_YOUNG_UP | 14/611 | 198/21355 | 0.0017 | 0.01716 | 0.01273 | 1387/54934/23370/1105/9736/4820/11320/51696/10144/9960/5295/120/3710/1911 | 14 |
| GSE37301_COMMON_LYMPHOID_PROGENITOR_VS_RAG2_KO_NK_CELL_DN | GSE37301_COMMON_LYMPHOID_PROGENITOR_VS_RAG2_KO_NK_CELL_DN | 14/611 | 198/21355 | 0.0017 | 0.01716 | 0.01273 | 2869/60468/23167/10390/1540/5820/2926/57634/51/23527/5094/57711/51176/23287 | 14 |
| GSE7460_FOXP3_MUT_VS_WT_ACT_TCONV_DN | GSE7460_FOXP3_MUT_VS_WT_ACT_TCONV_DN | 14/611 | 198/21355 | 0.0017 | 0.01716 | 0.01273 | 944/79828/4967/10425/817/324/148867/9712/10129/23013/51176/23607/2643/9953 | 14 |
| GSE21033_CTRL_VS_POLYIC_STIM_DC_24H_UP | GSE21033_CTRL_VS_POLYIC_STIM_DC_24H_UP | 12/611 | 156/21355 | 0.0018 | 0.01716 | 0.01273 | 85464/55690/1540/5820/89846/6249/84961/26207/4215/23248/552900/83451 | 12 |
| GSE11924_TFH_VS_TH2_CD4_TCELL_UP | GSE11924_TFH_VS_TH2_CD4_TCELL_UP | 14/611 | 199/21355 | 0.0018 | 0.01716 | 0.01273 | 57521/55690/22828/256435/3106/10006/9728/124245/84961/4627/55193/9020/10198/23524 | 14 |
| GSE14308_TH2_VS_TH17_DN | GSE14308_TH2_VS_TH17_DN | 14/611 | 199/21355 | 0.0018 | 0.01716 | 0.01273 | 65125/7799/79602/84301/3747/23077/7322/79663/6929/23141/10425/29123/55683/51720 | 14 |
| GSE15735_CTRL_VS_HDAC_INHIBITOR_TREATED_CD4_TCELL_2H_UP | GSE15735_CTRL_VS_HDAC_INHIBITOR_TREATED_CD4_TCELL_2H_UP | 14/611 | 199/21355 | 0.0018 | 0.01716 | 0.01273 | 4245/22992/91775/197322/83478/6929/126298/817/51460/30844/114804/9712/1871/11184 | 14 |
| GSE15930_NAIVE_VS_48H_IN_VITRO_STIM_IL12_CD8_TCELL_UP | GSE15930_NAIVE_VS_48H_IN_VITRO_STIM_IL12_CD8_TCELL_UP | 14/611 | 199/21355 | 0.0018 | 0.01716 | 0.01273 | 22828/1119/3784/727957/79828/1729/22955/9466/9807/29761/1911/4134/8428/473 | 14 |
| GSE17721_POLYIC_VS_PAM3CSK4_6H_BMDC_DN | GSE17721_POLYIC_VS_PAM3CSK4_6H_BMDC_DN | 14/611 | 199/21355 | 0.0018 | 0.01716 | 0.01273 | 27340/9736/3064/672/4733/2909/124245/51068/51271/1289/10767/1739/115/23450 | 14 |

| | | | | | | | | |
|---|---|---|---|---|---|---|---|---|
| GSE17974_1.5H_VS_72H_IL4_AND_ANTI_IL12_ACT_CD4_TCELL_DN | GSE17974_1.5H_VS_72H_IL4_AND_ANTI_IL12_ACT_CD4_TCELL_DN | 14/611 | 199/21355 | 0.0018 | 0.01716 | 0.01273 | 55526/586/7414/146712/79718/84376/84316/3660/3710/5690/23019/83854/120425/83990 | 14 |
| GSE17974_1H_VS_72H_UNTREATED_IN_VITRO_CD4_TCELL_DN | GSE17974_1H_VS_72H_UNTREATED_IN_VITRO_CD4_TCELL_DN | 14/611 | 199/21355 | 0.0018 | 0.01716 | 0.01273 | 3936/23511/79718/641/4733/2885/3710/23048/79801/8726/2175/10906/10198/10657 | 14 |
| GSE18281_PERIMEDULLARY_CORTICAL_REGION_VS_WHOLE_MEDULLA_THYMUS_UP | GSE18281_PERIMEDULLARY_CORTICAL_REGION_VS_WHOLE_MEDULLA_THYMUS_UP | 14/611 | 199/21355 | 0.0018 | 0.01716 | 0.01273 | 23085/5496/64324/7414/4775/23633/23369/79718/22848/10055/29123/23060/23604/65979 | 14 |
| GSE18281_SUBCAPSULAR_CORTICAL_REGION_VS_WHOLE_MEDULLA_THYMUS_UP | GSE18281_SUBCAPSULAR_CORTICAL_REGION_VS_WHOLE_MEDULLA_THYMUS_UP | 14/611 | 199/21355 | 0.0018 | 0.01716 | 0.01273 | 23085/23649/64324/23633/22861/4245/7554/26528/23476/22848/4215/22955/26133/23524 | 14 |
| GSE19888_CTRL_VS_A3R_ACTIVATION_MAST_CELL_UP | GSE19888_CTRL_VS_A3R_ACTIVATION_MAST_CELL_UP | 14/611 | 199/21355 | 0.0018 | 0.01716 | 0.01273 | 377/23370/23167/81669/25777/84166/9967/860/84961/5690/23019/283131/10521/23450 | 14 |
| GSE21546_WT_VS_SAP1A_KO_AND_ELK1_KO_ANTI_CD3_STIM_DP_THYMOCYTES_UP | GSE21546_WT_VS_SAP1A_KO_AND_ELK1_KO_ANTI_CD3_STIM_DP_THYMOCYTES_UP | 14/611 | 199/21355 | 0.0018 | 0.01716 | 0.01273 | 3936/7414/1540/55187/6249/5295/4627/120/2885/324/79745/283209/135112/8031 | 14 |
| GSE21670_STAT3_KO_VS_WT_CD4_TCELL_DN | GSE21670_STAT3_KO_VS_WT_CD4_TCELL_DN | 14/611 | 199/21355 | 0.0018 | 0.01716 | 0.01273 | 27340/1540/23633/401409/84830/157378/817/5770/114804/84937/22866/55500/493/6641 | 14 |
| GSE22025_PROGESTERONE_VS_TGFB1_AND_PROGESTERONE_TREATED_CD4_TCELL_UP | GSE22025_PROGESTERONE_VS_TGFB1_AND_PROGESTERONE_TREATED_CD4_TCELL_UP | 14/611 | 199/21355 | 0.0018 | 0.01716 | 0.01273 | 2869/9698/9736/6526/546/965/57634/23499/7699/976/9520/8726/10521/104 | 14 |
| GSE22886_NAIVE_CD4_TCELL_VS_12H_ACT_TH1_UP | GSE22886_NAIVE_CD4_TCELL_VS_12H_ACT_TH1_UP | 14/611 | 199/21355 | 0.0018 | 0.01716 | 0.01273 | 23370/26065/55187/23064/4820/22861/23077/10892/8631/120/23048/29761/23355/1911 | 14 |
| GSE22886_NAIVE_TCELL_VS_NEUTROPHIL_UP | GSE22886_NAIVE_TCELL_VS_NEUTROPHIL_UP | 14/611 | 199/21355 | 0.0018 | 0.01716 | 0.01273 | 171023/4670/64848/25777/79718/55619/2926/26528/10892/10055/5295/1606/50650/7267 | 14 |
| GSE23695_CD57_POS_VS_NEG_NK_CELL_UP | GSE23695_CD57_POS_VS_NEG_NK_CELL_UP | 14/611 | 199/21355 | 0.0018 | 0.01716 | 0.01273 | 9202/64324/58513/23077/55010/79828/51807/10479/55193/5515/8672/22834/5073/473 | 14 |
| GSE24972_MARGINAL_ZONE_BCELL_VS_FOLLICULAR_BCELL_IRF8_KO_DN | GSE24972_MARGINAL_ZONE_BCELL_VS_FOLLICULAR_BCELL_IRF8_KO_DN | 14/611 | 199/21355 | 0.0018 | 0.01716 | 0.01273 | 1119/54834/11320/23150/80196/202052/23059/549/10788/9274/25853/197135/54521/2289 | 14 |
| GSE25087_FETAL_VS_ADULT_TCONV_UP | GSE25087_FETAL_VS_ADULT_TCONV_UP | 14/611 | 199/21355 | 0.0018 | 0.01716 | 0.01273 | 9202/57102/5257/965/79828/4363/23406/6929/5305/3594/64968/4791/163486/10963 | 14 |
| GSE26030_TH1_VS_TH17_RESTIMULATED_DAY15_POST_POLARIZATION_UP | GSE26030_TH1_VS_TH17_RESTIMULATED_DAY15_POST_POLARIZATION_UP | 14/611 | 199/21355 | 0.0018 | 0.01716 | 0.01273 | 2869/26157/2909/51108/7182/54838/22955/7409/405/155038/25853/84766/114836/27244 | 14 |
| GSE29618_PRE_VS_DAY7_FLU_VACCINE_MONOCYTE_DN | GSE29618_PRE_VS_DAY7_FLU_VACCINE_MONOCYTE_DN | 14/611 | 199/21355 | 0.0018 | 0.01716 | 0.01273 | 64766/171023/23304/51455/8904/1954/9466/54878/9807/23019/8672/51719/2590/5514 | 14 |
| GSE33162_UNTREATED_VS_4H_LPS_STIM_HDAC3_HET_MACROPHAGE_DN | GSE33162_UNTREATED_VS_4H_LPS_STIM_HDAC3_HET_MACROPHAGE_DN | 14/611 | 199/21355 | 0.0018 | 0.01716 | 0.01273 | 60468/1387/1119/9567/81669/2272/3655/4215/753/26133/51735/473/27244/7267 | 14 |
| GSE360_CTRL_VS_T_GONDII_DC_UP | GSE360_CTRL_VS_T_GONDII_DC_UP | 14/611 | 199/21355 | 0.0018 | 0.01716 | 0.01273 | 51552/23370/7414/9976/1729/3707/10479/1432/4627/3077/1954/8742/23163/104 | 14 |
| GSE369_IFNG_KO_VS_WT_LIVER_UP | GSE369_IFNG_KO_VS_WT_LIVER_UP | 14/611 | 199/21355 | 0.0018 | 0.01716 | 0.01273 | 85464/60468/64421/23304/9267/146057/57690/546/126298/9918/79801/51176/65979/23032 | 14 |
| GSE37416_12H_VS_48H_F_TULARENSIS_LVS_NEUTROPHIL_UP | GSE37416_12H_VS_48H_F_TULARENSIS_LVS_NEUTROPHIL_UP | 14/611 | 199/21355 | 0.0018 | 0.01716 | 0.01273 | 11329/3936/1540/64848/965/6249/4627/10425/976/3454/23198/9020/51719/493 | 14 |
| GSE3982_DC_VS_BASOPHIL_DN | GSE3982_DC_VS_BASOPHIL_DN | 14/611 | 199/21355 | 0.0018 | 0.01716 | 0.01273 | 64766/60468/11329/1540/7322/10892/26207/10163/5977/23198/10198/9459/9129/10800 | 14 |
| GSE3982_MAST_CELL_VS_NKCELL_DN | GSE3982_MAST_CELL_VS_NKCELL_DN | 14/611 | 199/21355 | 0.0018 | 0.01716 | 0.01273 | 171023/7414/9267/79602/25777/9976/2081/5533/83478/3707/8631/9466/29123/2643 | 14 |
| GSE3982_MEMORY_CD4_TCELL_VS_BCELL_UP | GSE3982_MEMORY_CD4_TCELL_VS_BCELL_UP | 14/611 | 199/21355 | 0.0018 | 0.01716 | 0.01273 | 23150/8287/3655/1786/5588/3560/54878/976/10129/50852/9402/9949/23534/163486 | 14 |
| GSE41867_DAY15_EFFECTOR_VS_DAY30_EXHAUSTED_CD8_TCELL_LCMV_CLONE13_UP | GSE41867_DAY15_EFFECTOR_VS_DAY30_EXHAUSTED_CD8_TCELL_LCMV_CLONE13_UP | 14/611 | 199/21355 | 0.0018 | 0.01716 | 0.01273 | 84066/9267/585/401409/727957/23303/10163/138151/3560/9807/135112/10906/51317/473 | 14 |
| GSE6674_ANTI_IGM_VS_ANTI_IGM_AND_CPG_STIM_BCELL_UP | GSE6674_ANTI_IGM_VS_ANTI_IGM_AND_CPG_STIM_BCELL_UP | 14/611 | 199/21355 | 0.0018 | 0.01716 | 0.01273 | 11329/23370/1540/50807/4215/729852/1606/22806/120/5305/976/283209/51317/27244 | 14 |

| | | | | | | | | |
|---|---|---|---|---|---|---|---|---|
| GSE7460_WT_VS_FOXP3_HET_ACT_TCONV_UP | GSE7460_WT_VS_FOXP3_HET_ACT_TCONV_UP | 14/611 | 199/21355 | 0.0018 | 0.01716 | 0.01273 | 85464/155435/79828/5339/10425/148867/283209/9712/10129/50852/51176/284058/57337/2643 | 14 |
| GSE10239_NAIVE_VS_MEMORY_CD8_TCELL_UP | GSE10239_NAIVE_VS_MEMORY_CD8_TCELL_UP | 14/611 | 200/21355 | 0.0019 | 0.01716 | 0.01273 | 60468/51552/23304/10390/9782/51108/7187/6599/23527/10992/50852/6873/23607/80331 | 14 |
| GSE12001_MIR223_KO_VS_WT_NEUTROPHIL_DN | GSE12001_MIR223_KO_VS_WT_NEUTROPHIL_DN | 14/611 | 200/21355 | 0.0019 | 0.01716 | 0.01273 | 9135/51552/4204/26574/6938/57585/4733/23186/79956/57410/55818/7109/55500/79657 | 14 |
| GSE12392_WT_VS_IFNB_KO_CD8A_NEG_SPLEEN_DC_DN | GSE12392_WT_VS_IFNB_KO_CD8A_NEG_SPLEEN_DC_DN | 14/611 | 200/21355 | 0.0019 | 0.01716 | 0.01273 | 65125/85464/64766/1119/23077/8073/1729/5880/5900/84181/11064/114799/5906/27244 | 14 |
| GSE14308_TH17_VS_NATURAL_TREG_DN | GSE14308_TH17_VS_NATURAL_TREG_DN | 14/611 | 200/21355 | 0.0019 | 0.01716 | 0.01273 | 57521/23167/79602/23077/23515/4967/79956/22955/146691/3183/23355/114799/7267/51720 | 14 |
| GSE1432_1H_VS_24H_IFNG_MICROGLIA_DN | GSE1432_1H_VS_24H_IFNG_MICROGLIA_DN | 14/611 | 200/21355 | 0.0019 | 0.01716 | 0.01273 | 377/23304/1540/10111/2926/1786/976/5690/5900/2961/6891/6645/163486/79657 | 14 |
| GSE14350_IL2RB_KO_VS_WT_TEFF_UP | GSE14350_IL2RB_KO_VS_WT_TEFF_UP | 14/611 | 200/21355 | 0.0019 | 0.01716 | 0.01273 | 85464/60468/944/79602/3480/1520/23139/255231/135112/84937/4012/4791/84636/473 | 14 |
| GSE1460_DP_VS_CD4_THYMOCYTE_DN | GSE1460_DP_VS_CD4_THYMOCYTE_DN | 14/611 | 200/21355 | 0.0019 | 0.01716 | 0.01273 | 7414/146712/22861/3106/7322/1729/7187/23406/79956/7699/3560/5578/9263/104 | 14 |
| GSE14908_RESTING_VS_HDM_STIM_CD4_TCELL_NONATOPIC_PATIENT_DN | GSE14908_RESTING_VS_HDM_STIM_CD4_TCELL_NONATOPIC_PATIENT_DN | 14/611 | 200/21355 | 0.0019 | 0.01716 | 0.01273 | 57585/337867/5257/2909/727957/1729/5533/23186/54838/10163/1778/29761/84441/23032 | 14 |
| GSE15324_NAIVE_VS_ACTIVATED_ELF4_KO_CD8_TCELL_UP | GSE15324_NAIVE_VS_ACTIVATED_ELF4_KO_CD8_TCELL_UP | 14/611 | 200/21355 | 0.0019 | 0.01716 | 0.01273 | 1387/4204/9567/8729/9693/3710/9807/5900/11064/4700/79058/9020/10198/60685 | 14 |
| GSE16451_CTRL_VS_WEST_EQUINE_ENC_VIRUS_IMMATURE_NEURON_CELL_LINE_UP | GSE16451_CTRL_VS_WEST_EQUINE_ENC_VIRUS_IMMATURE_NEURON_CELL_LINE_UP | 14/611 | 200/21355 | 0.0019 | 0.01716 | 0.01273 | 150864/51108/6249/138151/5305/552900/2035/51176/1822/23607/64750/10198/54842/9953 | 14 |
| GSE16522_ANTI_CD3CD28_STIM_VS_UNSTIM_MEMORY_CD8_TCELL_DN | GSE16522_ANTI_CD3CD28_STIM_VS_UNSTIM_MEMORY_CD8_TCELL_DN | 14/611 | 200/21355 | 0.0019 | 0.01716 | 0.01273 | 64766/5257/1455/79828/1520/3655/23406/54899/7920/10956/10096/57337/55784/9694 | 14 |
| GSE17186_MEMORY_VS_CD21HIGH_TRANSITIONAL_BCELL_DN | GSE17186_MEMORY_VS_CD21HIGH_TRANSITIONAL_BCELL_DN | 14/611 | 200/21355 | 0.0019 | 0.01716 | 0.01273 | 150864/8826/4245/50807/4967/120/23048/50650/976/26036/3454/2643/51317/27244 | 14 |
| GSE17186_NAIVE_VS_CD21LOW_TRANSITIONAL_BCELL_CORD_BLOOD_DN | GSE17186_NAIVE_VS_CD21LOW_TRANSITIONAL_BCELL_CORD_BLOOD_DN | 14/611 | 200/21355 | 0.0019 | 0.01716 | 0.01273 | 64766/51552/7046/1119/8073/701/10892/9918/7813/51460/30844/5888/1871/9459 | 14 |
| GSE17721_0.5H_VS_24H_GARDIQUIMOD_BMDC_DN | GSE17721_0.5H_VS_24H_GARDIQUIMOD_BMDC_DN | 14/611 | 200/21355 | 0.0019 | 0.01716 | 0.01273 | 56261/23649/1105/51072/81669/4733/202052/7409/22908/51460/5515/1315/23365/8726 | 14 |
| GSE17721_CTRL_VS_PAM3CSK4_0.5H_BMDC_UP | GSE17721_CTRL_VS_PAM3CSK4_0.5H_BMDC_UP | 14/611 | 200/21355 | 0.0019 | 0.01716 | 0.01273 | 4670/1105/150864/84830/51271/10055/22908/5588/54815/5090/23198/283131/8031/54617 | 14 |
| GSE17721_CTRL_VS_POLYIC_24H_BMDC_DN | GSE17721_CTRL_VS_POLYIC_24H_BMDC_DN | 14/611 | 200/21355 | 0.0019 | 0.01716 | 0.01273 | 4204/1105/9567/257160/342945/124245/11165/9611/2885/817/5165/29761/23013/2643 | 14 |
| GSE17721_LPS_VS_POLYIC_8H_BMDC_UP | GSE17721_LPS_VS_POLYIC_8H_BMDC_UP | 14/611 | 200/21355 | 0.0019 | 0.01716 | 0.01273 | 1073/3683/117583/2909/5533/3157/4363/5880/1289/26133/117584/7982/9402/9694 | 14 |
| GSE17721_PAM3CSK4_VS_GADIQUIMOD_24H_BMDC_DN | GSE17721_PAM3CSK4_VS_GADIQUIMOD_24H_BMDC_DN | 14/611 | 200/21355 | 0.0019 | 0.01716 | 0.01273 | 23304/51072/9782/3106/2926/80196/10048/5977/10096/7982/128866/23607/8428/10963 | 14 |
| GSE17721_POLYIC_VS_CPG_6H_BMDC_UP | GSE17721_POLYIC_VS_CPG_6H_BMDC_UP | 14/611 | 200/21355 | 0.0019 | 0.01716 | 0.01273 | 10672/25777/5257/79828/6249/124245/22848/23515/2035/23013/51176/23607/2175/8428 | 14 |
| GSE18791_CTRL_VS_NEWCASTLE_VIRUS_DC_10H_DN | GSE18791_CTRL_VS_NEWCASTLE_VIRUS_DC_10H_DN | 14/611 | 200/21355 | 0.0019 | 0.01716 | 0.01273 | 10672/1540/56913/6774/23347/84166/5533/3660/135112/65979/2643/83990/388685/84441 | 14 |
| GSE18893_CTRL_VS_TNF_TREATED_TCONV_24H_DN | GSE18893_CTRL_VS_TNF_TREATED_TCONV_24H_DN | 14/611 | 200/21355 | 0.0019 | 0.01716 | 0.01273 | 23085/10390/23130/58513/10006/727957/50807/55870/6337/10425/549/324/79058/26043 | 14 |
| GSE19888_CTRL_VS_T_CELL_MEMBRANES_ACT_MAST_CELL_UP | GSE19888_CTRL_VS_T_CELL_MEMBRANES_ACT_MAST_CELL_UP | 14/611 | 200/21355 | 0.0019 | 0.01716 | 0.01273 | 9202/64421/64848/84301/10006/3480/6780/10048/10055/8904/3183/976/64783/9377 | 14 |
| GSE21063_3H_VS_16H_ANTI_IGM_STIM_BCELL_UP | GSE21063_3H_VS_16H_ANTI_IGM_STIM_BCELL_UP | 14/611 | 200/21355 | 0.0019 | 0.01716 | 0.01273 | 23131/55619/4297/51108/4363/54899/50852/219988/23607/57198/54617/4134/9949/55758 | 14 |

| | | | | | | | | |
|---|---|---|---|---|---|---|---|---|
| GSE21360_PRIMARY_VS_TERTIARY_MEMORY_CD8_TCELL_UP | GSE21360_PRIMARY_VS_TERTIARY_MEMORY_CD8_TCELL_UP | 14/611 | 200/21355 | 0.0019 | 0.01716 | 0.01273 | 4775/6938/10464/5339/9797/54838/4363/23303/5295/1432/5880/552900/23163/55818 | 14 |
| GSE21670_TGFB_VS_TGFB_AND_IL6_TREATED_CD4_TCELL_DN | GSE21670_TGFB_VS_TGFB_AND_IL6_TREATED_CD4_TCELL_DN | 14/611 | 200/21355 | 0.0019 | 0.01716 | 0.01273 | 23112/3683/6780/124245/4627/5690/22978/10956/29761/54870/50852/2035/51176/9953 | 14 |
| GSE21670_UNTREATED_VS_TGFB_IL6_TREATED_STAT3_KO_CD4_TCELL_DN | GSE21670_UNTREATED_VS_TGFB_IL6_TREATED_STAT3_KO_CD4_TCELL_DN | 14/611 | 200/21355 | 0.0019 | 0.01716 | 0.01273 | 7534/585/6774/3683/7326/29980/22848/120/11064/4700/10096/9219/84433/5906 | 14 |
| GSE22313_HEALTHY_VS_SLE_MOUSE_CD4_TCELL_DN | GSE22313_HEALTHY_VS_SLE_MOUSE_CD4_TCELL_DN | 14/611 | 200/21355 | 0.0019 | 0.01716 | 0.01273 | 23201/65059/944/23150/2909/11165/3077/79801/155038/120425/8031/84766/9459/27244 | 14 |
| GSE22601_DOUBLE_POSITIVE_VS_CD4_SINGLE_POSITIVE_THYMOCYTE_DN | GSE22601_DOUBLE_POSITIVE_VS_CD4_SINGLE_POSITIVE_THYMOCYTE_DN | 14/611 | 200/21355 | 0.0019 | 0.01716 | 0.01273 | 23047/672/29028/10111/84316/10892/165918/1060/6767/5933/9918/5094/10198/10963 | 14 |
| GSE22886_TH1_VS_TH2_48H_ACT_DN | GSE22886_TH1_VS_TH2_48H_ACT_DN | 14/611 | 200/21355 | 0.0019 | 0.01716 | 0.01273 | 23064/4245/2081/5257/5295/10163/5588/22978/23527/23248/5578/55023/51719/4012 | 14 |
| GSE23502_BM_VS_COLON_TUMOR_HDC_KO_MYELOID_DERIVED_SUPPRESSOR_CELL_DN | GSE23502_BM_VS_COLON_TUMOR_HDC_KO_MYELOID_DERIVED_SUPPRESSOR_CELL_DN | 14/611 | 200/21355 | 0.0019 | 0.01716 | 0.01273 | 4255/146712/23064/117583/155435/5295/1432/7040/9807/135112/10788/23013/104/84433 | 14 |
| GSE24726_WT_VS_E2_2_KO_PDC_UP | GSE24726_WT_VS_E2_2_KO_PDC_UP | 14/611 | 200/21355 | 0.0019 | 0.01716 | 0.01273 | 132789/7456/8073/4733/3655/79956/3660/5933/79801/64783/9459/4354/25938/7267 | 14 |
| GSE24814_STAT5_KO_VS_WT_PRE_BCELL_DN | GSE24814_STAT5_KO_VS_WT_PRE_BCELL_DN | 14/611 | 200/21355 | 0.0019 | 0.01716 | 0.01273 | 9202/4670/546/23150/4928/1203/9491/5090/6599/1739/9020/4354/124565/473 | 14 |
| GSE24972_WT_VS_IRF8_KO_MARGINAL_ZONE_SPLEEN_BCELL_UP | GSE24972_WT_VS_IRF8_KO_MARGINAL_ZONE_SPLEEN_BCELL_UP | 14/611 | 200/21355 | 0.0019 | 0.01716 | 0.01273 | 85464/23112/10390/7414/1265/9889/79602/55619/976/29761/23365/284058/64750/60685 | 14 |
| GSE25085_FETAL_LIVER_VS_ADULT_BM_SP4_THYMIC_IMPLANT_DN | GSE25085_FETAL_LIVER_VS_ADULT_BM_SP4_THYMIC_IMPLANT_DN | 14/611 | 200/21355 | 0.0019 | 0.01716 | 0.01273 | 11329/29028/79718/55010/55421/9960/10479/9918/54878/5090/10352/64333/7443/10198 | 14 |
| GSE25123_ROSIGLITAZONE_VS_IL4_AND_ROSIGLITAZONE_STIM_PPARG_KO_MACROPHAGE_DAY10_DN | GSE25123_ROSIGLITAZONE_VS_IL4_AND_ROSIGLITAZONE_STIM_PPARG_KO_MACROPHAGE_DAY10_DN | 14/611 | 200/21355 | 0.0019 | 0.01716 | 0.01273 | 65125/1265/79602/6938/84301/59269/128710/5880/1778/25862/9274/7109/55758/5906 | 14 |
| GSE2585_AIRE_KO_VS_WT_CD80_HIGH_MTEC_UP | GSE2585_AIRE_KO_VS_WT_CD80_HIGH_MTEC_UP | 14/611 | 200/21355 | 0.0019 | 0.01716 | 0.01273 | 51552/9202/9887/23167/4820/54834/79718/10111/10497/55010/8500/4850/22978/7267 | 14 |
| GSE26669_CTRL_VS_COSTIM_BLOCK_MLR_CD4_TCELL_DN | GSE26669_CTRL_VS_COSTIM_BLOCK_MLR_CD4_TCELL_DN | 14/611 | 200/21355 | 0.0019 | 0.01716 | 0.01273 | 146057/23064/89970/25777/83478/5925/120/23048/10905/117584/51176/115/4012/23032 | 14 |
| GSE26669_CTRL_VS_COSTIM_BLOCK_MLR_CD8_TCELL_DN | GSE26669_CTRL_VS_COSTIM_BLOCK_MLR_CD8_TCELL_DN | 14/611 | 200/21355 | 0.0019 | 0.01716 | 0.01273 | 1387/51552/9736/84301/25777/83478/55972/1432/10425/5305/54870/6645/9953/51720 | 14 |
| GSE27670_CTRL_VS_LMP1_TRANSDUCED_GC_BCELL_DN | GSE27670_CTRL_VS_LMP1_TRANSDUCED_GC_BCELL_DN | 14/611 | 200/21355 | 0.0019 | 0.01716 | 0.01273 | 54934/3069/8729/55852/54899/5880/4026/50650/51/5900/55818/7109/80728/4354 | 14 |
| GSE2770_UNTREATED_VS_IL12_TREATED_ACT_CD4_TCELL_48H_UP | GSE2770_UNTREATED_VS_IL12_TREATED_ACT_CD4_TCELL_48H_UP | 14/611 | 200/21355 | 0.0019 | 0.01716 | 0.01273 | 23112/64766/3936/4775/701/84316/342945/79956/976/8726/55500/114836/25938/5906 | 14 |
| GSE27786_BCELL_VS_NEUTROPHIL_UP | GSE27786_BCELL_VS_NEUTROPHIL_UP | 14/611 | 200/21355 | 0.0019 | 0.01716 | 0.01273 | 132789/9135/5496/4775/9736/9967/8500/10055/6767/324/135112/55683/64783/10963 | 14 |
| GSE27786_LSK_VS_CD4_TCELL_DN | GSE27786_LSK_VS_CD4_TCELL_DN | 14/611 | 200/21355 | 0.0019 | 0.01716 | 0.01273 | 55187/23064/55252/10006/84166/5925/3710/378938/29123/22866/4354/11184/23287/5906 | 14 |
| GSE29164_UNTREATED_VS_CD8_TCELL_AND_IL12_TREATED_MELANOMA_DAY3_DN | GSE29164_UNTREATED_VS_CD8_TCELL_AND_IL12_TREATED_MELANOMA_DAY3_DN | 14/611 | 200/21355 | 0.0019 | 0.01716 | 0.01273 | 23112/1540/89970/23347/8500/4967/5933/157680/3454/259230/56852/6873/56942/5906 | 14 |
| GSE29618_PRE_VS_DAY7_POST_TIV_FLU_VACCINE_PDC_DN | GSE29618_PRE_VS_DAY7_POST_TIV_FLU_VACCINE_PDC_DN | 14/611 | 200/21355 | 0.0019 | 0.01716 | 0.01273 | 9887/23077/546/23476/23059/4967/4026/23048/54815/10767/7040/283131/9949/9694 | 14 |
| GSE29949_MICROGLIA_BRAIN_VS_MONOCYTE_BONE_MARROW_DN | GSE29949_MICROGLIA_BRAIN_VS_MONOCYTE_BONE_MARROW_DN | 14/611 | 200/21355 | 0.0019 | 0.01716 | 0.01273 | 1387/4670/9736/55187/23047/9976/1786/1060/5925/976/10129/23355/23163/9694 | 14 |
| GSE30083_SP3_VS_SP4_THYMOCYTE_DN | GSE30083_SP3_VS_SP4_THYMOCYTE_DN | 14/611 | 200/21355 | 0.0019 | 0.01716 | 0.01273 | 727/3106/727957/7421/55852/3560/23443/817/976/23248/6645/9020/9953/81671 | 14 |
| GSE31082_DN_VS_DP_THYMOCYTE_DN | GSE31082_DN_VS_DP_THYMOCYTE_DN | 14/611 | 200/21355 | 0.0019 | 0.01716 | 0.01273 | 10672/23112/23181/64421/1105/6938/57690/9728/126298/57649/83891/84937/57337/80331 | 14 |

| | | | | | | | | |
|---|---|---|---|---|---|---|---|---|
| GSE32423_CTRL_VS_IL4_MEMORY_CD8_TCELL_UP | GSE32423_CTRL_VS_IL4_MEMORY_CD8_TCELL_UP | 14/611 | 200/21355 | 0.0019 | 0.01716 | 0.01273 | 65125/9698/987/3683/89970/55619/7322/283209/259230/10788/114792/25938/2590/9953 | 14 |
| GSE32986_UNSTIM_VS_CURDLAN_LOWDOSE_STIM_DC_DN | GSE32986_UNSTIM_VS_CURDLAN_LOWDOSE_STIM_DC_DN | 14/611 | 200/21355 | 0.0019 | 0.01716 | 0.01273 | 1119/3683/823/641/965/2909/23406/255231/23102/6645/23365/1871/8428/9844 | 14 |
| GSE33162_HDAC3_KO_VS_HDAC3_KO_MACROPHAGE_DN | GSE33162_HDAC3_KO_VS_HDAC3_KO_MACROPHAGE_DN | 14/611 | 200/21355 | 0.0019 | 0.01716 | 0.01273 | 5820/860/23139/221477/9960/7170/54870/50852/11163/5094/7531/1399/51719/81671 | 14 |
| GSE339_CD4POS_VS_CD8POS_DC_UP | GSE339_CD4POS_VS_CD8POS_DC_UP | 14/611 | 200/21355 | 0.0019 | 0.01716 | 0.01273 | 11329/54934/1119/1729/7187/4215/1432/5880/50650/57649/64333/1739/55500/27244 | 14 |
| GSE339_CD8POS_VS_CD4CD8DN_DC_IN_CULTURE_DN | GSE339_CD8POS_VS_CD4CD8DN_DC_IN_CULTURE_DN | 14/611 | 200/21355 | 0.0019 | 0.01716 | 0.01273 | 65125/377/9202/54934/23369/51068/6929/55193/23019/7170/64333/1315/8031/23534 | 14 |
| GSE34156_TLR1_TLR2_LIGAND_VS_NOD2_AND_TLR1_TLR2_LIGAND_24H_TREATED_MONOCYTE_DN | GSE34156_TLR1_TLR2_LIGAND_VS_NOD2_AND_TLR1_TLR2_LIGAND_24H_TREATED_MONOCYTE_DN | 14/611 | 200/21355 | 0.0019 | 0.01716 | 0.01273 | 23201/7534/6774/1729/83478/7813/405/5305/114804/7251/25853/54521/57534/83990 | 14 |
| GSE34515_CD16_POS_MONOCYTE_VS_DC_UP | GSE34515_CD16_POS_MONOCYTE_VS_DC_UP | 14/611 | 200/21355 | 0.0019 | 0.01716 | 0.01273 | 253461/3683/59269/701/6780/55852/22908/23048/2521/1739/8726/10521/10657/79657 | 14 |
| GSE35685_CD34POS_CD38NEG_VS_CD34POS_CD10POS_BONE_MARROW_DN | GSE35685_CD34POS_CD38NEG_VS_CD34POS_CD10POS_BONE_MARROW_DN | 14/611 | 200/21355 | 0.0019 | 0.01716 | 0.01273 | 6774/10497/84316/80196/255231/4215/50650/5770/128866/55500/10906/2643/473/81671 | 14 |
| GSE360_L_DONOVANI_VS_B_MALAYI_LOW_DOSE_DC_DN | GSE360_L_DONOVANI_VS_B_MALAYI_LOW_DOSE_DC_DN | 14/611 | 200/21355 | 0.0019 | 0.01716 | 0.01273 | 11329/3064/79602/3683/1729/7409/23443/5305/5090/23384/10129/6197/5094/9459 | 14 |
| GSE369_SOCS3_KO_VS_WT_LIVER_POST_IL6_INJECTION_DN | GSE369_SOCS3_KO_VS_WT_LIVER_POST_IL6_INJECTION_DN | 14/611 | 200/21355 | 0.0019 | 0.01716 | 0.01273 | 56261/60468/9267/4775/54834/337867/1520/23139/10048/23141/5925/10163/55784/9673 | 14 |
| GSE3720_UNSTIM_VS_LPS_STIM_VD1_GAMMADELTA_TCELL_UP | GSE3720_UNSTIM_VS_LPS_STIM_VD1_GAMMADELTA_TCELL_UP | 14/611 | 200/21355 | 0.0019 | 0.01716 | 0.01273 | 11329/4775/23064/4245/727957/6794/1431/1432/1778/55193/976/10129/55683/9953 | 14 |
| GSE39152_CD103_NEG_VS_POS_MEMORY_CD8_TCELL_DN | GSE39152_CD103_NEG_VS_POS_MEMORY_CD8_TCELL_DN | 14/611 | 200/21355 | 0.0019 | 0.01716 | 0.01273 | 377/6774/55095/51072/50807/8705/4363/5977/4026/405/5770/3454/6599/2590 | 14 |
| GSE39152_SPLEEN_CD103_NEG_VS_BRAIN_CD103_POS_MEMORY_CD8_TCELL_UP | GSE39152_SPLEEN_CD103_NEG_VS_BRAIN_CD103_POS_MEMORY_CD8_TCELL_UP | 14/611 | 200/21355 | 0.0019 | 0.01716 | 0.01273 | 377/337867/50807/7409/4026/57494/871/3594/5770/51/3454/80728/5073/2590 | 14 |
| GSE3982_DC_VS_MAC_DN | GSE3982_DC_VS_MAC_DN | 14/611 | 200/21355 | 0.0019 | 0.01716 | 0.01273 | 586/7046/6774/701/10497/3655/23303/138151/9962/3077/79801/7170/4642/4012 | 14 |
| GSE39864_WT_VS_GATA3_KO_TREG_DN | GSE39864_WT_VS_GATA3_KO_TREG_DN | 14/611 | 200/21355 | 0.0019 | 0.01716 | 0.01273 | 28977/54934/8073/4733/10111/84316/2926/23476/721/79956/23019/80728/115/473 | 14 |
| GSE40274_FOXP3_VS_FOXP3_AND_PBX1_TRANSDUCED_ACTIVATED_CD4_TCELL_UP | GSE40274_FOXP3_VS_FOXP3_AND_PBX1_TRANSDUCED_ACTIVATED_CD4_TCELL_UP | 14/611 | 200/21355 | 0.0019 | 0.01716 | 0.01273 | 3936/987/91775/6249/5775/22806/10905/10788/50852/9844/9459/9694/5906/27244 | 14 |
| GSE411_UNSTIM_VS_100MIN_IL6_STIM_MACROPHAGE_DN | GSE411_UNSTIM_VS_100MIN_IL6_STIM_MACROPHAGE_DN | 14/611 | 200/21355 | 0.0019 | 0.01716 | 0.01273 | 56261/55690/585/84376/23515/10464/7187/3454/64333/117584/92181/51176/9402/83451 | 14 |
| GSE41867_DAY6_EFFECTOR_VS_DAY30_MEMORY_CD8_TCELL_LCMV_ARMSTRONG_UP | GSE41867_DAY6_EFFECTOR_VS_DAY30_MEMORY_CD8_TCELL_LCMV_ARMSTRONG_UP | 14/611 | 200/21355 | 0.0019 | 0.01716 | 0.01273 | 64766/59269/401409/823/10006/23476/22848/3660/9962/135112/8428/84433/5906/473 | 14 |
| GSE41867_NAIVE_VS_DAY30_LCMV_CLONE13_EXHAUSTED_CD8_TCELL_DN | GSE41867_NAIVE_VS_DAY30_LCMV_CLONE13_EXHAUSTED_CD8_TCELL_DN | 14/611 | 200/21355 | 0.0019 | 0.01716 | 0.01273 | 56261/23181/23064/59269/337867/22848/3710/7813/23198/30844/117584/92912/10521/2590 | 14 |
| GSE42021_CD24LO_TREG_VS_CD24LO_TCONV_THYMUS_UP | GSE42021_CD24LO_TREG_VS_CD24LO_TCONV_THYMUS_UP | 14/611 | 200/21355 | 0.0019 | 0.01716 | 0.01273 | 65125/64324/23071/55870/26207/7409/1432/1778/11163/128866/1822/7109/23534/2590 | 14 |
| GSE42088_UNINF_VS_LEISHMANIA_INF_DC_24H_UP | GSE42088_UNINF_VS_LEISHMANIA_INF_DC_24H_UP | 14/611 | 200/21355 | 0.0019 | 0.01716 | 0.01273 | 23112/23130/1119/22861/23347/3784/1455/23406/1060/5295/5165/5900/5094/23287 | 14 |
| GSE43863_NAIVE_VS_MEMORY_LY6C_INT_CXCR5POS_CD4_TCELL_D150_LCMV_UP | GSE43863_NAIVE_VS_MEMORY_LY6C_INT_CXCR5POS_CD4_TCELL_D150_LCMV_UP | 14/611 | 200/21355 | 0.0019 | 0.01716 | 0.01273 | 57690/22992/51696/5339/5925/10163/3660/4627/120/7813/23102/259230/9020/27244 | 14 |
| GSE43955_TH0_VS_TGFB_IL6_TH17_ACT_CD4_TCELL_42H_UP | GSE43955_TH0_VS_TGFB_IL6_TH17_ACT_CD4_TCELL_42H_UP | 14/611 | 200/21355 | 0.0019 | 0.01716 | 0.01273 | 2869/26065/7326/25777/84316/1432/138151/7813/9274/1822/80728/6462/10906/7267 | 14 |
| GSE43955_TH0_VS_TGFB_IL6_TH17_ACT_CD4_TCELL_60H_UP | GSE43955_TH0_VS_TGFB_IL6_TH17_ACT_CD4_TCELL_60H_UP | 14/611 | 200/21355 | 0.0019 | 0.01716 | 0.01273 | 23047/59269/2926/4363/10163/5933/976/10767/6197/7251/8428/5073/11184/6641 | 14 |
| GSE46143_CTRL_VS_LMP2A_TRANSDUCED_CD10_POS_GC_BCELL_DN | GSE46143_CTRL_VS_LMP2A_TRANSDUCED_CD10_POS_GC_BCELL_DN | 14/611 | 200/21355 | 0.0019 | 0.01716 | 0.01273 | 2869/3936/4245/2909/2926/55870/26207/5880/8289/50650/8428/10330/473/7267 | 14 |

| | | | | | | | | |
|---|---|---|---|---|---|---|---|---|
| GSE46606_UNSTIM_VS_CD40L_IL2_IL5_1DAY_STIMULATED_IRF4MID_SORTED_BCELL_UP | GSE46606_UNSTIM_VS_CD40L_IL2_IL5_1DAY_STIMULATED_IRF4MID_SORTED_BCELL_UP | 14/611 | 200/21355 | 0.0019 | 0.01716 | 0.01273 | 9887/5048/55666/8729/4215/55236/22978/29761/23355/64783/10906/10198/84324/84636 | 14 |
| GSE5542_UNTREATED_VS_IFNA_TREATED_EPITHELIAL_CELLS_24H_UP | GSE5542_UNTREATED_VS_IFNA_TREATED_EPITHELIAL_CELLS_24H_UP | 14/611 | 200/21355 | 0.0019 | 0.01716 | 0.01273 | 8844/6777/377/4763/2909/4967/4627/3710/23048/57649/135112/23365/115/493 | 14 |
| GSE5589_LPS_AND_IL10_VS_LPS_AND_IL6_STIM_MACROPHAGE_45MIN_UP | GSE5589_LPS_AND_IL10_VS_LPS_AND_IL6_STIM_MACROPHAGE_45MIN_UP | 14/611 | 200/21355 | 0.0019 | 0.01716 | 0.01273 | 4763/23130/23064/55095/6938/23633/55252/9728/9851/84961/4363/126298/5305/7109 | 14 |
| GSE5679_RARA_AGONIST_AM580_VS_AM580_AND_ROSIGLITAZONE_TREATED_DC_DN | GSE5679_RARA_AGONIST_AM580_VS_AM580_AND_ROSIGLITAZONE_TREATED_DC_DN | 14/611 | 200/21355 | 0.0019 | 0.01716 | 0.01273 | 11329/23370/84196/55619/3655/26207/9960/5295/976/3454/64333/259230/51735/22834 | 14 |
| GSE6674_CPG_VS_PL2_3_STIM_BCELL_DN | GSE6674_CPG_VS_PL2_3_STIM_BCELL_DN | 14/611 | 200/21355 | 0.0019 | 0.01716 | 0.01273 | 65125/79954/3069/2081/55619/26528/23476/55852/55421/4363/3560/2643/55784/7403 | 14 |
| GSE6681_DELETED_FOXP3_VS_WT_TREG_DN | GSE6681_DELETED_FOXP3_VS_WT_TREG_DN | 14/611 | 200/21355 | 0.0019 | 0.01716 | 0.01273 | 1119/22887/11320/57690/23369/51807/54838/10615/549/7982/128866/63977/2175/23524 | 14 |
| GSE7460_WT_VS_FOXP3_HET_ACT_WITH_TGFB_TCONV_UP | GSE7460_WT_VS_FOXP3_HET_ACT_WITH_TGFB_TCONV_UP | 14/611 | 200/21355 | 0.0019 | 0.01716 | 0.01273 | 150864/81669/155435/9967/55421/26207/50650/324/976/148867/1739/50852/51176/9953 | 14 |
| GSE7852_LN_VS_FAT_TREG_DN | GSE7852_LN_VS_FAT_TREG_DN | 14/611 | 200/21355 | 0.0019 | 0.01716 | 0.01273 | 2869/79602/8826/117583/1729/5339/9693/3594/5090/5900/23355/2643/54842/2590 | 14 |
| GSE7852_LN_VS_FAT_TREG_UP | GSE7852_LN_VS_FAT_TREG_UP | 14/611 | 200/21355 | 0.0019 | 0.01716 | 0.01273 | 6777/171023/11329/56913/1265/4733/50650/23102/817/5305/22978/55023/4354/6654 | 14 |
| GSE8835_CD4_VS_CD8_TCELL_UP | GSE8835_CD4_VS_CD8_TCELL_UP | 14/611 | 200/21355 | 0.0019 | 0.01716 | 0.01273 | 23304/9698/55252/3480/9967/23139/23293/23059/9611/23019/23527/22834/9844/31 | 14 |
| GSE9037_CTRL_VS_LPS_4H_STIM_BMDM_DN | GSE9037_CTRL_VS_LPS_4H_STIM_BMDM_DN | 14/611 | 200/21355 | 0.0019 | 0.01716 | 0.01273 | 10390/23130/3683/4297/8500/6780/5533/9611/51460/29123/29761/6891/7251/7982 | 14 |
| GSE9037_WT_VS_IRAK4_KO_LPS_4H_STIM_BMDM_UP | GSE9037_WT_VS_IRAK4_KO_LPS_4H_STIM_BMDM_UP | 14/611 | 200/21355 | 0.0019 | 0.01716 | 0.01273 | 27340/59269/25777/5533/255231/57649/29123/2521/10788/56852/25853/23524/4791/26043 | 14 |
| KAECH_NAIVE_VS_DAY15_EFF_CD8_TCELL_UP | KAECH_NAIVE_VS_DAY15_EFF_CD8_TCELL_UP | 14/611 | 200/21355 | 0.0019 | 0.01716 | 0.01273 | 51108/3707/9960/1606/5305/54870/84937/51176/10198/4354/23450/2590/27244/7267 | 14 |
| QI_PBMC_ZOSTAVAX_AGE_50_75YO_CORRELATED_WITH_EXPANSION_OF_VZV_SPECIFIC_T_CELLS_TO_PEAK_AT_1DY_POSITIVE | QI_PBMC_ZOSTAVAX_AGE_50_75YO_CORRELATED_WITH_EXPANSION_OF_VZV_SPECIFIC_T_CELLS_TO_PEAK_AT_1DY_POSITIVE | 16/611 | 248/21355 | 0.0022 | 0.01952 | 0.01448 | 1105/6774/3106/155435/89846/860/1520/51271/57410/4627/23048/23527/9020/57198/51719/10800 | 16 |
| GSE35825_UNTREATED_VS_IFNG_STIM_MACROPHAGE_DN | GSE35825_UNTREATED_VS_IFNG_STIM_MACROPHAGE_DN | 13/611 | 181/21355 | 0.0022 | 0.01952 | 0.01448 | 64766/23064/546/8073/79718/10048/23048/5305/157680/23198/114804/4354/27244 | 13 |
| KANNAN_BLOOD_2012_2013_TIV_AGE_65PLS_REVACCINATED_IN_6_9_MO_VS_REVACCINATED_IN_12_13_MOS_DN | KANNAN_BLOOD_2012_2013_TIV_AGE_65PLS_REVACCINATED_IN_6_9_MO_VS_REVACCINATED_IN_12_13_MOS_DN | 20/611 | 343/21355 | 0.0022 | 0.01968 | 0.0146 | 65125/23181/6774/22861/84196/79230/55666/158358/23774/84193/10163/1432/405/25862/64333/5888/10956/197135/473/51720 | 20 |
| GSE21033_1H_VS_24H_POLYIC_STIM_DC_UP | GSE21033_1H_VS_24H_POLYIC_STIM_DC_UP | 12/611 | 161/21355 | 0.0024 | 0.02087 | 0.01549 | 60468/64421/6938/59269/23476/3655/23102/5900/114804/84181/23607/51719 | 12 |
| GSE13173_UNTREATED_VS_IL12_TREATED_ACT_CD8_TCELL_DN | GSE13173_UNTREATED_VS_IL12_TREATED_ACT_CD8_TCELL_DN | 11/611 | 141/21355 | 0.0025 | 0.02188 | 0.01624 | 10672/5820/10111/1060/9520/92912/65979/114836/23534/54842/31 | 11 |
| GARCIA_PINERES_PBMC_HPV_16_L1_VLP_AGE_18_25YO_2MO_UP | GARCIA_PINERES_PBMC_HPV_16_L1_VLP_AGE_18_25YO_2MO_UP | 19/611 | 324/21355 | 0.0027 | 0.02341 | 0.01737 | 727/59269/672/10111/9967/7421/10036/51068/55972/9693/3660/4277/5305/5690/10403/6891/2175/2643/4012 | 19 |
| GSE40274_FOXP3_VS_FOXP3_AND_HELIOS_TRANSDUCED_ACTIVATED_CD4_TCELL_DN | GSE40274_FOXP3_VS_FOXP3_AND_HELIOS_TRANSDUCED_ACTIVATED_CD4_TCELL_DN | 11/611 | 143/21355 | 0.0028 | 0.02433 | 0.01806 | 11052/7799/146057/1105/944/10111/378805/65117/84186/10198/114836 | 11 |
| NAKAYA_PBMC_FLUARIX_FLUVIRIN_AGE_18_50YO_CORRELATED_WITH_HAI_28DY_RESPONSE_AT_3DY_POSITIVE | NAKAYA_PBMC_FLUARIX_FLUVIRIN_AGE_18_50YO_CORRELATED_WITH_HAI_28DY_RESPONSE_AT_3DY_POSITIVE | 23/611 | 426/21355 | 0.0029 | 0.02538 | 0.01883 | 377/23304/79602/8826/3069/965/7421/54838/4967/7409/3660/54165/3454/64333/10956/10096/7531/92181/120425/124565/2590/80025/81671 | 23 |
| GSE37605_C57BL6_VS_NOD_FOXP3_FUSION_GFP_TCONV_UP | GSE37605_C57BL6_VS_NOD_FOXP3_FUSION_GFP_TCONV_UP | 12/611 | 167/21355 | 0.0032 | 0.02795 | 0.02074 | 4763/4255/586/59269/7421/83478/50807/23141/6613/51460/5578/84636 | 12 |
| GSE15624_3H_VS_6H_HALOFUGINONE_TREATED_CD4_TCELL_DN | GSE15624_3H_VS_6H_HALOFUGINONE_TREATED_CD4_TCELL_DN | 11/611 | 147/21355 | 0.0034 | 0.02994 | 0.02222 | 5496/3936/150864/6774/7173/3183/9962/3560/51/1911/7109 | 11 |
| NAKAYA_MONOCYTE_FLUMIST_AGE_18_50YO_7DY_DN | NAKAYA_MONOCYTE_FLUMIST_AGE_18_50YO_7DY_DN | 16/611 | 260/21355 | 0.0035 | 0.03052 | 0.02265 | 10672/11329/1540/1105/22861/860/1729/51271/23499/5880/7040/7170/1315/9020/64750/9949 | 16 |

| | | | | | | | | |
|---|---|---|---|---|---|---|---|---|
| GSE40274_CTRL_VS_FOXP3_AND_EOS_TRANSDUCED_ACTIVATED_CD4_TCELL_UP | GSE40274_CTRL_VS_FOXP3_AND_EOS_TRANSDUCED_ACTIVATED_CD4_TCELL_UP | 12/611 | 169/21355 | 0.0035 | 0.03052 | 0.02265 | 23370/860/5775/26207/1432/1606/5933/817/976/64333/9402/55784 | 12 |
| GSE4590_SMALL_VS_VPREB_POS_LARGE_PRE_BCELL_UP | GSE4590_SMALL_VS_VPREB_POS_LARGE_PRE_BCELL_UP | 12/611 | 169/21355 | 0.0035 | 0.03052 | 0.02265 | 4763/26574/3480/55619/79828/84961/26207/9466/22978/7798/51317/84636 | 12 |
| SCHERER_PBMC_APSV_WETVAX_AGE_18_32YO_50_TO_60DY_UP | SCHERER_PBMC_APSV_WETVAX_AGE_18_32YO_50_TO_60DY_UP | 12/611 | 169/21355 | 0.0035 | 0.03052 | 0.02265 | 23130/1105/9976/84196/23141/5295/65117/57649/26036/10403/7403/27244 | 12 |
| GSE26928_NAIVE_VS_CENT_MEMORY_CD4_TCELL_UP | GSE26928_NAIVE_VS_CENT_MEMORY_CD4_TCELL_UP | 13/611 | 192/21355 | 0.0037 | 0.03177 | 0.02358 | 64848/337867/727957/7182/3707/9797/4795/196074/259283/114804/22866/114792/4354 | 13 |
| GSE1460_NAIVE_CD4_TCELL_ADULT_BLOOD_VS_THYMIC_STROMAL_CELL_UP | GSE1460_NAIVE_CD4_TCELL_ADULT_BLOOD_VS_THYMIC_STROMAL_CELL_UP | 13/611 | 193/21355 | 0.0038 | 0.03303 | 0.02451 | 60468/23370/64421/3064/9976/1729/22848/1432/54878/50852/55818/51719/2643 | 13 |
| GSE26495_NAIVE_VS_PD1HIGH_CD8_TCELL_UP | GSE26495_NAIVE_VS_PD1HIGH_CD8_TCELL_UP | 13/611 | 193/21355 | 0.0038 | 0.03303 | 0.02451 | 150864/3480/1203/3655/6929/259283/114804/9712/5578/51176/22866/79632/84441 | 13 |
| GSE6259_33D1_POS_VS_DEC205_POS_SPLENIC_DC_UP | GSE6259_33D1_POS_VS_DEC205_POS_SPLENIC_DC_UP | 13/611 | 193/21355 | 0.0038 | 0.03303 | 0.02451 | 23112/150864/59269/155435/51696/4928/4297/84961/5588/23060/11064/25853/473 | 13 |
| GSE3565_CTRL_VS_LPS_INJECTED_DUSP1_KO_SPLENOCYTES_DN | GSE3565_CTRL_VS_LPS_INJECTED_DUSP1_KO_SPLENOCYTES_DN | 12/611 | 171/21355 | 0.0039 | 0.03321 | 0.02465 | 9873/53944/22955/1606/120/64333/51176/51735/23607/4354/27244/7267 | 12 |
| GSE40274_CTRL_VS_XBP1_TRANSDUCED_ACTIVATED_CD4_TCELL_UP | GSE40274_CTRL_VS_XBP1_TRANSDUCED_ACTIVATED_CD4_TCELL_UP | 12/611 | 171/21355 | 0.0039 | 0.03321 | 0.02465 | 10006/5533/5339/26207/54899/22806/3560/5305/29761/57337/51317/10800 | 12 |
| GSE21670_UNTREATED_VS_TGFB_IL6_TREATED_CD4_TCELL_UP | GSE21670_UNTREATED_VS_TGFB_IL6_TREATED_CD4_TCELL_UP | 13/611 | 194/21355 | 0.004 | 0.03355 | 0.02489 | 60468/64421/727/22828/9736/6938/29028/701/4297/4026/5090/6645/10198 | 13 |
| GSE22886_NAIVE_CD8_TCELL_VS_MEMORY_TCELL_UP | GSE22886_NAIVE_CD8_TCELL_VS_MEMORY_TCELL_UP | 13/611 | 194/21355 | 0.004 | 0.03355 | 0.02489 | 64421/6526/84316/9851/6929/54878/9807/7644/8672/23604/5205/10198/79157 | 13 |
| GSE14415_ACT_TCONV_VS_ACT_NATURAL_TREG_UP | GSE14415_ACT_TCONV_VS_ACT_NATURAL_TREG_UP | 12/611 | 172/21355 | 0.004 | 0.03355 | 0.02489 | 7046/944/57690/23347/51108/3655/146691/50852/51176/51735/23607/27244 | 12 |
| GSE1460_DP_THYMOCYTE_VS_THYMIC_STROMAL_CELL_UP | GSE1460_DP_THYMOCYTE_VS_THYMIC_STROMAL_CELL_UP | 13/611 | 195/21355 | 0.0042 | 0.03355 | 0.02489 | 6777/4255/81669/3784/701/1060/8631/3660/51/5165/22834/9129/9673 | 13 |
| GSE22611_UNSTIM_VS_6H_MDP_STIM_MUTANT_NOD2_TRANSDUCED_HEK293T_CELL_DN | GSE22611_UNSTIM_VS_6H_MDP_STIM_MUTANT_NOD2_TRANSDUCED_HEK293T_CELL_DN | 13/611 | 195/21355 | 0.0042 | 0.03355 | 0.02489 | 146712/7173/7421/753/138151/23102/54815/5305/976/23604/9263/23287/23032 | 13 |
| GSE46606_UNSTIM_VS_CD40L_IL2_IL5_1DAY_STIMULATED_IRF4_KO_BCELL_DN | GSE46606_UNSTIM_VS_CD40L_IL2_IL5_1DAY_STIMULATED_IRF4_KO_BCELL_DN | 13/611 | 195/21355 | 0.0042 | 0.03355 | 0.02489 | 65125/8844/9267/337867/55605/23774/165918/26207/8631/29123/5770/51735/7798 | 13 |
| OCONNOR_PBMC_MENVEO_ACWYVAX_AGE_30_70YO_7DY_AFTER_SECOND_DOSE_VS_7DY_AFTER_FIRST_DOSE_DN | OCONNOR_PBMC_MENVEO_ACWYVAX_AGE_30_70YO_7DY_AFTER_SECOND_DOSE_VS_7DY_AFTER_FIRST_DOSE_DN | 15/611 | 242/21355 | 0.0043 | 0.03355 | 0.02489 | 64766/90592/9698/22828/7456/150864/7326/84196/23515/22806/1315/92181/6873/54842/10657 | 15 |
| GSE14000_4H_VS_16H_LPS_DC_TRANSLATED_RNA_UP | GSE14000_4H_VS_16H_LPS_DC_TRANSLATED_RNA_UP | 13/611 | 196/21355 | 0.0043 | 0.03355 | 0.02489 | 4255/3064/22861/22848/91942/83478/6894/50807/7187/79956/135112/11184/7267 | 13 |
| GSE20727_CTRL_VS_H2O2_TREATED_DC_DN | GSE20727_CTRL_VS_H2O2_TREATED_DC_DN | 13/611 | 196/21355 | 0.0043 | 0.03355 | 0.02489 | 85464/9135/25777/23077/337867/23150/11165/342892/120/51586/5165/23384/79613 | 13 |
| GSE22886_UNSTIM_VS_IL15_STIM_NKCELL_UP | GSE22886_UNSTIM_VS_IL15_STIM_NKCELL_UP | 13/611 | 196/21355 | 0.0043 | 0.03355 | 0.02489 | 23130/4820/81669/22861/79663/6894/4215/1289/7769/23355/55818/79157/493 | 13 |
| GSE2770_IL12_AND_TGFB_ACT_VS_ACT_CD4_TCELL_2H_DN | GSE2770_IL12_AND_TGFB_ACT_VS_ACT_CD4_TCELL_2H_DN | 13/611 | 196/21355 | 0.0043 | 0.03355 | 0.02489 | 26065/256236/9797/11165/9960/5165/2035/22834/9844/11184/473/6641/7267 | 13 |
| GSE19401_UNSTIM_VS_PAM2CSK4_STIM_FOLLICULAR_DC_UP | GSE19401_UNSTIM_VS_PAM2CSK4_STIM_FOLLICULAR_DC_UP | 13/611 | 197/21355 | 0.0045 | 0.03355 | 0.02489 | 4763/7414/1105/9736/23071/3480/23476/7182/7187/4215/10403/9129/6654 | 13 |
| GSE21546_WT_VS_SAP1A_KO_DP_THYMOCYTES_UP | GSE21546_WT_VS_SAP1A_KO_DP_THYMOCYTES_UP | 13/611 | 197/21355 | 0.0045 | 0.03355 | 0.02489 | 1540/3106/84166/7322/53944/3660/83891/30844/6891/219988/9129/4012/493 | 13 |
| GSE22886_NAIVE_CD8_TCELL_VS_NKCELL_DN | GSE22886_NAIVE_CD8_TCELL_VS_NKCELL_DN | 13/611 | 197/21355 | 0.0045 | 0.03355 | 0.02489 | 51552/23064/23139/23101/3560/92170/23198/23365/55023/55784/493/51317/54842 | 13 |
| GSE2770_TGFB_AND_IL4_VS_IL12_TREATED_ACT_CD4_TCELL_2H_DN | GSE2770_TGFB_AND_IL4_VS_IL12_TREATED_ACT_CD4_TCELL_2H_DN | 13/611 | 197/21355 | 0.0045 | 0.03355 | 0.02489 | 65125/6774/79230/23303/7699/126298/10425/3560/23198/10800/9673/163486/5906 | 13 |
| GSE29618_PRE_VS_DAY7_POST_LAIV_FLU_VACCINE_MONOCYTE_UP | GSE29618_PRE_VS_DAY7_POST_LAIV_FLU_VACCINE_MONOCYTE_UP | 13/611 | 197/21355 | 0.0045 | 0.03355 | 0.02489 | 377/56005/1540/1105/10497/51271/23499/6613/55236/1315/283131/64750/54842 | 13 |
| GSE30971_WBP7_HET_VS_KO_MACROPHAGE_4H_LPS_STIM_DN | GSE30971_WBP7_HET_VS_KO_MACROPHAGE_4H_LPS_STIM_DN | 13/611 | 197/21355 | 0.0045 | 0.03355 | 0.02489 | 11329/727/23150/79663/1520/10144/5295/5925/5305/23355/10521/9459/4354 | 13 |
| GSE32986_GMCSF_VS_GMCSF_AND_CURDLAN_LOWDOSE_STIM_DC_DN | GSE32986_GMCSF_VS_GMCSF_AND_CURDLAN_LOWDOSE_STIM_DC_DN | 13/611 | 197/21355 | 0.0045 | 0.03355 | 0.02489 | 85464/54934/146057/3707/3655/51455/57494/7644/259283/114804/5578/4354/84441 | 13 |
| GSE37416_0H_VS_12H_F_TULARENSIS_LVS_NEUTROPHIL_UP | GSE37416_0H_VS_12H_F_TULARENSIS_LVS_NEUTROPHIL_UP | 13/611 | 197/21355 | 0.0045 | 0.03355 | 0.02489 | 7414/150864/8826/823/120/51/10403/10788/2035/51719/124446/1178/9673 | 13 |

| | | | | | | | | |
|---|---|---|---|---|---|---|---|---|
| GSE3982_EOSINOPHIL_VS_NKCELL_UP | GSE3982_EOSINOPHIL_VS_NKCELL_UP | 13/611 | 197/21355 | 0.0045 | 0.03355 | 0.02489 | 23214/23071/672/3106/3707/9712/4700/1871/283131/8031/23524/10800/6641 | 13 |
| GSE37301_PRO_BCELL_VS_GRANULOCYTE_MONOCYTE_PROGENITOR_DN | GSE37301_PRO_BCELL_VS_GRANULOCYTE_MONOCYTE_PROGENITOR_DN | 12/611 | 175/21355 | 0.0046 | 0.03355 | 0.02489 | 7414/146057/23303/2885/7813/976/3594/155038/493/124565/54842/2289 | 12 |
| GSE11864_UNTREATED_VS_CSF1_IFNG_PAM3CYS_IN_MAC_UP | GSE11864_UNTREATED_VS_CSF1_IFNG_PAM3CYS_IN_MAC_UP | 13/611 | 198/21355 | 0.0047 | 0.03355 | 0.02489 | 11329/54934/4670/9889/256435/2081/84196/2926/23406/65117/8289/23355/51720 | 13 |
| GSE13411_NAIVE_VS_SWITCHED_MEMORY_BCELL_DN | GSE13411_NAIVE_VS_SWITCHED_MEMORY_BCELL_DN | 13/611 | 198/21355 | 0.0047 | 0.03355 | 0.02489 | 5496/23167/7414/79718/5257/9491/4363/1778/5305/7644/11163/55683/7403 | 13 |
| GSE13738_TCR_VS_BYSTANDER_ACTIVATED_CD4_TCELL_DN | GSE13738_TCR_VS_BYSTANDER_ACTIVATED_CD4_TCELL_DN | 13/611 | 198/21355 | 0.0047 | 0.03355 | 0.02489 | 7046/9267/1540/1203/5775/3655/10144/57494/23365/55683/23013/83451/6654 | 13 |
| GSE1460_CD4_THYMOCYTE_VS_THYMIC_STROMAL_CELL_UP | GSE1460_CD4_THYMOCYTE_VS_THYMIC_STROMAL_CELL_UP | 13/611 | 198/21355 | 0.0047 | 0.03355 | 0.02489 | 171023/23085/26574/23130/80196/1729/9491/5339/1606/10425/405/22834/11184 | 13 |
| GSE16266_CTRL_VS_LPS_STIM_MEF_UP | GSE16266_CTRL_VS_LPS_STIM_MEF_UP | 13/611 | 198/21355 | 0.0047 | 0.03355 | 0.02489 | 10672/59269/965/378805/7421/860/23515/3707/165918/23198/117584/8031/64750 | 13 |
| GSE18791_CTRL_VS_NEWCASTLE_VIRUS_DC_6H_DN | GSE18791_CTRL_VS_NEWCASTLE_VIRUS_DC_6H_DN | 13/611 | 198/21355 | 0.0047 | 0.03355 | 0.02489 | 10672/1540/56913/23347/84166/23515/5533/135112/65979/219988/2643/83990/6654 | 13 |
| GSE18791_UNSTIM_VS_NEWCATSLE_VIRUS_DC_18H_UP | GSE18791_UNSTIM_VS_NEWCATSLE_VIRUS_DC_18H_UP | 13/611 | 198/21355 | 0.0047 | 0.03355 | 0.02489 | 3480/4733/342892/9611/138151/57410/6599/64968/4931/7982/51719/79692/9459 | 13 |
| GSE19923_WT_VS_E2A_KO_DP_THYMOCYTE_DN | GSE19923_WT_VS_E2A_KO_DP_THYMOCYTE_DN | 13/611 | 198/21355 | 0.0047 | 0.03355 | 0.02489 | 58513/11320/55010/2909/10892/1786/7699/10767/22978/26133/4012/9953/6654 | 13 |
| GSE21670_UNTREATED_VS_IL6_TREATED_CD4_TCELL_UP | GSE21670_UNTREATED_VS_IL6_TREATED_CD4_TCELL_UP | 13/611 | 198/21355 | 0.0047 | 0.03355 | 0.02489 | 9698/823/8729/23101/255231/23141/3560/9807/23019/92181/23163/4642/8428 | 13 |
| GSE22886_NAIVE_TCELL_VS_NKCELL_DN | GSE22886_NAIVE_TCELL_VS_NKCELL_DN | 13/611 | 198/21355 | 0.0047 | 0.03355 | 0.02489 | 23304/3064/23064/6526/23101/23499/50650/92170/23198/23365/64750/55784/51317 | 13 |
| GSE2770_IL12_AND_TGFB_VS_IL4_TREATED_ACT_CD4_TCELL_2H_DN | GSE2770_IL12_AND_TGFB_VS_IL4_TREATED_ACT_CD4_TCELL_2H_DN | 13/611 | 198/21355 | 0.0047 | 0.03355 | 0.02489 | 23112/23131/3064/2909/51108/54899/5295/7813/5090/23013/22834/57534/51317 | 13 |
| GSE27786_BCELL_VS_CD4_TCELL_DN | GSE27786_BCELL_VS_CD4_TCELL_DN | 13/611 | 198/21355 | 0.0047 | 0.03355 | 0.02489 | 23112/23214/3683/6526/158358/9611/84193/10479/6197/84937/80728/197135/114836 | 13 |
| GSE29949_CD8_POS_DC_SPLEEN_VS_DC_BRAIN_UP | GSE29949_CD8_POS_DC_SPLEEN_VS_DC_BRAIN_UP | 13/611 | 198/21355 | 0.0047 | 0.03355 | 0.02489 | 377/22828/987/6526/25777/23347/84830/6894/4363/5976/2521/5090/473 | 13 |
| GSE33162_UNTREATED_VS_4H_LPS_STIM_HDAC3_HET_MACROPHAGE_UP | GSE33162_UNTREATED_VS_4H_LPS_STIM_HDAC3_HET_MACROPHAGE_UP | 13/611 | 198/21355 | 0.0047 | 0.03355 | 0.02489 | 60468/23731/2272/3784/6780/3707/3655/817/9807/50852/55818/23607/9020 | 13 |
| GSE37301_HEMATOPOIETIC_STEM_CELL_VS_PRO_BCELL_UP | GSE37301_HEMATOPOIETIC_STEM_CELL_VS_PRO_BCELL_UP | 13/611 | 198/21355 | 0.0047 | 0.03355 | 0.02489 | 377/1119/8073/9851/23293/10767/3594/55291/8672/9274/50852/51735/64750 | 13 |
| NAKAYA_B_CELL_FLUARIX_FLUVIRIN_AGE_18_50YO_7DY_UP | NAKAYA_B_CELL_FLUARIX_FLUVIRIN_AGE_18_50YO_7DY_UP | 20/611 | 367/21355 | 0.0047 | 0.03355 | 0.02489 | 65125/377/1540/23071/3683/5257/965/10111/23139/6929/10048/6767/26036/5515/7769/10403/5094/8031/51281/23450 | 20 |
| GSE34156_UNTREATED_VS_6H_NOD2_LIGAND_TREATED_MONOCYTE_DN | GSE34156_UNTREATED_VS_6H_NOD2_LIGAND_TREATED_MONOCYTE_DN | 12/611 | 176/21355 | 0.0048 | 0.03355 | 0.02489 | 26065/6774/944/23150/80196/257160/23141/9693/3660/30844/10906/124446 | 12 |
| GSE10239_KLRG1INT_VS_KLRG1HIGH_EFF_CD8_TCELL_DN | GSE10239_KLRG1INT_VS_KLRG1HIGH_EFF_CD8_TCELL_DN | 13/611 | 199/21355 | 0.0049 | 0.03355 | 0.02489 | 132789/65125/79718/4297/3655/5933/50650/157680/1315/10788/8031/9459/79632 | 13 |
| GSE12366_PLASMA_CELL_VS_MEMORY_BCELL_DN | GSE12366_PLASMA_CELL_VS_MEMORY_BCELL_DN | 13/611 | 199/21355 | 0.0049 | 0.03355 | 0.02489 | 60468/51552/64421/57585/23369/84376/55619/23515/9797/4627/9466/283209/388685 | 13 |
| GSE12392_IFNAR_KO_VS_IFNB_KO_CD8_NEG_SPLEEN_DC_UP | GSE12392_IFNAR_KO_VS_IFNB_KO_CD8_NEG_SPLEEN_DC_UP | 13/611 | 199/21355 | 0.0049 | 0.03355 | 0.02489 | 10672/56261/9873/65059/9267/1119/5533/1786/65117/29123/30844/51735/55818 | 13 |
| GSE17301_IFNA2_VS_IFNA2_AND_ACD3_ACD28_STIM_CD8_TCELL_DN | GSE17301_IFNA2_VS_IFNA2_AND_ACD3_ACD28_STIM_CD8_TCELL_DN | 13/611 | 199/21355 | 0.0049 | 0.03355 | 0.02489 | 2869/51552/7414/2272/6780/2926/120/5578/8672/50852/4354/493/104 | 13 |
| GSE1791_CTRL_VS_NEUROMEDINU_IN_T_CELL_LINE_6H_DN | GSE1791_CTRL_VS_NEUROMEDINU_IN_T_CELL_LINE_6H_DN | 13/611 | 199/21355 | 0.0049 | 0.03355 | 0.02489 | 1105/337867/51696/9728/10479/5090/10905/7251/283131/25853/10521/79982/80331 | 13 |
| GSE19941_UNSTIM_VS_LPS_STIM_IL10_KO_NFKBP50_KO_MACROPHAGE_UP | GSE19941_UNSTIM_VS_LPS_STIM_IL10_KO_NFKBP50_KO_MACROPHAGE_UP | 13/611 | 199/21355 | 0.0049 | 0.03355 | 0.02489 | 7799/10390/6526/79718/641/701/55010/23406/7455/5977/55023/5906/27244 | 13 |
| GSE21379_WT_VS_SAP_KO_CD4_TCELL_DN | GSE21379_WT_VS_SAP_KO_CD4_TCELL_DN | 13/611 | 199/21355 | 0.0049 | 0.03355 | 0.02489 | 90273/586/23047/59269/55010/9728/54899/22806/9918/817/51176/114836/56942 | 13 |
| GSE21546_ELK1_KO_VS_SAP1A_KO_AND_ELK1_KO_ANTI_CD3_STIM_DP_THYMOCYTES_UP | GSE21546_ELK1_KO_VS_SAP1A_KO_AND_ELK1_KO_ANTI_CD3_STIM_DP_THYMOCYTES_UP | 13/611 | 199/21355 | 0.0049 | 0.03355 | 0.02489 | 6774/57102/23064/23369/55252/55236/157680/84181/54870/10454/1399/473/6641 | 13 |

| | | | | | | | | |
|---|---|---|---|---|---|---|---|---|
| GSE21546_SAP1A_KO_VS_SAP1A_KO_AND_ELK1_KO_DP_THYMOCYTES_DN | GSE21546_SAP1A_KO_VS_SAP1A_KO_AND_ELK1_KO_DP_THYMOCYTES_DN | 13/611 | 199/21355 | 0.0049 | 0.03355 | 0.02489 | 586/9267/701/4297/721/3707/6929/7455/51460/976/9520/5578/114836 | 13 |
| GSE21546_WT_VS_SAP1A_KO_AND_ELK1_KO_ANTI_CD3_STIM_DP_THYMOCYTES_DN | GSE21546_WT_VS_SAP1A_KO_AND_ELK1_KO_ANTI_CD3_STIM_DP_THYMOCYTES_DN | 13/611 | 199/21355 | 0.0049 | 0.03355 | 0.02489 | 586/23047/59269/337867/701/4297/79813/54899/7455/9918/5578/51176/114836 | 13 |
| GSE22196_HEALTHY_VS_OBESE_MOUSE_SKIN_GAMMADELTA_TCELL_UP | GSE22196_HEALTHY_VS_OBESE_MOUSE_SKIN_GAMMADELTA_TCELL_UP | 13/611 | 199/21355 | 0.0049 | 0.03355 | 0.02489 | 23201/23214/4245/549/23102/157680/9712/6891/23355/10906/2643/51317/473 | 13 |
| GSE23321_CD8_STEM_CELL_MEMORY_VS_EFFECTOR_MEMORY_CD8_TCELL_DN | GSE23321_CD8_STEM_CELL_MEMORY_VS_EFFECTOR_MEMORY_CD8_TCELL_DN | 13/611 | 199/21355 | 0.0049 | 0.03355 | 0.02489 | 10672/51552/23047/23150/7322/7421/4627/3710/5165/1315/9020/4134/10657 | 13 |
| GSE2405_0H_VS_12H_A_PHAGOCYTOPHILUM_STIM_NEUTROPHIL_UP | GSE2405_0H_VS_12H_A_PHAGOCYTOPHILUM_STIM_NEUTROPHIL_UP | 13/611 | 199/21355 | 0.0049 | 0.03355 | 0.02489 | 23370/1265/3106/91775/50807/8705/23406/4215/3560/259230/8428/84433/9953 | 13 |
| GSE24671_BAKIMULC_VS_SENDAI_VIRUS_INFECTED_MOUSE_SPLENOCYTES_DN | GSE24671_BAKIMULC_VS_SENDAI_VIRUS_INFECTED_MOUSE_SPLENOCYTES_DN | 13/611 | 199/21355 | 0.0049 | 0.03355 | 0.02489 | 23132/3064/124245/8705/5933/92170/5165/23198/79613/8065/8726/11184/81671 | 13 |
| GSE25123_CTRL_VS_ROSIGLITAZONE_STIM_MACROPHAGE_UP | GSE25123_CTRL_VS_ROSIGLITAZONE_STIM_MACROPHAGE_UP | 13/611 | 199/21355 | 0.0049 | 0.03355 | 0.02489 | 6777/9267/5048/22992/7322/79663/84961/5588/549/135112/7251/2035/9219 | 13 |
| GSE26156_DOUBLE_POSITIVE_VS_CD4_SINGLE_POSITIVE_THYMOCYTE_UP | GSE26156_DOUBLE_POSITIVE_VS_CD4_SINGLE_POSITIVE_THYMOCYTE_UP | 13/611 | 199/21355 | 0.0049 | 0.03355 | 0.02489 | 23201/9873/23370/9267/22861/9976/7182/3707/3655/5588/120/10956/115 | 13 |
| GSE26495_NAIVE_VS_PD1HIGH_CD8_TCELL_DN | GSE26495_NAIVE_VS_PD1HIGH_CD8_TCELL_DN | 13/611 | 199/21355 | 0.0049 | 0.03355 | 0.02489 | 253461/3683/9976/2081/8073/965/2909/255231/4026/6645/114836/55784/493 | 13 |
| GSE30971_CTRL_VS_LPS_STIM_MACROPHAGE_WBP7_HET_2H_DN | GSE30971_CTRL_VS_LPS_STIM_MACROPHAGE_WBP7_HET_2H_DN | 13/611 | 199/21355 | 0.0049 | 0.03355 | 0.02489 | 23201/10672/55619/55852/10144/23499/5305/10905/64783/4354/83451/163486/473 | 13 |
| GSE32986_UNSTIM_VS_GMCSF_AND_CURDLAN_HIGHDOSE_STIM_DC_DN | GSE32986_UNSTIM_VS_GMCSF_AND_CURDLAN_HIGHDOSE_STIM_DC_DN | 13/611 | 199/21355 | 0.0049 | 0.03355 | 0.02489 | 727/253461/4820/89845/5257/5339/8631/378938/7644/5578/375748/84441/7267 | 13 |
| GSE33292_WT_VS_TCF1_KO_DN3_THYMOCYTE_UP | GSE33292_WT_VS_TCF1_KO_DN3_THYMOCYTE_UP | 13/611 | 199/21355 | 0.0049 | 0.03355 | 0.02489 | 1387/9267/1540/79602/23633/10006/9728/23774/7813/10767/10788/388685/2590 | 13 |
| GSE360_CTRL_VS_L_DONOVANI_DC_UP | GSE360_CTRL_VS_L_DONOVANI_DC_UP | 13/611 | 199/21355 | 0.0049 | 0.03355 | 0.02489 | 79602/1729/7409/79036/9918/23102/5305/5090/259230/23163/4354/124565/7267 | 13 |
| GSE36009_WT_VS_NLRP10_KO_DC_LPS_STIM_DN | GSE36009_WT_VS_NLRP10_KO_DC_LPS_STIM_DN | 13/611 | 199/21355 | 0.0049 | 0.03355 | 0.02489 | 10390/55187/84301/89846/10464/6767/120/5305/23198/10905/7109/163486/7267 | 13 |
| GSE37416_12H_VS_24H_F_TULARENSIS_LVS_NEUTROPHIL_UP | GSE37416_12H_VS_24H_F_TULARENSIS_LVS_NEUTROPHIL_UP | 13/611 | 199/21355 | 0.0049 | 0.03355 | 0.02489 | 85464/64848/1520/165918/9797/11165/7920/4627/54815/3454/7170/9020/51719 | 13 |
| GSE37532_TREG_VS_TCONV_PPARG_KO_CD4_TCELL_FROM_LN_DN | GSE37532_TREG_VS_TCONV_PPARG_KO_CD4_TCELL_FROM_LN_DN | 13/611 | 199/21355 | 0.0049 | 0.03355 | 0.02489 | 23167/3936/6774/59269/23633/256236/200424/55253/25981/7040/7769/8672/9577 | 13 |
| GSE37533_PPARG1_FOXP3_VS_PPARG2_FOXP3_TRANSDUCED_CD4_TCELL_PIOGLITAZONE_TREATED_DN | GSE37533_PPARG1_FOXP3_VS_PPARG2_FOXP3_TRANSDUCED_CD4_TCELL_PIOGLITAZONE_TREATED_DN | 13/611 | 199/21355 | 0.0049 | 0.03355 | 0.02489 | 4763/23304/9267/1540/23064/3106/10892/3660/6891/6645/10906/2643/2289 | 13 |
| GSE3982_BASOPHIL_VS_TH1_UP | GSE3982_BASOPHIL_VS_TH1_UP | 13/611 | 199/21355 | 0.0049 | 0.03355 | 0.02489 | 6777/60468/7414/7456/26065/5090/29761/10788/23013/9129/23287/7403/473 | 13 |
| GSE3982_NEUTROPHIL_VS_EFF_MEMORY_CD4_TCELL_UP | GSE3982_NEUTROPHIL_VS_EFF_MEMORY_CD4_TCELL_UP | 13/611 | 199/21355 | 0.0049 | 0.03355 | 0.02489 | 23214/1105/23071/672/79663/23406/5976/26133/7251/23604/283131/55758/9673 | 13 |
| GSE40273_XBP1_KO_VS_WT_TREG_DN | GSE40273_XBP1_KO_VS_WT_TREG_DN | 13/611 | 199/21355 | 0.0049 | 0.03355 | 0.02489 | 2869/8844/23112/5820/84301/23077/5339/26207/22806/378938/5770/23365/57198 | 13 |
| GSE40277_EOS_AND_LEF1_TRANSDUCED_VS_GATA1_AND_SATB1_TRANSDUCED_CD4_TCELL_DN | GSE40277_EOS_AND_LEF1_TRANSDUCED_VS_GATA1_AND_SATB1_TRANSDUCED_CD4_TCELL_DN | 13/611 | 199/21355 | 0.0049 | 0.03355 | 0.02489 | 672/51108/6249/120/5933/79801/259230/1822/80728/55784/54842/2590/84441 | 13 |
| GSE40277_GATA1_AND_SATB1_TRANSDUCED_VS_CTRL_CD4_TCELL_DN | GSE40277_GATA1_AND_SATB1_TRANSDUCED_VS_CTRL_CD4_TCELL_DN | 13/611 | 199/21355 | 0.0049 | 0.03355 | 0.02489 | 6777/9202/727/23214/1119/79813/83478/5305/26036/114804/1911/23287/7267 | 13 |
| GSE40666_STAT1_KO_VS_STAT4_KO_CD8_TCELL_UP | GSE40666_STAT1_KO_VS_STAT4_KO_CD8_TCELL_UP | 13/611 | 199/21355 | 0.0049 | 0.03355 | 0.02489 | 23370/9735/84301/79828/7182/54838/4215/1432/26133/4931/284058/9459/27244 | 13 |
| GSE45739_UNSTIM_VS_ACD3_ACD28_STIM_WT_CD4_TCELL_DN | GSE45739_UNSTIM_VS_ACD3_ACD28_STIM_WT_CD4_TCELL_DN | 13/611 | 199/21355 | 0.0049 | 0.03355 | 0.02489 | 23130/79602/8826/3683/5775/26207/3560/54815/5305/5900/8672/1871/9263 | 13 |
| GSE4984_GALECTIN1_VS_VEHICLE_CTRL_TREATED_DC_DN | GSE4984_GALECTIN1_VS_VEHICLE_CTRL_TREATED_DC_DN | 13/611 | 199/21355 | 0.0049 | 0.03355 | 0.02489 | 9735/29980/29028/10615/255231/55236/7251/1871/10906/51719/2643/9694/5514 | 13 |
| GSE557_WT_VS_I_AB_KO_DC_DN | GSE557_WT_VS_I_AB_KO_DC_DN | 13/611 | 199/21355 | 0.0049 | 0.03355 | 0.02489 | 9873/150864/26065/23369/55666/55421/1431/3560/51460/5515/9844/84324/79657 | 13 |
| GSE5589_IL6_KO_VS_IL10_KO_LPS_AND_IL10_STIM_MACROPHAGE_45MIN_DN | GSE5589_IL6_KO_VS_IL10_KO_LPS_AND_IL10_STIM_MACROPHAGE_45MIN_DN | 13/611 | 199/21355 | 0.0049 | 0.03355 | 0.02489 | 56261/23731/6526/672/157378/10615/79956/5900/64968/54870/92912/284058/9459 | 13 |
| GSE7548_NAIVE_VS_DAY28_PCC_IMMUNIZATION_CD4_TCELL_UP | GSE7548_NAIVE_VS_DAY28_PCC_IMMUNIZATION_CD4_TCELL_UP | 13/611 | 199/21355 | 0.0049 | 0.03355 | 0.02489 | 23201/23132/146057/4775/9889/9728/10036/1432/1606/3077/9520/23384/23524 | 13 |
| GSE8621_UNSTIM_VS_LPS_STIM_MACROPHAGE_DN | GSE8621_UNSTIM_VS_LPS_STIM_MACROPHAGE_DN | 13/611 | 199/21355 | 0.0049 | 0.03355 | 0.02489 | 9736/6774/823/22887/89846/53944/22848/7187/79036/9466/4700/114799/10198 | 13 |

| | | | | | | | | |
|---|---|---|---|---|---|---|---|---|
| GSE8678_IL7R_LOW_VS_HIGH_EFF_CD8_TCELL_DN | GSE8678_IL7R_LOW_VS_HIGH_EFF_CD8_TCELL_DN | 13/611 | 199/21355 | 0.0049 | 0.03355 | 0.02489 | 9873/5820/1520/6223/255231/1606/5305/9466/64333/51176/23607/114836/27244 | 13 |
| GSE9988_LOW_LPS_VS_VEHICLE_TREATED_MONOCYTE_DN | GSE9988_LOW_LPS_VS_VEHICLE_TREATED_MONOCYTE_DN | 13/611 | 199/21355 | 0.0049 | 0.03355 | 0.02489 | 51696/8073/6794/54838/4215/6613/9466/155038/25853/8428/54521/10521/80331 | 13 |
| GSE11961_FOLLICULAR_BCELL_VS_PLASMA_CELL_DAY7_DN | GSE11961_FOLLICULAR_BCELL_VS_PLASMA_CELL_DAY7_DN | 13/611 | 200/21355 | 0.0051 | 0.03355 | 0.02489 | 6938/84316/53944/5533/5775/1606/549/23019/64333/10788/51176/7982/79632 | 13 |
| GSE13484_12H_VS_3H_YF17D_VACCINE_STIM_PBMC_UP | GSE13484_12H_VS_3H_YF17D_VACCINE_STIM_PBMC_UP | 13/611 | 200/21355 | 0.0051 | 0.03355 | 0.02489 | 585/546/10006/1786/4026/549/54878/9807/157680/11064/57711/8031/6654 | 13 |
| GSE13484_UNSTIM_VS_12H_YF17D_VACCINE_STIM_PBMC_DN | GSE13484_UNSTIM_VS_12H_YF17D_VACCINE_STIM_PBMC_DN | 13/611 | 200/21355 | 0.0051 | 0.03355 | 0.02489 | 28977/4763/56913/23071/6526/3157/23059/3660/22806/259230/55500/10906/4012 | 13 |
| GSE14308_TH2_VS_NAIVE_CD4_TCELL_DN | GSE14308_TH2_VS_NAIVE_CD4_TCELL_DN | 13/611 | 200/21355 | 0.0051 | 0.03355 | 0.02489 | 51552/9889/79602/11320/91775/89846/4215/324/157680/9712/11064/79692/79982 | 13 |
| GSE1432_CTRL_VS_IFNG_24H_MICROGLIA_UP | GSE1432_CTRL_VS_IFNG_24H_MICROGLIA_UP | 13/611 | 200/21355 | 0.0051 | 0.03355 | 0.02489 | 23201/65125/585/4775/7554/10497/3707/1606/8742/55818/23450/27244/7267 | 13 |
| GSE14699_NAIVE_VS_DELETIONAL_TOLERANCE_CD8_TCELL_DN | GSE14699_NAIVE_VS_DELETIONAL_TOLERANCE_CD8_TCELL_DN | 13/611 | 200/21355 | 0.0051 | 0.03355 | 0.02489 | 4775/89846/7421/51108/23476/26207/8904/1606/120/51176/9402/114836/55784 | 13 |
| GSE14769_UNSTIM_VS_240MIN_LPS_BMDM_DN | GSE14769_UNSTIM_VS_240MIN_LPS_BMDM_DN | 13/611 | 200/21355 | 0.0051 | 0.03355 | 0.02489 | 6774/3064/11320/55619/2909/80196/7009/148867/135112/25862/84937/51317/5514 | 13 |
| GSE15330_GRANULOCYTE_MONOCYTE_PROGENITOR_VS_PRO_BCELL_DN | GSE15330_GRANULOCYTE_MONOCYTE_PROGENITOR_VS_PRO_BCELL_DN | 13/611 | 200/21355 | 0.0051 | 0.03355 | 0.02489 | 65125/4775/256435/54899/9611/5515/51/10905/26133/54870/7109/54617/55023 | 13 |
| GSE15330_WT_VS_IKAROS_KO_GRANULOCYTE_MONOCYTE_PROGENITOR_DN | GSE15330_WT_VS_IKAROS_KO_GRANULOCYTE_MONOCYTE_PROGENITOR_DN | 13/611 | 200/21355 | 0.0051 | 0.03355 | 0.02489 | 6777/7414/9267/987/5048/860/5533/1432/3710/4026/3560/54815/283131 | 13 |
| GSE15930_STIM_VS_STIM_AND_IFNAB_72H_CD8_T_CELL_DN | GSE15930_STIM_VS_STIM_AND_IFNAB_72H_CD8_T_CELL_DN | 13/611 | 200/21355 | 0.0051 | 0.03355 | 0.02489 | 54934/987/9782/80196/55852/10615/5305/5090/1822/2643/9694/2289/81671 | 13 |
| GSE15930_STIM_VS_STIM_AND_TRICHOSTATINA_48H_CD8_T_CELL_DN | GSE15930_STIM_VS_STIM_AND_TRICHOSTATINA_48H_CD8_T_CELL_DN | 13/611 | 200/21355 | 0.0051 | 0.03355 | 0.02489 | 2869/9135/22828/23214/3106/1520/871/5578/29761/6891/55500/10906/55275 | 13 |
| GSE17186_NAIVE_VS_CD21HIGH_TRANSITIONAL_BCELL_DN | GSE17186_NAIVE_VS_CD21HIGH_TRANSITIONAL_BCELL_DN | 13/611 | 200/21355 | 0.0051 | 0.03355 | 0.02489 | 51552/29028/701/10615/1786/23443/55236/51460/92181/7798/1871/9459/10800 | 13 |
| GSE17186_NAIVE_VS_CD21LOW_TRANSITIONAL_BCELL_CORD_BLOOD_UP | GSE17186_NAIVE_VS_CD21LOW_TRANSITIONAL_BCELL_CORD_BLOOD_UP | 13/611 | 200/21355 | 0.0051 | 0.03355 | 0.02489 | 9267/23064/6938/51108/5339/26207/120/9466/10905/552900/117584/4012/2590 | 13 |
| GSE17301_CTRL_VS_48H_IFNA2_STIM_CD8_TCELL_UP | GSE17301_CTRL_VS_48H_IFNA2_STIM_CD8_TCELL_UP | 13/611 | 200/21355 | 0.0051 | 0.03355 | 0.02489 | 26065/3683/22861/23369/10892/10464/5339/5775/7187/10048/9466/79058/60685 | 13 |
| GSE17301_IFNA2_VS_IFNA2_AND_ACD3_ACD28_STIM_CD8_TCELL_UP | GSE17301_IFNA2_VS_IFNA2_AND_ACD3_ACD28_STIM_CD8_TCELL_UP | 13/611 | 200/21355 | 0.0051 | 0.03355 | 0.02489 | 1387/7456/9267/987/25777/11320/6249/1729/5775/26207/23102/259230/60685 | 13 |
| GSE17721_0.5H_VS_8H_PAM3CSK4_BMDC_UP | GSE17721_0.5H_VS_8H_PAM3CSK4_BMDC_UP | 13/611 | 200/21355 | 0.0051 | 0.03355 | 0.02489 | 4255/9267/80196/1786/79956/3183/7813/23048/30844/7982/9377/80331/27244 | 13 |
| GSE17721_12H_VS_24H_LPS_BMDC_UP | GSE17721_12H_VS_24H_LPS_BMDC_UP | 13/611 | 200/21355 | 0.0051 | 0.03355 | 0.02489 | 65125/7046/944/89970/1729/23515/5339/54899/8289/30844/10788/117584/128866 | 13 |
| GSE17721_CPG_VS_GARDIQUIMOD_4H_BMDC_DN | GSE17721_CPG_VS_GARDIQUIMOD_4H_BMDC_DN | 13/611 | 200/21355 | 0.0051 | 0.03355 | 0.02489 | 1265/55095/57585/29980/11165/79956/120/23048/6599/259230/9020/56942/80331 | 13 |
| GSE17721_CTRL_VS_CPG_8H_BMDC_DN | GSE17721_CTRL_VS_CPG_8H_BMDC_DN | 13/611 | 200/21355 | 0.0051 | 0.03355 | 0.02489 | 1105/55666/860/257160/51068/23515/54899/57494/5900/2961/9274/114836/9694 | 13 |
| GSE17721_CTRL_VS_GARDIQUIMOD_24H_BMDC_UP | GSE17721_CTRL_VS_GARDIQUIMOD_24H_BMDC_UP | 13/611 | 200/21355 | 0.0051 | 0.03355 | 0.02489 | 28977/23112/90273/7046/146712/8826/23150/3157/221477/7813/9807/7443/2590 | 13 |
| GSE17721_LPS_VS_CPG_1H_BMDC_DN | GSE17721_LPS_VS_CPG_1H_BMDC_DN | 13/611 | 200/21355 | 0.0051 | 0.03355 | 0.02489 | 64766/4204/1119/3106/6780/8904/50650/5090/259230/117584/51317/5514/6654 | 13 |
| GSE17721_LPS_VS_GARDIQUIMOD_2H_BMDC_DN | GSE17721_LPS_VS_GARDIQUIMOD_2H_BMDC_DN | 13/611 | 200/21355 | 0.0051 | 0.03355 | 0.02489 | 377/5496/51552/11329/54834/57585/1455/6249/23515/54838/11165/976/51501 | 13 |
| GSE17721_PAM3CSK4_VS_CPG_8H_BMDC_DN | GSE17721_PAM3CSK4_VS_CPG_8H_BMDC_DN | 13/611 | 200/21355 | 0.0051 | 0.03355 | 0.02489 | 23214/672/7173/3106/7322/80196/257160/26207/1606/50650/23443/128866/84324 | 13 |
| GSE17721_PAM3CSK4_VS_GADIQUIMOD_6H_BMDC_UP | GSE17721_PAM3CSK4_VS_GADIQUIMOD_6H_BMDC_UP | 13/611 | 200/21355 | 0.0051 | 0.03355 | 0.02489 | 3936/7414/79602/255231/5976/1289/7813/9962/976/26036/259230/1739/7109 | 13 |
| GSE17721_POLYIC_VS_CPG_2H_BMDC_UP | GSE17721_POLYIC_VS_CPG_2H_BMDC_UP | 13/611 | 200/21355 | 0.0051 | 0.03355 | 0.02489 | 7456/9267/4245/10464/11165/1432/23048/1911/7798/55818/115/55500/23032 | 13 |
| GSE17721_POLYIC_VS_GARDIQUIMOD_1H_BMDC_UP | GSE17721_POLYIC_VS_GARDIQUIMOD_1H_BMDC_UP | 13/611 | 200/21355 | 0.0051 | 0.03355 | 0.02489 | 1265/944/59269/4297/9967/80196/26528/5295/51455/378938/9274/54617/10330 | 13 |
| GSE17721_POLYIC_VS_GARDIQUIMOD_4H_BMDC_UP | GSE17721_POLYIC_VS_GARDIQUIMOD_4H_BMDC_UP | 13/611 | 200/21355 | 0.0051 | 0.03355 | 0.02489 | 79602/6938/3106/23077/80196/55852/26207/5925/9918/7813/3077/5165/473 | 13 |

| | | | | | | | | |
|---|---|---|---|---|---|---|---|---|
| GSE17721_POLYIC_VS_PAM3CSK4_4H_BMDC_DN | GSE17721_POLYIC_VS_PAM3CSK4_4H_BMDC_DN | 13/611 | 200/21355 | 0.0051 | 0.03355 | 0.02489 | 377/51072/54834/5048/7326/26528/50807/5339/4363/22908/57649/117584/51719 | 13 |
| GSE18281_PERIMEDULLARY_CORTICAL_REGION_VS_WHOLE_CORTEX_THYMUS_UP | GSE18281_PERIMEDULLARY_CORTICAL_REGION_VS_WHOLE_CORTEX_THYMUS_UP | 13/611 | 200/21355 | 0.0051 | 0.03355 | 0.02489 | 23085/64324/7414/23369/79718/5976/10055/5900/10403/26133/23060/23604/65979 | 13 |
| GSE19401_NAIVE_VS_IMMUNIZED_MOUSE_PLN_FOLLICULAR_DC_DN | GSE19401_NAIVE_VS_IMMUNIZED_MOUSE_PLN_FOLLICULAR_DC_DN | 13/611 | 200/21355 | 0.0051 | 0.03355 | 0.02489 | 23201/6777/23649/23214/4245/9976/1432/9466/26036/8742/157680/51/51317 | 13 |
| GSE19401_PAM2CSK4_VS_RETINOIC_ACID_STIM_FOLLICULAR_DC_UP | GSE19401_PAM2CSK4_VS_RETINOIC_ACID_STIM_FOLLICULAR_DC_UP | 13/611 | 200/21355 | 0.0051 | 0.03355 | 0.02489 | 2869/80264/7456/64848/23369/3157/6613/2035/7982/10906/10198/79157/79632 | 13 |
| GSE19923_HEB_KO_VS_HEB_AND_E2A_KO_DP_THYMOCYTE_DN | GSE19923_HEB_KO_VS_HEB_AND_E2A_KO_DP_THYMOCYTE_DN | 13/611 | 200/21355 | 0.0051 | 0.03355 | 0.02489 | 586/7414/944/89846/51068/1060/50650/3594/5770/10788/128866/2643/4791 | 13 |
| GSE19941_UNSTIM_VS_LPS_STIM_IL10_KO_MACROPHAGE_DN | GSE19941_UNSTIM_VS_LPS_STIM_IL10_KO_MACROPHAGE_DN | 13/611 | 200/21355 | 0.0051 | 0.03355 | 0.02489 | 11052/64766/7799/9567/23515/4967/55972/51460/135112/1911/9694/84636/27244 | 13 |
| GSE20366_CD103_POS_VS_NEG_TREG_KLRG1NEG_DN | GSE20366_CD103_POS_VS_NEG_TREG_KLRG1NEG_DN | 13/611 | 200/21355 | 0.0051 | 0.03355 | 0.02489 | 65125/7046/23150/6249/23303/7813/5090/83891/30844/114804/11163/128866/2643 | 13 |
| GSE20715_0H_VS_48H_OZONE_LUNG_DN | GSE20715_0H_VS_48H_OZONE_LUNG_DN | 13/611 | 200/21355 | 0.0051 | 0.03355 | 0.02489 | 23304/3607/9962/79745/871/64968/5888/7798/64783/2289/5514/10963/81671 | 13 |
| GSE21063_WT_VS_NFATC1_KO_BCELL_DN | GSE21063_WT_VS_NFATC1_KO_BCELL_DN | 13/611 | 200/21355 | 0.0051 | 0.03355 | 0.02489 | 9735/9267/89970/10464/1432/405/5094/84937/22834/55758/83990/4791/7403 | 13 |
| GSE22432_MULTIPOTENT_VS_COMMON_DC_PROGENITOR_DN | GSE22432_MULTIPOTENT_VS_COMMON_DC_PROGENITOR_DN | 13/611 | 200/21355 | 0.0051 | 0.03355 | 0.02489 | 55526/944/9567/401409/200424/753/1606/50650/10788/6891/2175/114836/80025 | 13 |
| GSE22432_PDC_VS_TGFB1_TREATEDCOMMON_DC_PROGENITOR_DN | GSE22432_PDC_VS_TGFB1_TREATEDCOMMON_DC_PROGENITOR_DN | 13/611 | 200/21355 | 0.0051 | 0.03355 | 0.02489 | 23731/10006/10892/5533/3707/23406/7920/5295/1432/5305/283209/55758/4354 | 13 |
| GSE22601_DOUBLE_NEGATIVE_VS_CD4_SINGLE_POSITIVE_THYMOCYTE_DN | GSE22601_DOUBLE_NEGATIVE_VS_CD4_SINGLE_POSITIVE_THYMOCYTE_DN | 13/611 | 200/21355 | 0.0051 | 0.03355 | 0.02489 | 132789/8844/6774/81669/54834/79954/2885/9466/5770/8726/283131/55784/10963 | 13 |
| GSE22886_DAY0_VS_DAY1_MONOCYTE_IN_CULTURE_UP | GSE22886_DAY0_VS_DAY1_MONOCYTE_IN_CULTURE_UP | 13/611 | 200/21355 | 0.0051 | 0.03355 | 0.02489 | 6777/545/11329/7414/22861/23077/10144/1432/138151/120/51317/23287/473 | 13 |
| GSE22886_NAIVE_CD4_TCELL_VS_MEMORY_TCELL_DN | GSE22886_NAIVE_CD4_TCELL_VS_MEMORY_TCELL_DN | 13/611 | 200/21355 | 0.0051 | 0.03355 | 0.02489 | 7534/6938/51696/80196/26528/5933/23527/6197/1399/9402/51719/9459/9694 | 13 |
| GSE23502_BM_VS_COLON_TUMOR_MYELOID_DERIVED_SUPPRESSOR_CELL_DN | GSE23502_BM_VS_COLON_TUMOR_MYELOID_DERIVED_SUPPRESSOR_CELL_DN | 13/611 | 200/21355 | 0.0051 | 0.03355 | 0.02489 | 11052/60468/9873/23047/23347/55619/701/23515/7182/8289/10521/84636/27244 | 13 |
| GSE24142_EARLY_THYMIC_PROGENITOR_VS_DN2_THYMOCYTE_ADULT_DN | GSE24142_EARLY_THYMIC_PROGENITOR_VS_DN2_THYMOCYTE_ADULT_DN | 13/611 | 200/21355 | 0.0051 | 0.03355 | 0.02489 | 586/6938/51807/5588/1606/5690/5578/56852/51176/9402/114836/4354/2289 | 13 |
| GSE24210_RESTING_TREG_VS_TCONV_UP | GSE24210_RESTING_TREG_VS_TCONV_UP | 13/611 | 200/21355 | 0.0051 | 0.03355 | 0.02489 | 9267/3683/57690/84458/7187/54899/3710/4026/79801/10129/51317/27244/51720 | 13 |
| GSE24574_BCL6_HIGH_TFH_VS_TFH_CD4_TCELL_DN | GSE24574_BCL6_HIGH_TFH_VS_TFH_CD4_TCELL_DN | 13/611 | 200/21355 | 0.0051 | 0.03355 | 0.02489 | 23201/7799/9267/26065/9736/3707/3655/5588/1606/4627/120/51176/9263 | 13 |
| GSE24634_NAIVE_CD4_TCELL_VS_DAY7_IL4_CONV_TREG_DN | GSE24634_NAIVE_CD4_TCELL_VS_DAY7_IL4_CONV_TREG_DN | 13/611 | 200/21355 | 0.0051 | 0.03355 | 0.02489 | 55526/9735/8847/55252/29028/641/965/55010/8705/5880/5690/5888/9577 | 13 |
| GSE24972_MARGINAL_ZONE_BCELL_VS_FOLLICULAR_BCELL_DN | GSE24972_MARGINAL_ZONE_BCELL_VS_FOLLICULAR_BCELL_DN | 13/611 | 200/21355 | 0.0051 | 0.03355 | 0.02489 | 27031/9735/11320/202052/8631/22806/549/3560/23048/3594/10788/9274/27244 | 13 |
| GSE25085_FETAL_LIVER_VS_FETAL_BM_SP4_THYMIC_IMPLANT_UP | GSE25085_FETAL_LIVER_VS_FETAL_BM_SP4_THYMIC_IMPLANT_UP | 13/611 | 200/21355 | 0.0051 | 0.03355 | 0.02489 | 9735/58513/5257/55619/6223/7745/26133/5888/10992/9459/9129/4354/27244 | 13 |
| GSE25890_CTRL_VS_IL33_IL7_TREATED_NUOCYTES_DN | GSE25890_CTRL_VS_IL33_IL7_TREATED_NUOCYTES_DN | 13/611 | 200/21355 | 0.0051 | 0.03355 | 0.02489 | 11329/23131/64848/84301/81669/8500/9797/51271/7040/2521/6197/92181/283989 | 13 |
| GSE26030_UNSTIM_VS_RESTIM_TH1_DAY15_POST_POLARIZATION_DN | GSE26030_UNSTIM_VS_RESTIM_TH1_DAY15_POST_POLARIZATION_DN | 13/611 | 200/21355 | 0.0051 | 0.03355 | 0.02489 | 7534/11329/10390/7046/4775/3683/401409/701/9918/3077/5305/259230/8428 | 13 |
| GSE2770_UNTREATED_VS_TGFB_AND_IL12_TREATED_ACT_CD4_TCELL_6H_UP | GSE2770_UNTREATED_VS_TGFB_AND_IL12_TREATED_ACT_CD4_TCELL_6H_UP | 13/611 | 200/21355 | 0.0051 | 0.03355 | 0.02489 | 9135/64421/25777/23150/55421/10464/55870/51455/7251/79058/55818/23607/163486 | 13 |
| GSE2770_UNTREATED_VS_TGFB_AND_IL4_TREATED_ACT_CD4_TCELL_4H_UP | GSE2770_UNTREATED_VS_TGFB_AND_IL4_TREATED_ACT_CD4_TCELL_4H_UP | 13/611 | 200/21355 | 0.0051 | 0.03355 | 0.02489 | 51742/23112/9267/51696/5339/54838/9807/3454/23527/51735/114836/4354/80331 | 13 |
| GSE27786_ERYTHROBLAST_VS_MONO_MAC_DN | GSE27786_ERYTHROBLAST_VS_MONO_MAC_DN | 13/611 | 200/21355 | 0.0051 | 0.03355 | 0.02489 | 11329/64421/23071/23077/546/9851/11165/50650/9520/6197/84937/4012/83451 | 13 |
| GSE27786_LIN_NEG_VS_CD4_TCELL_DN | GSE27786_LIN_NEG_VS_CD4_TCELL_DN | 13/611 | 200/21355 | 0.0051 | 0.03355 | 0.02489 | 7799/23167/10390/3683/5048/79718/84166/4297/6223/10048/3560/29123/10992 | 13 |
| GSE27786_LIN_NEG_VS_ERYTHROBLAST_UP | GSE27786_LIN_NEG_VS_ERYTHROBLAST_UP | 13/611 | 200/21355 | 0.0051 | 0.03355 | 0.02489 | 51552/27340/23167/10390/23214/117583/11320/4733/84166/10055/6197/2590/10657 | 13 |

| | | | | | | | | |
|---|---|---|---|---|---|---|---|---|
| GSE27786_LIN_NEG_VS_NEUTROPHIL_DN | GSE27786_LIN_NEG_VS_NEUTROPHIL_DN | 13/611 | 200/21355 | 0.0051 | 0.03355 | 0.02489 | 23130/79602/81669/51696/23515/5339/7187/2885/84937/9577/4354/23287/9694 | 13 |
| GSE28237_FOLLICULAR_VS_EARLY_GC_BCELL_UP | GSE28237_FOLLICULAR_VS_EARLY_GC_BCELL_UP | 13/611 | 200/21355 | 0.0051 | 0.03355 | 0.02489 | 23201/56261/22828/146712/23077/91775/6223/5295/65117/324/23355/9577/55500 | 13 |
| GSE2826_WT_VS_BTK_KO_BCELL_UP | GSE2826_WT_VS_BTK_KO_BCELL_UP | 13/611 | 200/21355 | 0.0051 | 0.03355 | 0.02489 | 4204/7456/23071/79718/5533/3707/5339/3655/255231/1060/1606/3183/4026 | 13 |
| GSE29164_CD8_TCELL_VS_CD8_TCELL_AND_IL12_TREATED_MELANOMA_DAY7_UP | GSE29164_CD8_TCELL_VS_CD8_TCELL_AND_IL12_TREATED_MELANOMA_DAY7_UP | 13/611 | 200/21355 | 0.0051 | 0.03355 | 0.02489 | 23112/23731/7187/55870/729852/7813/7745/283209/7170/23527/10788/9673/473 | 13 |
| GSE31082_DN_VS_DP_THYMOCYTE_UP | GSE31082_DN_VS_DP_THYMOCYTE_UP | 13/611 | 200/21355 | 0.0051 | 0.03355 | 0.02489 | 56005/10111/51108/51068/1203/1431/11165/148867/64968/30844/10788/83451/9694 | 13 |
| GSE31082_DP_VS_CD8_SP_THYMOCYTE_DN | GSE31082_DP_VS_CD8_SP_THYMOCYTE_DN | 13/611 | 200/21355 | 0.0051 | 0.03355 | 0.02489 | 10390/26574/59269/51696/55619/6249/5588/4627/50650/54815/155038/83451/9694 | 13 |
| GSE3203_INFLUENZA_INF_VS_IFNB_TREATED_LN_BCELL_DN | GSE3203_INFLUENZA_INF_VS_IFNB_TREATED_LN_BCELL_DN | 13/611 | 200/21355 | 0.0051 | 0.03355 | 0.02489 | 7414/1119/84301/3747/3480/51807/5339/23141/5588/549/552900/375748/55784 | 13 |
| GSE32128_INOS_DEPENDENT_VS_INOS_INDEPENDENT_ACTIVATED_TCELL_UP | GSE32128_INOS_DEPENDENT_VS_INOS_INDEPENDENT_ACTIVATED_TCELL_UP | 13/611 | 200/21355 | 0.0051 | 0.03355 | 0.02489 | 10672/56261/6777/79602/3106/257160/23186/10425/8289/10767/7040/8031/51719 | 13 |
| GSE32164_RESTING_DIFFERENTIATED_VS_ALTERNATIVELY_ACT_M2_MACROPHAGE_DN | GSE32164_RESTING_DIFFERENTIATED_VS_ALTERNATIVELY_ACT_M2_MACROPHAGE_DN | 13/611 | 200/21355 | 0.0051 | 0.03355 | 0.02489 | 10672/2869/51552/7456/3106/23077/80196/1729/221477/22908/1289/10906/51720 | 13 |
| GSE32423_IL7_VS_IL7_IL4_MEMORY_CD8_TCELL_UP | GSE32423_IL7_VS_IL7_IL4_MEMORY_CD8_TCELL_UP | 13/611 | 200/21355 | 0.0051 | 0.03355 | 0.02489 | 7534/23181/150864/7322/860/10892/4627/50650/9807/92912/283131/57198/27244 | 13 |
| GSE32423_IL7_VS_IL7_IL4_NAIVE_CD8_TCELL_DN | GSE32423_IL7_VS_IL7_IL4_NAIVE_CD8_TCELL_DN | 13/611 | 200/21355 | 0.0051 | 0.03355 | 0.02489 | 22828/4245/5257/5533/23186/8705/5933/65117/50852/5094/6873/63977/23032 | 13 |
| GSE32423_MEMORY_VS_NAIVE_CD8_TCELL_DN | GSE32423_MEMORY_VS_NAIVE_CD8_TCELL_DN | 13/611 | 200/21355 | 0.0051 | 0.03355 | 0.02489 | 23112/51552/22828/23633/9728/26207/5925/26036/84181/50852/1871/9129/27244 | 13 |
| GSE32986_CURDLAN_LOWDOSE_VS_GMCSF_AND_CURDLAN_LOWDOSE_STIM_DC_UP | GSE32986_CURDLAN_LOWDOSE_VS_GMCSF_AND_CURDLAN_LOWDOSE_STIM_DC_UP | 13/611 | 200/21355 | 0.0051 | 0.03355 | 0.02489 | 56261/11329/80196/6249/1786/23303/138151/5933/2885/9918/324/79157/473 | 13 |
| GSE33425_CD8_ALPHAALPHA_VS_ALPHABETA_CD161_HIGH_TCELL_DN | GSE33425_CD8_ALPHAALPHA_VS_ALPHABETA_CD161_HIGH_TCELL_DN | 13/611 | 200/21355 | 0.0051 | 0.03355 | 0.02489 | 3936/3683/51072/641/84316/7182/3707/1786/9960/5588/64333/10198/2590 | 13 |
| GSE34006_A2AR_KO_VS_A2AR_AGONIST_TREATED_TREG_DN | GSE34006_A2AR_KO_VS_A2AR_AGONIST_TREATED_TREG_DN | 13/611 | 200/21355 | 0.0051 | 0.03355 | 0.02489 | 132789/1073/23131/81669/9728/55852/22848/10163/3183/5305/1739/55818/51719 | 13 |
| GSE35685_CD34POS_CD10NEG_CD62LPOS_VS_CD34POS_CD10POS_BONE_MARROW_UP | GSE35685_CD34POS_CD10NEG_CD62LPOS_VS_CD34POS_CD10POS_BONE_MARROW_UP | 13/611 | 200/21355 | 0.0051 | 0.03355 | 0.02489 | 28977/90273/54934/987/23077/84196/4733/10111/2926/1432/80728/51719/7267 | 13 |
| GSE360_DC_VS_MAC_B_MALAYI_HIGH_DOSE_UP | GSE360_DC_VS_MAC_B_MALAYI_HIGH_DOSE_UP | 13/611 | 200/21355 | 0.0051 | 0.03355 | 0.02489 | 6938/3784/7421/1455/5339/10048/4627/549/9466/5515/2035/9020/1178 | 13 |
| GSE360_DC_VS_MAC_L_DONOVANI_DN | GSE360_DC_VS_MAC_L_DONOVANI_DN | 13/611 | 200/21355 | 0.0051 | 0.03355 | 0.02489 | 11329/79602/3683/23150/7009/7409/23443/6613/10788/6197/8065/55818/4354 | 13 |
| GSE360_L_DONOVANI_VS_B_MALAYI_HIGH_DOSE_MAC_DN | GSE360_L_DONOVANI_VS_B_MALAYI_HIGH_DOSE_MAC_DN | 13/611 | 200/21355 | 0.0051 | 0.03355 | 0.02489 | 5496/4255/23214/9567/3683/4297/1729/4967/7409/5933/10992/10906/10330 | 13 |
| GSE36009_UNSTIM_VS_LPS_STIM_DC_DN | GSE36009_UNSTIM_VS_LPS_STIM_DC_DN | 13/611 | 200/21355 | 0.0051 | 0.03355 | 0.02489 | 56261/7046/56913/51072/59269/55421/1203/4363/6873/7109/4791/9219/26043 | 13 |
| GSE36392_TYPE_2_MYELOID_VS_EOSINOPHIL_IL25_TREATED_LUNG_DN | GSE36392_TYPE_2_MYELOID_VS_EOSINOPHIL_IL25_TREATED_LUNG_DN | 13/611 | 200/21355 | 0.0051 | 0.03355 | 0.02489 | 56261/23112/3683/337867/3480/91775/84830/54899/2961/163702/2643/9673/2289 | 13 |
| GSE36527_CD62L_HIGH_VS_CD62L_LOW_TREG_CD69_NEG_KLRG1_NEG_UP | GSE36527_CD62L_HIGH_VS_CD62L_LOW_TREG_CD69_NEG_KLRG1_NEG_UP | 13/611 | 200/21355 | 0.0051 | 0.03355 | 0.02489 | 7534/7456/1105/81669/860/7009/405/117584/11163/128866/8428/10906/51720 | 13 |
| GSE36891_UNSTIM_VS_POLYIC_TLR3_STIM_PERITONEAL_MACROPHAGE_DN | GSE36891_UNSTIM_VS_POLYIC_TLR3_STIM_PERITONEAL_MACROPHAGE_DN | 13/611 | 200/21355 | 0.0051 | 0.03355 | 0.02489 | 9887/9267/11320/80196/23515/5933/50650/29761/23013/128866/10800/163486/10657 | 13 |
| GSE3691_IFN_PRODUCING_KILLER_DC_VS_CONVENTIONAL_DC_SPLEEN_UP | GSE3691_IFN_PRODUCING_KILLER_DC_VS_CONVENTIONAL_DC_SPLEEN_UP | 13/611 | 200/21355 | 0.0051 | 0.03355 | 0.02489 | 7046/146057/3064/79718/83478/22806/3710/9466/9807/51/4134/55758/4354 | 13 |
| GSE37301_MULTIPOTENT_PROGENITOR_VS_CD4_TCELL_DN | GSE37301_MULTIPOTENT_PROGENITOR_VS_CD4_TCELL_DN | 13/611 | 200/21355 | 0.0051 | 0.03355 | 0.02489 | 11329/727/7414/546/2909/80196/221477/1431/7813/54878/9807/23019/8031 | 13 |
| GSE37301_MULTIPOTENT_PROGENITOR_VS_RAG2_KO_NK_CELL_UP | GSE37301_MULTIPOTENT_PROGENITOR_VS_RAG2_KO_NK_CELL_UP | 13/611 | 200/21355 | 0.0051 | 0.03355 | 0.02489 | 54934/23731/727/944/84196/2926/79828/51271/22908/54870/7109/8031/51720 | 13 |
| GSE39110_DAY3_VS_DAY6_POST_IMMUNIZATION_CD8_TCELL_WITH_IL2_TREATMENT_UP | GSE39110_DAY3_VS_DAY6_POST_IMMUNIZATION_CD8_TCELL_WITH_IL2_TREATMENT_UP | 13/611 | 200/21355 | 0.0051 | 0.03355 | 0.02489 | 51072/401409/23347/701/22806/9918/4026/79801/10403/30844/10906/83990/2289 | 13 |
| GSE39556_UNTREATED_VS_3H_POLYIC_INJ_MOUSE_NK_CELL_DN | GSE39556_UNTREATED_VS_3H_POLYIC_INJ_MOUSE_NK_CELL_DN | 13/611 | 200/21355 | 0.0051 | 0.03355 | 0.02489 | 11329/146057/1105/25777/22887/2272/9728/23774/5533/9693/10788/4354/493 | 13 |

| | | | | | | | | |
|---|---|---|---|---|---|---|---|---|
| GSE3982_NEUTROPHIL_VS_CENT_MEMORY_CD4_TCELL_DN | GSE3982_NEUTROPHIL_VS_CENT_MEMORY_CD4_TCELL_DN | 13/611 | 200/21355 | 0.0051 | 0.03355 | 0.02489 | 9698/26065/89845/84830/10892/5533/120/50650/79745/23019/8726/55500/54521 | 13 |
| GSE3982_NEUTROPHIL_VS_CENT_MEMORY_CD4_TCELL_UP | GSE3982_NEUTROPHIL_VS_CENT_MEMORY_CD4_TCELL_UP | 13/611 | 200/21355 | 0.0051 | 0.03355 | 0.02489 | 9567/79663/8705/23406/9693/3660/976/8672/23355/1871/23287/7403/51720 | 13 |
| GSE3982_NKCELL_VS_TH2_DN | GSE3982_NKCELL_VS_TH2_DN | 13/611 | 200/21355 | 0.0051 | 0.03355 | 0.02489 | 23649/27340/59269/29028/23150/51068/6767/126298/9918/5515/30844/5888/1871 | 13 |
| GSE3994_WT_VS_PAC1_KO_ACTIVATED_MAST_CELL_DN | GSE3994_WT_VS_PAC1_KO_ACTIVATED_MAST_CELL_DN | 13/611 | 200/21355 | 0.0051 | 0.03355 | 0.02489 | 9202/10390/23130/944/5048/9491/23059/7920/8904/4026/817/157680/79657 | 13 |
| GSE40184_HEALTHY_VS_HCV_INFECTED_DONOR_PBMC_DN | GSE40184_HEALTHY_VS_HCV_INFECTED_DONOR_PBMC_DN | 13/611 | 200/21355 | 0.0051 | 0.03355 | 0.02489 | 65125/64421/84830/55852/5533/5976/6891/2035/22834/284058/8428/11184/9953 | 13 |
| GSE40274_FOXP3_VS_FOXP3_AND_EOS_TRANSDUCED_ACTIVATED_CD4_TCELL_UP | GSE40274_FOXP3_VS_FOXP3_AND_EOS_TRANSDUCED_ACTIVATED_CD4_TCELL_UP | 13/611 | 200/21355 | 0.0051 | 0.03355 | 0.02489 | 90273/585/1203/83478/9960/23384/64333/155038/10788/9402/8428/83451/84636 | 13 |
| GSE40274_HELIOS_VS_FOXP3_AND_HELIOS_TRANSDUCED_ACTIVATED_CD4_TCELL_UP | GSE40274_HELIOS_VS_FOXP3_AND_HELIOS_TRANSDUCED_ACTIVATED_CD4_TCELL_UP | 13/611 | 200/21355 | 0.0051 | 0.03355 | 0.02489 | 6777/586/256435/4297/2926/5339/4363/5925/146691/22806/3560/50650/27244 | 13 |
| GSE40655_FOXO1_KO_VS_WT_NTREG_UP | GSE40655_FOXO1_KO_VS_WT_NTREG_UP | 13/611 | 200/21355 | 0.0051 | 0.03355 | 0.02489 | 55690/25777/23077/84196/7187/26207/55291/23365/23163/51735/5205/493/54842 | 13 |
| GSE411_100MIN_VS_400MIN_IL6_STIM_MACROPHAGE_UP | GSE411_100MIN_VS_400MIN_IL6_STIM_MACROPHAGE_UP | 13/611 | 200/21355 | 0.0051 | 0.03355 | 0.02489 | 23181/150864/57585/337867/7182/1786/3660/4627/5933/23607/284058/9459/473 | 13 |
| GSE41176_UNSTIM_VS_ANTI_IGM_STIM_BCELL_24H_DN | GSE41176_UNSTIM_VS_ANTI_IGM_STIM_BCELL_24H_DN | 13/611 | 200/21355 | 0.0051 | 0.03355 | 0.02489 | 59269/54834/3106/84196/55852/10163/1432/1778/65117/8289/5515/80728/6654 | 13 |
| GSE41176_UNSTIM_VS_ANTI_IGM_STIM_TAK1_KO_BCELL_1H_UP | GSE41176_UNSTIM_VS_ANTI_IGM_STIM_TAK1_KO_BCELL_1H_UP | 13/611 | 200/21355 | 0.0051 | 0.03355 | 0.02489 | 57102/3747/23369/84316/6249/7182/92170/64333/9577/114836/84324/31/9694 | 13 |
| GSE41867_NAIVE_VS_DAY15_LCMV_EFFECTOR_CD8_TCELL_DN | GSE41867_NAIVE_VS_DAY15_LCMV_EFFECTOR_CD8_TCELL_DN | 13/611 | 200/21355 | 0.0051 | 0.03355 | 0.02489 | 85464/1387/23731/64324/4820/5048/23476/4627/120/84181/9712/259230/51317 | 13 |
| GSE41867_NAIVE_VS_DAY6_LCMV_EFFECTOR_CD8_TCELL_DN | GSE41867_NAIVE_VS_DAY6_LCMV_EFFECTOR_CD8_TCELL_DN | 13/611 | 200/21355 | 0.0051 | 0.03355 | 0.02489 | 85464/64324/9267/3683/23293/55870/4627/1778/51586/5900/84181/4012/163486 | 13 |
| GSE41867_NAIVE_VS_DAY8_LCMV_EFFECTOR_CD8_TCELL_DN | GSE41867_NAIVE_VS_DAY8_LCMV_EFFECTOR_CD8_TCELL_DN | 13/611 | 200/21355 | 0.0051 | 0.03355 | 0.02489 | 56261/23370/10390/64324/9267/55095/155435/337867/23293/10048/9960/4012/51317 | 13 |
| GSE43955_10H_VS_60H_ACT_CD4_TCELL_DN | GSE43955_10H_VS_60H_ACT_CD4_TCELL_DN | 13/611 | 200/21355 | 0.0051 | 0.03355 | 0.02489 | 10672/2869/4763/79602/7326/337867/3480/84316/255231/10767/11064/9274/55818 | 13 |
| GSE44649_NAIVE_VS_ACTIVATED_CD8_TCELL_DN | GSE44649_NAIVE_VS_ACTIVATED_CD8_TCELL_DN | 13/611 | 200/21355 | 0.0051 | 0.03355 | 0.02489 | 51696/29028/79718/641/6249/55852/55253/165918/23186/54165/9712/25853/57534 | 13 |
| GSE46606_UNSTIM_VS_CD40L_IL2_IL5_3DAY_STIMULATED_IRF4MID_SORTED_BCELL_DN | GSE46606_UNSTIM_VS_CD40L_IL2_IL5_3DAY_STIMULATED_IRF4MID_SORTED_BCELL_DN | 13/611 | 200/21355 | 0.0051 | 0.03355 | 0.02489 | 4763/90273/165918/4795/3183/5933/2885/50650/2961/10906/4791/5514/26043 | 13 |
| GSE4748_CTRL_VS_CYANOBACTERIUM_LPSLIKE_STIM_DC_1H_UP | GSE4748_CTRL_VS_CYANOBACTERIUM_LPSLIKE_STIM_DC_1H_UP | 13/611 | 200/21355 | 0.0051 | 0.03355 | 0.02489 | 10390/8826/25777/7322/57634/5339/84961/5690/23019/552900/8428/51719/2643 | 13 |
| GSE5142_HTERT_TRANSDUCED_VS_CTRL_CD8_TCELL_LATE_PASSAGE_CLONE_DN | GSE5142_HTERT_TRANSDUCED_VS_CTRL_CD8_TCELL_LATE_PASSAGE_CLONE_DN | 13/611 | 200/21355 | 0.0051 | 0.03355 | 0.02489 | 6777/3707/54838/23303/84193/8289/976/51176/54617/197135/4012/54842/2590 | 13 |
| GSE5589_LPS_AND_IL10_VS_LPS_AND_IL6_STIM_IL6_KO_MACROPHAGE_45MIN_DN | GSE5589_LPS_AND_IL10_VS_LPS_AND_IL6_STIM_IL6_KO_MACROPHAGE_45MIN_DN | 13/611 | 200/21355 | 0.0051 | 0.03355 | 0.02489 | 56261/23731/65059/7046/9736/1431/6767/283209/5900/64968/552900/4700/7109 | 13 |
| GSE5589_UNSTIM_VS_45MIN_LPS_AND_IL10_STIM_MACROPHAGE_UP | GSE5589_UNSTIM_VS_45MIN_LPS_AND_IL10_STIM_MACROPHAGE_UP | 13/611 | 200/21355 | 0.0051 | 0.03355 | 0.02489 | 3106/11320/23476/10048/7009/4795/5295/5977/57410/976/6891/8726/678655 | 13 |
| GSE7460_CTRL_VS_TGFB_TREATED_ACT_FOXP3_MUT_TCONV_UP | GSE7460_CTRL_VS_TGFB_TREATED_ACT_FOXP3_MUT_TCONV_UP | 13/611 | 200/21355 | 0.0051 | 0.03355 | 0.02489 | 2869/8844/171023/65059/150864/8073/5339/549/50650/259230/4134/84636/6654 | 13 |
| GSE7764_NKCELL_VS_SPLENOCYTE_DN | GSE7764_NKCELL_VS_SPLENOCYTE_DN | 13/611 | 200/21355 | 0.0051 | 0.03355 | 0.02489 | 7534/56261/6223/10048/54899/51271/5925/1606/79801/259230/9577/23607/9219 | 13 |
| GSE8621_LPS_STIM_VS_LPS_PRIMED_AND_LPS_STIM_MACROPHAGE_DN | GSE8621_LPS_STIM_VS_LPS_PRIMED_AND_LPS_STIM_MACROPHAGE_DN | 13/611 | 200/21355 | 0.0051 | 0.03355 | 0.02489 | 10672/56261/9887/3936/7456/987/55619/5533/165918/753/7813/56852/8726 | 13 |
| GSE8921_UNSTIM_0H_VS_TLR1_2_STIM_MONOCYTE_12H_DN | GSE8921_UNSTIM_0H_VS_TLR1_2_STIM_MONOCYTE_12H_DN | 13/611 | 200/21355 | 0.0051 | 0.03355 | 0.02489 | 145942/56913/57102/84301/81669/5775/54838/7409/8904/7251/10521/51317/51720 | 13 |
| GSE8921_UNSTIM_0H_VS_TLR1_2_STIM_MONOCYTE_24H_DN | GSE8921_UNSTIM_0H_VS_TLR1_2_STIM_MONOCYTE_24H_DN | 13/611 | 200/21355 | 0.0051 | 0.03355 | 0.02489 | 11052/10390/3064/53944/10048/3710/4026/51/8031/55500/80331/84636/81671 | 13 |
| GSE9650_GP33_VS_GP276_LCMV_SPECIFIC_EXHAUSTED_CD8_TCELL_DN | GSE9650_GP33_VS_GP276_LCMV_SPECIFIC_EXHAUSTED_CD8_TCELL_DN | 13/611 | 200/21355 | 0.0051 | 0.03355 | 0.02489 | 5496/11329/4670/3683/51108/80196/26207/10055/22908/10425/9962/51176/51501 | 13 |
| GSE21033_CTRL_VS_POLYIC_STIM_DC_6H_DN | GSE21033_CTRL_VS_POLYIC_STIM_DC_6H_DN | 11/611 | 157/21355 | 0.0056 | 0.03642 | 0.02703 | 1387/3683/55252/22806/1954/23102/23248/29761/84937/23607/163486 | 11 |
| GSE4590_PRE_BCELL_VS_VPREB_POS_LARGE_PRE_BCELL_DN | GSE4590_PRE_BCELL_VS_VPREB_POS_LARGE_PRE_BCELL_DN | 11/611 | 157/21355 | 0.0056 | 0.03642 | 0.02703 | 9135/23304/64324/7326/51108/84961/55972/2961/79613/63977/27244 | 11 |

| | | | | | | | | |
|---|---|---|---|---|---|---|---|---|
| GSE13547_WT_VS_ZFX_KO_BCELL_ANTI_IGM_STIM_2H_UP | GSE13547_WT_VS_ZFX_KO_BCELL_ANTI_IGM_STIM_2H_UP | 12/611 | 181/21355 | 0.006 | 0.03921 | 0.02909 | 11052/545/27340/51072/3157/9960/10055/23443/5690/5888/83990/2289 | 12 |
| GSE22611_NOD2_TRANSDUCED_VS_CTRL_HEK293T_STIMULATED_WITH_MDP_2H_DN | GSE22611_NOD2_TRANSDUCED_VS_CTRL_HEK293T_STIMULATED_WITH_MDP_2H_DN | 12/611 | 183/21355 | 0.0066 | 0.04264 | 0.03164 | 23085/9735/586/23511/3069/91775/221477/4363/1786/255231/23450/9377 | 12 |
| GSE24671_CTRL_VS_SENDAI_VIRUS_INFECTED_MOUSE_SPLENOCYTES_UP | GSE24671_CTRL_VS_SENDAI_VIRUS_INFECTED_MOUSE_SPLENOCYTES_UP | 11/611 | 161/21355 | 0.0067 | 0.04364 | 0.03238 | 23304/11320/3480/55619/1520/23499/378938/135112/120425/55023/55784 | 11 |
| GSE3565_DUSP1_VS_WT_SPLENOCYTES_POST_LPS_INJECTION_UP | GSE3565_DUSP1_VS_WT_SPLENOCYTES_POST_LPS_INJECTION_UP | 11/611 | 164/21355 | 0.0077 | 0.04982 | 0.03697 | 9873/944/23047/81669/53944/23476/51068/57649/64333/51735/55500 | 11 |
