## Appendix Table S3 for "Distinguishable topological properties of functional genome networks in HIV-1 reservoirs"

**Supplementary Table 3. List of enriched immunologic signatures harboring intact proviruses in elite controllers**

| ID | Description | GeneRatio | BgRatio | pvalue | p.adjust | qvalue | geneID | Count |
|---|---|---|---|---|---|---|---|---|
| GSE11057_CD4_EFF_MEM_VS_PBMC_UP | GSE11057_CD4_EFF_MEM_VS_PBMC_UP | 8/81 | 199/21355 | 8.88E-07 | 0.0028905 | 0.00251 | 114804/9252/51466/3835/6777/288/89894/54856 | 8 |
| GSE40225_WT_VS_RIP_B7X_DIABETIC_MOUSE_PANCREATIC_CD8_TCELL_UP | GSE40225_WT_VS_RIP_B7X_DIABETIC_MOUSE_PANCREATIC_CD8_TCELL_UP | 7/81 | 199/21355 | 1.07E-05 | 0.0100474 | 0.00874 | 4926/55770/63892/9873/51466/64853/23049 | 7 |
| GSE31082_DN_VS_CD4_SP_THYMOCYTE_DN | GSE31082_DN_VS_CD4_SP_THYMOCYTE_DN | 7/81 | 200/21355 | 1.1E-05 | 0.0100474 | 0.00874 | 8675/8897/55209/23214/83852/54664/116984 | 7 |
| NAKAYA_PBMC_FLUARIX_FLUVIRIN_AGE_18_50YO_CORRELATED_WITH_HAI_28DY_RESPONSE_AT_7DY_NEGATIVE | NAKAYA_PBMC_FLUARIX_FLUVIRIN_AGE_18_50YO_CORRELATED_WITH_HAI_28DY_RESPONSE_AT_7DY_NEGATIVE | 10/81 | 475/21355 | 1.23E-05 | 0.0100474 | 0.00874 | 4926/10782/6774/378938/1997/6777/7109/288/23049/54856 | 10 |
| GSE25146_UNSTIM_VS_HELIOBACTER_PYLORI_LPS_STIM_AGS_CELL_DN | GSE25146_UNSTIM_VS_HELIOBACTER_PYLORI_LPS_STIM_AGS_CELL_DN | 6/81 | 166/21355 | 4.04E-05 | 0.0263093 | 0.02288 | 27327/10521/953/55870/8418/54739 | 6 |
| GSE26488_CTRL_VS_PEPTIDE_INJECTION_OT2_THYMOCYTE_DN | GSE26488_CTRL_VS_PEPTIDE_INJECTION_OT2_THYMOCYTE_DN | 6/81 | 183/21355 | 6.95E-05 | 0.0263613 | 0.02293 | 64744/9873/8418/51735/57459/64853 | 6 |
| GSE5099_DAY3_VS_DAY7_MCSF_TREATED_MACROPHAGE_DN | GSE5099_DAY3_VS_DAY7_MCSF_TREATED_MACROPHAGE_DN | 6/81 | 183/21355 | 6.95E-05 | 0.0263613 | 0.02293 | 378938/57690/8418/23397/57459/1416 | 6 |
| GSE11057_NAIVE_CD4_VS_PBMC_CD4_TCELL_UP | GSE11057_NAIVE_CD4_VS_PBMC_CD4_TCELL_UP | 6/81 | 197/21355 | 0.000104 | 0.0263613 | 0.02293 | 55628/9252/51466/23214/6777/54856 | 6 |
| GSE16450_CTRL_VS_IFNA_12H_STIM_IMMATURE_NEURON_CELL_LINE_UP | GSE16450_CTRL_VS_IFNA_12H_STIM_IMMATURE_NEURON_CELL_LINE_UP | 6/81 | 197/21355 | 0.000104 | 0.0263613 | 0.02293 | 27327/55628/63892/9873/23315/64754 | 6 |
| GSE20366_TREG_VS_NAIVE_CD4_TCELL_HOMEOSTATIC_CONVERSION_DN | GSE20366_TREG_VS_NAIVE_CD4_TCELL_HOMEOSTATIC_CONVERSION_DN | 6/81 | 199/21355 | 0.00011 | 0.0263613 | 0.02293 | 57514/64744/9873/57690/23048/7109 | 6 |
| GSE22601_IMMATURE_CD4_SINGLE_POSITIVE_VS_CD8_SINGLE_POSITIVE_THYMOCYTE_DN | GSE22601_IMMATURE_CD4_SINGLE_POSITIVE_VS_CD8_SINGLE_POSITIVE_THYMOCYTE_DN | 6/81 | 200/21355 | 0.000113 | 0.0263613 | 0.02293 | 8897/6774/54739/3091/51735/116984 | 6 |
| GSE30971_WBP7_HET_VS_KO_MACROPHAGE_UP | GSE30971_WBP7_HET_VS_KO_MACROPHAGE_UP | 6/81 | 200/21355 | 0.000113 | 0.0263613 | 0.02293 | 114804/5034/57514/8897/6774/3091 | 6 |
| GSE5142_CTRL_VS_HTERT_TRANSDUCED_CD8_TCELL_EARLY_PASSAGE_CLONE_UP | GSE5142_CTRL_VS_HTERT_TRANSDUCED_CD8_TCELL_EARLY_PASSAGE_CLONE_UP | 6/81 | 200/21355 | 0.000113 | 0.0263613 | 0.02293 | 4926/114804/10521/81671/3091/6777 | 6 |
| GSE6674_ANTI_IGM_VS_CPG_STIM_BCELL_UP | GSE6674_ANTI_IGM_VS_CPG_STIM_BCELL_UP | 6/81 | 200/21355 | 0.000113 | 0.0263613 | 0.02293 | 10782/283237/9252/51466/7328/54664 | 6 |
