## Appendix Table S4 for "Distinguishable topological properties of functional genome networks in HIV-1 reservoirs"

**Supplementary Table 4. List of enriched immunologic signatures harboring defective proviruses in elite controllers**

| ID | Description | GeneRatio | BgRatio | pvalue | p.adjust | qvalue | geneID | Count |
|---|---|---|---|---|---|---|---|---|
| GSE27241_WT_VS_RORGT_KO_TH17_POLARIZED_CD4_TCELL_TREATED_WITH_DIGOXIN_UP | GSE27241_WT_VS_RORGT_KO_TH17_POLARIZED_CD4_TCELL_TREATED_WITH_DIGOXIN_UP | 10/95 | 180/21355 | 7.667E-09 | 2.6727E-05 | 2.506E-05 | 22992/9873/23085/1387/4763/171023/55677/23731/65125/9735 | 10 |
| GSE369_IFNG_KO_VS_WT_LIVER_UP | GSE369_IFNG_KO_VS_WT_LIVER_UP | 8/95 | 199/21355 | 3.002E-06 | 0.00271518 | 0.0025457 | 64421/221037/143/10743/8675/23304/60468/85464 | 8 |
| GSE16385_ROSIGLITAZONE_IL4_VS_IFNG_TNF_STIM_MACROPHAGE_UP | GSE16385_ROSIGLITAZONE_IL4_VS_IFNG_TNF_STIM_MACROPHAGE_UP | 8/95 | 200/21355 | 3.116E-06 | 0.00271518 | 0.0025457 | 22992/4753/1387/171023/8675/113791/60468/51742 | 8 |
| GSE21670_UNTREATED_VS_IL6_TREATED_CD4_TCELL_DN | GSE21670_UNTREATED_VS_IL6_TREATED_CD4_TCELL_DN | 8/95 | 200/21355 | 3.116E-06 | 0.00271518 | 0.0025457 | 55690/22992/51466/113791/5910/11329/10807/23112 | 8 |
| GSE11386_NAIVE_VS_MEMORY_BCELL_DN | GSE11386_NAIVE_VS_MEMORY_BCELL_DN | 7/95 | 159/21355 | 7.091E-06 | 0.00494392 | 0.0046353 | 221037/51466/113791/60468/1666/64766/64324 | 7 |
| GSE27291_0H_VS_7D_STIM_GAMMADELTA_TCELL_UP | GSE27291_0H_VS_7D_STIM_GAMMADELTA_TCELL_UP | 7/95 | 188/21355 | 2.107E-05 | 0.01058027 | 0.0099199 | 55526/90592/51278/10672/10260/64766/23131 | 7 |
| GSE12845_IGD_POS_BLOOD_VS_NAIVE_TONSIL_BCELL_DN | GSE12845_IGD_POS_BLOOD_VS_NAIVE_TONSIL_BCELL_DN | 7/95 | 199/21355 | 3.035E-05 | 0.01058027 | 0.0099199 | 23013/22992/54934/2885/2869/23112/9698 | 7 |
| GSE24574_BCL6_HIGH_VS_LOW_TFH_CD4_TCELL_UP | GSE24574_BCL6_HIGH_VS_LOW_TFH_CD4_TCELL_UP | 7/95 | 199/21355 | 3.035E-05 | 0.01058027 | 0.0099199 | 7799/22992/9873/23370/113791/7534/51742 | 7 |
| GSE27786_NKTCELL_VS_ERYTHROBLAST_UP | GSE27786_NKTCELL_VS_ERYTHROBLAST_UP | 7/95 | 199/21355 | 3.035E-05 | 0.01058027 | 0.0099199 | 9202/54802/8675/51278/132789/2869/28977 | 7 |
| GSE40273_GATA1_KO_VS_WT_TREG_DN | GSE40273_GATA1_KO_VS_WT_TREG_DN | 7/95 | 199/21355 | 3.035E-05 | 0.01058027 | 0.0099199 | 55690/9202/9742/55114/23370/8675/60468 | 7 |
| NAKAYA_PBMC_FLUARIX_FLUVIRIN_AGE_18_50YO_3DY_DN | NAKAYA_PBMC_FLUARIX_FLUVIRIN_AGE_18_50YO_3DY_DN | 9/95 | 433/21355 | 0.0001343 | 0.02713021 | 0.0254369 | 221037/7799/22992/54934/321/80264/171023/113791/4670 | 9 |
| GSE29618_BCELL_VS_MDC_DAY7_FLU_VACCINE_UP | GSE29618_BCELL_VS_MDC_DAY7_FLU_VACCINE_UP | 6/95 | 192/21355 | 0.0002187 | 0.02713021 | 0.0254369 | 7799/9873/23370/113791/60468/51742 | 6 |
| GSE29618_BCELL_VS_MDC_UP | GSE29618_BCELL_VS_MDC_UP | 6/95 | 194/21355 | 0.0002312 | 0.02713021 | 0.0254369 | 7799/9873/80264/113791/60468/2869 | 6 |
| GSE16450_IMMATURE_VS_MATURE_NEURON_CELL_LINE_UP | GSE16450_IMMATURE_VS_MATURE_NEURON_CELL_LINE_UP | 6/95 | 195/21355 | 0.0002377 | 0.02713021 | 0.0254369 | 321/4763/23370/113791/85464/23112 | 6 |
| GSE23321_CD8_STEM_CELL_MEMORY_VS_NAIVE_CD8_TCELL_DN | GSE23321_CD8_STEM_CELL_MEMORY_VS_NAIVE_CD8_TCELL_DN | 6/95 | 196/21355 | 0.0002444 | 0.02713021 | 0.0254369 | 4763/5116/80345/1666/85464/23112 | 6 |
| GSE29618_BCELL_VS_MONOCYTE_DAY7_FLU_VACCINE_UP | GSE29618_BCELL_VS_MONOCYTE_DAY7_FLU_VACCINE_UP | 6/95 | 196/21355 | 0.0002444 | 0.02713021 | 0.0254369 | 9873/9135/80264/23370/60468/51742 | 6 |
| GSE11057_NAIVE_CD4_VS_PBMC_CD4_TCELL_UP | GSE11057_NAIVE_CD4_VS_PBMC_CD4_TCELL_UP | 6/95 | 197/21355 | 0.0002512 | 0.02713021 | 0.0254369 | 51466/23370/171023/113791/23731/6777 | 6 |
| GSE11864_CSF1_VS_CSF1_PAM3CYS_IN_MAC_UP | GSE11864_CSF1_VS_CSF1_PAM3CYS_IN_MAC_UP | 6/95 | 197/21355 | 0.0002512 | 0.02713021 | 0.0254369 | 221037/121665/4542/80345/11329/1073 | 6 |
| GSE21063_3H_VS_16H_ANTI_IGM_STIM_BCELL_DN | GSE21063_3H_VS_16H_ANTI_IGM_STIM_BCELL_DN | 6/95 | 197/21355 | 0.0002512 | 0.02713021 | 0.0254369 | 23085/4753/9202/79932/545/1666 | 6 |
| GSE13411_IGM_MEMORY_BCELL_VS_PLASMA_CELL_UP | GSE13411_IGM_MEMORY_BCELL_VS_PLASMA_CELL_UP | 6/95 | 198/21355 | 0.0002581 | 0.02713021 | 0.0254369 | 51466/55114/80264/60468/3117/6777 | 6 |
| GSE19198_6H_VS_24H_IL21_TREATED_TCELL_DN | GSE19198_6H_VS_24H_IL21_TREATED_TCELL_DN | 6/95 | 198/21355 | 0.0002581 | 0.02713021 | 0.0254369 | 221037/4591/2885/113791/10807/85464 | 6 |
| GSE25088_IL4_VS_IL4_AND_ROSIGLITAZONE_STIM_STAT6_KO_MACROPHAGE_DAY10_UP | GSE25088_IL4_VS_IL4_AND_ROSIGLITAZONE_STIM_STAT6_KO_MACROPHAGE_DAY10_UP | 6/95 | 198/21355 | 0.0002581 | 0.02713021 | 0.0254369 | 55526/55690/4542/8675/85464/23112 | 6 |
| GSE36476_CTRL_VS_TSST_ACT_72H_MEMORY_CD4_TCELL_YOUNG_UP | GSE36476_CTRL_VS_TSST_ACT_72H_MEMORY_CD4_TCELL_YOUNG_UP | 6/95 | 198/21355 | 0.0002581 | 0.02713021 | 0.0254369 | 54934/55671/321/1387/23370/113791 | 6 |
| GSE10239_NAIVE_VS_KLRG1HIGH_EFF_CD8_TCELL_UP | GSE10239_NAIVE_VS_KLRG1HIGH_EFF_CD8_TCELL_UP | 6/95 | 199/21355 | 0.0002652 | 0.02713021 | 0.0254369 | 51466/113791/23304/60468/51278/85464 | 6 |
| GSE22886_NAIVE_TCELL_VS_DC_UP | GSE22886_NAIVE_TCELL_VS_DC_UP | 6/95 | 199/21355 | 0.0002652 | 0.02713021 | 0.0254369 | 4753/80264/171023/113791/11329/6777 | 6 |
| GSE3982_MAST_CELL_VS_BASOPHIL_DN | GSE3982_MAST_CELL_VS_BASOPHIL_DN | 6/95 | 199/21355 | 0.0002652 | 0.02713021 | 0.0254369 | 51466/4763/4591/60468/6777/5148 | 6 |
| GSE6092_UNSTIM_VS_IFNG_STIM_ENDOTHELIAL_CELL_DN | GSE6092_UNSTIM_VS_IFNG_STIM_ENDOTHELIAL_CELL_DN | 6/95 | 199/21355 | 0.0002652 | 0.02713021 | 0.0254369 | 55671/9742/4542/5116/60468/23112 | 6 |
| GSE12392_WT_VS_IFNB_KO_CD8A_NEG_SPLEEN_DC_DN | GSE12392_WT_VS_IFNB_KO_CD8A_NEG_SPLEEN_DC_DN | 6/95 | 200/21355 | 0.0002724 | 0.02713021 | 0.0254369 | 221037/9742/4542/65125/85464/64766 | 6 |
| GSE17301_ACD3_ACD28_VS_ACD3_ACD28_AND_IFNA2_STIM_CD8_TCELL_DN | GSE17301_ACD3_ACD28_VS_ACD3_ACD28_AND_IFNA2_STIM_CD8_TCELL_DN | 6/95 | 200/21355 | 0.0002724 | 0.02713021 | 0.0254369 | 4753/55114/1387/80345/10807/10260 | 6 |
| GSE17301_IFNA2_VS_IFNA2_AND_ACD3_ACD28_STIM_CD8_TCELL_UP | GSE17301_IFNA2_VS_IFNA2_AND_ACD3_ACD28_STIM_CD8_TCELL_UP | 6/95 | 200/21355 | 0.0002724 | 0.02713021 | 0.0254369 | 4753/55114/1387/80345/10807/10260 | 6 |
| GSE17721_CTRL_VS_GARDIQUIMOD_0.5H_BMDC_UP | GSE17721_CTRL_VS_GARDIQUIMOD_0.5H_BMDC_UP | 6/95 | 200/21355 | 0.0002724 | 0.02713021 | 0.0254369 | 143/10672/56261/2869/28977/4670 | 6 |
| GSE20715_0H_VS_24H_OZONE_LUNG_UP | GSE20715_0H_VS_24H_OZONE_LUNG_UP | 6/95 | 200/21355 | 0.0002724 | 0.02713021 | 0.0254369 | 55690/9202/51466/10743/4542/60468 | 6 |
| GSE25123_CTRL_VS_IL4_AND_ROSIGLITAZONE_STIM_MACROPHAGE_UP | GSE25123_CTRL_VS_IL4_AND_ROSIGLITAZONE_STIM_MACROPHAGE_UP | 6/95 | 200/21355 | 0.0002724 | 0.02713021 | 0.0254369 | 23085/51466/55671/4763/2885/113791 | 6 |
| GSE27786_LIN_NEG_VS_ERYTHROBLAST_UP | GSE27786_LIN_NEG_VS_ERYTHROBLAST_UP | 6/95 | 200/21355 | 0.0002724 | 0.02713021 | 0.0254369 | 221037/27340/10743/51552/23167/10390 | 6 |
| GSE8921_3H_VS_24H_TLR1_2_STIM_MONOCYTE_UP | GSE8921_3H_VS_24H_TLR1_2_STIM_MONOCYTE_UP | 6/95 | 200/21355 | 0.0002724 | 0.02713021 | 0.0254369 | 51466/9742/10743/64766/10390/64324 | 6 |
| NAKAYA_B_CELL_FLUARIX_FLUVIRIN_AGE_18_50YO_7DY_DN | NAKAYA_B_CELL_FLUARIX_FLUVIRIN_AGE_18_50YO_7DY_DN | 7/95 | 294/21355 | 0.0003431 | 0.03322241 | 0.0311488 | 4255/64421/23013/4763/63893/5116/60468 | 7 |
