## Appendix Table S5 for "Distinguishable topological properties of functional genome networks in HIV-1 reservoirs"

**Supplementary Table 5. A complete attribute list of enriched immunologic signatures harboring intact and defective proviruses in ART-treated patients and elite controllers**

| Description | Rich_factor | tpm | provirus_patient | provirus | patient | is_other_cell | is_CD4_T_cell | is_CD8_T_cell | is_B_cell | is_myeloid_cell | is_proinflammatory_facotr | Response | Count | atac_seq_count | HiC | id |
|---|---|---|---|---|---|---|---|---|---|---|---|---|---|---|---|---|
| GSE10240_IL22_VS_IL17_STIM_PRIMARY_BRONCHIAL_EPITHELIAL_CELLS_UP | 5.20853659 | 4.75808 | ART_intact | intact | ART | 1 | 0 | 0 | 0 | 0 | 1 | 2 | 8 | 352 | 19.909216 | 1 |
| GSE11057_CD4_CENT_MEM_VS_PBMC_UP | 5.9488362 | 14.30527 | ART_intact | intact | ART | 0 | 1 | 0 | 0 | 1 | 0 | 2 | 9 | 1076 | 39.824854 | 2 |
| GSE11057_CD4_EFF_MEM_VS_PBMC_UP | 5.23471014 | 10.74065 | ART_intact | intact | ART | 0 | 1 | 0 | 0 | 1 | 0 | 2 | 8 | 850 | 38.697303 | 3 |
| GSE11057_NAIVE_CD4_VS_PBMC_CD4_TCELL_UP | 5.9488362 | 17.06154 | ART_intact | intact | ART | 0 | 1 | 0 | 0 | 1 | 0 | 2 | 9 | 807 | 37.46477 | 4 |
| GSE1460_DP_VS_CD4_THYMOCYTE_DN | 5.20853659 | 14.02425 | ART_intact | intact | ART | 0 | 1 | 0 | 0 | 0 | 0 | 1 | 8 | 666 | 53.536364 | 5 |
| GSE16450_CTRL_VS_IFNA_12H_STIM_IMMATURE_NEURON_CELL_LINE_UP | 5.2878544 | 15.66052 | ART_intact | intact | ART | 1 | 0 | 0 | 0 | 0 | 1 | 2 | 8 | 928 | 33.645995 | 6 |
| GSE17974_1H_VS_72H_UNTREATED_IN_VITRO_CD4_TCELL_DN | 5.23471014 | 6.482483 | ART_intact | intact | ART | 0 | 1 | 0 | 0 | 0 | 0 | 1 | 8 | 587 | 53.109015 | 7 |
| GSE21927_BALBC_VS_C57BL6_MONOCYTE_SPLEEN_UP | 6.20063879 | 40.24171 | ART_intact | intact | ART | 0 | 0 | 0 | 0 | 1 | 0 | 2 | 9 | 437 | 15.517176 | 8 |
| GSE21927_C26GM_VS_4T1_TUMOR_MONOCYTE_BALBC_DN | 5.23471014 | 9.461119 | ART_intact | intact | ART | 0 | 0 | 0 | 0 | 1 | 0 | 1 | 8 | 826 | 24.369683 | 9 |
| GSE22601_IMMATURE_CD4_SINGLE_POSITIVE_VS_CD8_SINGLE_POSITIVE_THYMOCYTE_DN | 5.20853659 | 12.72376 | ART_intact | intact | ART | 0 | 1 | 1 | 0 | 0 | 0 | 1 | 8 | 488 | 35.162861 | 10 |
| GSE25087_TREG_VS_TCONV_FETUS_UP | 5.26114807 | 32.23903 | ART_intact | intact | ART | 1 | 0 | 0 | 0 | 0 | 0 | 2 | 8 | 882 | 41.119183 | 11 |
| GSE2770_IL4_ACT_VS_ACT_CD4_TCELL_2H_DN | 6.51067073 | 21.3547 | ART_intact | intact | ART | 0 | 1 | 0 | 0 | 0 | 1 | 1 | 10 | 646 | 30.034444 | 12 |
| GSE2935_UV_INACTIVATED_VS_LIVE_SENDAI_VIRUS_INF_MACROPHAGE_DN | 5.81959395 | 54.46699 | ART_intact | intact | ART | 0 | 0 | 0 | 0 | 1 | 1 | 1 | 8 | 583 | 31.160633 | 13 |
| GSE32255_UNSTIM_VS_4H_LPS_STIM_DC_UP | 5.60057697 | 12.62257 | ART_intact | intact | ART | 0 | 0 | 0 | 0 | 1 | 0 | 2 | 8 | 957 | 35.203061 | 14 |
| GSE3982_MAST_CELL_VS_BCELL_DN | 5.9488362 | 12.54422 | ART_intact | intact | ART | 0 | 0 | 0 | 1 | 1 | 0 | 1 | 9 | 983 | 33.316794 | 15 |
| GSE40225_WT_VS_RIP_B7X_DIABETIC_MOUSE_PANCREATIC_CD8_TCELL_UP | 6.54338767 | 13.50235 | ART_intact | intact | ART | 0 | 0 | 1 | 0 | 0 | 0 | 2 | 10 | 813 | 20.669456 | 16 |
| GSE5099_DAY3_VS_DAY7_MCSF_TREATED_MACROPHAGE_DN | 7.11548714 | 668.6613 | ART_intact | intact | ART | 1 | 0 | 0 | 0 | 1 | 0 | 1 | 10 | 570 | 28.215607 | 17 |
| GSE5589_WT_VS_IL6_KO_LPS_AND_IL10_STIM_MACROPHAGE_45MIN_UP | 5.20853659 | 16.91804 | ART_intact | intact | ART | 0 | 0 | 0 | 0 | 1 | 1 | 2 | 8 | 434 | 30.31491 | 18 |
| NAKAYA_PBMC_FLUARIX_FLUVIRIN_AGE_18_50YO_CORRELATED_WITH_HAI_28DY_RESPONSE_AT_3DY_NEGATIVE | 5.72366658 | 322.9337 | ART_intact | intact | ART | 0 | 0 | 0 | 0 | 1 | 0 | 0 | 16 | 1575 | 28.449661 | 19 |
| NAKAYA_PBMC_FLUARIX_FLUVIRIN_AGE_18_50YO_CORRELATED_WITH_HAI_28DY_RESPONSE_AT_7DY_NEGATIVE | 4.66026958 | 333.6429 | ART_intact | intact | ART | 0 | 0 | 0 | 0 | 1 | 0 | 0 | 17 | 1584 | 27.613844 | 20 |
| ANDERSON_BLOOD_CN54GP140_ADJUVANTED_WITH_GLA_AF_AGE_18_45YO_1DY_DN | 5.75661885 | 28.43847 | ART_defective | defective | ART | 0 | 0 | 0 | 0 | 0 | 0 | 1 | 14 | 943 | 28.492289 | 21 |
| GSE10094_LCMV_VS_LISTERIA_IND_EFF_CD4_TCELL_UP | 3.16138795 | 22.23804 | ART_defective | defective | ART | 0 | 1 | 0 | 0 | 0 | 0 | 2 | 18 | 1571 | 31.695706 | 22 |
| GSE10147_IL3_VS_IL3_AND_HIVP17_STIM_PDC_UP | 3.36876146 | 7.854994 | ART_defective | defective | ART | 0 | 0 | 0 | 0 | 1 | 1 | 2 | 16 | 1453 | 28.436496 | 23 |
| GSE10239_MEMORY_VS_KLRG1HIGH_EFF_CD8_TCELL_DN | 3.14558101 | 17.60299 | ART_defective | defective | ART | 0 | 0 | 1 | 0 | 0 | 0 | 1 | 18 | 1461 | 36.183725 | 24 |
| GSE10273_HIGH_VS_LOW_IL7_TREATED_IRF4_8_NULL_PRE_BCELL_DN | 3.35387426 | 28.81122 | ART_defective | defective | ART | 0 | 0 | 0 | 1 | 0 | 1 | 1 | 19 | 996 | 36.231608 | 25 |
| GSE10273_LOW_IL7_VS_HIGH_IL7_AND_IRF4_IN_IRF4_8_NULL_PRE_BCELL_DN | 2.97082651 | 12.35469 | ART_defective | defective | ART | 0 | 0 | 0 | 1 | 0 | 1 | 1 | 17 | 1083 | 28.768632 | 26 |
| GSE10325_CD4_TCELL_VS_BCELL_UP | 4.05995305 | 15.12604 | ART_defective | defective | ART | 0 | 1 | 0 | 1 | 0 | 0 | 2 | 23 | 2611 | 33.030103 | 27 |
| GSE10325_CD4_TCELL_VS_MYELOID_UP | 3.51265328 | 15.63126 | ART_defective | defective | ART | 0 | 1 | 0 | 0 | 1 | 0 | 2 | 20 | 2215 | 27.579806 | 28 |
| GSE10325_LUPUS_CD4_TCELL_VS_LUPUS_BCELL_UP | 4.3908166 | 14.48539 | ART_defective | defective | ART | 0 | 1 | 0 | 1 | 0 | 0 | 2 | 25 | 2593 | 32.470608 | 29 |
| GSE10325_LUPUS_CD4_TCELL_VS_LUPUS_MYELOID_UP | 3.33702062 | 12.1685 | ART_defective | defective | ART | 0 | 1 | 0 | 0 | 1 | 0 | 2 | 19 | 2035 | 39.051732 | 30 |
| GSE11057_CD4_CENT_MEM_VS_PBMC_UP | 3.90314621 | 25.35924 | ART_defective | defective | ART | 0 | 1 | 0 | 0 | 1 | 0 | 2 | 22 | 2113 | 31.129011 | 31 |
| GSE11057_CD4_EFF_MEM_VS_PBMC_UP | 2.81012263 | 16.49651 | ART_defective | defective | ART | 0 | 0 | 0 | 0 | 1 | 0 | 2 | 16 | 1846 | 37.901833 | 32 |
| GSE11057_EFF_MEM_VS_CENT_MEM_CD4_TCELL_DN | 4.275908 | 13.33656 | ART_defective | defective | ART | 0 | 1 | 0 | 0 | 0 | 0 | 1 | 23 | 2948 | 27.064002 | 33 |
| GSE11057_NAIVE_VS_CENT_MEMORY_CD4_TCELL_DN | 3.33702062 | 9.831558 | ART_defective | defective | ART | 0 | 1 | 0 | 0 | 0 | 0 | 1 | 19 | 1642 | 25.740791 | 34 |
| GSE11057_NAIVE_VS_MEMORY_CD4_TCELL_DN | 2.82431516 | 9.316935 | ART_defective | defective | ART | 0 | 1 | 0 | 0 | 0 | 0 | 1 | 16 | 1518 | 24.567943 | 35 |
| GSE11057_PBMC_VS_MEM_CD4_TCELL_DN | 3.19348326 | 20.886 | ART_defective | defective | ART | 0 | 1 | 0 | 0 | 1 | 0 | 1 | 18 | 1603 | 38.97863 | 36 |
| GSE11386_NAIVE_VS_MEMORY_BCELL_DN | 3.7368887 | 12.76804 | ART_defective | defective | ART | 0 | 0 | 0 | 1 | 0 | 0 | 1 | 17 | 2321 | 42.653313 | 37 |
| GSE11864_UNTREATED_VS_CSF1_PAM3CYS_IN_MAC_UP | 2.82431516 | 9.346312 | ART_defective | defective | ART | 0 | 0 | 0 | 0 | 1 | 0 | 2 | 16 | 1058 | 20.160698 | 38 |
| GSE11924_TFH_VS_TH1_CD4_TCELL_UP | 3.68828595 | 13.31208 | ART_defective | defective | ART | 1 | 1 | 0 | 0 | 0 | 0 | 2 | 21 | 1233 | 39.743023 | 39 |
| GSE11961_MARGINAL_ZONE_BCELL_VS_GERMINAL_CENTER_BCELL_DAY40_UP | 2.81012263 | 19.6714 | ART_defective | defective | ART | 0 | 0 | 0 | 1 | 0 | 0 | 2 | 16 | 1491 | 32.992315 | 40 |
| GSE12003_MIR223_KO_VS_WT_BM_PROGENITOR_4D_CULTURE_UP | 3.14558101 | 14.91895 | ART_defective | defective | ART | 1 | 0 | 0 | 0 | 0 | 0 | 2 | 18 | 1546 | 34.183014 | 41 |
| GSE12366_GC_BCELL_VS_PLASMA_CELL_UP | 3.90314621 | 8.912164 | ART_defective | defective | ART | 0 | 0 | 0 | 1 | 1 | 0 | 2 | 22 | 2277 | 27.649077 | 42 |
| GSE12366_PLASMA_CELL_VS_NAIVE_BCELL_DN | 3.86391861 | 14.58374 | ART_defective | defective | ART | 0 | 0 | 0 | 1 | 1 | 0 | 1 | 22 | 1666 | 28.94659 | 43 |
| GSE12392_CD8A_POS_VS_NEG_SPLEEN_DC_DN | 3.32033552 | 63.64586 | ART_defective | defective | ART | 0 | 0 | 0 | 0 | 1 | 0 | 1 | 19 | 1686 | 29.252079 | 44 |
| GSE12845_IGD_NEG_BLOOD_VS_DARKZONE_GC_TONSIL_BCELL_DN | 2.81012263 | 22.52888 | ART_defective | defective | ART | 0 | 0 | 0 | 1 | 0 | 0 | 1 | 16 | 1529 | 23.347865 | 45 |
| GSE12845_IGD_POS_BLOOD_VS_DARKZONE_GC_TONSIL_BCELL_DN | 3.51265328 | 16.16598 | ART_defective | defective | ART | 0 | 0 | 0 | 1 | 0 | 0 | 1 | 20 | 2244 | 26.406172 | 46 |
| GSE12845_IGD_POS_BLOOD_VS_NAIVE_TONSIL_BCELL_DN | 2.81012263 | 14.06371 | ART_defective | defective | ART | 0 | 0 | 0 | 1 | 0 | 0 | 1 | 16 | 1600 | 33.050096 | 47 |
| GSE13306_LAMINA_PROPRIA_VS_SPLEEN_TREG_UP | 3.33702062 | 27.71816 | ART_defective | defective | ART | 1 | 0 | 0 | 0 | 0 | 0 | 2 | 19 | 1944 | 35.94997 | 48 |
| GSE13306_TREG_VS_TCONV_SPLEEN_DN | 2.81012263 | 25.56522 | ART_defective | defective | ART | 1 | 0 | 0 | 0 | 0 | 0 | 1 | 16 | 1468 | 22.8744 | 49 |
| GSE13411_IGM_MEMORY_BCELL_VS_PLASMA_CELL_UP | 3.70691365 | 14.78428 | ART_defective | defective | ART | 0 | 0 | 0 | 1 | 1 | 1 | 2 | 21 | 1894 | 30.688868 | 50 |
| GSE13411_NAIVE_BCELL_VS_PLASMA_CELL_UP | 3.56641838 | 11.93549 | ART_defective | defective | ART | 0 | 0 | 0 | 1 | 1 | 0 | 2 | 20 | 1827 | 33.32393 | 51 |
| GSE13411_PLASMA_CELL_VS_MEMORY_BCELL_DN | 2.83865179 | 10.37248 | ART_defective | defective | ART | 0 | 0 | 0 | 1 | 1 | 0 | 1 | 16 | 1758 | 38.005175 | 52 |
| GSE13411_SWITCHED_MEMORY_BCELL_VS_PLASMA_CELL_UP | 4.36886252 | 12.06099 | ART_defective | defective | ART | 0 | 0 | 0 | 1 | 1 | 0 | 2 | 25 | 1673 | 26.503731 | 53 |
| GSE13484_UNSTIM_VS_3H_YF17D_VACCINE_STIM_PBMC_UP | 2.98575529 | 15.7742 | ART_defective | defective | ART | 0 | 0 | 0 | 0 | 1 | 0 | 2 | 17 | 1123 | 26.3394 | 54 |
| GSE13485_CTRL_VS_DAY1_YF17D_VACCINE_PBMC_UP | 3.03145563 | 31.68801 | ART_defective | defective | ART | 0 | 0 | 0 | 0 | 1 | 0 | 2 | 17 | 770 | 34.478353 | 55 |
| GSE13547_CTRL_VS_ANTI_IGM_STIM_ZFX_KO_BCELL_12H_DN | 3.02044816 | 9.497584 | ART_defective | defective | ART | 0 | 0 | 0 | 1 | 0 | 1 | 1 | 14 | 1895 | 34.50169 | 56 |
| GSE13738_TCR_VS_BYSTANDER_ACTIVATED_CD4_TCELL_UP | 2.82431516 | 8.264976 | ART_defective | defective | ART | 0 | 1 | 0 | 0 | 0 | 0 | 2 | 16 | 1364 | 31.023371 | 57 |
| GSE14350_TREG_VS_TEFF_UP | 4.21518394 | 11.43312 | ART_defective | defective | ART | 1 | 0 | 0 | 0 | 0 | 0 | 2 | 24 | 1929 | 31.956169 | 58 |
| GSE14415_FOXP3_KO_NATURAL_TREG_VS_TCONV_DN | 3.23245319 | 7.459579 | ART_defective | defective | ART | 1 | 0 | 0 | 0 | 0 | 0 | 1 | 16 | 1274 | 29.162278 | 59 |
| GSE14415_FOXP3_KO_NATURAL_TREG_VS_TCONV_UP | 2.86147721 | 10.86667 | ART_defective | defective | ART | 1 | 0 | 0 | 0 | 0 | 0 | 2 | 14 | 1477 | 28.407186 | 60 |
| GSE14415_INDUCED_TREG_VS_FOXP3_KO_INDUCED_TREG_UP | 3.25124653 | 7.822524 | ART_defective | defective | ART | 1 | 0 | 0 | 0 | 0 | 0 | 2 | 16 | 1488 | 34.683973 | 61 |
| GSE14415_INDUCED_TREG_VS_TCONV_DN | 2.98361343 | 6.541994 | ART_defective | defective | ART | 1 | 0 | 0 | 0 | 0 | 0 | 1 | 14 | 1786 | 35.372909 | 62 |
| GSE1460_INTRATHYMIC_T_PROGENITOR_VS_CD4_THYMOCYTE_DN | 3.16138795 | 56.24284 | ART_defective | defective | ART | 1 | 0 | 0 | 0 | 0 | 0 | 1 | 18 | 1675 | 37.499091 | 63 |
| GSE14699_NAIVE_VS_DELETIONAL_TOLERANCE_CD8_TCELL_UP | 3.67904212 | 14.82318 | ART_defective | defective | ART | 0 | 0 | 1 | 0 | 0 | 0 | 2 | 16 | 1238 | 22.863078 | 64 |
| GSE15330_HSC_VS_MEGAKARYOCYTE_ERYTHROID_PROGENITOR_DN | 3.95278038 | 8.068594 | ART_defective | defective | ART | 1 | 0 | 0 | 0 | 0 | 0 | 1 | 19 | 1311 | 28.566937 | 65 |
| GSE15330_HSC_VS_MEGAKARYOCYTE_ERYTHROID_PROGENITOR_UP | 3.88343335 | 7.81912 | ART_defective | defective | ART | 1 | 0 | 0 | 0 | 0 | 0 | 2 | 18 | 1644 | 24.378157 | 66 |
| GSE15330_LYMPHOID_MULTIPOTENT_VS_MEGAKARYOCYTE_ERYTHROID_PROGENITOR_IKAROS_KO_UP | 3.57930905 | 5.696579 | ART_defective | defective | ART | 1 | 0 | 0 | 0 | 0 | 0 | 2 | 17 | 1402 | 30.453318 | 67 |
| GSE15624_CTRL_VS_3H_HALOFUGINONE_TREATED_CD4_TCELL_UP | 4.80574877 | 17.1876 | ART_defective | defective | ART | 0 | 1 | 0 | 0 | 0 | 0 | 2 | 22 | 1623 | 24.738501 | 68 |
| GSE15733_BM_VS_SPLEEN_MEMORY_CD4_TCELL_DN | 2.81012263 | 16.23005 | ART_defective | defective | ART | 0 | 1 | 0 | 0 | 0 | 0 | 1 | 16 | 1755 | 36.793828 | 69 |

| | | | | | | | | | | | | | | | | |
|---|---|---|---|---|---|---|---|---|---|---|---|---|---|---|---|---|
| GSE16385_IL4_VS_ROSIGLITAZONE_STIM_MACROPHAGE_UP | 3.24286703 | 11.67789 | ART_defective | defective | ART | 0 | 0 | 0 | 0 | 1 | 1 | 2 | 18 | 1831 | 27.535275 | 70 |
| GSE16385_ROSIGLITAZONE_IL4_VS_IFNG_TNF_STIM_MACROPHAGE_UP | 3.49509002 | 14.54812 | ART_defective | defective | ART | 0 | 0 | 0 | 0 | 1 | 1 | 2 | 20 | 2253 | 37.092846 | 71 |
| GSE16450_CTRL_VS_IFNA_12H_STIM_IMMATURE_NEURON_CELL_LINE_UP | 3.90314621 | 10.95159 | ART_defective | defective | ART | 1 | 0 | 0 | 0 | 0 | 1 | 2 | 22 | 2470 | 40.43235 | 72 |
| GSE16450_CTRL_VS_IFNA_6H_STIM_IMMATURE_NEURON_CELL_LINE_DN | 3.22623694 | 19.31116 | ART_defective | defective | ART | 1 | 0 | 0 | 0 | 0 | 1 | 1 | 18 | 1467 | 32.452026 | 73 |
| GSE16450_CTRL_VS_IFNA_6H_STIM_MATURE_NEURON_CELL_LINE_DN | 2.97082651 | 341.7537 | ART_defective | defective | ART | 1 | 0 | 0 | 0 | 0 | 1 | 1 | 17 | 2029 | 36.191652 | 74 |
| GSE16450_IMMATURE_VS_MATURE_NEURON_CELL_LINE_UP | 4.30164925 | 251.1714 | ART_defective | defective | ART | 1 | 0 | 0 | 0 | 0 | 0 | 2 | 24 | 2380 | 36.484094 | 75 |
| GSE16522_ANTI_CD3CD28_STIM_VS_UNSTIM_NAIVE_CD8_TCELL_DN | 3.70691365 | 30.43462 | ART_defective | defective | ART | 0 | 0 | 1 | 0 | 0 | 0 | 1 | 21 | 1332 | 35.166012 | 76 |
| GSE17186_BLOOD_VS_CORD_BLOOD_CD21LOW_TRANSITIONAL_BCELL_UP | 2.97082651 | 19.56537 | ART_defective | defective | ART | 0 | 0 | 0 | 1 | 0 | 0 | 2 | 17 | 1741 | 38.922078 | 77 |
| GSE17301_ACD3_ACD28_VS_ACD3_ACD28_AND_IFNA2_STIM_CD8_TCELL_DN | 2.97082651 | 17.18927 | ART_defective | defective | ART | 0 | 0 | 1 | 0 | 0 | 1 | 1 | 17 | 1451 | 40.362761 | 78 |
| GSE17301_CTRL_VS_48H_ACD3_ACD28_IFNA5_STIM_CD8_TCELL_UP | 3.32033552 | 10.00725 | ART_defective | defective | ART | 0 | 0 | 1 | 0 | 0 | 1 | 2 | 19 | 2182 | 29.341386 | 79 |
| GSE17721_0.5H_VS_12H_GARDIQUIMOD_BMDC_DN | 3.14558101 | 16.32292 | ART_defective | defective | ART | 0 | 0 | 0 | 0 | 1 | 0 | 1 | 18 | 1560 | 29.12642 | 80 |
| GSE17721_0.5H_VS_8H_GARDIQUIMOD_BMDC_DN | 2.98575529 | 79.01212 | ART_defective | defective | ART | 0 | 0 | 0 | 0 | 1 | 0 | 1 | 17 | 1506 | 32.007368 | 81 |
| GSE17721_CTRL_VS_CPG_0.5H_BMDC_UP | 3.16138795 | 14.4053 | ART_defective | defective | ART | 0 | 0 | 0 | 0 | 1 | 0 | 2 | 18 | 1178 | 23.943437 | 82 |
| GSE17721_CTRL_VS_GARDIQUIMOD_0.5H_BMDC_UP | 2.97082651 | 20.61186 | ART_defective | defective | ART | 0 | 0 | 0 | 0 | 1 | 0 | 2 | 17 | 888 | 33.100537 | 83 |
| GSE17721_PAM3CSK4_VS_GADIQUIMOD_8H_BMDC_DN | 3.14558101 | 18.91739 | ART_defective | defective | ART | 0 | 0 | 0 | 0 | 1 | 0 | 1 | 18 | 1563 | 34.727325 | 84 |
| GSE17974_0.5H_VS_72H_IL4_AND_ANTI_IL12_ACT_CD4_TCELL_DN | 3.03145563 | 13.52337 | ART_defective | defective | ART | 0 | 1 | 0 | 0 | 0 | 1 | 1 | 17 | 1108 | 22.59851 | 85 |
| GSE17974_0.5H_VS_72H_IL4_AND_ANTI_IL12_ACT_CD4_TCELL_UP | 2.85313471 | 14.2427 | ART_defective | defective | ART | 0 | 1 | 0 | 0 | 0 | 1 | 2 | 16 | 1259 | 19.819466 | 86 |
| GSE17974_CTRL_VS_ACT_IL4_AND_ANTI_IL12_24H_CD4_TCELL_UP | 2.86776617 | 9.558902 | ART_defective | defective | ART | 0 | 1 | 0 | 0 | 0 | 1 | 2 | 16 | 1969 | 34.591498 | 87 |
| GSE17974_IL4_AND_ANTI_IL12_VS_UNTREATED_72H_ACT_CD4_TCELL_DN | 3.04700155 | 13.5166 | ART_defective | defective | ART | 0 | 1 | 0 | 0 | 0 | 1 | 1 | 17 | 1357 | 33.702208 | 88 |
| GSE18893_TCONV_VS_TREG_2H_CULTURE_DN | 2.97082651 | 10.61307 | ART_defective | defective | ART | 1 | 0 | 0 | 0 | 0 | 0 | 1 | 17 | 1243 | 30.490758 | 89 |
| GSE18893_TCONV_VS_TREG_2H_TNF_STIM_DN | 3.06270775 | 74.11514 | ART_defective | defective | ART | 1 | 0 | 0 | 0 | 0 | 1 | 1 | 17 | 850 | 20.413411 | 90 |
| GSE1925_CTRL_VS_IFNG_PRIMED_MACROPHAGE_3H_IFNG_STIM_UP | 3.8164776 | 13.1091 | ART_defective | defective | ART | 0 | 0 | 0 | 0 | 1 | 1 | 1 | 19 | 1256 | 28.886156 | 91 |
| GSE21033_1H_VS_12H_POLYIC_STIM_DC_DN | 3.22242342 | 20.24516 | ART_defective | defective | ART | 0 | 0 | 0 | 0 | 1 | 0 | 1 | 13 | 809 | 27.58567 | 92 |
| GSE21063_CTRL_VS_ANTI_IGM_STIM_BCELL_NFATC1_KO_3H_DN | 4.08999896 | 16.65121 | ART_defective | defective | ART | 0 | 0 | 0 | 1 | 0 | 1 | 1 | 22 | 1511 | 28.361498 | 93 |
| GSE21063_CTRL_VS_ANTI_IGM_STIM_BCELL_NFATC1_KO_8H_UP | 3.29380211 | 39.14549 | ART_defective | defective | ART | 0 | 0 | 0 | 1 | 0 | 1 | 2 | 18 | 1520 | 21.831395 | 94 |
| GSE21360_PRIMARY_VS_QUATERNARY_MEMORY_CD8_TCELL_DN | 2.82431516 | 14.36486 | ART_defective | defective | ART | 0 | 0 | 1 | 0 | 0 | 0 | 1 | 16 | 1522 | 31.623746 | 95 |
| GSE21546_ELK1_KO_VS_SAP1A_KO_AND_ELK1_KO_DP_THYMOCYTES_DN | 2.83865179 | 11.6082 | ART_defective | defective | ART | 1 | 0 | 0 | 0 | 0 | 0 | 1 | 16 | 830 | 25.613789 | 96 |
| GSE21546_UNSTIM_VS_ANTI_CD3_STIM_DP_THYMOCYTES_DN | 2.81012263 | 9.602695 | ART_defective | defective | ART | 1 | 0 | 0 | 0 | 0 | 0 | 1 | 16 | 1097 | 31.789474 | 97 |
| GSE21546_WT_VS_SAP1A_KO_AND_ELK1_KO_DP_THYMOCYTES_DN | 3.32033552 | 9.202481 | ART_defective | defective | ART | 1 | 0 | 0 | 0 | 0 | 0 | 1 | 19 | 1990 | 36.285202 | 98 |
| GSE21670_IL6_VS_TGFB_AND_IL6_TREATED_CD4_TCELL_UP | 3.0785767 | 13.68988 | ART_defective | defective | ART | 0 | 1 | 0 | 0 | 0 | 1 | 2 | 17 | 1983 | 35.847785 | 99 |
| GSE21670_UNTREATED_VS_IL6_TREATED_CD4_TCELL_DN | 3.66984452 | 26.02101 | ART_defective | defective | ART | 0 | 1 | 0 | 0 | 0 | 1 | 1 | 21 | 2387 | 34.838677 | 100 |
| GSE21670_UNTREATED_VS_TGFB_TREATED_STAT3_KO_CD4_TCELL_DN | 3.33702062 | 13.89168 | ART_defective | defective | ART | 0 | 1 | 0 | 0 | 0 | 1 | 1 | 19 | 1406 | 23.978593 | 101 |
| GSE21927_C26GM_VS_4T1_TUMOR_MONOCYTE_BALBC_DN | 4.9177146 | 29.55466 | ART_defective | defective | ART | 0 | 0 | 0 | 0 | 1 | 0 | 1 | 28 | 2354 | 33.73694 | 102 |
| GSE21927_SPLEEN_C57BL6_VS_EL4_TUMOR_BALBC_MONOCYTES_UP | 3.51265328 | 26.2793 | ART_defective | defective | ART | 0 | 0 | 0 | 1 | 0 | 0 | 2 | 20 | 1565 | 37.204954 | 103 |
| GSE21927_SPLENIC_VS_TUMOR_MONOCYTES_FROM_C26GM_TUMOROUS_MICE_BALBC_DN | 3.35387426 | 30.5253 | ART_defective | defective | ART | 0 | 0 | 0 | 0 | 1 | 0 | 1 | 19 | 1367 | 22.221656 | 104 |
| GSE22342_CD11C_HIGH_VS_LOW_DECIDUAL_MACROPHAGES_UP | 3.00083486 | 29.47789 | ART_defective | defective | ART | 0 | 0 | 0 | 0 | 1 | 0 | 2 | 17 | 1197 | 29.921093 | 105 |
| GSE22432_CONVENTIONAL_CDC_VS_PLASMACYTOID_PDC_UP | 2.81012263 | 9.36116 | ART_defective | defective | ART | 0 | 0 | 0 | 0 | 1 | 0 | 2 | 16 | 908 | 38.717635 | 106 |
| GSE22601_CD4_SINGLE_POSITIVE_VS_CD8_SINGLE_POSITIVE_THYMOCYTE_DN | 3.14558101 | 16.65893 | ART_defective | defective | ART | 1 | 0 | 0 | 0 | 0 | 0 | 1 | 18 | 1870 | 31.551767 | 107 |
| GSE22601_IMMATURE_CD4_SINGLE_POSITIVE_VS_DOUBLE_POSITIVE_THYMOCYTE_DN | 3.32033552 | 14.34829 | ART_defective | defective | ART | 0 | 1 | 0 | 0 | 0 | 0 | 1 | 19 | 2646 | 30.031113 | 108 |
| GSE22886_CD4_TCELL_VS_BCELL_NAIVE_UP | 3.17735456 | 25.96089 | ART_defective | defective | ART | 0 | 1 | 0 | 1 | 0 | 0 | 2 | 18 | 2327 | 28.945455 | 109 |
| GSE22886_CD8_TCELL_VS_BCELL_NAIVE_UP | 3.17735456 | 23.86332 | ART_defective | defective | ART | 0 | 0 | 1 | 1 | 0 | 0 | 2 | 18 | 1929 | 38.470259 | 110 |
| GSE22886_NAIVE_CD4_TCELL_VS_12H_ACT_TH2_UP | 2.98575529 | 33.27294 | ART_defective | defective | ART | 1 | 1 | 0 | 0 | 0 | 0 | 2 | 17 | 1228 | 32.521862 | 111 |
| GSE22886_NAIVE_CD4_TCELL_VS_MONOCYTE_UP | 3.19348326 | 13.63014 | ART_defective | defective | ART | 0 | 1 | 0 | 0 | 1 | 0 | 2 | 18 | 2020 | 31.234902 | 112 |
| GSE22886_NAIVE_CD8_TCELL_VS_DC_UP | 3.16138795 | 19.85809 | ART_defective | defective | ART | 0 | 0 | 1 | 0 | 1 | 0 | 2 | 18 | 1821 | 31.998995 | 113 |
| GSE22886_NAIVE_CD8_TCELL_VS_MEMORY_TCELL_DN | 2.98575529 | 13.799 | ART_defective | defective | ART | 1 | 0 | 1 | 0 | 0 | 0 | 1 | 17 | 1559 | 32.689934 | 114 |
| GSE22886_NAIVE_CD8_TCELL_VS_MONOCYTE_UP | 3.33702062 | 12.70896 | ART_defective | defective | ART | 0 | 0 | 1 | 0 | 1 | 0 | 2 | 19 | 2212 | 34.687608 | 115 |
| GSE22886_NAIVE_CD8_TCELL_VS_NKCELL_UP | 3.00083486 | 23.31347 | ART_defective | defective | ART | 1 | 0 | 1 | 0 | 0 | 0 | 2 | 17 | 1376 | 45.006024 | 116 |
| GSE22886_NAIVE_TCELL_VS_DC_UP | 3.33702062 | 24.78722 | ART_defective | defective | ART | 1 | 0 | 0 | 0 | 1 | 0 | 2 | 19 | 1420 | 33.257699 | 117 |
| GSE22886_NAIVE_TCELL_VS_MONOCYTE_UP | 3.17735456 | 13.01697 | ART_defective | defective | ART | 1 | 0 | 0 | 0 | 1 | 0 | 2 | 18 | 1915 | 31.819125 | 118 |
| GSE22886_NAIVE_TCELL_VS_NKCELL_UP | 2.82431516 | 28.58999 | ART_defective | defective | ART | 1 | 0 | 0 | 0 | 0 | 0 | 2 | 16 | 1439 | 43.552916 | 119 |
| GSE22886_NAIVE_VS_MEMORY_TCELL_DN | 2.98575529 | 26.77954 | ART_defective | defective | ART | 1 | 0 | 0 | 0 | 0 | 0 | 1 | 17 | 1700 | 34.562712 | 120 |
| GSE22886_TCELL_VS_BCELL_NAIVE_UP | 3.16138795 | 26.57056 | ART_defective | defective | ART | 1 | 0 | 0 | 1 | 0 | 0 | 2 | 18 | 2214 | 25.359101 | 121 |
| GSE22886_UNSTIM_VS_STIM_MEMORY_TCELL_UP | 2.82431516 | 25.23225 | ART_defective | defective | ART | 1 | 0 | 0 | 0 | 0 | 0 | 2 | 16 | 1400 | 28.834339 | 122 |
| GSE23321_CD8_STEM_CELL_MEMORY_VS_NAIVE_CD8_TCELL_DN | 3.56641838 | 18.86077 | ART_defective | defective | ART | 1 | 0 | 1 | 0 | 0 | 0 | 1 | 20 | 1729 | 30.044201 | 123 |
| GSE24026_PD1_LIGATION_VS_CTRL_IN_ACT_TCELL_LINE_DN | 2.97082651 | 14.96548 | ART_defective | defective | ART | 1 | 0 | 0 | 0 | 0 | 0 | 1 | 17 | 1098 | 23.126908 | 124 |
| GSE24210_CTRL_VS_IL35_TREATED_TCONV_CD4_TCELL_UP | 3.66984452 | 15.55299 | ART_defective | defective | ART | 0 | 1 | 0 | 0 | 0 | 1 | 2 | 21 | 1795 | 38.230458 | 125 |
| GSE24574_BCL6_HIGH_VS_LOW_TFH_CD4_TCELL_UP | 3.51265328 | 20.7617 | ART_defective | defective | ART | 0 | 1 | 0 | 0 | 0 | 0 | 2 | 20 | 2104 | 33.845655 | 126 |
| GSE24671_CTRL_VS_BAKIMULC_INFECTED_MOUSE_SPLENOCYTES_UP | 3.14558101 | 11.46958 | ART_defective | defective | ART | 1 | 0 | 0 | 0 | 0 | 0 | 2 | 18 | 1754 | 26.497541 | 127 |
| GSE24671_CTRL_VS_SENDAI_VIRUS_INFECTED_MOUSE_SPLENOCYTES_DN | 2.81012263 | 25.06864 | ART_defective | defective | ART | 1 | 0 | 0 | 0 | 0 | 0 | 1 | 16 | 1713 | 41.754634 | 128 |
| GSE24726_WT_VS_E2_2_KO_PDC_DAY6_POST_DELETION_DN | 2.97082651 | 327.5355 | ART_defective | defective | ART | 0 | 0 | 0 | 0 | 1 | 0 | 1 | 17 | 1719 | 35.29061 | 129 |
| GSE25087_TREG_VS_TCONV_ADULT_UP | 3.68828595 | 13.80642 | ART_defective | defective | ART | 1 | 0 | 0 | 0 | 0 | 0 | 2 | 21 | 1728 | 31.777091 | 130 |
| GSE25087_TREG_VS_TCONV_FETUS_UP | 3.00083486 | 19.49264 | ART_defective | defective | ART | 1 | 0 | 0 | 0 | 0 | 0 | 2 | 17 | 1579 | 27.338784 | 131 |
| GSE25088_CTRL_VS_IL4_AND_ROSIGLITAZONE_STIM_STAT6_KO_MACROPHAGE_UP | 3.54831474 | 18.9337 | ART_defective | defective | ART | 1 | 0 | 0 | 0 | 1 | 1 | 2 | 20 | 1496 | 23.947052 | 132 |
| GSE25088_IL4_VS_IL4_AND_ROSIGLITAZONE_STIM_STAT6_KO_MACROPHAGE_DAY10_UP | 2.82431516 | 13.12151 | ART_defective | defective | ART | 0 | 0 | 0 | 0 | 1 | 1 | 2 | 16 | 1963 | 35.709727 | 133 |
| GSE25146_UNSTIM_VS_HELIOBACTER_PYLORI_LPS_STIM_AGS_CELL_DN | 3.78985664 | 46.49262 | ART_defective | defective | ART | 1 | 0 | 0 | 0 | 0 | 0 | 1 | 18 | 1611 | 32.785031 | 134 |
| GSE25677_MPL_VS_MPL_AND_R848_STIM_BCELL_UP | 3.05820376 | 18.25125 | ART_defective | defective | ART | 0 | 0 | 0 | 1 | 0 | 0 | 2 | 14 | 1040 | 34.308293 | 135 |
| GSE25677_MPL_VS_R848_STIM_BCELL_DN | 4.6343735 | 23.17943 | ART_defective | defective | ART | 0 | 0 | 0 | 1 | 0 | 0 | 1 | 24 | 2394 | 39.412 | 136 |
| GSE26030_TH1_VS_TH17_DAY5_POST_POLARIZATION_DN | 3.01606753 | 24.45985 | ART_defective | defective | ART | 1 | 0 | 0 | 0 | 0 | 0 | 1 | 17 | 1741 | 42.141645 | 137 |
| GSE26030_UNSTIM_VS_RESTIM_TH1_DAY15_POST_POLARIZATION_UP | 2.97082651 | 71.12867 | ART_defective | defective | ART | 1 | 0 | 0 | 0 | 0 | 0 | 2 | 17 | 1639 | 26.075185 | 138 |
| GSE26488_CTRL_VS_PEPTIDE_INJECTION_OT2_THYMOCYTE_DN | 3.81977051 | 18.91523 | ART_defective | defective | ART | 1 | 0 | 0 | 0 | 0 | 0 | 1 | 20 | 2164 | 29.634043 | 139 |
| GSE26495_NAIVE_VS_PD1LOW_CD8_TCELL_DN | 3.33702062 | 19.18785 | ART_defective | defective | ART | 0 | 0 | 1 | 0 | 0 | 0 | 1 | 19 | 1684 | 23.727604 | 140 |
| GSE27241_WT_VS_RORGT_KO_TH17_POLARIZED_CD4_TCELL_TREATED_WITH_DIGOXIN_UP | 5.43680669 | 14.2914 | ART_defective | defective | ART | 0 | 1 | 0 | 0 | 0 | 0 | 2 | 28 | 2999 | 39.434688 | 141 |

| | | | | | | | | | | | | | | | | |
|---|---|---|---|---|---|---|---|---|---|---|---|---|---|---|---|---|
| GSE27291_0H_VS_7D_STIM_GAMMADELTA_TCELL_DN | 3.1271858 | 31.05123 | ART_defective | defective | ART | 1 | 0 | 0 | 0 | 0 | 0 | 1 | 17 | 1480 | 29.188725 | 142 |
| GSE27670_BLIMP1_VS_LMP1_TRANSDUCED_GC_BCELL_DN | 3.51265328 | 20.67179 | ART_defective | defective | ART | 0 | 0 | 0 | 1 | 0 | 0 | 1 | 20 | 1282 | 28.432373 | 143 |
| GSE2770_IL12_AND_TGFB_ACT_VS_ACT_CD4_TCELL_6H_DN | 2.82431516 | 15.37649 | ART_defective | defective | ART | 0 | 1 | 0 | 0 | 0 | 1 | 1 | 16 | 1156 | 29.070046 | 144 |
| GSE2770_IL12_VS_IL4_TREATED_ACT_CD4_TCELL_48H_DN | 3.16138795 | 21.23946 | ART_defective | defective | ART | 0 | 1 | 0 | 0 | 0 | 1 | 1 | 18 | 1184 | 24.071888 | 145 |
| GSE2770_IL12_VS_TGFB_AND_IL12_TREATED_ACT_CD4_TCELL_6H_DN | 3.33702062 | 13.09708 | ART_defective | defective | ART | 0 | 1 | 0 | 0 | 0 | 1 | 1 | 19 | 1155 | 29.651306 | 146 |
| GSE2770_IL4_ACT_VS_ACT_CD4_TCELL_2H_DN | 2.97082651 | 23.47961 | ART_defective | defective | ART | 0 | 1 | 0 | 0 | 0 | 1 | 1 | 17 | 1317 | 35.016383 | 147 |
| GSE2770_IL4_ACT_VS_ACT_CD4_TCELL_6H_UP | 2.82431516 | 22.01318 | ART_defective | defective | ART | 0 | 1 | 0 | 0 | 0 | 1 | 2 | 16 | 1135 | 19.437684 | 148 |
| GSE2770_TGFB_AND_IL4_VS_IL12_TREATED_ACT_CD4_TCELL_6H_UP | 2.97082651 | 21.07763 | ART_defective | defective | ART | 0 | 1 | 0 | 0 | 0 | 1 | 2 | 17 | 1631 | 38.611686 | 149 |
| GSE2770_UNTREATED_VS_IL12_TREATED_ACT_CD4_TCELL_2H_UP | 4.19410802 | 15.44843 | ART_defective | defective | ART | 0 | 1 | 0 | 0 | 0 | 1 | 2 | 24 | 2251 | 43.700565 | 150 |
| GSE2770_UNTREATED_VS_IL4_TREATED_ACT_CD4_TCELL_6H_DN | 2.97082651 | 14.57331 | ART_defective | defective | ART | 0 | 1 | 0 | 0 | 0 | 1 | 1 | 17 | 1607 | 22.727881 | 151 |
| GSE27786_LIN_NEG_VS_CD8_TCELL_DN | 2.97082651 | 350.1904 | ART_defective | defective | ART | 0 | 0 | 1 | 0 | 0 | 0 | 1 | 17 | 1208 | 28.541782 | 152 |
| GSE27896_HDAC6_KO_VS_WT_TREG_UP | 3.57452388 | 16.38767 | ART_defective | defective | ART | 1 | 0 | 0 | 0 | 0 | 0 | 2 | 18 | 1367 | 28.33789 | 153 |
| GSE28726_ACT_CD4_TCELL_VS_ACT_NKTCELL_DN | 2.81012263 | 16.79372 | ART_defective | defective | ART | 1 | 1 | 0 | 0 | 0 | 0 | 1 | 16 | 1498 | 28.299846 | 154 |
| GSE29618_PRE_VS_DAY7_FLU_VACCINE_MDC_DN | 3.17735456 | 9.477385 | ART_defective | defective | ART | 0 | 0 | 0 | 0 | 1 | 0 | 1 | 18 | 1872 | 30.844334 | 155 |
| GSE30083_SP1_VS_SP4_THYMOCYTE_DN | 3.84459902 | 19.6679 | ART_defective | defective | ART | 1 | 0 | 0 | 0 | 0 | 0 | 1 | 22 | 1951 | 31.142857 | 156 |
| GSE3039_CD4_TCELL_VS_NKT_CELL_UP | 3.49509002 | 9.325072 | ART_defective | defective | ART | 1 | 1 | 0 | 0 | 0 | 0 | 2 | 20 | 2073 | 31.069608 | 157 |
| GSE30962_ACUTE_VS_CHRONIC_LCMV_SECONDARY_INF_CD8_TCELL_UP | 3.32033552 | 9.903248 | ART_defective | defective | ART | 0 | 0 | 1 | 0 | 0 | 1 | 2 | 19 | 2715 | 43.130331 | 158 |
| GSE31082_CD4_VS_CD8_SP_THYMOCYTE_UP | 3.14558101 | 23.34712 | ART_defective | defective | ART | 1 | 0 | 0 | 0 | 0 | 0 | 2 | 18 | 1705 | 27.973172 | 159 |
| GSE31082_DN_VS_CD4_SP_THYMOCYTE_DN | 2.97082651 | 13.96759 | ART_defective | defective | ART | 1 | 0 | 0 | 0 | 0 | 0 | 1 | 17 | 1625 | 38.354113 | 160 |
| GSE31082_DN_VS_CD4_SP_THYMOCYTE_UP | 3.14558101 | 10.43666 | ART_defective | defective | ART | 1 | 0 | 0 | 0 | 0 | 0 | 2 | 18 | 916 | 27.287129 | 161 |
| GSE31082_DN_VS_CD8_SP_THYMOCYTE_DN | 3.32033552 | 17.29174 | ART_defective | defective | ART | 1 | 0 | 0 | 0 | 0 | 0 | 1 | 19 | 2103 | 43.915625 | 162 |
| GSE32255_WT_UNSTIM_VS_JMJD2D_KNOCKDOWN_4H_LPS_STIM_DC_UP | 3.3952303 | 15.20799 | ART_defective | defective | ART | 0 | 0 | 0 | 0 | 1 | 0 | 2 | 17 | 1554 | 33.223008 | 163 |
| GSE32533_WT_VS_MIR17_KO_ACT_CD4_TCELL_DN | 4.08056195 | 35.94272 | ART_defective | defective | ART | 0 | 1 | 0 | 0 | 0 | 0 | 1 | 23 | 1671 | 30.840332 | 164 |
| GSE32901_NAIVE_VS_TH17_ENRICHED_CD4_TCELL_UP | 2.95040066 | 414.7625 | ART_defective | defective | ART | 0 | 1 | 0 | 0 | 0 | 0 | 2 | 13 | 1602 | 33.291966 | 165 |
| GSE32901_TH1_VS_TH17_ENRICHED_CD4_TCELL_UP | 4.41907933 | 17.4117 | ART_defective | defective | ART | 0 | 1 | 0 | 0 | 0 | 0 | 2 | 22 | 2004 | 35.038062 | 166 |
| GSE32986_GMCSF_AND_CURDLAN_LOWDOSE_VS_GMCSF_AND_CURDLAN_HIGHDOSE_STIM_DC_UP | 3.16138795 | 8.642894 | ART_defective | defective | ART | 0 | 0 | 0 | 0 | 1 | 0 | 2 | 18 | 1625 | 34.966057 | 167 |
| GSE32986_GMCSF_VS_GMCSF_AND_CURDLAN_HIGHDOSE_STIM_DC_DN | 2.82431516 | 10.15659 | ART_defective | defective | ART | 0 | 0 | 0 | 0 | 1 | 0 | 1 | 16 | 1547 | 27.347607 | 168 |
| GSE33162_HDAC3_KO_VS_HDAC3_KO_4H_LPS_STIM_MACROPHAGE_UP | 3.14558101 | 22.9743 | ART_defective | defective | ART | 0 | 0 | 0 | 0 | 1 | 0 | 2 | 18 | 1320 | 30.233927 | 169 |
| GSE33425_CD161_HIGH_VS_INT_CD8_TCELL_UP | 2.97082651 | 10.53701 | ART_defective | defective | ART | 0 | 0 | 1 | 0 | 0 | 0 | 2 | 17 | 1379 | 20.683682 | 170 |
| GSE339_CD4POS_VS_CD4CD8DN_DC_UP | 2.97082651 | 22.6127 | ART_defective | defective | ART | 0 | 0 | 0 | 0 | 1 | 0 | 2 | 17 | 1165 | 19.740473 | 171 |
| GSE35435_RESTING_VS_IL4_TREATED_MACROPHAGE_UP | 3.16138795 | 19.73322 | ART_defective | defective | ART | 0 | 0 | 0 | 0 | 1 | 1 | 2 | 18 | 1200 | 22.255155 | 172 |
| GSE35685_CD34POS_CD38NEG_VS_CD34POS_CD10NEG_CD62LPOS_BONE_MARROW_UP | 3.66984452 | 16.65959 | ART_defective | defective | ART | 0 | 0 | 0 | 0 | 1 | 0 | 2 | 21 | 1593 | 37.311398 | 173 |
| GSE35825_IFNA_VS_IFNG_STIM_MACROPHAGE_DN | 3.49509002 | 14.81353 | ART_defective | defective | ART | 0 | 0 | 0 | 0 | 1 | 1 | 1 | 20 | 1585 | 25.661648 | 174 |
| GSE36476_CTRL_VS_TSST_ACT_16H_MEMORY_CD4_TCELL_OLD_UP | 2.82431516 | 15.61697 | ART_defective | defective | ART | 0 | 1 | 0 | 0 | 0 | 0 | 2 | 16 | 1556 | 25.895281 | 175 |
| GSE36476_CTRL_VS_TSST_ACT_16H_MEMORY_CD4_TCELL_YOUNG_UP | 3.70691365 | 35.70956 | ART_defective | defective | ART | 0 | 1 | 0 | 0 | 0 | 0 | 2 | 21 | 1943 | 25.686914 | 176 |
| GSE36476_CTRL_VS_TSST_ACT_40H_MEMORY_CD4_TCELL_YOUNG_UP | 2.82431516 | 19.44319 | ART_defective | defective | ART | 0 | 1 | 0 | 0 | 0 | 0 | 2 | 16 | 1831 | 38.312119 | 177 |
| GSE36476_YOUNG_VS_OLD_DONOR_MEMORY_CD4_TCELL_40H_TSST_ACT_DN | 2.83865179 | 10.76974 | ART_defective | defective | ART | 0 | 1 | 0 | 0 | 0 | 0 | 1 | 16 | 1390 | 42.75265 | 178 |
| GSE369_PRE_VS_POST_IL6_INJECTION_IFNG_WT_LIVER_DN | 2.97082651 | 12.12983 | ART_defective | defective | ART | 0 | 0 | 0 | 0 | 0 | 1 | 1 | 17 | 1608 | 31.04828 | 179 |
| GSE37301_COMMON_LYMPHOID_PROGENITOR_VS_CD4_TCELL_DN | 3.16138795 | 21.60245 | ART_defective | defective | ART | 1 | 1 | 0 | 0 | 0 | 0 | 1 | 18 | 1521 | 29.697328 | 180 |
| GSE37301_COMMON_LYMPHOID_PROGENITOR_VS_PRO_BCELL_DN | 3.68828595 | 26.09344 | ART_defective | defective | ART | 1 | 0 | 0 | 1 | 0 | 0 | 1 | 21 | 1737 | 30.240926 | 181 |
| GSE37301_PRO_BCELL_VS_CD4_TCELL_UP | 3.70068355 | 8.210544 | ART_defective | defective | ART | 0 | 1 | 0 | 1 | 0 | 0 | 2 | 18 | 2120 | 35.673825 | 182 |
| GSE37416_0H_VS_48H_F_TULARENSIS_LVS_NEUTROPHIL_UP | 2.83865179 | 26.31216 | ART_defective | defective | ART | 1 | 0 | 0 | 0 | 0 | 0 | 2 | 16 | 1554 | 29.000552 | 183 |
| GSE37532_WT_VS_PPARG_KO_LN_TCONV_UP | 5.06788052 | 10.65768 | ART_defective | defective | ART | 0 | 0 | 0 | 0 | 0 | 0 | 2 | 29 | 2948 | 27.843441 | 184 |
| GSE37533_PPARG1_FOXP3_VS_PPARG2_FOXP3_TRANSDUCED_CD4_TCELL_DN | 3.84459902 | 18.49607 | ART_defective | defective | ART | 0 | 1 | 0 | 0 | 0 | 0 | 1 | 22 | 1492 | 29.7012 | 185 |
| GSE37533_UNTREATED_VS_PIOGLIZATONE_TREATED_CD4_TCELL_FOXP3_TRASDUCED_CD4_TCELL_DN | 2.80470186 | 44.25219 | ART_defective | defective | ART | 0 | 1 | 0 | 0 | 0 | 0 | 1 | 13 | 998 | 23.643802 | 186 |
| GSE38696_LIGHT_ZONE_VS_DARK_ZONE_BCELL_UP | 3.79901089 | 21.84487 | ART_defective | defective | ART | 0 | 0 | 0 | 1 | 0 | 0 | 2 | 20 | 2395 | 43.752621 | 187 |
| GSE3982_EOSINOPHIL_VS_BASOPHIL_DN | 2.82431516 | 9.472439 | ART_defective | defective | ART | 1 | 0 | 0 | 0 | 0 | 0 | 1 | 16 | 1032 | 35.651127 | 188 |
| GSE3982_MAST_CELL_VS_TH1_DN | 2.82431516 | 13.60737 | ART_defective | defective | ART | 1 | 0 | 0 | 0 | 1 | 0 | 1 | 16 | 1526 | 38.954785 | 189 |
| GSE39820_CTRL_VS_TGFBETA1_IL6_IL23A_CD4_TCELL_UP | 3.68828595 | 66.33988 | ART_defective | defective | ART | 0 | 1 | 0 | 0 | 0 | 1 | 1 | 21 | 1814 | 29.130089 | 190 |
| GSE39820_CTRL_VS_TGFBETA3_IL6_IL23A_CD4_TCELL_UP | 3.32033552 | 8.001547 | ART_defective | defective | ART | 0 | 1 | 0 | 0 | 0 | 1 | 2 | 19 | 1656 | 25.4738 | 191 |
| GSE39820_TGFBETA1_IL6_VS_TGFBETA1_IL6_IL23A_TREATED_CD4_TCELL_UP | 3.14558101 | 14.12464 | ART_defective | defective | ART | 0 | 1 | 0 | 0 | 0 | 1 | 2 | 18 | 1624 | 29.250914 | 192 |
| GSE39820_TGFBETA3_IL6_VS_TGFBETA3_IL6_IL23A_TREATED_CD4_TCELL_UP | 2.81012263 | 5.484463 | ART_defective | defective | ART | 0 | 1 | 0 | 0 | 0 | 1 | 2 | 16 | 1244 | 34.787617 | 193 |
| GSE40068_BCL6_POS_VS_NEG_CXCR5_POS_TFH_DN | 2.81012263 | 75.79005 | ART_defective | defective | ART | 1 | 0 | 0 | 0 | 0 | 1 | 1 | 16 | 1369 | 27.347982 | 194 |
| GSE40273_EOS_KO_VS_WT_TREG_DN | 2.81012263 | 20.68757 | ART_defective | defective | ART | 1 | 0 | 0 | 0 | 0 | 0 | 1 | 16 | 1565 | 49.036765 | 195 |
| GSE40273_GATA1_KO_VS_WT_TREG_DN | 3.16138795 | 18.29952 | ART_defective | defective | ART | 1 | 0 | 0 | 0 | 0 | 0 | 1 | 18 | 2012 | 40.682327 | 196 |
| GSE40274_FOXP3_VS_FOXP3_AND_XBP1_TRANSDUCED_ACTIVATED_CD4_TCELL_DN | 2.85762077 | 3.723939 | ART_defective | defective | ART | 0 | 1 | 0 | 0 | 0 | 0 | 1 | 13 | 1104 | 30.818051 | 197 |
| GSE40274_IRF4_VS_FOXP3_AND_IRF4_TRANSDUCED_ACTIVATED_CD4_TCELL_UP | 2.91257501 | 36.06088 | ART_defective | defective | ART | 0 | 1 | 0 | 0 | 0 | 0 | 2 | 15 | 1179 | 33.065954 | 198 |
| GSE411_UNSTIM_VS_100MIN_IL6_STIM_SOCS3_KO_MACROPHAGE_DN | 2.97082651 | 9.078171 | ART_defective | defective | ART | 0 | 0 | 0 | 0 | 1 | 1 | 1 | 17 | 1425 | 37.871773 | 199 |
| GSE411_WT_VS_SOCS3_KO_MACROPHAGE_DN | 3.84459902 | 12.60777 | ART_defective | defective | ART | 0 | 0 | 0 | 0 | 1 | 0 | 1 | 22 | 1461 | 25.80718 | 200 |
| GSE41867_DAY6_EFFECTOR_VS_DAY30_MEMORY_CD8_TCELL_LCMV_ARMSTRONG_DN | 3.14558101 | 25.52355 | ART_defective | defective | ART | 0 | 0 | 1 | 0 | 0 | 0 | 1 | 18 | 1805 | 41.253607 | 201 |
| GSE41867_DAY8_VS_DAY15_LCMV_CLONE13_EFFECTOR_CD8_TCELL_UP | 2.97082651 | 13.40146 | ART_defective | defective | ART | 0 | 0 | 1 | 0 | 0 | 0 | 2 | 17 | 1592 | 36.98957 | 202 |
| GSE42724_MEMORY_BCELL_VS_PLASMABLAST_UP | 2.81012263 | 34.20753 | ART_defective | defective | ART | 0 | 0 | 0 | 1 | 1 | 0 | 2 | 16 | 898 | 24.555162 | 203 |
| GSE43863_LY6C_INT_CXCR5POS_VS_LY6C_LOW_CXCR5NEG_EFFECTOR_CD4_TCELL_UP | 2.98575529 | 14.88202 | ART_defective | defective | ART | 0 | 1 | 0 | 0 | 0 | 1 | 2 | 17 | 1344 | 29.36225 | 204 |
| GSE45365_WT_VS_IFNAR_KO_CD8A_DC_MCMV_INFECTION_UP | 3.68828595 | 19.54197 | ART_defective | defective | ART | 0 | 0 | 0 | 0 | 1 | 1 | 2 | 21 | 2012 | 34.53081 | 205 |
| GSE45739_UNSTIM_VS_ACD3_ACD28_STIM_NRAS_KO_CD4_TCELL_UP | 3.66984452 | 21.29092 | ART_defective | defective | ART | 0 | 1 | 0 | 0 | 0 | 0 | 2 | 21 | 1684 | 40.753276 | 206 |
| GSE45739_UNSTIM_VS_ACD3_ACD28_STIM_WT_CD4_TCELL_UP | 2.97082651 | 13.26679 | ART_defective | defective | ART | 0 | 1 | 0 | 0 | 0 | 0 | 2 | 17 | 1360 | 32.31044 | 207 |
| GSE45881_CXCR6HI_VS_CXCR1LO_COLONIC_LAMINA_PROPRIA_UP | 4.19410802 | 22.45634 | ART_defective | defective | ART | 0 | 0 | 0 | 0 | 0 | 1 | 2 | 24 | 1865 | 29.008148 | 208 |
| GSE4590_LARGE_PRE_BCELL_VS_VPREB_POS_LARGE_PRE_BCELL_DN | 2.91257501 | 11.32757 | ART_defective | defective | ART | 0 | 0 | 0 | 1 | 0 | 0 | 1 | 15 | 1762 | 34.597474 | 209 |
| GSE4984_GALECTIN1_VS_LPS_STIM_DC_UP | 3.07743775 | 17.36615 | ART_defective | defective | ART | 0 | 0 | 0 | 0 | 1 | 0 | 2 | 14 | 1259 | 42.94856 | 210 |
| GSE4984_LPS_VS_VEHICLE_CTRL_TREATED_DC_DN | 3.88343335 | 11.49518 | ART_defective | defective | ART | 0 | 0 | 0 | 0 | 1 | 0 | 1 | 22 | 2116 | 26.196275 | 211 |
| GSE5099_CLASSICAL_M1_VS_ALTERNATIVE_M2_MACROPHAGE_UP | 2.88254847 | 76.10618 | ART_defective | defective | ART | 0 | 0 | 0 | 0 | 1 | 0 | 2 | 16 | 1141 | 23.018286 | 212 |

| | | | | | | | | | | | | | | | | |
|---|---|---|---|---|---|---|---|---|---|---|---|---|---|---|---|---|
| GSE5503_PLN_DC_VS_SPLEEN_DC_ACTIVATED_ALLOGENIC_TCELL_DN | 2.97082651 | 14.81533 | ART_defective | defective | ART | 1 | 0 | 0 | 0 | 1 | 0 | 1 | 17 | 803 | 21.429437 | 213 |
| GSE5542_UNTREATED_VS_IFNA_AND_IFNG_TREATED_EPITHELIAL_CELLS_6H_DN | 3.86391861 | 18.70222 | ART_defective | defective | ART | 1 | 0 | 0 | 0 | 0 | 1 | 1 | 22 | 1546 | 26.29422 | 214 |
| GSE5542_UNTREATED_VS_IFNA_TREATED_EPITHELIAL_CELLS_6H_UP | 3.16138795 | 19.74873 | ART_defective | defective | ART | 1 | 0 | 0 | 0 | 0 | 1 | 2 | 18 | 1373 | 24.502758 | 215 |
| GSE5542_UNTREATED_VS_IFNG_TREATED_EPITHELIAL_CELLS_24H_UP | 2.82431516 | 13.03255 | ART_defective | defective | ART | 1 | 0 | 0 | 0 | 0 | 1 | 2 | 16 | 1417 | 24.8172 | 216 |
| GSE5542_UNTREATED_VS_IFNG_TREATED_EPITHELIAL_CELLS_6H_DN | 3.32033552 | 13.00535 | ART_defective | defective | ART | 1 | 0 | 0 | 0 | 0 | 1 | 1 | 19 | 1593 | 21.998062 | 217 |
| GSE5589_LPS_AND_IL10_VS_LPS_AND_IL6_STIM_MACROPHAGE_45MIN_DN | 3.14558101 | 15.16059 | ART_defective | defective | ART | 0 | 0 | 0 | 0 | 1 | 1 | 1 | 18 | 1418 | 21.97079 | 218 |
| GSE5679_CTRL_VS_PPARG_LIGAND_ROSIGLITAZONE_TREATED_DC_DN | 2.97082651 | 64.77403 | ART_defective | defective | ART | 0 | 0 | 0 | 0 | 1 | 0 | 1 | 17 | 1811 | 41.090169 | 219 |
| GSE5679_PPARG_LIGAND_ROSIGLITAZONE_VS_ROSIGLITAZONE_AND_RARA_AGONIST_AM580_TREATED_DC_UP | 2.97082651 | 3.867074 | ART_defective | defective | ART | 0 | 0 | 0 | 0 | 1 | 0 | 2 | 17 | 1080 | 34.645995 | 220 |
| GSE6092_UNSTIM_VS_IFNG_STIM_ENDOTHELIAL_CELL_DN | 3.16138795 | 29.21372 | ART_defective | defective | ART | 1 | 0 | 0 | 0 | 0 | 1 | 1 | 18 | 2332 | 45.292299 | 221 |
| GSE6674_ANTI_IGM_VS_CPG_STIM_BCELL_UP | 3.49509002 | 8.932795 | ART_defective | defective | ART | 0 | 0 | 0 | 1 | 0 | 1 | 2 | 20 | 2028 | 31.730084 | 222 |
| GSE7219_UNSTIM_VS_LPS_AND_ANTI_CD40_STIM_NIK_NFKB2_KO_DC_UP | 3.73071406 | 11.64773 | ART_defective | defective | ART | 0 | 0 | 0 | 0 | 1 | 1 | 2 | 19 | 2334 | 33.641684 | 223 |
| GSE7219_WT_VS_NIK_NFKB2_KO_DC_DN | 2.89534084 | 12.285 | ART_defective | defective | ART | 0 | 0 | 0 | 0 | 1 | 0 | 1 | 14 | 1864 | 33.761072 | 224 |
| GSE7348_LPS_VS_TOLERIZED_AND_LPS_STIM_MACROPHAGE_DN | 3.41474312 | 11.1207 | ART_defective | defective | ART | 0 | 0 | 0 | 0 | 1 | 0 | 1 | 17 | 1219 | 30.797486 | 225 |
| GSE7348_UNSTIM_VS_TOLERIZED_AND_LPS_STIM_MACROPHAGE_UP | 3.45194076 | 17.7758 | ART_defective | defective | ART | 0 | 0 | 0 | 0 | 1 | 0 | 2 | 16 | 1505 | 48.079663 | 226 |
| GSE7460_CD8_TCELL_VS_TREG_ACT_DN | 3.16138795 | 5.785245 | ART_defective | defective | ART | 1 | 0 | 1 | 0 | 0 | 0 | 1 | 18 | 1636 | 24.950196 | 227 |
| GSE7460_CTRL_VS_TGFB_TREATED_ACT_FOXP3HET_TCONV_UP | 3.14558101 | 22.68108 | ART_defective | defective | ART | 1 | 0 | 0 | 0 | 0 | 1 | 2 | 18 | 1317 | 31.500313 | 228 |
| GSE7460_FOXP3_MUT_VS_WT_ACT_WITH_TGFB_TCONV_DN | 3.49509002 | 8.791815 | ART_defective | defective | ART | 1 | 0 | 0 | 0 | 0 | 1 | 1 | 20 | 2358 | 39.325461 | 229 |
| GSE7509_UNSTIM_VS_FCGRIIB_STIM_MONOCYTE_DN | 2.94766628 | 25.28385 | ART_defective | defective | ART | 0 | 0 | 0 | 0 | 1 | 0 | 1 | 14 | 1426 | 44.930631 | 230 |
| GSE7568_CTRL_VS_3H_TGFB_TREATED_MACROPHAGES_WITH_IL4_AND_DEXAMETHASONE_DN | 2.93001558 | 14.07031 | ART_defective | defective | ART | 0 | 0 | 0 | 0 | 1 | 1 | 1 | 14 | 1134 | 33.482703 | 231 |
| GSE7568_IL4_TGFB_DEXAMETHASONE_VS_IL4_TGFB_TREATED_MACROPHAGE_DN | 3.3952303 | 4.989516 | ART_defective | defective | ART | 0 | 0 | 0 | 0 | 1 | 1 | 1 | 17 | 1328 | 25.669118 | 232 |
| GSE7568_IL4_VS_IL4_AND_TGFB_TREATED_MACROPHAGE_24H_UP | 3.13930241 | 10.76496 | ART_defective | defective | ART | 0 | 0 | 0 | 0 | 1 | 1 | 2 | 15 | 896 | 29.919094 | 233 |
| GSE7596_AKT_TRANSD_VS_CTRL_CD4_TCONV_WITH_TGFB_UP | 2.82212237 | 16.95831 | ART_defective | defective | ART | 0 | 1 | 0 | 0 | 0 | 1 | 2 | 13 | 802 | 21.519176 | 234 |
| GSE7768_OVA_ALONE_VS_OVA_WITH_LPS_IMMUNIZED_MOUSE_WHOLE_SPLEEN_6H_DN | 2.87830943 | 10.85188 | ART_defective | defective | ART | 0 | 0 | 0 | 0 | 0 | 0 | 1 | 14 | 1487 | 36.365 | 235 |
| GSE7768_OVA_ALONE_VS_OVA_WITH_MPL_IMMUNIZED_MOUSE_WHOLE_SPLEEN_6H_DN | 2.98361343 | 14.38027 | ART_defective | defective | ART | 0 | 0 | 0 | 0 | 0 | 0 | 1 | 14 | 896 | 31.929795 | 236 |
| GSE7831_UNSTIM_VS_CPG_STIM_PDC_4H_UP | 2.97082651 | 37.35867 | ART_defective | defective | ART | 0 | 0 | 0 | 0 | 1 | 0 | 2 | 17 | 1600 | 36.01534 | 237 |
| GSE9006_HEALTHY_VS_TYPE_1_DIABETES_PBMC_1MONTH_POST_DX_UP | 3.14558101 | 34.63416 | ART_defective | defective | ART | 0 | 0 | 0 | 0 | 1 | 0 | 2 | 18 | 1275 | 24.757445 | 238 |
| GSE9601_UNTREATED_VS_NFKB_INHIBITOR_TREATED_HCMV_INF_MONOCYTE_DN | 3.8164776 | 10.79661 | ART_defective | defective | ART | 0 | 0 | 0 | 0 | 1 | 1 | 1 | 19 | 2184 | 33.569796 | 239 |
| GSE9601_UNTREATED_VS_PI3K_INHIBITOR_TREATED_HCMV_INF_MONOCYTE_UP | 2.9787699 | 14.28206 | ART_defective | defective | ART | 0 | 0 | 0 | 0 | 1 | 1 | 2 | 15 | 1502 | 46.98842 | 240 |
| GSE9650_NAIVE_VS_EFF_CD8_TCELL_UP | 2.97082651 | 32.12113 | ART_defective | defective | ART | 0 | 0 | 1 | 0 | 0 | 0 | 2 | 17 | 1329 | 25.49485 | 241 |
| GSE9650_NAIVE_VS_MEMORY_CD8_TCELL_UP | 3.14558101 | 13.72964 | ART_defective | defective | ART | 0 | 0 | 1 | 0 | 0 | 0 | 2 | 18 | 1560 | 31.801565 | 242 |
| GSE9960_HEALTHY_VS_GRAM_NEG_AND_POS_SEPSIS_PBMC_DN | 3.03145563 | 10.82472 | ART_defective | defective | ART | 0 | 0 | 0 | 0 | 1 | 0 | 1 | 17 | 1408 | 30.483851 | 243 |
| GSE9988_ANTI_TREM1_AND_LPS_VS_CTRL_TREATED_MONOCYTES_DN | 2.83865179 | 33.32605 | ART_defective | defective | ART | 0 | 0 | 0 | 0 | 1 | 0 | 1 | 16 | 1072 | 30.094962 | 244 |
| GSE9988_ANTI_TREM1_VS_CTRL_TREATED_MONOCYTES_DN | 4.19410802 | 25.88527 | ART_defective | defective | ART | 0 | 0 | 0 | 0 | 1 | 0 | 1 | 24 | 1388 | 31.18617 | 245 |
| GSE9988_ANTI_TREM1_VS_VEHICLE_TREATED_MONOCYTES_DN | 3.54831474 | 26.68324 | ART_defective | defective | ART | 0 | 0 | 0 | 0 | 1 | 0 | 1 | 20 | 1483 | 32.04475 | 246 |
| GSE9988_LOW_LPS_VS_CTRL_TREATED_MONOCYTE_DN | 3.84459902 | 21.83586 | ART_defective | defective | ART | 0 | 0 | 0 | 0 | 1 | 0 | 1 | 22 | 1377 | 21.359286 | 247 |
| HOFT_CD4_POSITIVE_ALPHA_BETA_MEMORY_T_CELL_BCG_VACCINE_AGE_18_45YO_56D_TOP_100_DEG_AFTER_IN_VITRO_RE_STIMULATION_DN | 5.20545322 | 11.50684 | ART_defective | defective | ART | 0 | 1 | 0 | 0 | 0 | 0 | 1 | 7 | 819 | 24.224515 | 248 |
| KAECH_NAIVE_VS_DAY8_EFF_CD8_TCELL_UP | 3.49509002 | 27.43817 | ART_defective | defective | ART | 0 | 0 | 1 | 0 | 0 | 0 | 2 | 20 | 1782 | 28.183559 | 249 |
| KAECH_NAIVE_VS_MEMORY_CD8_TCELL_UP | 3.32033552 | 12.32811 | ART_defective | defective | ART | 0 | 0 | 1 | 0 | 0 | 0 | 2 | 19 | 1734 | 40.96638 | 250 |
| NAKAYA_PBMC_FLUARIX_FLUVIRIN_AGE_18_50YO_3DY_DN | 2.90584851 | 39.75762 | ART_defective | defective | ART | 0 | 0 | 0 | 0 | 1 | 0 | 1 | 36 | 3136 | 24.470139 | 251 |
| NAKAYA_PBMC_FLUARIX_FLUVIRIN_AGE_18_50YO_CORRELATED_WITH_HAI_28DY_RESPONSE_AT_3DY_NEGATIVE | 3.45668243 | 164.7889 | ART_defective | defective | ART | 0 | 0 | 0 | 0 | 1 | 0 | 0 | 36 | 3281 | 24.731514 | 252 |
| NAKAYA_PBMC_FLUARIX_FLUVIRIN_AGE_18_50YO_CORRELATED_WITH_HAI_28DY_RESPONSE_AT_7DY_NEGATIVE | 3.38471875 | 136.2366 | ART_defective | defective | ART | 0 | 0 | 0 | 0 | 1 | 0 | 0 | 46 | 3805 | 27.643409 | 253 |
| OCONNOR_PBMC_MENVEO_ACWYVAX_AGE_30_70YO_7DY_AFTER_SECOND_DOSE_VS_7DY_AFTER_FIRST_DOSE_UP | 5.54393589 | 21.42166 | ART_defective | defective | ART | 0 | 0 | 0 | 0 | 1 | 0 | 2 | 23 | 2469 | 25.959345 | 254 |
| QI_PBMC_ZOSTAVAX_AGE_50_75YO_CORRELATED_WITH_CONTRACTION_OF_VZV_SPECIFIC_T_CELLS_PEAK_TO_28DY_AT_1DYPOSITIVE | 3.09089593 | 56.31121 | ART_defective | defective | ART | 1 | 0 | 0 | 0 | 0 | 0 | 0 | 26 | 1983 | 37.225912 | 255 |
| THAKAR_PBMC_INACTIVATED_INFLUENZA_AGE_21_30YO_NONRESPONDER_28DY_UP | 3.01300863 | 21.58292 | ART_defective | defective | ART | 0 | 0 | 0 | 0 | 1 | 0 | 2 | 15 | 647 | 17.359028 | 256 |
| THAKAR_PBMC_INACTIVATED_INFLUENZA_AGE_21_30YO_NONRESPONDER_7DY_UP | 3.30244726 | 11.91183 | ART_defective | defective | ART | 0 | 0 | 0 | 0 | 1 | 0 | 2 | 12 | 435 | 14.353879 | 257 |
| ZAK_PBMC_MRKAD5_HIV_1_GAG_POL_NEF_AGE_20_50YO_1DY_ADDNL_EXON_LVL_UP | 3.71818087 | 18.39302 | ART_defective | defective | ART | 0 | 0 | 0 | 0 | 1 | 0 | 2 | 10 | 1192 | 43.322917 | 258 |
| ZAK_PBMC_MRKAD5_HIV_1_GAG_POL_NEF_AGE_20_50YO_CORRELATED_WITH_CD8_T_CELL_RESPONSE_3DY_POSITIVE | 3.7447393 | 12.30842 | ART_defective | defective | ART | 0 | 0 | 1 | 0 | 1 | 0 | 0 | 9 | 665 | 19.380723 | 259 |
| GSE11057_CD4_EFF_MEM_VS_PBMC_UP | 10.5986724 | 1.288995 | EC_intact | intact | EC | 0 | 1 | 0 | 0 | 1 | 0 | 2 | 8 | 791 | 43.007865 | 260 |
| GSE11057_NAIVE_CD4_VS_PBMC_CD4_TCELL_UP | 8.02970483 | 1.324372 | EC_intact | intact | EC | 0 | 1 | 0 | 0 | 1 | 0 | 2 | 6 | 729 | 46.910476 | 261 |
| GSE16450_CTRL_VS_IFNA_12H_STIM_IMMATURE_NEURON_CELL_LINE_UP | 8.02970483 | 0.167586 | EC_intact | intact | EC | 1 | 0 | 0 | 0 | 0 | 1 | 2 | 6 | 1187 | 41.175 | 262 |
| GSE20366_TREG_VS_NAIVE_CD4_TCELL_HOMEOSTATIC_CONVERSION_DN | 7.94900428 | 0.736233 | EC_intact | intact | EC | 0 | 1 | 0 | 0 | 0 | 0 | 1 | 6 | 622 | 38.040111 | 263 |
| GSE22601_IMMATURE_CD4_SINGLE_POSITIVE_VS_CD8_SINGLE_POSITIVE_THYMOCYTE_DN | 7.90925926 | 2.383826 | EC_intact | intact | EC | 0 | 1 | 1 | 0 | 0 | 0 | 1 | 6 | 301 | 37.544355 | 264 |
| GSE25146_UNSTIM_VS_HELIOBACTER_PYLORI_LPS_STIM_AGS_CELL_DN | 9.52922802 | 6.267102 | EC_intact | intact | EC | 1 | 0 | 0 | 0 | 0 | 0 | 1 | 6 | 403 | 40.981818 | 265 |
| GSE26488_CTRL_VS_PEPTIDE_INJECTION_OT2_THYMOCYTE_DN | 8.64399919 | 0.665595 | EC_intact | intact | EC | 1 | 0 | 0 | 0 | 0 | 0 | 1 | 6 | 371 | 23.458686 | 266 |
| GSE30971_WBP7_HET_VS_KO_MACROPHAGE_UP | 7.90925926 | 3.36305 | EC_intact | intact | EC | 0 | 0 | 0 | 0 | 1 | 0 | 2 | 6 | 334 | 45.288165 | 267 |
| GSE31082_DN_VS_CD4_SP_THYMOCYTE_DN | 9.22746914 | 1.907951 | EC_intact | intact | EC | 0 | 1 | 0 | 0 | 0 | 0 | 1 | 7 | 246 | 44.649123 | 268 |
| GSE40225_WT_VS_RIP_B7X_DIABETIC_MOUSE_PANCREATIC_CD8_TCELL_UP | 9.27383833 | 1.354557 | EC_intact | intact | EC | 0 | 0 | 1 | 0 | 0 | 0 | 2 | 7 | 665 | 25.718282 | 269 |
| GSE5099_DAY3_VS_DAY7_MCSF_TREATED_MACROPHAGE_DN | 8.64399919 | 1167.992 | EC_intact | intact | EC | 0 | 0 | 0 | 0 | 1 | 0 | 1 | 6 | 220 | 27.605549 | 270 |
| GSE5142_CTRL_VS_HTERT_TRANSDUCED_CD8_TCELL_EARLY_PASSAGE_CLONE_UP | 7.90925926 | 7.436006 | EC_intact | intact | EC | 0 | 0 | 1 | 0 | 0 | 0 | 2 | 6 | 501 | 37.21123 | 271 |
| GSE6674_ANTI_IGM_VS_CPG_STIM_BCELL_UP | 7.90925926 | 1.326208 | EC_intact | intact | EC | 0 | 0 | 0 | 1 | 0 | 1 | 2 | 6 | 345 | 31.971831 | 272 |
| NAKAYA_PBMC_FLUARIX_FLUVIRIN_AGE_18_50YO_CORRELATED_WITH_HAI_28DY_RESPONSE_AT_7DY_NEGATIVE | 5.55035737 | 468.4237 | EC_intact | intact | EC | 0 | 0 | 0 | 0 | 1 | 0 | 0 | 10 | 915 | 28.832028 | 273 |
| GSE10239_NAIVE_VS_KLRG1HIGH_EFF_CD8_TCELL_UP | 6.77757207 | 10.51001 | EC_defective | defective | EC | 0 | 0 | 1 | 0 | 0 | 0 | 2 | 6 | 690 | 38.978462 | 274 |
| GSE11057_NAIVE_CD4_VS_PBMC_CD4_TCELL_UP | 6.84637991 | 9.256685 | EC_defective | defective | EC | 0 | 1 | 0 | 0 | 1 | 0 | 2 | 6 | 575 | 72.367713 | 275 |
| GSE11386_NAIVE_VS_MEMORY_BCELL_DN | 9.89639192 | 7.899831 | EC_defective | defective | EC | 0 | 0 | 0 | 1 | 0 | 0 | 1 | 7 | 806 | 37.002903 | 276 |

| | | | | | | | | | | | | | | | |
|---|---|---|---|---|---|---|---|---|---|---|---|---|---|---|---|
| GSE11864_CSF1_VS_CSF1_PAM3CYS_IN_MAC_UP | 6.84637991 | 2.692919 | EC_defective | defective | EC | 0 | 0 | 0 | 0 | 1 | 0 | 2 | 6 | 437 | 22.937733 | 277 |
| GSE12392_WT_VS_IFNB_KO_CD8A_NEG_SPLEEN_DC_DN | 6.74368421 | 0.833038 | EC_defective | defective | EC | 0 | 0 | 0 | 0 | 1 | 1 | 1 | 6 | 747 | 38.937385 | 278 |
| GSE12845_IGD_POS_BLOOD_VS_NAIVE_TONSIL_BCELL_DN | 7.90716742 | 4.359668 | EC_defective | defective | EC | 0 | 0 | 0 | 1 | 0 | 0 | 1 | 7 | 771 | 38.989222 | 279 |
| GSE13411_IGM_MEMORY_BCELL_VS_PLASMA_CELL_UP | 6.81180223 | 0.826328 | EC_defective | defective | EC | 1 | 0 | 0 | 1 | 0 | 1 | 2 | 6 | 546 | 28.671001 | 280 |
| GSE16385_ROSIGLITAZONE_IL4_VS_IFNG_TNF_STIM_MACROPHAGE_UP | 8.99157895 | 7.448678 | EC_defective | defective | EC | 0 | 0 | 0 | 0 | 1 | 1 | 2 | 8 | 796 | 36.443503 | 281 |
| GSE16450_IMMATURE_VS_MATURE_NEURON_CELL_LINE_UP | 6.91659919 | 11.05271 | EC_defective | defective | EC | 1 | 0 | 0 | 0 | 0 | 0 | 2 | 6 | 916 | 72.693717 | 282 |
| GSE17301_ACD3_ACD28_VS_ACD3_ACD28_AND_IFNA2_STIM_CD8_TCELL_DN | 6.74368421 | 0.954911 | EC_defective | defective | EC | 0 | 0 | 1 | 0 | 0 | 1 | 1 | 6 | 281 | 22.8637 | 283 |
| GSE17301_IFNA2_VS_IFNA2_AND_ACD3_ACD28_STIM_CD8_TCELL_UP | 6.74368421 | 0.954911 | EC_defective | defective | EC | 0 | 0 | 1 | 0 | 0 | 1 | 2 | 6 | 281 | 22.8637 | 284 |
| GSE17721_CTRL_VS_GARDIQUIMOD_0.5H_BMDC_UP | 6.74368421 | 2.188483 | EC_defective | defective | EC | 0 | 0 | 0 | 0 | 1 | 0 | 2 | 6 | 391 | 37.969412 | 285 |
| GSE19198_6H_VS_24H_IL21_TREATED_TCELL_DN | 6.81180223 | 10.62257 | EC_defective | defective | EC | 1 | 0 | 0 | 0 | 0 | 1 | 1 | 6 | 657 | 43.740385 | 286 |
| GSE20715_0H_VS_24H_OZONE_LUNG_UP | 6.74368421 | 0.818558 | EC_defective | defective | EC | 0 | 0 | 0 | 0 | 0 | 0 | 2 | 6 | 736 | 35.787208 | 287 |
| GSE21063_3H_VS_16H_ANTI_IGM_STIM_BCELL_DN | 6.84637991 | 0.403675 | EC_defective | defective | EC | 0 | 0 | 0 | 1 | 0 | 1 | 1 | 6 | 572 | 28.417526 | 288 |
| GSE21670_UNTREATED_VS_IL6_TREATED_CD4_TCELL_DN | 8.99157895 | 9.689526 | EC_defective | defective | EC | 0 | 1 | 0 | 0 | 0 | 1 | 1 | 8 | 760 | 37.150393 | 289 |
| GSE22886_NAIVE_TCELL_VS_DC_UP | 6.77757207 | 9.533025 | EC_defective | defective | EC | 1 | 0 | 0 | 0 | 1 | 0 | 2 | 6 | 412 | 25.514706 | 290 |
| GSE23321_CD8_STEM_CELL_MEMORY_VS_NAIVE_CD8_TCELL_DN | 6.88131042 | 3.125561 | EC_defective | defective | EC | 0 | 0 | 1 | 0 | 0 | 0 | 1 | 6 | 791 | 34.433498 | 291 |
| GSE24574_BCL6_HIGH_VS_LOW_TFH_CD4_TCELL_UP | 7.90716742 | 12.8515 | EC_defective | defective | EC | 0 | 1 | 0 | 0 | 0 | 0 | 2 | 7 | 687 | 32.18018 | 292 |
| GSE25088_IL4_VS_IL4_AND_ROSIGLITAZONE_STIM_STAT6_KO_MACROPHAGE_DAY10_UP | 6.81180223 | 4.009084 | EC_defective | defective | EC | 0 | 0 | 0 | 0 | 1 | 1 | 2 | 6 | 815 | 49.55615 | 293 |
| GSE25123_CTRL_VS_IL4_AND_ROSIGLITAZONE_STIM_MACROPHAGE_UP | 6.74368421 | 10.8133 | EC_defective | defective | EC | 0 | 0 | 0 | 0 | 1 | 1 | 2 | 6 | 706 | 71.32491 | 294 |
| GSE27241_WT_VS_RORGT_KO_TH17_POLARIZED_CD4_TCELL_TREATED_WITH_DIGOXIN_UP | 12.4883041 | 0.710434 | EC_defective | defective | EC | 0 | 1 | 0 | 0 | 0 | 0 | 2 | 10 | 883 | 35.332649 | 295 |
| GSE27291_0H_VS_7D_STIM_GAMMADELTA_TCELL_UP | 8.36982083 | 1.814628 | EC_defective | defective | EC | 1 | 0 | 0 | 0 | 0 | 0 | 2 | 7 | 322 | 23.844691 | 296 |
| GSE27786_LIN_NEG_VS_ERYTHROBLAST_UP | 6.74368421 | 0.712847 | EC_defective | defective | EC | 1 | 0 | 0 | 0 | 0 | 0 | 2 | 6 | 403 | 25.635849 | 297 |
| GSE27786_NKTCELL_VS_ERYTHROBLAST_UP | 7.90716742 | 1.807833 | EC_defective | defective | EC | 1 | 0 | 0 | 0 | 0 | 0 | 2 | 7 | 297 | 25.558047 | 298 |
| GSE29618_BCELL_VS_MDC_DAY7_FLU_VACCINE_UP | 7.02467105 | 9.018799 | EC_defective | defective | EC | 0 | 0 | 0 | 1 | 1 | 0 | 2 | 6 | 696 | 39.662866 | 299 |
| GSE29618_BCELL_VS_MDC_UP | 6.95225176 | 9.019471 | EC_defective | defective | EC | 0 | 0 | 0 | 1 | 1 | 0 | 2 | 6 | 576 | 33.138365 | 300 |
| GSE29618_BCELL_VS_MONOCYTE_DAY7_FLU_VACCINE_UP | 6.88131042 | 0.835904 | EC_defective | defective | EC | 0 | 0 | 0 | 1 | 1 | 0 | 2 | 6 | 616 | 35.486804 | 301 |
| GSE36476_CTRL_VS_TSST_ACT_72H_MEMORY_CD4_TCELL_YOUNG_UP | 6.81180223 | 9.113697 | EC_defective | defective | EC | 0 | 1 | 0 | 0 | 0 | 0 | 2 | 6 | 498 | 57.444 | 302 |
| GSE369_IFNG_KO_VS_WT_LIVER_UP | 9.03676276 | 1.837716 | EC_defective | defective | EC | 0 | 0 | 0 | 0 | 0 | 1 | 2 | 8 | 893 | 38.440934 | 303 |
| GSE3982_MAST_CELL_VS_BASOPHIL_DN | 6.77757207 | 0.734007 | EC_defective | defective | EC | 1 | 0 | 0 | 0 | 1 | 0 | 1 | 6 | 639 | 43.550976 | 304 |
| GSE40273_GATA1_KO_VS_WT_TREG_DN | 7.90716742 | 0.942543 | EC_defective | defective | EC | 1 | 0 | 0 | 0 | 0 | 0 | 1 | 7 | 683 | 36.37707 | 305 |
| GSE6092_UNSTIM_VS_IFNG_STIM_ENDOTHELIAL_CELL_DN | 6.77757207 | 3.418504 | EC_defective | defective | EC | 1 | 0 | 0 | 0 | 0 | 1 | 1 | 6 | 924 | 63.317549 | 306 |
| GSE8921_3H_VS_24H_TLR1_2_STIM_MONOCYTE_UP | 6.74368421 | 0.892339 | EC_defective | defective | EC | 0 | 0 | 0 | 0 | 1 | 0 | 2 | 6 | 550 | 31.932624 | 307 |
| NAKAYA_B_CELL_FLUARIX_FLUVIRIN_AGE_18_50YO_7DY_DN | 5.35213033 | 0.554024 | EC_defective | defective | EC | 0 | 0 | 0 | 1 | 0 | 0 | 1 | 7 | 732 | 54.212291 | 308 |
| NAKAYA_PBMC_FLUARIX_FLUVIRIN_AGE_18_50YO_3DY_DN | 4.67229853 | 7.046374 | EC_defective | defective | EC | 0 | 0 | 0 | 0 | 1 | 0 | 1 | 9 | 794 | 29.592398 | 309 |
