## Appendix Table S6 for "Distinguishable topological properties of functional genome networks in HIV-1 reservoirs"

**Supplementary Table 6. A complete attribute list of enriched immunologic signatures in pretreatment HIV-1-infected individuals, patients subjected to short and long period of ART, and elite controll**

| Description | patient_period | patient | period | is_other_cell | is_CD4_T_cell | is_CD8_T_cell | is_B_cell | is_myeloid_cell | is_proinflammatory_factor | Response | Count | Rich_factor | tpm | HiC | atac_seq_count | id |
|---|---|---|---|---|---|---|---|---|---|---|---|---|---|---|---|---|
| ANDERSON_BLOOD_CN54GP140_ADJUVANTED_WITH_GLA_AF_AGE_18_45YO_1DY_DN | ART_long | ART | long | 0 | 0 | 0 | 0 | 0 | 0 | 1 | 5 | 7.432996867 | 16.307828 | 17.49023024 | 231 | 1 |
| GSE10273_HIGH_VS_LOW_IL7_TREATED_IRF4_8_NULL_PRE_BCELL_DN | ART_long | ART | long | 0 | 0 | 0 | 1 | 0 | 1 | 1 | 7 | 4.467306198 | 15.55269296 | 46.92087363 | 461 | 2 |
| GSE10325_CD4_TCELL_VS_BCELL_UP | ART_long | ART | long | 0 | 1 | 1 | 0 | 0 | 0 | 2 | 9 | 5.743679398 | 14.28236246 | 23.51377888 | 1120 | 3 |
| GSE10325_CD4_TCELL_VS_MYELOID_UP | ART_long | ART | long | 0 | 1 | 0 | 0 | 1 | 0 | 2 | 7 | 4.444857423 | 13.15808781 | 22.88719502 | 799 | 4 |
| GSE10325_LUPUS_CD4_TCELL_VS_LUPUS_BCELL_UP | ART_long | ART | long | 0 | 1 | 0 | 1 | 0 | 0 | 2 | 7 | 4.444857423 | 10.94407582 | 17.34005735 | 797 | 5 |
| GSE10325_LUPUS_CD4_TCELL_VS_LUPUS_MYELOID_UP | ART_long | ART | long | 0 | 1 | 0 | 0 | 1 | 0 | 2 | 10 | 6.349796319 | 14.14178586 | 24.24388861 | 1033 | 6 |
| GSE11057_CD4_CENT_MEM_VS_PBMC_UP | ART_long | ART | long | 0 | 1 | 0 | 0 | 1 | 0 | 2 | 7 | 4.489982879 | 18.0807373 | 33.13461386 | 823 | 7 |
| GSE11057_EFF_MEM_VS_CENT_MEM_CD4_TCELL_DN | ART_long | ART | long | 0 | 1 | 0 | 0 | 0 | 0 | 1 | 11 | 7.39345965 | 8.822993205 | 24.56996784 | 1383 | 8 |
| GSE11057_NAIVE_VS_CENT_MEMORY_CD4_TCELL_DN | ART_long | ART | long | 0 | 1 | 0 | 0 | 0 | 0 | 1 | 9 | 5.714816687 | 10.8776386 | 27.74837044 | 984 | 9 |
| GSE11057_NAIVE_VS_MEMORY_CD4_TCELL_DN | ART_long | ART | long | 0 | 1 | 0 | 0 | 0 | 0 | 1 | 9 | 5.743679398 | 14.30913243 | 27.81670883 | 924 | 10 |
| GSE11057_PBMC_VS_MEM_CD4_TCELL_DN | ART_long | ART | long | 0 | 1 | 0 | 0 | 1 | 0 | 1 | 7 | 4.489982879 | 17.91944125 | 24.07281899 | 620 | 11 |
| GSE11386_NAIVE_VS_MEMORY_BCELL_DN | ART_long | ART | long | 0 | 0 | 0 | 1 | 0 | 0 | 1 | 8 | 6.357783484 | 8.546682272 | 27.60963244 | 1106 | 12 |
| GSE11961_FOLLICULAR_BCELL_VS_GERMINAL_CENTER_BCELL_DAY40_UP | ART_long | ART | long | 0 | 0 | 0 | 1 | 0 | 0 | 2 | 9 | 5.686242604 | 13.55538397 | 29.29691111 | 1108 | 13 |
| GSE12003_MIR223_KO_VS_WT_BM_PROGENITOR_4D_CULTURE_UP | ART_long | ART | long | 1 | 0 | 0 | 0 | 0 | 0 | 2 | 7 | 4.422633136 | 14.55890202 | 19.25350474 | 911 | 14 |
| GSE12366_GC_BCELL_VS_PLASMA_CELL_UP | ART_long | ART | long | 0 | 0 | 0 | 1 | 0 | 0 | 2 | 7 | 4.489982879 | 11.73761252 | 24.1486718 | 1043 | 15 |
| GSE12366_PLASMA_CELL_VS_NAIVE_BCELL_DN | ART_long | ART | long | 0 | 0 | 0 | 1 | 0 | 0 | 1 | 7 | 4.444857423 | 14.9501187 | 23.79828611 | 491 | 16 |
| GSE12392_IFNAR_KO_VS_IFNB_KO_CD8_NEG_SPLEEN_DC_UP | ART_long | ART | long | 0 | 0 | 0 | 0 | 1 | 1 | 2 | 7 | 4.444857423 | 16.99956853 | 27.97517591 | 875 | 17 |
| GSE12845_IGD_POS_BLOOD_VS_DARKZONE_GC_TONSIL_BCELL_DN | ART_long | ART | long | 0 | 0 | 0 | 1 | 0 | 0 | 1 | 7 | 4.444857423 | 13.24724617 | 15.2980302 | 969 | 18 |
| GSE12845_IGD_POS_BLOOD_VS_NAIVE_TONSIL_BCELL_DN | ART_long | ART | long | 0 | 0 | 0 | 1 | 0 | 0 | 1 | 9 | 5.714816687 | 15.7050284 | 29.22788088 | 1090 | 19 |
| GSE13411_NAIVE_BCELL_VS_PLASMA_CELL_UP | ART_long | ART | long | 0 | 0 | 0 | 1 | 1 | 0 | 2 | 10 | 6.446987079 | 9.746694837 | 21.05608947 | 703 | 20 |
| GSE13411_SWITCHED_MEMORY_BCELL_VS_PLASMA_CELL_UP | ART_long | ART | long | 0 | 0 | 0 | 1 | 1 | 0 | 2 | 8 | 5.05443787 | 10.44090721 | 17.08141693 | 506 | 21 |
| GSE13738_RESTING_VS_BYSTANDER_ACTIVATED_CD4_TCELL_DN | ART_long | ART | long | 0 | 1 | 0 | 0 | 0 | 0 | 1 | 7 | 4.467306198 | 17.82727733 | 31.96474167 | 684 | 22 |
| GSE13738_TCR_VS_BYSTANDER_ACTIVATED_CD4_TCELL_DN | ART_long | ART | long | 0 | 1 | 0 | 0 | 0 | 0 | 1 | 8 | 5.105492798 | 11.51463141 | 23.09920297 | 765 | 23 |
| GSE14308_TH17_VS_NATURAL_TREG_DN | ART_long | ART | long | 1 | 0 | 0 | 0 | 0 | 0 | 2 | 7 | 4.422633136 | 6.342228374 | 37.76689288 | 763 | 24 |
| GSE14350_TREG_VS_TEFF_UP | ART_long | ART | long | 1 | 0 | 0 | 0 | 0 | 0 | 2 | 9 | 5.714816687 | 19.48031196 | 26.85422465 | 913 | 25 |
| GSE14415_INDUCED_TREG_VS_FOXP3_KO_INDUCED_TREG_UP | ART_long | ART | long | 1 | 0 | 0 | 0 | 0 | 1 | 2 | 7 | 5.14259667 | 12.16035155 | 22.47927762 | 621 | 26 |
| GSE1460_INTRATHYMIC_T_PROGENITOR_VS_CD4_THYMOCYTE_DN | ART_long | ART | long | 1 | 1 | 0 | 0 | 0 | 0 | 1 | 7 | 4.444857423 | 15.44292819 | 38.58825264 | 809 | 27 |
| GSE1460_INTRATHYMIC_T_PROGENITOR_VS_THYMIC_STROMAL_CELL_UP | ART_long | ART | long | 1 | 0 | 0 | 0 | 0 | 0 | 2 | 7 | 4.467306198 | 6.601586143 | 30.30346702 | 649 | 28 |
| GSE14699_NAIVE_VS_DELETIONAL_TOLERANCE_CD8_TCELL_UP | ART_long | ART | long | 0 | 0 | 1 | 0 | 0 | 0 | 2 | 6 | 4.987932108 | 7.391113618 | 20.20755394 | 480 | 29 |
| GSE15330_HSC_VS_MEGAKARYOCYTE_ERYTHROID_PROGENITOR_DN | ART_long | ART | long | 1 | 0 | 0 | 0 | 0 | 0 | 1 | 7 | 5.265039448 | 8.246032246 | 21.34112744 | 584 | 30 |
| GSE15330_HSC_VS_MEGAKARYOCYTE_ERYTHROID_PROGENITOR_UP | ART_long | ART | long | 1 | 0 | 0 | 0 | 0 | 0 | 2 | 9 | 7.020052597 | 6.473732154 | 19.85642459 | 875 | 31 |
| GSE15330_LYMPHOID_MULTIPOTENT_VS_GRANULOCYTE_MONOCYTE_PROGENITOR_UP | ART_long | ART | long | 1 | 0 | 0 | 0 | 1 | 0 | 2 | 8 | 5.079837055 | 13.96020499 | 38.38807942 | 918 | 32 |
| GSE15330_LYMPHOID_MULTIPOTENT_VS_MEGAKARYOCYTE_ERYTHROID_PROGENITOR_IKAROS_KO_UP | ART_long | ART | long | 1 | 0 | 0 | 0 | 0 | 0 | 2 | 9 | 6.850894703 | 6.507661645 | 21.7368793 | 843 | 33 |
| GSE15624_CTRL_VS_3H_HALOFUGINONE_TREATED_CD4_TCELL_UP | ART_long | ART | long | 0 | 1 | 0 | 0 | 0 | 0 | 2 | 6 | 4.738535503 | 6.568545241 | 19.953673 | 526 | 34 |
| GSE16385_MONOCYTE_VS_12H_IL4_TREATED_MACROPHAGE_DN | ART_long | ART | long | 0 | 0 | 0 | 0 | 1 | 1 | 1 | 7 | 4.422633136 | 16.06730701 | 25.28941501 | 485 | 35 |
| GSE16385_ROSIGLITAZONE_IL4_VS_IFNG_TNF_STIM_MACROPHAGE_UP | ART_long | ART | long | 0 | 0 | 0 | 0 | 1 | 1 | 2 | 7 | 4.422633136 | 18.34397432 | 29.61828098 | 1091 | 36 |
| GSE16450_IMMATURE_VS_MATURE_NEURON_CELL_LINE_12H_IFNA_STIM_UP | ART_long | ART | long | 1 | 0 | 0 | 0 | 0 | 1 | 2 | 8 | 5.05443787 | 29.16781529 | 22.67805136 | 611 | 37 |
| GSE16755_CTRL_VS_IFNA_TREATED_MAC_UP | ART_long | ART | long | 0 | 0 | 0 | 0 | 1 | 1 | 1 | 7 | 4.536033986 | 13.76320594 | 22.92831587 | 620 | 38 |
| GSE17186_NAIVE_VS_CD21HIGH_TRANSITIONAL_BCELL_DN | ART_long | ART | long | 0 | 0 | 0 | 1 | 0 | 0 | 1 | 8 | 5.05443787 | 6.086904713 | 18.74569031 | 597 | 39 |
| GSE17186_NAIVE_VS_CD21LOW_TRANSITIONAL_BCELL_DN | ART_long | ART | long | 0 | 0 | 0 | 1 | 0 | 0 | 1 | 9 | 5.686242604 | 56.86467364 | 30.75723719 | 723 | 40 |
| GSE17301_CTRL_VS_48H_ACD3_ACD28_IFNA5_STIM_CD8_TCELL_UP | ART_long | ART | long | 0 | 0 | 1 | 0 | 0 | 1 | 2 | 7 | 4.422633136 | 11.89486178 | 24.21941574 | 1017 | 41 |
| GSE17721_0.5H_VS_24H_CPG_BMDC_UP | ART_long | ART | long | 0 | 0 | 0 | 0 | 1 | 0 | 2 | 7 | 4.422633136 | 48.81580302 | 21.17392432 | 440 | 42 |
| GSE17721_CTRL_VS_CPG_24H_BMDC_UP | ART_long | ART | long | 0 | 0 | 0 | 0 | 1 | 0 | 2 | 8 | 5.05443787 | 16.21049698 | 24.6836936 | 986 | 43 |
| GSE17721_LPS_VS_POLYIC_16H_BMDC_DN | ART_long | ART | long | 0 | 0 | 0 | 0 | 1 | 0 | 1 | 7 | 4.422633136 | 33.10617907 | 23.85225565 | 573 | 44 |
| GSE17721_PAM3CSK4_VS_GADIQUIMOD_6H_BMDC_UP | ART_long | ART | long | 0 | 0 | 0 | 0 | 1 | 0 | 2 | 7 | 4.422633136 | 9.761479141 | 26.61134211 | 520 | 45 |
| GSE17721_PAM3CSK4_VS_GADIQUIMOD_8H_BMDC_DN | ART_long | ART | long | 0 | 0 | 0 | 0 | 1 | 0 | 1 | 7 | 4.422633136 | 14.64341432 | 24.93943212 | 519 | 46 |
| GSE18893_CTRL_VS_TNF_TREATED_TREG_2H_DN | ART_long | ART | long | 1 | 0 | 0 | 0 | 0 | 0 | 1 | 7 | 4.422633136 | 22.63651394 | 25.22830336 | 301 | 47 |
| GSE1925_3H_VS_24H_IFNG_STIM_IFNG_PRIMED_MACROPHAGE_DN | ART_long | ART | long | 0 | 0 | 0 | 0 | 1 | 1 | 1 | 7 | 4.422633136 | 15.87297192 | 35.13921845 | 601 | 48 |
| GSE20198_UNTREATED_VS_IL12_IL18_TREATED_ACT_CD4_TCELL_UP | ART_long | ART | long | 0 | 1 | 0 | 0 | 0 | 1 | 2 | 9 | 5.714816687 | 29.65064703 | 13.30628491 | 462 | 49 |
| GSE20484_MCSG_VS_CXCL4_MONOCYTE_DERIVED_MACROPHAGE_DN | ART_long | ART | long | 0 | 0 | 0 | 0 | 1 | 1 | 1 | 9 | 5.832043696 | 12.01018956 | 23.68873717 | 940 | 50 |
| GSE20715_WT_VS_TLR4_KO_LUNG_UP | ART_long | ART | long | 0 | 0 | 0 | 0 | 0 | 0 | 2 | 8 | 5.05443787 | 12.58389887 | 17.78434166 | 797 | 51 |
| GSE21033_1H_VS_12H_POLYIC_STIM_DC_DN | ART_long | ART | long | 0 | 0 | 0 | 0 | 1 | 0 | 1 | 6 | 5.377061564 | 10.45687988 | 21.46537295 | 539 | 52 |
| GSE21063_CTRL_VS_ANTI_IGM_STIM_BCELL_NFATC1_KO_3H_DN | ART_long | ART | long | 0 | 0 | 0 | 1 | 0 | 1 | 1 | 7 | 4.704928868 | 4.627574425 | 22.32130335 | 500 | 53 |
| GSE21546_WT_VS_SAP1A_KO_AND_ELK1_KO_DP_THYMOCYTES_DN | ART_long | ART | long | 1 | 0 | 0 | 0 | 0 | 0 | 1 | 9 | 5.686242604 | 6.555934412 | 33.59984141 | 1255 | 54 |
| GSE21546_WT_VS_SAP1A_KO_ANTI_CD3_STIM_DP_THYMOCYTES_DN | ART_long | ART | long | 1 | 0 | 0 | 0 | 0 | 0 | 1 | 7 | 4.422633136 | 21.56596233 | 29.34273482 | 470 | 55 |
| GSE21670_TGFB_VS_IL6_TREATED_STAT3_KO_CD4_TCELL_DN | ART_long | ART | long | 0 | 1 | 0 | 0 | 0 | 1 | 1 | 7 | 4.422633136 | 19.52953292 | 30.64508527 | 634 | 56 |
| GSE21670_UNTREATED_VS_IL6_TREATED_CD4_TCELL_DN | ART_long | ART | long | 0 | 1 | 0 | 0 | 0 | 1 | 1 | 7 | 4.422633136 | 17.02673978 | 34.36768429 | 1237 | 57 |
| GSE21774_CD62L_POS_CD56_BRIGHT_VS_CD62L_NEG_CD56_DIM_NK_CELL_DN | ART_long | ART | long | 1 | 0 | 0 | 0 | 0 | 0 | 1 | 7 | 4.422633136 | 4.721276512 | 21.40072439 | 778 | 58 |
| GSE21774_CD62L_POS_CD56_DIM_VS_CD62L_NEG_CD56_DIM_NK_CELL_UP | ART_long | ART | long | 1 | 0 | 0 | 0 | 0 | 0 | 2 | 7 | 4.444857423 | 5.873992544 | 23.54594379 | 821 | 59 |
| GSE21927_BALBC_VS_C57BL6_MONOCYTE_SPLEEN_UP | ART_long | ART | long | 0 | 0 | 0 | 0 | 1 | 0 | 1 | 7 | 4.680035065 | 7.70482472 | 18.09381145 | 736 | 60 |
| GSE21927_C26GM_VS_4T1_TUMOR_MONOCYTE_BALBC_DN | ART_long | ART | long | 0 | 0 | 0 | 0 | 1 | 0 | 1 | 8 | 5.079837055 | 6.106218169 | 28.85217428 | 818 | 61 |
| GSE22342_CD11C_HIGH_VS_LOW_DECIDUAL_MACROPHAGES_UP | ART_long | ART | long | 0 | 0 | 0 | 0 | 1 | 0 | 2 | 7 | 4.467306198 | 38.84919179 | 25.78448358 | 601 | 62 |
| GSE22886_CD4_TCELL_VS_BCELL_NAIVE_UP | ART_long | ART | long | 0 | 1 | 0 | 1 | 0 | 0 | 2 | 8 | 5.105492798 | 36.85938105 | 20.8609348 | 1116 | 63 |
| GSE22886_NAIVE_CD4_TCELL_VS_12H_ACT_TH1_UP | ART_long | ART | long | 1 | 1 | 0 | 0 | 0 | 0 | 2 | 7 | 4.444857423 | 12.48202529 | 36.14283751 | 464 | 64 |
| GSE22886_NAIVE_CD4_TCELL_VS_12H_ACT_TH2_UP | ART_long | ART | long | 0 | 1 | 0 | 0 | 0 | 0 | 2 | 8 | 5.079837055 | 35.28041707 | 27.92211281 | 557 | 65 |
| GSE22886_NAIVE_CD8_TCELL_VS_DC_UP | ART_long | ART | long | 0 | 0 | 1 | 0 | 1 | 0 | 2 | 7 | 4.444857423 | 20.17394065 | 33.79601845 | 1003 | 66 |
| GSE22886_NAIVE_CD8_TCELL_VS_MONOCYTE_UP | ART_long | ART | long | 0 | 0 | 1 | 0 | 1 | 0 | 2 | 7 | 4.444857423 | 14.15550525 | 23.78352046 | 837 | 67 |
| GSE22886_TCELL_VS_BCELL_NAIVE_UP | ART_long | ART | long | 1 | 0 | 0 | 1 | 0 | 0 | 2 | 10 | 6.349796319 | 34.7375274 | 20.03829307 | 1293 | 68 |
| GSE22886_UNSTIM_VS_STIM_MEMORY_TCELL_UP | ART_long | ART | long | 1 | 0 | 0 | 0 | 0 | 0 | 1 | 7 | 4.467306198 | 7.93105039 | 19.51122007 | 788 | 69 |
| GSE23505_UNTREATED_VS_4DAY_IL6_IL1_TGFB_TREATED_CD4_TCELL_UP | ART_long | ART | long | 0 | 1 | 0 | 0 | 0 | 1 | 2 | 8 | 5.079837055 | 2.984468048 | 16.25948843 | 1061 | 70 |
| GSE23568_CTRL_VS_ID3_TRANSDUCED_CD8_TCELL_UP | ART_long | ART | long | 0 | 0 | 1 | 0 | 0 | 0 | 2 | 7 | 4.422633136 | 12.82632719 | 20.4206426 | 539 | 71 |
| GSE24026_PD1_LIGATION_VS_CTRL_IN_ACT_TCELL_LINE_DN | ART_long | ART | long | 1 | 0 | 0 | 0 | 0 | 0 | 1 | 8 | 5.05443787 | 19.86231731 | 17.50272867 | 482 | 72 |
| GSE24142_ADULT_VS_FETAL_DN3_THYMOCYTE_UP | ART_long | ART | long | 1 | 0 | 0 | 0 | 0 | 0 | 2 | 7 | 4.422633136 | 9.004747459 | 20.04814945 | 913 | 73 |
| GSE24142_ADULT_VS_FETAL_EARLY_THYMIC_PROGENITOR_UP | ART_long | ART | long | 1 | 0 | 0 | 0 | 0 | 0 | 2 | 7 | 4.422633136 | 7.424740848 | 29.41997393 | 1095 | 74 |
| GSE24210_CTRL_VS_IL35_TREATED_TCONV_CD4_TCELL_UP | ART_long | ART | long | 0 | 1 | 0 | 0 | 0 | 1 | 2 | 9 | 5.686242604 | 21.39353271 | 27.33476598 | 958 | 75 |
| GSE24210_RESTING_TREG_VS_TCONV_UP | ART_long | ART | long | 1 | 0 | 0 | 0 | 0 | 0 | 2 | 8 | 5.05443787 | 38.31804476 | 34.75638504 | 793 | 76 |
| GSE24671_CTRL_VS_SENDAI_VIRUS_INFECTED_MOUSE_SPLENOCYTES_DN | ART_long | ART | long | 1 | 0 | 0 | 0 | 0 | 0 | 1 | 8 | 5.079837055 | 33.61348102 | 33.05169999 | 1010 | 77 |
| GSE25087_TREG_VS_TCONV_FETUS_UP | ART_long | ART | long | 1 | 0 | 0 | 0 | 0 | 0 | 2 | 7 | 4.467306198 | 42.69832348 | 35.31387048 | 1055 | 78 |
| GSE25677_MPL_VS_R848_STIM_BCELL_DN | ART_long | ART | long | 0 | 0 | 0 | 1 | 0 | 0 | 1 | 9 | 6.283140998 | 10.68337906 | 27.83080687 | 1269 | 79 |
| GSE26030_TH1_VS_TH17_DAY5_POST_POLARIZATION_DN | ART_long | ART | long | 1 | 0 | 0 | 0 | 0 | 0 | 1 | 8 | 5.131409005 | 38.78661428 | 36.24492344 | 939 | 80 |
| GSE26488_CTRL_VS_PEPTIDE_INJECTION_OT2_THYMOCYTE_DN | ART_long | ART | long | 1 | 0 | 0 | 0 | 0 | 0 | 1 | 9 | 6.214472791 | 12.91040433 | 24.85308343 | 1221 | 81 |
| GSE26495_NAIVE_VS_PD1LOW_CD8_TCELL_DN | ART_long | ART | long | 0 | 0 | 1 | 0 | 0 | 0 | 1 | 8 | 5.079837055 | 35.64902647 | 25.87417409 | 772 | 82 |
| GSE27291_0H_VS_7D_STIM_GAMMADELTA_TCELL_DN | ART_long | ART | long | 1 | 0 | 0 | 0 | 0 | 0 | 1 | 8 | 5.320460916 | 48.14964903 | 20.86782639 | 959 | 83 |
| GSE2770_UNTREATED_VS_IL12_TREATED_ACT_CD4_TCELL_2H_UP | ART_long | ART | long | 0 | 1 | 0 | 0 | 0 | 1 | 2 | 11 | 6.949852071 | 13.57523001 | 26.33471517 | 1025 | 84 |
| GSE2770_UNTREATED_VS_TGFB_AND_IL12_TREATED_ACT_CD4_TCELL_6H_UP | ART_long | ART | long | 0 | 1 | 0 | 0 | 0 | 1 | 2 | 7 | 4.422633136 | 20.79790224 | 19.36230854 | 366 | 85 |
| GSE27786_ERYTHROBLAST_VS_MONO_MAC_DN | ART_long | ART | long | 1 | 0 | 0 | 0 | 1 | 0 | 1 | 8 | 5.05443787 | 5.496296876 | 24.66029695 | 570 | 86 |
| GSE27786_ERYTHROBLAST_VS_NEUTROPHIL_DN | ART_long | ART | long | 1 | 0 | 0 | 0 | 0 | 0 | 1 | 7 | 4.422633136 | 6.945514458 | 24.24827488 | 430 | 87 |
| GSE27786_LIN_NEG_VS_CD4_TCELL_DN | ART_long | ART | long | 0 | 1 | 0 | 0 | 0 | 0 | 1 | 8 | 5.05443787 | 79.61258811 | 27.85218187 | 828 | 88 |
| GSE27786_LIN_NEG_VS_CD8_TCELL_DN | ART_long | ART | long | 0 | 0 | 1 | 0 | 0 | 0 | 1 | 7 | 4.422633136 | 785.9996791 | 21.11022898 | 403 | 89 |
| GSE27896_HDAC6_KO_VS_WT_TREG_UP | ART_long | ART | long | 1 | 0 | 0 | 0 | 0 | 0 | 2 | 8 | 5.743679398 | 9.349585298 | 21.95299459 | 371 | 90 |
| GSE28726_ACT_CD4_TCELL_VS_ACT_NKTCELL_DN | ART_long | ART | long | 1 | 1 | 0 | 0 | 0 | 0 | 1 | 8 | 5.079837055 | 17.49274571 | 18.92464524 | 718 | 91 |
| GSE29618_PRE_VS_DAY7_POST_TIV_FLU_VACCINE_MONOCYTE_UP | ART_long | ART | long | 0 | 0 | 0 | 0 | 1 | 0 | 2 | 7 | 4.467306198 | 5.257011369 | 18.70855255 | 373 | 92 |
| GSE29618_PRE_VS_DAY7_POST_TIV_FLU_VACCINE_PDC_DN | ART_long | ART | long | 0 | 0 | 0 | 0 | 0 | 0 | 1 | 8 | 5.05443787 | 11.81377205 | 27.8091197 | 1038 | 93 |
| GSE29949_MICROGLIA_BRAIN_VS_MONOCYTE_BONE_MARROW_DN | ART_long | ART | long | 1 | 0 | 0 | 0 | 1 | 0 | 1 | 8 | 5.05443787 | 14.53314589 | 27.36654091 | 653 | 94 |
| GSE30083_SP2_VS_SP4_THYMOCYTE_DN | ART_long | ART | long | 1 | 0 | 0 | 0 | 0 | 0 | 1 | 7 | 4.422633136 | 14.51497108 | 36.24738146 | 621 | 95 |
| GSE30962_ACUTE_VS_CHRONIC_LCMV_SECONDARY_INF_CD8_TCELL_UP | ART_long | ART | long | 0 | 0 | 1 | 0 | 0 | 1 | 2 | 8 | 5.05443787 | 9.806745996 | 46.01140718 | 1392 | 96 |
| GSE31082_DN_VS_CD8_SP_THYMOCYTE_UP | ART_long | ART | long | 1 | 0 | 0 | 0 | 0 | 0 | 2 | 7 | 4.422633136 | 8.143894099 | 20.15590355 | 400 | 97 |
| GSE32423_IL7_VS_IL7_IL4_NAIVE_CD8_TCELL_DN | ART_long | ART | long | 0 | 0 | 1 | 0 | 0 | 1 | 1 | 7 | 4.422633136 | 32.08548322 | 12.72228447 | 263 | 98 |
| GSE32986_UNSTIM_VS_CURDLAN_LOWDOSE_STIM_DC_DN | ART_long | ART | long | 0 | 0 | 0 | 0 | 1 | 0 | 1 | 8 | 5.05443787 | 36.79117266 | 20.18269013 | 617 | 99 |
| GSE33162_HDAC3_KO_VS_HDAC3_KO_4H_LPS_STIM_MACROPHAGE_UP | ART_long | ART | long | 0 | 0 | 0 | 0 | 1 | 0 | 2 | 7 | 4.422633136 | 33.71772471 | 30.80830898 | 399 | 100 |

| | | | | | | | | | | | | | | | | |
|---|---|---|---|---|---|---|---|---|---|---|---|---|---|---|---|---|
| GSE339_CD4POS_VS_CD4CD8DN_DC_UP | ART_long | ART | long | 0 | 0 | 0 | 0 | 1 | 0 | 2 | 7 | 4.422633136 | 41.34668398 | 22.41387326 | 587 | 101 |
| GSE34205_RSV_VS_FLU_INF_INFANT_PBMC_DN | ART_long | ART | long | 0 | 0 | 0 | 0 | 1 | 0 | 1 | 7 | 4.536033986 | 45.37403953 | 29.90340614 | 351 | 102 |
| GSE35685_CD34POS_CD38NEG_VS_CD34POS_CD10NEG_CD62LPOS_BONE_MARROW_UP | ART_long | ART | long | 0 | 0 | 0 | 0 | 0 | 0 | 2 | 7 | 4.422633136 | 15.87879358 | 19.78803029 | 168 | 103 |
| GSE35825_IFNA_VS_IFNG_STIM_MACROPHAGE_DN | ART_long | ART | long | 0 | 0 | 0 | 0 | 1 | 1 | 1 | 7 | 4.422633136 | 7.958323015 | 38.65041695 | 806 | 104 |
| GSE360_DC_VS_MAC_L_DONOVANI_DN | ART_long | ART | long | 0 | 0 | 0 | 0 | 1 | 0 | 1 | 9 | 5.686242604 | 43.46508461 | 18.92569069 | 640 | 105 |
| GSE360_L_DONOVANI_VS_B_MALAYI_LOW_DOSE_DC_DN | ART_long | ART | long | 0 | 0 | 0 | 0 | 1 | 0 | 1 | 7 | 4.422633136 | 65.11200593 | 27.30576237 | 663 | 106 |
| GSE36891_UNSTIM_VS_PAM_TLR2_STIM_PERITONEAL_MACROPHAGE_UP | ART_long | ART | long | 0 | 0 | 0 | 0 | 1 | 0 | 2 | 7 | 4.422633136 | 8.340571192 | 25.510101 | 917 | 107 |
| GSE369_IFNG_KO_VS_WT_LIVER_UP | ART_long | ART | long | 0 | 0 | 0 | 0 | 0 | 1 | 2 | 8 | 5.079837055 | 13.33337076 | 29.01249193 | 1101 | 108 |
| GSE37301_LYMPHOID_PRIMED_MPP_VS_GRAN_MONO_PROGENITOR_DN | ART_long | ART | long | 1 | 0 | 0 | 0 | 0 | 0 | 1 | 7 | 4.467306198 | 19.97108599 | 29.18757072 | 549 | 109 |
| GSE37301_MULTIPOTENT_PROGENITOR_VS_COMMON_LYMPHOID_PROGENITOR_UP | ART_long | ART | long | 1 | 0 | 0 | 0 | 0 | 0 | 2 | 7 | 5.265039448 | 12.55966546 | 19.47947628 | 1001 | 110 |
| GSE37416_0H_VS_12H_F_TULARENSIS_LVS_NEUTROPHIL_DN | ART_long | ART | long | 1 | 0 | 0 | 0 | 0 | 0 | 1 | 7 | 4.444857423 | 11.2238346 | 17.5198513 | 253 | 111 |
| GSE37532_WT_VS_PPARG_KO_LN_TCONV_UP | ART_long | ART | long | 1 | 0 | 0 | 0 | 0 | 0 | 2 | 9 | 5.686242604 | 10.57406405 | 21.9088994 | 988 | 112 |
| GSE37563_WT_VS_CTLA4_KO_CD4_TCELL_D4_POST_IMMUNIZATION_UP | ART_long | ART | long | 0 | 1 | 0 | 0 | 0 | 0 | 2 | 7 | 5.360767438 | 7.400047197 | 35.81601099 | 399 | 113 |
| GSE38696_LIGHT_ZONE_VS_DARK_ZONE_BCELL_UP | ART_long | ART | long | 0 | 0 | 0 | 1 | 0 | 0 | 2 | 10 | 6.867442758 | 6.102529285 | 27.81001355 | 1014 | 114 |
| GSE3920_IFNB_VS_IFNG_TREATED_ENDOTHELIAL_CELL_UP | ART_long | ART | long | 1 | 0 | 0 | 0 | 0 | 1 | 2 | 7 | 5.29656663 | 45.1270438 | 15.57450076 | 380 | 115 |
| GSE3982_MEMORY_CD4_TCELL_VS_BCELL_UP | ART_long | ART | long | 0 | 1 | 0 | 1 | 0 | 0 | 2 | 8 | 5.079837055 | 14.19278586 | 30.69570086 | 648 | 116 |
| GSE3982_NEUTROPHIL_VS_CENT_MEMORY_CD4_TCELL_UP | ART_long | ART | long | 1 | 1 | 0 | 0 | 0 | 0 | 2 | 7 | 4.422633136 | 9.59598859 | 20.53474347 | 492 | 117 |
| GSE40068_BCL6_POS_VS_NEG_CXCR5_POS_TFH_DN | ART_long | ART | long | 1 | 0 | 0 | 0 | 0 | 1 | 1 | 7 | 4.444857423 | 4.450039115 | 28.00146412 | 721 | 118 |
| GSE40273_EOS_KO_VS_WT_TREG_DN | ART_long | ART | long | 1 | 0 | 0 | 0 | 0 | 0 | 1 | 7 | 4.444857423 | 19.66935986 | 27.79238431 | 758 | 119 |
| GSE40273_GATA1_KO_VS_WT_TREG_DN | ART_long | ART | long | 1 | 0 | 0 | 0 | 0 | 0 | 1 | 8 | 5.079837055 | 13.25629316 | 31.05420312 | 1413 | 120 |
| GSE40666_UNTREATED_VS_IFNA_STIM_STAT4_KO_EFFECTOR_CD8_TCELL_90MIN_DN | ART_long | ART | long | 0 | 0 | 1 | 0 | 0 | 1 | 1 | 7 | 4.444857423 | 4.866832018 | 27.57629746 | 821 | 121 |
| GSE42021_CD24LO_TREG_VS_CD24LO_TCONV_THYMUS_UP | ART_long | ART | long | 1 | 0 | 0 | 0 | 0 | 0 | 2 | 7 | 4.422633136 | 9.372532187 | 35.18864805 | 739 | 122 |
| GSE43955_TH0_VS_TGFB_IL6_TH17_ACT_CD4_TCELL_60H_UP | ART_long | ART | long | 0 | 1 | 0 | 0 | 0 | 1 | 2 | 7 | 4.422633136 | 12.18648131 | 21.57262965 | 809 | 123 |
| GSE45365_NK_CELL_VS_CD8_TCELL_MCMV_INFECTION_UP | ART_long | ART | long | 1 | 0 | 1 | 0 | 0 | 0 | 2 | 7 | 4.467306198 | 3.919139894 | 20.18362354 | 966 | 124 |
| GSE45739_UNSTIM_VS_ACD3_ACD28_STIM_NRAS_KO_CD4_TCELL_UP | ART_long | ART | long | 0 | 1 | 0 | 0 | 0 | 0 | 2 | 10 | 6.318047337 | 8.2946795 | 23.28762347 | 755 | 125 |
| GSE45739_UNSTIM_VS_ACD3_ACD28_STIM_WT_CD4_TCELL_DN | ART_long | ART | long | 0 | 1 | 0 | 0 | 0 | 0 | 1 | 8 | 5.079837055 | 39.53142342 | 32.10041126 | 1096 | 126 |
| GSE45739_UNSTIM_VS_ACD3_ACD28_STIM_WT_CD4_TCELL_UP | ART_long | ART | long | 0 | 1 | 0 | 0 | 0 | 0 | 2 | 7 | 4.422633136 | 5.102682028 | 14.337136 | 481 | 127 |
| GSE4984_LPS_VS_VEHICLE_CTRL_TREATED_DC_DN | ART_long | ART | long | 0 | 0 | 0 | 0 | 1 | 0 | 1 | 7 | 4.467306198 | 9.625126461 | 22.09262041 | 706 | 128 |
| GSE5542_UNTREATED_VS_IFNA_TREATED_EPITHELIAL_CELLS_6H_UP | ART_long | ART | long | 1 | 0 | 0 | 0 | 0 | 1 | 2 | 9 | 5.714816687 | 22.49691535 | 30.3460105 | 686 | 129 |
| GSE5542_UNTREATED_VS_IFNG_TREATED_EPITHELIAL_CELLS_24H_UP | ART_long | ART | long | 1 | 0 | 0 | 0 | 0 | 1 | 2 | 8 | 5.105492798 | 13.87875831 | 27.85481328 | 740 | 130 |
| GSE5542_UNTREATED_VS_IFNG_TREATED_EPITHELIAL_CELLS_6H_DN | ART_long | ART | long | 1 | 0 | 0 | 0 | 0 | 1 | 1 | 7 | 4.422633136 | 2.009701022 | 39.22467992 | 702 | 131 |
| GSE5589_LPS_AND_IL10_VS_LPS_AND_IL6_STIM_IL10_KO_MACROPHAGE_45MIN_DN | ART_long | ART | long | 0 | 0 | 0 | 0 | 1 | 1 | 1 | 7 | 4.422633136 | 65.89094458 | 24.16183082 | 689 | 132 |
| GSE5589_UNSTIM_VS_45MIN_LPS_AND_IL10_STIM_MACROPHAGE_UP | ART_long | ART | long | 0 | 0 | 0 | 0 | 1 | 1 | 2 | 7 | 4.422633136 | 25.25255763 | 11.74260228 | 403 | 133 |
| GSE5589_UNSTIM_VS_45MIN_LPS_AND_IL6_STIM_MACROPHAGE_UP | ART_long | ART | long | 0 | 0 | 0 | 0 | 1 | 1 | 2 | 7 | 4.422633136 | 12.90316737 | 37.53985622 | 556 | 134 |
| GSE5679_CTRL_VS_PPARG_LIGAND_ROSIGLITAZONE_TREATED_DC_DN | ART_long | ART | long | 0 | 0 | 0 | 0 | 1 | 0 | 1 | 7 | 4.422633136 | 14.93113721 | 38.97160963 | 993 | 135 |
| GSE6092_IFNG_VS_IFNG_AND_B_BURGDORFERI_INF_ENDOTHELIAL_CELL_UP | ART_long | ART | long | 1 | 0 | 0 | 0 | 0 | 1 | 2 | 8 | 5.616042078 | 9.86603601 | 31.47240178 | 1046 | 136 |
| GSE6092_UNSTIM_VS_IFNG_STIM_ENDOTHELIAL_CELL_DN | ART_long | ART | long | 1 | 0 | 0 | 0 | 0 | 1 | 1 | 8 | 5.079837055 | 13.30411436 | 28.13417343 | 1243 | 137 |
| GSE7219_UNSTIM_VS_LPS_AND_ANTI_CD40_STIM_NIK_NFKB2_KO_DC_UP | ART_long | ART | long | 0 | 0 | 0 | 0 | 1 | 1 | 2 | 7 | 4.969250715 | 14.79957321 | 32.18070044 | 921 | 138 |
| GSE7348_UNSTIM_VS_TOLERIZED_AND_LPS_STIM_MACROPHAGE_UP | ART_long | ART | long | 0 | 0 | 0 | 0 | 1 | 0 | 2 | 7 | 5.460040909 | 31.3117176 | 36.14255495 | 782 | 139 |
| GSE7460_FOXP3_MUT_VS_WT_ACT_TCONV_DN | ART_long | ART | long | 1 | 0 | 0 | 0 | 0 | 0 | 1 | 8 | 5.105492798 | 18.28935018 | 24.637471 | 918 | 140 |
| GSE7460_FOXP3_MUT_VS_WT_ACT_WITH_TGFB_TCONV_DN | ART_long | ART | long | 1 | 0 | 0 | 0 | 0 | 1 | 1 | 8 | 5.05443787 | 11.75672868 | 22.80772188 | 1058 | 141 |
| GSE7460_WT_VS_FOXP3_HET_ACT_WITH_TGFB_TCONV_UP | ART_long | ART | long | 1 | 0 | 0 | 0 | 0 | 1 | 2 | 9 | 5.686242604 | 7.689726473 | 24.82338049 | 1083 | 142 |
| GSE7568_IL4_VS_IL4_AND_DEXAMETHASONE_TREATED_MACROPHAGE_DN | ART_long | ART | long | 0 | 0 | 0 | 0 | 1 | 1 | 1 | 7 | 5.857792233 | 10.593182 | 33.55267388 | 897 | 143 |
| GSE7764_IL15_NK_CELL_24H_VS_SPLENOCYTE_DN | ART_long | ART | long | 1 | 0 | 0 | 0 | 0 | 1 | 1 | 7 | 4.444857423 | 16.43291943 | 31.21112033 | 908 | 144 |
| GSE7768_OVA_ALONE_VS_OVA_WITH_LPS_IMMUNIZED_MOUSE_WHOLE_SPLEEN_6H_DN | ART_long | ART | long | 0 | 0 | 0 | 0 | 0 | 0 | 1 | 8 | 5.946397494 | 1.970686256 | 22.14575854 | 976 | 145 |
| GSE8685_IL2_STARVED_VS_IL2_ACT_IL2_STARVED_CD4_TCELL_UP | ART_long | ART | long | 0 | 1 | 0 | 0 | 0 | 1 | 2 | 8 | 5.079837055 | 19.30749679 | 28.76633629 | 917 | 146 |
| GSE9006_HEALTHY_VS_TYPE_1_DIABETES_PBMC_1MONTH_POST_DX_UP | ART_long | ART | long | 0 | 0 | 0 | 0 | 1 | 0 | 2 | 7 | 4.422633136 | 43.25244184 | 14.60860601 | 523 | 147 |
| GSE9037_CTRL_VS_LPS_4H_STIM_BMDM_DN | ART_long | ART | long | 0 | 0 | 0 | 0 | 1 | 0 | 1 | 8 | 5.05443787 | 56.93761895 | 19.30697447 | 782 | 148 |
| GSE9037_WT_VS_IRAK4_KO_LPS_4H_STIM_BMDM_UP | ART_long | ART | long | 0 | 0 | 0 | 0 | 1 | 0 | 2 | 7 | 4.422633136 | 14.84437741 | 25.34920943 | 951 | 149 |
| GSE9650_EFFECTOR_VS_MEMORY_CD8_TCELL_DN | ART_long | ART | long | 0 | 0 | 1 | 0 | 0 | 0 | 1 | 7 | 4.422633136 | 55.82344544 | 22.03083774 | 523 | 150 |
| NAKAYA_MONOCYTE_FLUMIST_AGE_18_50YO_7DY_UP | ART_long | ART | long | 0 | 0 | 0 | 0 | 1 | 0 | 2 | 13 | 3.436594786 | 10.69304678 | 29.52529138 | 797 | 151 |
| NAKAYA_PBMC_FLUARIX_FLUVIRIN_AGE_18_50YO_CORRELATED_WITH_HAI_28DY_RESPONSE_AT_3DY_NEGATIVE | ART_long | ART | long | 0 | 0 | 0 | 0 | 1 | 0 | 0 | 14 | 4.860036413 | 395.5355536 | 14.48144261 | 1362 | 152 |
| NAKAYA_PBMC_FLUARIX_FLUVIRIN_AGE_18_50YO_CORRELATED_WITH_HAI_28DY_RESPONSE_AT_7DY_NEGATIVE | ART_long | ART | long | 0 | 0 | 0 | 0 | 1 | 0 | 0 | 13 | 3.458299595 | 423.9811708 | 16.1054519 | 1185 | 153 |
| GSE10147_IL3_VS_IL3_AND_HIVP17_STIM_PDC_UP | ART_short | ART | short | 0 | 0 | 0 | 0 | 1 | 1 | 2 | 7 | 6.621412119 | 5.104864389 | 34.65971841 | 1156 | 154 |
| GSE10273_HIGH_VS_LOW_IL7_TREATED_IRF4_8_NULL_PRE_BCELL_DN | ART_short | ART | short | 0 | 0 | 0 | 1 | 0 | 1 | 1 | 7 | 5.551284908 | 21.48039134 | 47.54301201 | 319 | 155 |
| GSE10325_CD4_TCELL_VS_BCELL_UP | ART_short | ART | short | 0 | 1 | 0 | 1 | 0 | 0 | 2 | 11 | 8.723447712 | 22.13423943 | 33.71549329 | 1068 | 156 |
| GSE10325_CD4_TCELL_VS_MYELOID_UP | ART_short | ART | short | 0 | 1 | 0 | 0 | 1 | 0 | 2 | 12 | 9.468666864 | 20.99937944 | 31.11756831 | 1393 | 157 |
| GSE10325_LUPUS_CD4_TCELL_VS_LUPUS_BCELL_UP | ART_short | ART | short | 0 | 1 | 0 | 1 | 0 | 0 | 2 | 11 | 8.679611292 | 23.47540037 | 28.29355225 | 1036 | 158 |
| GSE10325_LUPUS_CD4_TCELL_VS_LUPUS_MYELOID_UP | ART_short | ART | short | 0 | 1 | 0 | 0 | 1 | 0 | 2 | 8 | 6.312444576 | 16.46100631 | 30.12739472 | 698 | 159 |
| GSE11057_CD4_CENT_MEM_VS_PBMC_UP | ART_short | ART | short | 0 | 1 | 0 | 0 | 1 | 0 | 2 | 7 | 5.579464019 | 22.36108073 | 20.71748913 | 871 | 160 |
| GSE11057_EFF_MEM_VS_CENT_MEM_CD4_TCELL_DN | ART_short | ART | short | 0 | 1 | 0 | 0 | 0 | 0 | 1 | 7 | 5.84656602 | 13.48454258 | 26.48958178 | 1080 | 161 |
| GSE11961_GERMINAL_CENTER_BCELL_DAY7_VS_PLASMA_CELL_DAY7_UP | ART_short | ART | short | 0 | 0 | 0 | 1 | 0 | 0 | 2 | 7 | 5.495772059 | 13.14466423 | 44.62626996 | 574 | 162 |
| GSE11961_MARGINAL_ZONE_BCELL_VS_GERMINAL_CENTER_BCELL_DAY40_UP | ART_short | ART | short | 0 | 0 | 0 | 1 | 0 | 0 | 2 | 8 | 6.312444576 | 13.5125517 | 39.88633667 | 739 | 163 |
| GSE11961_MARGINAL_ZONE_BCELL_VS_GERMINAL_CENTER_BCELL_DAY7_DN | ART_short | ART | short | 0 | 0 | 0 | 1 | 0 | 0 | 1 | 7 | 5.495772059 | 2.500288956 | 24.70854874 | 471 | 164 |
| GSE12366_GC_BCELL_VS_PLASMA_CELL_UP | ART_short | ART | short | 0 | 0 | 0 | 1 | 0 | 0 | 2 | 9 | 7.173596596 | 8.85686888 | 23.65853835 | 976 | 165 |
| GSE12392_WT_VS_IFNB_KO_CD8A_NEG_SPLEEN_DC_DN | ART_short | ART | short | 0 | 0 | 0 | 0 | 1 | 1 | 1 | 7 | 5.495772059 | 24.48972469 | 38.61909807 | 580 | 166 |
| GSE12845_IGD_POS_BLOOD_VS_DARKZONE_GC_TONSIL_BCELL_DN | ART_short | ART | short | 0 | 0 | 0 | 1 | 0 | 0 | 1 | 8 | 6.312444576 | 10.83093173 | 30.03406914 | 1096 | 167 |
| GSE13411_IGM_MEMORY_BCELL_VS_PLASMA_CELL_UP | ART_short | ART | short | 0 | 0 | 0 | 1 | 0 | 1 | 2 | 8 | 6.344325609 | 14.87659795 | 19.69956455 | 682 | 168 |
| GSE13411_SWITCHED_MEMORY_BCELL_VS_PLASMA_CELL_UP | ART_short | ART | short | 0 | 0 | 0 | 1 | 0 | 0 | 2 | 8 | 6.280882353 | 12.72533397 | 23.02519194 | 569 | 169 |
| GSE14000_4H_VS_16H_LPS_DC_TRANSLATED_RNA_UP | ART_short | ART | short | 0 | 0 | 0 | 0 | 1 | 0 | 2 | 7 | 5.607930672 | 9.173880714 | 38.31217763 | 638 | 170 |
| GSE14000_UNSTIM_VS_16H_LPS_DC_TRANSLATED_RNA_UP | ART_short | ART | short | 0 | 0 | 0 | 0 | 1 | 0 | 2 | 7 | 5.579464019 | 9.799497241 | 36.25495145 | 625 | 171 |
| GSE1460_INTRATHYMIC_T_PROGENITOR_VS_CD4_THYMOCYTE_DN | ART_short | ART | short | 1 | 1 | 0 | 0 | 0 | 0 | 1 | 7 | 5.523389004 | 23.40276859 | 30.26737016 | 631 | 172 |
| GSE15330_WT_VS_IKAROS_KO_HSC_UP | ART_short | ART | short | 1 | 0 | 0 | 0 | 0 | 0 | 2 | 7 | 5.495772059 | 30.29637551 | 31.9577386 | 518 | 173 |
| GSE16450_IMMATURE_VS_MATURE_NEURON_CELL_LINE_UP | ART_short | ART | short | 1 | 0 | 0 | 0 | 0 | 0 | 2 | 7 | 5.636689291 | 32.74479042 | 27.85563586 | 799 | 174 |
| GSE16522_ANTI_CD3CD28_STIM_VS_UNSTIM_NAIVE_CD8_TCELL_DN | ART_short | ART | short | 0 | 0 | 1 | 0 | 0 | 0 | 1 | 8 | 6.344325609 | 24.32515812 | 43.31851286 | 565 | 175 |
| GSE17721_0.5H_VS_12H_GARDIQUIMOD_BMDC_DN | ART_short | ART | short | 0 | 0 | 0 | 0 | 1 | 0 | 1 | 8 | 6.280882353 | 13.49678425 | 39.96179633 | 781 | 176 |
| GSE1925_CTRL_VS_IFNG_PRIMED_MACROPHAGE_3H_IFNG_STIM_UP | ART_short | ART | short | 0 | 0 | 0 | 0 | 1 | 1 | 2 | 7 | 6.316979378 | 25.30513473 | 18.80075622 | 442 | 177 |
| GSE21033_CTRL_VS_POLYIC_STIM_DC_24H_UP | ART_short | ART | short | 0 | 0 | 0 | 0 | 1 | 0 | 2 | 6 | 6.039309955 | 50.77826474 | 15.99689676 | 561 | 178 |
| GSE21063_CTRL_VS_ANTI_IGM_STIM_BCELL_NFATC1_KO_3H_DN | ART_short | ART | short | 0 | 0 | 0 | 1 | 0 | 1 | 1 | 7 | 5.84656602 | 23.17511864 | 37.46629552 | 479 | 179 |
| GSE21063_CTRL_VS_ANTI_IGM_STIM_BCELL_NFATC1_KO_8H_UP | ART_short | ART | short | 0 | 0 | 0 | 1 | 0 | 1 | 2 | 7 | 5.75473514 | 31.10120452 | 22.60631818 | 711 | 180 |
| GSE21360_PRIMARY_VS_TERTIARY_MEMORY_CD8_TCELL_UP | ART_short | ART | short | 0 | 0 | 1 | 0 | 0 | 0 | 2 | 8 | 6.280882353 | 50.72707379 | 26.83085801 | 599 | 181 |
| GSE21670_UNTREATED_VS_IL6_TREATED_CD4_TCELL_DN | ART_short | ART | short | 0 | 1 | 0 | 0 | 0 | 1 | 1 | 7 | 5.495772059 | 27.16564023 | 22.76472932 | 763 | 182 |
| GSE21927_C26GM_VS_4T1_TUMOR_MONOCYTE_BALBC_DN | ART_short | ART | short | 0 | 0 | 0 | 0 | 1 | 0 | 1 | 8 | 6.312444576 | 21.71445066 | 22.97409742 | 830 | 183 |
| GSE21927_SPLEEN_C57BL6_VS_EL4_TUMOR_BALBC_MONOCYTES_UP | ART_short | ART | short | 0 | 0 | 0 | 0 | 1 | 0 | 2 | 7 | 5.523389004 | 19.91412703 | 24.86395167 | 833 | 184 |
| GSE22601_CD4_SINGLE_POSITIVE_VS_CD8_SINGLE_POSITIVE_THYMOCYTE_DN | ART_short | ART | short | 0 | 1 | 1 | 0 | 0 | 0 | 1 | 8 | 6.280882353 | 25.71766049 | 36.73351515 | 892 | 185 |
| GSE22886_CD4_TCELL_VS_BCELL_NAIVE_UP | ART_short | ART | short | 0 | 1 | 0 | 1 | 0 | 0 | 2 | 9 | 7.13736631 | 24.11646288 | 36.60523973 | 1216 | 186 |
| GSE22886_IGM_MEMORY_BCELL_VS_BM_PLASMA_CELL_UP | ART_short | ART | short | 0 | 0 | 0 | 1 | 0 | 1 | 2 | 7 | 5.495772059 | 12.49973516 | 27.84304264 | 591 | 187 |
| GSE22886_NAIVE_CD4_TCELL_VS_MONOCYTE_UP | ART_short | ART | short | 0 | 1 | 0 | 0 | 1 | 0 | 2 | 8 | 6.376530308 | 25.77575386 | 29.5087378 | 753 | 188 |
| GSE22886_NAIVE_CD8_TCELL_VS_MONOCYTE_UP | ART_short | ART | short | 0 | 0 | 1 | 0 | 1 | 0 | 2 | 8 | 6.312444576 | 20.93118722 | 36.42818862 | 920 | 189 |
| GSE22886_NAIVE_TCELL_VS_DC_UP | ART_short | ART | short | 1 | 0 | 0 | 0 | 1 | 0 | 2 | 7 | 5.523389004 | 18.53862584 | 25.72405674 | 653 | 190 |
| GSE22886_NAIVE_TCELL_VS_MONOCYTE_UP | ART_short | ART | short | 1 | 0 | 0 | 0 | 1 | 0 | 2 | 7 | 5.551284908 | 24.19056951 | 32.45649928 | 582 | 191 |
| GSE22886_TCELL_VS_BCELL_NAIVE_UP | ART_short | ART | short | 1 | 0 | 0 | 2 | 0 | 0 | 2 | 7 | 5.523389004 | 24.83757227 | 38.54797755 | 1040 | 192 |
| GSE24210_CTRL_VS_IL35_TREATED_TCONV_CD4_TCELL_UP | ART_short | ART | short | 0 | 1 | 0 | 0 | 0 | 1 | 2 | 10 | 7.851102941 | 19.8172836 | 45.58606033 | 685 | 193 |
| GSE24671_CTRL_VS_BAKIMULC_INFECTED_MOUSE_SPLENOCYTES_UP | ART_short | ART | short | 1 | 0 | 0 | 0 | 0 | 0 | 2 | 7 | 5.495772059 | 14.73294435 | 30.2452189 | 669 | 194 |
| GSE24671_CTRL_VS_SENDAI_VIRUS_INFECTED_MOUSE_SPLENOCYTES_DN | ART_short | ART | short | 1 | 0 | 0 | 0 | 0 | 0 | 1 | 7 | 5.523389004 | 12.6332133 | 40.24892534 | 512 | 195 |
| GSE25677_MPL_VS_MPL_AND_R848_STIM_BCELL_UP | ART_short | ART | short | 0 | 0 | 0 | 1 | 0 | 0 | 2 | 6 | 5.888327206 | 20.96897259 | 48.08996316 | 336 | 196 |
| GSE25677_MPL_VS_R848_STIM_BCELL_DN | ART_short | ART | short | 0 | 0 | 0 | 1 | 0 | 0 | 1 | 7 | 6.072676308 | 23.96375768 | 29.16215636 | 589 | 197 |

| | | | | | | | | | | | | | | | | |
|---|---|---|---|---|---|---|---|---|---|---|---|---|---|---|---|---|
| GSE27241_WT_VS_RORGT_KO_TH17_POLARIZED_CD4_TCELL_TREATED_WITH_DIGOXIN_UP | ART_short | ART | short | 0 | 1 | 0 | 0 | 0 | 0 | 2 | 9 | 7.851102941 | 19.02224175 | 33.78673015 | 1039 | 198 |
| GSE2770_TGFB_AND_IL4_VS_IL12_TREATED_ACT_CD4_TCELL_6H_UP | ART_short | ART | short | 0 | 1 | 0 | 0 | 0 | 1 | 2 | 8 | 6.280882353 | 36.59804794 | 32.85597951 | 643 | 199 |
| GSE31082_DN_VS_CD8_SP_THYMOCYTE_DN | ART_short | ART | short | 1 | 0 | 0 | 0 | 0 | 0 | 1 | 7 | 5.495772059 | 18.84161872 | 56.75767559 | 522 | 200 |
| GSE32533_WT_VS_MIR17_KO_ACT_CD4_TCELL_DN | ART_short | ART | short | 0 | 1 | 0 | 0 | 0 | 0 | 1 | 9 | 7.173596596 | 10.38905069 | 24.98262218 | 513 | 201 |
| GSE35685_CD34POS_CD38NEG_VS_CD34POS_CD10NEG_CD62LPOS_BONE_MARROW_UP | ART_short | ART | short | 0 | 0 | 0 | 0 | 0 | 0 | 2 | 8 | 6.280882353 | 25.29694556 | 26.52871992 | 604 | 202 |
| GSE360_CTRL_VS_B_MALAYI_HIGH_DOSE_MAC_DN | ART_short | ART | short | 0 | 0 | 0 | 0 | 1 | 0 | 1 | 8 | 6.280882353 | 24.78642261 | 45.40494983 | 730 | 203 |
| GSE360_L_MAJOR_VS_B_MALAYI_HIGH_DOSE_DC_DN | ART_short | ART | short | 0 | 0 | 0 | 0 | 1 | 0 | 1 | 7 | 5.551284908 | 15.64668901 | 43.21670155 | 628 | 204 |
| GSE3982_MAST_CELL_VS_CENT_MEMORY_CD4_TCELL_DN | ART_short | ART | short | 0 | 1 | 0 | 0 | 1 | 0 | 1 | 7 | 5.551284908 | 13.50226776 | 34.40505147 | 1017 | 205 |
| GSE40273_GATA1_KO_VS_WT_TREG_DN | ART_short | ART | short | 1 | 0 | 0 | 0 | 0 | 0 | 1 | 8 | 6.312444576 | 23.05617369 | 41.85837107 | 974 | 206 |
| GSE40274_CTRL_VS_FOXP3_AND_HELIOS_TRANSDUCED_ACTIVATED_CD4_TCELL_UP | ART_short | ART | short | 0 | 1 | 0 | 0 | 0 | 0 | 2 | 7 | 5.579464019 | 17.43069224 | 27.3761844 | 590 | 207 |
| GSE41867_DAY8_VS_DAY15_LCMV_CLONE13_EFFECTOR_CD8_TCELL_UP | ART_short | ART | short | 0 | 0 | 1 | 0 | 0 | 0 | 2 | 7 | 5.495772059 | 9.702900108 | 49.77187227 | 682 | 208 |
| GSE41867_NAIVE_VS_DAY15_LCMV_EFFECTOR_CD8_TCELL_DN | ART_short | ART | short | 0 | 0 | 1 | 0 | 0 | 0 | 1 | 7 | 5.495772059 | 21.23241142 | 38.5252212 | 613 | 209 |
| GSE41867_NAIVE_VS_DAY6_LCMV_EFFECTOR_CD8_TCELL_DN | ART_short | ART | short | 0 | 0 | 1 | 0 | 0 | 0 | 1 | 7 | 5.495772059 | 28.31317203 | 65.35105712 | 748 | 210 |
| GSE42021_CD24HI_TREG_VS_CD24HI_TCONV_THYMUS_DN | ART_short | ART | short | 1 | 0 | 0 | 0 | 0 | 0 | 1 | 7 | 5.495772059 | 19.43063433 | 21.68813656 | 462 | 211 |
| GSE43863_NAIVE_VS_MEMORY_LY6C_INT_CXCR5POS_CD4_TCELL_D150_LCMV_UP | ART_short | ART | short | 0 | 1 | 0 | 0 | 0 | 1 | 2 | 7 | 5.495772059 | 27.32738124 | 45.65478656 | 986 | 212 |
| GSE45365_WT_VS_IFNAR_KO_CD8A_DC_MCMV_INFECTION_UP | ART_short | ART | short | 0 | 0 | 0 | 0 | 1 | 1 | 2 | 9 | 7.101500148 | 23.36697387 | 28.72832819 | 949 | 213 |
| GSE5589_LPS_VS_LPS_AND_IL10_STIM_IL10_KO_MACROPHAGE_45MIN_DN | ART_short | ART | short | 0 | 0 | 0 | 0 | 1 | 1 | 1 | 8 | 6.280882353 | 37.90958122 | 29.18419652 | 698 | 214 |
| GSE6259_BCELL_VS_CD8_TCELL_DN | ART_short | ART | short | 0 | 0 | 1 | 1 | 0 | 0 | 1 | 8 | 6.344325609 | 3.964711718 | 18.65926544 | 602 | 215 |
| GSE7219_WT_VS_NIK_NFKB2_KO_DC_DN | ART_short | ART | short | 0 | 0 | 0 | 1 | 0 | 0 | 1 | 6 | 5.574747651 | 26.1393384 | 24.96108294 | 960 | 216 |
| GSE7768_OVA_ALONE_VS_OVA_WITH_MPL_IMMUNIZED_MOUSE_WHOLE_SPLEEN_6H_DN | ART_short | ART | short | 0 | 0 | 0 | 0 | 0 | 0 | 1 | 7 | 6.702161047 | 4.612619975 | 29.98991821 | 435 | 217 |
| GSE7852_LN_VS_THYMUS_TCONV_DN | ART_short | ART | short | 1 | 0 | 0 | 0 | 0 | 0 | 1 | 7 | 5.495772059 | 21.06515046 | 25.89152962 | 623 | 218 |
| GSE12003_4D_VS_8D_CULTURE_MIR223_KO_BM_PROGENITOR_DN | ART_untreat | ART | untreat | 1 | 0 | 0 | 0 | 0 | 0 | 1 | 7 | 8.016141141 | 12.62607688 | 21.06152545 | 582 | 219 |
| GSE13411_SWITCHED_MEMORY_BCELL_VS_PLASMA_CELL_UP | ART_untreat | ART | untreat | 0 | 0 | 0 | 1 | 0 | 0 | 2 | 7 | 6.733558559 | 7.211434195 | 27.52248418 | 544 | 220 |
| GSE16450_IMMATURE_VS_MATURE_NEURON_CELL_LINE_UP | ART_untreat | ART | untreat | 1 | 0 | 0 | 0 | 0 | 0 | 2 | 8 | 7.892815893 | 677.5661781 | 31.44721001 | 539 | 221 |
| GSE17974_0.5H_VS_72H_IL4_AND_ANTI_IL12_ACT_CD4_TCELL_DN | ART_untreat | ART | untreat | 0 | 1 | 0 | 0 | 0 | 1 | 1 | 7 | 6.870978121 | 8.561303039 | 26.25005133 | 365 | 222 |
| GSE21927_SPLENIC_VS_TUMOR_MONOCYTES_FROM_C26GM_TUMOROUS_MICE_BALBC_DN | ART_untreat | ART | untreat | 0 | 0 | 0 | 0 | 1 | 0 | 1 | 7 | 6.801574302 | 23.2583329 | 29.47425367 | 693 | 223 |
| GSE22886_NAIVE_CD8_TCELL_VS_NKCELL_UP | ART_untreat | ART | untreat | 1 | 0 | 1 | 0 | 0 | 0 | 2 | 7 | 6.801574302 | 33.03591728 | 35.09437329 | 668 | 224 |
| GSE3982_EOSINOPHIL_VS_BASOPHIL_DN | ART_untreat | ART | untreat | 1 | 0 | 0 | 0 | 0 | 0 | 1 | 7 | 6.801574302 | 10.20945548 | 28.28276606 | 563 | 225 |
| GSE9988_ANTI_TREM1_VS_CTRL_TREATED_MONOCYTES_DN | ART_untreat | ART | untreat | 0 | 0 | 0 | 0 | 1 | 0 | 1 | 8 | 7.695495495 | 24.23319028 | 28.01528839 | 472 | 226 |
| GSE10147_IL3_VS_IL3_AND_HIVP17_STIM_PDC_UP | EC_long | EC | long | 0 | 1 | 0 | 0 | 0 | 0 | 2 | 6 | 8.771221249 | 0.303820149 | 45.55603909 | 1059 | 227 |
| GSE11057_NAIVE_CD4_VS_PBMC_CD4_TCELL_UP | EC_long | EC | long | 0 | 1 | 0 | 0 | 0 | 0 | 2 | 6 | 7.390978311 | 9.062356029 | 49.09591792 | 660 | 228 |
| GSE12845_IGD_NEG_BLOOD_VS_NAIVE_TONSIL_BCELL_DN | EC_long | EC | long | 0 | 0 | 0 | 1 | 0 | 0 | 1 | 8 | 9.854637748 | 1.047475913 | 36.62581534 | 755 | 229 |
| GSE13411_IGM_MEMORY_BCELL_VS_PLASMA_CELL_UP | EC_long | EC | long | 0 | 0 | 0 | 1 | 0 | 1 | 2 | 6 | 7.353650138 | 0.689160554 | 27.97946358 | 661 | 230 |
| GSE13738_RESTING_VS_TCR_ACTIVATED_CD4_TCELL_UP | EC_long | EC | long | 0 | 1 | 0 | 0 | 0 | 0 | 2 | 6 | 7.466783217 | 9.615784953 | 22.91771724 | 411 | 231 |
| GSE16385_ROSIGLITAZONE_IL4_VS_IFNG_TNF_STIM_MACROPHAGE_UP | EC_long | EC | long | 0 | 0 | 0 | 0 | 1 | 0 | 2 | 7 | 8.493465909 | 8.248595637 | 32.95583357 | 667 | 232 |
| GSE16450_CTRL_VS_IFNA_12H_STIM_IMMATURE_NEURON_CELL_LINE_UP | EC_long | EC | long | 1 | 0 | 0 | 0 | 0 | 1 | 2 | 6 | 7.390978311 | 0.485616628 | 52.89932988 | 748 | 233 |
| GSE17301_ACD3_ACD28_VS_ACD3_ACD28_AND_IFNA2_STIM_CD8_TCELL_DN | EC_long | EC | long | 0 | 0 | 1 | 0 | 0 | 1 | 1 | 6 | 7.280113636 | 0.812974992 | 34.84495654 | 259 | 234 |
| GSE17301_IFNA2_VS_IFNA2_AND_ACD3_ACD28_STIM_CD8_TCELL_UP | EC_long | EC | long | 0 | 0 | 1 | 0 | 0 | 1 | 2 | 6 | 7.280113636 | 0.812974992 | 34.84495654 | 259 | 235 |
| GSE20715_0H_VS_24H_OZONE_LUNG_UP | EC_long | EC | long | 0 | 0 | 0 | 0 | 0 | 0 | 2 | 6 | 7.280113636 | 0.818557748 | 30.57529684 | 736 | 236 |
| GSE21033_1H_VS_24H_POLYIC_STIM_DC_UP | EC_long | EC | long | 0 | 0 | 0 | 0 | 1 | 0 | 2 | 6 | 9.043619424 | 0.38133305 | 24.83524412 | 577 | 237 |
| GSE21063_3H_VS_16H_ANTI_IGM_STIM_BCELL_DN | EC_long | EC | long | 0 | 0 | 0 | 1 | 0 | 1 | 1 | 6 | 7.390978311 | 0.40367544 | 19.49144545 | 572 | 238 |
| GSE21670_UNTREATED_VS_IL6_TREATED_CD4_TCELL_DN | EC_long | EC | long | 0 | 1 | 0 | 0 | 0 | 1 | 1 | 7 | 8.493465909 | 8.842076 | 34.89482631 | 533 | 239 |
| GSE25088_WT_VS_STAT6_KO_MACROPHAGE_IL4_STIM_UP | EC_long | EC | long | 0 | 0 | 0 | 0 | 1 | 1 | 2 | 6 | 7.316697122 | 8.934161359 | 37.77871547 | 412 | 240 |
| GSE25123_CTRL_VS_IL4_AND_ROSIGLITAZONE_STIM_MACROPHAGE_UP | EC_long | EC | long | 0 | 0 | 0 | 0 | 1 | 1 | 2 | 6 | 7.280113636 | 10.81329822 | 50.20293319 | 706 | 241 |
| GSE27241_WT_VS_RORGT_KO_TH17_POLARIZED_CD4_TCELL_TREATED_WITH_DIGOXIN_UP | EC_long | EC | long | 0 | 1 | 0 | 0 | 0 | 0 | 2 | 9 | 12.13352273 | 0.721989665 | 32.02378775 | 887 | 242 |
| GSE36476_CTRL_VS_TSST_ACT_72H_MEMORY_CD4_TCELL_YOUNG_UP | EC_long | EC | long | 0 | 1 | 0 | 0 | 0 | 0 | 2 | 6 | 7.353650138 | 9.113697015 | 43.11850638 | 498 | 243 |
| GSE369_IFNG_KO_VS_WT_LIVER_UP | EC_long | EC | long | 0 | 0 | 0 | 0 | 0 | 1 | 2 | 8 | 9.755596163 | 1.946835308 | 21.49135932 | 746 | 244 |
| GSE40273_GATA1_KO_VS_WT_TREG_DN | EC_long | EC | long | 1 | 0 | 0 | 0 | 0 | 0 | 1 | 9 | 10.97504568 | 1.088476719 | 31.40176743 | 1216 | 245 |
| GSE5503_MLN_DC_VS_PLN_DC_ACTIVATED_ALLOGENIC_TCELL_DN | EC_long | EC | long | 1 | 0 | 0 | 0 | 1 | 0 | 1 | 6 | 7.280113636 | 0.81715191 | 43.08764197 | 634 | 246 |
| NAKAYA_B_CELL_FLUARIX_FLUVIRIN_AGE_18_50YO_7DY_DN | EC_long | EC | long | 0 | 0 | 0 | 1 | 0 | 0 | 1 | 9 | 7.428687384 | 0.60748616 | 46.7644176 | 1389 | 247 |
| NAKAYA_PBMC_FLUARIX_FLUVIRIN_AGE_18_50YO_3DY_DN | EC_long | EC | long | 0 | 0 | 0 | 0 | 1 | 0 | 1 | 10 | 5.604398488 | 6.096863895 | 28.95377746 | 1181 | 248 |
