## Appendix Table S7 for "Distinguishable topological properties of functional genome networks in HIV-1 reservoirs"

**Supplementary Table 7. A complete edge list of the topological interaction among networks in HIV-1 infected individuals in a longitudinal order and in elite controllers (tetrapartitie graph)**

| row | column | correlation_coefficient | cat | id_from | id_to | patient_from | patient_to | type_of_a_walk |
|---|---|---|---|---|---|---|---|---|
| 4 | 7 | 0.999943662 | cat1 | from_1 | to_1 | art_long | art_long | M |
| 8 | 9 | 0.902324265 | cat1 | from_2 | to_2 | art_long | art_long | M |
| 8 | 10 | 0.904335594 | cat1 | from_3 | to_3 | art_long | art_long | M |
| 9 | 10 | 0.99996863 | cat1 | from_4 | to_4 | art_long | art_long | M |
| 4 | 11 | 0.800934224 | cat1 | from_5 | to_5 | art_long | art_long | M |
| 7 | 11 | 0.801088536 | cat1 | from_6 | to_6 | art_long | art_long | M |
| 12 | 13 | 0.73438152 | cat1 | from_7 | to_7 | art_long | art_long | M |
| 5 | 15 | 0.745317889 | cat1 | from_8 | to_8 | art_long | art_long | M |
| 13 | 15 | 0.824305339 | cat1 | from_9 | to_9 | art_long | art_long | M |
| 2 | 16 | 0.756237291 | cat1 | from_10 | to_10 | art_long | art_long | M |
| 12 | 16 | 0.845387517 | cat1 | from_11 | to_11 | art_long | art_long | M |
| 13 | 16 | 0.668687335 | cat1 | from_12 | to_12 | art_long | art_long | M |
| 15 | 16 | 0.78044594 | cat1 | from_13 | to_13 | art_long | art_long | M |
| 2 | 18 | 0.756237291 | cat1 | from_14 | to_14 | art_long | art_long | M |
| 12 | 18 | 0.845387517 | cat1 | from_15 | to_15 | art_long | art_long | M |
| 13 | 18 | 0.668687335 | cat1 | from_16 | to_16 | art_long | art_long | M |
| 15 | 18 | 0.78044594 | cat1 | from_17 | to_17 | art_long | art_long | M |
| 16 | 18 | 1 | cat1 | from_18 | to_18 | art_long | art_long | M |
| 12 | 19 | 0.944817401 | cat1 | from_19 | to_19 | art_long | art_long | M |
| 13 | 19 | 0.796939036 | cat1 | from_20 | to_20 | art_long | art_long | M |
| 16 | 19 | 0.818442311 | cat1 | from_21 | to_21 | art_long | art_long | M |
| 18 | 19 | 0.818442311 | cat1 | from_22 | to_22 | art_long | art_long | M |
| 13 | 20 | 0.772336835 | cat1 | from_23 | to_23 | art_long | art_long | M |
| 19 | 20 | 0.677496493 | cat1 | from_24 | to_24 | art_long | art_long | M |
| 13 | 21 | 0.669364957 | cat1 | from_25 | to_25 | art_long | art_long | M |
| 15 | 21 | 0.743399074 | cat1 | from_26 | to_26 | art_long | art_long | M |
| 20 | 21 | 0.841236001 | cat1 | from_27 | to_27 | art_long | art_long | M |
| 9 | 22 | 0.780750497 | cat1 | from_28 | to_28 | art_long | art_long | M |
| 10 | 22 | 0.77848359 | cat1 | from_29 | to_29 | art_long | art_long | M |
| 11 | 22 | 0.723578405 | cat1 | from_30 | to_30 | art_long | art_long | M |
| 8 | 23 | 0.7254786 | cat1 | from_31 | to_31 | art_long | art_long | M |
| 9 | 23 | 0.950286551 | cat1 | from_32 | to_32 | art_long | art_long | M |
| 10 | 23 | 0.949167232 | cat1 | from_33 | to_33 | art_long | art_long | M |
| 22 | 23 | 0.936482622 | cat1 | from_34 | to_34 | art_long | art_long | M |
| 14 | 24 | 1 | cat1 | from_35 | to_35 | art_long | art_long | M |
| 14 | 25 | 0.742067098 | cat1 | from_36 | to_36 | art_long | art_long | M |
| 24 | 25 | 0.742067098 | cat1 | from_37 | to_37 | art_long | art_long | M |
| 22 | 27 | 0.721470809 | cat1 | from_38 | to_38 | art_long | art_long | M |
| 14 | 28 | 0.999913472 | cat1 | from_39 | to_39 | art_long | art_long | M |
| 24 | 28 | 0.999913472 | cat1 | from_40 | to_40 | art_long | art_long | M |
| 25 | 28 | 0.745265448 | cat1 | from_41 | to_41 | art_long | art_long | M |
| 3 | 29 | 0.685166239 | cat1 | from_42 | to_42 | art_long | art_long | M |
| 25 | 31 | 0.928192513 | cat1 | from_43 | to_43 | art_long | art_long | M |
| 25 | 33 | 0.944302055 | cat1 | from_44 | to_44 | art_long | art_long | M |
| 31 | 33 | 0.998944866 | cat1 | from_45 | to_45 | art_long | art_long | M |
| 4 | 34 | 0.716738189 | cat1 | from_46 | to_46 | art_long | art_long | M |
| 5 | 34 | 0.6741564 | cat1 | from_47 | to_47 | art_long | art_long | M |
| 7 | 34 | 0.717604401 | cat1 | from_48 | to_48 | art_long | art_long | M |
| 22 | 34 | 0.680997094 | cat1 | from_49 | to_49 | art_long | art_long | M |
| 17 | 35 | 0.792328802 | cat1 | from_50 | to_50 | art_long | art_long | M |
| 17 | 36 | 0.99998586 | cat1 | from_51 | to_51 | art_long | art_long | M |
| 35 | 36 | 0.792323955 | cat1 | from_52 | to_52 | art_long | art_long | M |
| 26 | 37 | 0.957778227 | cat1 | from_53 | to_53 | art_long | art_long | M |
| 17 | 38 | 0.787936067 | cat1 | from_54 | to_54 | art_long | art_long | M |
| 35 | 38 | 0.999605701 | cat1 | from_55 | to_55 | art_long | art_long | M |
| 36 | 38 | 0.78778631 | cat1 | from_56 | to_56 | art_long | art_long | M |
| 2 | 39 | 0.678467169 | cat1 | from_57 | to_57 | art_long | art_long | M |
| 12 | 39 | 0.93832786 | cat1 | from_58 | to_58 | art_long | art_long | M |
| 13 | 39 | 0.77090952 | cat1 | from_59 | to_59 | art_long | art_long | M |
| 15 | 39 | 0.696586548 | cat1 | from_60 | to_60 | art_long | art_long | M |
| 16 | 39 | 0.950670778 | cat1 | from_61 | to_61 | art_long | art_long | M |
| 18 | 39 | 0.950670778 | cat1 | from_62 | to_62 | art_long | art_long | M |
| 19 | 39 | 0.956272131 | cat1 | from_63 | to_63 | art_long | art_long | M |
| 12 | 40 | 0.943272135 | cat1 | from_64 | to_64 | art_long | art_long | M |
| 13 | 40 | 0.79748072 | cat1 | from_65 | to_65 | art_long | art_long | M |
| 16 | 40 | 0.820311053 | cat1 | from_66 | to_66 | art_long | art_long | M |
| 18 | 40 | 0.820311053 | cat1 | from_67 | to_67 | art_long | art_long | M |
| 19 | 40 | 0.999974175 | cat1 | from_68 | to_68 | art_long | art_long | M |
| 20 | 40 | 0.677504491 | cat1 | from_69 | to_69 | art_long | art_long | M |
| 39 | 40 | 0.957248731 | cat1 | from_70 | to_70 | art_long | art_long | M |
| 29 | 41 | 0.818536547 | cat1 | from_71 | to_71 | art_long | art_long | M |
| 42 | 43 | 0.92933654 | cat1 | from_72 | to_72 | art_long | art_long | M |
| 42 | 45 | 1 | cat1 | from_73 | to_73 | art_long | art_long | M |
| 43 | 45 | 0.92933654 | cat1 | from_74 | to_74 | art_long | art_long | M |
| 44 | 46 | 1 | cat1 | from_75 | to_75 | art_long | art_long | M |
| 14 | 47 | 0.669654246 | cat1 | from_76 | to_76 | art_long | art_long | M |
| 24 | 47 | 0.669654246 | cat1 | from_77 | to_77 | art_long | art_long | M |
| 28 | 47 | 0.669026472 | cat1 | from_78 | to_78 | art_long | art_long | M |
| 30 | 47 | 0.957512643 | cat1 | from_79 | to_79 | art_long | art_long | M |
| 17 | 48 | 0.792328802 | cat1 | from_80 | to_80 | art_long | art_long | M |
| 35 | 48 | 1 | cat1 | from_81 | to_81 | art_long | art_long | M |
| 36 | 48 | 0.792323955 | cat1 | from_82 | to_82 | art_long | art_long | M |
| 38 | 48 | 0.999605701 | cat1 | from_83 | to_83 | art_long | art_long | M |
| 35 | 50 | 0.812968063 | cat1 | from_84 | to_84 | art_long | art_long | M |
| 38 | 50 | 0.819679063 | cat1 | from_85 | to_85 | art_long | art_long | M |
| 48 | 50 | 0.812968063 | cat1 | from_86 | to_86 | art_long | art_long | M |
| 44 | 52 | 0.870308282 | cat1 | from_87 | to_87 | art_long | art_long | M |
| 46 | 52 | 0.870308282 | cat1 | from_88 | to_88 | art_long | art_long | M |
| 2 | 53 | 0.99865413 | cat1 | from_89 | to_89 | art_long | art_long | M |
| 16 | 53 | 0.753715911 | cat1 | from_90 | to_90 | art_long | art_long | M |
| 18 | 53 | 0.753715911 | cat1 | from_91 | to_91 | art_long | art_long | M |
| 39 | 53 | 0.681906698 | cat1 | from_92 | to_92 | art_long | art_long | M |
| 25 | 54 | 0.716622404 | cat1 | from_93 | to_93 | art_long | art_long | M |
| 30 | 54 | 0.781330617 | cat1 | from_94 | to_94 | art_long | art_long | M |
| 31 | 54 | 0.694342264 | cat1 | from_95 | to_95 | art_long | art_long | M |
| 33 | 54 | 0.702513227 | cat1 | from_96 | to_96 | art_long | art_long | M |
| 47 | 54 | 0.734272204 | cat1 | from_97 | to_97 | art_long | art_long | M |
| 14 | 55 | 0.669654246 | cat1 | from_98 | to_98 | art_long | art_long | M |
| 24 | 55 | 0.669654246 | cat1 | from_99 | to_99 | art_long | art_long | M |
| 28 | 55 | 0.669026472 | cat1 | from_100 | to_100 | art_long | art_long | M |
| 30 | 55 | 0.957512643 | cat1 | from_101 | to_101 | art_long | art_long | M |
| 47 | 55 | 1 | cat1 | from_102 | to_102 | art_long | art_long | M |

| | | | | | | | | |
|---|---|---|---|---|---|---|---|---|
| 54 | 55 | 0.734272204 | cat1 | from_103 | to_103 | art_long | art_long | M |
| 22 | 56 | 0.712778951 | cat1 | from_104 | to_104 | art_long | art_long | M |
| 49 | 56 | 0.709321832 | cat1 | from_105 | to_105 | art_long | art_long | M |
| 22 | 57 | 0.712778951 | cat1 | from_106 | to_106 | art_long | art_long | M |
| 49 | 57 | 0.709321832 | cat1 | from_107 | to_107 | art_long | art_long | M |
| 56 | 57 | 1 | cat1 | from_108 | to_108 | art_long | art_long | M |
| 14 | 58 | 0.669654246 | cat1 | from_109 | to_109 | art_long | art_long | M |
| 24 | 58 | 0.669654246 | cat1 | from_110 | to_110 | art_long | art_long | M |
| 28 | 58 | 0.669026472 | cat1 | from_111 | to_111 | art_long | art_long | M |
| 30 | 58 | 0.957512643 | cat1 | from_112 | to_112 | art_long | art_long | M |
| 47 | 58 | 1 | cat1 | from_113 | to_113 | art_long | art_long | M |
| 54 | 58 | 0.734272204 | cat1 | from_114 | to_114 | art_long | art_long | M |
| 55 | 58 | 1 | cat1 | from_115 | to_115 | art_long | art_long | M |
| 14 | 59 | 0.999978687 | cat1 | from_116 | to_116 | art_long | art_long | M |
| 24 | 59 | 0.999978687 | cat1 | from_117 | to_117 | art_long | art_long | M |
| 25 | 59 | 0.743670517 | cat1 | from_118 | to_118 | art_long | art_long | M |
| 28 | 59 | 0.999978046 | cat1 | from_119 | to_119 | art_long | art_long | M |
| 47 | 59 | 0.669357162 | cat1 | from_120 | to_120 | art_long | art_long | M |
| 55 | 59 | 0.669357162 | cat1 | from_121 | to_121 | art_long | art_long | M |
| 58 | 59 | 0.669357162 | cat1 | from_122 | to_122 | art_long | art_long | M |
| 44 | 60 | 0.996052292 | cat1 | from_123 | to_123 | art_long | art_long | M |
| 46 | 60 | 0.996052292 | cat1 | from_124 | to_124 | art_long | art_long | M |
| 52 | 60 | 0.898219959 | cat1 | from_125 | to_125 | art_long | art_long | M |
| 44 | 61 | 0.915387767 | cat1 | from_126 | to_126 | art_long | art_long | M |
| 46 | 61 | 0.915387767 | cat1 | from_127 | to_127 | art_long | art_long | M |
| 52 | 61 | 0.730235289 | cat1 | from_128 | to_128 | art_long | art_long | M |
| 60 | 61 | 0.926658368 | cat1 | from_129 | to_129 | art_long | art_long | M |
| 42 | 62 | 0.999910267 | cat1 | from_130 | to_130 | art_long | art_long | M |
| 43 | 62 | 0.93101938 | cat1 | from_131 | to_131 | art_long | art_long | M |
| 45 | 62 | 0.999910267 | cat1 | from_132 | to_132 | art_long | art_long | M |
| 5 | 63 | 0.966316623 | cat1 | from_133 | to_133 | art_long | art_long | M |
| 15 | 63 | 0.717178259 | cat1 | from_134 | to_134 | art_long | art_long | M |
| 27 | 64 | 0.807243734 | cat1 | from_135 | to_135 | art_long | art_long | M |
| 34 | 64 | 0.713439948 | cat1 | from_136 | to_136 | art_long | art_long | M |
| 22 | 65 | 0.699946548 | cat1 | from_137 | to_137 | art_long | art_long | M |
| 23 | 65 | 0.711383841 | cat1 | from_138 | to_138 | art_long | art_long | M |
| 34 | 65 | 0.830475455 | cat1 | from_139 | to_139 | art_long | art_long | M |
| 49 | 65 | 0.67356238 | cat1 | from_140 | to_140 | art_long | art_long | M |
| 29 | 66 | 0.831587614 | cat1 | from_141 | to_141 | art_long | art_long | M |
| 41 | 66 | 0.666916561 | cat1 | from_142 | to_142 | art_long | art_long | M |
| 29 | 67 | 0.831587614 | cat1 | from_143 | to_143 | art_long | art_long | M |
| 41 | 67 | 0.666916561 | cat1 | from_144 | to_144 | art_long | art_long | M |
| 66 | 67 | 1 | cat1 | from_145 | to_145 | art_long | art_long | M |
| 13 | 68 | 0.770278283 | cat1 | from_146 | to_146 | art_long | art_long | M |
| 19 | 68 | 0.674928148 | cat1 | from_147 | to_147 | art_long | art_long | M |
| 40 | 68 | 0.675104381 | cat1 | from_148 | to_148 | art_long | art_long | M |
| 14 | 69 | 0.66706457 | cat1 | from_149 | to_149 | art_long | art_long | M |
| 24 | 69 | 0.66706457 | cat1 | from_150 | to_150 | art_long | art_long | M |
| 28 | 69 | 0.666624217 | cat1 | from_151 | to_151 | art_long | art_long | M |
| 30 | 69 | 0.961669434 | cat1 | from_152 | to_152 | art_long | art_long | M |
| 47 | 69 | 0.999890826 | cat1 | from_153 | to_153 | art_long | art_long | M |
| 54 | 69 | 0.738201599 | cat1 | from_154 | to_154 | art_long | art_long | M |
| 55 | 69 | 0.999890826 | cat1 | from_155 | to_155 | art_long | art_long | M |
| 58 | 69 | 0.999890826 | cat1 | from_156 | to_156 | art_long | art_long | M |
| 59 | 69 | 0.666860448 | cat1 | from_157 | to_157 | art_long | art_long | M |
| 49 | 70 | 0.955085217 | cat1 | from_158 | to_158 | art_long | art_long | M |
| 56 | 70 | 0.800339111 | cat1 | from_159 | to_159 | art_long | art_long | M |
| 57 | 70 | 0.800339111 | cat1 | from_160 | to_160 | art_long | art_long | M |
| 65 | 70 | 0.672520754 | cat1 | from_161 | to_161 | art_long | art_long | M |
| 3 | 71 | 0.757877932 | cat1 | from_162 | to_162 | art_long | art_long | M |
| 29 | 71 | 0.969012896 | cat1 | from_163 | to_163 | art_long | art_long | M |
| 41 | 71 | 0.837966519 | cat1 | from_164 | to_164 | art_long | art_long | M |
| 66 | 71 | 0.851063462 | cat1 | from_165 | to_165 | art_long | art_long | M |
| 67 | 71 | 0.851063462 | cat1 | from_166 | to_166 | art_long | art_long | M |
| 30 | 72 | 0.926894994 | cat1 | from_167 | to_167 | art_long | art_long | M |
| 47 | 72 | 0.921173246 | cat1 | from_168 | to_168 | art_long | art_long | M |
| 54 | 72 | 0.940570252 | cat1 | from_169 | to_169 | art_long | art_long | M |
| 55 | 72 | 0.921173246 | cat1 | from_170 | to_170 | art_long | art_long | M |
| 58 | 72 | 0.921173246 | cat1 | from_171 | to_171 | art_long | art_long | M |
| 69 | 72 | 0.923371151 | cat1 | from_172 | to_172 | art_long | art_long | M |
| 14 | 73 | 1 | cat1 | from_173 | to_173 | art_long | art_long | M |
| 24 | 73 | 1 | cat1 | from_174 | to_174 | art_long | art_long | M |
| 25 | 73 | 0.742067098 | cat1 | from_175 | to_175 | art_long | art_long | M |
| 28 | 73 | 0.999913472 | cat1 | from_176 | to_176 | art_long | art_long | M |
| 47 | 73 | 0.669654246 | cat1 | from_177 | to_177 | art_long | art_long | M |
| 55 | 73 | 0.669654246 | cat1 | from_178 | to_178 | art_long | art_long | M |
| 58 | 73 | 0.669654246 | cat1 | from_179 | to_179 | art_long | art_long | M |
| 59 | 73 | 0.999978687 | cat1 | from_180 | to_180 | art_long | art_long | M |
| 69 | 73 | 0.66706457 | cat1 | from_181 | to_181 | art_long | art_long | M |
| 14 | 74 | 1 | cat1 | from_182 | to_182 | art_long | art_long | M |
| 24 | 74 | 1 | cat1 | from_183 | to_183 | art_long | art_long | M |
| 25 | 74 | 0.742067098 | cat1 | from_184 | to_184 | art_long | art_long | M |
| 28 | 74 | 0.999913472 | cat1 | from_185 | to_185 | art_long | art_long | M |
| 47 | 74 | 0.669654246 | cat1 | from_186 | to_186 | art_long | art_long | M |
| 55 | 74 | 0.669654246 | cat1 | from_187 | to_187 | art_long | art_long | M |
| 58 | 74 | 0.669654246 | cat1 | from_188 | to_188 | art_long | art_long | M |
| 59 | 74 | 0.999978687 | cat1 | from_189 | to_189 | art_long | art_long | M |
| 69 | 74 | 0.66706457 | cat1 | from_190 | to_190 | art_long | art_long | M |
| 73 | 74 | 1 | cat1 | from_191 | to_191 | art_long | art_long | M |
| 49 | 75 | 0.999973484 | cat1 | from_192 | to_192 | art_long | art_long | M |
| 56 | 75 | 0.71075463 | cat1 | from_193 | to_193 | art_long | art_long | M |
| 57 | 75 | 0.71075463 | cat1 | from_194 | to_194 | art_long | art_long | M |
| 65 | 75 | 0.674285399 | cat1 | from_195 | to_195 | art_long | art_long | M |
| 70 | 75 | 0.95606854 | cat1 | from_196 | to_196 | art_long | art_long | M |
| 14 | 76 | 0.932211263 | cat1 | from_197 | to_197 | art_long | art_long | M |
| 24 | 76 | 0.932211263 | cat1 | from_198 | to_198 | art_long | art_long | M |
| 25 | 76 | 0.93434828 | cat1 | from_199 | to_199 | art_long | art_long | M |
| 28 | 76 | 0.933841755 | cat1 | from_200 | to_200 | art_long | art_long | M |
| 31 | 76 | 0.80096256 | cat1 | from_201 | to_201 | art_long | art_long | M |
| 33 | 76 | 0.823678887 | cat1 | from_202 | to_202 | art_long | art_long | M |
| 59 | 76 | 0.933040655 | cat1 | from_203 | to_203 | art_long | art_long | M |
| 73 | 76 | 0.932211263 | cat1 | from_204 | to_204 | art_long | art_long | M |
| 74 | 76 | 0.932211263 | cat1 | from_205 | to_205 | art_long | art_long | M |
| 30 | 77 | 0.927822312 | cat1 | from_206 | to_206 | art_long | art_long | M |

| | | | | | | | | |
|---|---|---|---|---|---|---|---|---|
| 47 | 77 | 0.919578628 | cat1 | from_207 | to_207 | art_long | art_long | M |
| 54 | 77 | 0.941896354 | cat1 | from_208 | to_208 | art_long | art_long | M |
| 55 | 77 | 0.919578628 | cat1 | from_209 | to_209 | art_long | art_long | M |
| 58 | 77 | 0.919578628 | cat1 | from_210 | to_210 | art_long | art_long | M |
| 69 | 77 | 0.921902451 | cat1 | from_211 | to_211 | art_long | art_long | M |
| 72 | 77 | 0.999962474 | cat1 | from_212 | to_212 | art_long | art_long | M |
| 14 | 78 | 0.999913472 | cat1 | from_213 | to_213 | art_long | art_long | M |
| 24 | 78 | 0.999913472 | cat1 | from_214 | to_214 | art_long | art_long | M |
| 25 | 78 | 0.745265448 | cat1 | from_215 | to_215 | art_long | art_long | M |
| 28 | 78 | 1 | cat1 | from_216 | to_216 | art_long | art_long | M |
| 47 | 78 | 0.669026472 | cat1 | from_217 | to_217 | art_long | art_long | M |
| 55 | 78 | 0.669026472 | cat1 | from_218 | to_218 | art_long | art_long | M |
| 58 | 78 | 0.669026472 | cat1 | from_219 | to_219 | art_long | art_long | M |
| 59 | 78 | 0.999978046 | cat1 | from_220 | to_220 | art_long | art_long | M |
| 69 | 78 | 0.666624217 | cat1 | from_221 | to_221 | art_long | art_long | M |
| 73 | 78 | 0.999913472 | cat1 | from_222 | to_222 | art_long | art_long | M |
| 74 | 78 | 0.999913472 | cat1 | from_223 | to_223 | art_long | art_long | M |
| 76 | 78 | 0.933841755 | cat1 | from_224 | to_224 | art_long | art_long | M |
| 12 | 79 | 0.965187295 | cat1 | from_225 | to_225 | art_long | art_long | M |
| 13 | 79 | 0.778019164 | cat1 | from_226 | to_226 | art_long | art_long | M |
| 16 | 79 | 0.773332328 | cat1 | from_227 | to_227 | art_long | art_long | M |
| 18 | 79 | 0.773332328 | cat1 | from_228 | to_228 | art_long | art_long | M |
| 19 | 79 | 0.990063918 | cat1 | from_229 | to_229 | art_long | art_long | M |
| 20 | 79 | 0.670266 | cat1 | from_230 | to_230 | art_long | art_long | M |
| 39 | 79 | 0.927178918 | cat1 | from_231 | to_231 | art_long | art_long | M |
| 40 | 79 | 0.989027761 | cat1 | from_232 | to_232 | art_long | art_long | M |
| 30 | 80 | 0.929494503 | cat1 | from_233 | to_233 | art_long | art_long | M |
| 47 | 80 | 0.916120086 | cat1 | from_234 | to_234 | art_long | art_long | M |
| 54 | 80 | 0.944376888 | cat1 | from_235 | to_235 | art_long | art_long | M |
| 55 | 80 | 0.916120086 | cat1 | from_236 | to_236 | art_long | art_long | M |
| 58 | 80 | 0.916120086 | cat1 | from_237 | to_237 | art_long | art_long | M |
| 69 | 80 | 0.918699639 | cat1 | from_238 | to_238 | art_long | art_long | M |
| 72 | 80 | 0.999654291 | cat1 | from_239 | to_239 | art_long | art_long | M |
| 77 | 80 | 0.999844556 | cat1 | from_240 | to_240 | art_long | art_long | M |
| 25 | 81 | 0.697543207 | cat1 | from_241 | to_241 | art_long | art_long | M |
| 30 | 81 | 0.767177907 | cat1 | from_242 | to_242 | art_long | art_long | M |
| 31 | 81 | 0.733466582 | cat1 | from_243 | to_243 | art_long | art_long | M |
| 33 | 81 | 0.734759387 | cat1 | from_244 | to_244 | art_long | art_long | M |
| 47 | 81 | 0.676478548 | cat1 | from_245 | to_245 | art_long | art_long | M |
| 54 | 81 | 0.988044105 | cat1 | from_246 | to_246 | art_long | art_long | M |
| 55 | 81 | 0.676478548 | cat1 | from_247 | to_247 | art_long | art_long | M |
| 58 | 81 | 0.676478548 | cat1 | from_248 | to_248 | art_long | art_long | M |
| 69 | 81 | 0.682524455 | cat1 | from_249 | to_249 | art_long | art_long | M |
| 72 | 81 | 0.904805116 | cat1 | from_250 | to_250 | art_long | art_long | M |
| 77 | 81 | 0.907434674 | cat1 | from_251 | to_251 | art_long | art_long | M |
| 80 | 81 | 0.91257599 | cat1 | from_252 | to_252 | art_long | art_long | M |
| 3 | 82 | 0.742206192 | cat1 | from_253 | to_253 | art_long | art_long | M |
| 29 | 82 | 0.754912335 | cat1 | from_254 | to_254 | art_long | art_long | M |
| 41 | 82 | 0.676653357 | cat1 | from_255 | to_255 | art_long | art_long | M |
| 66 | 82 | 0.685152099 | cat1 | from_256 | to_256 | art_long | art_long | M |
| 67 | 82 | 0.685152099 | cat1 | from_257 | to_257 | art_long | art_long | M |
| 71 | 82 | 0.816603387 | cat1 | from_258 | to_258 | art_long | art_long | M |
| 25 | 83 | 0.666543824 | cat1 | from_259 | to_259 | art_long | art_long | M |
| 30 | 83 | 0.933158572 | cat1 | from_260 | to_260 | art_long | art_long | M |
| 47 | 83 | 0.900931255 | cat1 | from_261 | to_261 | art_long | art_long | M |
| 54 | 83 | 0.950976964 | cat1 | from_262 | to_262 | art_long | art_long | M |
| 55 | 83 | 0.900931255 | cat1 | from_263 | to_263 | art_long | art_long | M |
| 58 | 83 | 0.900931255 | cat1 | from_264 | to_264 | art_long | art_long | M |
| 69 | 83 | 0.904445361 | cat1 | from_265 | to_265 | art_long | art_long | M |
| 72 | 83 | 0.995839521 | cat1 | from_266 | to_266 | art_long | art_long | M |
| 77 | 83 | 0.996591579 | cat1 | from_267 | to_267 | art_long | art_long | M |
| 80 | 83 | 0.997891142 | cat1 | from_268 | to_268 | art_long | art_long | M |
| 81 | 83 | 0.929056982 | cat1 | from_269 | to_269 | art_long | art_long | M |
| 8 | 84 | 0.732331819 | cat1 | from_270 | to_270 | art_long | art_long | M |
| 49 | 84 | 0.885600856 | cat1 | from_271 | to_271 | art_long | art_long | M |
| 70 | 84 | 0.708206658 | cat1 | from_272 | to_272 | art_long | art_long | M |
| 75 | 84 | 0.884037069 | cat1 | from_273 | to_273 | art_long | art_long | M |
| 34 | 85 | 0.702141191 | cat1 | from_274 | to_274 | art_long | art_long | M |
| 49 | 85 | 0.83416185 | cat1 | from_275 | to_275 | art_long | art_long | M |
| 56 | 85 | 0.820402628 | cat1 | from_276 | to_276 | art_long | art_long | M |
| 57 | 85 | 0.820402628 | cat1 | from_277 | to_277 | art_long | art_long | M |
| 70 | 85 | 0.960122287 | cat1 | from_278 | to_278 | art_long | art_long | M |
| 75 | 85 | 0.836047119 | cat1 | from_279 | to_279 | art_long | art_long | M |
| 32 | 86 | 0.74489146 | cat1 | from_280 | to_280 | art_long | art_long | M |
| 14 | 87 | 0.669654246 | cat1 | from_281 | to_281 | art_long | art_long | M |
| 24 | 87 | 0.669654246 | cat1 | from_282 | to_282 | art_long | art_long | M |
| 28 | 87 | 0.669026472 | cat1 | from_283 | to_283 | art_long | art_long | M |
| 30 | 87 | 0.957512643 | cat1 | from_284 | to_284 | art_long | art_long | M |
| 47 | 87 | 1 | cat1 | from_285 | to_285 | art_long | art_long | M |
| 54 | 87 | 0.734272204 | cat1 | from_286 | to_286 | art_long | art_long | M |
| 55 | 87 | 1 | cat1 | from_287 | to_287 | art_long | art_long | M |
| 58 | 87 | 1 | cat1 | from_288 | to_288 | art_long | art_long | M |
| 59 | 87 | 0.669357162 | cat1 | from_289 | to_289 | art_long | art_long | M |
| 69 | 87 | 0.999890826 | cat1 | from_290 | to_290 | art_long | art_long | M |
| 72 | 87 | 0.921173246 | cat1 | from_291 | to_291 | art_long | art_long | M |
| 73 | 87 | 0.669654246 | cat1 | from_292 | to_292 | art_long | art_long | M |
| 74 | 87 | 0.669654246 | cat1 | from_293 | to_293 | art_long | art_long | M |
| 77 | 87 | 0.919578628 | cat1 | from_294 | to_294 | art_long | art_long | M |
| 78 | 87 | 0.669026472 | cat1 | from_295 | to_295 | art_long | art_long | M |
| 80 | 87 | 0.916120086 | cat1 | from_296 | to_296 | art_long | art_long | M |
| 81 | 87 | 0.676478548 | cat1 | from_297 | to_297 | art_long | art_long | M |
| 83 | 87 | 0.900931255 | cat1 | from_298 | to_298 | art_long | art_long | M |
| 8 | 88 | 0.719617017 | cat1 | from_299 | to_299 | art_long | art_long | M |
| 9 | 88 | 0.948042025 | cat1 | from_300 | to_300 | art_long | art_long | M |
| 10 | 88 | 0.946800503 | cat1 | from_301 | to_301 | art_long | art_long | M |
| 22 | 88 | 0.938876814 | cat1 | from_302 | to_302 | art_long | art_long | M |
| 23 | 88 | 0.999878248 | cat1 | from_303 | to_303 | art_long | art_long | M |
| 65 | 88 | 0.71323495 | cat1 | from_304 | to_304 | art_long | art_long | M |
| 3 | 89 | 0.709881465 | cat1 | from_305 | to_305 | art_long | art_long | M |
| 29 | 89 | 0.836485115 | cat1 | from_306 | to_306 | art_long | art_long | M |
| 41 | 89 | 0.751450276 | cat1 | from_307 | to_307 | art_long | art_long | M |
| 66 | 89 | 0.760218664 | cat1 | from_308 | to_308 | art_long | art_long | M |
| 67 | 89 | 0.760218664 | cat1 | from_309 | to_309 | art_long | art_long | M |
| 71 | 89 | 0.869789995 | cat1 | from_310 | to_310 | art_long | art_long | M |

| | | | | | | | | |
|---|---|---|---|---|---|---|---|---|
| 82 | 89 | 0.970679347 | cat1 | from_311 | to_311 | art_long | art_long | M |
| 14 | 90 | 0.870862898 | cat1 | from_312 | to_312 | art_long | art_long | M |
| 24 | 90 | 0.870862898 | cat1 | from_313 | to_313 | art_long | art_long | M |
| 25 | 90 | 0.947781534 | cat1 | from_314 | to_314 | art_long | art_long | M |
| 28 | 90 | 0.875397828 | cat1 | from_315 | to_315 | art_long | art_long | M |
| 31 | 90 | 0.898414391 | cat1 | from_316 | to_316 | art_long | art_long | M |
| 33 | 90 | 0.911521796 | cat1 | from_317 | to_317 | art_long | art_long | M |
| 59 | 90 | 0.873132459 | cat1 | from_318 | to_318 | art_long | art_long | M |
| 73 | 90 | 0.870862898 | cat1 | from_319 | to_319 | art_long | art_long | M |
| 74 | 90 | 0.870862898 | cat1 | from_320 | to_320 | art_long | art_long | M |
| 76 | 90 | 0.973809375 | cat1 | from_321 | to_321 | art_long | art_long | M |
| 78 | 90 | 0.875397828 | cat1 | from_322 | to_322 | art_long | art_long | M |
| 22 | 91 | 0.695766707 | cat1 | from_323 | to_323 | art_long | art_long | M |
| 23 | 91 | 0.700589898 | cat1 | from_324 | to_324 | art_long | art_long | M |
| 27 | 91 | 0.957849107 | cat1 | from_325 | to_325 | art_long | art_long | M |
| 64 | 91 | 0.722677238 | cat1 | from_326 | to_326 | art_long | art_long | M |
| 88 | 91 | 0.70221173 | cat1 | from_327 | to_327 | art_long | art_long | M |
| 42 | 92 | 0.999910267 | cat1 | from_328 | to_328 | art_long | art_long | M |
| 43 | 92 | 0.93101938 | cat1 | from_329 | to_329 | art_long | art_long | M |
| 45 | 92 | 0.999910267 | cat1 | from_330 | to_330 | art_long | art_long | M |
| 62 | 92 | 1 | cat1 | from_331 | to_331 | art_long | art_long | M |
| 8 | 93 | 0.747079028 | cat1 | from_332 | to_332 | art_long | art_long | M |
| 32 | 94 | 0.74489146 | cat1 | from_333 | to_333 | art_long | art_long | M |
| 86 | 94 | 1 | cat1 | from_334 | to_334 | art_long | art_long | M |
| 14 | 95 | 0.669654246 | cat1 | from_335 | to_335 | art_long | art_long | M |
| 24 | 95 | 0.669654246 | cat1 | from_336 | to_336 | art_long | art_long | M |
| 28 | 95 | 0.669026472 | cat1 | from_337 | to_337 | art_long | art_long | M |
| 30 | 95 | 0.957512643 | cat1 | from_338 | to_338 | art_long | art_long | M |
| 47 | 95 | 1 | cat1 | from_339 | to_339 | art_long | art_long | M |
| 54 | 95 | 0.734272204 | cat1 | from_340 | to_340 | art_long | art_long | M |
| 55 | 95 | 1 | cat1 | from_341 | to_341 | art_long | art_long | M |
| 58 | 95 | 1 | cat1 | from_342 | to_342 | art_long | art_long | M |
| 59 | 95 | 0.669357162 | cat1 | from_343 | to_343 | art_long | art_long | M |
| 69 | 95 | 0.999890826 | cat1 | from_344 | to_344 | art_long | art_long | M |
| 72 | 95 | 0.921173246 | cat1 | from_345 | to_345 | art_long | art_long | M |
| 73 | 95 | 0.669654246 | cat1 | from_346 | to_346 | art_long | art_long | M |
| 74 | 95 | 0.669654246 | cat1 | from_347 | to_347 | art_long | art_long | M |
| 77 | 95 | 0.919578628 | cat1 | from_348 | to_348 | art_long | art_long | M |
| 78 | 95 | 0.669026472 | cat1 | from_349 | to_349 | art_long | art_long | M |
| 80 | 95 | 0.916120086 | cat1 | from_350 | to_350 | art_long | art_long | M |
| 81 | 95 | 0.676478548 | cat1 | from_351 | to_351 | art_long | art_long | M |
| 83 | 95 | 0.900931255 | cat1 | from_352 | to_352 | art_long | art_long | M |
| 87 | 95 | 1 | cat1 | from_353 | to_353 | art_long | art_long | M |
| 29 | 96 | 0.776160842 | cat1 | from_354 | to_354 | art_long | art_long | M |
| 41 | 96 | 0.97648978 | cat1 | from_355 | to_355 | art_long | art_long | M |
| 71 | 96 | 0.823915995 | cat1 | from_356 | to_356 | art_long | art_long | M |
| 82 | 96 | 0.729606795 | cat1 | from_357 | to_357 | art_long | art_long | M |
| 89 | 96 | 0.753050965 | cat1 | from_358 | to_358 | art_long | art_long | M |
| 14 | 97 | 1 | cat1 | from_359 | to_359 | art_long | art_long | M |
| 24 | 97 | 1 | cat1 | from_360 | to_360 | art_long | art_long | M |
| 25 | 97 | 0.742067098 | cat1 | from_361 | to_361 | art_long | art_long | M |
| 28 | 97 | 0.999913472 | cat1 | from_362 | to_362 | art_long | art_long | M |
| 47 | 97 | 0.669654246 | cat1 | from_363 | to_363 | art_long | art_long | M |
| 55 | 97 | 0.669654246 | cat1 | from_364 | to_364 | art_long | art_long | M |
| 58 | 97 | 0.669654246 | cat1 | from_365 | to_365 | art_long | art_long | M |
| 59 | 97 | 0.999978687 | cat1 | from_366 | to_366 | art_long | art_long | M |
| 69 | 97 | 0.66706457 | cat1 | from_367 | to_367 | art_long | art_long | M |
| 73 | 97 | 1 | cat1 | from_368 | to_368 | art_long | art_long | M |
| 74 | 97 | 1 | cat1 | from_369 | to_369 | art_long | art_long | M |
| 76 | 97 | 0.932211263 | cat1 | from_370 | to_370 | art_long | art_long | M |
| 78 | 97 | 0.999913472 | cat1 | from_371 | to_371 | art_long | art_long | M |
| 87 | 97 | 0.669654246 | cat1 | from_372 | to_372 | art_long | art_long | M |
| 90 | 97 | 0.870862898 | cat1 | from_373 | to_373 | art_long | art_long | M |
| 95 | 97 | 0.669654246 | cat1 | from_374 | to_374 | art_long | art_long | M |
| 29 | 98 | 0.670903602 | cat1 | from_375 | to_375 | art_long | art_long | M |
| 41 | 98 | 0.892024684 | cat1 | from_376 | to_376 | art_long | art_long | M |
| 71 | 98 | 0.691608371 | cat1 | from_377 | to_377 | art_long | art_long | M |
| 82 | 98 | 0.790438249 | cat1 | from_378 | to_378 | art_long | art_long | M |
| 89 | 98 | 0.841718207 | cat1 | from_379 | to_379 | art_long | art_long | M |
| 96 | 98 | 0.883165601 | cat1 | from_380 | to_380 | art_long | art_long | M |
| 44 | 99 | 0.917068621 | cat1 | from_381 | to_381 | art_long | art_long | M |
| 46 | 99 | 0.917068621 | cat1 | from_382 | to_382 | art_long | art_long | M |
| 52 | 99 | 0.728730331 | cat1 | from_383 | to_383 | art_long | art_long | M |
| 60 | 99 | 0.927559009 | cat1 | from_384 | to_384 | art_long | art_long | M |
| 61 | 99 | 0.999960565 | cat1 | from_385 | to_385 | art_long | art_long | M |
| 42 | 100 | 1 | cat1 | from_386 | to_386 | art_long | art_long | M |
| 43 | 100 | 0.92933654 | cat1 | from_387 | to_387 | art_long | art_long | M |
| 45 | 100 | 1 | cat1 | from_388 | to_388 | art_long | art_long | M |
| 62 | 100 | 0.999910267 | cat1 | from_389 | to_389 | art_long | art_long | M |
| 92 | 100 | 0.999910267 | cat1 | from_390 | to_390 | art_long | art_long | M |
| 42 | 101 | 1 | cat1 | from_391 | to_391 | art_long | art_long | M |
| 43 | 101 | 0.92933654 | cat1 | from_392 | to_392 | art_long | art_long | M |
| 45 | 101 | 1 | cat1 | from_393 | to_393 | art_long | art_long | M |
| 62 | 101 | 0.999910267 | cat1 | from_394 | to_394 | art_long | art_long | M |
| 92 | 101 | 0.999910267 | cat1 | from_395 | to_395 | art_long | art_long | M |
| 100 | 101 | 1 | cat1 | from_396 | to_396 | art_long | art_long | M |
| 44 | 102 | 0.999251827 | cat1 | from_397 | to_397 | art_long | art_long | M |
| 46 | 102 | 0.999251827 | cat1 | from_398 | to_398 | art_long | art_long | M |
| 52 | 102 | 0.883314852 | cat1 | from_399 | to_399 | art_long | art_long | M |
| 60 | 102 | 0.998740227 | cat1 | from_400 | to_400 | art_long | art_long | M |
| 61 | 102 | 0.921187815 | cat1 | from_401 | to_401 | art_long | art_long | M |
| 99 | 102 | 0.922530372 | cat1 | from_402 | to_402 | art_long | art_long | M |
| 51 | 103 | 0.843143104 | cat1 | from_403 | to_403 | art_long | art_long | M |
| 17 | 104 | 0.792328802 | cat1 | from_404 | to_404 | art_long | art_long | M |
| 35 | 104 | 1 | cat1 | from_405 | to_405 | art_long | art_long | M |
| 36 | 104 | 0.792323955 | cat1 | from_406 | to_406 | art_long | art_long | M |
| 38 | 104 | 0.999605701 | cat1 | from_407 | to_407 | art_long | art_long | M |
| 48 | 104 | 1 | cat1 | from_408 | to_408 | art_long | art_long | M |
| 50 | 104 | 0.812968063 | cat1 | from_409 | to_409 | art_long | art_long | M |
| 44 | 105 | 0.723041928 | cat1 | from_410 | to_410 | art_long | art_long | M |
| 46 | 105 | 0.723041928 | cat1 | from_411 | to_411 | art_long | art_long | M |
| 50 | 105 | 0.681276847 | cat1 | from_412 | to_412 | art_long | art_long | M |
| 60 | 105 | 0.744634563 | cat1 | from_413 | to_413 | art_long | art_long | M |
| 61 | 105 | 0.939909943 | cat1 | from_414 | to_414 | art_long | art_long | M |

| | | | | | | | | |
|---|---|---|---|---|---|---|---|---|
| 99 | 105 | 0.938522996 | cat1 | from_415 | to_415 | art_long | art_long | M |
| 102 | 105 | 0.733152249 | cat1 | from_416 | to_416 | art_long | art_long | M |
| 44 | 106 | 1 | cat1 | from_417 | to_417 | art_long | art_long | M |
| 46 | 106 | 1 | cat1 | from_418 | to_418 | art_long | art_long | M |
| 52 | 106 | 0.870308282 | cat1 | from_419 | to_419 | art_long | art_long | M |
| 60 | 106 | 0.996052292 | cat1 | from_420 | to_420 | art_long | art_long | M |
| 61 | 106 | 0.915387767 | cat1 | from_421 | to_421 | art_long | art_long | M |
| 99 | 106 | 0.917068621 | cat1 | from_422 | to_422 | art_long | art_long | M |
| 102 | 106 | 0.999251827 | cat1 | from_423 | to_423 | art_long | art_long | M |
| 105 | 106 | 0.723041928 | cat1 | from_424 | to_424 | art_long | art_long | M |
| 42 | 107 | 1 | cat1 | from_425 | to_425 | art_long | art_long | M |
| 43 | 107 | 0.92933654 | cat1 | from_426 | to_426 | art_long | art_long | M |
| 45 | 107 | 1 | cat1 | from_427 | to_427 | art_long | art_long | M |
| 62 | 107 | 0.999910267 | cat1 | from_428 | to_428 | art_long | art_long | M |
| 92 | 107 | 0.999910267 | cat1 | from_429 | to_429 | art_long | art_long | M |
| 100 | 107 | 1 | cat1 | from_430 | to_430 | art_long | art_long | M |
| 101 | 107 | 1 | cat1 | from_431 | to_431 | art_long | art_long | M |
| 14 | 109 | 0.66706457 | cat1 | from_432 | to_432 | art_long | art_long | M |
| 24 | 109 | 0.66706457 | cat1 | from_433 | to_433 | art_long | art_long | M |
| 28 | 109 | 0.666624217 | cat1 | from_434 | to_434 | art_long | art_long | M |
| 30 | 109 | 0.961669434 | cat1 | from_435 | to_435 | art_long | art_long | M |
| 47 | 109 | 0.999890826 | cat1 | from_436 | to_436 | art_long | art_long | M |
| 54 | 109 | 0.738201599 | cat1 | from_437 | to_437 | art_long | art_long | M |
| 55 | 109 | 0.999890826 | cat1 | from_438 | to_438 | art_long | art_long | M |
| 58 | 109 | 0.999890826 | cat1 | from_439 | to_439 | art_long | art_long | M |
| 59 | 109 | 0.666860448 | cat1 | from_440 | to_440 | art_long | art_long | M |
| 69 | 109 | 1 | cat1 | from_441 | to_441 | art_long | art_long | M |
| 72 | 109 | 0.923371151 | cat1 | from_442 | to_442 | art_long | art_long | M |
| 73 | 109 | 0.66706457 | cat1 | from_443 | to_443 | art_long | art_long | M |
| 74 | 109 | 0.66706457 | cat1 | from_444 | to_444 | art_long | art_long | M |
| 77 | 109 | 0.921902451 | cat1 | from_445 | to_445 | art_long | art_long | M |
| 78 | 109 | 0.666624217 | cat1 | from_446 | to_446 | art_long | art_long | M |
| 80 | 109 | 0.918699639 | cat1 | from_447 | to_447 | art_long | art_long | M |
| 81 | 109 | 0.682524455 | cat1 | from_448 | to_448 | art_long | art_long | M |
| 83 | 109 | 0.904445361 | cat1 | from_449 | to_449 | art_long | art_long | M |
| 87 | 109 | 0.999890826 | cat1 | from_450 | to_450 | art_long | art_long | M |
| 95 | 109 | 0.999890826 | cat1 | from_451 | to_451 | art_long | art_long | M |
| 97 | 109 | 0.66706457 | cat1 | from_452 | to_452 | art_long | art_long | M |
| 14 | 110 | 0.965027172 | cat1 | from_453 | to_453 | art_long | art_long | M |
| 24 | 110 | 0.965027172 | cat1 | from_454 | to_454 | art_long | art_long | M |
| 25 | 110 | 0.781131536 | cat1 | from_455 | to_455 | art_long | art_long | M |
| 28 | 110 | 0.968392195 | cat1 | from_456 | to_456 | art_long | art_long | M |
| 31 | 110 | 0.699238981 | cat1 | from_457 | to_457 | art_long | art_long | M |
| 33 | 110 | 0.714805687 | cat1 | from_458 | to_458 | art_long | art_long | M |
| 59 | 110 | 0.966718136 | cat1 | from_459 | to_459 | art_long | art_long | M |
| 73 | 110 | 0.965027172 | cat1 | from_460 | to_460 | art_long | art_long | M |
| 74 | 110 | 0.965027172 | cat1 | from_461 | to_461 | art_long | art_long | M |
| 76 | 110 | 0.933709258 | cat1 | from_462 | to_462 | art_long | art_long | M |
| 78 | 110 | 0.968392195 | cat1 | from_463 | to_463 | art_long | art_long | M |
| 90 | 110 | 0.932280569 | cat1 | from_464 | to_464 | art_long | art_long | M |
| 97 | 110 | 0.965027172 | cat1 | from_465 | to_465 | art_long | art_long | M |
| 14 | 111 | 0.668386791 | cat1 | from_466 | to_466 | art_long | art_long | M |
| 24 | 111 | 0.668386791 | cat1 | from_467 | to_467 | art_long | art_long | M |
| 28 | 111 | 0.667852051 | cat1 | from_468 | to_468 | art_long | art_long | M |
| 30 | 111 | 0.959602555 | cat1 | from_469 | to_469 | art_long | art_long | M |
| 47 | 111 | 0.99997309 | cat1 | from_470 | to_470 | art_long | art_long | M |
| 54 | 111 | 0.736243125 | cat1 | from_471 | to_471 | art_long | art_long | M |
| 55 | 111 | 0.99997309 | cat1 | from_472 | to_472 | art_long | art_long | M |
| 58 | 111 | 0.99997309 | cat1 | from_473 | to_473 | art_long | art_long | M |
| 59 | 111 | 0.668135854 | cat1 | from_474 | to_474 | art_long | art_long | M |
| 69 | 111 | 0.999972319 | cat1 | from_475 | to_475 | art_long | art_long | M |
| 72 | 111 | 0.922289612 | cat1 | from_476 | to_476 | art_long | art_long | M |
| 73 | 111 | 0.668386791 | cat1 | from_477 | to_477 | art_long | art_long | M |
| 74 | 111 | 0.668386791 | cat1 | from_478 | to_478 | art_long | art_long | M |
| 77 | 111 | 0.920757466 | cat1 | from_479 | to_479 | art_long | art_long | M |
| 78 | 111 | 0.667852051 | cat1 | from_480 | to_480 | art_long | art_long | M |
| 80 | 111 | 0.917425796 | cat1 | from_481 | to_481 | art_long | art_long | M |
| 81 | 111 | 0.679498705 | cat1 | from_482 | to_482 | art_long | art_long | M |
| 83 | 111 | 0.902700541 | cat1 | from_483 | to_483 | art_long | art_long | M |
| 87 | 111 | 0.99997309 | cat1 | from_484 | to_484 | art_long | art_long | M |
| 95 | 111 | 0.99997309 | cat1 | from_485 | to_485 | art_long | art_long | M |
| 97 | 111 | 0.668386791 | cat1 | from_486 | to_486 | art_long | art_long | M |
| 109 | 111 | 0.999972319 | cat1 | from_487 | to_487 | art_long | art_long | M |
| 14 | 112 | 0.744892559 | cat1 | from_488 | to_488 | art_long | art_long | M |
| 24 | 112 | 0.744892559 | cat1 | from_489 | to_489 | art_long | art_long | M |
| 25 | 112 | 0.999962251 | cat1 | from_490 | to_490 | art_long | art_long | M |
| 28 | 112 | 0.747982594 | cat1 | from_491 | to_491 | art_long | art_long | M |
| 31 | 112 | 0.924924322 | cat1 | from_492 | to_492 | art_long | art_long | M |
| 33 | 112 | 0.941407056 | cat1 | from_493 | to_493 | art_long | art_long | M |
| 54 | 112 | 0.715913999 | cat1 | from_494 | to_494 | art_long | art_long | M |
| 59 | 112 | 0.746442282 | cat1 | from_495 | to_495 | art_long | art_long | M |
| 73 | 112 | 0.744892559 | cat1 | from_496 | to_496 | art_long | art_long | M |
| 74 | 112 | 0.744892559 | cat1 | from_497 | to_497 | art_long | art_long | M |
| 76 | 112 | 0.935861007 | cat1 | from_498 | to_498 | art_long | art_long | M |
| 78 | 112 | 0.747982594 | cat1 | from_499 | to_499 | art_long | art_long | M |
| 80 | 112 | 0.666760941 | cat1 | from_500 | to_500 | art_long | art_long | M |
| 81 | 112 | 0.695508401 | cat1 | from_501 | to_501 | art_long | art_long | M |
| 83 | 112 | 0.666467553 | cat1 | from_502 | to_502 | art_long | art_long | M |
| 90 | 112 | 0.947309309 | cat1 | from_503 | to_503 | art_long | art_long | M |
| 97 | 112 | 0.744892559 | cat1 | from_504 | to_504 | art_long | art_long | M |
| 110 | 112 | 0.781704539 | cat1 | from_505 | to_505 | art_long | art_long | M |
| 4 | 113 | 0.667878223 | cat1 | from_506 | to_506 | art_long | art_long | M |
| 7 | 113 | 0.669965884 | cat1 | from_507 | to_507 | art_long | art_long | M |
| 22 | 113 | 0.701034412 | cat1 | from_508 | to_508 | art_long | art_long | M |
| 34 | 113 | 0.953031861 | cat1 | from_509 | to_509 | art_long | art_long | M |
| 65 | 113 | 0.942968194 | cat1 | from_510 | to_510 | art_long | art_long | M |
| 12 | 114 | 0.743873879 | cat1 | from_511 | to_511 | art_long | art_long | M |
| 13 | 114 | 0.945992459 | cat1 | from_512 | to_512 | art_long | art_long | M |
| 19 | 114 | 0.808483229 | cat1 | from_513 | to_513 | art_long | art_long | M |
| 20 | 114 | 0.78981352 | cat1 | from_514 | to_514 | art_long | art_long | M |
| 39 | 114 | 0.694489249 | cat1 | from_515 | to_515 | art_long | art_long | M |
| 40 | 114 | 0.807292982 | cat1 | from_516 | to_516 | art_long | art_long | M |
| 68 | 114 | 0.78129351 | cat1 | from_517 | to_517 | art_long | art_long | M |
| 79 | 114 | 0.823330239 | cat1 | from_518 | to_518 | art_long | art_long | M |

| | | | | | | | | |
|---|---|---|---|---|---|---|---|---|
| 26 | 115 | 0.999162246 | cat1 | from_519 | to_519 | art_long | art_long | M |
| 37 | 115 | 0.95384964 | cat1 | from_520 | to_520 | art_long | art_long | M |
| 5 | 116 | 0.966963769 | cat1 | from_521 | to_521 | art_long | art_long | M |
| 15 | 116 | 0.717876585 | cat1 | from_522 | to_522 | art_long | art_long | M |
| 63 | 116 | 0.999982635 | cat1 | from_523 | to_523 | art_long | art_long | M |
| 27 | 117 | 0.807182183 | cat1 | from_524 | to_524 | art_long | art_long | M |
| 34 | 117 | 0.713010523 | cat1 | from_525 | to_525 | art_long | art_long | M |
| 64 | 117 | 0.999986782 | cat1 | from_526 | to_526 | art_long | art_long | M |
| 91 | 117 | 0.722136503 | cat1 | from_527 | to_527 | art_long | art_long | M |
| 26 | 118 | 0.785608821 | cat1 | from_528 | to_528 | art_long | art_long | M |
| 37 | 118 | 0.773626792 | cat1 | from_529 | to_529 | art_long | art_long | M |
| 47 | 118 | 0.667479727 | cat1 | from_530 | to_530 | art_long | art_long | M |
| 55 | 118 | 0.667479727 | cat1 | from_531 | to_531 | art_long | art_long | M |
| 58 | 118 | 0.667479727 | cat1 | from_532 | to_532 | art_long | art_long | M |
| 87 | 118 | 0.667479727 | cat1 | from_533 | to_533 | art_long | art_long | M |
| 95 | 118 | 0.667479727 | cat1 | from_534 | to_534 | art_long | art_long | M |
| 111 | 118 | 0.666634228 | cat1 | from_535 | to_535 | art_long | art_long | M |
| 115 | 118 | 0.779505692 | cat1 | from_536 | to_536 | art_long | art_long | M |
| 14 | 119 | 0.668386791 | cat1 | from_537 | to_537 | art_long | art_long | M |
| 24 | 119 | 0.668386791 | cat1 | from_538 | to_538 | art_long | art_long | M |
| 28 | 119 | 0.667852051 | cat1 | from_539 | to_539 | art_long | art_long | M |
| 30 | 119 | 0.959602555 | cat1 | from_540 | to_540 | art_long | art_long | M |
| 47 | 119 | 0.99997309 | cat1 | from_541 | to_541 | art_long | art_long | M |
| 54 | 119 | 0.736243125 | cat1 | from_542 | to_542 | art_long | art_long | M |
| 55 | 119 | 0.99997309 | cat1 | from_543 | to_543 | art_long | art_long | M |
| 58 | 119 | 0.99997309 | cat1 | from_544 | to_544 | art_long | art_long | M |
| 59 | 119 | 0.668135854 | cat1 | from_545 | to_545 | art_long | art_long | M |
| 69 | 119 | 0.999972319 | cat1 | from_546 | to_546 | art_long | art_long | M |
| 72 | 119 | 0.922289612 | cat1 | from_547 | to_547 | art_long | art_long | M |
| 73 | 119 | 0.668386791 | cat1 | from_548 | to_548 | art_long | art_long | M |
| 74 | 119 | 0.668386791 | cat1 | from_549 | to_549 | art_long | art_long | M |
| 77 | 119 | 0.920757466 | cat1 | from_550 | to_550 | art_long | art_long | M |
| 78 | 119 | 0.667852051 | cat1 | from_551 | to_551 | art_long | art_long | M |
| 80 | 119 | 0.917425796 | cat1 | from_552 | to_552 | art_long | art_long | M |
| 81 | 119 | 0.679498705 | cat1 | from_553 | to_553 | art_long | art_long | M |
| 83 | 119 | 0.902700541 | cat1 | from_554 | to_554 | art_long | art_long | M |
| 87 | 119 | 0.99997309 | cat1 | from_555 | to_555 | art_long | art_long | M |
| 95 | 119 | 0.99997309 | cat1 | from_556 | to_556 | art_long | art_long | M |
| 97 | 119 | 0.668386791 | cat1 | from_557 | to_557 | art_long | art_long | M |
| 109 | 119 | 0.999972319 | cat1 | from_558 | to_558 | art_long | art_long | M |
| 111 | 119 | 1 | cat1 | from_559 | to_559 | art_long | art_long | M |
| 118 | 119 | 0.666634228 | cat1 | from_560 | to_560 | art_long | art_long | M |
| 30 | 120 | 0.927822312 | cat1 | from_561 | to_561 | art_long | art_long | M |
| 47 | 120 | 0.919578628 | cat1 | from_562 | to_562 | art_long | art_long | M |
| 54 | 120 | 0.941896354 | cat1 | from_563 | to_563 | art_long | art_long | M |
| 55 | 120 | 0.919578628 | cat1 | from_564 | to_564 | art_long | art_long | M |
| 58 | 120 | 0.919578628 | cat1 | from_565 | to_565 | art_long | art_long | M |
| 69 | 120 | 0.921902451 | cat1 | from_566 | to_566 | art_long | art_long | M |
| 72 | 120 | 0.999962474 | cat1 | from_567 | to_567 | art_long | art_long | M |
| 77 | 120 | 1 | cat1 | from_568 | to_568 | art_long | art_long | M |
| 80 | 120 | 0.999844556 | cat1 | from_569 | to_569 | art_long | art_long | M |
| 81 | 120 | 0.907434674 | cat1 | from_570 | to_570 | art_long | art_long | M |
| 83 | 120 | 0.996591579 | cat1 | from_571 | to_571 | art_long | art_long | M |
| 87 | 120 | 0.919578628 | cat1 | from_572 | to_572 | art_long | art_long | M |
| 95 | 120 | 0.919578628 | cat1 | from_573 | to_573 | art_long | art_long | M |
| 109 | 120 | 0.921902451 | cat1 | from_574 | to_574 | art_long | art_long | M |
| 111 | 120 | 0.920757466 | cat1 | from_575 | to_575 | art_long | art_long | M |
| 119 | 120 | 0.920757466 | cat1 | from_576 | to_576 | art_long | art_long | M |
| 29 | 121 | 0.671098849 | cat1 | from_577 | to_577 | art_long | art_long | M |
| 41 | 121 | 0.891647016 | cat1 | from_578 | to_578 | art_long | art_long | M |
| 71 | 121 | 0.691231079 | cat1 | from_579 | to_579 | art_long | art_long | M |
| 82 | 121 | 0.790845239 | cat1 | from_580 | to_580 | art_long | art_long | M |
| 89 | 121 | 0.841740096 | cat1 | from_581 | to_581 | art_long | art_long | M |
| 96 | 121 | 0.883112742 | cat1 | from_582 | to_582 | art_long | art_long | M |
| 98 | 121 | 0.999992127 | cat1 | from_583 | to_583 | art_long | art_long | M |
| 14 | 122 | 1 | cat1 | from_584 | to_584 | art_long | art_long | M |
| 24 | 122 | 1 | cat1 | from_585 | to_585 | art_long | art_long | M |
| 25 | 122 | 0.742067098 | cat1 | from_586 | to_586 | art_long | art_long | M |
| 28 | 122 | 0.999913472 | cat1 | from_587 | to_587 | art_long | art_long | M |
| 47 | 122 | 0.669654246 | cat1 | from_588 | to_588 | art_long | art_long | M |
| 55 | 122 | 0.669654246 | cat1 | from_589 | to_589 | art_long | art_long | M |
| 58 | 122 | 0.669654246 | cat1 | from_590 | to_590 | art_long | art_long | M |
| 59 | 122 | 0.999978687 | cat1 | from_591 | to_591 | art_long | art_long | M |
| 69 | 122 | 0.66706457 | cat1 | from_592 | to_592 | art_long | art_long | M |
| 73 | 122 | 1 | cat1 | from_593 | to_593 | art_long | art_long | M |
| 74 | 122 | 1 | cat1 | from_594 | to_594 | art_long | art_long | M |
| 76 | 122 | 0.932211263 | cat1 | from_595 | to_595 | art_long | art_long | M |
| 78 | 122 | 0.999913472 | cat1 | from_596 | to_596 | art_long | art_long | M |
| 87 | 122 | 0.669654246 | cat1 | from_597 | to_597 | art_long | art_long | M |
| 90 | 122 | 0.870862898 | cat1 | from_598 | to_598 | art_long | art_long | M |
| 95 | 122 | 0.669654246 | cat1 | from_599 | to_599 | art_long | art_long | M |
| 97 | 122 | 1 | cat1 | from_600 | to_600 | art_long | art_long | M |
| 109 | 122 | 0.66706457 | cat1 | from_601 | to_601 | art_long | art_long | M |
| 110 | 122 | 0.965027172 | cat1 | from_602 | to_602 | art_long | art_long | M |
| 111 | 122 | 0.668386791 | cat1 | from_603 | to_603 | art_long | art_long | M |
| 112 | 122 | 0.744892559 | cat1 | from_604 | to_604 | art_long | art_long | M |
| 119 | 122 | 0.668386791 | cat1 | from_605 | to_605 | art_long | art_long | M |
| 34 | 123 | 0.702141191 | cat1 | from_606 | to_606 | art_long | art_long | M |
| 49 | 123 | 0.83416185 | cat1 | from_607 | to_607 | art_long | art_long | M |
| 56 | 123 | 0.820402628 | cat1 | from_608 | to_608 | art_long | art_long | M |
| 57 | 123 | 0.820402628 | cat1 | from_609 | to_609 | art_long | art_long | M |
| 70 | 123 | 0.960122287 | cat1 | from_610 | to_610 | art_long | art_long | M |
| 75 | 123 | 0.836047119 | cat1 | from_611 | to_611 | art_long | art_long | M |
| 85 | 123 | 1 | cat1 | from_612 | to_612 | art_long | art_long | M |
| 29 | 124 | 0.829224009 | cat1 | from_613 | to_613 | art_long | art_long | M |
| 66 | 124 | 0.674713923 | cat1 | from_614 | to_614 | art_long | art_long | M |
| 67 | 124 | 0.674713923 | cat1 | from_615 | to_615 | art_long | art_long | M |
| 71 | 124 | 0.847953948 | cat1 | from_616 | to_616 | art_long | art_long | M |
| 82 | 124 | 0.683880041 | cat1 | from_617 | to_617 | art_long | art_long | M |
| 89 | 124 | 0.758396109 | cat1 | from_618 | to_618 | art_long | art_long | M |
| 6 | 125 | 0.772619705 | cat1 | from_619 | to_619 | art_long | art_long | M |
| 8 | 125 | 0.809364035 | cat1 | from_620 | to_620 | art_long | art_long | M |
| 9 | 125 | 0.788905939 | cat1 | from_621 | to_621 | art_long | art_long | M |
| 10 | 125 | 0.789230334 | cat1 | from_622 | to_622 | art_long | art_long | M |

| | | | | | | | | |
|---|---|---|---|---|---|---|---|---|
| 23 | 125 | 0.673995277 | cat1 | from_623 | to_623 | art_long | art_long | M |
| 65 | 125 | 0.827652221 | cat1 | from_624 | to_624 | art_long | art_long | M |
| 84 | 125 | 0.795888019 | cat1 | from_625 | to_625 | art_long | art_long | M |
| 88 | 125 | 0.671644218 | cat1 | from_626 | to_626 | art_long | art_long | M |
| 8 | 126 | 0.722552965 | cat1 | from_627 | to_627 | art_long | art_long | M |
| 9 | 126 | 0.949186732 | cat1 | from_628 | to_628 | art_long | art_long | M |
| 10 | 126 | 0.948005924 | cat1 | from_629 | to_629 | art_long | art_long | M |
| 22 | 126 | 0.937715137 | cat1 | from_630 | to_630 | art_long | art_long | M |
| 23 | 126 | 0.999969211 | cat1 | from_631 | to_631 | art_long | art_long | M |
| 65 | 126 | 0.712336395 | cat1 | from_632 | to_632 | art_long | art_long | M |
| 88 | 126 | 0.99996991 | cat1 | from_633 | to_633 | art_long | art_long | M |
| 91 | 126 | 0.701426822 | cat1 | from_634 | to_634 | art_long | art_long | M |
| 125 | 126 | 0.672833474 | cat1 | from_635 | to_635 | art_long | art_long | M |
| 4 | 127 | 0.731462073 | cat1 | from_636 | to_636 | art_long | art_long | M |
| 5 | 127 | 0.686857279 | cat1 | from_637 | to_637 | art_long | art_long | M |
| 7 | 127 | 0.730796075 | cat1 | from_638 | to_638 | art_long | art_long | M |
| 22 | 127 | 0.728245552 | cat1 | from_639 | to_639 | art_long | art_long | M |
| 34 | 127 | 0.952559569 | cat1 | from_640 | to_640 | art_long | art_long | M |
| 64 | 127 | 0.728049513 | cat1 | from_641 | to_641 | art_long | art_long | M |
| 65 | 127 | 0.942722305 | cat1 | from_642 | to_642 | art_long | art_long | M |
| 70 | 127 | 0.679163144 | cat1 | from_643 | to_643 | art_long | art_long | M |
| 85 | 127 | 0.716959213 | cat1 | from_644 | to_644 | art_long | art_long | M |
| 113 | 127 | 0.963714391 | cat1 | from_645 | to_645 | art_long | art_long | M |
| 117 | 127 | 0.728361201 | cat1 | from_646 | to_646 | art_long | art_long | M |
| 123 | 127 | 0.716959213 | cat1 | from_647 | to_647 | art_long | art_long | M |
| 44 | 128 | 0.999885344 | cat1 | from_648 | to_648 | art_long | art_long | M |
| 46 | 128 | 0.999885344 | cat1 | from_649 | to_649 | art_long | art_long | M |
| 52 | 128 | 0.87555592 | cat1 | from_650 | to_650 | art_long | art_long | M |
| 60 | 128 | 0.997282276 | cat1 | from_651 | to_651 | art_long | art_long | M |
| 61 | 128 | 0.917821861 | cat1 | from_652 | to_652 | art_long | art_long | M |
| 99 | 128 | 0.919370561 | cat1 | from_653 | to_653 | art_long | art_long | M |
| 102 | 128 | 0.999722904 | cat1 | from_654 | to_654 | art_long | art_long | M |
| 105 | 128 | 0.727129339 | cat1 | from_655 | to_655 | art_long | art_long | M |
| 106 | 128 | 0.999885344 | cat1 | from_656 | to_656 | art_long | art_long | M |
| 26 | 129 | 0.835736339 | cat1 | from_657 | to_657 | art_long | art_long | M |
| 37 | 129 | 0.948286777 | cat1 | from_658 | to_658 | art_long | art_long | M |
| 115 | 129 | 0.837048721 | cat1 | from_659 | to_659 | art_long | art_long | M |
| 118 | 129 | 0.671771882 | cat1 | from_660 | to_660 | art_long | art_long | M |
| 26 | 130 | 0.958465101 | cat1 | from_661 | to_661 | art_long | art_long | M |
| 37 | 130 | 0.999906932 | cat1 | from_662 | to_662 | art_long | art_long | M |
| 115 | 130 | 0.955091958 | cat1 | from_663 | to_663 | art_long | art_long | M |
| 118 | 130 | 0.772399897 | cat1 | from_664 | to_664 | art_long | art_long | M |
| 129 | 130 | 0.950095135 | cat1 | from_665 | to_665 | art_long | art_long | M |
| 26 | 131 | 0.785460564 | cat1 | from_666 | to_666 | art_long | art_long | M |
| 37 | 131 | 0.773871767 | cat1 | from_667 | to_667 | art_long | art_long | M |
| 47 | 131 | 0.667914204 | cat1 | from_668 | to_668 | art_long | art_long | M |
| 55 | 131 | 0.667914204 | cat1 | from_669 | to_669 | art_long | art_long | M |
| 58 | 131 | 0.667914204 | cat1 | from_670 | to_670 | art_long | art_long | M |
| 87 | 131 | 0.667914204 | cat1 | from_671 | to_671 | art_long | art_long | M |
| 95 | 131 | 0.667914204 | cat1 | from_672 | to_672 | art_long | art_long | M |
| 111 | 131 | 0.667030231 | cat1 | from_673 | to_673 | art_long | art_long | M |
| 115 | 131 | 0.779140478 | cat1 | from_674 | to_674 | art_long | art_long | M |
| 118 | 131 | 0.999985421 | cat1 | from_675 | to_675 | art_long | art_long | M |
| 119 | 131 | 0.667030231 | cat1 | from_676 | to_676 | art_long | art_long | M |
| 129 | 131 | 0.671351146 | cat1 | from_677 | to_677 | art_long | art_long | M |
| 130 | 131 | 0.772572633 | cat1 | from_678 | to_678 | art_long | art_long | M |
| 17 | 132 | 0.792328802 | cat1 | from_679 | to_679 | art_long | art_long | M |
| 35 | 132 | 1 | cat1 | from_680 | to_680 | art_long | art_long | M |
| 36 | 132 | 0.792323955 | cat1 | from_681 | to_681 | art_long | art_long | M |
| 38 | 132 | 0.999605701 | cat1 | from_682 | to_682 | art_long | art_long | M |
| 48 | 132 | 1 | cat1 | from_683 | to_683 | art_long | art_long | M |
| 50 | 132 | 0.812968063 | cat1 | from_684 | to_684 | art_long | art_long | M |
| 104 | 132 | 1 | cat1 | from_685 | to_685 | art_long | art_long | M |
| 17 | 133 | 0.99998586 | cat1 | from_686 | to_686 | art_long | art_long | M |
| 35 | 133 | 0.792323955 | cat1 | from_687 | to_687 | art_long | art_long | M |
| 36 | 133 | 1 | cat1 | from_688 | to_688 | art_long | art_long | M |
| 38 | 133 | 0.78778631 | cat1 | from_689 | to_689 | art_long | art_long | M |
| 48 | 133 | 0.792323955 | cat1 | from_690 | to_690 | art_long | art_long | M |
| 104 | 133 | 0.792323955 | cat1 | from_691 | to_691 | art_long | art_long | M |
| 132 | 133 | 0.792323955 | cat1 | from_692 | to_692 | art_long | art_long | M |
| 17 | 134 | 0.99998586 | cat1 | from_693 | to_693 | art_long | art_long | M |
| 35 | 134 | 0.792323955 | cat1 | from_694 | to_694 | art_long | art_long | M |
| 36 | 134 | 1 | cat1 | from_695 | to_695 | art_long | art_long | M |
| 38 | 134 | 0.78778631 | cat1 | from_696 | to_696 | art_long | art_long | M |
| 48 | 134 | 0.792323955 | cat1 | from_697 | to_697 | art_long | art_long | M |
| 104 | 134 | 0.792323955 | cat1 | from_698 | to_698 | art_long | art_long | M |
| 132 | 134 | 0.792323955 | cat1 | from_699 | to_699 | art_long | art_long | M |
| 133 | 134 | 1 | cat1 | from_700 | to_700 | art_long | art_long | M |
| 44 | 135 | 1 | cat1 | from_701 | to_701 | art_long | art_long | M |
| 46 | 135 | 1 | cat1 | from_702 | to_702 | art_long | art_long | M |
| 52 | 135 | 0.870308282 | cat1 | from_703 | to_703 | art_long | art_long | M |
| 60 | 135 | 0.996052292 | cat1 | from_704 | to_704 | art_long | art_long | M |
| 61 | 135 | 0.915387767 | cat1 | from_705 | to_705 | art_long | art_long | M |
| 99 | 135 | 0.917068621 | cat1 | from_706 | to_706 | art_long | art_long | M |
| 102 | 135 | 0.999251827 | cat1 | from_707 | to_707 | art_long | art_long | M |
| 105 | 135 | 0.723041928 | cat1 | from_708 | to_708 | art_long | art_long | M |
| 106 | 135 | 1 | cat1 | from_709 | to_709 | art_long | art_long | M |
| 128 | 135 | 0.999885344 | cat1 | from_710 | to_710 | art_long | art_long | M |
| 26 | 136 | 0.955094431 | cat1 | from_711 | to_711 | art_long | art_long | M |
| 37 | 136 | 0.988040658 | cat1 | from_712 | to_712 | art_long | art_long | M |
| 115 | 136 | 0.957486398 | cat1 | from_713 | to_713 | art_long | art_long | M |
| 118 | 136 | 0.751321948 | cat1 | from_714 | to_714 | art_long | art_long | M |
| 129 | 136 | 0.958381664 | cat1 | from_715 | to_715 | art_long | art_long | M |
| 130 | 136 | 0.990052346 | cat1 | from_716 | to_716 | art_long | art_long | M |
| 131 | 136 | 0.750747798 | cat1 | from_717 | to_717 | art_long | art_long | M |
| 26 | 137 | 0.714150925 | cat1 | from_718 | to_718 | art_long | art_long | M |
| 37 | 137 | 0.771004388 | cat1 | from_719 | to_719 | art_long | art_long | M |
| 115 | 137 | 0.712560363 | cat1 | from_720 | to_720 | art_long | art_long | M |
| 118 | 137 | 0.95543664 | cat1 | from_721 | to_721 | art_long | art_long | M |
| 129 | 137 | 0.763676779 | cat1 | from_722 | to_722 | art_long | art_long | M |
| 130 | 137 | 0.771483527 | cat1 | from_723 | to_723 | art_long | art_long | M |
| 131 | 137 | 0.95481243 | cat1 | from_724 | to_724 | art_long | art_long | M |
| 136 | 137 | 0.768009974 | cat1 | from_725 | to_725 | art_long | art_long | M |
| 17 | 138 | 0.991336988 | cat1 | from_726 | to_726 | art_long | art_long | M |

| | | | | | | | | |
|---|---|---|---|---|---|---|---|---|
| 35 | 138 | 0.785307858 | cat1 | from_727 | to_727 | art_long | art_long | M |
| 36 | 138 | 0.990624499 | cat1 | from_728 | to_728 | art_long | art_long | M |
| 38 | 138 | 0.784533736 | cat1 | from_729 | to_729 | art_long | art_long | M |
| 48 | 138 | 0.785307858 | cat1 | from_730 | to_730 | art_long | art_long | M |
| 104 | 138 | 0.785307858 | cat1 | from_731 | to_731 | art_long | art_long | M |
| 132 | 138 | 0.785307858 | cat1 | from_732 | to_732 | art_long | art_long | M |
| 133 | 138 | 0.990624499 | cat1 | from_733 | to_733 | art_long | art_long | M |
| 134 | 138 | 0.990624499 | cat1 | from_734 | to_734 | art_long | art_long | M |
| 42 | 139 | 0.943764559 | cat1 | from_735 | to_735 | art_long | art_long | M |
| 43 | 139 | 0.920665622 | cat1 | from_736 | to_736 | art_long | art_long | M |
| 45 | 139 | 0.943764559 | cat1 | from_737 | to_737 | art_long | art_long | M |
| 62 | 139 | 0.948108887 | cat1 | from_738 | to_738 | art_long | art_long | M |
| 92 | 139 | 0.948108887 | cat1 | from_739 | to_739 | art_long | art_long | M |
| 100 | 139 | 0.943764559 | cat1 | from_740 | to_740 | art_long | art_long | M |
| 101 | 139 | 0.943764559 | cat1 | from_741 | to_741 | art_long | art_long | M |
| 107 | 139 | 0.943764559 | cat1 | from_742 | to_742 | art_long | art_long | M |
| 30 | 140 | 0.928689652 | cat1 | from_743 | to_743 | art_long | art_long | M |
| 47 | 140 | 0.917895047 | cat1 | from_744 | to_744 | art_long | art_long | M |
| 54 | 140 | 0.943166087 | cat1 | from_745 | to_745 | art_long | art_long | M |
| 55 | 140 | 0.917895047 | cat1 | from_746 | to_746 | art_long | art_long | M |
| 58 | 140 | 0.917895047 | cat1 | from_747 | to_747 | art_long | art_long | M |
| 69 | 140 | 0.920346089 | cat1 | from_748 | to_748 | art_long | art_long | M |
| 72 | 140 | 0.99984813 | cat1 | from_749 | to_749 | art_long | art_long | M |
| 77 | 140 | 0.999961587 | cat1 | from_750 | to_750 | art_long | art_long | M |
| 80 | 140 | 0.999960686 | cat1 | from_751 | to_751 | art_long | art_long | M |
| 81 | 140 | 0.910025801 | cat1 | from_752 | to_752 | art_long | art_long | M |
| 83 | 140 | 0.99727635 | cat1 | from_753 | to_753 | art_long | art_long | M |
| 87 | 140 | 0.917895047 | cat1 | from_754 | to_754 | art_long | art_long | M |
| 95 | 140 | 0.917895047 | cat1 | from_755 | to_755 | art_long | art_long | M |
| 109 | 140 | 0.920346089 | cat1 | from_756 | to_756 | art_long | art_long | M |
| 111 | 140 | 0.919137002 | cat1 | from_757 | to_757 | art_long | art_long | M |
| 112 | 140 | 0.666582724 | cat1 | from_758 | to_758 | art_long | art_long | M |
| 119 | 140 | 0.919137002 | cat1 | from_759 | to_759 | art_long | art_long | M |
| 120 | 140 | 0.999961587 | cat1 | from_760 | to_760 | art_long | art_long | M |
| 26 | 141 | 0.714693452 | cat1 | from_761 | to_761 | art_long | art_long | M |
| 37 | 141 | 0.771925959 | cat1 | from_762 | to_762 | art_long | art_long | M |
| 115 | 141 | 0.712834115 | cat1 | from_763 | to_763 | art_long | art_long | M |
| 118 | 141 | 0.956280892 | cat1 | from_764 | to_764 | art_long | art_long | M |
| 129 | 141 | 0.763612616 | cat1 | from_765 | to_765 | art_long | art_long | M |
| 130 | 141 | 0.772316022 | cat1 | from_766 | to_766 | art_long | art_long | M |
| 131 | 141 | 0.955692131 | cat1 | from_767 | to_767 | art_long | art_long | M |
| 136 | 141 | 0.767914739 | cat1 | from_768 | to_768 | art_long | art_long | M |
| 137 | 141 | 0.999978029 | cat1 | from_769 | to_769 | art_long | art_long | M |
| 26 | 142 | 0.836503686 | cat1 | from_770 | to_770 | art_long | art_long | M |
| 37 | 142 | 0.949458853 | cat1 | from_771 | to_771 | art_long | art_long | M |
| 115 | 142 | 0.837501451 | cat1 | from_772 | to_772 | art_long | art_long | M |
| 118 | 142 | 0.673374107 | cat1 | from_773 | to_773 | art_long | art_long | M |
| 129 | 142 | 0.999970131 | cat1 | from_774 | to_774 | art_long | art_long | M |
| 130 | 142 | 0.951162946 | cat1 | from_775 | to_775 | art_long | art_long | M |
| 131 | 142 | 0.672993989 | cat1 | from_776 | to_776 | art_long | art_long | M |
| 136 | 142 | 0.958362536 | cat1 | from_777 | to_777 | art_long | art_long | M |
| 137 | 142 | 0.764245719 | cat1 | from_778 | to_778 | art_long | art_long | M |
| 141 | 142 | 0.764232465 | cat1 | from_779 | to_779 | art_long | art_long | M |
| 17 | 143 | 0.683163883 | cat1 | from_780 | to_780 | art_long | art_long | M |
| 35 | 143 | 0.929749797 | cat1 | from_781 | to_781 | art_long | art_long | M |
| 36 | 143 | 0.681259199 | cat1 | from_782 | to_782 | art_long | art_long | M |
| 38 | 143 | 0.939721741 | cat1 | from_783 | to_783 | art_long | art_long | M |
| 48 | 143 | 0.929749797 | cat1 | from_784 | to_784 | art_long | art_long | M |
| 50 | 143 | 0.848058915 | cat1 | from_785 | to_785 | art_long | art_long | M |
| 104 | 143 | 0.929749797 | cat1 | from_786 | to_786 | art_long | art_long | M |
| 132 | 143 | 0.929749797 | cat1 | from_787 | to_787 | art_long | art_long | M |
| 133 | 143 | 0.681259199 | cat1 | from_788 | to_788 | art_long | art_long | M |
| 134 | 143 | 0.681259199 | cat1 | from_789 | to_789 | art_long | art_long | M |
| 138 | 143 | 0.724049341 | cat1 | from_790 | to_790 | art_long | art_long | M |
| 26 | 144 | 0.785608821 | cat1 | from_791 | to_791 | art_long | art_long | M |
| 37 | 144 | 0.773626792 | cat1 | from_792 | to_792 | art_long | art_long | M |
| 47 | 144 | 0.667479727 | cat1 | from_793 | to_793 | art_long | art_long | M |
| 55 | 144 | 0.667479727 | cat1 | from_794 | to_794 | art_long | art_long | M |
| 58 | 144 | 0.667479727 | cat1 | from_795 | to_795 | art_long | art_long | M |
| 87 | 144 | 0.667479727 | cat1 | from_796 | to_796 | art_long | art_long | M |
| 95 | 144 | 0.667479727 | cat1 | from_797 | to_797 | art_long | art_long | M |
| 111 | 144 | 0.666634228 | cat1 | from_798 | to_798 | art_long | art_long | M |
| 115 | 144 | 0.779505692 | cat1 | from_799 | to_799 | art_long | art_long | M |
| 118 | 144 | 1 | cat1 | from_800 | to_800 | art_long | art_long | M |
| 119 | 144 | 0.666634228 | cat1 | from_801 | to_801 | art_long | art_long | M |
| 129 | 144 | 0.671771882 | cat1 | from_802 | to_802 | art_long | art_long | M |
| 130 | 144 | 0.772399897 | cat1 | from_803 | to_803 | art_long | art_long | M |
| 131 | 144 | 0.999985421 | cat1 | from_804 | to_804 | art_long | art_long | M |
| 136 | 144 | 0.751321948 | cat1 | from_805 | to_805 | art_long | art_long | M |
| 137 | 144 | 0.95543664 | cat1 | from_806 | to_806 | art_long | art_long | M |
| 141 | 144 | 0.956280892 | cat1 | from_807 | to_807 | art_long | art_long | M |
| 142 | 144 | 0.673374107 | cat1 | from_808 | to_808 | art_long | art_long | M |
| 8 | 145 | 0.780573001 | cat1 | from_809 | to_809 | art_long | art_long | M |
| 93 | 145 | 0.847393097 | cat1 | from_810 | to_810 | art_long | art_long | M |
| 49 | 146 | 0.955085217 | cat1 | from_811 | to_811 | art_long | art_long | M |
| 56 | 146 | 0.800339111 | cat1 | from_812 | to_812 | art_long | art_long | M |
| 57 | 146 | 0.800339111 | cat1 | from_813 | to_813 | art_long | art_long | M |
| 65 | 146 | 0.672520754 | cat1 | from_814 | to_814 | art_long | art_long | M |
| 70 | 146 | 1 | cat1 | from_815 | to_815 | art_long | art_long | M |
| 75 | 146 | 0.95606854 | cat1 | from_816 | to_816 | art_long | art_long | M |
| 84 | 146 | 0.708206658 | cat1 | from_817 | to_817 | art_long | art_long | M |
| 85 | 146 | 0.960122287 | cat1 | from_818 | to_818 | art_long | art_long | M |
| 123 | 146 | 0.960122287 | cat1 | from_819 | to_819 | art_long | art_long | M |
| 127 | 146 | 0.679163144 | cat1 | from_820 | to_820 | art_long | art_long | M |
| 42 | 147 | 1 | cat1 | from_821 | to_821 | art_long | art_long | M |
| 43 | 147 | 0.92933654 | cat1 | from_822 | to_822 | art_long | art_long | M |
| 45 | 147 | 1 | cat1 | from_823 | to_823 | art_long | art_long | M |
| 62 | 147 | 0.999910267 | cat1 | from_824 | to_824 | art_long | art_long | M |
| 92 | 147 | 0.999910267 | cat1 | from_825 | to_825 | art_long | art_long | M |
| 100 | 147 | 1 | cat1 | from_826 | to_826 | art_long | art_long | M |
| 101 | 147 | 1 | cat1 | from_827 | to_827 | art_long | art_long | M |
| 107 | 147 | 1 | cat1 | from_828 | to_828 | art_long | art_long | M |
| 139 | 147 | 0.943764559 | cat1 | from_829 | to_829 | art_long | art_long | M |
| 44 | 148 | 0.917068621 | cat1 | from_830 | to_830 | art_long | art_long | M |

| | | | | | | | | |
|---|---|---|---|---|---|---|---|---|
| 46 | 148 | 0.917068621 | cat1 | from_831 | to_831 | art_long | art_long | M |
| 52 | 148 | 0.728730331 | cat1 | from_832 | to_832 | art_long | art_long | M |
| 60 | 148 | 0.927559009 | cat1 | from_833 | to_833 | art_long | art_long | M |
| 61 | 148 | 0.999960565 | cat1 | from_834 | to_834 | art_long | art_long | M |
| 99 | 148 | 1 | cat1 | from_835 | to_835 | art_long | art_long | M |
| 102 | 148 | 0.922530372 | cat1 | from_836 | to_836 | art_long | art_long | M |
| 105 | 148 | 0.938522996 | cat1 | from_837 | to_837 | art_long | art_long | M |
| 106 | 148 | 0.917068621 | cat1 | from_838 | to_838 | art_long | art_long | M |
| 128 | 148 | 0.919370561 | cat1 | from_839 | to_839 | art_long | art_long | M |
| 135 | 148 | 0.917068621 | cat1 | from_840 | to_840 | art_long | art_long | M |
| 42 | 149 | 1 | cat1 | from_841 | to_841 | art_long | art_long | M |
| 43 | 149 | 0.92933654 | cat1 | from_842 | to_842 | art_long | art_long | M |
| 45 | 149 | 1 | cat1 | from_843 | to_843 | art_long | art_long | M |
| 62 | 149 | 0.999910267 | cat1 | from_844 | to_844 | art_long | art_long | M |
| 92 | 149 | 0.999910267 | cat1 | from_845 | to_845 | art_long | art_long | M |
| 100 | 149 | 1 | cat1 | from_846 | to_846 | art_long | art_long | M |
| 101 | 149 | 1 | cat1 | from_847 | to_847 | art_long | art_long | M |
| 107 | 149 | 1 | cat1 | from_848 | to_848 | art_long | art_long | M |
| 139 | 149 | 0.943764559 | cat1 | from_849 | to_849 | art_long | art_long | M |
| 147 | 149 | 1 | cat1 | from_850 | to_850 | art_long | art_long | M |
| 3 | 150 | 0.709881465 | cat1 | from_851 | to_851 | art_long | art_long | M |
| 29 | 150 | 0.836485115 | cat1 | from_852 | to_852 | art_long | art_long | M |
| 41 | 150 | 0.751450276 | cat1 | from_853 | to_853 | art_long | art_long | M |
| 66 | 150 | 0.760218664 | cat1 | from_854 | to_854 | art_long | art_long | M |
| 67 | 150 | 0.760218664 | cat1 | from_855 | to_855 | art_long | art_long | M |
| 71 | 150 | 0.869789995 | cat1 | from_856 | to_856 | art_long | art_long | M |
| 82 | 150 | 0.970679347 | cat1 | from_857 | to_857 | art_long | art_long | M |
| 89 | 150 | 1 | cat1 | from_858 | to_858 | art_long | art_long | M |
| 96 | 150 | 0.753050965 | cat1 | from_859 | to_859 | art_long | art_long | M |
| 98 | 150 | 0.841718207 | cat1 | from_860 | to_860 | art_long | art_long | M |
| 121 | 150 | 0.841740096 | cat1 | from_861 | to_861 | art_long | art_long | M |
| 124 | 150 | 0.758396109 | cat1 | from_862 | to_862 | art_long | art_long | M |
| 1 | 151 | -0.834770795 | cat1 | from_863 | to_863 | art_long | art_long | M |
| 93 | 152 | 0.870844083 | cat1 | from_864 | to_864 | art_long | art_long | M |
| 105 | 152 | 0.823935814 | cat1 | from_865 | to_865 | art_long | art_long | M |
| 151 | 152 | 0.766479675 | cat1 | from_866 | to_866 | art_long | art_long | M |
| 61 | 153 | 0.684419483 | cat1 | from_867 | to_867 | art_long | art_long | M |
| 93 | 153 | 0.821058974 | cat1 | from_868 | to_868 | art_long | art_long | M |
| 99 | 153 | 0.686308825 | cat1 | from_869 | to_869 | art_long | art_long | M |
| 105 | 153 | 0.805110176 | cat1 | from_870 | to_870 | art_long | art_long | M |
| 148 | 153 | 0.686308825 | cat1 | from_871 | to_871 | art_long | art_long | M |
| 151 | 153 | 0.785485677 | cat1 | from_872 | to_872 | art_long | art_long | M |
| 152 | 153 | 0.983237036 | cat1 | from_873 | to_873 | art_long | art_long | M |
| 17 | 154 | 0.833979915 | cat1 | from_874 | to_874 | art_long | art_short | J |
| 36 | 154 | 0.831033725 | cat1 | from_875 | to_875 | art_long | art_short | J |
| 38 | 154 | 0.671505867 | cat1 | from_876 | to_876 | art_long | art_short | J |
| 91 | 154 | -0.680455596 | cat1 | from_877 | to_877 | art_long | art_short | J |
| 133 | 154 | 0.831033725 | cat1 | from_878 | to_878 | art_long | art_short | J |
| 134 | 154 | 0.831033725 | cat1 | from_879 | to_879 | art_long | art_short | J |
| 138 | 154 | 0.899229507 | cat1 | from_880 | to_880 | art_long | art_short | J |
| 143 | 154 | 0.766375546 | cat1 | from_881 | to_881 | art_long | art_short | J |
| 2 | 155 | 0.970245621 | cat1 | from_882 | to_882 | art_long | art_short | J |
| 16 | 155 | 0.726716636 | cat1 | from_883 | to_883 | art_long | art_short | J |
| 18 | 155 | 0.726716636 | cat1 | from_884 | to_884 | art_long | art_short | J |
| 39 | 155 | 0.678599617 | cat1 | from_885 | to_885 | art_long | art_short | J |
| 53 | 155 | 0.981497378 | cat1 | from_886 | to_886 | art_long | art_short | J |
| 8 | 156 | 0.746450949 | cat1 | from_887 | to_887 | art_long | art_short | J |
| 79 | 156 | 0.688255799 | cat1 | from_888 | to_888 | art_long | art_short | J |
| 114 | 156 | 0.801802061 | cat1 | from_889 | to_889 | art_long | art_short | J |
| 125 | 156 | 0.733881235 | cat1 | from_890 | to_890 | art_long | art_short | J |
| 6 | 157 | 0.839915778 | cat1 | from_891 | to_891 | art_long | art_short | J |
| 8 | 157 | 0.814793452 | cat1 | from_892 | to_892 | art_long | art_short | J |
| 125 | 157 | 0.756791062 | cat1 | from_893 | to_893 | art_long | art_short | J |
| 156 | 157 | 0.677747253 | cat1 | from_894 | to_894 | art_short | art_short | H |
| 8 | 158 | 0.74634914 | cat1 | from_895 | to_895 | art_long | art_short | J |
| 13 | 158 | 0.666893501 | cat1 | from_896 | to_896 | art_long | art_short | J |
| 79 | 158 | 0.688955774 | cat1 | from_897 | to_897 | art_long | art_short | J |
| 114 | 158 | 0.802742799 | cat1 | from_898 | to_898 | art_long | art_short | J |
| 125 | 158 | 0.735469481 | cat1 | from_899 | to_899 | art_long | art_short | J |
| 156 | 158 | 0.999973879 | cat1 | from_900 | to_900 | art_short | art_short | H |
| 157 | 158 | 0.676167354 | cat1 | from_901 | to_901 | art_short | art_short | H |
| 4 | 159 | 0.860440995 | cat1 | from_902 | to_902 | art_long | art_short | J |
| 6 | 159 | 0.859039832 | cat1 | from_903 | to_903 | art_long | art_short | J |
| 7 | 159 | 0.86489273 | cat1 | from_904 | to_904 | art_long | art_short | J |
| 11 | 159 | 0.708682976 | cat1 | from_905 | to_905 | art_long | art_short | J |
| 137 | 159 | -0.66910886 | cat1 | from_906 | to_906 | art_long | art_short | J |
| 141 | 159 | -0.668850587 | cat1 | from_907 | to_907 | art_long | art_short | J |
| 4 | 160 | 0.958593283 | cat1 | from_908 | to_908 | art_long | art_short | J |
| 6 | 160 | 0.677409622 | cat1 | from_909 | to_909 | art_long | art_short | J |
| 7 | 160 | 0.961562144 | cat1 | from_910 | to_910 | art_long | art_short | J |
| 11 | 160 | 0.773120739 | cat1 | from_911 | to_911 | art_long | art_short | J |
| 34 | 160 | 0.711382909 | cat1 | from_912 | to_912 | art_long | art_short | J |
| 113 | 160 | 0.697241832 | cat1 | from_913 | to_913 | art_long | art_short | J |
| 127 | 160 | 0.684412498 | cat1 | from_914 | to_914 | art_long | art_short | J |
| 159 | 160 | 0.945546783 | cat1 | from_915 | to_915 | art_short | art_short | H |
| 9 | 161 | 0.814010689 | cat1 | from_916 | to_916 | art_long | art_short | J |
| 10 | 161 | 0.815096198 | cat1 | from_917 | to_917 | art_long | art_short | J |
| 22 | 161 | 0.909800809 | cat1 | from_918 | to_918 | art_long | art_short | J |
| 23 | 161 | 0.911705633 | cat1 | from_919 | to_919 | art_long | art_short | J |
| 88 | 161 | 0.907486772 | cat1 | from_920 | to_920 | art_long | art_short | J |
| 113 | 161 | 0.683769301 | cat1 | from_921 | to_921 | art_long | art_short | J |
| 126 | 161 | 0.909611771 | cat1 | from_922 | to_922 | art_long | art_short | J |
| 5 | 162 | 0.685411892 | cat1 | from_923 | to_923 | art_long | art_short | J |
| 12 | 162 | 0.671618199 | cat1 | from_924 | to_924 | art_long | art_short | J |
| 13 | 162 | 0.848207314 | cat1 | from_925 | to_925 | art_long | art_short | J |
| 15 | 162 | 0.965191372 | cat1 | from_926 | to_926 | art_long | art_short | J |
| 16 | 162 | 0.752297685 | cat1 | from_927 | to_927 | art_long | art_short | J |
| 18 | 162 | 0.752297685 | cat1 | from_928 | to_928 | art_long | art_short | J |
| 21 | 162 | 0.705902301 | cat1 | from_929 | to_929 | art_long | art_short | J |
| 39 | 162 | 0.700126698 | cat1 | from_930 | to_930 | art_long | art_short | J |
| 63 | 162 | 0.683752746 | cat1 | from_931 | to_931 | art_long | art_short | J |
| 114 | 162 | 0.689032751 | cat1 | from_932 | to_932 | art_long | art_short | J |
| 116 | 162 | 0.682919344 | cat1 | from_933 | to_933 | art_long | art_short | J |
| 12 | 163 | 0.772628351 | cat1 | from_934 | to_934 | art_long | art_short | J |

| | | | | | | | | |
|---|---|---|---|---|---|---|---|---|
| 13 | 163 | 0.943049548 | cat1 | from_935 | to_935 | art_long | art_short | J |
| 15 | 163 | 0.861298418 | cat1 | from_936 | to_936 | art_long | art_short | J |
| 16 | 163 | 0.684374438 | cat1 | from_937 | to_937 | art_long | art_short | J |
| 18 | 163 | 0.684374438 | cat1 | from_938 | to_938 | art_long | art_short | J |
| 19 | 163 | 0.723770807 | cat1 | from_939 | to_939 | art_long | art_short | J |
| 20 | 163 | 0.68593669 | cat1 | from_940 | to_940 | art_long | art_short | J |
| 39 | 163 | 0.737542264 | cat1 | from_941 | to_941 | art_long | art_short | J |
| 40 | 163 | 0.722828087 | cat1 | from_942 | to_942 | art_long | art_short | J |
| 68 | 163 | 0.680372636 | cat1 | from_943 | to_943 | art_long | art_short | J |
| 79 | 163 | 0.734659135 | cat1 | from_944 | to_944 | art_long | art_short | J |
| 114 | 163 | 0.882205961 | cat1 | from_945 | to_945 | art_long | art_short | J |
| 162 | 163 | 0.942369257 | cat1 | from_946 | to_946 | art_short | art_short | H |
| 2 | 164 | 0.69778056 | cat1 | from_947 | to_947 | art_long | art_short | J |
| 12 | 164 | 0.940518887 | cat1 | from_948 | to_948 | art_long | art_short | J |
| 13 | 164 | 0.672575295 | cat1 | from_949 | to_949 | art_long | art_short | J |
| 15 | 164 | 0.715841415 | cat1 | from_950 | to_950 | art_long | art_short | J |
| 16 | 164 | 0.958338963 | cat1 | from_951 | to_951 | art_long | art_short | J |
| 18 | 164 | 0.958338963 | cat1 | from_952 | to_952 | art_long | art_short | J |
| 19 | 164 | 0.851573308 | cat1 | from_953 | to_953 | art_long | art_short | J |
| 39 | 164 | 0.945740347 | cat1 | from_954 | to_954 | art_long | art_short | J |
| 40 | 164 | 0.851382469 | cat1 | from_955 | to_955 | art_long | art_short | J |
| 53 | 164 | 0.709973904 | cat1 | from_956 | to_956 | art_long | art_short | J |
| 79 | 164 | 0.846415718 | cat1 | from_957 | to_957 | art_long | art_short | J |
| 155 | 164 | 0.738325744 | cat1 | from_958 | to_958 | art_short | art_short | H |
| 162 | 164 | 0.763408514 | cat1 | from_959 | to_959 | art_short | art_short | H |
| 163 | 164 | 0.748343949 | cat1 | from_960 | to_960 | art_short | art_short | H |
| 12 | 165 | 0.787572356 | cat1 | from_961 | to_961 | art_long | art_short | J |
| 13 | 165 | 0.931890519 | cat1 | from_962 | to_962 | art_long | art_short | J |
| 15 | 165 | 0.677102864 | cat1 | from_963 | to_963 | art_long | art_short | J |
| 19 | 165 | 0.77096072 | cat1 | from_964 | to_964 | art_long | art_short | J |
| 20 | 165 | 0.74133149 | cat1 | from_965 | to_965 | art_long | art_short | J |
| 39 | 165 | 0.695066256 | cat1 | from_966 | to_966 | art_long | art_short | J |
| 40 | 165 | 0.768866712 | cat1 | from_967 | to_967 | art_long | art_short | J |
| 68 | 165 | 0.73085183 | cat1 | from_968 | to_968 | art_long | art_short | J |
| 79 | 165 | 0.803882748 | cat1 | from_969 | to_969 | art_long | art_short | J |
| 114 | 165 | 0.968284166 | cat1 | from_970 | to_970 | art_long | art_short | J |
| 156 | 165 | 0.774420938 | cat1 | from_971 | to_971 | art_short | art_short | H |
| 158 | 165 | 0.774298126 | cat1 | from_972 | to_972 | art_short | art_short | H |
| 162 | 165 | 0.794354615 | cat1 | from_973 | to_973 | art_short | art_short | H |
| 163 | 165 | 0.951791596 | cat1 | from_974 | to_974 | art_short | art_short | H |
| 17 | 166 | 0.721445665 | cat1 | from_975 | to_975 | art_long | art_short | J |
| 35 | 166 | 0.961157336 | cat1 | from_976 | to_976 | art_long | art_short | J |
| 36 | 166 | 0.72001661 | cat1 | from_977 | to_977 | art_long | art_short | J |
| 38 | 166 | 0.968528247 | cat1 | from_978 | to_978 | art_long | art_short | J |
| 48 | 166 | 0.961157336 | cat1 | from_979 | to_979 | art_long | art_short | J |
| 50 | 166 | 0.850505945 | cat1 | from_980 | to_980 | art_long | art_short | J |
| 104 | 166 | 0.961157336 | cat1 | from_981 | to_981 | art_long | art_short | J |
| 132 | 166 | 0.961157336 | cat1 | from_982 | to_982 | art_long | art_short | J |
| 133 | 166 | 0.72001661 | cat1 | from_983 | to_983 | art_long | art_short | J |
| 134 | 166 | 0.72001661 | cat1 | from_984 | to_984 | art_long | art_short | J |
| 138 | 166 | 0.750238912 | cat1 | from_985 | to_985 | art_long | art_short | J |
| 143 | 166 | 0.995257201 | cat1 | from_986 | to_986 | art_long | art_short | J |
| 154 | 166 | 0.748893553 | cat1 | from_987 | to_987 | art_short | art_short | H |
| 12 | 167 | 0.999931192 | cat1 | from_988 | to_988 | art_long | art_short | J |
| 13 | 167 | 0.737106999 | cat1 | from_989 | to_989 | art_long | art_short | J |
| 16 | 167 | 0.850670413 | cat1 | from_990 | to_990 | art_long | art_short | J |
| 18 | 167 | 0.850670413 | cat1 | from_991 | to_991 | art_long | art_short | J |
| 19 | 167 | 0.947118025 | cat1 | from_992 | to_992 | art_long | art_short | J |
| 39 | 167 | 0.942319164 | cat1 | from_993 | to_993 | art_long | art_short | J |
| 40 | 167 | 0.945655195 | cat1 | from_994 | to_994 | art_long | art_short | J |
| 79 | 167 | 0.96585092 | cat1 | from_995 | to_995 | art_long | art_short | J |
| 114 | 167 | 0.743703662 | cat1 | from_996 | to_996 | art_long | art_short | J |
| 162 | 167 | 0.673944576 | cat1 | from_997 | to_997 | art_short | art_short | H |
| 163 | 167 | 0.773001454 | cat1 | from_998 | to_998 | art_short | art_short | H |
| 164 | 167 | 0.942599325 | cat1 | from_999 | to_999 | art_short | art_short | H |
| 165 | 167 | 0.786027512 | cat1 | from_1000 | to_1000 | art_short | art_short | H |
| 2 | 168 | 0.767896747 | cat1 | from_1001 | to_1001 | art_long | art_short | J |
| 53 | 168 | 0.781119807 | cat1 | from_1002 | to_1002 | art_long | art_short | J |
| 155 | 168 | 0.811603304 | cat1 | from_1003 | to_1003 | art_short | art_short | H |
| 162 | 168 | 0.704101307 | cat1 | from_1004 | to_1004 | art_short | art_short | H |
| 163 | 168 | 0.71171954 | cat1 | from_1005 | to_1005 | art_short | art_short | H |
| 12 | 169 | 0.771495643 | cat1 | from_1006 | to_1006 | art_long | art_short | J |
| 13 | 169 | 0.944684457 | cat1 | from_1007 | to_1007 | art_long | art_short | J |
| 15 | 169 | 0.864910863 | cat1 | from_1008 | to_1008 | art_long | art_short | J |
| 16 | 169 | 0.687240867 | cat1 | from_1009 | to_1009 | art_long | art_short | J |
| 18 | 169 | 0.687240867 | cat1 | from_1010 | to_1010 | art_long | art_short | J |
| 19 | 169 | 0.724367588 | cat1 | from_1011 | to_1011 | art_long | art_short | J |
| 20 | 169 | 0.686632206 | cat1 | from_1012 | to_1012 | art_long | art_short | J |
| 39 | 169 | 0.739368423 | cat1 | from_1013 | to_1013 | art_long | art_short | J |
| 40 | 169 | 0.723485259 | cat1 | from_1014 | to_1014 | art_long | art_short | J |
| 68 | 169 | 0.681262367 | cat1 | from_1015 | to_1015 | art_long | art_short | J |
| 79 | 169 | 0.73406809 | cat1 | from_1016 | to_1016 | art_long | art_short | J |
| 114 | 169 | 0.881689022 | cat1 | from_1017 | to_1017 | art_long | art_short | J |
| 162 | 169 | 0.943853315 | cat1 | from_1018 | to_1018 | art_short | art_short | H |
| 163 | 169 | 0.999964091 | cat1 | from_1019 | to_1019 | art_short | art_short | H |
| 164 | 169 | 0.748826652 | cat1 | from_1020 | to_1020 | art_short | art_short | H |
| 165 | 169 | 0.950301578 | cat1 | from_1021 | to_1021 | art_short | art_short | H |
| 167 | 169 | 0.771966844 | cat1 | from_1022 | to_1022 | art_short | art_short | H |
| 168 | 169 | 0.712132869 | cat1 | from_1023 | to_1023 | art_short | art_short | H |
| 42 | 170 | 0.925927703 | cat1 | from_1024 | to_1024 | art_long | art_short | J |
| 43 | 170 | 0.910296823 | cat1 | from_1025 | to_1025 | art_long | art_short | J |
| 45 | 170 | 0.925927703 | cat1 | from_1026 | to_1026 | art_long | art_short | J |
| 62 | 170 | 0.930904356 | cat1 | from_1027 | to_1027 | art_long | art_short | J |
| 92 | 170 | 0.930904356 | cat1 | from_1028 | to_1028 | art_long | art_short | J |
| 100 | 170 | 0.925927703 | cat1 | from_1029 | to_1029 | art_long | art_short | J |
| 101 | 170 | 0.925927703 | cat1 | from_1030 | to_1030 | art_long | art_short | J |
| 107 | 170 | 0.925927703 | cat1 | from_1031 | to_1031 | art_long | art_short | J |
| 139 | 170 | 0.998732542 | cat1 | from_1032 | to_1032 | art_long | art_short | J |
| 147 | 170 | 0.925927703 | cat1 | from_1033 | to_1033 | art_long | art_short | J |
| 149 | 170 | 0.925927703 | cat1 | from_1034 | to_1034 | art_long | art_short | J |
| 42 | 171 | 0.929543808 | cat1 | from_1035 | to_1035 | art_long | art_short | J |
| 43 | 171 | 0.912472223 | cat1 | from_1036 | to_1036 | art_long | art_short | J |
| 45 | 171 | 0.929543808 | cat1 | from_1037 | to_1037 | art_long | art_short | J |
| 62 | 171 | 0.934399717 | cat1 | from_1038 | to_1038 | art_long | art_short | J |

| | | | | | | | | |
|---|---|---|---|---|---|---|---|---|
| 92 | 171 | 0.934399717 | cat1 | from_1039 | to_1039 | art_long | art_short | J |
| 100 | 171 | 0.929543808 | cat1 | from_1040 | to_1040 | art_long | art_short | J |
| 101 | 171 | 0.929543808 | cat1 | from_1041 | to_1041 | art_long | art_short | J |
| 107 | 171 | 0.929543808 | cat1 | from_1042 | to_1042 | art_long | art_short | J |
| 139 | 171 | 0.999173332 | cat1 | from_1043 | to_1043 | art_long | art_short | J |
| 147 | 171 | 0.929543808 | cat1 | from_1044 | to_1044 | art_long | art_short | J |
| 149 | 171 | 0.929543808 | cat1 | from_1045 | to_1045 | art_long | art_short | J |
| 170 | 171 | 0.99995306 | cat1 | from_1046 | to_1046 | art_short | art_short | H |
| 22 | 172 | 0.686190251 | cat1 | from_1047 | to_1047 | art_long | art_short | J |
| 27 | 172 | 0.963729205 | cat1 | from_1048 | to_1048 | art_long | art_short | J |
| 64 | 172 | 0.740443769 | cat1 | from_1049 | to_1049 | art_long | art_short | J |
| 91 | 172 | 0.952574782 | cat1 | from_1050 | to_1050 | art_long | art_short | J |
| 117 | 172 | 0.739047995 | cat1 | from_1051 | to_1051 | art_long | art_short | J |
| 161 | 172 | 0.685965402 | cat1 | from_1052 | to_1052 | art_short | art_short | H |
| 14 | 173 | 0.942132612 | cat1 | from_1053 | to_1053 | art_long | art_short | J |
| 24 | 173 | 0.942132612 | cat1 | from_1054 | to_1054 | art_long | art_short | J |
| 25 | 173 | 0.782269655 | cat1 | from_1055 | to_1055 | art_long | art_short | J |
| 28 | 173 | 0.946461107 | cat1 | from_1056 | to_1056 | art_long | art_short | J |
| 31 | 173 | 0.728778342 | cat1 | from_1057 | to_1057 | art_long | art_short | J |
| 33 | 173 | 0.741070326 | cat1 | from_1058 | to_1058 | art_long | art_short | J |
| 59 | 173 | 0.944301259 | cat1 | from_1059 | to_1059 | art_long | art_short | J |
| 73 | 173 | 0.942132612 | cat1 | from_1060 | to_1060 | art_long | art_short | J |
| 74 | 173 | 0.942132612 | cat1 | from_1061 | to_1061 | art_long | art_short | J |
| 76 | 173 | 0.921874199 | cat1 | from_1062 | to_1062 | art_long | art_short | J |
| 78 | 173 | 0.946461107 | cat1 | from_1063 | to_1063 | art_long | art_short | J |
| 90 | 173 | 0.937958184 | cat1 | from_1064 | to_1064 | art_long | art_short | J |
| 97 | 173 | 0.942132612 | cat1 | from_1065 | to_1065 | art_long | art_short | J |
| 110 | 173 | 0.997066805 | cat1 | from_1066 | to_1066 | art_long | art_short | J |
| 112 | 173 | 0.782177509 | cat1 | from_1067 | to_1067 | art_long | art_short | J |
| 122 | 173 | 0.942132612 | cat1 | from_1068 | to_1068 | art_long | art_short | J |
| 14 | 174 | 0.92537369 | cat1 | from_1069 | to_1069 | art_long | art_short | J |
| 24 | 174 | 0.92537369 | cat1 | from_1070 | to_1070 | art_long | art_short | J |
| 25 | 174 | 0.780700278 | cat1 | from_1071 | to_1071 | art_long | art_short | J |
| 28 | 174 | 0.930280021 | cat1 | from_1072 | to_1072 | art_long | art_short | J |
| 31 | 174 | 0.744767167 | cat1 | from_1073 | to_1073 | art_long | art_short | J |
| 33 | 174 | 0.755016907 | cat1 | from_1074 | to_1074 | art_long | art_short | J |
| 59 | 174 | 0.92782876 | cat1 | from_1075 | to_1075 | art_long | art_short | J |
| 73 | 174 | 0.92537369 | cat1 | from_1076 | to_1076 | art_long | art_short | J |
| 74 | 174 | 0.92537369 | cat1 | from_1077 | to_1077 | art_long | art_short | J |
| 76 | 174 | 0.911950821 | cat1 | from_1078 | to_1078 | art_long | art_short | J |
| 78 | 174 | 0.930280021 | cat1 | from_1079 | to_1079 | art_long | art_short | J |
| 90 | 174 | 0.938719301 | cat1 | from_1080 | to_1080 | art_long | art_short | J |
| 97 | 174 | 0.92537369 | cat1 | from_1081 | to_1081 | art_long | art_short | J |
| 110 | 174 | 0.992380267 | cat1 | from_1082 | to_1082 | art_long | art_short | J |
| 112 | 174 | 0.78020082 | cat1 | from_1083 | to_1083 | art_long | art_short | J |
| 122 | 174 | 0.92537369 | cat1 | from_1084 | to_1084 | art_long | art_short | J |
| 173 | 174 | 0.998899662 | cat1 | from_1085 | to_1085 | art_short | art_short | H |
| 3 | 175 | 0.703205196 | cat1 | from_1086 | to_1086 | art_long | art_short | J |
| 29 | 175 | 0.716914727 | cat1 | from_1087 | to_1087 | art_long | art_short | J |
| 71 | 175 | 0.740152156 | cat1 | from_1088 | to_1088 | art_long | art_short | J |
| 82 | 175 | 0.965513443 | cat1 | from_1089 | to_1089 | art_long | art_short | J |
| 89 | 175 | 0.908553182 | cat1 | from_1090 | to_1090 | art_long | art_short | J |
| 98 | 175 | 0.717792795 | cat1 | from_1091 | to_1091 | art_long | art_short | J |
| 121 | 175 | 0.71920157 | cat1 | from_1092 | to_1092 | art_long | art_short | J |
| 150 | 175 | 0.908553182 | cat1 | from_1093 | to_1093 | art_long | art_short | J |
| 44 | 176 | 0.761120107 | cat1 | from_1094 | to_1094 | art_long | art_short | J |
| 46 | 176 | 0.761120107 | cat1 | from_1095 | to_1095 | art_long | art_short | J |
| 52 | 176 | 0.735626954 | cat1 | from_1096 | to_1096 | art_long | art_short | J |
| 60 | 176 | 0.806642941 | cat1 | from_1097 | to_1097 | art_long | art_short | J |
| 61 | 176 | 0.916136877 | cat1 | from_1098 | to_1098 | art_long | art_short | J |
| 99 | 176 | 0.912540734 | cat1 | from_1099 | to_1099 | art_long | art_short | J |
| 102 | 176 | 0.781693552 | cat1 | from_1100 | to_1100 | art_long | art_short | J |
| 105 | 176 | 0.922017596 | cat1 | from_1101 | to_1101 | art_long | art_short | J |
| 106 | 176 | 0.761120107 | cat1 | from_1102 | to_1102 | art_long | art_short | J |
| 128 | 176 | 0.769310949 | cat1 | from_1103 | to_1103 | art_long | art_short | J |
| 135 | 176 | 0.761120107 | cat1 | from_1104 | to_1104 | art_long | art_short | J |
| 148 | 176 | 0.912540734 | cat1 | from_1105 | to_1105 | art_long | art_short | J |
| 17 | 177 | 0.876747825 | cat1 | from_1106 | to_1106 | art_long | art_short | J |
| 35 | 177 | 0.694097688 | cat1 | from_1107 | to_1107 | art_long | art_short | J |
| 36 | 177 | 0.874177775 | cat1 | from_1108 | to_1108 | art_long | art_short | J |
| 38 | 177 | 0.703357623 | cat1 | from_1109 | to_1109 | art_long | art_short | J |
| 48 | 177 | 0.694097688 | cat1 | from_1110 | to_1110 | art_long | art_short | J |
| 91 | 177 | -0.675372386 | cat1 | from_1111 | to_1111 | art_long | art_short | J |
| 104 | 177 | 0.694097688 | cat1 | from_1112 | to_1112 | art_long | art_short | J |
| 132 | 177 | 0.694097688 | cat1 | from_1113 | to_1113 | art_long | art_short | J |
| 133 | 177 | 0.874177775 | cat1 | from_1114 | to_1114 | art_long | art_short | J |
| 134 | 177 | 0.874177775 | cat1 | from_1115 | to_1115 | art_long | art_short | J |
| 138 | 177 | 0.932321999 | cat1 | from_1116 | to_1116 | art_long | art_short | J |
| 143 | 177 | 0.770347813 | cat1 | from_1117 | to_1117 | art_long | art_short | J |
| 154 | 177 | 0.996575986 | cat1 | from_1118 | to_1118 | art_short | art_short | H |
| 166 | 177 | 0.76084653 | cat1 | from_1119 | to_1119 | art_short | art_short | H |
| 42 | 178 | 0.806953795 | cat1 | from_1120 | to_1120 | art_long | art_short | J |
| 43 | 178 | 0.712400686 | cat1 | from_1121 | to_1121 | art_long | art_short | J |
| 45 | 178 | 0.806953795 | cat1 | from_1122 | to_1122 | art_long | art_short | J |
| 52 | 178 | 0.699486888 | cat1 | from_1123 | to_1123 | art_long | art_short | J |
| 62 | 178 | 0.813675253 | cat1 | from_1124 | to_1124 | art_long | art_short | J |
| 92 | 178 | 0.813675253 | cat1 | from_1125 | to_1125 | art_long | art_short | J |
| 93 | 178 | -0.672493919 | cat1 | from_1126 | to_1126 | art_long | art_short | J |
| 100 | 178 | 0.806953795 | cat1 | from_1127 | to_1127 | art_long | art_short | J |
| 101 | 178 | 0.806953795 | cat1 | from_1128 | to_1128 | art_long | art_short | J |
| 107 | 178 | 0.806953795 | cat1 | from_1129 | to_1129 | art_long | art_short | J |
| 139 | 178 | 0.929247626 | cat1 | from_1130 | to_1130 | art_long | art_short | J |
| 147 | 178 | 0.806953795 | cat1 | from_1131 | to_1131 | art_long | art_short | J |
| 149 | 178 | 0.806953795 | cat1 | from_1132 | to_1132 | art_long | art_short | J |
| 170 | 178 | 0.938732019 | cat1 | from_1133 | to_1133 | art_short | art_short | H |
| 171 | 178 | 0.93709122 | cat1 | from_1134 | to_1134 | art_short | art_short | H |
| 2 | 179 | 0.951439821 | cat1 | from_1135 | to_1135 | art_long | art_short | J |
| 12 | 179 | 0.668735615 | cat1 | from_1136 | to_1136 | art_long | art_short | J |
| 16 | 179 | 0.710589957 | cat1 | from_1137 | to_1137 | art_long | art_short | J |
| 18 | 179 | 0.710589957 | cat1 | from_1138 | to_1138 | art_long | art_short | J |
| 39 | 179 | 0.67135529 | cat1 | from_1139 | to_1139 | art_long | art_short | J |
| 53 | 179 | 0.966125011 | cat1 | from_1140 | to_1140 | art_long | art_short | J |
| 155 | 179 | 0.997664135 | cat1 | from_1141 | to_1141 | art_short | art_short | H |
| 164 | 179 | 0.741842178 | cat1 | from_1142 | to_1142 | art_short | art_short | H |

| | | | | | | | | |
|---|---|---|---|---|---|---|---|---|
| 167 | 179 | 0.670177784 | cat1 | from_1143 | to_1143 | art_short | art_short | H |
| 168 | 179 | 0.815225355 | cat1 | from_1144 | to_1144 | art_short | art_short | H |
| 2 | 180 | 0.812064054 | cat1 | from_1145 | to_1145 | art_long | art_short | J |
| 15 | 180 | 0.711774 | cat1 | from_1146 | to_1146 | art_long | art_short | J |
| 53 | 180 | 0.819831239 | cat1 | from_1147 | to_1147 | art_long | art_short | J |
| 155 | 180 | 0.829263899 | cat1 | from_1148 | to_1148 | art_short | art_short | H |
| 162 | 180 | 0.729232602 | cat1 | from_1149 | to_1149 | art_short | art_short | H |
| 168 | 180 | 0.96176494 | cat1 | from_1150 | to_1150 | art_short | art_short | H |
| 179 | 180 | 0.825218101 | cat1 | from_1151 | to_1151 | art_short | art_short | H |
| 3 | 181 | 0.784960594 | cat1 | from_1152 | to_1152 | art_long | art_short | J |
| 29 | 181 | 0.889603668 | cat1 | from_1153 | to_1153 | art_long | art_short | J |
| 41 | 181 | 0.710833204 | cat1 | from_1154 | to_1154 | art_long | art_short | J |
| 66 | 181 | 0.72411782 | cat1 | from_1155 | to_1155 | art_long | art_short | J |
| 67 | 181 | 0.72411782 | cat1 | from_1156 | to_1156 | art_long | art_short | J |
| 71 | 181 | 0.913507829 | cat1 | from_1157 | to_1157 | art_long | art_short | J |
| 82 | 181 | 0.846226754 | cat1 | from_1158 | to_1158 | art_long | art_short | J |
| 89 | 181 | 0.814709013 | cat1 | from_1159 | to_1159 | art_long | art_short | J |
| 96 | 181 | 0.769052835 | cat1 | from_1160 | to_1160 | art_long | art_short | J |
| 124 | 181 | 0.722716325 | cat1 | from_1161 | to_1161 | art_long | art_short | J |
| 150 | 181 | 0.814709013 | cat1 | from_1162 | to_1162 | art_long | art_short | J |
| 175 | 181 | 0.862053479 | cat1 | from_1163 | to_1163 | art_short | art_short | H |
| 22 | 182 | 0.679421891 | cat1 | from_1164 | to_1164 | art_long | art_short | J |
| 49 | 182 | 0.70688716 | cat1 | from_1165 | to_1165 | art_long | art_short | J |
| 56 | 182 | 0.966656398 | cat1 | from_1166 | to_1166 | art_long | art_short | J |
| 57 | 182 | 0.966656398 | cat1 | from_1167 | to_1167 | art_long | art_short | J |
| 70 | 182 | 0.766542811 | cat1 | from_1168 | to_1168 | art_long | art_short | J |
| 75 | 182 | 0.706451354 | cat1 | from_1169 | to_1169 | art_long | art_short | J |
| 85 | 182 | 0.758703081 | cat1 | from_1170 | to_1170 | art_long | art_short | J |
| 123 | 182 | 0.758703081 | cat1 | from_1171 | to_1171 | art_long | art_short | J |
| 146 | 182 | 0.766542811 | cat1 | from_1172 | to_1172 | art_long | art_short | J |
| 161 | 182 | 0.674769276 | cat1 | from_1173 | to_1173 | art_short | art_short | H |
| 44 | 183 | 0.75563409 | cat1 | from_1174 | to_1174 | art_long | art_short | J |
| 46 | 183 | 0.75563409 | cat1 | from_1175 | to_1175 | art_long | art_short | J |
| 52 | 183 | 0.734183749 | cat1 | from_1176 | to_1176 | art_long | art_short | J |
| 60 | 183 | 0.801914866 | cat1 | from_1177 | to_1177 | art_long | art_short | J |
| 61 | 183 | 0.912076992 | cat1 | from_1178 | to_1178 | art_long | art_short | J |
| 99 | 183 | 0.90839972 | cat1 | from_1179 | to_1179 | art_long | art_short | J |
| 102 | 183 | 0.776532431 | cat1 | from_1180 | to_1180 | art_long | art_short | J |
| 105 | 183 | 0.919592981 | cat1 | from_1181 | to_1181 | art_long | art_short | J |
| 106 | 183 | 0.75563409 | cat1 | from_1182 | to_1182 | art_long | art_short | J |
| 128 | 183 | 0.763951162 | cat1 | from_1183 | to_1183 | art_long | art_short | J |
| 135 | 183 | 0.75563409 | cat1 | from_1184 | to_1184 | art_long | art_short | J |
| 148 | 183 | 0.90839972 | cat1 | from_1185 | to_1185 | art_long | art_short | J |
| 176 | 183 | 0.999949866 | cat1 | from_1186 | to_1186 | art_short | art_short | H |
| 42 | 184 | 0.936414302 | cat1 | from_1187 | to_1187 | art_long | art_short | J |
| 43 | 184 | 0.916508342 | cat1 | from_1188 | to_1188 | art_long | art_short | J |
| 45 | 184 | 0.936414302 | cat1 | from_1189 | to_1189 | art_long | art_short | J |
| 62 | 184 | 0.941030937 | cat1 | from_1190 | to_1190 | art_long | art_short | J |
| 92 | 184 | 0.941030937 | cat1 | from_1191 | to_1191 | art_long | art_short | J |
| 100 | 184 | 0.936414302 | cat1 | from_1192 | to_1192 | art_long | art_short | J |
| 101 | 184 | 0.936414302 | cat1 | from_1193 | to_1193 | art_long | art_short | J |
| 107 | 184 | 0.936414302 | cat1 | from_1194 | to_1194 | art_long | art_short | J |
| 139 | 184 | 0.999767387 | cat1 | from_1195 | to_1195 | art_long | art_short | J |
| 147 | 184 | 0.936414302 | cat1 | from_1196 | to_1196 | art_long | art_short | J |
| 149 | 184 | 0.936414302 | cat1 | from_1197 | to_1197 | art_long | art_short | J |
| 170 | 184 | 0.999585777 | cat1 | from_1198 | to_1198 | art_short | art_short | H |
| 171 | 184 | 0.999817705 | cat1 | from_1199 | to_1199 | art_short | art_short | H |
| 178 | 184 | 0.933600338 | cat1 | from_1200 | to_1200 | art_short | art_short | H |
| 3 | 185 | 0.860862618 | cat1 | from_1201 | to_1201 | art_long | art_short | J |
| 82 | 185 | 0.810229465 | cat1 | from_1202 | to_1202 | art_long | art_short | J |
| 89 | 185 | 0.785799735 | cat1 | from_1203 | to_1203 | art_long | art_short | J |
| 150 | 185 | 0.785799735 | cat1 | from_1204 | to_1204 | art_long | art_short | J |
| 175 | 185 | 0.820529294 | cat1 | from_1205 | to_1205 | art_short | art_short | H |
| 5 | 186 | 0.737395976 | cat1 | from_1206 | to_1206 | art_long | art_short | J |
| 13 | 186 | 0.698612944 | cat1 | from_1207 | to_1207 | art_long | art_short | J |
| 63 | 186 | 0.871000756 | cat1 | from_1208 | to_1208 | art_long | art_short | J |
| 114 | 186 | 0.710916863 | cat1 | from_1209 | to_1209 | art_long | art_short | J |
| 116 | 186 | 0.86846809 | cat1 | from_1210 | to_1210 | art_long | art_short | J |
| 156 | 186 | 0.90620835 | cat1 | from_1211 | to_1211 | art_short | art_short | H |
| 158 | 186 | 0.90738337 | cat1 | from_1212 | to_1212 | art_short | art_short | H |
| 163 | 186 | 0.715844585 | cat1 | from_1213 | to_1213 | art_short | art_short | H |
| 165 | 186 | 0.737977755 | cat1 | from_1214 | to_1214 | art_short | art_short | H |
| 169 | 186 | 0.71500714 | cat1 | from_1215 | to_1215 | art_short | art_short | H |
| 2 | 187 | 0.824342985 | cat1 | from_1216 | to_1216 | art_long | art_short | J |
| 15 | 187 | 0.725975355 | cat1 | from_1217 | to_1217 | art_long | art_short | J |
| 53 | 187 | 0.828957547 | cat1 | from_1218 | to_1218 | art_long | art_short | J |
| 155 | 187 | 0.826537018 | cat1 | from_1219 | to_1219 | art_short | art_short | H |
| 162 | 187 | 0.727243669 | cat1 | from_1220 | to_1220 | art_short | art_short | H |
| 168 | 187 | 0.953123926 | cat1 | from_1221 | to_1221 | art_short | art_short | H |
| 179 | 187 | 0.818286888 | cat1 | from_1222 | to_1222 | art_short | art_short | H |
| 180 | 187 | 0.998092626 | cat1 | from_1223 | to_1223 | art_short | art_short | H |
| 4 | 188 | 0.852462005 | cat1 | from_1224 | to_1224 | art_long | art_short | J |
| 6 | 188 | 0.857914561 | cat1 | from_1225 | to_1225 | art_long | art_short | J |
| 7 | 188 | 0.857085143 | cat1 | from_1226 | to_1226 | art_long | art_short | J |
| 11 | 188 | 0.702576973 | cat1 | from_1227 | to_1227 | art_long | art_short | J |
| 137 | 188 | -0.671124707 | cat1 | from_1228 | to_1228 | art_long | art_short | J |
| 141 | 188 | -0.670984569 | cat1 | from_1229 | to_1229 | art_long | art_short | J |
| 159 | 188 | 0.999838384 | cat1 | from_1230 | to_1230 | art_short | art_short | H |
| 160 | 188 | 0.94248458 | cat1 | from_1231 | to_1231 | art_short | art_short | H |
| 29 | 189 | 0.779229354 | cat1 | from_1232 | to_1232 | art_long | art_short | J |
| 66 | 189 | 0.920493218 | cat1 | from_1233 | to_1233 | art_long | art_short | J |
| 67 | 189 | 0.920493218 | cat1 | from_1234 | to_1234 | art_long | art_short | J |
| 71 | 189 | 0.791653171 | cat1 | from_1235 | to_1235 | art_long | art_short | J |
| 82 | 189 | 0.734985129 | cat1 | from_1236 | to_1236 | art_long | art_short | J |
| 89 | 189 | 0.730215356 | cat1 | from_1237 | to_1237 | art_long | art_short | J |
| 150 | 189 | 0.730215356 | cat1 | from_1238 | to_1238 | art_long | art_short | J |
| 175 | 189 | 0.73424377 | cat1 | from_1239 | to_1239 | art_short | art_short | H |
| 181 | 189 | 0.829254233 | cat1 | from_1240 | to_1240 | art_short | art_short | H |
| 32 | 190 | 0.939553041 | cat1 | from_1241 | to_1241 | art_long | art_short | J |
| 86 | 190 | 0.673042578 | cat1 | from_1242 | to_1242 | art_long | art_short | J |
| 94 | 190 | 0.673042578 | cat1 | from_1243 | to_1243 | art_long | art_short | J |
| 32 | 191 | 0.938097914 | cat1 | from_1244 | to_1244 | art_long | art_short | J |
| 86 | 191 | 0.672124146 | cat1 | from_1245 | to_1245 | art_long | art_short | J |
| 94 | 191 | 0.672124146 | cat1 | from_1246 | to_1246 | art_long | art_short | J |

| | | | | | | | | |
|---|---|---|---|---|---|---|---|---|
| 190 | 191 | 0.999967647 | cat1 | from_1247 | to_1247 | art_short | art_short | H |
| 12 | 192 | 0.694663171 | cat1 | from_1248 | to_1248 | art_long | art_short | J |
| 13 | 192 | 0.694377053 | cat1 | from_1249 | to_1249 | art_long | art_short | J |
| 15 | 192 | 0.856560696 | cat1 | from_1250 | to_1250 | art_long | art_short | J |
| 16 | 192 | 0.83499754 | cat1 | from_1251 | to_1251 | art_long | art_short | J |
| 18 | 192 | 0.83499754 | cat1 | from_1252 | to_1252 | art_long | art_short | J |
| 39 | 192 | 0.758451318 | cat1 | from_1253 | to_1253 | art_long | art_short | J |
| 68 | 192 | 0.686877844 | cat1 | from_1254 | to_1254 | art_long | art_short | J |
| 162 | 192 | 0.860174773 | cat1 | from_1255 | to_1255 | art_short | art_short | H |
| 163 | 192 | 0.772306779 | cat1 | from_1256 | to_1256 | art_short | art_short | H |
| 164 | 192 | 0.819755172 | cat1 | from_1257 | to_1257 | art_short | art_short | H |
| 167 | 192 | 0.698233176 | cat1 | from_1258 | to_1258 | art_short | art_short | H |
| 169 | 192 | 0.774396077 | cat1 | from_1259 | to_1259 | art_short | art_short | H |
| 8 | 193 | 0.689188199 | cat1 | from_1260 | to_1260 | art_long | art_short | J |
| 49 | 193 | 0.866231514 | cat1 | from_1261 | to_1261 | art_long | art_short | J |
| 70 | 193 | 0.706990516 | cat1 | from_1262 | to_1262 | art_long | art_short | J |
| 75 | 193 | 0.86295226 | cat1 | from_1263 | to_1263 | art_long | art_short | J |
| 84 | 193 | 0.953892933 | cat1 | from_1264 | to_1264 | art_long | art_short | J |
| 125 | 193 | 0.738567152 | cat1 | from_1265 | to_1265 | art_long | art_short | J |
| 146 | 193 | 0.706990516 | cat1 | from_1266 | to_1266 | art_long | art_short | J |
| 14 | 194 | 0.942132612 | cat1 | from_1267 | to_1267 | art_long | art_short | J |
| 24 | 194 | 0.942132612 | cat1 | from_1268 | to_1268 | art_long | art_short | J |
| 25 | 194 | 0.782269655 | cat1 | from_1269 | to_1269 | art_long | art_short | J |
| 28 | 194 | 0.946461107 | cat1 | from_1270 | to_1270 | art_long | art_short | J |
| 31 | 194 | 0.728778342 | cat1 | from_1271 | to_1271 | art_long | art_short | J |
| 33 | 194 | 0.741070326 | cat1 | from_1272 | to_1272 | art_long | art_short | J |
| 59 | 194 | 0.944301259 | cat1 | from_1273 | to_1273 | art_long | art_short | J |
| 73 | 194 | 0.942132612 | cat1 | from_1274 | to_1274 | art_long | art_short | J |
| 74 | 194 | 0.942132612 | cat1 | from_1275 | to_1275 | art_long | art_short | J |
| 76 | 194 | 0.921874199 | cat1 | from_1276 | to_1276 | art_long | art_short | J |
| 78 | 194 | 0.946461107 | cat1 | from_1277 | to_1277 | art_long | art_short | J |
| 90 | 194 | 0.937958184 | cat1 | from_1278 | to_1278 | art_long | art_short | J |
| 97 | 194 | 0.942132612 | cat1 | from_1279 | to_1279 | art_long | art_short | J |
| 110 | 194 | 0.997066805 | cat1 | from_1280 | to_1280 | art_long | art_short | J |
| 112 | 194 | 0.782177509 | cat1 | from_1281 | to_1281 | art_long | art_short | J |
| 122 | 194 | 0.942132612 | cat1 | from_1282 | to_1282 | art_long | art_short | J |
| 173 | 194 | 1 | cat1 | from_1283 | to_1283 | art_short | art_short | H |
| 174 | 194 | 0.998899662 | cat1 | from_1284 | to_1284 | art_short | art_short | H |
| 30 | 195 | 0.995815108 | cat1 | from_1285 | to_1285 | art_long | art_short | J |
| 47 | 195 | 0.927149255 | cat1 | from_1286 | to_1286 | art_long | art_short | J |
| 54 | 195 | 0.782453596 | cat1 | from_1287 | to_1287 | art_long | art_short | J |
| 55 | 195 | 0.927149255 | cat1 | from_1288 | to_1288 | art_long | art_short | J |
| 58 | 195 | 0.927149255 | cat1 | from_1289 | to_1289 | art_long | art_short | J |
| 69 | 195 | 0.93258456 | cat1 | from_1290 | to_1290 | art_long | art_short | J |
| 72 | 195 | 0.912349372 | cat1 | from_1291 | to_1291 | art_long | art_short | J |
| 77 | 195 | 0.914059521 | cat1 | from_1292 | to_1292 | art_long | art_short | J |
| 80 | 195 | 0.917328121 | cat1 | from_1293 | to_1293 | art_long | art_short | J |
| 81 | 195 | 0.782380418 | cat1 | from_1294 | to_1294 | art_long | art_short | J |
| 83 | 195 | 0.926901977 | cat1 | from_1295 | to_1295 | art_long | art_short | J |
| 87 | 195 | 0.927149255 | cat1 | from_1296 | to_1296 | art_long | art_short | J |
| 95 | 195 | 0.927149255 | cat1 | from_1297 | to_1297 | art_long | art_short | J |
| 109 | 195 | 0.93258456 | cat1 | from_1298 | to_1298 | art_long | art_short | J |
| 111 | 195 | 0.929873101 | cat1 | from_1299 | to_1299 | art_long | art_short | J |
| 119 | 195 | 0.929873101 | cat1 | from_1300 | to_1300 | art_long | art_short | J |
| 120 | 195 | 0.914059521 | cat1 | from_1301 | to_1301 | art_long | art_short | J |
| 140 | 195 | 0.915719937 | cat1 | from_1302 | to_1302 | art_long | art_short | J |
| 13 | 196 | 0.675090107 | cat1 | from_1303 | to_1303 | art_long | art_short | J |
| 15 | 196 | 0.890437109 | cat1 | from_1304 | to_1304 | art_long | art_short | J |
| 16 | 196 | 0.680956242 | cat1 | from_1305 | to_1305 | art_long | art_short | J |
| 18 | 196 | 0.680956242 | cat1 | from_1306 | to_1306 | art_long | art_short | J |
| 162 | 196 | 0.958099146 | cat1 | from_1307 | to_1307 | art_short | art_short | H |
| 163 | 196 | 0.856034787 | cat1 | from_1308 | to_1308 | art_short | art_short | H |
| 164 | 196 | 0.729070495 | cat1 | from_1309 | to_1309 | art_short | art_short | H |
| 165 | 196 | 0.678284253 | cat1 | from_1310 | to_1310 | art_short | art_short | H |
| 169 | 196 | 0.856268418 | cat1 | from_1311 | to_1311 | art_short | art_short | H |
| 180 | 196 | 0.728206308 | cat1 | from_1312 | to_1312 | art_short | art_short | H |
| 187 | 196 | 0.718047798 | cat1 | from_1313 | to_1313 | art_short | art_short | H |
| 192 | 196 | 0.828918222 | cat1 | from_1314 | to_1314 | art_short | art_short | H |
| 12 | 197 | 0.959617585 | cat1 | from_1315 | to_1315 | art_long | art_short | J |
| 16 | 197 | 0.901514063 | cat1 | from_1316 | to_1316 | art_long | art_short | J |
| 18 | 197 | 0.901514063 | cat1 | from_1317 | to_1317 | art_long | art_short | J |
| 19 | 197 | 0.839691488 | cat1 | from_1318 | to_1318 | art_long | art_short | J |
| 39 | 197 | 0.909578305 | cat1 | from_1319 | to_1319 | art_long | art_short | J |
| 40 | 197 | 0.838373762 | cat1 | from_1320 | to_1320 | art_long | art_short | J |
| 79 | 197 | 0.856706926 | cat1 | from_1321 | to_1321 | art_long | art_short | J |
| 155 | 197 | 0.718603087 | cat1 | from_1322 | to_1322 | art_short | art_short | H |
| 162 | 197 | 0.742524485 | cat1 | from_1323 | to_1323 | art_short | art_short | H |
| 163 | 197 | 0.757087218 | cat1 | from_1324 | to_1324 | art_short | art_short | H |
| 164 | 197 | 0.987563845 | cat1 | from_1325 | to_1325 | art_short | art_short | H |
| 165 | 197 | 0.695484801 | cat1 | from_1326 | to_1326 | art_short | art_short | H |
| 167 | 197 | 0.959861746 | cat1 | from_1327 | to_1327 | art_short | art_short | H |
| 169 | 197 | 0.756240798 | cat1 | from_1328 | to_1328 | art_short | art_short | H |
| 179 | 197 | 0.732807129 | cat1 | from_1329 | to_1329 | art_short | art_short | H |
| 192 | 197 | 0.782372977 | cat1 | from_1330 | to_1330 | art_short | art_short | H |
| 196 | 197 | 0.729768354 | cat1 | from_1331 | to_1331 | art_short | art_short | H |
| 8 | 198 | 0.748845425 | cat1 | from_1332 | to_1332 | art_long | art_short | J |
| 9 | 198 | 0.688657824 | cat1 | from_1333 | to_1333 | art_long | art_short | J |
| 10 | 198 | 0.692257797 | cat1 | from_1334 | to_1334 | art_long | art_short | J |
| 65 | 198 | 0.726006037 | cat1 | from_1335 | to_1335 | art_long | art_short | J |
| 84 | 198 | 0.677059622 | cat1 | from_1336 | to_1336 | art_long | art_short | J |
| 113 | 198 | 0.678969567 | cat1 | from_1337 | to_1337 | art_long | art_short | J |
| 125 | 198 | 0.885173384 | cat1 | from_1338 | to_1338 | art_long | art_short | J |
| 156 | 198 | 0.758377941 | cat1 | from_1339 | to_1339 | art_short | art_short | H |
| 157 | 198 | 0.777593828 | cat1 | from_1340 | to_1340 | art_short | art_short | H |
| 158 | 198 | 0.756836404 | cat1 | from_1341 | to_1341 | art_short | art_short | H |
| 193 | 198 | 0.76185265 | cat1 | from_1342 | to_1342 | art_short | art_short | H |
| 49 | 199 | 0.946412994 | cat1 | from_1343 | to_1343 | art_long | art_short | J |
| 56 | 199 | 0.734452987 | cat1 | from_1344 | to_1344 | art_long | art_short | J |
| 57 | 199 | 0.734452987 | cat1 | from_1345 | to_1345 | art_long | art_short | J |
| 70 | 199 | 0.950289027 | cat1 | from_1346 | to_1346 | art_long | art_short | J |
| 75 | 199 | 0.945102055 | cat1 | from_1347 | to_1347 | art_long | art_short | J |
| 84 | 199 | 0.763678009 | cat1 | from_1348 | to_1348 | art_long | art_short | J |
| 85 | 199 | 0.874461252 | cat1 | from_1349 | to_1349 | art_long | art_short | J |
| 123 | 199 | 0.874461252 | cat1 | from_1350 | to_1350 | art_long | art_short | J |

| | | | | | | | | |
|---|---|---|---|---|---|---|---|---|
| 146 | 199 | 0.950289027 | cat1 | from_1351 | to_1351 | art_long | art_short | J |
| 182 | 199 | 0.781660426 | cat1 | from_1352 | to_1352 | art_short | art_short | H |
| 193 | 199 | 0.838484473 | cat1 | from_1353 | to_1353 | art_short | art_short | H |
| 30 | 200 | 0.996656683 | cat1 | from_1354 | to_1354 | art_long | art_short | J |
| 47 | 200 | 0.930748807 | cat1 | from_1355 | to_1355 | art_long | art_short | J |
| 54 | 200 | 0.782645518 | cat1 | from_1356 | to_1356 | art_long | art_short | J |
| 55 | 200 | 0.930748807 | cat1 | from_1357 | to_1357 | art_long | art_short | J |
| 58 | 200 | 0.930748807 | cat1 | from_1358 | to_1358 | art_long | art_short | J |
| 69 | 200 | 0.936050246 | cat1 | from_1359 | to_1359 | art_long | art_short | J |
| 72 | 200 | 0.914260076 | cat1 | from_1360 | to_1360 | art_long | art_short | J |
| 77 | 200 | 0.915887615 | cat1 | from_1361 | to_1361 | art_long | art_short | J |
| 80 | 200 | 0.918987663 | cat1 | from_1362 | to_1362 | art_long | art_short | J |
| 81 | 200 | 0.781074296 | cat1 | from_1363 | to_1363 | art_long | art_short | J |
| 83 | 200 | 0.927936548 | cat1 | from_1364 | to_1364 | art_long | art_short | J |
| 87 | 200 | 0.930748807 | cat1 | from_1365 | to_1365 | art_long | art_short | J |
| 95 | 200 | 0.930748807 | cat1 | from_1366 | to_1366 | art_long | art_short | J |
| 109 | 200 | 0.936050246 | cat1 | from_1367 | to_1367 | art_long | art_short | J |
| 111 | 200 | 0.933406288 | cat1 | from_1368 | to_1368 | art_long | art_short | J |
| 119 | 200 | 0.933406288 | cat1 | from_1369 | to_1369 | art_long | art_short | J |
| 120 | 200 | 0.915887615 | cat1 | from_1370 | to_1370 | art_long | art_short | J |
| 140 | 200 | 0.917464311 | cat1 | from_1371 | to_1371 | art_long | art_short | J |
| 195 | 200 | 0.999952724 | cat1 | from_1372 | to_1372 | art_short | art_short | H |
| 8 | 201 | 0.934086491 | cat1 | from_1373 | to_1373 | art_long | art_short | J |
| 9 | 201 | 0.931114463 | cat1 | from_1374 | to_1374 | art_long | art_short | J |
| 10 | 201 | 0.933974162 | cat1 | from_1375 | to_1375 | art_long | art_short | J |
| 23 | 201 | 0.834656792 | cat1 | from_1376 | to_1376 | art_long | art_short | J |
| 88 | 201 | 0.826936534 | cat1 | from_1377 | to_1377 | art_long | art_short | J |
| 125 | 201 | 0.750638741 | cat1 | from_1378 | to_1378 | art_long | art_short | J |
| 126 | 201 | 0.830799776 | cat1 | from_1379 | to_1379 | art_long | art_short | J |
| 145 | 201 | 0.66661947 | cat1 | from_1380 | to_1380 | art_long | art_short | J |
| 156 | 201 | 0.700464996 | cat1 | from_1381 | to_1381 | art_short | art_short | H |
| 157 | 201 | 0.725215792 | cat1 | from_1382 | to_1382 | art_short | art_short | H |
| 158 | 201 | 0.699363848 | cat1 | from_1383 | to_1383 | art_short | art_short | H |
| 161 | 201 | 0.809097496 | cat1 | from_1384 | to_1384 | art_short | art_short | H |
| 193 | 201 | 0.672372617 | cat1 | from_1385 | to_1385 | art_short | art_short | H |
| 198 | 201 | 0.807982367 | cat1 | from_1386 | to_1386 | art_short | art_short | H |
| 51 | 202 | 0.846259279 | cat1 | from_1387 | to_1387 | art_long | art_short | J |
| 44 | 203 | 0.761120107 | cat1 | from_1388 | to_1388 | art_long | art_short | J |
| 46 | 203 | 0.761120107 | cat1 | from_1389 | to_1389 | art_long | art_short | J |
| 52 | 203 | 0.735626954 | cat1 | from_1390 | to_1390 | art_long | art_short | J |
| 60 | 203 | 0.806642941 | cat1 | from_1391 | to_1391 | art_long | art_short | J |
| 61 | 203 | 0.916136877 | cat1 | from_1392 | to_1392 | art_long | art_short | J |
| 99 | 203 | 0.912540734 | cat1 | from_1393 | to_1393 | art_long | art_short | J |
| 102 | 203 | 0.781693552 | cat1 | from_1394 | to_1394 | art_long | art_short | J |
| 105 | 203 | 0.922017596 | cat1 | from_1395 | to_1395 | art_long | art_short | J |
| 106 | 203 | 0.761120107 | cat1 | from_1396 | to_1396 | art_long | art_short | J |
| 128 | 203 | 0.769310949 | cat1 | from_1397 | to_1397 | art_long | art_short | J |
| 135 | 203 | 0.761120107 | cat1 | from_1398 | to_1398 | art_long | art_short | J |
| 148 | 203 | 0.912540734 | cat1 | from_1399 | to_1399 | art_long | art_short | J |
| 176 | 203 | 1 | cat1 | from_1400 | to_1400 | art_short | art_short | H |
| 183 | 203 | 0.999949866 | cat1 | from_1401 | to_1401 | art_short | art_short | H |
| 44 | 204 | 0.919286768 | cat1 | from_1402 | to_1402 | art_long | art_short | J |
| 46 | 204 | 0.919286768 | cat1 | from_1403 | to_1403 | art_long | art_short | J |
| 52 | 204 | 0.939053464 | cat1 | from_1404 | to_1404 | art_long | art_short | J |
| 60 | 204 | 0.950595912 | cat1 | from_1405 | to_1405 | art_long | art_short | J |
| 61 | 204 | 0.907498976 | cat1 | from_1406 | to_1406 | art_long | art_short | J |
| 99 | 204 | 0.905614218 | cat1 | from_1407 | to_1407 | art_long | art_short | J |
| 102 | 204 | 0.933821174 | cat1 | from_1408 | to_1408 | art_long | art_short | J |
| 105 | 204 | 0.773077887 | cat1 | from_1409 | to_1409 | art_long | art_short | J |
| 106 | 204 | 0.919286768 | cat1 | from_1410 | to_1410 | art_long | art_short | J |
| 128 | 204 | 0.925141333 | cat1 | from_1411 | to_1411 | art_long | art_short | J |
| 135 | 204 | 0.919286768 | cat1 | from_1412 | to_1412 | art_long | art_short | J |
| 148 | 204 | 0.905614218 | cat1 | from_1413 | to_1413 | art_long | art_short | J |
| 176 | 204 | 0.914852767 | cat1 | from_1414 | to_1414 | art_short | art_short | H |
| 183 | 204 | 0.913074144 | cat1 | from_1415 | to_1415 | art_short | art_short | H |
| 203 | 204 | 0.914852767 | cat1 | from_1416 | to_1416 | art_short | art_short | H |
| 4 | 205 | 0.731345158 | cat1 | from_1417 | to_1417 | art_long | art_short | J |
| 7 | 205 | 0.734251845 | cat1 | from_1418 | to_1418 | art_long | art_short | J |
| 11 | 205 | 0.963758359 | cat1 | from_1419 | to_1419 | art_long | art_short | J |
| 22 | 205 | 0.686798846 | cat1 | from_1420 | to_1420 | art_long | art_short | J |
| 129 | 205 | -0.679105211 | cat1 | from_1421 | to_1421 | art_long | art_short | J |
| 142 | 205 | -0.678350345 | cat1 | from_1422 | to_1422 | art_long | art_short | J |
| 159 | 205 | 0.759051628 | cat1 | from_1423 | to_1423 | art_short | art_short | H |
| 160 | 205 | 0.780149949 | cat1 | from_1424 | to_1424 | art_short | art_short | H |
| 161 | 205 | 0.690163656 | cat1 | from_1425 | to_1425 | art_short | art_short | H |
| 188 | 205 | 0.757670195 | cat1 | from_1426 | to_1426 | art_short | art_short | H |
| 30 | 206 | 0.891217204 | cat1 | from_1427 | to_1427 | art_long | art_short | J |
| 31 | 206 | 0.70269235 | cat1 | from_1428 | to_1428 | art_long | art_short | J |
| 33 | 206 | 0.698572854 | cat1 | from_1429 | to_1429 | art_long | art_short | J |
| 47 | 206 | 0.766895457 | cat1 | from_1430 | to_1430 | art_long | art_short | J |
| 54 | 206 | 0.92255135 | cat1 | from_1431 | to_1431 | art_long | art_short | J |
| 55 | 206 | 0.766895457 | cat1 | from_1432 | to_1432 | art_long | art_short | J |
| 58 | 206 | 0.766895457 | cat1 | from_1433 | to_1433 | art_long | art_short | J |
| 69 | 206 | 0.774851019 | cat1 | from_1434 | to_1434 | art_long | art_short | J |
| 72 | 206 | 0.912492799 | cat1 | from_1435 | to_1435 | art_long | art_short | J |
| 77 | 206 | 0.91600261 | cat1 | from_1436 | to_1436 | art_long | art_short | J |
| 80 | 206 | 0.922933433 | cat1 | from_1437 | to_1437 | art_long | art_short | J |
| 81 | 206 | 0.95140419 | cat1 | from_1438 | to_1438 | art_long | art_short | J |
| 83 | 206 | 0.945974709 | cat1 | from_1439 | to_1439 | art_long | art_short | J |
| 87 | 206 | 0.766895457 | cat1 | from_1440 | to_1440 | art_long | art_short | J |
| 95 | 206 | 0.766895457 | cat1 | from_1441 | to_1441 | art_long | art_short | J |
| 109 | 206 | 0.774851019 | cat1 | from_1442 | to_1442 | art_long | art_short | J |
| 111 | 206 | 0.770866195 | cat1 | from_1443 | to_1443 | art_long | art_short | J |
| 119 | 206 | 0.770866195 | cat1 | from_1444 | to_1444 | art_long | art_short | J |
| 120 | 206 | 0.91600261 | cat1 | from_1445 | to_1445 | art_long | art_short | J |
| 140 | 206 | 0.919483674 | cat1 | from_1446 | to_1446 | art_long | art_short | J |
| 195 | 206 | 0.914885421 | cat1 | from_1447 | to_1447 | art_short | art_short | H |
| 200 | 206 | 0.91273131 | cat1 | from_1448 | to_1448 | art_short | art_short | H |
| 22 | 207 | 0.686542199 | cat1 | from_1449 | to_1449 | art_long | art_short | J |
| 34 | 207 | 0.942562468 | cat1 | from_1450 | to_1450 | art_long | art_short | J |
| 65 | 207 | 0.932554792 | cat1 | from_1451 | to_1451 | art_long | art_short | J |
| 113 | 207 | 0.997834187 | cat1 | from_1452 | to_1452 | art_long | art_short | J |
| 127 | 207 | 0.944068296 | cat1 | from_1453 | to_1453 | art_long | art_short | J |
| 160 | 207 | 0.692658698 | cat1 | from_1454 | to_1454 | art_short | art_short | H |

| | | | | | | | | |
|---|---|---|---|---|---|---|---|---|
| 161 | 207 | 0.69722495 | cat1 | from_1455 | to_1455 | art_short | art_short | H |
| 198 | 207 | 0.712807407 | cat1 | from_1456 | to_1456 | art_short | art_short | H |
| 3 | 208 | 0.757069314 | cat1 | from_1457 | to_1457 | art_long | art_short | J |
| 29 | 208 | 0.976660434 | cat1 | from_1458 | to_1458 | art_long | art_short | J |
| 41 | 208 | 0.797151371 | cat1 | from_1459 | to_1459 | art_long | art_short | J |
| 66 | 208 | 0.810683824 | cat1 | from_1460 | to_1460 | art_long | art_short | J |
| 67 | 208 | 0.810683824 | cat1 | from_1461 | to_1461 | art_long | art_short | J |
| 71 | 208 | 0.976146794 | cat1 | from_1462 | to_1462 | art_long | art_short | J |
| 82 | 208 | 0.823796616 | cat1 | from_1463 | to_1463 | art_long | art_short | J |
| 89 | 208 | 0.854857188 | cat1 | from_1464 | to_1464 | art_long | art_short | J |
| 96 | 208 | 0.800935605 | cat1 | from_1465 | to_1465 | art_long | art_short | J |
| 124 | 208 | 0.808585158 | cat1 | from_1466 | to_1466 | art_long | art_short | J |
| 150 | 208 | 0.854857188 | cat1 | from_1467 | to_1467 | art_long | art_short | J |
| 175 | 208 | 0.803707485 | cat1 | from_1468 | to_1468 | art_short | art_short | H |
| 181 | 208 | 0.966218489 | cat1 | from_1469 | to_1469 | art_short | art_short | H |
| 189 | 208 | 0.824906268 | cat1 | from_1470 | to_1470 | art_short | art_short | H |
| 3 | 209 | 0.697185198 | cat1 | from_1471 | to_1471 | art_long | art_short | J |
| 29 | 209 | 0.827439305 | cat1 | from_1472 | to_1472 | art_long | art_short | J |
| 41 | 209 | 0.696711476 | cat1 | from_1473 | to_1473 | art_long | art_short | J |
| 66 | 209 | 0.705636519 | cat1 | from_1474 | to_1474 | art_long | art_short | J |
| 67 | 209 | 0.705636519 | cat1 | from_1475 | to_1475 | art_long | art_short | J |
| 71 | 209 | 0.828792701 | cat1 | from_1476 | to_1476 | art_long | art_short | J |
| 82 | 209 | 0.96944271 | cat1 | from_1477 | to_1477 | art_long | art_short | J |
| 89 | 209 | 0.975329425 | cat1 | from_1478 | to_1478 | art_long | art_short | J |
| 96 | 209 | 0.716711804 | cat1 | from_1479 | to_1479 | art_long | art_short | J |
| 98 | 209 | 0.803199191 | cat1 | from_1480 | to_1480 | art_long | art_short | J |
| 121 | 209 | 0.804088254 | cat1 | from_1481 | to_1481 | art_long | art_short | J |
| 124 | 209 | 0.704898483 | cat1 | from_1482 | to_1482 | art_long | art_short | J |
| 150 | 209 | 0.975329425 | cat1 | from_1483 | to_1483 | art_long | art_short | J |
| 175 | 209 | 0.965178168 | cat1 | from_1484 | to_1484 | art_short | art_short | H |
| 181 | 209 | 0.852317095 | cat1 | from_1485 | to_1485 | art_short | art_short | H |
| 185 | 209 | 0.817746287 | cat1 | from_1486 | to_1486 | art_short | art_short | H |
| 189 | 209 | 0.750871418 | cat1 | from_1487 | to_1487 | art_short | art_short | H |
| 208 | 209 | 0.862181664 | cat1 | from_1488 | to_1488 | art_short | art_short | H |
| 3 | 210 | 0.697185198 | cat1 | from_1489 | to_1489 | art_long | art_short | J |
| 29 | 210 | 0.827439305 | cat1 | from_1490 | to_1490 | art_long | art_short | J |
| 41 | 210 | 0.696711476 | cat1 | from_1491 | to_1491 | art_long | art_short | J |
| 66 | 210 | 0.705636519 | cat1 | from_1492 | to_1492 | art_long | art_short | J |
| 67 | 210 | 0.705636519 | cat1 | from_1493 | to_1493 | art_long | art_short | J |
| 71 | 210 | 0.828792701 | cat1 | from_1494 | to_1494 | art_long | art_short | J |
| 82 | 210 | 0.96944271 | cat1 | from_1495 | to_1495 | art_long | art_short | J |
| 89 | 210 | 0.975329425 | cat1 | from_1496 | to_1496 | art_long | art_short | J |
| 96 | 210 | 0.716711804 | cat1 | from_1497 | to_1497 | art_long | art_short | J |
| 98 | 210 | 0.803199191 | cat1 | from_1498 | to_1498 | art_long | art_short | J |
| 121 | 210 | 0.804088254 | cat1 | from_1499 | to_1499 | art_long | art_short | J |
| 124 | 210 | 0.704898483 | cat1 | from_1500 | to_1500 | art_long | art_short | J |
| 150 | 210 | 0.975329425 | cat1 | from_1501 | to_1501 | art_long | art_short | J |
| 175 | 210 | 0.965178168 | cat1 | from_1502 | to_1502 | art_short | art_short | H |
| 181 | 210 | 0.852317095 | cat1 | from_1503 | to_1503 | art_short | art_short | H |
| 185 | 210 | 0.817746287 | cat1 | from_1504 | to_1504 | art_short | art_short | H |
| 189 | 210 | 0.750871418 | cat1 | from_1505 | to_1505 | art_short | art_short | H |
| 208 | 210 | 0.862181664 | cat1 | from_1506 | to_1506 | art_short | art_short | H |
| 209 | 210 | 1 | cat1 | from_1507 | to_1507 | art_short | art_short | H |
| 30 | 211 | 0.996656683 | cat1 | from_1508 | to_1508 | art_long | art_short | J |
| 47 | 211 | 0.930748807 | cat1 | from_1509 | to_1509 | art_long | art_short | J |
| 54 | 211 | 0.782645518 | cat1 | from_1510 | to_1510 | art_long | art_short | J |
| 55 | 211 | 0.930748807 | cat1 | from_1511 | to_1511 | art_long | art_short | J |
| 58 | 211 | 0.930748807 | cat1 | from_1512 | to_1512 | art_long | art_short | J |
| 69 | 211 | 0.936050246 | cat1 | from_1513 | to_1513 | art_long | art_short | J |
| 72 | 211 | 0.914260076 | cat1 | from_1514 | to_1514 | art_long | art_short | J |
| 77 | 211 | 0.915887615 | cat1 | from_1515 | to_1515 | art_long | art_short | J |
| 80 | 211 | 0.918987663 | cat1 | from_1516 | to_1516 | art_long | art_short | J |
| 81 | 211 | 0.781074296 | cat1 | from_1517 | to_1517 | art_long | art_short | J |
| 83 | 211 | 0.927936548 | cat1 | from_1518 | to_1518 | art_long | art_short | J |
| 87 | 211 | 0.930748807 | cat1 | from_1519 | to_1519 | art_long | art_short | J |
| 95 | 211 | 0.930748807 | cat1 | from_1520 | to_1520 | art_long | art_short | J |
| 109 | 211 | 0.936050246 | cat1 | from_1521 | to_1521 | art_long | art_short | J |
| 111 | 211 | 0.933406288 | cat1 | from_1522 | to_1522 | art_long | art_short | J |
| 119 | 211 | 0.933406288 | cat1 | from_1523 | to_1523 | art_long | art_short | J |
| 120 | 211 | 0.915887615 | cat1 | from_1524 | to_1524 | art_long | art_short | J |
| 140 | 211 | 0.917464311 | cat1 | from_1525 | to_1525 | art_long | art_short | J |
| 195 | 211 | 0.999952724 | cat1 | from_1526 | to_1526 | art_short | art_short | H |
| 200 | 211 | 1 | cat1 | from_1527 | to_1527 | art_short | art_short | H |
| 206 | 211 | 0.91273131 | cat1 | from_1528 | to_1528 | art_short | art_short | H |
| 34 | 212 | 0.697809941 | cat1 | from_1529 | to_1529 | art_long | art_short | J |
| 49 | 212 | 0.855662273 | cat1 | from_1530 | to_1530 | art_long | art_short | J |
| 56 | 212 | 0.796342184 | cat1 | from_1531 | to_1531 | art_long | art_short | J |
| 57 | 212 | 0.796342184 | cat1 | from_1532 | to_1532 | art_long | art_short | J |
| 70 | 212 | 0.953640024 | cat1 | from_1533 | to_1533 | art_long | art_short | J |
| 75 | 212 | 0.855683793 | cat1 | from_1534 | to_1534 | art_long | art_short | J |
| 85 | 212 | 0.966522189 | cat1 | from_1535 | to_1535 | art_long | art_short | J |
| 113 | 212 | 0.681168276 | cat1 | from_1536 | to_1536 | art_long | art_short | J |
| 123 | 212 | 0.966522189 | cat1 | from_1537 | to_1537 | art_long | art_short | J |
| 127 | 212 | 0.675196095 | cat1 | from_1538 | to_1538 | art_long | art_short | J |
| 146 | 212 | 0.953640024 | cat1 | from_1539 | to_1539 | art_long | art_short | J |
| 182 | 212 | 0.800742871 | cat1 | from_1540 | to_1540 | art_short | art_short | H |
| 199 | 212 | 0.948677837 | cat1 | from_1541 | to_1541 | art_short | art_short | H |
| 207 | 212 | 0.675073908 | cat1 | from_1542 | to_1542 | art_short | art_short | H |
| 27 | 213 | -0.709691787 | cat1 | from_1543 | to_1543 | art_long | art_short | J |
| 50 | 213 | 0.757958152 | cat1 | from_1544 | to_1544 | art_long | art_short | J |
| 138 | 213 | 0.713769529 | cat1 | from_1545 | to_1545 | art_long | art_short | J |
| 154 | 213 | 0.837247219 | cat1 | from_1546 | to_1546 | art_short | art_short | H |
| 172 | 213 | -0.669679274 | cat1 | from_1547 | to_1547 | art_short | art_short | H |
| 177 | 213 | 0.826989195 | cat1 | from_1548 | to_1548 | art_short | art_short | H |
| 35 | 214 | 0.859809388 | cat1 | from_1549 | to_1549 | art_long | art_short | J |
| 38 | 214 | 0.871511344 | cat1 | from_1550 | to_1550 | art_long | art_short | J |
| 48 | 214 | 0.859809388 | cat1 | from_1551 | to_1551 | art_long | art_short | J |
| 50 | 214 | 0.952109858 | cat1 | from_1552 | to_1552 | art_long | art_short | J |
| 64 | 214 | -0.670162364 | cat1 | from_1553 | to_1553 | art_long | art_short | J |
| 104 | 214 | 0.859809388 | cat1 | from_1554 | to_1554 | art_long | art_short | J |
| 117 | 214 | -0.669645382 | cat1 | from_1555 | to_1555 | art_long | art_short | J |
| 132 | 214 | 0.859809388 | cat1 | from_1556 | to_1556 | art_long | art_short | J |
| 143 | 214 | 0.957296277 | cat1 | from_1557 | to_1557 | art_long | art_short | J |
| 154 | 214 | 0.70456838 | cat1 | from_1558 | to_1558 | art_short | art_short | H |

| | | | | | | | | |
|---|---|---|---|---|---|---|---|---|
| 166 | 214 | 0.944767317 | cat1 | from_1559 | to_1559 | art_short | art_short | H |
| 177 | 214 | 0.701073698 | cat1 | from_1560 | to_1560 | art_short | art_short | H |
| 213 | 214 | 0.758610788 | cat1 | from_1561 | to_1561 | art_short | art_short | H |
| 82 | 215 | 0.789827591 | cat1 | from_1562 | to_1562 | art_long | art_short | J |
| 89 | 215 | 0.767096097 | cat1 | from_1563 | to_1563 | art_long | art_short | J |
| 150 | 215 | 0.767096097 | cat1 | from_1564 | to_1564 | art_long | art_short | J |
| 175 | 215 | 0.801038072 | cat1 | from_1565 | to_1565 | art_short | art_short | H |
| 209 | 215 | 0.799269688 | cat1 | from_1566 | to_1566 | art_short | art_short | H |
| 210 | 215 | 0.799269688 | cat1 | from_1567 | to_1567 | art_short | art_short | H |
| 2 | 216 | 0.6818802 | cat1 | from_1568 | to_1568 | art_long | art_short | J |
| 12 | 216 | 0.839143784 | cat1 | from_1569 | to_1569 | art_long | art_short | J |
| 15 | 216 | 0.70052665 | cat1 | from_1570 | to_1570 | art_long | art_short | J |
| 16 | 216 | 0.905873079 | cat1 | from_1571 | to_1571 | art_long | art_short | J |
| 18 | 216 | 0.905873079 | cat1 | from_1572 | to_1572 | art_long | art_short | J |
| 19 | 216 | 0.675978827 | cat1 | from_1573 | to_1573 | art_long | art_short | J |
| 39 | 216 | 0.823349593 | cat1 | from_1574 | to_1574 | art_long | art_short | J |
| 40 | 216 | 0.675373062 | cat1 | from_1575 | to_1575 | art_long | art_short | J |
| 53 | 216 | 0.696599769 | cat1 | from_1576 | to_1576 | art_long | art_short | J |
| 79 | 216 | 0.68077324 | cat1 | from_1577 | to_1577 | art_long | art_short | J |
| 155 | 216 | 0.734591939 | cat1 | from_1578 | to_1578 | art_short | art_short | H |
| 162 | 216 | 0.761291237 | cat1 | from_1579 | to_1579 | art_short | art_short | H |
| 163 | 216 | 0.680144891 | cat1 | from_1580 | to_1580 | art_short | art_short | H |
| 164 | 216 | 0.961112292 | cat1 | from_1581 | to_1581 | art_short | art_short | H |
| 167 | 216 | 0.840306041 | cat1 | from_1582 | to_1582 | art_short | art_short | H |
| 169 | 216 | 0.680100752 | cat1 | from_1583 | to_1583 | art_short | art_short | H |
| 179 | 216 | 0.741580991 | cat1 | from_1584 | to_1584 | art_short | art_short | H |
| 192 | 216 | 0.820124237 | cat1 | from_1585 | to_1585 | art_short | art_short | H |
| 196 | 216 | 0.802647794 | cat1 | from_1586 | to_1586 | art_short | art_short | H |
| 197 | 216 | 0.957525196 | cat1 | from_1587 | to_1587 | art_short | art_short | H |
| 1 | 217 | 0.784691196 | cat1 | from_1588 | to_1588 | art_long | art_short | J |
| 145 | 217 | 0.730238663 | cat1 | from_1589 | to_1589 | art_long | art_short | J |
| 30 | 218 | 0.996656683 | cat1 | from_1590 | to_1590 | art_long | art_short | J |
| 47 | 218 | 0.930748807 | cat1 | from_1591 | to_1591 | art_long | art_short | J |
| 54 | 218 | 0.782645518 | cat1 | from_1592 | to_1592 | art_long | art_short | J |
| 55 | 218 | 0.930748807 | cat1 | from_1593 | to_1593 | art_long | art_short | J |
| 58 | 218 | 0.930748807 | cat1 | from_1594 | to_1594 | art_long | art_short | J |
| 69 | 218 | 0.936050246 | cat1 | from_1595 | to_1595 | art_long | art_short | J |
| 72 | 218 | 0.914260076 | cat1 | from_1596 | to_1596 | art_long | art_short | J |
| 77 | 218 | 0.915887615 | cat1 | from_1597 | to_1597 | art_long | art_short | J |
| 80 | 218 | 0.918987663 | cat1 | from_1598 | to_1598 | art_long | art_short | J |
| 81 | 218 | 0.781074296 | cat1 | from_1599 | to_1599 | art_long | art_short | J |
| 83 | 218 | 0.927936548 | cat1 | from_1600 | to_1600 | art_long | art_short | J |
| 87 | 218 | 0.930748807 | cat1 | from_1601 | to_1601 | art_long | art_short | J |
| 95 | 218 | 0.930748807 | cat1 | from_1602 | to_1602 | art_long | art_short | J |
| 109 | 218 | 0.936050246 | cat1 | from_1603 | to_1603 | art_long | art_short | J |
| 111 | 218 | 0.933406288 | cat1 | from_1604 | to_1604 | art_long | art_short | J |
| 119 | 218 | 0.933406288 | cat1 | from_1605 | to_1605 | art_long | art_short | J |
| 120 | 218 | 0.915887615 | cat1 | from_1606 | to_1606 | art_long | art_short | J |
| 140 | 218 | 0.917464311 | cat1 | from_1607 | to_1607 | art_long | art_short | J |
| 195 | 218 | 0.999952724 | cat1 | from_1608 | to_1608 | art_short | art_short | H |
| 200 | 218 | 1 | cat1 | from_1609 | to_1609 | art_short | art_short | H |
| 206 | 218 | 0.91273131 | cat1 | from_1610 | to_1610 | art_short | art_short | H |
| 211 | 218 | 1 | cat1 | from_1611 | to_1611 | art_short | art_short | H |
| 30 | 219 | 0.720210313 | cat1 | from_1612 | to_1612 | art_long | art_untreat | E |
| 81 | 219 | 0.692304759 | cat1 | from_1613 | to_1613 | art_long | art_untreat | E |
| 195 | 219 | 0.780599181 | cat1 | from_1614 | to_1614 | art_short | art_untreat | C |
| 200 | 219 | 0.774484672 | cat1 | from_1615 | to_1615 | art_short | art_untreat | C |
| 206 | 219 | 0.849843566 | cat1 | from_1616 | to_1616 | art_short | art_untreat | C |
| 211 | 219 | 0.774484672 | cat1 | from_1617 | to_1617 | art_short | art_untreat | C |
| 217 | 219 | 0.694742672 | cat1 | from_1618 | to_1618 | art_short | art_untreat | C |
| 218 | 219 | 0.774484672 | cat1 | from_1619 | to_1619 | art_short | art_untreat | C |
| 12 | 220 | 0.722698239 | cat1 | from_1620 | to_1620 | art_long | art_untreat | E |
| 13 | 220 | 0.796424227 | cat1 | from_1621 | to_1621 | art_long | art_untreat | E |
| 15 | 220 | 0.830168101 | cat1 | from_1622 | to_1622 | art_long | art_untreat | E |
| 114 | 220 | 0.721771804 | cat1 | from_1623 | to_1623 | art_long | art_untreat | E |
| 162 | 220 | 0.947085684 | cat1 | from_1624 | to_1624 | art_short | art_untreat | C |
| 163 | 220 | 0.95174168 | cat1 | from_1625 | to_1625 | art_short | art_untreat | C |
| 164 | 220 | 0.748778046 | cat1 | from_1626 | to_1626 | art_short | art_untreat | C |
| 165 | 220 | 0.862285202 | cat1 | from_1627 | to_1627 | art_short | art_untreat | C |
| 167 | 220 | 0.721270601 | cat1 | from_1628 | to_1628 | art_short | art_untreat | C |
| 168 | 220 | 0.692872538 | cat1 | from_1629 | to_1629 | art_short | art_untreat | C |
| 169 | 220 | 0.95047175 | cat1 | from_1630 | to_1630 | art_short | art_untreat | C |
| 180 | 220 | 0.680919817 | cat1 | from_1631 | to_1631 | art_short | art_untreat | C |
| 192 | 220 | 0.782348865 | cat1 | from_1632 | to_1632 | art_short | art_untreat | C |
| 196 | 220 | 0.94951131 | cat1 | from_1633 | to_1633 | art_short | art_untreat | C |
| 197 | 220 | 0.778972619 | cat1 | from_1634 | to_1634 | art_short | art_untreat | C |
| 216 | 220 | 0.763059877 | cat1 | from_1635 | to_1635 | art_short | art_untreat | C |
| 25 | 221 | 0.743984247 | cat1 | from_1636 | to_1636 | art_long | art_untreat | E |
| 31 | 221 | 0.920850157 | cat1 | from_1637 | to_1637 | art_long | art_untreat | E |
| 33 | 221 | 0.906210488 | cat1 | from_1638 | to_1638 | art_long | art_untreat | E |
| 90 | 221 | 0.815559728 | cat1 | from_1639 | to_1639 | art_long | art_untreat | E |
| 110 | 221 | 0.694550179 | cat1 | from_1640 | to_1640 | art_long | art_untreat | E |
| 112 | 221 | 0.738578664 | cat1 | from_1641 | to_1641 | art_long | art_untreat | E |
| 173 | 221 | 0.743373944 | cat1 | from_1642 | to_1642 | art_short | art_untreat | C |
| 174 | 221 | 0.771137224 | cat1 | from_1643 | to_1643 | art_short | art_untreat | C |
| 194 | 221 | 0.743373944 | cat1 | from_1644 | to_1644 | art_short | art_untreat | C |
| 206 | 221 | 0.676966946 | cat1 | from_1645 | to_1645 | art_short | art_untreat | C |
| 219 | 221 | 0.7700942 | cat1 | from_1646 | to_1646 | art_untreat | art_untreat | A |
| 32 | 222 | -0.700954002 | cat1 | from_1647 | to_1647 | art_long | art_untreat | E |
| 56 | 222 | 0.829933542 | cat1 | from_1648 | to_1648 | art_long | art_untreat | E |
| 57 | 222 | 0.829933542 | cat1 | from_1649 | to_1649 | art_long | art_untreat | E |
| 161 | 222 | 0.705009029 | cat1 | from_1650 | to_1650 | art_short | art_untreat | C |
| 182 | 222 | 0.945116242 | cat1 | from_1651 | to_1651 | art_short | art_untreat | C |
| 199 | 222 | 0.765738146 | cat1 | from_1652 | to_1652 | art_short | art_untreat | C |
| 212 | 222 | 0.728343349 | cat1 | from_1653 | to_1653 | art_short | art_untreat | C |
| 44 | 223 | 0.691037992 | cat1 | from_1654 | to_1654 | art_long | art_untreat | E |
| 46 | 223 | 0.691037992 | cat1 | from_1655 | to_1655 | art_long | art_untreat | E |
| 52 | 223 | 0.856669922 | cat1 | from_1656 | to_1656 | art_long | art_untreat | E |
| 60 | 223 | 0.752473423 | cat1 | from_1657 | to_1657 | art_long | art_untreat | E |
| 61 | 223 | 0.753766631 | cat1 | from_1658 | to_1658 | art_long | art_untreat | E |
| 99 | 223 | 0.74862913 | cat1 | from_1659 | to_1659 | art_long | art_untreat | E |
| 102 | 223 | 0.718476273 | cat1 | from_1660 | to_1660 | art_long | art_untreat | E |
| 105 | 223 | 0.698715038 | cat1 | from_1661 | to_1661 | art_long | art_untreat | E |
| 106 | 223 | 0.691037992 | cat1 | from_1662 | to_1662 | art_long | art_untreat | E |

| | | | | | | | | |
|---|---|---|---|---|---|---|---|---|
| 128 | 223 | 0.70190414 | cat1 | from_1663 | to_1663 | art_long | art_untreat | E |
| 135 | 223 | 0.691037992 | cat1 | from_1664 | to_1664 | art_long | art_untreat | E |
| 148 | 223 | 0.74862913 | cat1 | from_1665 | to_1665 | art_long | art_untreat | E |
| 176 | 223 | 0.921110008 | cat1 | from_1666 | to_1666 | art_short | art_untreat | C |
| 183 | 223 | 0.923314343 | cat1 | from_1667 | to_1667 | art_short | art_untreat | C |
| 203 | 223 | 0.921110008 | cat1 | from_1668 | to_1668 | art_short | art_untreat | C |
| 204 | 223 | 0.919755044 | cat1 | from_1669 | to_1669 | art_short | art_untreat | C |
| 29 | 224 | 0.807273907 | cat1 | from_1670 | to_1670 | art_long | art_untreat | E |
| 71 | 224 | 0.759302147 | cat1 | from_1671 | to_1671 | art_long | art_untreat | E |
| 89 | 224 | 0.694158236 | cat1 | from_1672 | to_1672 | art_long | art_untreat | E |
| 124 | 224 | 0.89652885 | cat1 | from_1673 | to_1673 | art_long | art_untreat | E |
| 150 | 224 | 0.694158236 | cat1 | from_1674 | to_1674 | art_long | art_untreat | E |
| 175 | 224 | 0.698069089 | cat1 | from_1675 | to_1675 | art_short | art_untreat | C |
| 181 | 224 | 0.793797689 | cat1 | from_1676 | to_1676 | art_short | art_untreat | C |
| 208 | 224 | 0.818812664 | cat1 | from_1677 | to_1677 | art_short | art_untreat | C |
| 209 | 224 | 0.741503328 | cat1 | from_1678 | to_1678 | art_short | art_untreat | C |
| 210 | 224 | 0.741503328 | cat1 | from_1679 | to_1679 | art_short | art_untreat | C |
| 30 | 225 | 0.879247336 | cat1 | from_1680 | to_1680 | art_long | art_untreat | E |
| 47 | 225 | 0.704510767 | cat1 | from_1681 | to_1681 | art_long | art_untreat | E |
| 54 | 225 | 0.709879723 | cat1 | from_1682 | to_1682 | art_long | art_untreat | E |
| 55 | 225 | 0.704510767 | cat1 | from_1683 | to_1683 | art_long | art_untreat | E |
| 58 | 225 | 0.704510767 | cat1 | from_1684 | to_1684 | art_long | art_untreat | E |
| 69 | 225 | 0.714920421 | cat1 | from_1685 | to_1685 | art_long | art_untreat | E |
| 72 | 225 | 0.759370984 | cat1 | from_1686 | to_1686 | art_long | art_untreat | E |
| 77 | 225 | 0.764286986 | cat1 | from_1687 | to_1687 | art_long | art_untreat | E |
| 80 | 225 | 0.774114858 | cat1 | from_1688 | to_1688 | art_long | art_untreat | E |
| 81 | 225 | 0.770515474 | cat1 | from_1689 | to_1689 | art_long | art_untreat | E |
| 83 | 225 | 0.808220605 | cat1 | from_1690 | to_1690 | art_long | art_untreat | E |
| 87 | 225 | 0.704510767 | cat1 | from_1691 | to_1691 | art_long | art_untreat | E |
| 95 | 225 | 0.704510767 | cat1 | from_1692 | to_1692 | art_long | art_untreat | E |
| 109 | 225 | 0.714920421 | cat1 | from_1693 | to_1693 | art_long | art_untreat | E |
| 111 | 225 | 0.709698219 | cat1 | from_1694 | to_1694 | art_long | art_untreat | E |
| 119 | 225 | 0.709698219 | cat1 | from_1695 | to_1695 | art_long | art_untreat | E |
| 120 | 225 | 0.764286986 | cat1 | from_1696 | to_1696 | art_long | art_untreat | E |
| 140 | 225 | 0.769202341 | cat1 | from_1697 | to_1697 | art_long | art_untreat | E |
| 195 | 225 | 0.91910319 | cat1 | from_1698 | to_1698 | art_short | art_untreat | C |
| 200 | 225 | 0.915228448 | cat1 | from_1699 | to_1699 | art_short | art_untreat | C |
| 206 | 225 | 0.92640908 | cat1 | from_1700 | to_1700 | art_short | art_untreat | C |
| 211 | 225 | 0.915228448 | cat1 | from_1701 | to_1701 | art_short | art_untreat | C |
| 218 | 225 | 0.915228448 | cat1 | from_1702 | to_1702 | art_short | art_untreat | C |
| 219 | 225 | 0.963724349 | cat1 | from_1703 | to_1703 | art_untreat | art_untreat | A |
| 221 | 225 | 0.720600053 | cat1 | from_1704 | to_1704 | art_untreat | art_untreat | A |
| 61 | 226 | 0.706637476 | cat1 | from_1705 | to_1705 | art_long | art_untreat | E |
| 99 | 226 | 0.700325757 | cat1 | from_1706 | to_1706 | art_long | art_untreat | E |
| 105 | 226 | 0.77172687 | cat1 | from_1707 | to_1707 | art_long | art_untreat | E |
| 148 | 226 | 0.700325757 | cat1 | from_1708 | to_1708 | art_long | art_untreat | E |
| 176 | 226 | 0.931019432 | cat1 | from_1709 | to_1709 | art_short | art_untreat | C |
| 183 | 226 | 0.934627278 | cat1 | from_1710 | to_1710 | art_short | art_untreat | C |
| 203 | 226 | 0.931019432 | cat1 | from_1711 | to_1711 | art_short | art_untreat | C |
| 204 | 226 | 0.78858878 | cat1 | from_1712 | to_1712 | art_short | art_untreat | C |
| 223 | 226 | 0.939599761 | cat1 | from_1713 | to_1713 | art_untreat | art_untreat | A |
| 1 | 227 | 0.691943956 | cat1 | from_1714 | to_1714 | art_long | ec | N |
| 113 | 227 | 0.668260988 | cat1 | from_1715 | to_1715 | art_long | ec | N |
| 198 | 227 | 0.733877451 | cat1 | from_1716 | to_1716 | art_short | ec | K |
| 207 | 227 | 0.713824426 | cat1 | from_1717 | to_1717 | art_short | ec | K |
| 34 | 228 | 0.779602261 | cat1 | from_1718 | to_1718 | art_long | ec | N |
| 113 | 228 | 0.816945165 | cat1 | from_1719 | to_1719 | art_long | ec | N |
| 198 | 228 | 0.711161313 | cat1 | from_1720 | to_1720 | art_short | ec | K |
| 207 | 228 | 0.849475512 | cat1 | from_1721 | to_1721 | art_short | ec | K |
| 227 | 228 | 0.969483354 | cat1 | from_1722 | to_1722 | ec | ec | P |
| 12 | 229 | 0.807493863 | cat1 | from_1723 | to_1723 | art_long | ec | N |
| 79 | 229 | 0.742673184 | cat1 | from_1724 | to_1724 | art_long | ec | N |
| 165 | 229 | 0.710916725 | cat1 | from_1725 | to_1725 | art_short | ec | K |
| 167 | 229 | 0.800518615 | cat1 | from_1726 | to_1726 | art_short | ec | K |
| 197 | 229 | 0.759287548 | cat1 | from_1727 | to_1727 | art_short | ec | K |
| 217 | 229 | 0.842201938 | cat1 | from_1728 | to_1728 | art_short | ec | K |
| 53 | 230 | 0.67171766 | cat1 | from_1729 | to_1729 | art_long | ec | N |
| 155 | 230 | 0.749596997 | cat1 | from_1730 | to_1730 | art_short | ec | K |
| 168 | 230 | 0.860587963 | cat1 | from_1731 | to_1731 | art_short | ec | K |
| 179 | 230 | 0.770681953 | cat1 | from_1732 | to_1732 | art_short | ec | K |
| 180 | 230 | 0.907933692 | cat1 | from_1733 | to_1733 | art_short | ec | K |
| 187 | 230 | 0.88380754 | cat1 | from_1734 | to_1734 | art_short | ec | K |
| 196 | 230 | 0.763860911 | cat1 | from_1735 | to_1735 | art_short | ec | K |
| 220 | 230 | 0.736576287 | cat1 | from_1736 | to_1736 | art_untreat | ec | F |
| 34 | 231 | 0.769532503 | cat1 | from_1737 | to_1737 | art_long | ec | N |
| 113 | 231 | 0.808804782 | cat1 | from_1738 | to_1738 | art_long | ec | N |
| 198 | 231 | 0.713955802 | cat1 | from_1739 | to_1739 | art_short | ec | K |
| 207 | 231 | 0.842241873 | cat1 | from_1740 | to_1740 | art_short | ec | K |
| 227 | 231 | 0.97326363 | cat1 | from_1741 | to_1741 | ec | ec | P |
| 228 | 231 | 0.999873252 | cat1 | from_1742 | to_1742 | ec | ec | P |
| 139 | 232 | 0.687193407 | cat1 | from_1743 | to_1743 | art_long | ec | N |
| 170 | 232 | 0.722887322 | cat1 | from_1744 | to_1744 | art_short | ec | K |
| 171 | 232 | 0.716158574 | cat1 | from_1745 | to_1745 | art_short | ec | K |
| 178 | 232 | 0.792467008 | cat1 | from_1746 | to_1746 | art_short | ec | K |
| 184 | 232 | 0.70270205 | cat1 | from_1747 | to_1747 | art_short | ec | K |
| 223 | 232 | 0.733526988 | cat1 | from_1748 | to_1748 | art_untreat | ec | F |
| 226 | 232 | 0.735425899 | cat1 | from_1749 | to_1749 | art_untreat | ec | F |
| 26 | 233 | 0.805047772 | cat1 | from_1750 | to_1750 | art_long | ec | N |
| 115 | 233 | 0.826762205 | cat1 | from_1751 | to_1751 | art_long | ec | N |
| 130 | 233 | 0.673443662 | cat1 | from_1752 | to_1752 | art_long | ec | N |
| 136 | 233 | 0.746007988 | cat1 | from_1753 | to_1753 | art_long | ec | N |
| 153 | 233 | -0.689471286 | cat1 | from_1754 | to_1754 | art_long | ec | N |
| 41 | 234 | 0.719826945 | cat1 | from_1755 | to_1755 | art_long | ec | N |
| 82 | 234 | 0.672477792 | cat1 | from_1756 | to_1756 | art_long | ec | N |
| 89 | 234 | 0.70478219 | cat1 | from_1757 | to_1757 | art_long | ec | N |
| 96 | 234 | 0.718499776 | cat1 | from_1758 | to_1758 | art_long | ec | N |
| 98 | 234 | 0.841683294 | cat1 | from_1759 | to_1759 | art_long | ec | N |
| 121 | 234 | 0.843698626 | cat1 | from_1760 | to_1760 | art_long | ec | N |
| 150 | 234 | 0.70478219 | cat1 | from_1761 | to_1761 | art_long | ec | N |
| 175 | 234 | 0.740326276 | cat1 | from_1762 | to_1762 | art_short | ec | K |
| 209 | 234 | 0.783526332 | cat1 | from_1763 | to_1763 | art_short | ec | K |
| 210 | 234 | 0.783526332 | cat1 | from_1764 | to_1764 | art_short | ec | K |
| 29 | 235 | 0.788285407 | cat1 | from_1765 | to_1765 | art_long | ec | N |
| 41 | 235 | 0.832078752 | cat1 | from_1766 | to_1766 | art_long | ec | N |

| | | | | | | | | |
|---|---|---|---|---|---|---|---|---|
| 71 | 235 | 0.691529697 | cat1 | from_1767 | to_1767 | art_long | ec | N |
| 96 | 235 | 0.816860458 | cat1 | from_1768 | to_1768 | art_long | ec | N |
| 98 | 235 | 0.74980927 | cat1 | from_1769 | to_1769 | art_long | ec | N |
| 121 | 235 | 0.751541975 | cat1 | from_1770 | to_1770 | art_long | ec | N |
| 181 | 235 | 0.719643997 | cat1 | from_1771 | to_1771 | art_short | ec | K |
| 208 | 235 | 0.769362246 | cat1 | from_1772 | to_1772 | art_short | ec | K |
| 209 | 235 | 0.694614391 | cat1 | from_1773 | to_1773 | art_short | ec | K |
| 210 | 235 | 0.694614391 | cat1 | from_1774 | to_1774 | art_short | ec | K |
| 234 | 235 | 0.903294163 | cat1 | from_1775 | to_1775 | ec | ec | P |
| 1 | 236 | 0.698930723 | cat1 | from_1776 | to_1776 | art_long | ec | N |
| 152 | 236 | -0.669195234 | cat1 | from_1777 | to_1777 | art_long | ec | N |
| 153 | 236 | -0.780200364 | cat1 | from_1778 | to_1778 | art_long | ec | N |
| 227 | 236 | 0.772338652 | cat1 | from_1779 | to_1779 | ec | ec | P |
| 228 | 236 | 0.689290665 | cat1 | from_1780 | to_1780 | ec | ec | P |
| 231 | 236 | 0.695962663 | cat1 | from_1781 | to_1781 | ec | ec | P |
| 232 | 236 | 0.721222051 | cat1 | from_1782 | to_1782 | ec | ec | P |
| 1 | 237 | 0.712021124 | cat1 | from_1783 | to_1783 | art_long | ec | N |
| 178 | 237 | 0.80147014 | cat1 | from_1784 | to_1784 | art_short | ec | K |
| 223 | 237 | 0.707856274 | cat1 | from_1785 | to_1785 | art_untreat | ec | F |
| 232 | 237 | 0.976043603 | cat1 | from_1786 | to_1786 | ec | ec | P |
| 236 | 237 | 0.791463763 | cat1 | from_1787 | to_1787 | ec | ec | P |
| 2 | 238 | 0.77186424 | cat1 | from_1788 | to_1788 | art_long | ec | N |
| 53 | 238 | 0.80200823 | cat1 | from_1789 | to_1789 | art_long | ec | N |
| 155 | 238 | 0.894470839 | cat1 | from_1790 | to_1790 | art_short | ec | K |
| 164 | 238 | 0.68341347 | cat1 | from_1791 | to_1791 | art_short | ec | K |
| 168 | 238 | 0.731416521 | cat1 | from_1792 | to_1792 | art_short | ec | K |
| 179 | 238 | 0.919463761 | cat1 | from_1793 | to_1793 | art_short | ec | K |
| 180 | 238 | 0.754963202 | cat1 | from_1794 | to_1794 | art_short | ec | K |
| 187 | 238 | 0.727959268 | cat1 | from_1795 | to_1795 | art_short | ec | K |
| 197 | 238 | 0.727696163 | cat1 | from_1796 | to_1796 | art_short | ec | K |
| 216 | 238 | 0.756087472 | cat1 | from_1797 | to_1797 | art_short | ec | K |
| 230 | 238 | 0.869447337 | cat1 | from_1798 | to_1798 | ec | ec | P |
| 32 | 239 | -0.686524942 | cat1 | from_1799 | to_1799 | art_long | ec | N |
| 182 | 239 | 0.795066018 | cat1 | from_1800 | to_1800 | art_short | ec | K |
| 199 | 239 | 0.671551741 | cat1 | from_1801 | to_1801 | art_short | ec | K |
| 222 | 239 | 0.949601629 | cat1 | from_1802 | to_1802 | art_untreat | ec | F |
| 17 | 240 | 0.701627411 | cat1 | from_1803 | to_1803 | art_long | ec | N |
| 36 | 240 | 0.69803622 | cat1 | from_1804 | to_1804 | art_long | ec | N |
| 133 | 240 | 0.69803622 | cat1 | from_1805 | to_1805 | art_long | ec | N |
| 134 | 240 | 0.69803622 | cat1 | from_1806 | to_1806 | art_long | ec | N |
| 138 | 240 | 0.784000188 | cat1 | from_1807 | to_1807 | art_long | ec | N |
| 143 | 240 | 0.709883469 | cat1 | from_1808 | to_1808 | art_long | ec | N |
| 154 | 240 | 0.956741214 | cat1 | from_1809 | to_1809 | art_short | ec | K |
| 166 | 240 | 0.676419523 | cat1 | from_1810 | to_1810 | art_short | ec | K |
| 177 | 240 | 0.939039214 | cat1 | from_1811 | to_1811 | art_short | ec | K |
| 178 | 240 | 0.707197479 | cat1 | from_1812 | to_1812 | art_short | ec | K |
| 213 | 240 | 0.691028523 | cat1 | from_1813 | to_1813 | art_short | ec | K |
| 237 | 240 | 0.704632841 | cat1 | from_1814 | to_1814 | ec | ec | P |
| 17 | 241 | 0.7069649 | cat1 | from_1815 | to_1815 | art_long | ec | N |
| 36 | 241 | 0.703404761 | cat1 | from_1816 | to_1816 | art_long | ec | N |
| 133 | 241 | 0.703404761 | cat1 | from_1817 | to_1817 | art_long | ec | N |
| 134 | 241 | 0.703404761 | cat1 | from_1818 | to_1818 | art_long | ec | N |
| 138 | 241 | 0.788522649 | cat1 | from_1819 | to_1819 | art_long | ec | N |
| 143 | 241 | 0.711415323 | cat1 | from_1820 | to_1820 | art_long | ec | N |
| 154 | 241 | 0.957962754 | cat1 | from_1821 | to_1821 | art_short | ec | K |
| 166 | 241 | 0.678679743 | cat1 | from_1822 | to_1822 | art_short | ec | K |
| 177 | 241 | 0.940903701 | cat1 | from_1823 | to_1823 | art_short | ec | K |
| 178 | 241 | 0.707726269 | cat1 | from_1824 | to_1824 | art_short | ec | K |
| 213 | 241 | 0.69096866 | cat1 | from_1825 | to_1825 | art_short | ec | K |
| 237 | 241 | 0.70047002 | cat1 | from_1826 | to_1826 | ec | ec | P |
| 240 | 241 | 0.999968373 | cat1 | from_1827 | to_1827 | ec | ec | P |
| 157 | 242 | 0.716339803 | cat1 | from_1828 | to_1828 | art_short | ec | K |
| 198 | 242 | 0.858014017 | cat1 | from_1829 | to_1829 | art_short | ec | K |
| 201 | 242 | 0.702903437 | cat1 | from_1830 | to_1830 | art_short | ec | K |
| 217 | 242 | 0.772304632 | cat1 | from_1831 | to_1831 | art_short | ec | K |
| 227 | 242 | 0.812934289 | cat1 | from_1832 | to_1832 | ec | ec | P |
| 228 | 242 | 0.680793919 | cat1 | from_1833 | to_1833 | ec | ec | P |
| 231 | 242 | 0.690638321 | cat1 | from_1834 | to_1834 | ec | ec | P |
| 34 | 243 | 0.784551281 | cat1 | from_1835 | to_1835 | art_long | ec | N |
| 113 | 243 | 0.820928254 | cat1 | from_1836 | to_1836 | art_long | ec | N |
| 161 | 243 | 0.66695857 | cat1 | from_1837 | to_1837 | art_short | ec | K |
| 198 | 243 | 0.709700007 | cat1 | from_1838 | to_1838 | art_short | ec | K |
| 207 | 243 | 0.853003227 | cat1 | from_1839 | to_1839 | art_short | ec | K |
| 227 | 243 | 0.96750585 | cat1 | from_1840 | to_1840 | ec | ec | P |
| 228 | 243 | 0.999968465 | cat1 | from_1841 | to_1841 | ec | ec | P |
| 231 | 243 | 0.999715284 | cat1 | from_1842 | to_1842 | ec | ec | P |
| 236 | 243 | 0.685897293 | cat1 | from_1843 | to_1843 | ec | ec | P |
| 242 | 243 | 0.675818926 | cat1 | from_1844 | to_1844 | ec | ec | P |
| 236 | 244 | 0.673806222 | cat1 | from_1845 | to_1845 | ec | ec | P |
| 242 | 244 | 0.707062675 | cat1 | from_1846 | to_1846 | ec | ec | P |
| 81 | 245 | 0.690481313 | cat1 | from_1847 | to_1847 | art_long | ec | N |
| 145 | 245 | 0.714602137 | cat1 | from_1848 | to_1848 | art_long | ec | N |
| 206 | 245 | 0.738930826 | cat1 | from_1849 | to_1849 | art_short | ec | K |
| 217 | 245 | 0.83931829 | cat1 | from_1850 | to_1850 | art_short | ec | K |
| 219 | 245 | 0.890716177 | cat1 | from_1851 | to_1851 | art_untreat | ec | F |
| 221 | 245 | 0.764045833 | cat1 | from_1852 | to_1852 | art_untreat | ec | F |
| 225 | 245 | 0.773298368 | cat1 | from_1853 | to_1853 | art_untreat | ec | F |
| 229 | 245 | 0.686820926 | cat1 | from_1854 | to_1854 | ec | ec | P |
| 242 | 245 | 0.727845689 | cat1 | from_1855 | to_1855 | ec | ec | P |
| 49 | 246 | -0.676891691 | cat1 | from_1856 | to_1856 | art_long | ec | N |
| 52 | 246 | 0.670239036 | cat1 | from_1857 | to_1857 | art_long | ec | N |
| 75 | 246 | -0.679413701 | cat1 | from_1858 | to_1858 | art_long | ec | N |
| 86 | 246 | 0.673278653 | cat1 | from_1859 | to_1859 | art_long | ec | N |
| 94 | 246 | 0.673278653 | cat1 | from_1860 | to_1860 | art_long | ec | N |
| 219 | 246 | 0.698300165 | cat1 | from_1861 | to_1861 | art_untreat | ec | F |
| 223 | 246 | 0.687312873 | cat1 | from_1862 | to_1862 | art_untreat | ec | F |
| 225 | 246 | 0.696890508 | cat1 | from_1863 | to_1863 | art_untreat | ec | F |
| 12 | 247 | 0.952596754 | cat1 | from_1864 | to_1864 | art_long | ec | N |
| 13 | 247 | 0.703459817 | cat1 | from_1865 | to_1865 | art_long | ec | N |
| 19 | 247 | 0.920956059 | cat1 | from_1866 | to_1866 | art_long | ec | N |
| 39 | 247 | 0.826394325 | cat1 | from_1867 | to_1867 | art_long | ec | N |
| 40 | 247 | 0.918131836 | cat1 | from_1868 | to_1868 | art_long | ec | N |
| 79 | 247 | 0.966599502 | cat1 | from_1869 | to_1869 | art_long | ec | N |
| 114 | 247 | 0.807980714 | cat1 | from_1870 | to_1870 | art_long | ec | N |

| | | | | | | | | |
|---|---|---|---|---|---|---|---|---|
| 145 | 247 | 0.700776203 | cat1 | from_1871 | to_1871 | art_long | ec | N |
| 156 | 247 | 0.736252336 | cat1 | from_1872 | to_1872 | art_short | ec | K |
| 158 | 247 | 0.735164432 | cat1 | from_1873 | to_1873 | art_short | ec | K |
| 163 | 247 | 0.71666194 | cat1 | from_1874 | to_1874 | art_short | ec | K |
| 164 | 247 | 0.793416508 | cat1 | from_1875 | to_1875 | art_short | ec | K |
| 165 | 247 | 0.822478643 | cat1 | from_1876 | to_1876 | art_short | ec | K |
| 167 | 247 | 0.950242589 | cat1 | from_1877 | to_1877 | art_short | ec | K |
| 169 | 247 | 0.713936397 | cat1 | from_1878 | to_1878 | art_short | ec | K |
| 197 | 247 | 0.843590359 | cat1 | from_1879 | to_1879 | art_short | ec | K |
| 229 | 247 | 0.884298369 | cat1 | from_1880 | to_1880 | ec | ec | P |
| 61 | 248 | 0.858027382 | cat1 | from_1881 | to_1881 | art_long | ec | N |
| 99 | 248 | 0.857289398 | cat1 | from_1882 | to_1882 | art_long | ec | N |
| 105 | 248 | 0.973657525 | cat1 | from_1883 | to_1883 | art_long | ec | N |
| 148 | 248 | 0.857289398 | cat1 | from_1884 | to_1884 | art_long | ec | N |
| 151 | 248 | 0.764767542 | cat1 | from_1885 | to_1885 | art_long | ec | N |
| 152 | 248 | 0.916638307 | cat1 | from_1886 | to_1886 | art_long | ec | N |
| 153 | 248 | 0.893172948 | cat1 | from_1887 | to_1887 | art_long | ec | N |
| 176 | 248 | 0.817854567 | cat1 | from_1888 | to_1888 | art_short | ec | K |
| 183 | 248 | 0.815096833 | cat1 | from_1889 | to_1889 | art_short | ec | K |
| 203 | 248 | 0.817854567 | cat1 | from_1890 | to_1890 | art_short | ec | K |
| 4 | 7 | 0.999742218 | cat2 | from_1891 | to_1891 | art_long | art_long | M |
| 6 | 8 | 0.645803529 | cat2 | from_1892 | to_1892 | art_long | art_long | M |
| 8 | 9 | 0.900526435 | cat2 | from_1893 | to_1893 | art_long | art_long | M |
| 8 | 10 | 0.901211849 | cat2 | from_1894 | to_1894 | art_long | art_long | M |
| 9 | 10 | 0.999846186 | cat2 | from_1895 | to_1895 | art_long | art_long | M |
| 4 | 11 | 0.797633075 | cat2 | from_1896 | to_1896 | art_long | art_long | M |
| 7 | 11 | 0.799085935 | cat2 | from_1897 | to_1897 | art_long | art_long | M |
| 12 | 13 | 0.734675369 | cat2 | from_1898 | to_1898 | art_long | art_long | M |
| 5 | 15 | 0.743922825 | cat2 | from_1899 | to_1899 | art_long | art_long | M |
| 13 | 15 | 0.821714796 | cat2 | from_1900 | to_1900 | art_long | art_long | M |
| 2 | 16 | 0.753316046 | cat2 | from_1901 | to_1901 | art_long | art_long | M |
| 12 | 16 | 0.83880115 | cat2 | from_1902 | to_1902 | art_long | art_long | M |
| 13 | 16 | 0.655681778 | cat2 | from_1903 | to_1903 | art_long | art_long | M |
| 15 | 16 | 0.776813743 | cat2 | from_1904 | to_1904 | art_long | art_long | M |
| 2 | 18 | 0.753836303 | cat2 | from_1905 | to_1905 | art_long | art_long | M |
| 12 | 18 | 0.839656321 | cat2 | from_1906 | to_1906 | art_long | art_long | M |
| 13 | 18 | 0.656914239 | cat2 | from_1907 | to_1907 | art_long | art_long | M |
| 15 | 18 | 0.777409834 | cat2 | from_1908 | to_1908 | art_long | art_long | M |
| 16 | 18 | 0.9999672 | cat2 | from_1909 | to_1909 | art_long | art_long | M |
| 12 | 19 | 0.944654806 | cat2 | from_1910 | to_1910 | art_long | art_long | M |
| 13 | 19 | 0.794979246 | cat2 | from_1911 | to_1911 | art_long | art_long | M |
| 16 | 19 | 0.81425755 | cat2 | from_1912 | to_1912 | art_long | art_long | M |
| 18 | 19 | 0.81491429 | cat2 | from_1913 | to_1913 | art_long | art_long | M |
| 13 | 20 | 0.772712406 | cat2 | from_1914 | to_1914 | art_long | art_long | M |
| 19 | 20 | 0.666350672 | cat2 | from_1915 | to_1915 | art_long | art_long | M |
| 13 | 21 | 0.674321871 | cat2 | from_1916 | to_1916 | art_long | art_long | M |
| 15 | 21 | 0.738331917 | cat2 | from_1917 | to_1917 | art_long | art_long | M |
| 20 | 21 | 0.844213578 | cat2 | from_1918 | to_1918 | art_long | art_long | M |
| 9 | 22 | 0.771361336 | cat2 | from_1919 | to_1919 | art_long | art_long | M |
| 10 | 22 | 0.771135386 | cat2 | from_1920 | to_1920 | art_long | art_long | M |
| 11 | 22 | 0.720358771 | cat2 | from_1921 | to_1921 | art_long | art_long | M |
| 8 | 23 | 0.719370703 | cat2 | from_1922 | to_1922 | art_long | art_long | M |
| 9 | 23 | 0.948832165 | cat2 | from_1923 | to_1923 | art_long | art_long | M |
| 10 | 23 | 0.948420641 | cat2 | from_1924 | to_1924 | art_long | art_long | M |
| 22 | 23 | 0.932613091 | cat2 | from_1925 | to_1925 | art_long | art_long | M |
| 14 | 24 | 0.998987674 | cat2 | from_1926 | to_1926 | art_long | art_long | M |
| 14 | 25 | 0.73898095 | cat2 | from_1927 | to_1927 | art_long | art_long | M |
| 24 | 25 | 0.742181291 | cat2 | from_1928 | to_1928 | art_long | art_long | M |
| 14 | 26 | 0.641317939 | cat2 | from_1929 | to_1929 | art_long | art_long | M |
| 24 | 26 | 0.646052629 | cat2 | from_1930 | to_1930 | art_long | art_long | M |
| 22 | 27 | 0.717188323 | cat2 | from_1931 | to_1931 | art_long | art_long | M |
| 14 | 28 | 0.998886463 | cat2 | from_1932 | to_1932 | art_long | art_long | M |
| 24 | 28 | 0.999913553 | cat2 | from_1933 | to_1933 | art_long | art_long | M |
| 25 | 28 | 0.745383727 | cat2 | from_1934 | to_1934 | art_long | art_long | M |
| 26 | 28 | 0.646304219 | cat2 | from_1935 | to_1935 | art_long | art_long | M |
| 3 | 29 | 0.68194502 | cat2 | from_1936 | to_1936 | art_long | art_long | M |
| 25 | 31 | 0.924691024 | cat2 | from_1937 | to_1937 | art_long | art_long | M |
| 14 | 32 | 0.633323671 | cat2 | from_1938 | to_1938 | art_long | art_long | M |
| 24 | 32 | 0.639292009 | cat2 | from_1939 | to_1939 | art_long | art_long | M |
| 28 | 32 | 0.638570903 | cat2 | from_1940 | to_1940 | art_long | art_long | M |
| 25 | 33 | 0.940703801 | cat2 | from_1941 | to_1941 | art_long | art_long | M |
| 31 | 33 | 0.998976021 | cat2 | from_1942 | to_1942 | art_long | art_long | M |
| 4 | 34 | 0.715442801 | cat2 | from_1943 | to_1943 | art_long | art_long | M |
| 5 | 34 | 0.672525433 | cat2 | from_1944 | to_1944 | art_long | art_long | M |
| 7 | 34 | 0.717061105 | cat2 | from_1945 | to_1945 | art_long | art_long | M |
| 22 | 34 | 0.675836566 | cat2 | from_1946 | to_1946 | art_long | art_long | M |
| 17 | 35 | 0.790463689 | cat2 | from_1947 | to_1947 | art_long | art_long | M |
| 17 | 36 | 0.999962479 | cat2 | from_1948 | to_1948 | art_long | art_long | M |
| 35 | 36 | 0.79081234 | cat2 | from_1949 | to_1949 | art_long | art_long | M |
| 26 | 37 | 0.956951603 | cat2 | from_1950 | to_1950 | art_long | art_long | M |
| 17 | 38 | 0.78695135 | cat2 | from_1951 | to_1951 | art_long | art_long | M |
| 35 | 38 | 0.999510916 | cat2 | from_1952 | to_1952 | art_long | art_long | M |
| 36 | 38 | 0.787059298 | cat2 | from_1953 | to_1953 | art_long | art_long | M |
| 2 | 39 | 0.678575665 | cat2 | from_1954 | to_1954 | art_long | art_long | M |
| 12 | 39 | 0.938079872 | cat2 | from_1955 | to_1955 | art_long | art_long | M |
| 13 | 39 | 0.768792787 | cat2 | from_1956 | to_1956 | art_long | art_long | M |
| 15 | 39 | 0.69677759 | cat2 | from_1957 | to_1957 | art_long | art_long | M |
| 16 | 39 | 0.94647313 | cat2 | from_1958 | to_1958 | art_long | art_long | M |
| 18 | 39 | 0.947167774 | cat2 | from_1959 | to_1959 | art_long | art_long | M |
| 19 | 39 | 0.9562991 | cat2 | from_1960 | to_1960 | art_long | art_long | M |
| 12 | 40 | 0.923214796 | cat2 | from_1961 | to_1961 | art_long | art_long | M |
| 13 | 40 | 0.766783516 | cat2 | from_1962 | to_1962 | art_long | art_long | M |
| 16 | 40 | 0.819199564 | cat2 | from_1963 | to_1963 | art_long | art_long | M |
| 18 | 40 | 0.818391592 | cat2 | from_1964 | to_1964 | art_long | art_long | M |
| 19 | 40 | 0.983578382 | cat2 | from_1965 | to_1965 | art_long | art_long | M |
| 39 | 40 | 0.942013925 | cat2 | from_1966 | to_1966 | art_long | art_long | M |
| 29 | 41 | 0.818093502 | cat2 | from_1967 | to_1967 | art_long | art_long | M |
| 17 | 42 | 0.637178301 | cat2 | from_1968 | to_1968 | art_long | art_long | M |
| 36 | 42 | 0.639368627 | cat2 | from_1969 | to_1969 | art_long | art_long | M |
| 42 | 43 | 0.901556008 | cat2 | from_1970 | to_1970 | art_long | art_long | M |
| 11 | 44 | 0.634727518 | cat2 | from_1971 | to_1971 | art_long | art_long | M |
| 35 | 44 | 0.641463373 | cat2 | from_1972 | to_1972 | art_long | art_long | M |
| 38 | 44 | 0.635825037 | cat2 | from_1973 | to_1973 | art_long | art_long | M |
| 42 | 44 | 0.668807416 | cat2 | from_1974 | to_1974 | art_long | art_long | M |

| | | | | | | | | |
|---|---|---|---|---|---|---|---|---|
| 4 | 45 | 0.655215435 | cat2 | from_1975 | to_1975 | art_long | art_long | M |
| 7 | 45 | 0.654541789 | cat2 | from_1976 | to_1976 | art_long | art_long | M |
| 17 | 45 | 0.664514252 | cat2 | from_1977 | to_1977 | art_long | art_long | M |
| 36 | 45 | 0.665255524 | cat2 | from_1978 | to_1978 | art_long | art_long | M |
| 42 | 45 | 0.976810147 | cat2 | from_1979 | to_1979 | art_long | art_long | M |
| 43 | 45 | 0.929072947 | cat2 | from_1980 | to_1980 | art_long | art_long | M |
| 44 | 45 | 0.634738876 | cat2 | from_1981 | to_1981 | art_long | art_long | M |
| 11 | 46 | 0.645925315 | cat2 | from_1982 | to_1982 | art_long | art_long | M |
| 35 | 46 | 0.653177964 | cat2 | from_1983 | to_1983 | art_long | art_long | M |
| 38 | 46 | 0.648994592 | cat2 | from_1984 | to_1984 | art_long | art_long | M |
| 42 | 46 | 0.658950122 | cat2 | from_1985 | to_1985 | art_long | art_long | M |
| 44 | 46 | 0.993879162 | cat2 | from_1986 | to_1986 | art_long | art_long | M |
| 45 | 46 | 0.648528092 | cat2 | from_1987 | to_1987 | art_long | art_long | M |
| 14 | 47 | 0.663038341 | cat2 | from_1988 | to_1988 | art_long | art_long | M |
| 24 | 47 | 0.655841327 | cat2 | from_1989 | to_1989 | art_long | art_long | M |
| 27 | 47 | 0.652577961 | cat2 | from_1990 | to_1990 | art_long | art_long | M |
| 28 | 47 | 0.655163901 | cat2 | from_1991 | to_1991 | art_long | art_long | M |
| 30 | 47 | 0.951254414 | cat2 | from_1992 | to_1992 | art_long | art_long | M |
| 17 | 48 | 0.790523528 | cat2 | from_1993 | to_1993 | art_long | art_long | M |
| 35 | 48 | 0.999999616 | cat2 | from_1994 | to_1994 | art_long | art_long | M |
| 36 | 48 | 0.790866187 | cat2 | from_1995 | to_1995 | art_long | art_long | M |
| 38 | 48 | 0.999522606 | cat2 | from_1996 | to_1996 | art_long | art_long | M |
| 44 | 48 | 0.641274179 | cat2 | from_1997 | to_1997 | art_long | art_long | M |
| 46 | 48 | 0.653084509 | cat2 | from_1998 | to_1998 | art_long | art_long | M |
| 35 | 50 | 0.807299993 | cat2 | from_1999 | to_1999 | art_long | art_long | M |
| 38 | 50 | 0.815510501 | cat2 | from_2000 | to_2000 | art_long | art_long | M |
| 48 | 50 | 0.807403054 | cat2 | from_2001 | to_2001 | art_long | art_long | M |
| 44 | 52 | 0.864306126 | cat2 | from_2002 | to_2002 | art_long | art_long | M |
| 46 | 52 | 0.87108964 | cat2 | from_2003 | to_2003 | art_long | art_long | M |
| 2 | 53 | 0.997432922 | cat2 | from_2004 | to_2004 | art_long | art_long | M |
| 16 | 53 | 0.745945118 | cat2 | from_2005 | to_2005 | art_long | art_long | M |
| 18 | 53 | 0.746862949 | cat2 | from_2006 | to_2006 | art_long | art_long | M |
| 39 | 53 | 0.681899311 | cat2 | from_2007 | to_2007 | art_long | art_long | M |
| 25 | 54 | 0.718051048 | cat2 | from_2008 | to_2008 | art_long | art_long | M |
| 30 | 54 | 0.777063825 | cat2 | from_2009 | to_2009 | art_long | art_long | M |
| 31 | 54 | 0.693176831 | cat2 | from_2010 | to_2010 | art_long | art_long | M |
| 33 | 54 | 0.701355711 | cat2 | from_2011 | to_2011 | art_long | art_long | M |
| 47 | 54 | 0.714800152 | cat2 | from_2012 | to_2012 | art_long | art_long | M |
| 14 | 55 | 0.663616622 | cat2 | from_2013 | to_2013 | art_long | art_long | M |
| 24 | 55 | 0.656699188 | cat2 | from_2014 | to_2014 | art_long | art_long | M |
| 27 | 55 | 0.65322939 | cat2 | from_2015 | to_2015 | art_long | art_long | M |
| 28 | 55 | 0.656023212 | cat2 | from_2016 | to_2016 | art_long | art_long | M |
| 30 | 55 | 0.951925872 | cat2 | from_2017 | to_2017 | art_long | art_long | M |
| 47 | 55 | 0.999980207 | cat2 | from_2018 | to_2018 | art_long | art_long | M |
| 54 | 55 | 0.715879146 | cat2 | from_2019 | to_2019 | art_long | art_long | M |
| 22 | 56 | 0.709977707 | cat2 | from_2020 | to_2020 | art_long | art_long | M |
| 49 | 56 | 0.703727788 | cat2 | from_2021 | to_2021 | art_long | art_long | M |
| 22 | 57 | 0.70895525 | cat2 | from_2022 | to_2022 | art_long | art_long | M |
| 49 | 57 | 0.704951677 | cat2 | from_2023 | to_2023 | art_long | art_long | M |
| 56 | 57 | 0.999945099 | cat2 | from_2024 | to_2024 | art_long | art_long | M |
| 14 | 58 | 0.669273006 | cat2 | from_2025 | to_2025 | art_long | art_long | M |
| 24 | 58 | 0.666837007 | cat2 | from_2026 | to_2026 | art_long | art_long | M |
| 27 | 58 | 0.660103994 | cat2 | from_2027 | to_2027 | art_long | art_long | M |
| 28 | 58 | 0.66618767 | cat2 | from_2028 | to_2028 | art_long | art_long | M |
| 30 | 58 | 0.957527001 | cat2 | from_2029 | to_2029 | art_long | art_long | M |
| 47 | 58 | 0.994336224 | cat2 | from_2030 | to_2030 | art_long | art_long | M |
| 54 | 58 | 0.729216967 | cat2 | from_2031 | to_2031 | art_long | art_long | M |
| 55 | 58 | 0.994985233 | cat2 | from_2032 | to_2032 | art_long | art_long | M |
| 14 | 59 | 0.99880583 | cat2 | from_2033 | to_2033 | art_long | art_long | M |
| 24 | 59 | 0.999972814 | cat2 | from_2034 | to_2034 | art_long | art_long | M |
| 25 | 59 | 0.743960427 | cat2 | from_2035 | to_2035 | art_long | art_long | M |
| 26 | 59 | 0.646484694 | cat2 | from_2036 | to_2036 | art_long | art_long | M |
| 28 | 59 | 0.999973244 | cat2 | from_2037 | to_2037 | art_long | art_long | M |
| 32 | 59 | 0.639331578 | cat2 | from_2038 | to_2038 | art_long | art_long | M |
| 47 | 59 | 0.654943204 | cat2 | from_2039 | to_2039 | art_long | art_long | M |
| 55 | 59 | 0.655822088 | cat2 | from_2040 | to_2040 | art_long | art_long | M |
| 58 | 59 | 0.666298776 | cat2 | from_2041 | to_2041 | art_long | art_long | M |
| 11 | 60 | 0.63691726 | cat2 | from_2042 | to_2042 | art_long | art_long | M |
| 35 | 60 | 0.642643395 | cat2 | from_2043 | to_2043 | art_long | art_long | M |
| 38 | 60 | 0.641580232 | cat2 | from_2044 | to_2044 | art_long | art_long | M |
| 42 | 60 | 0.633719735 | cat2 | from_2045 | to_2045 | art_long | art_long | M |
| 44 | 60 | 0.983050078 | cat2 | from_2046 | to_2046 | art_long | art_long | M |
| 45 | 60 | 0.633979893 | cat2 | from_2047 | to_2047 | art_long | art_long | M |
| 46 | 60 | 0.994762607 | cat2 | from_2048 | to_2048 | art_long | art_long | M |
| 48 | 60 | 0.642594815 | cat2 | from_2049 | to_2049 | art_long | art_long | M |
| 52 | 60 | 0.898397052 | cat2 | from_2050 | to_2050 | art_long | art_long | M |
| 44 | 61 | 0.891016179 | cat2 | from_2051 | to_2051 | art_long | art_long | M |
| 46 | 61 | 0.908749095 | cat2 | from_2052 | to_2052 | art_long | art_long | M |
| 50 | 61 | 0.651715038 | cat2 | from_2053 | to_2053 | art_long | art_long | M |
| 52 | 61 | 0.726831256 | cat2 | from_2054 | to_2054 | art_long | art_long | M |
| 60 | 61 | 0.924458309 | cat2 | from_2055 | to_2055 | art_long | art_long | M |
| 4 | 62 | 0.632964197 | cat2 | from_2056 | to_2056 | art_long | art_long | M |
| 7 | 62 | 0.635634673 | cat2 | from_2057 | to_2057 | art_long | art_long | M |
| 17 | 62 | 0.644974053 | cat2 | from_2058 | to_2058 | art_long | art_long | M |
| 36 | 62 | 0.646734335 | cat2 | from_2059 | to_2059 | art_long | art_long | M |
| 42 | 62 | 0.998400781 | cat2 | from_2060 | to_2060 | art_long | art_long | M |
| 43 | 62 | 0.914289834 | cat2 | from_2061 | to_2061 | art_long | art_long | M |
| 44 | 62 | 0.662314209 | cat2 | from_2062 | to_2062 | art_long | art_long | M |
| 45 | 62 | 0.987014145 | cat2 | from_2063 | to_2063 | art_long | art_long | M |
| 46 | 62 | 0.658471246 | cat2 | from_2064 | to_2064 | art_long | art_long | M |
| 60 | 62 | 0.637000823 | cat2 | from_2065 | to_2065 | art_long | art_long | M |
| 5 | 63 | 0.964993174 | cat2 | from_2066 | to_2066 | art_long | art_long | M |
| 13 | 63 | 0.643623596 | cat2 | from_2067 | to_2067 | art_long | art_long | M |
| 15 | 63 | 0.717353552 | cat2 | from_2068 | to_2068 | art_long | art_long | M |
| 14 | 64 | 0.662607617 | cat2 | from_2069 | to_2069 | art_long | art_long | M |
| 24 | 64 | 0.666135326 | cat2 | from_2070 | to_2070 | art_long | art_long | M |
| 27 | 64 | 0.80463147 | cat2 | from_2071 | to_2071 | art_long | art_long | M |
| 28 | 64 | 0.664218091 | cat2 | from_2072 | to_2072 | art_long | art_long | M |
| 34 | 64 | 0.712017975 | cat2 | from_2073 | to_2073 | art_long | art_long | M |
| 50 | 64 | -0.633610531 | cat2 | from_2074 | to_2074 | art_long | art_long | M |
| 59 | 64 | 0.665402093 | cat2 | from_2075 | to_2075 | art_long | art_long | M |
| 9 | 65 | 0.642844959 | cat2 | from_2076 | to_2076 | art_long | art_long | M |
| 10 | 65 | 0.642256387 | cat2 | from_2077 | to_2077 | art_long | art_long | M |
| 22 | 65 | 0.700550607 | cat2 | from_2078 | to_2078 | art_long | art_long | M |

| | | | | | | | | |
|---|---|---|---|---|---|---|---|---|
| 23 | 65 | 0.710872528 | cat2 | from_2079 | to_2079 | art_long | art_long | M |
| 34 | 65 | 0.828480697 | cat2 | from_2080 | to_2080 | art_long | art_long | M |
| 49 | 65 | 0.662543035 | cat2 | from_2081 | to_2081 | art_long | art_long | M |
| 63 | 65 | 0.635750412 | cat2 | from_2082 | to_2082 | art_long | art_long | M |
| 29 | 66 | 0.831818044 | cat2 | from_2083 | to_2083 | art_long | art_long | M |
| 41 | 66 | 0.668986062 | cat2 | from_2084 | to_2084 | art_long | art_long | M |
| 29 | 67 | 0.831349117 | cat2 | from_2085 | to_2085 | art_long | art_long | M |
| 41 | 67 | 0.669895414 | cat2 | from_2086 | to_2086 | art_long | art_long | M |
| 66 | 67 | 0.999795627 | cat2 | from_2087 | to_2087 | art_long | art_long | M |
| 13 | 68 | 0.773406036 | cat2 | from_2088 | to_2088 | art_long | art_long | M |
| 19 | 68 | 0.672866834 | cat2 | from_2089 | to_2089 | art_long | art_long | M |
| 25 | 68 | 0.667368688 | cat2 | from_2090 | to_2090 | art_long | art_long | M |
| 33 | 68 | 0.641005732 | cat2 | from_2091 | to_2091 | art_long | art_long | M |
| 40 | 68 | 0.642504178 | cat2 | from_2092 | to_2092 | art_long | art_long | M |
| 14 | 69 | 0.666174115 | cat2 | from_2093 | to_2093 | art_long | art_long | M |
| 24 | 69 | 0.662942475 | cat2 | from_2094 | to_2094 | art_long | art_long | M |
| 27 | 69 | 0.657676958 | cat2 | from_2095 | to_2095 | art_long | art_long | M |
| 28 | 69 | 0.662474893 | cat2 | from_2096 | to_2096 | art_long | art_long | M |
| 30 | 69 | 0.961380852 | cat2 | from_2097 | to_2097 | art_long | art_long | M |
| 47 | 69 | 0.99595932 | cat2 | from_2098 | to_2098 | art_long | art_long | M |
| 54 | 69 | 0.731296569 | cat2 | from_2099 | to_2099 | art_long | art_long | M |
| 55 | 69 | 0.996496971 | cat2 | from_2100 | to_2100 | art_long | art_long | M |
| 58 | 69 | 0.999733377 | cat2 | from_2101 | to_2101 | art_long | art_long | M |
| 59 | 69 | 0.662436745 | cat2 | from_2102 | to_2102 | art_long | art_long | M |
| 49 | 70 | 0.953583245 | cat2 | from_2103 | to_2103 | art_long | art_long | M |
| 56 | 70 | 0.784930615 | cat2 | from_2104 | to_2104 | art_long | art_long | M |
| 57 | 70 | 0.786900907 | cat2 | from_2105 | to_2105 | art_long | art_long | M |
| 65 | 70 | 0.650704866 | cat2 | from_2106 | to_2106 | art_long | art_long | M |
| 3 | 71 | 0.752545492 | cat2 | from_2107 | to_2107 | art_long | art_long | M |
| 29 | 71 | 0.969041146 | cat2 | from_2108 | to_2108 | art_long | art_long | M |
| 41 | 71 | 0.83696334 | cat2 | from_2109 | to_2109 | art_long | art_long | M |
| 66 | 71 | 0.850977593 | cat2 | from_2110 | to_2110 | art_long | art_long | M |
| 67 | 71 | 0.850338238 | cat2 | from_2111 | to_2111 | art_long | art_long | M |
| 25 | 72 | 0.655667394 | cat2 | from_2112 | to_2112 | art_long | art_long | M |
| 30 | 72 | 0.926457451 | cat2 | from_2113 | to_2113 | art_long | art_long | M |
| 47 | 72 | 0.919028228 | cat2 | from_2114 | to_2114 | art_long | art_long | M |
| 54 | 72 | 0.93177923 | cat2 | from_2115 | to_2115 | art_long | art_long | M |
| 55 | 72 | 0.919440149 | cat2 | from_2116 | to_2116 | art_long | art_long | M |
| 58 | 72 | 0.921088477 | cat2 | from_2117 | to_2117 | art_long | art_long | M |
| 69 | 72 | 0.923659296 | cat2 | from_2118 | to_2118 | art_long | art_long | M |
| 14 | 73 | 0.99953714 | cat2 | from_2119 | to_2119 | art_long | art_long | M |
| 24 | 73 | 0.999893814 | cat2 | from_2120 | to_2120 | art_long | art_long | M |
| 25 | 73 | 0.741309136 | cat2 | from_2121 | to_2121 | art_long | art_long | M |
| 26 | 73 | 0.644662111 | cat2 | from_2122 | to_2122 | art_long | art_long | M |
| 28 | 73 | 0.999802565 | cat2 | from_2123 | to_2123 | art_long | art_long | M |
| 32 | 73 | 0.637500358 | cat2 | from_2124 | to_2124 | art_long | art_long | M |
| 47 | 73 | 0.658318201 | cat2 | from_2125 | to_2125 | art_long | art_long | M |
| 55 | 73 | 0.659085684 | cat2 | from_2126 | to_2126 | art_long | art_long | M |
| 58 | 73 | 0.66777401 | cat2 | from_2127 | to_2127 | art_long | art_long | M |
| 59 | 73 | 0.999816521 | cat2 | from_2128 | to_2128 | art_long | art_long | M |
| 64 | 73 | 0.665140257 | cat2 | from_2129 | to_2129 | art_long | art_long | M |
| 69 | 73 | 0.664136359 | cat2 | from_2130 | to_2130 | art_long | art_long | M |
| 14 | 74 | 0.99923661 | cat2 | from_2131 | to_2131 | art_long | art_long | M |
| 24 | 74 | 0.999982457 | cat2 | from_2132 | to_2132 | art_long | art_long | M |
| 25 | 74 | 0.741845797 | cat2 | from_2133 | to_2133 | art_long | art_long | M |
| 26 | 74 | 0.645503972 | cat2 | from_2134 | to_2134 | art_long | art_long | M |
| 28 | 74 | 0.999894056 | cat2 | from_2135 | to_2135 | art_long | art_long | M |
| 32 | 74 | 0.63858013 | cat2 | from_2136 | to_2136 | art_long | art_long | M |
| 47 | 74 | 0.65686491 | cat2 | from_2137 | to_2137 | art_long | art_long | M |
| 55 | 74 | 0.657686057 | cat2 | from_2138 | to_2138 | art_long | art_long | M |
| 58 | 74 | 0.667234956 | cat2 | from_2139 | to_2139 | art_long | art_long | M |
| 59 | 74 | 0.999934903 | cat2 | from_2140 | to_2140 | art_long | art_long | M |
| 64 | 74 | 0.665747923 | cat2 | from_2141 | to_2141 | art_long | art_long | M |
| 69 | 74 | 0.66344474 | cat2 | from_2142 | to_2142 | art_long | art_long | M |
| 73 | 74 | 0.999962591 | cat2 | from_2143 | to_2143 | art_long | art_long | M |
| 49 | 75 | 0.999339877 | cat2 | from_2144 | to_2144 | art_long | art_long | M |
| 56 | 75 | 0.701527069 | cat2 | from_2145 | to_2145 | art_long | art_long | M |
| 57 | 75 | 0.703120756 | cat2 | from_2146 | to_2146 | art_long | art_long | M |
| 65 | 75 | 0.658324778 | cat2 | from_2147 | to_2147 | art_long | art_long | M |
| 70 | 75 | 0.956672907 | cat2 | from_2148 | to_2148 | art_long | art_long | M |
| 14 | 76 | 0.929591247 | cat2 | from_2149 | to_2149 | art_long | art_long | M |
| 24 | 76 | 0.925242017 | cat2 | from_2150 | to_2150 | art_long | art_long | M |
| 25 | 76 | 0.921241161 | cat2 | from_2151 | to_2151 | art_long | art_long | M |
| 28 | 76 | 0.926827419 | cat2 | from_2152 | to_2152 | art_long | art_long | M |
| 31 | 76 | 0.770198972 | cat2 | from_2153 | to_2153 | art_long | art_long | M |
| 33 | 76 | 0.793052775 | cat2 | from_2154 | to_2154 | art_long | art_long | M |
| 47 | 76 | 0.63391457 | cat2 | from_2155 | to_2155 | art_long | art_long | M |
| 55 | 76 | 0.633986373 | cat2 | from_2156 | to_2156 | art_long | art_long | M |
| 59 | 76 | 0.925658502 | cat2 | from_2157 | to_2157 | art_long | art_long | M |
| 69 | 76 | 0.632099006 | cat2 | from_2158 | to_2158 | art_long | art_long | M |
| 72 | 76 | 0.649079133 | cat2 | from_2159 | to_2159 | art_long | art_long | M |
| 73 | 76 | 0.926856105 | cat2 | from_2160 | to_2160 | art_long | art_long | M |
| 74 | 76 | 0.925921803 | cat2 | from_2161 | to_2161 | art_long | art_long | M |
| 25 | 77 | 0.643454517 | cat2 | from_2162 | to_2162 | art_long | art_long | M |
| 30 | 77 | 0.921848925 | cat2 | from_2163 | to_2163 | art_long | art_long | M |
| 47 | 77 | 0.921368292 | cat2 | from_2164 | to_2164 | art_long | art_long | M |
| 54 | 77 | 0.919550015 | cat2 | from_2165 | to_2165 | art_long | art_long | M |
| 55 | 77 | 0.921271854 | cat2 | from_2166 | to_2166 | art_long | art_long | M |
| 58 | 77 | 0.914828904 | cat2 | from_2167 | to_2167 | art_long | art_long | M |
| 69 | 77 | 0.918936858 | cat2 | from_2168 | to_2168 | art_long | art_long | M |
| 72 | 77 | 0.996704892 | cat2 | from_2169 | to_2169 | art_long | art_long | M |
| 76 | 77 | 0.648705534 | cat2 | from_2170 | to_2170 | art_long | art_long | M |
| 14 | 78 | 0.988377039 | cat2 | from_2171 | to_2171 | art_long | art_long | M |
| 24 | 78 | 0.980562805 | cat2 | from_2172 | to_2172 | art_long | art_long | M |
| 25 | 78 | 0.720307953 | cat2 | from_2173 | to_2173 | art_long | art_long | M |
| 28 | 78 | 0.980584167 | cat2 | from_2174 | to_2174 | art_long | art_long | M |
| 47 | 78 | 0.676748176 | cat2 | from_2175 | to_2175 | art_long | art_long | M |
| 55 | 78 | 0.676375882 | cat2 | from_2176 | to_2176 | art_long | art_long | M |
| 58 | 78 | 0.666852739 | cat2 | from_2177 | to_2177 | art_long | art_long | M |
| 59 | 78 | 0.979947691 | cat2 | from_2178 | to_2178 | art_long | art_long | M |
| 64 | 78 | 0.638910931 | cat2 | from_2179 | to_2179 | art_long | art_long | M |
| 69 | 78 | 0.666664453 | cat2 | from_2180 | to_2180 | art_long | art_long | M |
| 73 | 78 | 0.983311632 | cat2 | from_2181 | to_2181 | art_long | art_long | M |
| 74 | 78 | 0.981705248 | cat2 | from_2182 | to_2182 | art_long | art_long | M |

| | | | | | | | | |
|---|---|---|---|---|---|---|---|---|
| 76 | 78 | 0.931946511 | cat2 | from_2183 | to_2183 | art_long | art_long | M |
| 12 | 79 | 0.965213133 | cat2 | from_2184 | to_2184 | art_long | art_long | M |
| 13 | 79 | 0.778890797 | cat2 | from_2185 | to_2185 | art_long | art_long | M |
| 16 | 79 | 0.765419274 | cat2 | from_2186 | to_2186 | art_long | art_long | M |
| 18 | 79 | 0.766350872 | cat2 | from_2187 | to_2187 | art_long | art_long | M |
| 19 | 79 | 0.989406359 | cat2 | from_2188 | to_2188 | art_long | art_long | M |
| 20 | 79 | 0.666500913 | cat2 | from_2189 | to_2189 | art_long | art_long | M |
| 39 | 79 | 0.926537951 | cat2 | from_2190 | to_2190 | art_long | art_long | M |
| 40 | 79 | 0.965440714 | cat2 | from_2191 | to_2191 | art_long | art_long | M |
| 68 | 79 | 0.666019412 | cat2 | from_2192 | to_2192 | art_long | art_long | M |
| 25 | 80 | 0.638540954 | cat2 | from_2193 | to_2193 | art_long | art_long | M |
| 30 | 80 | 0.920088967 | cat2 | from_2194 | to_2194 | art_long | art_long | M |
| 47 | 80 | 0.917948873 | cat2 | from_2195 | to_2195 | art_long | art_long | M |
| 54 | 80 | 0.915741942 | cat2 | from_2196 | to_2196 | art_long | art_long | M |
| 55 | 80 | 0.917669696 | cat2 | from_2197 | to_2197 | art_long | art_long | M |
| 58 | 80 | 0.908340903 | cat2 | from_2198 | to_2198 | art_long | art_long | M |
| 69 | 80 | 0.913196904 | cat2 | from_2199 | to_2199 | art_long | art_long | M |
| 72 | 80 | 0.993687519 | cat2 | from_2200 | to_2200 | art_long | art_long | M |
| 76 | 80 | 0.645435495 | cat2 | from_2201 | to_2201 | art_long | art_long | M |
| 77 | 80 | 0.99943956 | cat2 | from_2202 | to_2202 | art_long | art_long | M |
| 25 | 81 | 0.6983509 | cat2 | from_2203 | to_2203 | art_long | art_long | M |
| 30 | 81 | 0.764229561 | cat2 | from_2204 | to_2204 | art_long | art_long | M |
| 31 | 81 | 0.729220835 | cat2 | from_2205 | to_2205 | art_long | art_long | M |
| 33 | 81 | 0.730710002 | cat2 | from_2206 | to_2206 | art_long | art_long | M |
| 47 | 81 | 0.660885894 | cat2 | from_2207 | to_2207 | art_long | art_long | M |
| 54 | 81 | 0.987945039 | cat2 | from_2208 | to_2208 | art_long | art_long | M |
| 55 | 81 | 0.661809032 | cat2 | from_2209 | to_2209 | art_long | art_long | M |
| 58 | 81 | 0.672957329 | cat2 | from_2210 | to_2210 | art_long | art_long | M |
| 69 | 81 | 0.677498347 | cat2 | from_2211 | to_2211 | art_long | art_long | M |
| 72 | 81 | 0.898186274 | cat2 | from_2212 | to_2212 | art_long | art_long | M |
| 77 | 81 | 0.888888095 | cat2 | from_2213 | to_2213 | art_long | art_long | M |
| 80 | 81 | 0.888324061 | cat2 | from_2214 | to_2214 | art_long | art_long | M |
| 3 | 82 | 0.718903906 | cat2 | from_2215 | to_2215 | art_long | art_long | M |
| 29 | 82 | 0.749859462 | cat2 | from_2216 | to_2216 | art_long | art_long | M |
| 41 | 82 | 0.665700707 | cat2 | from_2217 | to_2217 | art_long | art_long | M |
| 66 | 82 | 0.67756735 | cat2 | from_2218 | to_2218 | art_long | art_long | M |
| 67 | 82 | 0.675171587 | cat2 | from_2219 | to_2219 | art_long | art_long | M |
| 71 | 82 | 0.812029012 | cat2 | from_2220 | to_2220 | art_long | art_long | M |
| 30 | 83 | 0.916287492 | cat2 | from_2221 | to_2221 | art_long | art_long | M |
| 47 | 83 | 0.901576762 | cat2 | from_2222 | to_2222 | art_long | art_long | M |
| 54 | 83 | 0.910054214 | cat2 | from_2223 | to_2223 | art_long | art_long | M |
| 55 | 83 | 0.900984861 | cat2 | from_2224 | to_2224 | art_long | art_long | M |
| 58 | 83 | 0.886773533 | cat2 | from_2225 | to_2225 | art_long | art_long | M |
| 69 | 83 | 0.893365063 | cat2 | from_2226 | to_2226 | art_long | art_long | M |
| 72 | 83 | 0.983641175 | cat2 | from_2227 | to_2227 | art_long | art_long | M |
| 76 | 83 | 0.632113763 | cat2 | from_2228 | to_2228 | art_long | art_long | M |
| 77 | 83 | 0.993802522 | cat2 | from_2229 | to_2229 | art_long | art_long | M |
| 80 | 83 | 0.996838175 | cat2 | from_2230 | to_2230 | art_long | art_long | M |
| 81 | 83 | 0.893334087 | cat2 | from_2231 | to_2231 | art_long | art_long | M |
| 8 | 84 | 0.735455943 | cat2 | from_2232 | to_2232 | art_long | art_long | M |
| 9 | 84 | 0.64658614 | cat2 | from_2233 | to_2233 | art_long | art_long | M |
| 10 | 84 | 0.643997947 | cat2 | from_2234 | to_2234 | art_long | art_long | M |
| 49 | 84 | 0.883400103 | cat2 | from_2235 | to_2235 | art_long | art_long | M |
| 70 | 84 | 0.719797922 | cat2 | from_2236 | to_2236 | art_long | art_long | M |
| 75 | 84 | 0.885614655 | cat2 | from_2237 | to_2237 | art_long | art_long | M |
| 34 | 85 | 0.701935151 | cat2 | from_2238 | to_2238 | art_long | art_long | M |
| 49 | 85 | 0.833648641 | cat2 | from_2239 | to_2239 | art_long | art_long | M |
| 56 | 85 | 0.818764755 | cat2 | from_2240 | to_2240 | art_long | art_long | M |
| 57 | 85 | 0.819353386 | cat2 | from_2241 | to_2241 | art_long | art_long | M |
| 70 | 85 | 0.952049132 | cat2 | from_2242 | to_2242 | art_long | art_long | M |
| 75 | 85 | 0.833487457 | cat2 | from_2243 | to_2243 | art_long | art_long | M |
| 32 | 86 | 0.747853117 | cat2 | from_2244 | to_2244 | art_long | art_long | M |
| 58 | 86 | 0.634482264 | cat2 | from_2245 | to_2245 | art_long | art_long | M |
| 61 | 86 | 0.632637935 | cat2 | from_2246 | to_2246 | art_long | art_long | M |
| 69 | 86 | 0.631971559 | cat2 | from_2247 | to_2247 | art_long | art_long | M |
| 72 | 86 | 0.639107083 | cat2 | from_2248 | to_2248 | art_long | art_long | M |
| 14 | 87 | 0.668907235 | cat2 | from_2249 | to_2249 | art_long | art_long | M |
| 24 | 87 | 0.665872911 | cat2 | from_2250 | to_2250 | art_long | art_long | M |
| 27 | 87 | 0.65957065 | cat2 | from_2251 | to_2251 | art_long | art_long | M |
| 28 | 87 | 0.665219649 | cat2 | from_2252 | to_2252 | art_long | art_long | M |
| 30 | 87 | 0.957335548 | cat2 | from_2253 | to_2253 | art_long | art_long | M |
| 47 | 87 | 0.99566391 | cat2 | from_2254 | to_2254 | art_long | art_long | M |
| 54 | 87 | 0.72786277 | cat2 | from_2255 | to_2255 | art_long | art_long | M |
| 55 | 87 | 0.996229485 | cat2 | from_2256 | to_2256 | art_long | art_long | M |
| 58 | 87 | 0.99991125 | cat2 | from_2257 | to_2257 | art_long | art_long | M |
| 59 | 87 | 0.665289232 | cat2 | from_2258 | to_2258 | art_long | art_long | M |
| 69 | 87 | 0.999881345 | cat2 | from_2259 | to_2259 | art_long | art_long | M |
| 72 | 87 | 0.921402422 | cat2 | from_2260 | to_2260 | art_long | art_long | M |
| 73 | 87 | 0.667003525 | cat2 | from_2261 | to_2261 | art_long | art_long | M |
| 74 | 87 | 0.666349533 | cat2 | from_2262 | to_2262 | art_long | art_long | M |
| 76 | 87 | 0.632412985 | cat2 | from_2263 | to_2263 | art_long | art_long | M |
| 77 | 87 | 0.916216959 | cat2 | from_2264 | to_2264 | art_long | art_long | M |
| 78 | 87 | 0.668507439 | cat2 | from_2265 | to_2265 | art_long | art_long | M |
| 80 | 87 | 0.910109586 | cat2 | from_2266 | to_2266 | art_long | art_long | M |
| 81 | 87 | 0.671862185 | cat2 | from_2267 | to_2267 | art_long | art_long | M |
| 83 | 87 | 0.889180065 | cat2 | from_2268 | to_2268 | art_long | art_long | M |
| 86 | 87 | 0.63272624 | cat2 | from_2269 | to_2269 | art_long | art_long | M |
| 9 | 88 | 0.869838141 | cat2 | from_2270 | to_2270 | art_long | art_long | M |
| 10 | 88 | 0.874725836 | cat2 | from_2271 | to_2271 | art_long | art_long | M |
| 22 | 88 | 0.909393968 | cat2 | from_2272 | to_2272 | art_long | art_long | M |
| 23 | 88 | 0.938662111 | cat2 | from_2273 | to_2273 | art_long | art_long | M |
| 65 | 88 | 0.684235033 | cat2 | from_2274 | to_2274 | art_long | art_long | M |
| 14 | 90 | 0.86459322 | cat2 | from_2275 | to_2275 | art_long | art_long | M |
| 24 | 90 | 0.869036604 | cat2 | from_2276 | to_2276 | art_long | art_long | M |
| 25 | 90 | 0.94785226 | cat2 | from_2277 | to_2277 | art_long | art_long | M |
| 28 | 90 | 0.873564679 | cat2 | from_2278 | to_2278 | art_long | art_long | M |
| 31 | 90 | 0.898331105 | cat2 | from_2279 | to_2279 | art_long | art_long | M |
| 33 | 90 | 0.911272109 | cat2 | from_2280 | to_2280 | art_long | art_long | M |
| 59 | 90 | 0.871556717 | cat2 | from_2281 | to_2281 | art_long | art_long | M |
| 73 | 90 | 0.867789901 | cat2 | from_2282 | to_2282 | art_long | art_long | M |
| 74 | 90 | 0.868552117 | cat2 | from_2283 | to_2283 | art_long | art_long | M |
| 76 | 90 | 0.955169444 | cat2 | from_2284 | to_2284 | art_long | art_long | M |
| 78 | 90 | 0.841159623 | cat2 | from_2285 | to_2285 | art_long | art_long | M |
| 22 | 91 | 0.686939526 | cat2 | from_2286 | to_2286 | art_long | art_long | M |

| | | | | | | | | |
|---|---|---|---|---|---|---|---|---|
| 23 | 91 | 0.699437602 | cat2 | from_2287 | to_2287 | art_long | art_long | M |
| 27 | 91 | 0.957109426 | cat2 | from_2288 | to_2288 | art_long | art_long | M |
| 64 | 91 | 0.723183155 | cat2 | from_2289 | to_2289 | art_long | art_long | M |
| 88 | 91 | 0.640117712 | cat2 | from_2290 | to_2290 | art_long | art_long | M |
| 4 | 92 | 0.654947343 | cat2 | from_2291 | to_2291 | art_long | art_long | M |
| 7 | 92 | 0.653865556 | cat2 | from_2292 | to_2292 | art_long | art_long | M |
| 17 | 92 | 0.66370337 | cat2 | from_2293 | to_2293 | art_long | art_long | M |
| 36 | 92 | 0.66419134 | cat2 | from_2294 | to_2294 | art_long | art_long | M |
| 42 | 92 | 0.970592474 | cat2 | from_2295 | to_2295 | art_long | art_long | M |
| 43 | 92 | 0.931166485 | cat2 | from_2296 | to_2296 | art_long | art_long | M |
| 45 | 92 | 0.999546629 | cat2 | from_2297 | to_2297 | art_long | art_long | M |
| 46 | 92 | 0.644348414 | cat2 | from_2298 | to_2298 | art_long | art_long | M |
| 60 | 92 | 0.632333119 | cat2 | from_2299 | to_2299 | art_long | art_long | M |
| 62 | 92 | 0.982426393 | cat2 | from_2300 | to_2300 | art_long | art_long | M |
| 8 | 93 | 0.684619672 | cat2 | from_2301 | to_2301 | art_long | art_long | M |
| 32 | 94 | 0.744098474 | cat2 | from_2302 | to_2302 | art_long | art_long | M |
| 58 | 94 | 0.638585871 | cat2 | from_2303 | to_2303 | art_long | art_long | M |
| 61 | 94 | 0.634257274 | cat2 | from_2304 | to_2304 | art_long | art_long | M |
| 69 | 94 | 0.636852445 | cat2 | from_2305 | to_2305 | art_long | art_long | M |
| 72 | 94 | 0.644391352 | cat2 | from_2306 | to_2306 | art_long | art_long | M |
| 77 | 94 | 0.63313233 | cat2 | from_2307 | to_2307 | art_long | art_long | M |
| 86 | 94 | 0.999027714 | cat2 | from_2308 | to_2308 | art_long | art_long | M |
| 87 | 94 | 0.637413365 | cat2 | from_2309 | to_2309 | art_long | art_long | M |
| 14 | 95 | 0.666784465 | cat2 | from_2310 | to_2310 | art_long | art_long | M |
| 24 | 95 | 0.661726542 | cat2 | from_2311 | to_2311 | art_long | art_long | M |
| 27 | 95 | 0.656892332 | cat2 | from_2312 | to_2312 | art_long | art_long | M |
| 28 | 95 | 0.661060844 | cat2 | from_2313 | to_2313 | art_long | art_long | M |
| 30 | 95 | 0.955422652 | cat2 | from_2314 | to_2314 | art_long | art_long | M |
| 47 | 95 | 0.998848455 | cat2 | from_2315 | to_2315 | art_long | art_long | M |
| 54 | 95 | 0.722312574 | cat2 | from_2316 | to_2316 | art_long | art_long | M |
| 55 | 95 | 0.999130543 | cat2 | from_2317 | to_2317 | art_long | art_long | M |
| 58 | 95 | 0.998290173 | cat2 | from_2318 | to_2318 | art_long | art_long | M |
| 59 | 95 | 0.660989619 | cat2 | from_2319 | to_2319 | art_long | art_long | M |
| 69 | 95 | 0.999062498 | cat2 | from_2320 | to_2320 | art_long | art_long | M |
| 72 | 95 | 0.921249661 | cat2 | from_2321 | to_2321 | art_long | art_long | M |
| 73 | 95 | 0.663511773 | cat2 | from_2322 | to_2322 | art_long | art_long | M |
| 74 | 95 | 0.662469142 | cat2 | from_2323 | to_2323 | art_long | art_long | M |
| 76 | 95 | 0.633827792 | cat2 | from_2324 | to_2324 | art_long | art_long | M |
| 77 | 95 | 0.919710989 | cat2 | from_2325 | to_2325 | art_long | art_long | M |
| 78 | 95 | 0.673232159 | cat2 | from_2326 | to_2326 | art_long | art_long | M |
| 80 | 95 | 0.914901554 | cat2 | from_2327 | to_2327 | art_long | art_long | M |
| 81 | 95 | 0.667263812 | cat2 | from_2328 | to_2328 | art_long | art_long | M |
| 83 | 95 | 0.896161208 | cat2 | from_2329 | to_2329 | art_long | art_long | M |
| 87 | 95 | 0.99898032 | cat2 | from_2330 | to_2330 | art_long | art_long | M |
| 94 | 95 | 0.632598317 | cat2 | from_2331 | to_2331 | art_long | art_long | M |
| 29 | 96 | 0.773393717 | cat2 | from_2332 | to_2332 | art_long | art_long | M |
| 41 | 96 | 0.975638748 | cat2 | from_2333 | to_2333 | art_long | art_long | M |
| 71 | 96 | 0.81978811 | cat2 | from_2334 | to_2334 | art_long | art_long | M |
| 82 | 96 | 0.710721861 | cat2 | from_2335 | to_2335 | art_long | art_long | M |
| 14 | 97 | 0.999382651 | cat2 | from_2336 | to_2336 | art_long | art_long | M |
| 24 | 97 | 0.999951392 | cat2 | from_2337 | to_2337 | art_long | art_long | M |
| 25 | 97 | 0.741608444 | cat2 | from_2338 | to_2338 | art_long | art_long | M |
| 26 | 97 | 0.645126829 | cat2 | from_2339 | to_2339 | art_long | art_long | M |
| 28 | 97 | 0.999861694 | cat2 | from_2340 | to_2340 | art_long | art_long | M |
| 32 | 97 | 0.638094649 | cat2 | from_2341 | to_2341 | art_long | art_long | M |
| 47 | 97 | 0.657532397 | cat2 | from_2342 | to_2342 | art_long | art_long | M |
| 55 | 97 | 0.658329129 | cat2 | from_2343 | to_2343 | art_long | art_long | M |
| 58 | 97 | 0.66748647 | cat2 | from_2344 | to_2344 | art_long | art_long | M |
| 59 | 97 | 0.999890303 | cat2 | from_2345 | to_2345 | art_long | art_long | M |
| 64 | 97 | 0.66547755 | cat2 | from_2346 | to_2346 | art_long | art_long | M |
| 69 | 97 | 0.663765651 | cat2 | from_2347 | to_2347 | art_long | art_long | M |
| 73 | 97 | 0.999988893 | cat2 | from_2348 | to_2348 | art_long | art_long | M |
| 74 | 97 | 0.999992252 | cat2 | from_2349 | to_2349 | art_long | art_long | M |
| 76 | 97 | 0.926355594 | cat2 | from_2350 | to_2350 | art_long | art_long | M |
| 78 | 97 | 0.982445422 | cat2 | from_2351 | to_2351 | art_long | art_long | M |
| 87 | 97 | 0.666653347 | cat2 | from_2352 | to_2352 | art_long | art_long | M |
| 90 | 97 | 0.86821329 | cat2 | from_2353 | to_2353 | art_long | art_long | M |
| 95 | 97 | 0.662949789 | cat2 | from_2354 | to_2354 | art_long | art_long | M |
| 29 | 98 | 0.669282433 | cat2 | from_2355 | to_2355 | art_long | art_long | M |
| 41 | 98 | 0.885588908 | cat2 | from_2356 | to_2356 | art_long | art_long | M |
| 71 | 98 | 0.690287645 | cat2 | from_2357 | to_2357 | art_long | art_long | M |
| 82 | 98 | 0.789915955 | cat2 | from_2358 | to_2358 | art_long | art_long | M |
| 96 | 98 | 0.870435795 | cat2 | from_2359 | to_2359 | art_long | art_long | M |
| 44 | 99 | 0.9193764 | cat2 | from_2360 | to_2360 | art_long | art_long | M |
| 46 | 99 | 0.916439191 | cat2 | from_2361 | to_2361 | art_long | art_long | M |
| 52 | 99 | 0.729160907 | cat2 | from_2362 | to_2362 | art_long | art_long | M |
| 60 | 99 | 0.921574694 | cat2 | from_2363 | to_2363 | art_long | art_long | M |
| 61 | 99 | 0.982306933 | cat2 | from_2364 | to_2364 | art_long | art_long | M |
| 4 | 100 | 0.639584037 | cat2 | from_2365 | to_2365 | art_long | art_long | M |
| 7 | 100 | 0.641579895 | cat2 | from_2366 | to_2366 | art_long | art_long | M |
| 17 | 100 | 0.651196936 | cat2 | from_2367 | to_2367 | art_long | art_long | M |
| 36 | 100 | 0.65284269 | cat2 | from_2368 | to_2368 | art_long | art_long | M |
| 42 | 100 | 0.99661167 | cat2 | from_2369 | to_2369 | art_long | art_long | M |
| 43 | 100 | 0.91710359 | cat2 | from_2370 | to_2370 | art_long | art_long | M |
| 44 | 100 | 0.659411718 | cat2 | from_2371 | to_2371 | art_long | art_long | M |
| 45 | 100 | 0.991110869 | cat2 | from_2372 | to_2372 | art_long | art_long | M |
| 46 | 100 | 0.65858397 | cat2 | from_2373 | to_2373 | art_long | art_long | M |
| 60 | 100 | 0.637317938 | cat2 | from_2374 | to_2374 | art_long | art_long | M |
| 62 | 100 | 0.999537906 | cat2 | from_2375 | to_2375 | art_long | art_long | M |
| 92 | 100 | 0.987073267 | cat2 | from_2376 | to_2376 | art_long | art_long | M |
| 4 | 101 | 0.632316 | cat2 | from_2377 | to_2377 | art_long | art_long | M |
| 7 | 101 | 0.635144497 | cat2 | from_2378 | to_2378 | art_long | art_long | M |
| 17 | 101 | 0.644614375 | cat2 | from_2379 | to_2379 | art_long | art_long | M |
| 36 | 101 | 0.646538682 | cat2 | from_2380 | to_2380 | art_long | art_long | M |
| 42 | 101 | 0.999181186 | cat2 | from_2381 | to_2381 | art_long | art_long | M |
| 43 | 101 | 0.909968344 | cat2 | from_2382 | to_2382 | art_long | art_long | M |
| 44 | 101 | 0.66475274 | cat2 | from_2383 | to_2383 | art_long | art_long | M |
| 45 | 101 | 0.984672958 | cat2 | from_2384 | to_2384 | art_long | art_long | M |
| 46 | 101 | 0.659328744 | cat2 | from_2385 | to_2385 | art_long | art_long | M |
| 60 | 101 | 0.63602704 | cat2 | from_2386 | to_2386 | art_long | art_long | M |
| 62 | 101 | 0.999806694 | cat2 | from_2387 | to_2387 | art_long | art_long | M |
| 92 | 101 | 0.979522396 | cat2 | from_2388 | to_2388 | art_long | art_long | M |
| 100 | 101 | 0.999123439 | cat2 | from_2389 | to_2389 | art_long | art_long | M |
| 42 | 102 | 0.663919099 | cat2 | from_2390 | to_2390 | art_long | art_long | M |

| | | | | | | | | |
|---|---|---|---|---|---|---|---|---|
| 44 | 102 | 0.996881794 | cat2 | from_2391 | to_2391 | art_long | art_long | M |
| 46 | 102 | 0.983086087 | cat2 | from_2392 | to_2392 | art_long | art_long | M |
| 52 | 102 | 0.867057992 | cat2 | from_2393 | to_2393 | art_long | art_long | M |
| 60 | 102 | 0.97209536 | cat2 | from_2394 | to_2394 | art_long | art_long | M |
| 61 | 102 | 0.879839779 | cat2 | from_2395 | to_2395 | art_long | art_long | M |
| 62 | 102 | 0.65409168 | cat2 | from_2396 | to_2396 | art_long | art_long | M |
| 99 | 102 | 0.920620248 | cat2 | from_2397 | to_2397 | art_long | art_long | M |
| 100 | 102 | 0.648812942 | cat2 | from_2398 | to_2398 | art_long | art_long | M |
| 101 | 102 | 0.657051289 | cat2 | from_2399 | to_2399 | art_long | art_long | M |
| 51 | 103 | 0.838596464 | cat2 | from_2400 | to_2400 | art_long | art_long | M |
| 17 | 104 | 0.792443044 | cat2 | from_2401 | to_2401 | art_long | art_long | M |
| 35 | 104 | 0.999331397 | cat2 | from_2402 | to_2402 | art_long | art_long | M |
| 36 | 104 | 0.792541654 | cat2 | from_2403 | to_2403 | art_long | art_long | M |
| 38 | 104 | 0.999346126 | cat2 | from_2404 | to_2404 | art_long | art_long | M |
| 44 | 104 | 0.633155604 | cat2 | from_2405 | to_2405 | art_long | art_long | M |
| 46 | 104 | 0.648854759 | cat2 | from_2406 | to_2406 | art_long | art_long | M |
| 48 | 104 | 0.99936307 | cat2 | from_2407 | to_2407 | art_long | art_long | M |
| 50 | 104 | 0.8110707 | cat2 | from_2408 | to_2408 | art_long | art_long | M |
| 60 | 104 | 0.640198279 | cat2 | from_2409 | to_2409 | art_long | art_long | M |
| 44 | 105 | 0.728137326 | cat2 | from_2410 | to_2410 | art_long | art_long | M |
| 46 | 105 | 0.726667626 | cat2 | from_2411 | to_2411 | art_long | art_long | M |
| 50 | 105 | 0.65535061 | cat2 | from_2412 | to_2412 | art_long | art_long | M |
| 60 | 105 | 0.744336119 | cat2 | from_2413 | to_2413 | art_long | art_long | M |
| 61 | 105 | 0.93010304 | cat2 | from_2414 | to_2414 | art_long | art_long | M |
| 99 | 105 | 0.938998394 | cat2 | from_2415 | to_2415 | art_long | art_long | M |
| 102 | 105 | 0.734112774 | cat2 | from_2416 | to_2416 | art_long | art_long | M |
| 42 | 106 | 0.667433747 | cat2 | from_2417 | to_2417 | art_long | art_long | M |
| 44 | 106 | 0.984083521 | cat2 | from_2418 | to_2418 | art_long | art_long | M |
| 46 | 106 | 0.958428387 | cat2 | from_2419 | to_2419 | art_long | art_long | M |
| 52 | 106 | 0.831127489 | cat2 | from_2420 | to_2420 | art_long | art_long | M |
| 60 | 106 | 0.93888348 | cat2 | from_2421 | to_2421 | art_long | art_long | M |
| 61 | 106 | 0.839536195 | cat2 | from_2422 | to_2422 | art_long | art_long | M |
| 62 | 106 | 0.651433158 | cat2 | from_2423 | to_2423 | art_long | art_long | M |
| 99 | 106 | 0.900415791 | cat2 | from_2424 | to_2424 | art_long | art_long | M |
| 100 | 106 | 0.64375516 | cat2 | from_2425 | to_2425 | art_long | art_long | M |
| 101 | 106 | 0.656352117 | cat2 | from_2426 | to_2426 | art_long | art_long | M |
| 102 | 106 | 0.993391459 | cat2 | from_2427 | to_2427 | art_long | art_long | M |
| 105 | 106 | 0.711742871 | cat2 | from_2428 | to_2428 | art_long | art_long | M |
| 4 | 107 | 0.655780047 | cat2 | from_2429 | to_2429 | art_long | art_long | M |
| 7 | 107 | 0.654947471 | cat2 | from_2430 | to_2430 | art_long | art_long | M |
| 17 | 107 | 0.66493554 | cat2 | from_2431 | to_2431 | art_long | art_long | M |
| 36 | 107 | 0.665622418 | cat2 | from_2432 | to_2432 | art_long | art_long | M |
| 42 | 107 | 0.975079474 | cat2 | from_2433 | to_2433 | art_long | art_long | M |
| 43 | 107 | 0.929265111 | cat2 | from_2434 | to_2434 | art_long | art_long | M |
| 44 | 107 | 0.632909739 | cat2 | from_2435 | to_2435 | art_long | art_long | M |
| 45 | 107 | 0.999968482 | cat2 | from_2436 | to_2436 | art_long | art_long | M |
| 46 | 107 | 0.64756351 | cat2 | from_2437 | to_2437 | art_long | art_long | M |
| 60 | 107 | 0.633424391 | cat2 | from_2438 | to_2438 | art_long | art_long | M |
| 62 | 107 | 0.985712058 | cat2 | from_2439 | to_2439 | art_long | art_long | M |
| 92 | 107 | 0.999729264 | cat2 | from_2440 | to_2440 | art_long | art_long | M |
| 100 | 107 | 0.990023384 | cat2 | from_2441 | to_2441 | art_long | art_long | M |
| 101 | 107 | 0.983257204 | cat2 | from_2442 | to_2442 | art_long | art_long | M |
| 37 | 108 | 0.665851984 | cat2 | from_2443 | to_2443 | art_long | art_long | M |
| 49 | 108 | 0.646601876 | cat2 | from_2444 | to_2444 | art_long | art_long | M |
| 70 | 108 | 0.640940141 | cat2 | from_2445 | to_2445 | art_long | art_long | M |
| 75 | 108 | 0.646610629 | cat2 | from_2446 | to_2446 | art_long | art_long | M |
| 14 | 109 | 0.661959762 | cat2 | from_2447 | to_2447 | art_long | art_long | M |
| 24 | 109 | 0.655513317 | cat2 | from_2448 | to_2448 | art_long | art_long | M |
| 27 | 109 | 0.652584696 | cat2 | from_2449 | to_2449 | art_long | art_long | M |
| 28 | 109 | 0.655025491 | cat2 | from_2450 | to_2450 | art_long | art_long | M |
| 30 | 109 | 0.957084475 | cat2 | from_2451 | to_2451 | art_long | art_long | M |
| 47 | 109 | 0.999751303 | cat2 | from_2452 | to_2452 | art_long | art_long | M |
| 54 | 109 | 0.721510058 | cat2 | from_2453 | to_2453 | art_long | art_long | M |
| 55 | 109 | 0.999837323 | cat2 | from_2454 | to_2454 | art_long | art_long | M |
| 58 | 109 | 0.995876257 | cat2 | from_2455 | to_2455 | art_long | art_long | M |
| 59 | 109 | 0.654763941 | cat2 | from_2456 | to_2456 | art_long | art_long | M |
| 69 | 109 | 0.997417408 | cat2 | from_2457 | to_2457 | art_long | art_long | M |
| 72 | 109 | 0.922222125 | cat2 | from_2458 | to_2458 | art_long | art_long | M |
| 73 | 109 | 0.657746975 | cat2 | from_2459 | to_2459 | art_long | art_long | M |
| 74 | 109 | 0.656438031 | cat2 | from_2460 | to_2460 | art_long | art_long | M |
| 76 | 109 | 0.633543692 | cat2 | from_2461 | to_2461 | art_long | art_long | M |
| 77 | 109 | 0.923306065 | cat2 | from_2462 | to_2462 | art_long | art_long | M |
| 78 | 109 | 0.673340512 | cat2 | from_2463 | to_2463 | art_long | art_long | M |
| 80 | 109 | 0.919636716 | cat2 | from_2464 | to_2464 | art_long | art_long | M |
| 81 | 109 | 0.669282741 | cat2 | from_2465 | to_2465 | art_long | art_long | M |
| 83 | 109 | 0.903307545 | cat2 | from_2466 | to_2466 | art_long | art_long | M |
| 87 | 109 | 0.996980675 | cat2 | from_2467 | to_2467 | art_long | art_long | M |
| 95 | 109 | 0.999406872 | cat2 | from_2468 | to_2468 | art_long | art_long | M |
| 97 | 109 | 0.657039822 | cat2 | from_2469 | to_2469 | art_long | art_long | M |
| 14 | 110 | 0.964148278 | cat2 | from_2470 | to_2470 | art_long | art_long | M |
| 24 | 110 | 0.965074526 | cat2 | from_2471 | to_2471 | art_long | art_long | M |
| 25 | 110 | 0.780944716 | cat2 | from_2472 | to_2472 | art_long | art_long | M |
| 28 | 110 | 0.968434229 | cat2 | from_2473 | to_2473 | art_long | art_long | M |
| 31 | 110 | 0.693770078 | cat2 | from_2474 | to_2474 | art_long | art_long | M |
| 33 | 110 | 0.709307097 | cat2 | from_2475 | to_2475 | art_long | art_long | M |
| 58 | 110 | 0.632287889 | cat2 | from_2476 | to_2476 | art_long | art_long | M |
| 59 | 110 | 0.966753232 | cat2 | from_2477 | to_2477 | art_long | art_long | M |
| 69 | 110 | 0.63228075 | cat2 | from_2478 | to_2478 | art_long | art_long | M |
| 73 | 110 | 0.96498848 | cat2 | from_2479 | to_2479 | art_long | art_long | M |
| 74 | 110 | 0.965064274 | cat2 | from_2480 | to_2480 | art_long | art_long | M |
| 76 | 110 | 0.927024722 | cat2 | from_2481 | to_2481 | art_long | art_long | M |
| 78 | 110 | 0.949940295 | cat2 | from_2482 | to_2482 | art_long | art_long | M |
| 90 | 110 | 0.929817451 | cat2 | from_2483 | to_2483 | art_long | art_long | M |
| 97 | 110 | 0.965038733 | cat2 | from_2484 | to_2484 | art_long | art_long | M |
| 14 | 111 | 0.666642754 | cat2 | from_2485 | to_2485 | art_long | art_long | M |
| 24 | 111 | 0.662493251 | cat2 | from_2486 | to_2486 | art_long | art_long | M |
| 27 | 111 | 0.657441031 | cat2 | from_2487 | to_2487 | art_long | art_long | M |
| 28 | 111 | 0.661925641 | cat2 | from_2488 | to_2488 | art_long | art_long | M |
| 30 | 111 | 0.958604679 | cat2 | from_2489 | to_2489 | art_long | art_long | M |
| 47 | 111 | 0.997643906 | cat2 | from_2490 | to_2490 | art_long | art_long | M |
| 54 | 111 | 0.726947531 | cat2 | from_2491 | to_2491 | art_long | art_long | M |
| 55 | 111 | 0.998053347 | cat2 | from_2492 | to_2492 | art_long | art_long | M |
| 58 | 111 | 0.999243287 | cat2 | from_2493 | to_2493 | art_long | art_long | M |
| 59 | 111 | 0.66187109 | cat2 | from_2494 | to_2494 | art_long | art_long | M |

| | | | | | | | | |
|---|---|---|---|---|---|---|---|---|
| 69 | 111 | 0.999763419 | cat2 | from_2495 | to_2495 | art_long | art_long | M |
| 72 | 111 | 0.922660767 | cat2 | from_2496 | to_2496 | art_long | art_long | M |
| 73 | 111 | 0.663984373 | cat2 | from_2497 | to_2497 | art_long | art_long | M |
| 74 | 111 | 0.663116323 | cat2 | from_2498 | to_2498 | art_long | art_long | M |
| 76 | 111 | 0.63311652 | cat2 | from_2499 | to_2499 | art_long | art_long | M |
| 77 | 111 | 0.919535766 | cat2 | from_2500 | to_2500 | art_long | art_long | M |
| 78 | 111 | 0.670124302 | cat2 | from_2501 | to_2501 | art_long | art_long | M |
| 80 | 111 | 0.91426119 | cat2 | from_2502 | to_2502 | art_long | art_long | M |
| 81 | 111 | 0.672503845 | cat2 | from_2503 | to_2503 | art_long | art_long | M |
| 83 | 111 | 0.894970776 | cat2 | from_2504 | to_2504 | art_long | art_long | M |
| 87 | 111 | 0.999663555 | cat2 | from_2505 | to_2505 | art_long | art_long | M |
| 94 | 111 | 0.634865804 | cat2 | from_2506 | to_2506 | art_long | art_long | M |
| 95 | 111 | 0.999767764 | cat2 | from_2507 | to_2507 | art_long | art_long | M |
| 97 | 111 | 0.663517525 | cat2 | from_2508 | to_2508 | art_long | art_long | M |
| 109 | 111 | 0.998650052 | cat2 | from_2509 | to_2509 | art_long | art_long | M |
| 14 | 112 | 0.737888702 | cat2 | from_2510 | to_2510 | art_long | art_long | M |
| 24 | 112 | 0.743146544 | cat2 | from_2511 | to_2511 | art_long | art_long | M |
| 25 | 112 | 0.998905632 | cat2 | from_2512 | to_2512 | art_long | art_long | M |
| 28 | 112 | 0.74624096 | cat2 | from_2513 | to_2513 | art_long | art_long | M |
| 31 | 112 | 0.92536551 | cat2 | from_2514 | to_2514 | art_long | art_long | M |
| 33 | 112 | 0.941544748 | cat2 | from_2515 | to_2515 | art_long | art_long | M |
| 54 | 112 | 0.716840878 | cat2 | from_2516 | to_2516 | art_long | art_long | M |
| 59 | 112 | 0.745021801 | cat2 | from_2517 | to_2517 | art_long | art_long | M |
| 68 | 112 | 0.669815143 | cat2 | from_2518 | to_2518 | art_long | art_long | M |
| 72 | 112 | 0.649962791 | cat2 | from_2519 | to_2519 | art_long | art_long | M |
| 73 | 112 | 0.741608084 | cat2 | from_2520 | to_2520 | art_long | art_long | M |
| 74 | 112 | 0.74254024 | cat2 | from_2521 | to_2521 | art_long | art_long | M |
| 76 | 112 | 0.91447152 | cat2 | from_2522 | to_2522 | art_long | art_long | M |
| 77 | 112 | 0.634028767 | cat2 | from_2523 | to_2523 | art_long | art_long | M |
| 78 | 112 | 0.7121828 | cat2 | from_2524 | to_2524 | art_long | art_long | M |
| 81 | 112 | 0.695155677 | cat2 | from_2525 | to_2525 | art_long | art_long | M |
| 90 | 112 | 0.947957513 | cat2 | from_2526 | to_2526 | art_long | art_long | M |
| 97 | 112 | 0.742122904 | cat2 | from_2527 | to_2527 | art_long | art_long | M |
| 110 | 112 | 0.779372723 | cat2 | from_2528 | to_2528 | art_long | art_long | M |
| 4 | 113 | 0.669845983 | cat2 | from_2529 | to_2529 | art_long | art_long | M |
| 7 | 113 | 0.671519836 | cat2 | from_2530 | to_2530 | art_long | art_long | M |
| 22 | 113 | 0.68876236 | cat2 | from_2531 | to_2531 | art_long | art_long | M |
| 34 | 113 | 0.951700179 | cat2 | from_2532 | to_2532 | art_long | art_long | M |
| 64 | 113 | 0.667151023 | cat2 | from_2533 | to_2533 | art_long | art_long | M |
| 65 | 113 | 0.935899571 | cat2 | from_2534 | to_2534 | art_long | art_long | M |
| 70 | 113 | 0.645193449 | cat2 | from_2535 | to_2535 | art_long | art_long | M |
| 85 | 113 | 0.655579521 | cat2 | from_2536 | to_2536 | art_long | art_long | M |
| 12 | 114 | 0.740208574 | cat2 | from_2537 | to_2537 | art_long | art_long | M |
| 13 | 114 | 0.94476887 | cat2 | from_2538 | to_2538 | art_long | art_long | M |
| 19 | 114 | 0.800322954 | cat2 | from_2539 | to_2539 | art_long | art_long | M |
| 20 | 114 | 0.797438157 | cat2 | from_2540 | to_2540 | art_long | art_long | M |
| 39 | 114 | 0.687637745 | cat2 | from_2541 | to_2541 | art_long | art_long | M |
| 40 | 114 | 0.757947291 | cat2 | from_2542 | to_2542 | art_long | art_long | M |
| 68 | 114 | 0.785257012 | cat2 | from_2543 | to_2543 | art_long | art_long | M |
| 79 | 114 | 0.820110231 | cat2 | from_2544 | to_2544 | art_long | art_long | M |
| 14 | 115 | 0.636655901 | cat2 | from_2545 | to_2545 | art_long | art_long | M |
| 24 | 115 | 0.63461955 | cat2 | from_2546 | to_2546 | art_long | art_long | M |
| 26 | 115 | 0.987810199 | cat2 | from_2547 | to_2547 | art_long | art_long | M |
| 28 | 115 | 0.635341534 | cat2 | from_2548 | to_2548 | art_long | art_long | M |
| 37 | 115 | 0.94880953 | cat2 | from_2549 | to_2549 | art_long | art_long | M |
| 59 | 115 | 0.634792792 | cat2 | from_2550 | to_2550 | art_long | art_long | M |
| 73 | 115 | 0.635419945 | cat2 | from_2551 | to_2551 | art_long | art_long | M |
| 74 | 115 | 0.634961146 | cat2 | from_2552 | to_2552 | art_long | art_long | M |
| 78 | 115 | 0.634729732 | cat2 | from_2553 | to_2553 | art_long | art_long | M |
| 97 | 115 | 0.635175836 | cat2 | from_2554 | to_2554 | art_long | art_long | M |
| 5 | 116 | 0.966722139 | cat2 | from_2555 | to_2555 | art_long | art_long | M |
| 13 | 116 | 0.647884727 | cat2 | from_2556 | to_2556 | art_long | art_long | M |
| 15 | 116 | 0.714410089 | cat2 | from_2557 | to_2557 | art_long | art_long | M |
| 63 | 116 | 0.995712846 | cat2 | from_2558 | to_2558 | art_long | art_long | M |
| 14 | 117 | 0.662357776 | cat2 | from_2559 | to_2559 | art_long | art_long | M |
| 24 | 117 | 0.666408375 | cat2 | from_2560 | to_2560 | art_long | art_long | M |
| 27 | 117 | 0.803756367 | cat2 | from_2561 | to_2561 | art_long | art_long | M |
| 28 | 117 | 0.664433109 | cat2 | from_2562 | to_2562 | art_long | art_long | M |
| 34 | 117 | 0.710998102 | cat2 | from_2563 | to_2563 | art_long | art_long | M |
| 59 | 117 | 0.665684395 | cat2 | from_2564 | to_2564 | art_long | art_long | M |
| 64 | 117 | 0.999918779 | cat2 | from_2565 | to_2565 | art_long | art_long | M |
| 73 | 117 | 0.66524398 | cat2 | from_2566 | to_2566 | art_long | art_long | M |
| 74 | 117 | 0.665952153 | cat2 | from_2567 | to_2567 | art_long | art_long | M |
| 78 | 117 | 0.636843119 | cat2 | from_2568 | to_2568 | art_long | art_long | M |
| 91 | 117 | 0.722387735 | cat2 | from_2569 | to_2569 | art_long | art_long | M |
| 97 | 117 | 0.665636041 | cat2 | from_2570 | to_2570 | art_long | art_long | M |
| 113 | 117 | 0.666228199 | cat2 | from_2571 | to_2571 | art_long | art_long | M |
| 26 | 118 | 0.785400703 | cat2 | from_2572 | to_2572 | art_long | art_long | M |
| 37 | 118 | 0.774369698 | cat2 | from_2573 | to_2573 | art_long | art_long | M |
| 47 | 118 | 0.651113543 | cat2 | from_2574 | to_2574 | art_long | art_long | M |
| 55 | 118 | 0.652058416 | cat2 | from_2575 | to_2575 | art_long | art_long | M |
| 58 | 118 | 0.663604208 | cat2 | from_2576 | to_2576 | art_long | art_long | M |
| 69 | 118 | 0.66030623 | cat2 | from_2577 | to_2577 | art_long | art_long | M |
| 87 | 118 | 0.662450702 | cat2 | from_2578 | to_2578 | art_long | art_long | M |
| 95 | 118 | 0.657666977 | cat2 | from_2579 | to_2579 | art_long | art_long | M |
| 109 | 118 | 0.651861793 | cat2 | from_2580 | to_2580 | art_long | art_long | M |
| 111 | 118 | 0.659139686 | cat2 | from_2581 | to_2581 | art_long | art_long | M |
| 115 | 118 | 0.776064692 | cat2 | from_2582 | to_2582 | art_long | art_long | M |
| 14 | 119 | 0.663369254 | cat2 | from_2583 | to_2583 | art_long | art_long | M |
| 24 | 119 | 0.656975366 | cat2 | from_2584 | to_2584 | art_long | art_long | M |
| 27 | 119 | 0.653560715 | cat2 | from_2585 | to_2585 | art_long | art_long | M |
| 28 | 119 | 0.656394342 | cat2 | from_2586 | to_2586 | art_long | art_long | M |
| 30 | 119 | 0.955149155 | cat2 | from_2587 | to_2587 | art_long | art_long | M |
| 47 | 119 | 0.999810969 | cat2 | from_2588 | to_2588 | art_long | art_long | M |
| 54 | 119 | 0.71976713 | cat2 | from_2589 | to_2589 | art_long | art_long | M |
| 55 | 119 | 0.999904629 | cat2 | from_2590 | to_2590 | art_long | art_long | M |
| 58 | 119 | 0.996064644 | cat2 | from_2591 | to_2591 | art_long | art_long | M |
| 59 | 119 | 0.656183482 | cat2 | from_2592 | to_2592 | art_long | art_long | M |
| 69 | 119 | 0.997475652 | cat2 | from_2593 | to_2593 | art_long | art_long | M |
| 72 | 119 | 0.921215748 | cat2 | from_2594 | to_2594 | art_long | art_long | M |
| 73 | 119 | 0.659192322 | cat2 | from_2595 | to_2595 | art_long | art_long | M |
| 74 | 119 | 0.657893328 | cat2 | from_2596 | to_2596 | art_long | art_long | M |
| 76 | 119 | 0.633818645 | cat2 | from_2597 | to_2597 | art_long | art_long | M |
| 77 | 119 | 0.922149813 | cat2 | from_2598 | to_2598 | art_long | art_long | M |

| | | | | | | | | |
|---|---|---|---|---|---|---|---|---|
| 78 | 119 | 0.674459495 | cat2 | from_2599 | to_2599 | art_long | art_long | M |
| 80 | 119 | 0.918325242 | cat2 | from_2600 | to_2600 | art_long | art_long | M |
| 81 | 119 | 0.666452632 | cat2 | from_2601 | to_2601 | art_long | art_long | M |
| 83 | 119 | 0.901493864 | cat2 | from_2602 | to_2602 | art_long | art_long | M |
| 87 | 119 | 0.997153043 | cat2 | from_2603 | to_2603 | art_long | art_long | M |
| 95 | 119 | 0.999524729 | cat2 | from_2604 | to_2604 | art_long | art_long | M |
| 97 | 119 | 0.658490604 | cat2 | from_2605 | to_2605 | art_long | art_long | M |
| 109 | 119 | 0.999972076 | cat2 | from_2606 | to_2606 | art_long | art_long | M |
| 111 | 119 | 0.998738496 | cat2 | from_2607 | to_2607 | art_long | art_long | M |
| 118 | 119 | 0.652916056 | cat2 | from_2608 | to_2608 | art_long | art_long | M |
| 25 | 120 | 0.660713796 | cat2 | from_2609 | to_2609 | art_long | art_long | M |
| 30 | 120 | 0.927882156 | cat2 | from_2610 | to_2610 | art_long | art_long | M |
| 47 | 120 | 0.913233908 | cat2 | from_2611 | to_2611 | art_long | art_long | M |
| 54 | 120 | 0.937612718 | cat2 | from_2612 | to_2612 | art_long | art_long | M |
| 55 | 120 | 0.913898027 | cat2 | from_2613 | to_2613 | art_long | art_long | M |
| 58 | 120 | 0.919585153 | cat2 | from_2614 | to_2614 | art_long | art_long | M |
| 69 | 120 | 0.921567662 | cat2 | from_2615 | to_2615 | art_long | art_long | M |
| 72 | 120 | 0.999141222 | cat2 | from_2616 | to_2616 | art_long | art_long | M |
| 76 | 120 | 0.64587849 | cat2 | from_2617 | to_2617 | art_long | art_long | M |
| 77 | 120 | 0.99265483 | cat2 | from_2618 | to_2618 | art_long | art_long | M |
| 80 | 120 | 0.988635161 | cat2 | from_2619 | to_2619 | art_long | art_long | M |
| 81 | 120 | 0.904579774 | cat2 | from_2620 | to_2620 | art_long | art_long | M |
| 83 | 120 | 0.977214733 | cat2 | from_2621 | to_2621 | art_long | art_long | M |
| 86 | 120 | 0.643437952 | cat2 | from_2622 | to_2622 | art_long | art_long | M |
| 87 | 120 | 0.919360269 | cat2 | from_2623 | to_2623 | art_long | art_long | M |
| 94 | 120 | 0.64694351 | cat2 | from_2624 | to_2624 | art_long | art_long | M |
| 95 | 120 | 0.917384228 | cat2 | from_2625 | to_2625 | art_long | art_long | M |
| 109 | 120 | 0.917232146 | cat2 | from_2626 | to_2626 | art_long | art_long | M |
| 111 | 120 | 0.919677597 | cat2 | from_2627 | to_2627 | art_long | art_long | M |
| 112 | 120 | 0.656778827 | cat2 | from_2628 | to_2628 | art_long | art_long | M |
| 119 | 120 | 0.916209075 | cat2 | from_2629 | to_2629 | art_long | art_long | M |
| 29 | 121 | 0.671893465 | cat2 | from_2630 | to_2630 | art_long | art_long | M |
| 41 | 121 | 0.891990179 | cat2 | from_2631 | to_2631 | art_long | art_long | M |
| 71 | 121 | 0.691676158 | cat2 | from_2632 | to_2632 | art_long | art_long | M |
| 82 | 121 | 0.782978624 | cat2 | from_2633 | to_2633 | art_long | art_long | M |
| 96 | 121 | 0.881527398 | cat2 | from_2634 | to_2634 | art_long | art_long | M |
| 98 | 121 | 0.996091708 | cat2 | from_2635 | to_2635 | art_long | art_long | M |
| 14 | 122 | 0.999596384 | cat2 | from_2636 | to_2636 | art_long | art_long | M |
| 24 | 122 | 0.999862437 | cat2 | from_2637 | to_2637 | art_long | art_long | M |
| 25 | 122 | 0.741176217 | cat2 | from_2638 | to_2638 | art_long | art_long | M |
| 26 | 122 | 0.644459166 | cat2 | from_2639 | to_2639 | art_long | art_long | M |
| 28 | 122 | 0.999770526 | cat2 | from_2640 | to_2640 | art_long | art_long | M |
| 32 | 122 | 0.637242094 | cat2 | from_2641 | to_2641 | art_long | art_long | M |
| 47 | 122 | 0.658649522 | cat2 | from_2642 | to_2642 | art_long | art_long | M |
| 55 | 122 | 0.659404501 | cat2 | from_2643 | to_2643 | art_long | art_long | M |
| 58 | 122 | 0.667892355 | cat2 | from_2644 | to_2644 | art_long | art_long | M |
| 59 | 122 | 0.99977822 | cat2 | from_2645 | to_2645 | art_long | art_long | M |
| 64 | 122 | 0.664991624 | cat2 | from_2646 | to_2646 | art_long | art_long | M |
| 69 | 122 | 0.664290267 | cat2 | from_2647 | to_2647 | art_long | art_long | M |
| 73 | 122 | 0.999997972 | cat2 | from_2648 | to_2648 | art_long | art_long | M |
| 74 | 122 | 0.999943143 | cat2 | from_2649 | to_2649 | art_long | art_long | M |
| 76 | 122 | 0.927063698 | cat2 | from_2650 | to_2650 | art_long | art_long | M |
| 78 | 122 | 0.98367511 | cat2 | from_2651 | to_2651 | art_long | art_long | M |
| 87 | 122 | 0.667148641 | cat2 | from_2652 | to_2652 | art_long | art_long | M |
| 90 | 122 | 0.867603105 | cat2 | from_2653 | to_2653 | art_long | art_long | M |
| 95 | 122 | 0.663747417 | cat2 | from_2654 | to_2654 | art_long | art_long | M |
| 97 | 122 | 0.999977372 | cat2 | from_2655 | to_2655 | art_long | art_long | M |
| 109 | 122 | 0.658044691 | cat2 | from_2656 | to_2656 | art_long | art_long | M |
| 110 | 122 | 0.964960469 | cat2 | from_2657 | to_2657 | art_long | art_long | M |
| 111 | 122 | 0.664179363 | cat2 | from_2658 | to_2658 | art_long | art_long | M |
| 112 | 122 | 0.741383074 | cat2 | from_2659 | to_2659 | art_long | art_long | M |
| 115 | 122 | 0.63551995 | cat2 | from_2660 | to_2660 | art_long | art_long | M |
| 117 | 122 | 0.665071942 | cat2 | from_2661 | to_2661 | art_long | art_long | M |
| 119 | 122 | 0.659487706 | cat2 | from_2662 | to_2662 | art_long | art_long | M |
| 34 | 123 | 0.700718163 | cat2 | from_2663 | to_2663 | art_long | art_long | M |
| 49 | 123 | 0.835634538 | cat2 | from_2664 | to_2664 | art_long | art_long | M |
| 56 | 123 | 0.816352012 | cat2 | from_2665 | to_2665 | art_long | art_long | M |
| 57 | 123 | 0.817318263 | cat2 | from_2666 | to_2666 | art_long | art_long | M |
| 70 | 123 | 0.956295667 | cat2 | from_2667 | to_2667 | art_long | art_long | M |
| 75 | 123 | 0.836750915 | cat2 | from_2668 | to_2668 | art_long | art_long | M |
| 85 | 123 | 0.999347061 | cat2 | from_2669 | to_2669 | art_long | art_long | M |
| 113 | 123 | 0.656436305 | cat2 | from_2670 | to_2670 | art_long | art_long | M |
| 29 | 124 | 0.82739818 | cat2 | from_2671 | to_2671 | art_long | art_long | M |
| 41 | 124 | 0.667781288 | cat2 | from_2672 | to_2672 | art_long | art_long | M |
| 66 | 124 | 0.676585422 | cat2 | from_2673 | to_2673 | art_long | art_long | M |
| 67 | 124 | 0.678022847 | cat2 | from_2674 | to_2674 | art_long | art_long | M |
| 71 | 124 | 0.845334825 | cat2 | from_2675 | to_2675 | art_long | art_long | M |
| 82 | 124 | 0.669129041 | cat2 | from_2676 | to_2676 | art_long | art_long | M |
| 6 | 125 | 0.779582514 | cat2 | from_2677 | to_2677 | art_long | art_long | M |
| 8 | 125 | 0.812264182 | cat2 | from_2678 | to_2678 | art_long | art_long | M |
| 9 | 125 | 0.785127979 | cat2 | from_2679 | to_2679 | art_long | art_long | M |
| 10 | 125 | 0.783367174 | cat2 | from_2680 | to_2680 | art_long | art_long | M |
| 23 | 125 | 0.663617104 | cat2 | from_2681 | to_2681 | art_long | art_long | M |
| 49 | 125 | 0.656771942 | cat2 | from_2682 | to_2682 | art_long | art_long | M |
| 65 | 125 | 0.807380294 | cat2 | from_2683 | to_2683 | art_long | art_long | M |
| 75 | 125 | 0.658290956 | cat2 | from_2684 | to_2684 | art_long | art_long | M |
| 84 | 125 | 0.801917276 | cat2 | from_2685 | to_2685 | art_long | art_long | M |
| 113 | 125 | 0.651993193 | cat2 | from_2686 | to_2686 | art_long | art_long | M |
| 8 | 126 | 0.689702202 | cat2 | from_2687 | to_2687 | art_long | art_long | M |
| 9 | 126 | 0.928864608 | cat2 | from_2688 | to_2688 | art_long | art_long | M |
| 10 | 126 | 0.930739807 | cat2 | from_2689 | to_2689 | art_long | art_long | M |
| 22 | 126 | 0.937061738 | cat2 | from_2690 | to_2690 | art_long | art_long | M |
| 23 | 126 | 0.988685622 | cat2 | from_2691 | to_2691 | art_long | art_long | M |
| 65 | 126 | 0.71037969 | cat2 | from_2692 | to_2692 | art_long | art_long | M |
| 88 | 126 | 0.979766084 | cat2 | from_2693 | to_2693 | art_long | art_long | M |
| 91 | 126 | 0.684538153 | cat2 | from_2694 | to_2694 | art_long | art_long | M |
| 4 | 127 | 0.732095007 | cat2 | from_2695 | to_2695 | art_long | art_long | M |
| 5 | 127 | 0.687552996 | cat2 | from_2696 | to_2696 | art_long | art_long | M |
| 7 | 127 | 0.731567472 | cat2 | from_2697 | to_2697 | art_long | art_long | M |
| 22 | 127 | 0.718915293 | cat2 | from_2698 | to_2698 | art_long | art_long | M |
| 34 | 127 | 0.952114121 | cat2 | from_2699 | to_2699 | art_long | art_long | M |
| 63 | 127 | 0.649101703 | cat2 | from_2700 | to_2700 | art_long | art_long | M |
| 64 | 127 | 0.728653554 | cat2 | from_2701 | to_2701 | art_long | art_long | M |
| 65 | 127 | 0.937995159 | cat2 | from_2702 | to_2702 | art_long | art_long | M |

| | | | | | | | | |
|---|---|---|---|---|---|---|---|---|
| 70 | 127 | 0.675622646 | cat2 | from_2703 | to_2703 | art_long | art_long | M |
| 85 | 127 | 0.717708355 | cat2 | from_2704 | to_2704 | art_long | art_long | M |
| 113 | 127 | 0.963606957 | cat2 | from_2705 | to_2705 | art_long | art_long | M |
| 116 | 127 | 0.649330091 | cat2 | from_2706 | to_2706 | art_long | art_long | M |
| 117 | 127 | 0.728726056 | cat2 | from_2707 | to_2707 | art_long | art_long | M |
| 123 | 127 | 0.717604987 | cat2 | from_2708 | to_2708 | art_long | art_long | M |
| 11 | 128 | 0.646095639 | cat2 | from_2709 | to_2709 | art_long | art_long | M |
| 35 | 128 | 0.653190209 | cat2 | from_2710 | to_2710 | art_long | art_long | M |
| 38 | 128 | 0.649851017 | cat2 | from_2711 | to_2711 | art_long | art_long | M |
| 42 | 128 | 0.651882277 | cat2 | from_2712 | to_2712 | art_long | art_long | M |
| 44 | 128 | 0.989710485 | cat2 | from_2713 | to_2713 | art_long | art_long | M |
| 45 | 128 | 0.648300871 | cat2 | from_2714 | to_2714 | art_long | art_long | M |
| 46 | 128 | 0.999371271 | cat2 | from_2715 | to_2715 | art_long | art_long | M |
| 48 | 128 | 0.65312478 | cat2 | from_2716 | to_2716 | art_long | art_long | M |
| 52 | 128 | 0.876279261 | cat2 | from_2717 | to_2717 | art_long | art_long | M |
| 60 | 128 | 0.997108602 | cat2 | from_2718 | to_2718 | art_long | art_long | M |
| 61 | 128 | 0.914243594 | cat2 | from_2719 | to_2719 | art_long | art_long | M |
| 62 | 128 | 0.653318726 | cat2 | from_2720 | to_2720 | art_long | art_long | M |
| 92 | 128 | 0.645175723 | cat2 | from_2721 | to_2721 | art_long | art_long | M |
| 99 | 128 | 0.915744966 | cat2 | from_2722 | to_2722 | art_long | art_long | M |
| 100 | 128 | 0.654104978 | cat2 | from_2723 | to_2723 | art_long | art_long | M |
| 101 | 128 | 0.653528369 | cat2 | from_2724 | to_2724 | art_long | art_long | M |
| 102 | 128 | 0.977274976 | cat2 | from_2725 | to_2725 | art_long | art_long | M |
| 104 | 128 | 0.650035642 | cat2 | from_2726 | to_2726 | art_long | art_long | M |
| 105 | 128 | 0.728615676 | cat2 | from_2727 | to_2727 | art_long | art_long | M |
| 106 | 128 | 0.948672738 | cat2 | from_2728 | to_2728 | art_long | art_long | M |
| 107 | 128 | 0.647590151 | cat2 | from_2729 | to_2729 | art_long | art_long | M |
| 25 | 129 | 0.648939934 | cat2 | from_2730 | to_2730 | art_long | art_long | M |
| 26 | 129 | 0.838112446 | cat2 | from_2731 | to_2731 | art_long | art_long | M |
| 37 | 129 | 0.945593997 | cat2 | from_2732 | to_2732 | art_long | art_long | M |
| 108 | 129 | 0.665650089 | cat2 | from_2733 | to_2733 | art_long | art_long | M |
| 112 | 129 | 0.653505248 | cat2 | from_2734 | to_2734 | art_long | art_long | M |
| 115 | 129 | 0.82207357 | cat2 | from_2735 | to_2735 | art_long | art_long | M |
| 118 | 129 | 0.671309075 | cat2 | from_2736 | to_2736 | art_long | art_long | M |
| 26 | 130 | 0.958818662 | cat2 | from_2737 | to_2737 | art_long | art_long | M |
| 37 | 130 | 0.997068155 | cat2 | from_2738 | to_2738 | art_long | art_long | M |
| 108 | 130 | 0.665327358 | cat2 | from_2739 | to_2739 | art_long | art_long | M |
| 115 | 130 | 0.939927563 | cat2 | from_2740 | to_2740 | art_long | art_long | M |
| 118 | 130 | 0.771081795 | cat2 | from_2741 | to_2741 | art_long | art_long | M |
| 129 | 130 | 0.95112754 | cat2 | from_2742 | to_2742 | art_long | art_long | M |
| 26 | 131 | 0.785961933 | cat2 | from_2743 | to_2743 | art_long | art_long | M |
| 37 | 131 | 0.774833093 | cat2 | from_2744 | to_2744 | art_long | art_long | M |
| 47 | 131 | 0.649647864 | cat2 | from_2745 | to_2745 | art_long | art_long | M |
| 55 | 131 | 0.65065561 | cat2 | from_2746 | to_2746 | art_long | art_long | M |
| 58 | 131 | 0.663208393 | cat2 | from_2747 | to_2747 | art_long | art_long | M |
| 69 | 131 | 0.65965512 | cat2 | from_2748 | to_2748 | art_long | art_long | M |
| 87 | 131 | 0.661920529 | cat2 | from_2749 | to_2749 | art_long | art_long | M |
| 95 | 131 | 0.656682194 | cat2 | from_2750 | to_2750 | art_long | art_long | M |
| 109 | 131 | 0.6504888 | cat2 | from_2751 | to_2751 | art_long | art_long | M |
| 111 | 131 | 0.658321039 | cat2 | from_2752 | to_2752 | art_long | art_long | M |
| 115 | 131 | 0.774889095 | cat2 | from_2753 | to_2753 | art_long | art_long | M |
| 118 | 131 | 0.999934339 | cat2 | from_2754 | to_2754 | art_long | art_long | M |
| 119 | 131 | 0.651593191 | cat2 | from_2755 | to_2755 | art_long | art_long | M |
| 129 | 131 | 0.672002799 | cat2 | from_2756 | to_2756 | art_long | art_long | M |
| 130 | 131 | 0.772232887 | cat2 | from_2757 | to_2757 | art_long | art_long | M |
| 17 | 132 | 0.756173633 | cat2 | from_2758 | to_2758 | art_long | art_long | M |
| 31 | 132 | -0.647321897 | cat2 | from_2759 | to_2759 | art_long | art_long | M |
| 33 | 132 | -0.64278646 | cat2 | from_2760 | to_2760 | art_long | art_long | M |
| 35 | 132 | 0.975652151 | cat2 | from_2761 | to_2761 | art_long | art_long | M |
| 36 | 132 | 0.758012316 | cat2 | from_2762 | to_2762 | art_long | art_long | M |
| 38 | 132 | 0.972154705 | cat2 | from_2763 | to_2763 | art_long | art_long | M |
| 44 | 132 | 0.673108384 | cat2 | from_2764 | to_2764 | art_long | art_long | M |
| 46 | 132 | 0.660588456 | cat2 | from_2765 | to_2765 | art_long | art_long | M |
| 48 | 132 | 0.975459473 | cat2 | from_2766 | to_2766 | art_long | art_long | M |
| 50 | 132 | 0.761786652 | cat2 | from_2767 | to_2767 | art_long | art_long | M |
| 60 | 132 | 0.639086497 | cat2 | from_2768 | to_2768 | art_long | art_long | M |
| 102 | 132 | 0.671948218 | cat2 | from_2769 | to_2769 | art_long | art_long | M |
| 104 | 132 | 0.966980977 | cat2 | from_2770 | to_2770 | art_long | art_long | M |
| 106 | 132 | 0.675907031 | cat2 | from_2771 | to_2771 | art_long | art_long | M |
| 128 | 132 | 0.653590019 | cat2 | from_2772 | to_2772 | art_long | art_long | M |
| 17 | 133 | 0.999262003 | cat2 | from_2773 | to_2773 | art_long | art_long | M |
| 35 | 133 | 0.791953625 | cat2 | from_2774 | to_2774 | art_long | art_long | M |
| 36 | 133 | 0.999513159 | cat2 | from_2775 | to_2775 | art_long | art_long | M |
| 38 | 133 | 0.787764687 | cat2 | from_2776 | to_2776 | art_long | art_long | M |
| 42 | 133 | 0.646389735 | cat2 | from_2777 | to_2777 | art_long | art_long | M |
| 45 | 133 | 0.665608235 | cat2 | from_2778 | to_2778 | art_long | art_long | M |
| 48 | 133 | 0.791980141 | cat2 | from_2779 | to_2779 | art_long | art_long | M |
| 62 | 133 | 0.652071466 | cat2 | from_2780 | to_2780 | art_long | art_long | M |
| 92 | 133 | 0.663699822 | cat2 | from_2781 | to_2781 | art_long | art_long | M |
| 100 | 133 | 0.657340848 | cat2 | from_2782 | to_2782 | art_long | art_long | M |
| 101 | 133 | 0.652324698 | cat2 | from_2783 | to_2783 | art_long | art_long | M |
| 104 | 133 | 0.792542526 | cat2 | from_2784 | to_2784 | art_long | art_long | M |
| 107 | 133 | 0.665727541 | cat2 | from_2785 | to_2785 | art_long | art_long | M |
| 132 | 133 | 0.765962261 | cat2 | from_2786 | to_2786 | art_long | art_long | M |
| 17 | 134 | 0.999830245 | cat2 | from_2787 | to_2787 | art_long | art_long | M |
| 35 | 134 | 0.789377028 | cat2 | from_2788 | to_2788 | art_long | art_long | M |
| 36 | 134 | 0.999700041 | cat2 | from_2789 | to_2789 | art_long | art_long | M |
| 38 | 134 | 0.785968714 | cat2 | from_2790 | to_2790 | art_long | art_long | M |
| 42 | 134 | 0.633421083 | cat2 | from_2791 | to_2791 | art_long | art_long | M |
| 45 | 134 | 0.66452487 | cat2 | from_2792 | to_2792 | art_long | art_long | M |
| 48 | 134 | 0.789452293 | cat2 | from_2793 | to_2793 | art_long | art_long | M |
| 62 | 134 | 0.642103652 | cat2 | from_2794 | to_2794 | art_long | art_long | M |
| 92 | 134 | 0.664124113 | cat2 | from_2795 | to_2795 | art_long | art_long | M |
| 100 | 134 | 0.648866416 | cat2 | from_2796 | to_2796 | art_long | art_long | M |
| 101 | 134 | 0.641555767 | cat2 | from_2797 | to_2797 | art_long | art_long | M |
| 104 | 134 | 0.792000363 | cat2 | from_2798 | to_2798 | art_long | art_long | M |
| 107 | 134 | 0.665085865 | cat2 | from_2799 | to_2799 | art_long | art_long | M |
| 132 | 134 | 0.751254724 | cat2 | from_2800 | to_2800 | art_long | art_long | M |
| 133 | 134 | 0.998449212 | cat2 | from_2801 | to_2801 | art_long | art_long | M |
| 11 | 135 | 0.64580939 | cat2 | from_2802 | to_2802 | art_long | art_long | M |
| 35 | 135 | 0.653054533 | cat2 | from_2803 | to_2803 | art_long | art_long | M |
| 38 | 135 | 0.648847678 | cat2 | from_2804 | to_2804 | art_long | art_long | M |
| 42 | 135 | 0.659169491 | cat2 | from_2805 | to_2805 | art_long | art_long | M |
| 44 | 135 | 0.994071261 | cat2 | from_2806 | to_2806 | art_long | art_long | M |

| | | | | | | | | |
|---|---|---|---|---|---|---|---|---|
| 45 | 135 | 0.648371303 | cat2 | from_2807 | to_2807 | art_long | art_long | M |
| 46 | 135 | 0.999998464 | cat2 | from_2808 | to_2808 | art_long | art_long | M |
| 48 | 135 | 0.652959551 | cat2 | from_2809 | to_2809 | art_long | art_long | M |
| 52 | 135 | 0.87106527 | cat2 | from_2810 | to_2810 | art_long | art_long | M |
| 60 | 135 | 0.994671856 | cat2 | from_2811 | to_2811 | art_long | art_long | M |
| 61 | 135 | 0.908554607 | cat2 | from_2812 | to_2812 | art_long | art_long | M |
| 62 | 135 | 0.658595149 | cat2 | from_2813 | to_2813 | art_long | art_long | M |
| 92 | 135 | 0.644144896 | cat2 | from_2814 | to_2814 | art_long | art_long | M |
| 99 | 135 | 0.916573379 | cat2 | from_2815 | to_2815 | art_long | art_long | M |
| 100 | 135 | 0.658660046 | cat2 | from_2816 | to_2816 | art_long | art_long | M |
| 101 | 135 | 0.659477813 | cat2 | from_2817 | to_2817 | art_long | art_long | M |
| 102 | 135 | 0.983398919 | cat2 | from_2818 | to_2818 | art_long | art_long | M |
| 104 | 135 | 0.648667699 | cat2 | from_2819 | to_2819 | art_long | art_long | M |
| 105 | 135 | 0.726760394 | cat2 | from_2820 | to_2820 | art_long | art_long | M |
| 106 | 135 | 0.958927018 | cat2 | from_2821 | to_2821 | art_long | art_long | M |
| 107 | 135 | 0.647392912 | cat2 | from_2822 | to_2822 | art_long | art_long | M |
| 128 | 135 | 0.999313512 | cat2 | from_2823 | to_2823 | art_long | art_long | M |
| 132 | 135 | 0.660850226 | cat2 | from_2824 | to_2824 | art_long | art_long | M |
| 26 | 136 | 0.953987853 | cat2 | from_2825 | to_2825 | art_long | art_long | M |
| 37 | 136 | 0.981703097 | cat2 | from_2826 | to_2826 | art_long | art_long | M |
| 108 | 136 | 0.654838505 | cat2 | from_2827 | to_2827 | art_long | art_long | M |
| 115 | 136 | 0.934896988 | cat2 | from_2828 | to_2828 | art_long | art_long | M |
| 118 | 136 | 0.747571306 | cat2 | from_2829 | to_2829 | art_long | art_long | M |
| 129 | 136 | 0.95900085 | cat2 | from_2830 | to_2830 | art_long | art_long | M |
| 130 | 136 | 0.989545898 | cat2 | from_2831 | to_2831 | art_long | art_long | M |
| 131 | 136 | 0.748394235 | cat2 | from_2832 | to_2832 | art_long | art_long | M |
| 26 | 137 | 0.715349325 | cat2 | from_2833 | to_2833 | art_long | art_long | M |
| 37 | 137 | 0.771995997 | cat2 | from_2834 | to_2834 | art_long | art_long | M |
| 72 | 137 | 0.632542174 | cat2 | from_2835 | to_2835 | art_long | art_long | M |
| 115 | 137 | 0.708401543 | cat2 | from_2836 | to_2836 | art_long | art_long | M |
| 118 | 137 | 0.95553186 | cat2 | from_2837 | to_2837 | art_long | art_long | M |
| 120 | 137 | 0.635498253 | cat2 | from_2838 | to_2838 | art_long | art_long | M |
| 129 | 137 | 0.763217193 | cat2 | from_2839 | to_2839 | art_long | art_long | M |
| 130 | 137 | 0.771252173 | cat2 | from_2840 | to_2840 | art_long | art_long | M |
| 131 | 137 | 0.954998101 | cat2 | from_2841 | to_2841 | art_long | art_long | M |
| 136 | 137 | 0.765473032 | cat2 | from_2842 | to_2842 | art_long | art_long | M |
| 17 | 138 | 0.990974776 | cat2 | from_2843 | to_2843 | art_long | art_long | M |
| 35 | 138 | 0.781559284 | cat2 | from_2844 | to_2844 | art_long | art_long | M |
| 36 | 138 | 0.990040417 | cat2 | from_2845 | to_2845 | art_long | art_long | M |
| 38 | 138 | 0.782058998 | cat2 | from_2846 | to_2846 | art_long | art_long | M |
| 45 | 138 | 0.643286308 | cat2 | from_2847 | to_2847 | art_long | art_long | M |
| 48 | 138 | 0.781644569 | cat2 | from_2848 | to_2848 | art_long | art_long | M |
| 92 | 138 | 0.644960862 | cat2 | from_2849 | to_2849 | art_long | art_long | M |
| 104 | 138 | 0.784605525 | cat2 | from_2850 | to_2850 | art_long | art_long | M |
| 107 | 138 | 0.643941969 | cat2 | from_2851 | to_2851 | art_long | art_long | M |
| 132 | 138 | 0.741121786 | cat2 | from_2852 | to_2852 | art_long | art_long | M |
| 133 | 138 | 0.988418533 | cat2 | from_2853 | to_2853 | art_long | art_long | M |
| 134 | 138 | 0.990638237 | cat2 | from_2854 | to_2854 | art_long | art_long | M |
| 42 | 139 | 0.936789642 | cat2 | from_2855 | to_2855 | art_long | art_long | M |
| 43 | 139 | 0.914411705 | cat2 | from_2856 | to_2856 | art_long | art_long | M |
| 45 | 139 | 0.940058198 | cat2 | from_2857 | to_2857 | art_long | art_long | M |
| 62 | 139 | 0.946080933 | cat2 | from_2858 | to_2858 | art_long | art_long | M |
| 92 | 139 | 0.941711504 | cat2 | from_2859 | to_2859 | art_long | art_long | M |
| 100 | 139 | 0.943216547 | cat2 | from_2860 | to_2860 | art_long | art_long | M |
| 101 | 139 | 0.94074537 | cat2 | from_2861 | to_2861 | art_long | art_long | M |
| 107 | 139 | 0.939341404 | cat2 | from_2862 | to_2862 | art_long | art_long | M |
| 25 | 140 | 0.65773199 | cat2 | from_2863 | to_2863 | art_long | art_long | M |
| 30 | 140 | 0.928539382 | cat2 | from_2864 | to_2864 | art_long | art_long | M |
| 47 | 140 | 0.914770923 | cat2 | from_2865 | to_2865 | art_long | art_long | M |
| 54 | 140 | 0.935772775 | cat2 | from_2866 | to_2866 | art_long | art_long | M |
| 55 | 140 | 0.915250427 | cat2 | from_2867 | to_2867 | art_long | art_long | M |
| 58 | 140 | 0.917995475 | cat2 | from_2868 | to_2868 | art_long | art_long | M |
| 69 | 140 | 0.920623093 | cat2 | from_2869 | to_2869 | art_long | art_long | M |
| 72 | 140 | 0.999785296 | cat2 | from_2870 | to_2870 | art_long | art_long | M |
| 76 | 140 | 0.646174019 | cat2 | from_2871 | to_2871 | art_long | art_long | M |
| 77 | 140 | 0.995732134 | cat2 | from_2872 | to_2872 | art_long | art_long | M |
| 80 | 140 | 0.992695794 | cat2 | from_2873 | to_2873 | art_long | art_long | M |
| 81 | 140 | 0.904587615 | cat2 | from_2874 | to_2874 | art_long | art_long | M |
| 83 | 140 | 0.983210976 | cat2 | from_2875 | to_2875 | art_long | art_long | M |
| 86 | 140 | 0.638615016 | cat2 | from_2876 | to_2876 | art_long | art_long | M |
| 87 | 140 | 0.918161548 | cat2 | from_2877 | to_2877 | art_long | art_long | M |
| 94 | 140 | 0.643399429 | cat2 | from_2878 | to_2878 | art_long | art_long | M |
| 95 | 140 | 0.91751196 | cat2 | from_2879 | to_2879 | art_long | art_long | M |
| 109 | 140 | 0.918398066 | cat2 | from_2880 | to_2880 | art_long | art_long | M |
| 111 | 140 | 0.919270997 | cat2 | from_2881 | to_2881 | art_long | art_long | M |
| 112 | 140 | 0.652386891 | cat2 | from_2882 | to_2882 | art_long | art_long | M |
| 119 | 140 | 0.917277712 | cat2 | from_2883 | to_2883 | art_long | art_long | M |
| 120 | 140 | 0.999533394 | cat2 | from_2884 | to_2884 | art_long | art_long | M |
| 26 | 141 | 0.716314586 | cat2 | from_2885 | to_2885 | art_long | art_long | M |
| 37 | 141 | 0.773006327 | cat2 | from_2886 | to_2886 | art_long | art_long | M |
| 115 | 141 | 0.708149943 | cat2 | from_2887 | to_2887 | art_long | art_long | M |
| 118 | 141 | 0.956301344 | cat2 | from_2888 | to_2888 | art_long | art_long | M |
| 120 | 141 | 0.634925041 | cat2 | from_2889 | to_2889 | art_long | art_long | M |
| 129 | 141 | 0.763769631 | cat2 | from_2890 | to_2890 | art_long | art_long | M |
| 130 | 141 | 0.772643051 | cat2 | from_2891 | to_2891 | art_long | art_long | M |
| 131 | 141 | 0.955866639 | cat2 | from_2892 | to_2892 | art_long | art_long | M |
| 136 | 141 | 0.766196565 | cat2 | from_2893 | to_2893 | art_long | art_long | M |
| 137 | 141 | 0.999958097 | cat2 | from_2894 | to_2894 | art_long | art_long | M |
| 25 | 142 | 0.646141142 | cat2 | from_2895 | to_2895 | art_long | art_long | M |
| 26 | 142 | 0.836354687 | cat2 | from_2896 | to_2896 | art_long | art_long | M |
| 37 | 142 | 0.939568324 | cat2 | from_2897 | to_2897 | art_long | art_long | M |
| 108 | 142 | 0.662595033 | cat2 | from_2898 | to_2898 | art_long | art_long | M |
| 112 | 142 | 0.653831013 | cat2 | from_2899 | to_2899 | art_long | art_long | M |
| 115 | 142 | 0.809953635 | cat2 | from_2900 | to_2900 | art_long | art_long | M |
| 118 | 142 | 0.668189223 | cat2 | from_2901 | to_2901 | art_long | art_long | M |
| 129 | 142 | 0.997699679 | cat2 | from_2902 | to_2902 | art_long | art_long | M |
| 130 | 142 | 0.950029205 | cat2 | from_2903 | to_2903 | art_long | art_long | M |
| 131 | 142 | 0.669600703 | cat2 | from_2904 | to_2904 | art_long | art_long | M |
| 136 | 142 | 0.959391495 | cat2 | from_2905 | to_2905 | art_long | art_long | M |
| 137 | 142 | 0.758706038 | cat2 | from_2906 | to_2906 | art_long | art_long | M |
| 141 | 142 | 0.759731186 | cat2 | from_2907 | to_2907 | art_long | art_long | M |
| 17 | 143 | 0.684659976 | cat2 | from_2908 | to_2908 | art_long | art_long | M |
| 35 | 143 | 0.927548809 | cat2 | from_2909 | to_2909 | art_long | art_long | M |
| 36 | 143 | 0.682597592 | cat2 | from_2910 | to_2910 | art_long | art_long | M |

| | | | | | | | | |
|---|---|---|---|---|---|---|---|---|
| 38 | 143 | 0.938355282 | cat2 | from_2911 | to_2911 | art_long | art_long | M |
| 48 | 143 | 0.927608753 | cat2 | from_2912 | to_2912 | art_long | art_long | M |
| 50 | 143 | 0.848259868 | cat2 | from_2913 | to_2913 | art_long | art_long | M |
| 104 | 143 | 0.929443118 | cat2 | from_2914 | to_2914 | art_long | art_long | M |
| 132 | 143 | 0.889881372 | cat2 | from_2915 | to_2915 | art_long | art_long | M |
| 133 | 143 | 0.681432782 | cat2 | from_2916 | to_2916 | art_long | art_long | M |
| 134 | 143 | 0.68304633 | cat2 | from_2917 | to_2917 | art_long | art_long | M |
| 138 | 143 | 0.725600345 | cat2 | from_2918 | to_2918 | art_long | art_long | M |
| 26 | 144 | 0.780331423 | cat2 | from_2919 | to_2919 | art_long | art_long | M |
| 37 | 144 | 0.771898585 | cat2 | from_2920 | to_2920 | art_long | art_long | M |
| 47 | 144 | 0.660008581 | cat2 | from_2921 | to_2921 | art_long | art_long | M |
| 55 | 144 | 0.660622105 | cat2 | from_2922 | to_2922 | art_long | art_long | M |
| 58 | 144 | 0.66685473 | cat2 | from_2923 | to_2923 | art_long | art_long | M |
| 69 | 144 | 0.664500139 | cat2 | from_2924 | to_2924 | art_long | art_long | M |
| 87 | 144 | 0.666410805 | cat2 | from_2925 | to_2925 | art_long | art_long | M |
| 95 | 144 | 0.664026482 | cat2 | from_2926 | to_2926 | art_long | art_long | M |
| 109 | 144 | 0.659867066 | cat2 | from_2927 | to_2927 | art_long | art_long | M |
| 111 | 144 | 0.664422077 | cat2 | from_2928 | to_2928 | art_long | art_long | M |
| 115 | 144 | 0.779031317 | cat2 | from_2929 | to_2929 | art_long | art_long | M |
| 118 | 144 | 0.998574606 | cat2 | from_2930 | to_2930 | art_long | art_long | M |
| 119 | 144 | 0.660859138 | cat2 | from_2931 | to_2931 | art_long | art_long | M |
| 129 | 144 | 0.664318107 | cat2 | from_2932 | to_2932 | art_long | art_long | M |
| 130 | 144 | 0.764599769 | cat2 | from_2933 | to_2933 | art_long | art_long | M |
| 131 | 144 | 0.997969255 | cat2 | from_2934 | to_2934 | art_long | art_long | M |
| 136 | 144 | 0.738986148 | cat2 | from_2935 | to_2935 | art_long | art_long | M |
| 137 | 144 | 0.953448148 | cat2 | from_2936 | to_2936 | art_long | art_long | M |
| 141 | 144 | 0.953881182 | cat2 | from_2937 | to_2937 | art_long | art_long | M |
| 142 | 144 | 0.657651361 | cat2 | from_2938 | to_2938 | art_long | art_long | M |
| 8 | 145 | 0.766153506 | cat2 | from_2939 | to_2939 | art_long | art_long | M |
| 93 | 145 | 0.832432853 | cat2 | from_2940 | to_2940 | art_long | art_long | M |
| 49 | 146 | 0.95567901 | cat2 | from_2941 | to_2941 | art_long | art_long | M |
| 56 | 146 | 0.794206544 | cat2 | from_2942 | to_2942 | art_long | art_long | M |
| 57 | 146 | 0.795424862 | cat2 | from_2943 | to_2943 | art_long | art_long | M |
| 65 | 146 | 0.661600712 | cat2 | from_2944 | to_2944 | art_long | art_long | M |
| 70 | 146 | 0.997363951 | cat2 | from_2945 | to_2945 | art_long | art_long | M |
| 75 | 146 | 0.956196329 | cat2 | from_2946 | to_2946 | art_long | art_long | M |
| 84 | 146 | 0.711461137 | cat2 | from_2947 | to_2947 | art_long | art_long | M |
| 85 | 146 | 0.958622549 | cat2 | from_2948 | to_2948 | art_long | art_long | M |
| 108 | 146 | 0.643278214 | cat2 | from_2949 | to_2949 | art_long | art_long | M |
| 113 | 146 | 0.647114731 | cat2 | from_2950 | to_2950 | art_long | art_long | M |
| 123 | 146 | 0.960254457 | cat2 | from_2951 | to_2951 | art_long | art_long | M |
| 127 | 146 | 0.679509534 | cat2 | from_2952 | to_2952 | art_long | art_long | M |
| 7 | 147 | 0.633376427 | cat2 | from_2953 | to_2953 | art_long | art_long | M |
| 17 | 147 | 0.642807244 | cat2 | from_2954 | to_2954 | art_long | art_long | M |
| 36 | 147 | 0.644800102 | cat2 | from_2955 | to_2955 | art_long | art_long | M |
| 42 | 147 | 0.999547335 | cat2 | from_2956 | to_2956 | art_long | art_long | M |
| 43 | 147 | 0.907952189 | cat2 | from_2957 | to_2957 | art_long | art_long | M |
| 44 | 147 | 0.665896853 | cat2 | from_2958 | to_2958 | art_long | art_long | M |
| 45 | 147 | 0.982809459 | cat2 | from_2959 | to_2959 | art_long | art_long | M |
| 46 | 147 | 0.659334591 | cat2 | from_2960 | to_2960 | art_long | art_long | M |
| 60 | 147 | 0.635534418 | cat2 | from_2961 | to_2961 | art_long | art_long | M |
| 62 | 147 | 0.999602159 | cat2 | from_2962 | to_2962 | art_long | art_long | M |
| 92 | 147 | 0.977384312 | cat2 | from_2963 | to_2963 | art_long | art_long | M |
| 100 | 147 | 0.998635076 | cat2 | from_2964 | to_2964 | art_long | art_long | M |
| 101 | 147 | 0.999946123 | cat2 | from_2965 | to_2965 | art_long | art_long | M |
| 102 | 147 | 0.658915944 | cat2 | from_2966 | to_2966 | art_long | art_long | M |
| 106 | 147 | 0.659297773 | cat2 | from_2967 | to_2967 | art_long | art_long | M |
| 107 | 147 | 0.981312688 | cat2 | from_2968 | to_2968 | art_long | art_long | M |
| 128 | 147 | 0.653208124 | cat2 | from_2969 | to_2969 | art_long | art_long | M |
| 133 | 147 | 0.6509039 | cat2 | from_2970 | to_2970 | art_long | art_long | M |
| 134 | 147 | 0.639568921 | cat2 | from_2971 | to_2971 | art_long | art_long | M |
| 135 | 147 | 0.659501719 | cat2 | from_2972 | to_2972 | art_long | art_long | M |
| 139 | 147 | 0.939877421 | cat2 | from_2973 | to_2973 | art_long | art_long | M |
| 44 | 148 | 0.919621791 | cat2 | from_2974 | to_2974 | art_long | art_long | M |
| 46 | 148 | 0.903791538 | cat2 | from_2975 | to_2975 | art_long | art_long | M |
| 52 | 148 | 0.718497984 | cat2 | from_2976 | to_2976 | art_long | art_long | M |
| 60 | 148 | 0.90270517 | cat2 | from_2977 | to_2977 | art_long | art_long | M |
| 61 | 148 | 0.953675113 | cat2 | from_2978 | to_2978 | art_long | art_long | M |
| 99 | 148 | 0.993132553 | cat2 | from_2979 | to_2979 | art_long | art_long | M |
| 102 | 148 | 0.928832008 | cat2 | from_2980 | to_2980 | art_long | art_long | M |
| 105 | 148 | 0.928605548 | cat2 | from_2981 | to_2981 | art_long | art_long | M |
| 106 | 148 | 0.921394639 | cat2 | from_2982 | to_2982 | art_long | art_long | M |
| 128 | 148 | 0.89931874 | cat2 | from_2983 | to_2983 | art_long | art_long | M |
| 135 | 148 | 0.904129071 | cat2 | from_2984 | to_2984 | art_long | art_long | M |
| 4 | 149 | 0.652856632 | cat2 | from_2985 | to_2985 | art_long | art_long | M |
| 7 | 149 | 0.652752055 | cat2 | from_2986 | to_2986 | art_long | art_long | M |
| 17 | 149 | 0.662663522 | cat2 | from_2987 | to_2987 | art_long | art_long | M |
| 36 | 149 | 0.663599056 | cat2 | from_2988 | to_2988 | art_long | art_long | M |
| 42 | 149 | 0.982498749 | cat2 | from_2989 | to_2989 | art_long | art_long | M |
| 43 | 149 | 0.927905492 | cat2 | from_2990 | to_2990 | art_long | art_long | M |
| 44 | 149 | 0.640956486 | cat2 | from_2991 | to_2991 | art_long | art_long | M |
| 45 | 149 | 0.999596339 | cat2 | from_2992 | to_2992 | art_long | art_long | M |
| 46 | 149 | 0.651644842 | cat2 | from_2993 | to_2993 | art_long | art_long | M |
| 60 | 149 | 0.635640298 | cat2 | from_2994 | to_2994 | art_long | art_long | M |
| 62 | 149 | 0.991163824 | cat2 | from_2995 | to_2995 | art_long | art_long | M |
| 92 | 149 | 0.998376874 | cat2 | from_2996 | to_2996 | art_long | art_long | M |
| 100 | 149 | 0.994490491 | cat2 | from_2997 | to_2997 | art_long | art_long | M |
| 101 | 149 | 0.989230591 | cat2 | from_2998 | to_2998 | art_long | art_long | M |
| 107 | 149 | 0.999339272 | cat2 | from_2999 | to_2999 | art_long | art_long | M |
| 128 | 149 | 0.65050931 | cat2 | from_3000 | to_3000 | art_long | art_long | M |
| 133 | 149 | 0.664837559 | cat2 | from_3001 | to_3001 | art_long | art_long | M |
| 134 | 149 | 0.662174203 | cat2 | from_3002 | to_3002 | art_long | art_long | M |
| 135 | 149 | 0.651537547 | cat2 | from_3003 | to_3003 | art_long | art_long | M |
| 138 | 149 | 0.640607855 | cat2 | from_3004 | to_3004 | art_long | art_long | M |
| 139 | 149 | 0.9421377 | cat2 | from_3005 | to_3005 | art_long | art_long | M |
| 147 | 149 | 0.987657969 | cat2 | from_3006 | to_3006 | art_long | art_long | M |
| 3 | 150 | 0.659454587 | cat2 | from_3007 | to_3007 | art_long | art_long | M |
| 29 | 150 | 0.815590562 | cat2 | from_3008 | to_3008 | art_long | art_long | M |
| 41 | 150 | 0.719554776 | cat2 | from_3009 | to_3009 | art_long | art_long | M |
| 66 | 150 | 0.734796012 | cat2 | from_3010 | to_3010 | art_long | art_long | M |
| 67 | 150 | 0.730102156 | cat2 | from_3011 | to_3011 | art_long | art_long | M |
| 71 | 150 | 0.849786533 | cat2 | from_3012 | to_3012 | art_long | art_long | M |
| 82 | 150 | 0.965177516 | cat2 | from_3013 | to_3013 | art_long | art_long | M |
| 96 | 150 | 0.708132992 | cat2 | from_3014 | to_3014 | art_long | art_long | M |

| | | | | | | | | |
|---|---|---|---|---|---|---|---|---|
| 98 | 150 | 0.832291198 | cat2 | from_3015 | to_3015 | art_long | art_long | M |
| 121 | 150 | 0.815278176 | cat2 | from_3016 | to_3016 | art_long | art_long | M |
| 124 | 150 | 0.719519605 | cat2 | from_3017 | to_3017 | art_long | art_long | M |
| 1 | 151 | -0.835511836 | cat2 | from_3018 | to_3018 | art_long | art_long | M |
| 93 | 152 | 0.831410413 | cat2 | from_3019 | to_3019 | art_long | art_long | M |
| 105 | 152 | 0.734269283 | cat2 | from_3020 | to_3020 | art_long | art_long | M |
| 148 | 152 | 0.675847839 | cat2 | from_3021 | to_3021 | art_long | art_long | M |
| 89 | 153 | 0.643583971 | cat2 | from_3022 | to_3022 | art_long | art_long | M |
| 93 | 153 | 0.76533951 | cat2 | from_3023 | to_3023 | art_long | art_long | M |
| 105 | 153 | 0.682053241 | cat2 | from_3024 | to_3024 | art_long | art_long | M |
| 148 | 153 | 0.682087755 | cat2 | from_3025 | to_3025 | art_long | art_long | M |
| 152 | 153 | 0.985686239 | cat2 | from_3026 | to_3026 | art_long | art_long | M |
| 17 | 154 | 0.82964948 | cat2 | from_3027 | to_3027 | art_long | art_short | J |
| 35 | 154 | 0.647769787 | cat2 | from_3028 | to_3028 | art_long | art_short | J |
| 36 | 154 | 0.825813172 | cat2 | from_3029 | to_3029 | art_long | art_short | J |
| 38 | 154 | 0.661525734 | cat2 | from_3030 | to_3030 | art_long | art_short | J |
| 48 | 154 | 0.647938423 | cat2 | from_3031 | to_3031 | art_long | art_short | J |
| 91 | 154 | -0.657937628 | cat2 | from_3032 | to_3032 | art_long | art_short | J |
| 104 | 154 | 0.654379026 | cat2 | from_3033 | to_3033 | art_long | art_short | J |
| 133 | 154 | 0.820984346 | cat2 | from_3034 | to_3034 | art_long | art_short | J |
| 134 | 154 | 0.829040394 | cat2 | from_3035 | to_3035 | art_long | art_short | J |
| 138 | 154 | 0.896979691 | cat2 | from_3036 | to_3036 | art_long | art_short | J |
| 143 | 154 | 0.763494 | cat2 | from_3037 | to_3037 | art_long | art_short | J |
| 2 | 155 | 0.970245817 | cat2 | from_3038 | to_3038 | art_long | art_short | J |
| 12 | 155 | 0.652102246 | cat2 | from_3039 | to_3039 | art_long | art_short | J |
| 16 | 155 | 0.723494832 | cat2 | from_3040 | to_3040 | art_long | art_short | J |
| 18 | 155 | 0.724037108 | cat2 | from_3041 | to_3041 | art_long | art_short | J |
| 39 | 155 | 0.678768078 | cat2 | from_3042 | to_3042 | art_long | art_short | J |
| 53 | 155 | 0.98053428 | cat2 | from_3043 | to_3043 | art_long | art_short | J |
| 8 | 156 | 0.749645462 | cat2 | from_3044 | to_3044 | art_long | art_short | J |
| 13 | 156 | 0.668893827 | cat2 | from_3045 | to_3045 | art_long | art_short | J |
| 19 | 156 | 0.634106974 | cat2 | from_3046 | to_3046 | art_long | art_short | J |
| 79 | 156 | 0.685405807 | cat2 | from_3047 | to_3047 | art_long | art_short | J |
| 114 | 156 | 0.808873435 | cat2 | from_3048 | to_3048 | art_long | art_short | J |
| 125 | 156 | 0.742219005 | cat2 | from_3049 | to_3049 | art_long | art_short | J |
| 6 | 157 | 0.847191091 | cat2 | from_3050 | to_3050 | art_long | art_short | J |
| 8 | 157 | 0.815754545 | cat2 | from_3051 | to_3051 | art_long | art_short | J |
| 125 | 157 | 0.764325384 | cat2 | from_3052 | to_3052 | art_long | art_short | J |
| 156 | 157 | 0.692196673 | cat2 | from_3053 | to_3053 | art_short | art_short | H |
| 8 | 158 | 0.749704977 | cat2 | from_3054 | to_3054 | art_long | art_short | J |
| 13 | 158 | 0.671365455 | cat2 | from_3055 | to_3055 | art_long | art_short | J |
| 19 | 158 | 0.63617578 | cat2 | from_3056 | to_3056 | art_long | art_short | J |
| 79 | 158 | 0.68641889 | cat2 | from_3057 | to_3057 | art_long | art_short | J |
| 114 | 158 | 0.809729462 | cat2 | from_3058 | to_3058 | art_long | art_short | J |
| 125 | 158 | 0.743710267 | cat2 | from_3059 | to_3059 | art_long | art_short | J |
| 156 | 158 | 0.999965973 | cat2 | from_3060 | to_3060 | art_short | art_short | H |
| 157 | 158 | 0.690381091 | cat2 | from_3061 | to_3061 | art_short | art_short | H |
| 4 | 159 | 0.859422817 | cat2 | from_3062 | to_3062 | art_long | art_short | J |
| 6 | 159 | 0.863344924 | cat2 | from_3063 | to_3063 | art_long | art_short | J |
| 7 | 159 | 0.861691552 | cat2 | from_3064 | to_3064 | art_long | art_short | J |
| 11 | 159 | 0.697745148 | cat2 | from_3065 | to_3065 | art_long | art_short | J |
| 113 | 159 | 0.66058522 | cat2 | from_3066 | to_3066 | art_long | art_short | J |
| 118 | 159 | -0.632186579 | cat2 | from_3067 | to_3067 | art_long | art_short | J |
| 137 | 159 | -0.646941142 | cat2 | from_3068 | to_3068 | art_long | art_short | J |
| 141 | 159 | -0.645328801 | cat2 | from_3069 | to_3069 | art_long | art_short | J |
| 144 | 159 | -0.642196687 | cat2 | from_3070 | to_3070 | art_long | art_short | J |
| 4 | 160 | 0.95888708 | cat2 | from_3071 | to_3071 | art_long | art_short | J |
| 6 | 160 | 0.677350565 | cat2 | from_3072 | to_3072 | art_long | art_short | J |
| 7 | 160 | 0.9616363 | cat2 | from_3073 | to_3073 | art_long | art_short | J |
| 11 | 160 | 0.770138358 | cat2 | from_3074 | to_3074 | art_long | art_short | J |
| 34 | 160 | 0.710242387 | cat2 | from_3075 | to_3075 | art_long | art_short | J |
| 113 | 160 | 0.698988275 | cat2 | from_3076 | to_3076 | art_long | art_short | J |
| 127 | 160 | 0.685306357 | cat2 | from_3077 | to_3077 | art_long | art_short | J |
| 159 | 160 | 0.942690841 | cat2 | from_3078 | to_3078 | art_short | art_short | H |
| 9 | 161 | 0.811365716 | cat2 | from_3079 | to_3079 | art_long | art_short | J |
| 10 | 161 | 0.813411059 | cat2 | from_3080 | to_3080 | art_long | art_short | J |
| 22 | 161 | 0.907841571 | cat2 | from_3081 | to_3081 | art_long | art_short | J |
| 23 | 161 | 0.911512475 | cat2 | from_3082 | to_3082 | art_long | art_short | J |
| 88 | 161 | 0.859439315 | cat2 | from_3083 | to_3083 | art_long | art_short | J |
| 113 | 161 | 0.679652451 | cat2 | from_3084 | to_3084 | art_long | art_short | J |
| 126 | 161 | 0.902499753 | cat2 | from_3085 | to_3085 | art_long | art_short | J |
| 5 | 162 | 0.686119944 | cat2 | from_3086 | to_3086 | art_long | art_short | J |
| 12 | 162 | 0.672463223 | cat2 | from_3087 | to_3087 | art_long | art_short | J |
| 13 | 162 | 0.847435058 | cat2 | from_3088 | to_3088 | art_long | art_short | J |
| 15 | 162 | 0.964935109 | cat2 | from_3089 | to_3089 | art_long | art_short | J |
| 16 | 162 | 0.746382554 | cat2 | from_3090 | to_3090 | art_long | art_short | J |
| 18 | 162 | 0.747164699 | cat2 | from_3091 | to_3091 | art_long | art_short | J |
| 21 | 162 | 0.704392116 | cat2 | from_3092 | to_3092 | art_long | art_short | J |
| 39 | 162 | 0.700316955 | cat2 | from_3093 | to_3093 | art_long | art_short | J |
| 63 | 162 | 0.684403074 | cat2 | from_3094 | to_3094 | art_long | art_short | J |
| 114 | 162 | 0.685872433 | cat2 | from_3095 | to_3095 | art_long | art_short | J |
| 116 | 162 | 0.682508813 | cat2 | from_3096 | to_3096 | art_long | art_short | J |
| 12 | 163 | 0.772838842 | cat2 | from_3097 | to_3097 | art_long | art_short | J |
| 13 | 163 | 0.943682105 | cat2 | from_3098 | to_3098 | art_long | art_short | J |
| 15 | 163 | 0.858922273 | cat2 | from_3099 | to_3099 | art_long | art_short | J |
| 16 | 163 | 0.672167475 | cat2 | from_3100 | to_3100 | art_long | art_short | J |
| 18 | 163 | 0.673354742 | cat2 | from_3101 | to_3101 | art_long | art_short | J |
| 19 | 163 | 0.722610577 | cat2 | from_3102 | to_3102 | art_long | art_short | J |
| 20 | 163 | 0.688045913 | cat2 | from_3103 | to_3103 | art_long | art_short | J |
| 21 | 163 | 0.660624537 | cat2 | from_3104 | to_3104 | art_long | art_short | J |
| 39 | 163 | 0.73599498 | cat2 | from_3105 | to_3105 | art_long | art_short | J |
| 40 | 163 | 0.69494878 | cat2 | from_3106 | to_3106 | art_long | art_short | J |
| 63 | 163 | 0.63537762 | cat2 | from_3107 | to_3107 | art_long | art_short | J |
| 68 | 163 | 0.684420327 | cat2 | from_3108 | to_3108 | art_long | art_short | J |
| 79 | 163 | 0.735974249 | cat2 | from_3109 | to_3109 | art_long | art_short | J |
| 114 | 163 | 0.881833842 | cat2 | from_3110 | to_3110 | art_long | art_short | J |
| 116 | 163 | 0.637891254 | cat2 | from_3111 | to_3111 | art_long | art_short | J |
| 158 | 163 | 0.632215988 | cat2 | from_3112 | to_3112 | art_short | art_short | H |
| 162 | 163 | 0.941251815 | cat2 | from_3113 | to_3113 | art_short | art_short | H |
| 2 | 164 | 0.697881149 | cat2 | from_3114 | to_3114 | art_long | art_short | J |
| 12 | 164 | 0.94023572 | cat2 | from_3115 | to_3115 | art_long | art_short | J |
| 13 | 164 | 0.670990452 | cat2 | from_3116 | to_3116 | art_long | art_short | J |
| 15 | 164 | 0.71601335 | cat2 | from_3117 | to_3117 | art_long | art_short | J |
| 16 | 164 | 0.954221684 | cat2 | from_3118 | to_3118 | art_long | art_short | J |

| | | | | | | | | |
|---|---|---|---|---|---|---|---|---|
| 18 | 164 | 0.954911388 | cat2 | from_3119 | to_3119 | art_long | art_short | J |
| 19 | 164 | 0.851672238 | cat2 | from_3120 | to_3120 | art_long | art_short | J |
| 39 | 164 | 0.94577455 | cat2 | from_3121 | to_3121 | art_long | art_short | J |
| 40 | 164 | 0.837589239 | cat2 | from_3122 | to_3122 | art_long | art_short | J |
| 53 | 164 | 0.709843178 | cat2 | from_3123 | to_3123 | art_long | art_short | J |
| 79 | 164 | 0.845930917 | cat2 | from_3124 | to_3124 | art_long | art_short | J |
| 155 | 164 | 0.738458134 | cat2 | from_3125 | to_3125 | art_short | art_short | H |
| 162 | 164 | 0.763467889 | cat2 | from_3126 | to_3126 | art_short | art_short | H |
| 163 | 164 | 0.746641381 | cat2 | from_3127 | to_3127 | art_short | art_short | H |
| 12 | 165 | 0.784453909 | cat2 | from_3128 | to_3128 | art_long | art_short | J |
| 13 | 165 | 0.931771304 | cat2 | from_3129 | to_3129 | art_long | art_short | J |
| 15 | 165 | 0.671758465 | cat2 | from_3130 | to_3130 | art_long | art_short | J |
| 19 | 165 | 0.765052806 | cat2 | from_3131 | to_3131 | art_long | art_short | J |
| 20 | 165 | 0.749418032 | cat2 | from_3132 | to_3132 | art_long | art_short | J |
| 39 | 165 | 0.689684742 | cat2 | from_3133 | to_3133 | art_long | art_short | J |
| 40 | 165 | 0.724988218 | cat2 | from_3134 | to_3134 | art_long | art_short | J |
| 68 | 165 | 0.736014662 | cat2 | from_3135 | to_3135 | art_long | art_short | J |
| 79 | 165 | 0.802159078 | cat2 | from_3136 | to_3136 | art_long | art_short | J |
| 114 | 165 | 0.969095494 | cat2 | from_3137 | to_3137 | art_long | art_short | J |
| 156 | 165 | 0.781325564 | cat2 | from_3138 | to_3138 | art_short | art_short | H |
| 158 | 165 | 0.781200027 | cat2 | from_3139 | to_3139 | art_short | art_short | H |
| 162 | 165 | 0.790742989 | cat2 | from_3140 | to_3140 | art_short | art_short | H |
| 163 | 165 | 0.950869654 | cat2 | from_3141 | to_3141 | art_short | art_short | H |
| 164 | 165 | 0.654244296 | cat2 | from_3142 | to_3142 | art_short | art_short | H |
| 17 | 166 | 0.719215144 | cat2 | from_3143 | to_3143 | art_long | art_short | J |
| 35 | 166 | 0.961130506 | cat2 | from_3144 | to_3144 | art_long | art_short | J |
| 36 | 166 | 0.718161732 | cat2 | from_3145 | to_3145 | art_long | art_short | J |
| 38 | 166 | 0.968306524 | cat2 | from_3146 | to_3146 | art_long | art_short | J |
| 48 | 166 | 0.961123588 | cat2 | from_3147 | to_3147 | art_long | art_short | J |
| 50 | 166 | 0.843671678 | cat2 | from_3148 | to_3148 | art_long | art_short | J |
| 104 | 166 | 0.960214822 | cat2 | from_3149 | to_3149 | art_long | art_short | J |
| 132 | 166 | 0.93936711 | cat2 | from_3150 | to_3150 | art_long | art_short | J |
| 133 | 166 | 0.719431574 | cat2 | from_3151 | to_3151 | art_long | art_short | J |
| 134 | 166 | 0.716675062 | cat2 | from_3152 | to_3152 | art_long | art_short | J |
| 138 | 166 | 0.745907893 | cat2 | from_3153 | to_3153 | art_long | art_short | J |
| 143 | 166 | 0.992359507 | cat2 | from_3154 | to_3154 | art_long | art_short | J |
| 154 | 166 | 0.733414554 | cat2 | from_3155 | to_3155 | art_short | art_short | H |
| 12 | 167 | 0.999860738 | cat2 | from_3156 | to_3156 | art_long | art_short | J |
| 13 | 167 | 0.736473999 | cat2 | from_3157 | to_3157 | art_long | art_short | J |
| 16 | 167 | 0.84533064 | cat2 | from_3158 | to_3158 | art_long | art_short | J |
| 18 | 167 | 0.846092795 | cat2 | from_3159 | to_3159 | art_long | art_short | J |
| 19 | 167 | 0.947123754 | cat2 | from_3160 | to_3160 | art_long | art_short | J |
| 39 | 167 | 0.942277818 | cat2 | from_3161 | to_3161 | art_long | art_short | J |
| 40 | 167 | 0.927878674 | cat2 | from_3162 | to_3162 | art_long | art_short | J |
| 79 | 167 | 0.965617774 | cat2 | from_3163 | to_3163 | art_long | art_short | J |
| 114 | 167 | 0.738316498 | cat2 | from_3164 | to_3164 | art_long | art_short | J |
| 155 | 167 | 0.654429702 | cat2 | from_3165 | to_3165 | art_short | art_short | H |
| 162 | 167 | 0.674570686 | cat2 | from_3166 | to_3166 | art_short | art_short | H |
| 163 | 167 | 0.772401423 | cat2 | from_3167 | to_3167 | art_short | art_short | H |
| 164 | 167 | 0.942539319 | cat2 | from_3168 | to_3168 | art_short | art_short | H |
| 165 | 167 | 0.781415933 | cat2 | from_3169 | to_3169 | art_short | art_short | H |
| 2 | 168 | 0.759443912 | cat2 | from_3170 | to_3170 | art_long | art_short | J |
| 13 | 168 | 0.663722089 | cat2 | from_3171 | to_3171 | art_long | art_short | J |
| 15 | 168 | 0.652768902 | cat2 | from_3172 | to_3172 | art_long | art_short | J |
| 53 | 168 | 0.779299814 | cat2 | from_3173 | to_3173 | art_long | art_short | J |
| 155 | 168 | 0.803220229 | cat2 | from_3174 | to_3174 | art_short | art_short | H |
| 162 | 168 | 0.701392809 | cat2 | from_3175 | to_3175 | art_short | art_short | H |
| 163 | 168 | 0.715224631 | cat2 | from_3176 | to_3176 | art_short | art_short | H |
| 165 | 168 | 0.656636814 | cat2 | from_3177 | to_3177 | art_short | art_short | H |
| 12 | 169 | 0.771629702 | cat2 | from_3178 | to_3178 | art_long | art_short | J |
| 13 | 169 | 0.945330095 | cat2 | from_3179 | to_3179 | art_long | art_short | J |
| 15 | 169 | 0.862320392 | cat2 | from_3180 | to_3180 | art_long | art_short | J |
| 16 | 169 | 0.674608743 | cat2 | from_3181 | to_3181 | art_long | art_short | J |
| 18 | 169 | 0.675820686 | cat2 | from_3182 | to_3182 | art_long | art_short | J |
| 19 | 169 | 0.723067864 | cat2 | from_3183 | to_3183 | art_long | art_short | J |
| 20 | 169 | 0.689167689 | cat2 | from_3184 | to_3184 | art_long | art_short | J |
| 21 | 169 | 0.663881666 | cat2 | from_3185 | to_3185 | art_long | art_short | J |
| 39 | 169 | 0.737659449 | cat2 | from_3186 | to_3186 | art_long | art_short | J |
| 40 | 169 | 0.694939981 | cat2 | from_3187 | to_3187 | art_long | art_short | J |
| 63 | 169 | 0.638167696 | cat2 | from_3188 | to_3188 | art_long | art_short | J |
| 68 | 169 | 0.685460812 | cat2 | from_3189 | to_3189 | art_long | art_short | J |
| 79 | 169 | 0.735355731 | cat2 | from_3190 | to_3190 | art_long | art_short | J |
| 114 | 169 | 0.881580599 | cat2 | from_3191 | to_3191 | art_long | art_short | J |
| 116 | 169 | 0.640971532 | cat2 | from_3192 | to_3192 | art_long | art_short | J |
| 162 | 169 | 0.942580168 | cat2 | from_3193 | to_3193 | art_short | art_short | H |
| 163 | 169 | 0.999960279 | cat2 | from_3194 | to_3194 | art_short | art_short | H |
| 164 | 169 | 0.746964059 | cat2 | from_3195 | to_3195 | art_short | art_short | H |
| 165 | 169 | 0.949593993 | cat2 | from_3196 | to_3196 | art_short | art_short | H |
| 167 | 169 | 0.771254618 | cat2 | from_3197 | to_3197 | art_short | art_short | H |
| 168 | 169 | 0.715905605 | cat2 | from_3198 | to_3198 | art_short | art_short | H |
| 42 | 170 | 0.892456934 | cat2 | from_3199 | to_3199 | art_long | art_short | J |
| 43 | 170 | 0.910334999 | cat2 | from_3200 | to_3200 | art_long | art_short | J |
| 45 | 170 | 0.924878035 | cat2 | from_3201 | to_3201 | art_long | art_short | J |
| 62 | 170 | 0.909660395 | cat2 | from_3202 | to_3202 | art_long | art_short | J |
| 92 | 170 | 0.930883209 | cat2 | from_3203 | to_3203 | art_long | art_short | J |
| 100 | 170 | 0.909838253 | cat2 | from_3204 | to_3204 | art_long | art_short | J |
| 101 | 170 | 0.901763575 | cat2 | from_3205 | to_3205 | art_long | art_short | J |
| 107 | 170 | 0.925255792 | cat2 | from_3206 | to_3206 | art_long | art_short | J |
| 139 | 170 | 0.989287993 | cat2 | from_3207 | to_3207 | art_long | art_short | J |
| 147 | 170 | 0.899516693 | cat2 | from_3208 | to_3208 | art_long | art_short | J |
| 149 | 170 | 0.923048617 | cat2 | from_3209 | to_3209 | art_long | art_short | J |
| 42 | 171 | 0.897217397 | cat2 | from_3210 | to_3210 | art_long | art_short | J |
| 43 | 171 | 0.912574979 | cat2 | from_3211 | to_3211 | art_long | art_short | J |
| 45 | 171 | 0.928689073 | cat2 | from_3212 | to_3212 | art_long | art_short | J |
| 62 | 171 | 0.914080523 | cat2 | from_3213 | to_3213 | art_long | art_short | J |
| 92 | 171 | 0.934443361 | cat2 | from_3214 | to_3214 | art_long | art_short | J |
| 100 | 171 | 0.914260266 | cat2 | from_3215 | to_3215 | art_long | art_short | J |
| 101 | 171 | 0.90636159 | cat2 | from_3216 | to_3216 | art_long | art_short | J |
| 107 | 171 | 0.929028226 | cat2 | from_3217 | to_3217 | art_long | art_short | J |
| 139 | 171 | 0.990371084 | cat2 | from_3218 | to_3218 | art_long | art_short | J |
| 147 | 171 | 0.904157104 | cat2 | from_3219 | to_3219 | art_long | art_short | J |
| 149 | 171 | 0.926995825 | cat2 | from_3220 | to_3220 | art_long | art_short | J |
| 170 | 171 | 0.999941833 | cat2 | from_3221 | to_3221 | art_short | art_short | H |
| 22 | 172 | 0.682552729 | cat2 | from_3222 | to_3222 | art_long | art_short | J |

| | | | | | | | | |
|---|---|---|---|---|---|---|---|---|
| 27 | 172 | 0.963719724 | cat2 | from_3223 | to_3223 | art_long | art_short | J |
| 64 | 172 | 0.737706772 | cat2 | from_3224 | to_3224 | art_long | art_short | J |
| 91 | 172 | 0.951647082 | cat2 | from_3225 | to_3225 | art_long | art_short | J |
| 117 | 172 | 0.735525186 | cat2 | from_3226 | to_3226 | art_long | art_short | J |
| 161 | 172 | 0.685590896 | cat2 | from_3227 | to_3227 | art_short | art_short | H |
| 14 | 173 | 0.940865111 | cat2 | from_3228 | to_3228 | art_long | art_short | J |
| 24 | 173 | 0.937547233 | cat2 | from_3229 | to_3229 | art_long | art_short | J |
| 25 | 173 | 0.773380181 | cat2 | from_3230 | to_3230 | art_long | art_short | J |
| 28 | 173 | 0.941834014 | cat2 | from_3231 | to_3231 | art_long | art_short | J |
| 30 | 173 | 0.637496986 | cat2 | from_3232 | to_3232 | art_long | art_short | J |
| 31 | 173 | 0.704820438 | cat2 | from_3233 | to_3233 | art_long | art_short | J |
| 33 | 173 | 0.717380631 | cat2 | from_3234 | to_3234 | art_long | art_short | J |
| 59 | 173 | 0.93937863 | cat2 | from_3235 | to_3235 | art_long | art_short | J |
| 73 | 173 | 0.938829948 | cat2 | from_3236 | to_3236 | art_long | art_short | J |
| 74 | 173 | 0.93809264 | cat2 | from_3237 | to_3237 | art_long | art_short | J |
| 76 | 173 | 0.921994416 | cat2 | from_3238 | to_3238 | art_long | art_short | J |
| 78 | 173 | 0.942241844 | cat2 | from_3239 | to_3239 | art_long | art_short | J |
| 90 | 173 | 0.923786172 | cat2 | from_3240 | to_3240 | art_long | art_short | J |
| 97 | 173 | 0.938436892 | cat2 | from_3241 | to_3241 | art_long | art_short | J |
| 110 | 173 | 0.992585955 | cat2 | from_3242 | to_3242 | art_long | art_short | J |
| 112 | 173 | 0.766816342 | cat2 | from_3243 | to_3243 | art_long | art_short | J |
| 122 | 173 | 0.938991544 | cat2 | from_3244 | to_3244 | art_long | art_short | J |
| 14 | 174 | 0.923667043 | cat2 | from_3245 | to_3245 | art_long | art_short | J |
| 24 | 174 | 0.919958871 | cat2 | from_3246 | to_3246 | art_long | art_short | J |
| 25 | 174 | 0.770633732 | cat2 | from_3247 | to_3247 | art_long | art_short | J |
| 28 | 174 | 0.924816607 | cat2 | from_3248 | to_3248 | art_long | art_short | J |
| 30 | 174 | 0.63786069 | cat2 | from_3249 | to_3249 | art_long | art_short | J |
| 31 | 174 | 0.718472011 | cat2 | from_3250 | to_3250 | art_long | art_short | J |
| 33 | 174 | 0.729050981 | cat2 | from_3251 | to_3251 | art_long | art_short | J |
| 59 | 174 | 0.922046209 | cat2 | from_3252 | to_3252 | art_long | art_short | J |
| 73 | 174 | 0.92136412 | cat2 | from_3253 | to_3253 | art_long | art_short | J |
| 74 | 174 | 0.920553634 | cat2 | from_3254 | to_3254 | art_long | art_short | J |
| 76 | 174 | 0.912337432 | cat2 | from_3255 | to_3255 | art_long | art_short | J |
| 78 | 174 | 0.927181928 | cat2 | from_3256 | to_3256 | art_long | art_short | J |
| 90 | 174 | 0.923002573 | cat2 | from_3257 | to_3257 | art_long | art_short | J |
| 97 | 174 | 0.920931026 | cat2 | from_3258 | to_3258 | art_long | art_short | J |
| 110 | 174 | 0.987024673 | cat2 | from_3259 | to_3259 | art_long | art_short | J |
| 112 | 174 | 0.763266953 | cat2 | from_3260 | to_3260 | art_long | art_short | J |
| 122 | 174 | 0.921542942 | cat2 | from_3261 | to_3261 | art_long | art_short | J |
| 173 | 174 | 0.99886365 | cat2 | from_3262 | to_3262 | art_short | art_short | H |
| 3 | 175 | 0.694057638 | cat2 | from_3263 | to_3263 | art_long | art_short | J |
| 29 | 175 | 0.716522365 | cat2 | from_3264 | to_3264 | art_long | art_short | J |
| 71 | 175 | 0.739892694 | cat2 | from_3265 | to_3265 | art_long | art_short | J |
| 82 | 175 | 0.963076609 | cat2 | from_3266 | to_3266 | art_long | art_short | J |
| 96 | 175 | 0.656004496 | cat2 | from_3267 | to_3267 | art_long | art_short | J |
| 98 | 175 | 0.717500654 | cat2 | from_3268 | to_3268 | art_long | art_short | J |
| 121 | 175 | 0.717970657 | cat2 | from_3269 | to_3269 | art_long | art_short | J |
| 150 | 175 | 0.893692121 | cat2 | from_3270 | to_3270 | art_long | art_short | J |
| 44 | 176 | 0.740167596 | cat2 | from_3271 | to_3271 | art_long | art_short | J |
| 46 | 176 | 0.755255743 | cat2 | from_3272 | to_3272 | art_long | art_short | J |
| 52 | 176 | 0.732119279 | cat2 | from_3273 | to_3273 | art_long | art_short | J |
| 60 | 176 | 0.804571739 | cat2 | from_3274 | to_3274 | art_long | art_short | J |
| 61 | 176 | 0.916151605 | cat2 | from_3275 | to_3275 | art_long | art_short | J |
| 99 | 176 | 0.896053176 | cat2 | from_3276 | to_3276 | art_long | art_short | J |
| 102 | 176 | 0.745769164 | cat2 | from_3277 | to_3277 | art_long | art_short | J |
| 105 | 176 | 0.912227572 | cat2 | from_3278 | to_3278 | art_long | art_short | J |
| 106 | 176 | 0.696828226 | cat2 | from_3279 | to_3279 | art_long | art_short | J |
| 128 | 176 | 0.766074187 | cat2 | from_3280 | to_3280 | art_long | art_short | J |
| 135 | 176 | 0.755088545 | cat2 | from_3281 | to_3281 | art_long | art_short | J |
| 148 | 176 | 0.869685647 | cat2 | from_3282 | to_3282 | art_long | art_short | J |
| 17 | 177 | 0.877297173 | cat2 | from_3283 | to_3283 | art_long | art_short | J |
| 35 | 177 | 0.691224574 | cat2 | from_3284 | to_3284 | art_long | art_short | J |
| 36 | 177 | 0.874511082 | cat2 | from_3285 | to_3285 | art_long | art_short | J |
| 38 | 177 | 0.701631789 | cat2 | from_3286 | to_3286 | art_long | art_short | J |
| 48 | 177 | 0.69130403 | cat2 | from_3287 | to_3287 | art_long | art_short | J |
| 91 | 177 | -0.667565986 | cat2 | from_3288 | to_3288 | art_long | art_short | J |
| 104 | 177 | 0.694086707 | cat2 | from_3289 | to_3289 | art_long | art_short | J |
| 132 | 177 | 0.654453254 | cat2 | from_3290 | to_3290 | art_long | art_short | J |
| 133 | 177 | 0.872931448 | cat2 | from_3291 | to_3291 | art_long | art_short | J |
| 134 | 177 | 0.875154541 | cat2 | from_3292 | to_3292 | art_long | art_short | J |
| 138 | 177 | 0.932901954 | cat2 | from_3293 | to_3293 | art_long | art_short | J |
| 143 | 177 | 0.771595025 | cat2 | from_3294 | to_3294 | art_long | art_short | J |
| 154 | 177 | 0.991377159 | cat2 | from_3295 | to_3295 | art_short | art_short | H |
| 166 | 177 | 0.756996437 | cat2 | from_3296 | to_3296 | art_short | art_short | H |
| 42 | 178 | 0.813294488 | cat2 | from_3297 | to_3297 | art_long | art_short | J |
| 43 | 178 | 0.6928738 | cat2 | from_3298 | to_3298 | art_long | art_short | J |
| 45 | 178 | 0.79118318 | cat2 | from_3299 | to_3299 | art_long | art_short | J |
| 52 | 178 | 0.700490788 | cat2 | from_3300 | to_3300 | art_long | art_short | J |
| 62 | 178 | 0.817783493 | cat2 | from_3301 | to_3301 | art_long | art_short | J |
| 92 | 178 | 0.792411382 | cat2 | from_3302 | to_3302 | art_long | art_short | J |
| 100 | 178 | 0.809289836 | cat2 | from_3303 | to_3303 | art_long | art_short | J |
| 101 | 178 | 0.812014194 | cat2 | from_3304 | to_3304 | art_long | art_short | J |
| 107 | 178 | 0.789657975 | cat2 | from_3305 | to_3305 | art_long | art_short | J |
| 139 | 178 | 0.924857852 | cat2 | from_3306 | to_3306 | art_long | art_short | J |
| 147 | 178 | 0.812469506 | cat2 | from_3307 | to_3307 | art_long | art_short | J |
| 149 | 178 | 0.796232399 | cat2 | from_3308 | to_3308 | art_long | art_short | J |
| 170 | 178 | 0.910255487 | cat2 | from_3309 | to_3309 | art_short | art_short | H |
| 171 | 178 | 0.909766645 | cat2 | from_3310 | to_3310 | art_short | art_short | H |
| 2 | 179 | 0.951443891 | cat2 | from_3311 | to_3311 | art_long | art_short | J |
| 12 | 179 | 0.668804153 | cat2 | from_3312 | to_3312 | art_long | art_short | J |
| 16 | 179 | 0.707346793 | cat2 | from_3313 | to_3313 | art_long | art_short | J |
| 18 | 179 | 0.707886843 | cat2 | from_3314 | to_3314 | art_long | art_short | J |
| 39 | 179 | 0.671536293 | cat2 | from_3315 | to_3315 | art_long | art_short | J |
| 53 | 179 | 0.96523256 | cat2 | from_3316 | to_3316 | art_long | art_short | J |
| 155 | 179 | 0.997664942 | cat2 | from_3317 | to_3317 | art_short | art_short | H |
| 164 | 179 | 0.74197925 | cat2 | from_3318 | to_3318 | art_short | art_short | H |
| 167 | 179 | 0.67035417 | cat2 | from_3319 | to_3319 | art_short | art_short | H |
| 168 | 179 | 0.806902407 | cat2 | from_3320 | to_3320 | art_short | art_short | H |
| 2 | 180 | 0.811665258 | cat2 | from_3321 | to_3321 | art_long | art_short | J |
| 15 | 180 | 0.71184392 | cat2 | from_3322 | to_3322 | art_long | art_short | J |
| 53 | 180 | 0.820412079 | cat2 | from_3323 | to_3323 | art_long | art_short | J |
| 155 | 180 | 0.829013577 | cat2 | from_3324 | to_3324 | art_short | art_short | H |
| 162 | 180 | 0.729943902 | cat2 | from_3325 | to_3325 | art_short | art_short | H |
| 163 | 180 | 0.646072167 | cat2 | from_3326 | to_3326 | art_short | art_short | H |

| | | | | | | | | |
|---|---|---|---|---|---|---|---|---|
| 168 | 180 | 0.955562597 | cat2 | from_3327 | to_3327 | art_short | art_short | H |
| 169 | 180 | 0.647307574 | cat2 | from_3328 | to_3328 | art_short | art_short | H |
| 179 | 180 | 0.82500148 | cat2 | from_3329 | to_3329 | art_short | art_short | H |
| 3 | 181 | 0.767315514 | cat2 | from_3330 | to_3330 | art_long | art_short | J |
| 29 | 181 | 0.886806829 | cat2 | from_3331 | to_3331 | art_long | art_short | J |
| 41 | 181 | 0.703710912 | cat2 | from_3332 | to_3332 | art_long | art_short | J |
| 66 | 181 | 0.719584956 | cat2 | from_3333 | to_3333 | art_long | art_short | J |
| 67 | 181 | 0.717739665 | cat2 | from_3334 | to_3334 | art_long | art_short | J |
| 71 | 181 | 0.911155237 | cat2 | from_3335 | to_3335 | art_long | art_short | J |
| 82 | 181 | 0.846150507 | cat2 | from_3336 | to_3336 | art_long | art_short | J |
| 96 | 181 | 0.755300218 | cat2 | from_3337 | to_3337 | art_long | art_short | J |
| 124 | 181 | 0.712475877 | cat2 | from_3338 | to_3338 | art_long | art_short | J |
| 150 | 181 | 0.807764788 | cat2 | from_3339 | to_3339 | art_long | art_short | J |
| 175 | 181 | 0.861322501 | cat2 | from_3340 | to_3340 | art_short | art_short | H |
| 22 | 182 | 0.677136421 | cat2 | from_3341 | to_3341 | art_long | art_short | J |
| 32 | 182 | -0.635636079 | cat2 | from_3342 | to_3342 | art_long | art_short | J |
| 49 | 182 | 0.700848917 | cat2 | from_3343 | to_3343 | art_long | art_short | J |
| 56 | 182 | 0.966652171 | cat2 | from_3344 | to_3344 | art_long | art_short | J |
| 57 | 182 | 0.966556939 | cat2 | from_3345 | to_3345 | art_long | art_short | J |
| 70 | 182 | 0.75099671 | cat2 | from_3346 | to_3346 | art_long | art_short | J |
| 75 | 182 | 0.696679187 | cat2 | from_3347 | to_3347 | art_long | art_short | J |
| 85 | 182 | 0.756926215 | cat2 | from_3348 | to_3348 | art_long | art_short | J |
| 123 | 182 | 0.754538997 | cat2 | from_3349 | to_3349 | art_long | art_short | J |
| 146 | 182 | 0.760174106 | cat2 | from_3350 | to_3350 | art_long | art_short | J |
| 161 | 182 | 0.6747624 | cat2 | from_3351 | to_3351 | art_short | art_short | H |
| 44 | 183 | 0.74460609 | cat2 | from_3352 | to_3352 | art_long | art_short | J |
| 46 | 183 | 0.754458179 | cat2 | from_3353 | to_3353 | art_long | art_short | J |
| 52 | 183 | 0.734088398 | cat2 | from_3354 | to_3354 | art_long | art_short | J |
| 60 | 183 | 0.802080775 | cat2 | from_3355 | to_3355 | art_long | art_short | J |
| 61 | 183 | 0.911072696 | cat2 | from_3356 | to_3356 | art_long | art_short | J |
| 99 | 183 | 0.899766544 | cat2 | from_3357 | to_3357 | art_long | art_short | J |
| 102 | 183 | 0.753776087 | cat2 | from_3358 | to_3358 | art_long | art_short | J |
| 105 | 183 | 0.915546693 | cat2 | from_3359 | to_3359 | art_long | art_short | J |
| 106 | 183 | 0.709575106 | cat2 | from_3360 | to_3360 | art_long | art_short | J |
| 128 | 183 | 0.763879593 | cat2 | from_3361 | to_3361 | art_long | art_short | J |
| 135 | 183 | 0.754373977 | cat2 | from_3362 | to_3362 | art_long | art_short | J |
| 148 | 183 | 0.87888082 | cat2 | from_3363 | to_3363 | art_long | art_short | J |
| 176 | 183 | 0.998845309 | cat2 | from_3364 | to_3364 | art_short | art_short | H |
| 42 | 184 | 0.918022216 | cat2 | from_3365 | to_3365 | art_long | art_short | J |
| 43 | 184 | 0.915632652 | cat2 | from_3366 | to_3366 | art_long | art_short | J |
| 45 | 184 | 0.936275205 | cat2 | from_3367 | to_3367 | art_long | art_short | J |
| 62 | 184 | 0.931356982 | cat2 | from_3368 | to_3368 | art_long | art_short | J |
| 92 | 184 | 0.940110559 | cat2 | from_3369 | to_3369 | art_long | art_short | J |
| 100 | 184 | 0.930101888 | cat2 | from_3370 | to_3370 | art_long | art_short | J |
| 101 | 184 | 0.924742641 | cat2 | from_3371 | to_3371 | art_long | art_short | J |
| 107 | 184 | 0.936117427 | cat2 | from_3372 | to_3372 | art_long | art_short | J |
| 139 | 184 | 0.997293583 | cat2 | from_3373 | to_3373 | art_long | art_short | J |
| 147 | 184 | 0.923162874 | cat2 | from_3374 | to_3374 | art_long | art_short | J |
| 149 | 184 | 0.936356265 | cat2 | from_3375 | to_3375 | art_long | art_short | J |
| 170 | 184 | 0.997327806 | cat2 | from_3376 | to_3376 | art_short | art_short | H |
| 171 | 184 | 0.997868998 | cat2 | from_3377 | to_3377 | art_short | art_short | H |
| 178 | 184 | 0.919075113 | cat2 | from_3378 | to_3378 | art_short | art_short | H |
| 3 | 185 | 0.85751424 | cat2 | from_3379 | to_3379 | art_long | art_short | J |
| 82 | 185 | 0.802733578 | cat2 | from_3380 | to_3380 | art_long | art_short | J |
| 150 | 185 | 0.761172699 | cat2 | from_3381 | to_3381 | art_long | art_short | J |
| 175 | 185 | 0.819238101 | cat2 | from_3382 | to_3382 | art_short | art_short | H |
| 181 | 185 | 0.651029535 | cat2 | from_3383 | to_3383 | art_short | art_short | H |
| 5 | 186 | 0.739749617 | cat2 | from_3384 | to_3384 | art_long | art_short | J |
| 13 | 186 | 0.702998204 | cat2 | from_3385 | to_3385 | art_long | art_short | J |
| 63 | 186 | 0.864668499 | cat2 | from_3386 | to_3386 | art_long | art_short | J |
| 114 | 186 | 0.719768992 | cat2 | from_3387 | to_3387 | art_long | art_short | J |
| 116 | 186 | 0.870911006 | cat2 | from_3388 | to_3388 | art_long | art_short | J |
| 125 | 186 | 0.639976691 | cat2 | from_3389 | to_3389 | art_long | art_short | J |
| 156 | 186 | 0.908878361 | cat2 | from_3390 | to_3390 | art_short | art_short | H |
| 158 | 186 | 0.910087404 | cat2 | from_3391 | to_3391 | art_short | art_short | H |
| 163 | 186 | 0.719242715 | cat2 | from_3392 | to_3392 | art_short | art_short | H |
| 165 | 186 | 0.745305366 | cat2 | from_3393 | to_3393 | art_short | art_short | H |
| 169 | 186 | 0.718700859 | cat2 | from_3394 | to_3394 | art_short | art_short | H |
| 2 | 187 | 0.819948049 | cat2 | from_3395 | to_3395 | art_long | art_short | J |
| 15 | 187 | 0.723205754 | cat2 | from_3396 | to_3396 | art_long | art_short | J |
| 53 | 187 | 0.828961876 | cat2 | from_3397 | to_3397 | art_long | art_short | J |
| 155 | 187 | 0.82262212 | cat2 | from_3398 | to_3398 | art_short | art_short | H |
| 162 | 187 | 0.726875869 | cat2 | from_3399 | to_3399 | art_short | art_short | H |
| 163 | 187 | 0.636431102 | cat2 | from_3400 | to_3400 | art_short | art_short | H |
| 168 | 187 | 0.9530702 | cat2 | from_3401 | to_3401 | art_short | art_short | H |
| 169 | 187 | 0.638369426 | cat2 | from_3402 | to_3402 | art_short | art_short | H |
| 179 | 187 | 0.814516271 | cat2 | from_3403 | to_3403 | art_short | art_short | H |
| 180 | 187 | 0.995654061 | cat2 | from_3404 | to_3404 | art_short | art_short | H |
| 4 | 188 | 0.85336149 | cat2 | from_3405 | to_3405 | art_long | art_short | J |
| 6 | 188 | 0.85857634 | cat2 | from_3406 | to_3406 | art_long | art_short | J |
| 7 | 188 | 0.856680956 | cat2 | from_3407 | to_3407 | art_long | art_short | J |
| 11 | 188 | 0.696398212 | cat2 | from_3408 | to_3408 | art_long | art_short | J |
| 113 | 188 | 0.659694177 | cat2 | from_3409 | to_3409 | art_long | art_short | J |
| 118 | 188 | -0.642852195 | cat2 | from_3410 | to_3410 | art_long | art_short | J |
| 131 | 188 | -0.64063615 | cat2 | from_3411 | to_3411 | art_long | art_short | J |
| 137 | 188 | -0.656000255 | cat2 | from_3412 | to_3412 | art_long | art_short | J |
| 141 | 188 | -0.654770363 | cat2 | from_3413 | to_3413 | art_long | art_short | J |
| 144 | 188 | -0.65059297 | cat2 | from_3414 | to_3414 | art_long | art_short | J |
| 159 | 188 | 0.998907809 | cat2 | from_3415 | to_3415 | art_short | art_short | H |
| 160 | 188 | 0.942166752 | cat2 | from_3416 | to_3416 | art_short | art_short | H |
| 29 | 189 | 0.776893803 | cat2 | from_3417 | to_3417 | art_long | art_short | J |
| 66 | 189 | 0.918972253 | cat2 | from_3418 | to_3418 | art_long | art_short | J |
| 67 | 189 | 0.920330753 | cat2 | from_3419 | to_3419 | art_long | art_short | J |
| 71 | 189 | 0.788430229 | cat2 | from_3420 | to_3420 | art_long | art_short | J |
| 82 | 189 | 0.717177004 | cat2 | from_3421 | to_3421 | art_long | art_short | J |
| 150 | 189 | 0.687749957 | cat2 | from_3422 | to_3422 | art_long | art_short | J |
| 175 | 189 | 0.727894263 | cat2 | from_3423 | to_3423 | art_short | art_short | H |
| 181 | 189 | 0.815840721 | cat2 | from_3424 | to_3424 | art_short | art_short | H |
| 24 | 190 | 0.634272367 | cat2 | from_3425 | to_3425 | art_long | art_short | J |
| 28 | 190 | 0.635870498 | cat2 | from_3426 | to_3426 | art_long | art_short | J |
| 32 | 190 | 0.939499952 | cat2 | from_3427 | to_3427 | art_long | art_short | J |
| 59 | 190 | 0.635319879 | cat2 | from_3428 | to_3428 | art_long | art_short | J |
| 73 | 190 | 0.633117087 | cat2 | from_3429 | to_3429 | art_long | art_short | J |
| 74 | 190 | 0.633819026 | cat2 | from_3430 | to_3430 | art_long | art_short | J |

| | | | | | | | | |
|---|---|---|---|---|---|---|---|---|
| 86 | 190 | 0.676301568 | cat2 | from_3431 | to_3431 | art_long | art_short | J |
| 94 | 190 | 0.674239358 | cat2 | from_3432 | to_3432 | art_long | art_short | J |
| 97 | 190 | 0.633505453 | cat2 | from_3433 | to_3433 | art_long | art_short | J |
| 110 | 190 | 0.644488648 | cat2 | from_3434 | to_3434 | art_long | art_short | J |
| 122 | 190 | 0.632946846 | cat2 | from_3435 | to_3435 | art_long | art_short | J |
| 24 | 191 | 0.632405661 | cat2 | from_3436 | to_3436 | art_long | art_short | J |
| 28 | 191 | 0.634099537 | cat2 | from_3437 | to_3437 | art_long | art_short | J |
| 32 | 191 | 0.936328374 | cat2 | from_3438 | to_3438 | art_long | art_short | J |
| 59 | 191 | 0.633410958 | cat2 | from_3439 | to_3439 | art_long | art_short | J |
| 74 | 191 | 0.632114599 | cat2 | from_3440 | to_3440 | art_long | art_short | J |
| 86 | 191 | 0.674221932 | cat2 | from_3441 | to_3441 | art_long | art_short | J |
| 94 | 191 | 0.673368498 | cat2 | from_3442 | to_3442 | art_long | art_short | J |
| 97 | 191 | 0.631908904 | cat2 | from_3443 | to_3443 | art_long | art_short | J |
| 110 | 191 | 0.644803254 | cat2 | from_3444 | to_3444 | art_long | art_short | J |
| 173 | 191 | 0.632549834 | cat2 | from_3445 | to_3445 | art_short | art_short | H |
| 190 | 191 | 0.999588542 | cat2 | from_3446 | to_3446 | art_short | art_short | H |
| 12 | 192 | 0.695511927 | cat2 | from_3447 | to_3447 | art_long | art_short | J |
| 13 | 192 | 0.697617922 | cat2 | from_3448 | to_3448 | art_long | art_short | J |
| 15 | 192 | 0.854595815 | cat2 | from_3449 | to_3449 | art_long | art_short | J |
| 16 | 192 | 0.822572539 | cat2 | from_3450 | to_3450 | art_long | art_short | J |
| 18 | 192 | 0.82378676 | cat2 | from_3451 | to_3451 | art_long | art_short | J |
| 39 | 192 | 0.75709808 | cat2 | from_3452 | to_3452 | art_long | art_short | J |
| 68 | 192 | 0.690473477 | cat2 | from_3453 | to_3453 | art_long | art_short | J |
| 162 | 192 | 0.859833936 | cat2 | from_3454 | to_3454 | art_short | art_short | H |
| 163 | 192 | 0.774602135 | cat2 | from_3455 | to_3455 | art_short | art_short | H |
| 164 | 192 | 0.818003766 | cat2 | from_3456 | to_3456 | art_short | art_short | H |
| 167 | 192 | 0.698284327 | cat2 | from_3457 | to_3457 | art_short | art_short | H |
| 169 | 192 | 0.776716606 | cat2 | from_3458 | to_3458 | art_short | art_short | H |
| 8 | 193 | 0.693992758 | cat2 | from_3459 | to_3459 | art_long | art_short | J |
| 49 | 193 | 0.865355001 | cat2 | from_3460 | to_3460 | art_long | art_short | J |
| 70 | 193 | 0.718326121 | cat2 | from_3461 | to_3461 | art_long | art_short | J |
| 75 | 193 | 0.865654641 | cat2 | from_3462 | to_3462 | art_long | art_short | J |
| 84 | 193 | 0.956010533 | cat2 | from_3463 | to_3463 | art_long | art_short | J |
| 125 | 193 | 0.74675349 | cat2 | from_3464 | to_3464 | art_long | art_short | J |
| 146 | 193 | 0.710674186 | cat2 | from_3465 | to_3465 | art_long | art_short | J |
| 14 | 194 | 0.941421839 | cat2 | from_3466 | to_3466 | art_long | art_short | J |
| 24 | 194 | 0.942187357 | cat2 | from_3467 | to_3467 | art_long | art_short | J |
| 25 | 194 | 0.781855119 | cat2 | from_3468 | to_3468 | art_long | art_short | J |
| 28 | 194 | 0.946508725 | cat2 | from_3469 | to_3469 | art_long | art_short | J |
| 30 | 194 | 0.635218442 | cat2 | from_3470 | to_3470 | art_long | art_short | J |
| 31 | 194 | 0.722174476 | cat2 | from_3471 | to_3471 | art_long | art_short | J |
| 33 | 194 | 0.734517768 | cat2 | from_3472 | to_3472 | art_long | art_short | J |
| 59 | 194 | 0.94433291 | cat2 | from_3473 | to_3473 | art_long | art_short | J |
| 73 | 194 | 0.942148303 | cat2 | from_3474 | to_3474 | art_long | art_short | J |
| 74 | 194 | 0.94219562 | cat2 | from_3475 | to_3475 | art_long | art_short | J |
| 76 | 194 | 0.915683622 | cat2 | from_3476 | to_3476 | art_long | art_short | J |
| 78 | 194 | 0.929046156 | cat2 | from_3477 | to_3477 | art_long | art_short | J |
| 90 | 194 | 0.935135166 | cat2 | from_3478 | to_3478 | art_long | art_short | J |
| 97 | 194 | 0.942182827 | cat2 | from_3479 | to_3479 | art_long | art_short | J |
| 110 | 194 | 0.997062535 | cat2 | from_3480 | to_3480 | art_long | art_short | J |
| 112 | 194 | 0.779437161 | cat2 | from_3481 | to_3481 | art_long | art_short | J |
| 122 | 194 | 0.942127168 | cat2 | from_3482 | to_3482 | art_long | art_short | J |
| 173 | 194 | 0.995878772 | cat2 | from_3483 | to_3483 | art_short | art_short | H |
| 174 | 194 | 0.993926786 | cat2 | from_3484 | to_3484 | art_short | art_short | H |
| 190 | 194 | 0.638797573 | cat2 | from_3485 | to_3485 | art_short | art_short | H |
| 191 | 194 | 0.639820418 | cat2 | from_3486 | to_3486 | art_short | art_short | H |
| 30 | 195 | 0.99568176 | cat2 | from_3487 | to_3487 | art_long | art_short | J |
| 47 | 195 | 0.923103803 | cat2 | from_3488 | to_3488 | art_long | art_short | J |
| 54 | 195 | 0.77654315 | cat2 | from_3489 | to_3489 | art_long | art_short | J |
| 55 | 195 | 0.923642448 | cat2 | from_3490 | to_3490 | art_long | art_short | J |
| 58 | 195 | 0.927282932 | cat2 | from_3491 | to_3491 | art_long | art_short | J |
| 69 | 195 | 0.932734462 | cat2 | from_3492 | to_3492 | art_long | art_short | J |
| 72 | 195 | 0.912481362 | cat2 | from_3493 | to_3493 | art_long | art_short | J |
| 77 | 195 | 0.910124917 | cat2 | from_3494 | to_3494 | art_long | art_short | J |
| 80 | 195 | 0.910473497 | cat2 | from_3495 | to_3495 | art_long | art_short | J |
| 81 | 195 | 0.778043536 | cat2 | from_3496 | to_3496 | art_long | art_short | J |
| 83 | 195 | 0.913328457 | cat2 | from_3497 | to_3497 | art_long | art_short | J |
| 87 | 195 | 0.927335115 | cat2 | from_3498 | to_3498 | art_long | art_short | J |
| 95 | 195 | 0.926287462 | cat2 | from_3499 | to_3499 | art_long | art_short | J |
| 109 | 195 | 0.929843865 | cat2 | from_3500 | to_3500 | art_long | art_short | J |
| 111 | 195 | 0.929703184 | cat2 | from_3501 | to_3501 | art_long | art_short | J |
| 119 | 195 | 0.927247054 | cat2 | from_3502 | to_3502 | art_long | art_short | J |
| 120 | 195 | 0.913977934 | cat2 | from_3503 | to_3503 | art_long | art_short | J |
| 140 | 195 | 0.915933927 | cat2 | from_3504 | to_3504 | art_long | art_short | J |
| 173 | 195 | 0.633330833 | cat2 | from_3505 | to_3505 | art_short | art_short | H |
| 174 | 195 | 0.638117642 | cat2 | from_3506 | to_3506 | art_short | art_short | H |
| 5 | 196 | 0.642326227 | cat2 | from_3507 | to_3507 | art_long | art_short | J |
| 13 | 196 | 0.670133435 | cat2 | from_3508 | to_3508 | art_long | art_short | J |
| 15 | 196 | 0.889969049 | cat2 | from_3509 | to_3509 | art_long | art_short | J |
| 16 | 196 | 0.679892936 | cat2 | from_3510 | to_3510 | art_long | art_short | J |
| 18 | 196 | 0.680181762 | cat2 | from_3511 | to_3511 | art_long | art_short | J |
| 162 | 196 | 0.95654995 | cat2 | from_3512 | to_3512 | art_short | art_short | H |
| 163 | 196 | 0.850623812 | cat2 | from_3513 | to_3513 | art_short | art_short | H |
| 164 | 196 | 0.728646289 | cat2 | from_3514 | to_3514 | art_short | art_short | H |
| 165 | 196 | 0.667906149 | cat2 | from_3515 | to_3515 | art_short | art_short | H |
| 168 | 196 | 0.644136292 | cat2 | from_3516 | to_3516 | art_short | art_short | H |
| 169 | 196 | 0.850573435 | cat2 | from_3517 | to_3517 | art_short | art_short | H |
| 180 | 196 | 0.72673418 | cat2 | from_3518 | to_3518 | art_short | art_short | H |
| 187 | 196 | 0.711608652 | cat2 | from_3519 | to_3519 | art_short | art_short | H |
| 192 | 196 | 0.824217456 | cat2 | from_3520 | to_3520 | art_short | art_short | H |
| 2 | 197 | 0.638644558 | cat2 | from_3521 | to_3521 | art_long | art_short | J |
| 12 | 197 | 0.953335323 | cat2 | from_3522 | to_3522 | art_long | art_short | J |
| 13 | 197 | 0.639029858 | cat2 | from_3523 | to_3523 | art_long | art_short | J |
| 15 | 197 | 0.652071566 | cat2 | from_3524 | to_3524 | art_long | art_short | J |
| 16 | 197 | 0.901898011 | cat2 | from_3525 | to_3525 | art_long | art_short | J |
| 18 | 197 | 0.901866204 | cat2 | from_3526 | to_3526 | art_long | art_short | J |
| 19 | 197 | 0.835975451 | cat2 | from_3527 | to_3527 | art_long | art_short | J |
| 39 | 197 | 0.906018069 | cat2 | from_3528 | to_3528 | art_long | art_short | J |
| 40 | 197 | 0.836395343 | cat2 | from_3529 | to_3529 | art_long | art_short | J |
| 53 | 197 | 0.654111847 | cat2 | from_3530 | to_3530 | art_long | art_short | J |
| 79 | 197 | 0.84918709 | cat2 | from_3531 | to_3531 | art_long | art_short | J |
| 155 | 197 | 0.715825541 | cat2 | from_3532 | to_3532 | art_short | art_short | H |
| 162 | 197 | 0.737254118 | cat2 | from_3533 | to_3533 | art_short | art_short | H |
| 163 | 197 | 0.745283897 | cat2 | from_3534 | to_3534 | art_short | art_short | H |

| | | | | | | | | |
|---|---|---|---|---|---|---|---|---|
| 164 | 197 | 0.983927356 | cat2 | from_3535 | to_3535 | art_short | art_short | H |
| 165 | 197 | 0.674857839 | cat2 | from_3536 | to_3536 | art_short | art_short | H |
| 167 | 197 | 0.954829967 | cat2 | from_3537 | to_3537 | art_short | art_short | H |
| 169 | 197 | 0.744037461 | cat2 | from_3538 | to_3538 | art_short | art_short | H |
| 179 | 197 | 0.729943127 | cat2 | from_3539 | to_3539 | art_short | art_short | H |
| 192 | 197 | 0.771249202 | cat2 | from_3540 | to_3540 | art_short | art_short | H |
| 196 | 197 | 0.728842726 | cat2 | from_3541 | to_3541 | art_short | art_short | H |
| 6 | 198 | 0.657930269 | cat2 | from_3542 | to_3542 | art_long | art_short | J |
| 8 | 198 | 0.752746017 | cat2 | from_3543 | to_3543 | art_long | art_short | J |
| 9 | 198 | 0.688084349 | cat2 | from_3544 | to_3544 | art_long | art_short | J |
| 10 | 198 | 0.689998799 | cat2 | from_3545 | to_3545 | art_long | art_short | J |
| 65 | 198 | 0.712160668 | cat2 | from_3546 | to_3546 | art_long | art_short | J |
| 84 | 198 | 0.683783164 | cat2 | from_3547 | to_3547 | art_long | art_short | J |
| 113 | 198 | 0.680710786 | cat2 | from_3548 | to_3548 | art_long | art_short | J |
| 125 | 198 | 0.88722659 | cat2 | from_3549 | to_3549 | art_long | art_short | J |
| 156 | 198 | 0.76277318 | cat2 | from_3550 | to_3550 | art_short | art_short | H |
| 157 | 198 | 0.781250131 | cat2 | from_3551 | to_3551 | art_short | art_short | H |
| 158 | 198 | 0.761389912 | cat2 | from_3552 | to_3552 | art_short | art_short | H |
| 186 | 198 | 0.666590051 | cat2 | from_3553 | to_3553 | art_short | art_short | H |
| 193 | 198 | 0.766074373 | cat2 | from_3554 | to_3554 | art_short | art_short | H |
| 49 | 199 | 0.946036768 | cat2 | from_3555 | to_3555 | art_long | art_short | J |
| 56 | 199 | 0.731903942 | cat2 | from_3556 | to_3556 | art_long | art_short | J |
| 57 | 199 | 0.732691164 | cat2 | from_3557 | to_3557 | art_long | art_short | J |
| 70 | 199 | 0.944822135 | cat2 | from_3558 | to_3558 | art_long | art_short | J |
| 75 | 199 | 0.942871812 | cat2 | from_3559 | to_3559 | art_long | art_short | J |
| 84 | 199 | 0.759008906 | cat2 | from_3560 | to_3560 | art_long | art_short | J |
| 85 | 199 | 0.874804441 | cat2 | from_3561 | to_3561 | art_long | art_short | J |
| 86 | 199 | -0.65241145 | cat2 | from_3562 | to_3562 | art_long | art_short | J |
| 94 | 199 | -0.657936687 | cat2 | from_3563 | to_3563 | art_long | art_short | J |
| 113 | 199 | 0.648178441 | cat2 | from_3564 | to_3564 | art_long | art_short | J |
| 123 | 199 | 0.87517234 | cat2 | from_3565 | to_3565 | art_long | art_short | J |
| 146 | 199 | 0.949902308 | cat2 | from_3566 | to_3566 | art_long | art_short | J |
| 182 | 199 | 0.778676555 | cat2 | from_3567 | to_3567 | art_short | art_short | H |
| 193 | 199 | 0.83282391 | cat2 | from_3568 | to_3568 | art_short | art_short | H |
| 30 | 200 | 0.995087905 | cat2 | from_3569 | to_3569 | art_long | art_short | J |
| 47 | 200 | 0.929994673 | cat2 | from_3570 | to_3570 | art_long | art_short | J |
| 54 | 200 | 0.772009701 | cat2 | from_3571 | to_3571 | art_long | art_short | J |
| 55 | 200 | 0.930288964 | cat2 | from_3572 | to_3572 | art_long | art_short | J |
| 58 | 200 | 0.930009466 | cat2 | from_3573 | to_3573 | art_long | art_short | J |
| 69 | 200 | 0.936011085 | cat2 | from_3574 | to_3574 | art_long | art_short | J |
| 72 | 200 | 0.91460473 | cat2 | from_3575 | to_3575 | art_long | art_short | J |
| 77 | 200 | 0.915329716 | cat2 | from_3576 | to_3576 | art_long | art_short | J |
| 80 | 200 | 0.916619675 | cat2 | from_3577 | to_3577 | art_long | art_short | J |
| 81 | 200 | 0.77267373 | cat2 | from_3578 | to_3578 | art_long | art_short | J |
| 83 | 200 | 0.920693606 | cat2 | from_3579 | to_3579 | art_long | art_short | J |
| 87 | 200 | 0.930585359 | cat2 | from_3580 | to_3580 | art_long | art_short | J |
| 95 | 200 | 0.931308162 | cat2 | from_3581 | to_3581 | art_long | art_short | J |
| 109 | 200 | 0.935933391 | cat2 | from_3582 | to_3582 | art_long | art_short | J |
| 111 | 200 | 0.933856736 | cat2 | from_3583 | to_3583 | art_long | art_short | J |
| 119 | 200 | 0.933363796 | cat2 | from_3584 | to_3584 | art_long | art_short | J |
| 120 | 200 | 0.914417816 | cat2 | from_3585 | to_3585 | art_long | art_short | J |
| 140 | 200 | 0.917439168 | cat2 | from_3586 | to_3586 | art_long | art_short | J |
| 173 | 200 | 0.634629121 | cat2 | from_3587 | to_3587 | art_short | art_short | H |
| 174 | 200 | 0.639305586 | cat2 | from_3588 | to_3588 | art_short | art_short | H |
| 195 | 200 | 0.999172664 | cat2 | from_3589 | to_3589 | art_short | art_short | H |
| 8 | 201 | 0.934233739 | cat2 | from_3590 | to_3590 | art_long | art_short | J |
| 9 | 201 | 0.930701048 | cat2 | from_3591 | to_3591 | art_long | art_short | J |
| 10 | 201 | 0.932780275 | cat2 | from_3592 | to_3592 | art_long | art_short | J |
| 23 | 201 | 0.830896794 | cat2 | from_3593 | to_3593 | art_long | art_short | J |
| 84 | 201 | 0.632747065 | cat2 | from_3594 | to_3594 | art_long | art_short | J |
| 88 | 201 | 0.741071659 | cat2 | from_3595 | to_3595 | art_long | art_short | J |
| 125 | 201 | 0.751822804 | cat2 | from_3596 | to_3596 | art_long | art_short | J |
| 126 | 201 | 0.803919295 | cat2 | from_3597 | to_3597 | art_long | art_short | J |
| 145 | 201 | 0.657584988 | cat2 | from_3598 | to_3598 | art_long | art_short | J |
| 156 | 201 | 0.700355286 | cat2 | from_3599 | to_3599 | art_short | art_short | H |
| 157 | 201 | 0.724166835 | cat2 | from_3600 | to_3600 | art_short | art_short | H |
| 158 | 201 | 0.699540775 | cat2 | from_3601 | to_3601 | art_short | art_short | H |
| 161 | 201 | 0.803256232 | cat2 | from_3602 | to_3602 | art_short | art_short | H |
| 193 | 201 | 0.672860168 | cat2 | from_3603 | to_3603 | art_short | art_short | H |
| 198 | 201 | 0.809270643 | cat2 | from_3604 | to_3604 | art_short | art_short | H |
| 51 | 202 | 0.846499142 | cat2 | from_3605 | to_3605 | art_long | art_short | J |
| 44 | 203 | 0.753505771 | cat2 | from_3606 | to_3606 | art_long | art_short | J |
| 46 | 203 | 0.761336764 | cat2 | from_3607 | to_3607 | art_long | art_short | J |
| 52 | 203 | 0.736455646 | cat2 | from_3608 | to_3608 | art_long | art_short | J |
| 60 | 203 | 0.80719785 | cat2 | from_3609 | to_3609 | art_long | art_short | J |
| 61 | 203 | 0.914123675 | cat2 | from_3610 | to_3610 | art_long | art_short | J |
| 99 | 203 | 0.906464919 | cat2 | from_3611 | to_3611 | art_long | art_short | J |
| 102 | 203 | 0.76356358 | cat2 | from_3612 | to_3612 | art_long | art_short | J |
| 105 | 203 | 0.919713406 | cat2 | from_3613 | to_3613 | art_long | art_short | J |
| 106 | 203 | 0.721496649 | cat2 | from_3614 | to_3614 | art_long | art_short | J |
| 128 | 203 | 0.770022756 | cat2 | from_3615 | to_3615 | art_long | art_short | J |
| 135 | 203 | 0.761285286 | cat2 | from_3616 | to_3616 | art_long | art_short | J |
| 148 | 203 | 0.887687726 | cat2 | from_3617 | to_3617 | art_long | art_short | J |
| 176 | 203 | 0.997801269 | cat2 | from_3618 | to_3618 | art_short | art_short | H |
| 183 | 203 | 0.999763351 | cat2 | from_3619 | to_3619 | art_short | art_short | H |
| 44 | 204 | 0.910699389 | cat2 | from_3620 | to_3620 | art_long | art_short | J |
| 46 | 204 | 0.919312107 | cat2 | from_3621 | to_3621 | art_long | art_short | J |
| 52 | 204 | 0.939361059 | cat2 | from_3622 | to_3622 | art_long | art_short | J |
| 60 | 204 | 0.950670699 | cat2 | from_3623 | to_3623 | art_long | art_short | J |
| 61 | 204 | 0.904225434 | cat2 | from_3624 | to_3624 | art_long | art_short | J |
| 99 | 204 | 0.901855266 | cat2 | from_3625 | to_3625 | art_long | art_short | J |
| 102 | 204 | 0.913479796 | cat2 | from_3626 | to_3626 | art_long | art_short | J |
| 105 | 204 | 0.773784981 | cat2 | from_3627 | to_3627 | art_long | art_short | J |
| 106 | 204 | 0.873383088 | cat2 | from_3628 | to_3628 | art_long | art_short | J |
| 128 | 204 | 0.925529662 | cat2 | from_3629 | to_3629 | art_long | art_short | J |
| 135 | 204 | 0.919263325 | cat2 | from_3630 | to_3630 | art_long | art_short | J |
| 148 | 204 | 0.88539799 | cat2 | from_3631 | to_3631 | art_long | art_short | J |
| 176 | 204 | 0.911480063 | cat2 | from_3632 | to_3632 | art_short | art_short | H |
| 183 | 204 | 0.912625125 | cat2 | from_3633 | to_3633 | art_short | art_short | H |
| 203 | 204 | 0.915012179 | cat2 | from_3634 | to_3634 | art_short | art_short | H |
| 4 | 205 | 0.731900335 | cat2 | from_3635 | to_3635 | art_long | art_short | J |
| 7 | 205 | 0.734964091 | cat2 | from_3636 | to_3636 | art_long | art_short | J |
| 11 | 205 | 0.962492438 | cat2 | from_3637 | to_3637 | art_long | art_short | J |
| 22 | 205 | 0.677924813 | cat2 | from_3638 | to_3638 | art_long | art_short | J |

| | | | | | | | | |
|---|---|---|---|---|---|---|---|---|
| 129 | 205 | -0.663677198 | cat2 | from_3639 | to_3639 | art_long | art_short | J |
| 136 | 205 | -0.635833876 | cat2 | from_3640 | to_3640 | art_long | art_short | J |
| 142 | 205 | -0.651455063 | cat2 | from_3641 | to_3641 | art_long | art_short | J |
| 159 | 205 | 0.75523592 | cat2 | from_3642 | to_3642 | art_short | art_short | H |
| 160 | 205 | 0.780525019 | cat2 | from_3643 | to_3643 | art_short | art_short | H |
| 161 | 205 | 0.687686156 | cat2 | from_3644 | to_3644 | art_short | art_short | H |
| 188 | 205 | 0.756710088 | cat2 | from_3645 | to_3645 | art_short | art_short | H |
| 30 | 206 | 0.891343181 | cat2 | from_3646 | to_3646 | art_long | art_short | J |
| 31 | 206 | 0.683412051 | cat2 | from_3647 | to_3647 | art_long | art_short | J |
| 33 | 206 | 0.679789393 | cat2 | from_3648 | to_3648 | art_long | art_short | J |
| 47 | 206 | 0.763521136 | cat2 | from_3649 | to_3649 | art_long | art_short | J |
| 54 | 206 | 0.917374945 | cat2 | from_3650 | to_3650 | art_long | art_short | J |
| 55 | 206 | 0.763984182 | cat2 | from_3651 | to_3651 | art_long | art_short | J |
| 58 | 206 | 0.767273749 | cat2 | from_3652 | to_3652 | art_long | art_short | J |
| 69 | 206 | 0.775186823 | cat2 | from_3653 | to_3653 | art_long | art_short | J |
| 72 | 206 | 0.912294848 | cat2 | from_3654 | to_3654 | art_long | art_short | J |
| 77 | 206 | 0.910798731 | cat2 | from_3655 | to_3655 | art_long | art_short | J |
| 80 | 206 | 0.914417713 | cat2 | from_3656 | to_3656 | art_long | art_short | J |
| 81 | 206 | 0.947689599 | cat2 | from_3657 | to_3657 | art_long | art_short | J |
| 83 | 206 | 0.9298009 | cat2 | from_3658 | to_3658 | art_long | art_short | J |
| 87 | 206 | 0.767279995 | cat2 | from_3659 | to_3659 | art_long | art_short | J |
| 95 | 206 | 0.766288016 | cat2 | from_3660 | to_3660 | art_long | art_short | J |
| 109 | 206 | 0.772572089 | cat2 | from_3661 | to_3661 | art_long | art_short | J |
| 111 | 206 | 0.770881591 | cat2 | from_3662 | to_3662 | art_long | art_short | J |
| 119 | 206 | 0.768701171 | cat2 | from_3663 | to_3663 | art_long | art_short | J |
| 120 | 206 | 0.916051851 | cat2 | from_3664 | to_3664 | art_long | art_short | J |
| 140 | 206 | 0.919489043 | cat2 | from_3665 | to_3665 | art_long | art_short | J |
| 195 | 206 | 0.914989736 | cat2 | from_3666 | to_3666 | art_short | art_short | H |
| 200 | 206 | 0.911849519 | cat2 | from_3667 | to_3667 | art_short | art_short | H |
| 4 | 207 | 0.644400612 | cat2 | from_3668 | to_3668 | art_long | art_short | J |
| 7 | 207 | 0.647607746 | cat2 | from_3669 | to_3669 | art_long | art_short | J |
| 22 | 207 | 0.680374228 | cat2 | from_3670 | to_3670 | art_long | art_short | J |
| 34 | 207 | 0.942553259 | cat2 | from_3671 | to_3671 | art_long | art_short | J |
| 64 | 207 | 0.641759926 | cat2 | from_3672 | to_3672 | art_long | art_short | J |
| 65 | 207 | 0.929821162 | cat2 | from_3673 | to_3673 | art_long | art_short | J |
| 113 | 207 | 0.996810133 | cat2 | from_3674 | to_3674 | art_long | art_short | J |
| 117 | 207 | 0.64001671 | cat2 | from_3675 | to_3675 | art_long | art_short | J |
| 125 | 207 | 0.652861993 | cat2 | from_3676 | to_3676 | art_long | art_short | J |
| 127 | 207 | 0.943882685 | cat2 | from_3677 | to_3677 | art_long | art_short | J |
| 159 | 207 | 0.660465099 | cat2 | from_3678 | to_3678 | art_short | art_short | H |
| 160 | 207 | 0.692175336 | cat2 | from_3679 | to_3679 | art_short | art_short | H |
| 161 | 207 | 0.695933253 | cat2 | from_3680 | to_3680 | art_short | art_short | H |
| 188 | 207 | 0.662723701 | cat2 | from_3681 | to_3681 | art_short | art_short | H |
| 198 | 207 | 0.709711275 | cat2 | from_3682 | to_3682 | art_short | art_short | H |
| 199 | 207 | 0.651301461 | cat2 | from_3683 | to_3683 | art_short | art_short | H |
| 3 | 208 | 0.756636748 | cat2 | from_3684 | to_3684 | art_long | art_short | J |
| 29 | 208 | 0.976268945 | cat2 | from_3685 | to_3685 | art_long | art_short | J |
| 41 | 208 | 0.798430411 | cat2 | from_3686 | to_3686 | art_long | art_short | J |
| 66 | 208 | 0.811683637 | cat2 | from_3687 | to_3687 | art_long | art_short | J |
| 67 | 208 | 0.811874661 | cat2 | from_3688 | to_3688 | art_long | art_short | J |
| 71 | 208 | 0.975514371 | cat2 | from_3689 | to_3689 | art_long | art_short | J |
| 82 | 208 | 0.815058028 | cat2 | from_3690 | to_3690 | art_long | art_short | J |
| 96 | 208 | 0.800988748 | cat2 | from_3691 | to_3691 | art_long | art_short | J |
| 98 | 208 | 0.656460638 | cat2 | from_3692 | to_3692 | art_long | art_short | J |
| 121 | 208 | 0.662107414 | cat2 | from_3693 | to_3693 | art_long | art_short | J |
| 124 | 208 | 0.809347667 | cat2 | from_3694 | to_3694 | art_long | art_short | J |
| 150 | 208 | 0.826789518 | cat2 | from_3695 | to_3695 | art_long | art_short | J |
| 175 | 208 | 0.801971554 | cat2 | from_3696 | to_3696 | art_short | art_short | H |
| 181 | 208 | 0.960505494 | cat2 | from_3697 | to_3697 | art_short | art_short | H |
| 185 | 208 | 0.63427706 | cat2 | from_3698 | to_3698 | art_short | art_short | H |
| 189 | 208 | 0.824931967 | cat2 | from_3699 | to_3699 | art_short | art_short | H |
| 3 | 209 | 0.682233365 | cat2 | from_3700 | to_3700 | art_long | art_short | J |
| 29 | 209 | 0.82521851 | cat2 | from_3701 | to_3701 | art_long | art_short | J |
| 41 | 209 | 0.69050679 | cat2 | from_3702 | to_3702 | art_long | art_short | J |
| 66 | 209 | 0.701804419 | cat2 | from_3703 | to_3703 | art_long | art_short | J |
| 67 | 209 | 0.700131079 | cat2 | from_3704 | to_3704 | art_long | art_short | J |
| 71 | 209 | 0.826978627 | cat2 | from_3705 | to_3705 | art_long | art_short | J |
| 82 | 209 | 0.968805759 | cat2 | from_3706 | to_3706 | art_long | art_short | J |
| 96 | 209 | 0.704754214 | cat2 | from_3707 | to_3707 | art_long | art_short | J |
| 98 | 209 | 0.803278222 | cat2 | from_3708 | to_3708 | art_long | art_short | J |
| 121 | 209 | 0.800111566 | cat2 | from_3709 | to_3709 | art_long | art_short | J |
| 124 | 209 | 0.695838624 | cat2 | from_3710 | to_3710 | art_long | art_short | J |
| 150 | 209 | 0.964888043 | cat2 | from_3711 | to_3711 | art_long | art_short | J |
| 175 | 209 | 0.964640273 | cat2 | from_3712 | to_3712 | art_short | art_short | H |
| 181 | 209 | 0.852420833 | cat2 | from_3713 | to_3713 | art_short | art_short | H |
| 185 | 209 | 0.814045231 | cat2 | from_3714 | to_3714 | art_short | art_short | H |
| 189 | 209 | 0.73943682 | cat2 | from_3715 | to_3715 | art_short | art_short | H |
| 208 | 209 | 0.857525089 | cat2 | from_3716 | to_3716 | art_short | art_short | H |
| 3 | 210 | 0.67707965 | cat2 | from_3717 | to_3717 | art_long | art_short | J |
| 29 | 210 | 0.823233328 | cat2 | from_3718 | to_3718 | art_long | art_short | J |
| 41 | 210 | 0.687229157 | cat2 | from_3719 | to_3719 | art_long | art_short | J |
| 66 | 210 | 0.699249256 | cat2 | from_3720 | to_3720 | art_long | art_short | J |
| 67 | 210 | 0.697052544 | cat2 | from_3721 | to_3721 | art_long | art_short | J |
| 71 | 210 | 0.825170375 | cat2 | from_3722 | to_3722 | art_long | art_short | J |
| 82 | 210 | 0.969477866 | cat2 | from_3723 | to_3723 | art_long | art_short | J |
| 96 | 210 | 0.700182576 | cat2 | from_3724 | to_3724 | art_long | art_short | J |
| 98 | 210 | 0.80296403 | cat2 | from_3725 | to_3725 | art_long | art_short | J |
| 121 | 210 | 0.797499942 | cat2 | from_3726 | to_3726 | art_long | art_short | J |
| 124 | 210 | 0.691860673 | cat2 | from_3727 | to_3727 | art_long | art_short | J |
| 150 | 210 | 0.968397936 | cat2 | from_3728 | to_3728 | art_long | art_short | J |
| 175 | 210 | 0.963472241 | cat2 | from_3729 | to_3729 | art_short | art_short | H |
| 181 | 210 | 0.852414289 | cat2 | from_3730 | to_3730 | art_short | art_short | H |
| 185 | 210 | 0.811520327 | cat2 | from_3731 | to_3731 | art_short | art_short | H |
| 189 | 210 | 0.735005193 | cat2 | from_3732 | to_3732 | art_short | art_short | H |
| 208 | 210 | 0.854701253 | cat2 | from_3733 | to_3733 | art_short | art_short | H |
| 209 | 210 | 0.999660704 | cat2 | from_3734 | to_3734 | art_short | art_short | H |
| 30 | 211 | 0.994878317 | cat2 | from_3735 | to_3735 | art_long | art_short | J |
| 47 | 211 | 0.930228316 | cat2 | from_3736 | to_3736 | art_long | art_short | J |
| 54 | 211 | 0.771515289 | cat2 | from_3737 | to_3737 | art_long | art_short | J |
| 55 | 211 | 0.930499835 | cat2 | from_3738 | to_3738 | art_long | art_short | J |
| 58 | 211 | 0.92985721 | cat2 | from_3739 | to_3739 | art_long | art_short | J |
| 69 | 211 | 0.935921452 | cat2 | from_3740 | to_3740 | art_long | art_short | J |
| 72 | 211 | 0.914553329 | cat2 | from_3741 | to_3741 | art_long | art_short | J |
| 77 | 211 | 0.91557035 | cat2 | from_3742 | to_3742 | art_long | art_short | J |

| | | | | | | | | |
|---|---|---|---|---|---|---|---|---|
| 80 | 211 | 0.916962682 | cat2 | from_3743 | to_3743 | art_long | art_short | J |
| 81 | 211 | 0.772239695 | cat2 | from_3744 | to_3744 | art_long | art_short | J |
| 83 | 211 | 0.9212056 | cat2 | from_3745 | to_3745 | art_long | art_short | J |
| 87 | 211 | 0.930481382 | cat2 | from_3746 | to_3746 | art_long | art_short | J |
| 95 | 211 | 0.931367932 | cat2 | from_3747 | to_3747 | art_long | art_short | J |
| 109 | 211 | 0.936104352 | cat2 | from_3748 | to_3748 | art_long | art_short | J |
| 111 | 211 | 0.933842111 | cat2 | from_3749 | to_3749 | art_long | art_short | J |
| 119 | 211 | 0.933531268 | cat2 | from_3750 | to_3750 | art_long | art_short | J |
| 120 | 211 | 0.914219018 | cat2 | from_3751 | to_3751 | art_long | art_short | J |
| 140 | 211 | 0.917345874 | cat2 | from_3752 | to_3752 | art_long | art_short | J |
| 173 | 211 | 0.634629773 | cat2 | from_3753 | to_3753 | art_short | art_short | H |
| 174 | 211 | 0.639337337 | cat2 | from_3754 | to_3754 | art_short | art_short | H |
| 195 | 211 | 0.999022806 | cat2 | from_3755 | to_3755 | art_short | art_short | H |
| 200 | 211 | 0.999993419 | cat2 | from_3756 | to_3756 | art_short | art_short | H |
| 206 | 211 | 0.911687185 | cat2 | from_3757 | to_3757 | art_short | art_short | H |
| 34 | 212 | 0.697498435 | cat2 | from_3758 | to_3758 | art_long | art_short | J |
| 49 | 212 | 0.855307466 | cat2 | from_3759 | to_3759 | art_long | art_short | J |
| 56 | 212 | 0.794477378 | cat2 | from_3760 | to_3760 | art_long | art_short | J |
| 57 | 212 | 0.795117808 | cat2 | from_3761 | to_3761 | art_long | art_short | J |
| 70 | 212 | 0.946353697 | cat2 | from_3762 | to_3762 | art_long | art_short | J |
| 75 | 212 | 0.85338221 | cat2 | from_3763 | to_3763 | art_long | art_short | J |
| 85 | 212 | 0.966618669 | cat2 | from_3764 | to_3764 | art_long | art_short | J |
| 113 | 212 | 0.682472519 | cat2 | from_3765 | to_3765 | art_long | art_short | J |
| 123 | 212 | 0.966234027 | cat2 | from_3766 | to_3766 | art_long | art_short | J |
| 127 | 212 | 0.676111316 | cat2 | from_3767 | to_3767 | art_long | art_short | J |
| 146 | 212 | 0.952494958 | cat2 | from_3768 | to_3768 | art_long | art_short | J |
| 182 | 212 | 0.798617487 | cat2 | from_3769 | to_3769 | art_short | art_short | H |
| 199 | 212 | 0.948807473 | cat2 | from_3770 | to_3770 | art_short | art_short | H |
| 207 | 212 | 0.675192909 | cat2 | from_3771 | to_3771 | art_short | art_short | H |
| 17 | 213 | 0.649072149 | cat2 | from_3772 | to_3772 | art_long | art_short | J |
| 27 | 213 | -0.696513575 | cat2 | from_3773 | to_3773 | art_long | art_short | J |
| 36 | 213 | 0.645304831 | cat2 | from_3774 | to_3774 | art_long | art_short | J |
| 50 | 213 | 0.761563326 | cat2 | from_3775 | to_3775 | art_long | art_short | J |
| 133 | 213 | 0.640550638 | cat2 | from_3776 | to_3776 | art_long | art_short | J |
| 134 | 213 | 0.648596597 | cat2 | from_3777 | to_3777 | art_long | art_short | J |
| 138 | 213 | 0.716268579 | cat2 | from_3778 | to_3778 | art_long | art_short | J |
| 143 | 213 | 0.656377532 | cat2 | from_3779 | to_3779 | art_long | art_short | J |
| 154 | 213 | 0.842900685 | cat2 | from_3780 | to_3780 | art_short | art_short | H |
| 172 | 213 | -0.658020764 | cat2 | from_3781 | to_3781 | art_short | art_short | H |
| 177 | 213 | 0.826365405 | cat2 | from_3782 | to_3782 | art_short | art_short | H |
| 35 | 214 | 0.859706296 | cat2 | from_3783 | to_3783 | art_long | art_short | J |
| 38 | 214 | 0.871111146 | cat2 | from_3784 | to_3784 | art_long | art_short | J |
| 48 | 214 | 0.859692399 | cat2 | from_3785 | to_3785 | art_long | art_short | J |
| 50 | 214 | 0.943532837 | cat2 | from_3786 | to_3786 | art_long | art_short | J |
| 64 | 214 | -0.668774892 | cat2 | from_3787 | to_3787 | art_long | art_short | J |
| 104 | 214 | 0.858565772 | cat2 | from_3788 | to_3788 | art_long | art_short | J |
| 117 | 214 | -0.667694491 | cat2 | from_3789 | to_3789 | art_long | art_short | J |
| 132 | 214 | 0.842167904 | cat2 | from_3790 | to_3790 | art_long | art_short | J |
| 143 | 214 | 0.953851086 | cat2 | from_3791 | to_3791 | art_long | art_short | J |
| 154 | 214 | 0.688314768 | cat2 | from_3792 | to_3792 | art_short | art_short | H |
| 166 | 214 | 0.944743354 | cat2 | from_3793 | to_3793 | art_short | art_short | H |
| 177 | 214 | 0.696680679 | cat2 | from_3794 | to_3794 | art_short | art_short | H |
| 213 | 214 | 0.743055982 | cat2 | from_3795 | to_3795 | art_short | art_short | H |
| 82 | 215 | 0.771979708 | cat2 | from_3796 | to_3796 | art_long | art_short | J |
| 150 | 215 | 0.72470583 | cat2 | from_3797 | to_3797 | art_long | art_short | J |
| 175 | 215 | 0.794390856 | cat2 | from_3798 | to_3798 | art_short | art_short | H |
| 209 | 215 | 0.787825024 | cat2 | from_3799 | to_3799 | art_short | art_short | H |
| 210 | 215 | 0.783437108 | cat2 | from_3800 | to_3800 | art_short | art_short | H |
| 2 | 216 | 0.675600047 | cat2 | from_3801 | to_3801 | art_long | art_short | J |
| 12 | 216 | 0.826779731 | cat2 | from_3802 | to_3802 | art_long | art_short | J |
| 15 | 216 | 0.693129804 | cat2 | from_3803 | to_3803 | art_long | art_short | J |
| 16 | 216 | 0.905645703 | cat2 | from_3804 | to_3804 | art_long | art_short | J |
| 18 | 216 | 0.905196723 | cat2 | from_3805 | to_3805 | art_long | art_short | J |
| 19 | 216 | 0.668077811 | cat2 | from_3806 | to_3806 | art_long | art_short | J |
| 39 | 216 | 0.814946125 | cat2 | from_3807 | to_3807 | art_long | art_short | J |
| 40 | 216 | 0.680375562 | cat2 | from_3808 | to_3808 | art_long | art_short | J |
| 53 | 216 | 0.683250436 | cat2 | from_3809 | to_3809 | art_long | art_short | J |
| 79 | 216 | 0.667485001 | cat2 | from_3810 | to_3810 | art_long | art_short | J |
| 155 | 216 | 0.72739058 | cat2 | from_3811 | to_3811 | art_short | art_short | H |
| 162 | 216 | 0.750005245 | cat2 | from_3812 | to_3812 | art_short | art_short | H |
| 163 | 216 | 0.660499658 | cat2 | from_3813 | to_3813 | art_short | art_short | H |
| 164 | 216 | 0.951939354 | cat2 | from_3814 | to_3814 | art_short | art_short | H |
| 167 | 216 | 0.829729109 | cat2 | from_3815 | to_3815 | art_short | art_short | H |
| 169 | 216 | 0.659925054 | cat2 | from_3816 | to_3816 | art_short | art_short | H |
| 179 | 216 | 0.734238247 | cat2 | from_3817 | to_3817 | art_short | art_short | H |
| 192 | 216 | 0.800041638 | cat2 | from_3818 | to_3818 | art_short | art_short | H |
| 196 | 216 | 0.798358381 | cat2 | from_3819 | to_3819 | art_short | art_short | H |
| 197 | 216 | 0.956529862 | cat2 | from_3820 | to_3820 | art_short | art_short | H |
| 1 | 217 | 0.786290273 | cat2 | from_3821 | to_3821 | art_long | art_short | J |
| 145 | 217 | 0.731437181 | cat2 | from_3822 | to_3822 | art_long | art_short | J |
| 30 | 218 | 0.994229043 | cat2 | from_3823 | to_3823 | art_long | art_short | J |
| 47 | 218 | 0.93081183 | cat2 | from_3824 | to_3824 | art_long | art_short | J |
| 54 | 218 | 0.770092038 | cat2 | from_3825 | to_3825 | art_long | art_short | J |
| 55 | 218 | 0.931020223 | cat2 | from_3826 | to_3826 | art_long | art_short | J |
| 58 | 218 | 0.929371296 | cat2 | from_3827 | to_3827 | art_long | art_short | J |
| 69 | 218 | 0.935608667 | cat2 | from_3828 | to_3828 | art_long | art_short | J |
| 72 | 218 | 0.914347971 | cat2 | from_3829 | to_3829 | art_long | art_short | J |
| 77 | 218 | 0.916174248 | cat2 | from_3830 | to_3830 | art_long | art_short | J |
| 80 | 218 | 0.917850193 | cat2 | from_3831 | to_3831 | art_long | art_short | J |
| 81 | 218 | 0.770983719 | cat2 | from_3832 | to_3832 | art_long | art_short | J |
| 83 | 218 | 0.922561137 | cat2 | from_3833 | to_3833 | art_long | art_short | J |
| 87 | 218 | 0.930129223 | cat2 | from_3834 | to_3834 | art_long | art_short | J |
| 95 | 218 | 0.931469507 | cat2 | from_3835 | to_3835 | art_long | art_short | J |
| 109 | 218 | 0.93651375 | cat2 | from_3836 | to_3836 | art_long | art_short | J |
| 111 | 218 | 0.933737344 | cat2 | from_3837 | to_3837 | art_long | art_short | J |
| 119 | 218 | 0.933931175 | cat2 | from_3838 | to_3838 | art_long | art_short | J |
| 120 | 218 | 0.913605196 | cat2 | from_3839 | to_3839 | art_long | art_short | J |
| 140 | 218 | 0.917024225 | cat2 | from_3840 | to_3840 | art_long | art_short | J |
| 173 | 218 | 0.634587925 | cat2 | from_3841 | to_3841 | art_short | art_short | H |
| 174 | 218 | 0.639381351 | cat2 | from_3842 | to_3842 | art_short | art_short | H |
| 195 | 218 | 0.998538781 | cat2 | from_3843 | to_3843 | art_short | art_short | H |
| 200 | 218 | 0.999906393 | cat2 | from_3844 | to_3844 | art_short | art_short | H |
| 206 | 218 | 0.911174594 | cat2 | from_3845 | to_3845 | art_short | art_short | H |
| 211 | 218 | 0.999949452 | cat2 | from_3846 | to_3846 | art_short | art_short | H |

| | | | | | | | | |
|---|---|---|---|---|---|---|---|---|
| 30 | 219 | 0.718232518 | cat2 | from_3847 | to_3847 | art_long | art_untreat | E |
| 81 | 219 | 0.692756166 | cat2 | from_3848 | to_3848 | art_long | art_untreat | E |
| 195 | 219 | 0.777426618 | cat2 | from_3849 | to_3849 | art_short | art_untreat | C |
| 200 | 219 | 0.767827769 | cat2 | from_3850 | to_3850 | art_short | art_untreat | C |
| 206 | 219 | 0.847365745 | cat2 | from_3851 | to_3851 | art_short | art_untreat | C |
| 211 | 219 | 0.767446584 | cat2 | from_3852 | to_3852 | art_short | art_untreat | C |
| 217 | 219 | 0.68814497 | cat2 | from_3853 | to_3853 | art_short | art_untreat | C |
| 218 | 219 | 0.766337403 | cat2 | from_3854 | to_3854 | art_short | art_untreat | C |
| 12 | 220 | 0.723109491 | cat2 | from_3855 | to_3855 | art_long | art_untreat | E |
| 13 | 220 | 0.798883332 | cat2 | from_3856 | to_3856 | art_long | art_untreat | E |
| 15 | 220 | 0.827613559 | cat2 | from_3857 | to_3857 | art_long | art_untreat | E |
| 16 | 220 | 0.633145078 | cat2 | from_3858 | to_3858 | art_long | art_untreat | E |
| 18 | 220 | 0.634357125 | cat2 | from_3859 | to_3859 | art_long | art_untreat | E |
| 39 | 220 | 0.636308754 | cat2 | from_3860 | to_3860 | art_long | art_untreat | E |
| 114 | 220 | 0.725424309 | cat2 | from_3861 | to_3861 | art_long | art_untreat | E |
| 162 | 220 | 0.945649424 | cat2 | from_3862 | to_3862 | art_short | art_untreat | C |
| 163 | 220 | 0.952275224 | cat2 | from_3863 | to_3863 | art_short | art_untreat | C |
| 164 | 220 | 0.746767005 | cat2 | from_3864 | to_3864 | art_short | art_untreat | C |
| 165 | 220 | 0.863450309 | cat2 | from_3865 | to_3865 | art_short | art_untreat | C |
| 167 | 220 | 0.720790254 | cat2 | from_3866 | to_3866 | art_short | art_untreat | C |
| 168 | 220 | 0.697285234 | cat2 | from_3867 | to_3867 | art_short | art_untreat | C |
| 169 | 220 | 0.951046665 | cat2 | from_3868 | to_3868 | art_short | art_untreat | C |
| 180 | 220 | 0.681746631 | cat2 | from_3869 | to_3869 | art_short | art_untreat | C |
| 186 | 220 | 0.657795483 | cat2 | from_3870 | to_3870 | art_short | art_untreat | C |
| 187 | 220 | 0.663793476 | cat2 | from_3871 | to_3871 | art_short | art_untreat | C |
| 192 | 220 | 0.784620461 | cat2 | from_3872 | to_3872 | art_short | art_untreat | C |
| 196 | 220 | 0.94299065 | cat2 | from_3873 | to_3873 | art_short | art_untreat | C |
| 197 | 220 | 0.766215621 | cat2 | from_3874 | to_3874 | art_short | art_untreat | C |
| 216 | 220 | 0.741375011 | cat2 | from_3875 | to_3875 | art_short | art_untreat | C |
| 89 | 221 | 0.861539741 | cat2 | from_3876 | to_3876 | art_long | art_untreat | E |
| 32 | 222 | -0.67527832 | cat2 | from_3877 | to_3877 | art_long | art_untreat | E |
| 49 | 222 | 0.639198987 | cat2 | from_3878 | to_3878 | art_long | art_untreat | E |
| 56 | 222 | 0.824748779 | cat2 | from_3879 | to_3879 | art_long | art_untreat | E |
| 57 | 222 | 0.825845616 | cat2 | from_3880 | to_3880 | art_long | art_untreat | E |
| 70 | 222 | 0.653176797 | cat2 | from_3881 | to_3881 | art_long | art_untreat | E |
| 75 | 222 | 0.637110973 | cat2 | from_3882 | to_3882 | art_long | art_untreat | E |
| 146 | 222 | 0.652841906 | cat2 | from_3883 | to_3883 | art_long | art_untreat | E |
| 161 | 222 | 0.697942062 | cat2 | from_3884 | to_3884 | art_short | art_untreat | C |
| 182 | 222 | 0.938893939 | cat2 | from_3885 | to_3885 | art_short | art_untreat | C |
| 193 | 222 | 0.642776764 | cat2 | from_3886 | to_3886 | art_short | art_untreat | C |
| 199 | 222 | 0.767182127 | cat2 | from_3887 | to_3887 | art_short | art_untreat | C |
| 212 | 222 | 0.729314508 | cat2 | from_3888 | to_3888 | art_short | art_untreat | C |
| 44 | 223 | 0.686477376 | cat2 | from_3889 | to_3889 | art_long | art_untreat | E |
| 46 | 223 | 0.692247866 | cat2 | from_3890 | to_3890 | art_long | art_untreat | E |
| 52 | 223 | 0.857183267 | cat2 | from_3891 | to_3891 | art_long | art_untreat | E |
| 60 | 223 | 0.753280319 | cat2 | from_3892 | to_3892 | art_long | art_untreat | E |
| 61 | 223 | 0.75158083 | cat2 | from_3893 | to_3893 | art_long | art_untreat | E |
| 99 | 223 | 0.74626769 | cat2 | from_3894 | to_3894 | art_long | art_untreat | E |
| 102 | 223 | 0.704816044 | cat2 | from_3895 | to_3895 | art_long | art_untreat | E |
| 105 | 223 | 0.699301396 | cat2 | from_3896 | to_3896 | art_long | art_untreat | E |
| 106 | 223 | 0.6595096 | cat2 | from_3897 | to_3897 | art_long | art_untreat | E |
| 128 | 223 | 0.703153482 | cat2 | from_3898 | to_3898 | art_long | art_untreat | E |
| 135 | 223 | 0.692222475 | cat2 | from_3899 | to_3899 | art_long | art_untreat | E |
| 148 | 223 | 0.732787487 | cat2 | from_3900 | to_3900 | art_long | art_untreat | E |
| 176 | 223 | 0.918541504 | cat2 | from_3901 | to_3901 | art_short | art_untreat | C |
| 183 | 223 | 0.923136105 | cat2 | from_3902 | to_3902 | art_short | art_untreat | C |
| 203 | 223 | 0.921331642 | cat2 | from_3903 | to_3903 | art_short | art_untreat | C |
| 204 | 223 | 0.920038237 | cat2 | from_3904 | to_3904 | art_short | art_untreat | C |
| 29 | 224 | 0.807399037 | cat2 | from_3905 | to_3905 | art_long | art_untreat | E |
| 71 | 224 | 0.759285661 | cat2 | from_3906 | to_3906 | art_long | art_untreat | E |
| 82 | 224 | 0.657590919 | cat2 | from_3907 | to_3907 | art_long | art_untreat | E |
| 124 | 224 | 0.896714524 | cat2 | from_3908 | to_3908 | art_long | art_untreat | E |
| 150 | 224 | 0.667449299 | cat2 | from_3909 | to_3909 | art_long | art_untreat | E |
| 175 | 224 | 0.696321806 | cat2 | from_3910 | to_3910 | art_short | art_untreat | C |
| 181 | 224 | 0.788037574 | cat2 | from_3911 | to_3911 | art_short | art_untreat | C |
| 208 | 224 | 0.819906503 | cat2 | from_3912 | to_3912 | art_short | art_untreat | C |
| 209 | 224 | 0.736674198 | cat2 | from_3913 | to_3913 | art_short | art_untreat | C |
| 210 | 224 | 0.733800999 | cat2 | from_3914 | to_3914 | art_short | art_untreat | C |
| 30 | 225 | 0.878228171 | cat2 | from_3915 | to_3915 | art_long | art_untreat | E |
| 47 | 225 | 0.694554349 | cat2 | from_3916 | to_3916 | art_long | art_untreat | E |
| 54 | 225 | 0.709189646 | cat2 | from_3917 | to_3917 | art_long | art_untreat | E |
| 55 | 225 | 0.695285693 | cat2 | from_3918 | to_3918 | art_long | art_untreat | E |
| 58 | 225 | 0.703206593 | cat2 | from_3919 | to_3919 | art_long | art_untreat | E |
| 69 | 225 | 0.71270479 | cat2 | from_3920 | to_3920 | art_long | art_untreat | E |
| 72 | 225 | 0.756529861 | cat2 | from_3921 | to_3921 | art_long | art_untreat | E |
| 77 | 225 | 0.753583543 | cat2 | from_3922 | to_3922 | art_long | art_untreat | E |
| 80 | 225 | 0.759291057 | cat2 | from_3923 | to_3923 | art_long | art_untreat | E |
| 81 | 225 | 0.770194804 | cat2 | from_3924 | to_3924 | art_long | art_untreat | E |
| 83 | 225 | 0.784450977 | cat2 | from_3925 | to_3925 | art_long | art_untreat | E |
| 87 | 225 | 0.70255885 | cat2 | from_3926 | to_3926 | art_long | art_untreat | E |
| 95 | 225 | 0.699436096 | cat2 | from_3927 | to_3927 | art_long | art_untreat | E |
| 109 | 225 | 0.706766666 | cat2 | from_3928 | to_3928 | art_long | art_untreat | E |
| 111 | 225 | 0.706184836 | cat2 | from_3929 | to_3929 | art_long | art_untreat | E |
| 119 | 225 | 0.70171943 | cat2 | from_3930 | to_3930 | art_long | art_untreat | E |
| 120 | 225 | 0.763505702 | cat2 | from_3931 | to_3931 | art_long | art_untreat | E |
| 140 | 225 | 0.767041816 | cat2 | from_3932 | to_3932 | art_long | art_untreat | E |
| 195 | 225 | 0.917188193 | cat2 | from_3933 | to_3933 | art_short | art_untreat | C |
| 200 | 225 | 0.910261774 | cat2 | from_3934 | to_3934 | art_short | art_untreat | C |
| 206 | 225 | 0.92510687 | cat2 | from_3935 | to_3935 | art_short | art_untreat | C |
| 211 | 225 | 0.909909745 | cat2 | from_3936 | to_3936 | art_short | art_untreat | C |
| 218 | 225 | 0.908871568 | cat2 | from_3937 | to_3937 | art_short | art_untreat | C |
| 219 | 225 | 0.96352926 | cat2 | from_3938 | to_3938 | art_untreat | art_untreat | A |
| 61 | 226 | 0.706614913 | cat2 | from_3939 | to_3939 | art_long | art_untreat | E |
| 99 | 226 | 0.689058645 | cat2 | from_3940 | to_3940 | art_long | art_untreat | E |
| 105 | 226 | 0.764719695 | cat2 | from_3941 | to_3941 | art_long | art_untreat | E |
| 148 | 226 | 0.669686223 | cat2 | from_3942 | to_3942 | art_long | art_untreat | E |
| 176 | 226 | 0.930967841 | cat2 | from_3943 | to_3943 | art_short | art_untreat | C |
| 183 | 226 | 0.934008971 | cat2 | from_3944 | to_3944 | art_short | art_untreat | C |
| 203 | 226 | 0.929574807 | cat2 | from_3945 | to_3945 | art_short | art_untreat | C |
| 204 | 226 | 0.786384762 | cat2 | from_3946 | to_3946 | art_short | art_untreat | C |
| 217 | 226 | 0.638326797 | cat2 | from_3947 | to_3947 | art_short | art_untreat | C |
| 223 | 226 | 0.937730028 | cat2 | from_3948 | to_3948 | art_untreat | art_untreat | A |
| 1 | 227 | 0.659426682 | cat2 | from_3949 | to_3949 | art_long | ec | N |
| 113 | 227 | 0.670240471 | cat2 | from_3950 | to_3950 | art_long | ec | N |

| | | | | | | | | |
|---|---|---|---|---|---|---|---|---|
| 198 | 227 | 0.738265139 | cat2 | from_3951 | to_3951 | art_short | ec | K |
| 207 | 227 | 0.711227815 | cat2 | from_3952 | to_3952 | art_short | ec | K |
| 34 | 228 | 0.778544026 | cat2 | from_3953 | to_3953 | art_long | ec | N |
| 113 | 228 | 0.817927279 | cat2 | from_3954 | to_3954 | art_long | ec | N |
| 127 | 228 | 0.649194977 | cat2 | from_3955 | to_3955 | art_long | ec | N |
| 161 | 228 | 0.661384672 | cat2 | from_3956 | to_3956 | art_short | ec | K |
| 198 | 228 | 0.712923451 | cat2 | from_3957 | to_3957 | art_short | ec | K |
| 207 | 228 | 0.848701576 | cat2 | from_3958 | to_3958 | art_short | ec | K |
| 227 | 228 | 0.968600877 | cat2 | from_3959 | to_3959 | ec | ec | P |
| 12 | 229 | 0.806640462 | cat2 | from_3960 | to_3960 | art_long | ec | N |
| 19 | 229 | 0.642371056 | cat2 | from_3961 | to_3961 | art_long | ec | N |
| 79 | 229 | 0.743543203 | cat2 | from_3962 | to_3962 | art_long | ec | N |
| 145 | 229 | 0.646358112 | cat2 | from_3963 | to_3963 | art_long | ec | N |
| 156 | 229 | 0.64841036 | cat2 | from_3964 | to_3964 | art_short | ec | K |
| 158 | 229 | 0.644468342 | cat2 | from_3965 | to_3965 | art_short | ec | K |
| 164 | 229 | 0.649414801 | cat2 | from_3966 | to_3966 | art_short | ec | K |
| 165 | 229 | 0.716246649 | cat2 | from_3967 | to_3967 | art_short | ec | K |
| 167 | 229 | 0.79869077 | cat2 | from_3968 | to_3968 | art_short | ec | K |
| 197 | 229 | 0.744424995 | cat2 | from_3969 | to_3969 | art_short | ec | K |
| 217 | 229 | 0.820571578 | cat2 | from_3970 | to_3970 | art_short | ec | K |
| 220 | 229 | 0.657511489 | cat2 | from_3971 | to_3971 | art_untreat | ec | F |
| 2 | 230 | 0.639848204 | cat2 | from_3972 | to_3972 | art_long | ec | N |
| 53 | 230 | 0.67180924 | cat2 | from_3973 | to_3973 | art_long | ec | N |
| 155 | 230 | 0.742317407 | cat2 | from_3974 | to_3974 | art_short | ec | K |
| 162 | 230 | 0.662232204 | cat2 | from_3975 | to_3975 | art_short | ec | K |
| 168 | 230 | 0.864569401 | cat2 | from_3976 | to_3976 | art_short | ec | K |
| 179 | 230 | 0.763210477 | cat2 | from_3977 | to_3977 | art_short | ec | K |
| 180 | 230 | 0.902769276 | cat2 | from_3978 | to_3978 | art_short | ec | K |
| 187 | 230 | 0.885333165 | cat2 | from_3979 | to_3979 | art_short | ec | K |
| 196 | 230 | 0.751736174 | cat2 | from_3980 | to_3980 | art_short | ec | K |
| 220 | 230 | 0.740123591 | cat2 | from_3981 | to_3981 | art_untreat | ec | F |
| 34 | 231 | 0.76854363 | cat2 | from_3982 | to_3982 | art_long | ec | N |
| 113 | 231 | 0.809821548 | cat2 | from_3983 | to_3983 | art_long | ec | N |
| 127 | 231 | 0.637787945 | cat2 | from_3984 | to_3984 | art_long | ec | N |
| 161 | 231 | 0.659067396 | cat2 | from_3985 | to_3985 | art_short | ec | K |
| 198 | 231 | 0.715610722 | cat2 | from_3986 | to_3986 | art_short | ec | K |
| 207 | 231 | 0.841527072 | cat2 | from_3987 | to_3987 | art_short | ec | K |
| 227 | 231 | 0.972291779 | cat2 | from_3988 | to_3988 | ec | ec | P |
| 228 | 231 | 0.999873528 | cat2 | from_3989 | to_3989 | ec | ec | P |
| 139 | 232 | 0.671071921 | cat2 | from_3990 | to_3990 | art_long | ec | N |
| 170 | 232 | 0.724060866 | cat2 | from_3991 | to_3991 | art_short | ec | K |
| 171 | 232 | 0.717016135 | cat2 | from_3992 | to_3992 | art_short | ec | K |
| 178 | 232 | 0.751557906 | cat2 | from_3993 | to_3993 | art_short | ec | K |
| 184 | 232 | 0.697102741 | cat2 | from_3994 | to_3994 | art_short | ec | K |
| 223 | 232 | 0.718399224 | cat2 | from_3995 | to_3995 | art_untreat | ec | F |
| 226 | 232 | 0.729684907 | cat2 | from_3996 | to_3996 | art_untreat | ec | F |
| 26 | 233 | 0.807601265 | cat2 | from_3997 | to_3997 | art_long | ec | N |
| 37 | 233 | 0.665741917 | cat2 | from_3998 | to_3998 | art_long | ec | N |
| 115 | 233 | 0.807664364 | cat2 | from_3999 | to_3999 | art_long | ec | N |
| 130 | 233 | 0.681157206 | cat2 | from_4000 | to_4000 | art_long | ec | N |
| 136 | 233 | 0.753971184 | cat2 | from_4001 | to_4001 | art_long | ec | N |
| 29 | 234 | 0.637345342 | cat2 | from_4002 | to_4002 | art_long | ec | N |
| 41 | 234 | 0.722860201 | cat2 | from_4003 | to_4003 | art_long | ec | N |
| 82 | 234 | 0.659894238 | cat2 | from_4004 | to_4004 | art_long | ec | N |
| 96 | 234 | 0.722648947 | cat2 | from_4005 | to_4005 | art_long | ec | N |
| 98 | 234 | 0.834168487 | cat2 | from_4006 | to_4006 | art_long | ec | N |
| 121 | 234 | 0.844146715 | cat2 | from_4007 | to_4007 | art_long | ec | N |
| 150 | 234 | 0.670775668 | cat2 | from_4008 | to_4008 | art_long | ec | N |
| 175 | 234 | 0.736526292 | cat2 | from_4009 | to_4009 | art_short | ec | K |
| 209 | 234 | 0.775524699 | cat2 | from_4010 | to_4010 | art_short | ec | K |
| 210 | 234 | 0.771839256 | cat2 | from_4011 | to_4011 | art_short | ec | K |
| 29 | 235 | 0.784105045 | cat2 | from_4012 | to_4012 | art_long | ec | N |
| 41 | 235 | 0.833019844 | cat2 | from_4013 | to_4013 | art_long | ec | N |
| 71 | 235 | 0.68781777 | cat2 | from_4014 | to_4014 | art_long | ec | N |
| 96 | 235 | 0.821202177 | cat2 | from_4015 | to_4015 | art_long | ec | N |
| 98 | 235 | 0.73640094 | cat2 | from_4016 | to_4016 | art_long | ec | N |
| 121 | 235 | 0.75094149 | cat2 | from_4017 | to_4017 | art_long | ec | N |
| 181 | 235 | 0.704148406 | cat2 | from_4018 | to_4018 | art_short | ec | K |
| 208 | 235 | 0.769123651 | cat2 | from_4019 | to_4019 | art_short | ec | K |
| 209 | 235 | 0.680766978 | cat2 | from_4020 | to_4020 | art_short | ec | K |
| 210 | 235 | 0.675807774 | cat2 | from_4021 | to_4021 | art_short | ec | K |
| 224 | 235 | 0.647123898 | cat2 | from_4022 | to_4022 | art_untreat | ec | F |
| 234 | 235 | 0.903787488 | cat2 | from_4023 | to_4023 | ec | ec | P |
| 1 | 236 | 0.685134786 | cat2 | from_4024 | to_4024 | art_long | ec | N |
| 202 | 236 | 0.641354415 | cat2 | from_4025 | to_4025 | art_short | ec | K |
| 227 | 236 | 0.770108554 | cat2 | from_4026 | to_4026 | ec | ec | P |
| 228 | 236 | 0.689492905 | cat2 | from_4027 | to_4027 | ec | ec | P |
| 231 | 236 | 0.696159182 | cat2 | from_4028 | to_4028 | ec | ec | P |
| 232 | 236 | 0.719192308 | cat2 | from_4029 | to_4029 | ec | ec | P |
| 1 | 237 | 0.678959118 | cat2 | from_4030 | to_4030 | art_long | ec | N |
| 170 | 237 | 0.662060864 | cat2 | from_4031 | to_4031 | art_short | ec | K |
| 171 | 237 | 0.654715353 | cat2 | from_4032 | to_4032 | art_short | ec | K |
| 178 | 237 | 0.761263342 | cat2 | from_4033 | to_4033 | art_short | ec | K |
| 184 | 237 | 0.634310888 | cat2 | from_4034 | to_4034 | art_short | ec | K |
| 223 | 237 | 0.693487979 | cat2 | from_4035 | to_4035 | art_untreat | ec | F |
| 226 | 237 | 0.660676424 | cat2 | from_4036 | to_4036 | art_untreat | ec | F |
| 232 | 237 | 0.976426076 | cat2 | from_4037 | to_4037 | ec | ec | P |
| 236 | 237 | 0.789030661 | cat2 | from_4038 | to_4038 | ec | ec | P |
| 2 | 238 | 0.768931301 | cat2 | from_4039 | to_4039 | art_long | ec | N |
| 53 | 238 | 0.802674915 | cat2 | from_4040 | to_4040 | art_long | ec | N |
| 155 | 238 | 0.89120973 | cat2 | from_4041 | to_4041 | art_short | ec | K |
| 164 | 238 | 0.681807505 | cat2 | from_4042 | to_4042 | art_short | ec | K |
| 168 | 238 | 0.735053514 | cat2 | from_4043 | to_4043 | art_short | ec | K |
| 179 | 238 | 0.916113631 | cat2 | from_4044 | to_4044 | art_short | ec | K |
| 180 | 238 | 0.755234767 | cat2 | from_4045 | to_4045 | art_short | ec | K |
| 187 | 238 | 0.731570474 | cat2 | from_4046 | to_4046 | art_short | ec | K |
| 197 | 238 | 0.71532158 | cat2 | from_4047 | to_4047 | art_short | ec | K |
| 216 | 238 | 0.734461317 | cat2 | from_4048 | to_4048 | art_short | ec | K |
| 229 | 238 | 0.642919512 | cat2 | from_4049 | to_4049 | ec | ec | P |
| 230 | 238 | 0.870327263 | cat2 | from_4050 | to_4050 | ec | ec | P |
| 32 | 239 | -0.656670918 | cat2 | from_4051 | to_4051 | art_long | ec | N |
| 161 | 239 | 0.652465369 | cat2 | from_4052 | to_4052 | art_short | ec | K |
| 182 | 239 | 0.787625011 | cat2 | from_4053 | to_4053 | art_short | ec | K |
| 193 | 239 | 0.661183876 | cat2 | from_4054 | to_4054 | art_short | ec | K |

| | | | | | | | | |
|---|---|---|---|---|---|---|---|---|
| 199 | 239 | 0.673700592 | cat2 | from_4055 | to_4055 | art_short | ec | K |
| 222 | 239 | 0.950048125 | cat2 | from_4056 | to_4056 | art_untreat | ec | F |
| 227 | 239 | 0.639764902 | cat2 | from_4057 | to_4057 | ec | ec | P |
| 17 | 240 | 0.70267162 | cat2 | from_4058 | to_4058 | art_long | ec | N |
| 36 | 240 | 0.698473007 | cat2 | from_4059 | to_4059 | art_long | ec | N |
| 133 | 240 | 0.695089946 | cat2 | from_4060 | to_4060 | art_long | ec | N |
| 134 | 240 | 0.700652195 | cat2 | from_4061 | to_4061 | art_long | ec | N |
| 138 | 240 | 0.786232906 | cat2 | from_4062 | to_4062 | art_long | ec | N |
| 143 | 240 | 0.710843657 | cat2 | from_4063 | to_4063 | art_long | ec | N |
| 154 | 240 | 0.956574717 | cat2 | from_4064 | to_4064 | art_short | ec | K |
| 166 | 240 | 0.668646839 | cat2 | from_4065 | to_4065 | art_short | ec | K |
| 177 | 240 | 0.938649548 | cat2 | from_4066 | to_4066 | art_short | ec | K |
| 178 | 240 | 0.665177321 | cat2 | from_4067 | to_4067 | art_short | ec | K |
| 213 | 240 | 0.698317511 | cat2 | from_4068 | to_4068 | art_short | ec | K |
| 232 | 240 | 0.658608234 | cat2 | from_4069 | to_4069 | ec | ec | P |
| 237 | 240 | 0.709765914 | cat2 | from_4070 | to_4070 | ec | ec | P |
| 17 | 241 | 0.708171591 | cat2 | from_4071 | to_4071 | art_long | ec | N |
| 36 | 241 | 0.704056958 | cat2 | from_4072 | to_4072 | art_long | ec | N |
| 133 | 241 | 0.700931287 | cat2 | from_4073 | to_4073 | art_long | ec | N |
| 134 | 241 | 0.706030293 | cat2 | from_4074 | to_4074 | art_long | ec | N |
| 138 | 241 | 0.790772895 | cat2 | from_4075 | to_4075 | art_long | ec | N |
| 143 | 241 | 0.712587994 | cat2 | from_4076 | to_4076 | art_long | ec | N |
| 154 | 241 | 0.957258231 | cat2 | from_4077 | to_4077 | art_short | ec | K |
| 166 | 241 | 0.671689782 | cat2 | from_4078 | to_4078 | art_short | ec | K |
| 177 | 241 | 0.940792384 | cat2 | from_4079 | to_4079 | art_short | ec | K |
| 178 | 241 | 0.66784305 | cat2 | from_4080 | to_4080 | art_short | ec | K |
| 213 | 241 | 0.697555144 | cat2 | from_4081 | to_4081 | art_short | ec | K |
| 232 | 241 | 0.654287409 | cat2 | from_4082 | to_4082 | ec | ec | P |
| 237 | 241 | 0.705439043 | cat2 | from_4083 | to_4083 | ec | ec | P |
| 240 | 241 | 0.999936552 | cat2 | from_4084 | to_4084 | ec | ec | P |
| 156 | 242 | 0.676766094 | cat2 | from_4085 | to_4085 | art_short | ec | K |
| 157 | 242 | 0.727078384 | cat2 | from_4086 | to_4086 | art_short | ec | K |
| 158 | 242 | 0.67196637 | cat2 | from_4087 | to_4087 | art_short | ec | K |
| 193 | 242 | 0.644316473 | cat2 | from_4088 | to_4088 | art_short | ec | K |
| 198 | 242 | 0.860343898 | cat2 | from_4089 | to_4089 | art_short | ec | K |
| 201 | 242 | 0.704189619 | cat2 | from_4090 | to_4090 | art_short | ec | K |
| 217 | 242 | 0.739117897 | cat2 | from_4091 | to_4091 | art_short | ec | K |
| 227 | 242 | 0.815812918 | cat2 | from_4092 | to_4092 | ec | ec | P |
| 228 | 242 | 0.681279983 | cat2 | from_4093 | to_4093 | ec | ec | P |
| 229 | 242 | 0.637256661 | cat2 | from_4094 | to_4094 | ec | ec | P |
| 231 | 242 | 0.690806284 | cat2 | from_4095 | to_4095 | ec | ec | P |
| 236 | 242 | 0.638844418 | cat2 | from_4096 | to_4096 | ec | ec | P |
| 34 | 243 | 0.783515722 | cat2 | from_4097 | to_4097 | art_long | ec | N |
| 113 | 243 | 0.821880542 | cat2 | from_4098 | to_4098 | art_long | ec | N |
| 127 | 243 | 0.654823526 | cat2 | from_4099 | to_4099 | art_long | ec | N |
| 161 | 243 | 0.662586564 | cat2 | from_4100 | to_4100 | art_short | ec | K |
| 198 | 243 | 0.711408303 | cat2 | from_4101 | to_4101 | art_short | ec | K |
| 207 | 243 | 0.852252569 | cat2 | from_4102 | to_4102 | art_short | ec | K |
| 227 | 243 | 0.966597838 | cat2 | from_4103 | to_4103 | ec | ec | P |
| 228 | 243 | 0.999968243 | cat2 | from_4104 | to_4104 | ec | ec | P |
| 231 | 243 | 0.999716889 | cat2 | from_4105 | to_4105 | ec | ec | P |
| 236 | 243 | 0.686127243 | cat2 | from_4106 | to_4106 | ec | ec | P |
| 242 | 243 | 0.676284458 | cat2 | from_4107 | to_4107 | ec | ec | P |
| 193 | 244 | 0.632245566 | cat2 | from_4108 | to_4108 | art_short | ec | K |
| 236 | 244 | 0.66875462 | cat2 | from_4109 | to_4109 | ec | ec | P |
| 242 | 244 | 0.71658283 | cat2 | from_4110 | to_4110 | ec | ec | P |
| 31 | 245 | 0.657193367 | cat2 | from_4111 | to_4111 | art_long | ec | N |
| 81 | 245 | 0.689710709 | cat2 | from_4112 | to_4112 | art_long | ec | N |
| 145 | 245 | 0.697612276 | cat2 | from_4113 | to_4113 | art_long | ec | N |
| 206 | 245 | 0.725769721 | cat2 | from_4114 | to_4114 | art_short | ec | K |
| 217 | 245 | 0.81462815 | cat2 | from_4115 | to_4115 | art_short | ec | K |
| 219 | 245 | 0.886302444 | cat2 | from_4116 | to_4116 | art_untreat | ec | F |
| 225 | 245 | 0.767563491 | cat2 | from_4117 | to_4117 | art_untreat | ec | F |
| 229 | 245 | 0.692181486 | cat2 | from_4118 | to_4118 | ec | ec | P |
| 242 | 245 | 0.734095709 | cat2 | from_4119 | to_4119 | ec | ec | P |
| 49 | 246 | -0.656516091 | cat2 | from_4120 | to_4120 | art_long | ec | N |
| 52 | 246 | 0.656054488 | cat2 | from_4121 | to_4121 | art_long | ec | N |
| 75 | 246 | -0.652095576 | cat2 | from_4122 | to_4122 | art_long | ec | N |
| 86 | 246 | 0.676712804 | cat2 | from_4123 | to_4123 | art_long | ec | N |
| 94 | 246 | 0.674400832 | cat2 | from_4124 | to_4124 | art_long | ec | N |
| 190 | 246 | 0.645758018 | cat2 | from_4125 | to_4125 | art_short | ec | K |
| 191 | 246 | 0.648331907 | cat2 | from_4126 | to_4126 | art_short | ec | K |
| 219 | 246 | 0.697657769 | cat2 | from_4127 | to_4127 | art_untreat | ec | F |
| 223 | 246 | 0.675997478 | cat2 | from_4128 | to_4128 | art_untreat | ec | F |
| 225 | 246 | 0.694402226 | cat2 | from_4129 | to_4129 | art_untreat | ec | F |
| 12 | 247 | 0.950067419 | cat2 | from_4130 | to_4130 | art_long | ec | N |
| 13 | 247 | 0.707572346 | cat2 | from_4131 | to_4131 | art_long | ec | N |
| 16 | 247 | 0.635649782 | cat2 | from_4132 | to_4132 | art_long | ec | N |
| 18 | 247 | 0.637026382 | cat2 | from_4133 | to_4133 | art_long | ec | N |
| 19 | 247 | 0.916629319 | cat2 | from_4134 | to_4134 | art_long | ec | N |
| 39 | 247 | 0.822512659 | cat2 | from_4135 | to_4135 | art_long | ec | N |
| 40 | 247 | 0.880355338 | cat2 | from_4136 | to_4136 | art_long | ec | N |
| 79 | 247 | 0.96494988 | cat2 | from_4137 | to_4137 | art_long | ec | N |
| 114 | 247 | 0.811410696 | cat2 | from_4138 | to_4138 | art_long | ec | N |
| 145 | 247 | 0.685537563 | cat2 | from_4139 | to_4139 | art_long | ec | N |
| 156 | 247 | 0.740975209 | cat2 | from_4140 | to_4140 | art_short | ec | K |
| 158 | 247 | 0.740025345 | cat2 | from_4141 | to_4141 | art_short | ec | K |
| 163 | 247 | 0.720233737 | cat2 | from_4142 | to_4142 | art_short | ec | K |
| 164 | 247 | 0.789703203 | cat2 | from_4143 | to_4143 | art_short | ec | K |
| 165 | 247 | 0.825748849 | cat2 | from_4144 | to_4144 | art_short | ec | K |
| 167 | 247 | 0.946719918 | cat2 | from_4145 | to_4145 | art_short | ec | K |
| 169 | 247 | 0.717703099 | cat2 | from_4146 | to_4146 | art_short | ec | K |
| 186 | 247 | 0.635405644 | cat2 | from_4147 | to_4147 | art_short | ec | K |
| 197 | 247 | 0.82706647 | cat2 | from_4148 | to_4148 | art_short | ec | K |
| 220 | 247 | 0.642535643 | cat2 | from_4149 | to_4149 | art_untreat | ec | F |
| 229 | 247 | 0.886151172 | cat2 | from_4150 | to_4150 | ec | ec | P |
| 50 | 248 | 0.663992137 | cat2 | from_4151 | to_4151 | art_long | ec | N |
| 61 | 248 | 0.856048296 | cat2 | from_4152 | to_4152 | art_long | ec | N |
| 99 | 248 | 0.826686481 | cat2 | from_4153 | to_4153 | art_long | ec | N |
| 105 | 248 | 0.951088645 | cat2 | from_4154 | to_4154 | art_long | ec | N |
| 148 | 248 | 0.793979246 | cat2 | from_4155 | to_4155 | art_long | ec | N |
| 151 | 248 | 0.766539377 | cat2 | from_4156 | to_4156 | art_long | ec | N |
| 152 | 248 | 0.684657952 | cat2 | from_4157 | to_4157 | art_long | ec | N |
| 157 | 248 | 0.641188854 | cat2 | from_4158 | to_4158 | art_short | ec | K |

| | | | | | | | | |
|---|---|---|---|---|---|---|---|---|
| 176 | 248 | 0.816088572 | cat2 | from_4159 | to_4159 | art_short | ec | K |
| 183 | 248 | 0.809055938 | cat2 | from_4160 | to_4160 | art_short | ec | K |
| 203 | 248 | 0.809520353 | cat2 | from_4161 | to_4161 | art_short | ec | K |
| 226 | 248 | 0.660365383 | cat2 | from_4162 | to_4162 | art_untreat | ec | F |
| 2 | 6 | -0.626555667 | cat3 | from_4163 | to_4163 | art_long | art_long | M |
| 4 | 6 | 0.651439942 | cat3 | from_4164 | to_4164 | art_long | art_long | M |
| 4 | 7 | 0.927325024 | cat3 | from_4165 | to_4165 | art_long | art_long | M |
| 6 | 8 | 0.65595039 | cat3 | from_4166 | to_4166 | art_long | art_long | M |
| 6 | 9 | 0.620173701 | cat3 | from_4167 | to_4167 | art_long | art_long | M |
| 8 | 9 | 0.895123548 | cat3 | from_4168 | to_4168 | art_long | art_long | M |
| 6 | 10 | 0.618164286 | cat3 | from_4169 | to_4169 | art_long | art_long | M |
| 8 | 10 | 0.878651596 | cat3 | from_4170 | to_4170 | art_long | art_long | M |
| 9 | 10 | 0.997012563 | cat3 | from_4171 | to_4171 | art_long | art_long | M |
| 4 | 11 | 0.789840258 | cat3 | from_4172 | to_4172 | art_long | art_long | M |
| 7 | 11 | 0.756988339 | cat3 | from_4173 | to_4173 | art_long | art_long | M |
| 12 | 13 | 0.778151542 | cat3 | from_4174 | to_4174 | art_long | art_long | M |
| 5 | 15 | 0.691069108 | cat3 | from_4175 | to_4175 | art_long | art_long | M |
| 12 | 15 | 0.650113021 | cat3 | from_4176 | to_4176 | art_long | art_long | M |
| 13 | 15 | 0.847571767 | cat3 | from_4177 | to_4177 | art_long | art_long | M |
| 2 | 16 | 0.617962658 | cat3 | from_4178 | to_4178 | art_long | art_long | M |
| 12 | 16 | 0.633517878 | cat3 | from_4179 | to_4179 | art_long | art_long | M |
| 15 | 16 | 0.610978408 | cat3 | from_4180 | to_4180 | art_long | art_long | M |
| 5 | 18 | 0.634413039 | cat3 | from_4181 | to_4181 | art_long | art_long | M |
| 12 | 18 | 0.818635658 | cat3 | from_4182 | to_4182 | art_long | art_long | M |
| 13 | 18 | 0.681423013 | cat3 | from_4183 | to_4183 | art_long | art_long | M |
| 15 | 18 | 0.80044173 | cat3 | from_4184 | to_4184 | art_long | art_long | M |
| 16 | 18 | 0.762711168 | cat3 | from_4185 | to_4185 | art_long | art_long | M |
| 12 | 19 | 0.952653043 | cat3 | from_4186 | to_4186 | art_long | art_long | M |
| 13 | 19 | 0.826479818 | cat3 | from_4187 | to_4187 | art_long | art_long | M |
| 15 | 19 | 0.617408184 | cat3 | from_4188 | to_4188 | art_long | art_long | M |
| 16 | 19 | 0.635208589 | cat3 | from_4189 | to_4189 | art_long | art_long | M |
| 18 | 19 | 0.778226284 | cat3 | from_4190 | to_4190 | art_long | art_long | M |
| 16 | 21 | 0.603116851 | cat3 | from_4191 | to_4191 | art_long | art_long | M |
| 20 | 21 | 0.866062529 | cat3 | from_4192 | to_4192 | art_long | art_long | M |
| 9 | 22 | 0.707352447 | cat3 | from_4193 | to_4193 | art_long | art_long | M |
| 10 | 22 | 0.721796658 | cat3 | from_4194 | to_4194 | art_long | art_long | M |
| 11 | 22 | 0.644024009 | cat3 | from_4195 | to_4195 | art_long | art_long | M |
| 8 | 23 | 0.697138249 | cat3 | from_4196 | to_4196 | art_long | art_long | M |
| 9 | 23 | 0.919837731 | cat3 | from_4197 | to_4197 | art_long | art_long | M |
| 10 | 23 | 0.930460821 | cat3 | from_4198 | to_4198 | art_long | art_long | M |
| 11 | 23 | 0.611656506 | cat3 | from_4199 | to_4199 | art_long | art_long | M |
| 22 | 23 | 0.831762918 | cat3 | from_4200 | to_4200 | art_long | art_long | M |
| 14 | 24 | 0.656463324 | cat3 | from_4201 | to_4201 | art_long | art_long | M |
| 14 | 25 | 0.746707354 | cat3 | from_4202 | to_4202 | art_long | art_long | M |
| 20 | 27 | -0.634855404 | cat3 | from_4203 | to_4203 | art_long | art_long | M |
| 22 | 27 | 0.748142023 | cat3 | from_4204 | to_4204 | art_long | art_long | M |
| 14 | 28 | 0.791288732 | cat3 | from_4205 | to_4205 | art_long | art_long | M |
| 24 | 28 | 0.940654389 | cat3 | from_4206 | to_4206 | art_long | art_long | M |
| 25 | 28 | 0.671246136 | cat3 | from_4207 | to_4207 | art_long | art_long | M |
| 3 | 29 | 0.613797639 | cat3 | from_4208 | to_4208 | art_long | art_long | M |
| 2 | 31 | -0.628846854 | cat3 | from_4209 | to_4209 | art_long | art_long | M |
| 14 | 31 | 0.629983929 | cat3 | from_4210 | to_4210 | art_long | art_long | M |
| 25 | 31 | 0.888280072 | cat3 | from_4211 | to_4211 | art_long | art_long | M |
| 24 | 32 | 0.688661489 | cat3 | from_4212 | to_4212 | art_long | art_long | M |
| 28 | 32 | 0.625606919 | cat3 | from_4213 | to_4213 | art_long | art_long | M |
| 14 | 33 | 0.633888383 | cat3 | from_4214 | to_4214 | art_long | art_long | M |
| 25 | 33 | 0.914263876 | cat3 | from_4215 | to_4215 | art_long | art_long | M |
| 31 | 33 | 0.99598544 | cat3 | from_4216 | to_4216 | art_long | art_long | M |
| 4 | 34 | 0.709160178 | cat3 | from_4217 | to_4217 | art_long | art_long | M |
| 5 | 34 | 0.678328085 | cat3 | from_4218 | to_4218 | art_long | art_long | M |
| 7 | 34 | 0.625995765 | cat3 | from_4219 | to_4219 | art_long | art_long | M |
| 17 | 35 | 0.715589752 | cat3 | from_4220 | to_4220 | art_long | art_long | M |
| 17 | 36 | 0.965977074 | cat3 | from_4221 | to_4221 | art_long | art_long | M |
| 26 | 37 | 0.961940021 | cat3 | from_4222 | to_4222 | art_long | art_long | M |
| 17 | 38 | 0.756507478 | cat3 | from_4223 | to_4223 | art_long | art_long | M |
| 35 | 38 | 0.980235252 | cat3 | from_4224 | to_4224 | art_long | art_long | M |
| 36 | 38 | 0.668697556 | cat3 | from_4225 | to_4225 | art_long | art_long | M |
| 12 | 39 | 0.745430965 | cat3 | from_4226 | to_4226 | art_long | art_long | M |
| 13 | 39 | 0.640983496 | cat3 | from_4227 | to_4227 | art_long | art_long | M |
| 16 | 39 | 0.917352035 | cat3 | from_4228 | to_4228 | art_long | art_long | M |
| 18 | 39 | 0.833060217 | cat3 | from_4229 | to_4229 | art_long | art_long | M |
| 19 | 39 | 0.770942741 | cat3 | from_4230 | to_4230 | art_long | art_long | M |
| 20 | 39 | 0.651395947 | cat3 | from_4231 | to_4231 | art_long | art_long | M |
| 21 | 39 | 0.634906131 | cat3 | from_4232 | to_4232 | art_long | art_long | M |
| 12 | 40 | 0.823239032 | cat3 | from_4233 | to_4233 | art_long | art_long | M |
| 13 | 40 | 0.720512042 | cat3 | from_4234 | to_4234 | art_long | art_long | M |
| 16 | 40 | 0.792122428 | cat3 | from_4235 | to_4235 | art_long | art_long | M |
| 18 | 40 | 0.643736441 | cat3 | from_4236 | to_4236 | art_long | art_long | M |
| 19 | 40 | 0.90150055 | cat3 | from_4237 | to_4237 | art_long | art_long | M |
| 39 | 40 | 0.868737788 | cat3 | from_4238 | to_4238 | art_long | art_long | M |
| 29 | 41 | 0.742863375 | cat3 | from_4239 | to_4239 | art_long | art_long | M |
| 4 | 42 | 0.612234728 | cat3 | from_4240 | to_4240 | art_long | art_long | M |
| 21 | 42 | 0.665994351 | cat3 | from_4241 | to_4241 | art_long | art_long | M |
| 36 | 43 | 0.604286829 | cat3 | from_4242 | to_4242 | art_long | art_long | M |
| 42 | 43 | 0.658515454 | cat3 | from_4243 | to_4243 | art_long | art_long | M |
| 11 | 44 | 0.655935644 | cat3 | from_4244 | to_4244 | art_long | art_long | M |
| 35 | 44 | 0.662652642 | cat3 | from_4245 | to_4245 | art_long | art_long | M |
| 38 | 44 | 0.659289119 | cat3 | from_4246 | to_4246 | art_long | art_long | M |
| 42 | 44 | 0.644612989 | cat3 | from_4247 | to_4247 | art_long | art_long | M |
| 7 | 45 | 0.604618335 | cat3 | from_4248 | to_4248 | art_long | art_long | M |
| 42 | 45 | 0.967961003 | cat3 | from_4249 | to_4249 | art_long | art_long | M |
| 43 | 45 | 0.699269899 | cat3 | from_4250 | to_4250 | art_long | art_long | M |
| 44 | 45 | 0.657840873 | cat3 | from_4251 | to_4251 | art_long | art_long | M |
| 11 | 46 | 0.649777212 | cat3 | from_4252 | to_4252 | art_long | art_long | M |
| 35 | 46 | 0.675804926 | cat3 | from_4253 | to_4253 | art_long | art_long | M |
| 38 | 46 | 0.650210644 | cat3 | from_4254 | to_4254 | art_long | art_long | M |
| 42 | 46 | 0.655944134 | cat3 | from_4255 | to_4255 | art_long | art_long | M |
| 44 | 46 | 0.993540123 | cat3 | from_4256 | to_4256 | art_long | art_long | M |
| 45 | 46 | 0.674819057 | cat3 | from_4257 | to_4257 | art_long | art_long | M |
| 28 | 47 | 0.617339946 | cat3 | from_4258 | to_4258 | art_long | art_long | M |
| 30 | 47 | 0.85438588 | cat3 | from_4259 | to_4259 | art_long | art_long | M |
| 17 | 48 | 0.702364573 | cat3 | from_4260 | to_4260 | art_long | art_long | M |
| 31 | 48 | -0.677958491 | cat3 | from_4261 | to_4261 | art_long | art_long | M |
| 33 | 48 | -0.667974211 | cat3 | from_4262 | to_4262 | art_long | art_long | M |

| | | | | | | | | |
|---|---|---|---|---|---|---|---|---|
| 35 | 48 | 0.928159089 | cat3 | from_4263 | to_4263 | art_long | art_long | M |
| 36 | 48 | 0.630331721 | cat3 | from_4264 | to_4264 | art_long | art_long | M |
| 38 | 48 | 0.891920123 | cat3 | from_4265 | to_4265 | art_long | art_long | M |
| 44 | 48 | 0.614528432 | cat3 | from_4266 | to_4266 | art_long | art_long | M |
| 46 | 48 | 0.637954594 | cat3 | from_4267 | to_4267 | art_long | art_long | M |
| 35 | 50 | 0.702198115 | cat3 | from_4268 | to_4268 | art_long | art_long | M |
| 38 | 50 | 0.781874327 | cat3 | from_4269 | to_4269 | art_long | art_long | M |
| 48 | 50 | 0.630090817 | cat3 | from_4270 | to_4270 | art_long | art_long | M |
| 44 | 52 | 0.868921947 | cat3 | from_4271 | to_4271 | art_long | art_long | M |
| 46 | 52 | 0.861407826 | cat3 | from_4272 | to_4272 | art_long | art_long | M |
| 2 | 53 | 0.738647595 | cat3 | from_4273 | to_4273 | art_long | art_long | M |
| 16 | 53 | 0.764634327 | cat3 | from_4274 | to_4274 | art_long | art_long | M |
| 39 | 53 | 0.695679706 | cat3 | from_4275 | to_4275 | art_long | art_long | M |
| 25 | 54 | 0.638545639 | cat3 | from_4276 | to_4276 | art_long | art_long | M |
| 42 | 54 | -0.606265244 | cat3 | from_4277 | to_4277 | art_long | art_long | M |
| 24 | 55 | 0.634338222 | cat3 | from_4278 | to_4278 | art_long | art_long | M |
| 27 | 55 | 0.622482694 | cat3 | from_4279 | to_4279 | art_long | art_long | M |
| 28 | 55 | 0.673783176 | cat3 | from_4280 | to_4280 | art_long | art_long | M |
| 30 | 55 | 0.850665704 | cat3 | from_4281 | to_4281 | art_long | art_long | M |
| 47 | 55 | 0.956164227 | cat3 | from_4282 | to_4282 | art_long | art_long | M |
| 22 | 56 | 0.703927666 | cat3 | from_4283 | to_4283 | art_long | art_long | M |
| 22 | 57 | 0.639537649 | cat3 | from_4284 | to_4284 | art_long | art_long | M |
| 56 | 57 | 0.795415565 | cat3 | from_4285 | to_4285 | art_long | art_long | M |
| 14 | 58 | 0.703828342 | cat3 | from_4286 | to_4286 | art_long | art_long | M |
| 30 | 58 | 0.89751696 | cat3 | from_4287 | to_4287 | art_long | art_long | M |
| 47 | 58 | 0.698983049 | cat3 | from_4288 | to_4288 | art_long | art_long | M |
| 54 | 58 | 0.642328169 | cat3 | from_4289 | to_4289 | art_long | art_long | M |
| 55 | 58 | 0.779625174 | cat3 | from_4290 | to_4290 | art_long | art_long | M |
| 14 | 59 | 0.974844266 | cat3 | from_4291 | to_4291 | art_long | art_long | M |
| 24 | 59 | 0.797240821 | cat3 | from_4292 | to_4292 | art_long | art_long | M |
| 25 | 59 | 0.75832106 | cat3 | from_4293 | to_4293 | art_long | art_long | M |
| 28 | 59 | 0.907423832 | cat3 | from_4294 | to_4294 | art_long | art_long | M |
| 33 | 59 | 0.611125577 | cat3 | from_4295 | to_4295 | art_long | art_long | M |
| 58 | 59 | 0.693351097 | cat3 | from_4296 | to_4296 | art_long | art_long | M |
| 11 | 60 | 0.603315525 | cat3 | from_4297 | to_4297 | art_long | art_long | M |
| 38 | 60 | 0.621544737 | cat3 | from_4298 | to_4298 | art_long | art_long | M |
| 43 | 60 | 0.63121775 | cat3 | from_4299 | to_4299 | art_long | art_long | M |
| 44 | 60 | 0.906923628 | cat3 | from_4300 | to_4300 | art_long | art_long | M |
| 46 | 60 | 0.856020043 | cat3 | from_4301 | to_4301 | art_long | art_long | M |
| 52 | 60 | 0.836972159 | cat3 | from_4302 | to_4302 | art_long | art_long | M |
| 43 | 61 | 0.614913438 | cat3 | from_4303 | to_4303 | art_long | art_long | M |
| 44 | 61 | 0.842661251 | cat3 | from_4304 | to_4304 | art_long | art_long | M |
| 46 | 61 | 0.815986587 | cat3 | from_4305 | to_4305 | art_long | art_long | M |
| 50 | 61 | 0.618717756 | cat3 | from_4306 | to_4306 | art_long | art_long | M |
| 52 | 61 | 0.632886286 | cat3 | from_4307 | to_4307 | art_long | art_long | M |
| 60 | 61 | 0.814721384 | cat3 | from_4308 | to_4308 | art_long | art_long | M |
| 4 | 62 | 0.630489472 | cat3 | from_4309 | to_4309 | art_long | art_long | M |
| 7 | 62 | 0.619514703 | cat3 | from_4310 | to_4310 | art_long | art_long | M |
| 17 | 62 | 0.626506192 | cat3 | from_4311 | to_4311 | art_long | art_long | M |
| 42 | 62 | 0.958300543 | cat3 | from_4312 | to_4312 | art_long | art_long | M |
| 43 | 62 | 0.777556156 | cat3 | from_4313 | to_4313 | art_long | art_long | M |
| 44 | 62 | 0.66160169 | cat3 | from_4314 | to_4314 | art_long | art_long | M |
| 45 | 62 | 0.991226636 | cat3 | from_4315 | to_4315 | art_long | art_long | M |
| 46 | 62 | 0.664081459 | cat3 | from_4316 | to_4316 | art_long | art_long | M |
| 5 | 63 | 0.927717049 | cat3 | from_4317 | to_4317 | art_long | art_long | M |
| 13 | 63 | 0.66308378 | cat3 | from_4318 | to_4318 | art_long | art_long | M |
| 15 | 63 | 0.750660717 | cat3 | from_4319 | to_4319 | art_long | art_long | M |
| 18 | 63 | 0.700211056 | cat3 | from_4320 | to_4320 | art_long | art_long | M |
| 24 | 64 | 0.624368111 | cat3 | from_4321 | to_4321 | art_long | art_long | M |
| 27 | 64 | 0.732216452 | cat3 | from_4322 | to_4322 | art_long | art_long | M |
| 28 | 64 | 0.659971739 | cat3 | from_4323 | to_4323 | art_long | art_long | M |
| 34 | 64 | 0.618397693 | cat3 | from_4324 | to_4324 | art_long | art_long | M |
| 50 | 64 | -0.686263453 | cat3 | from_4325 | to_4325 | art_long | art_long | M |
| 22 | 65 | 0.632789327 | cat3 | from_4326 | to_4326 | art_long | art_long | M |
| 23 | 65 | 0.638688631 | cat3 | from_4327 | to_4327 | art_long | art_long | M |
| 34 | 65 | 0.814555822 | cat3 | from_4328 | to_4328 | art_long | art_long | M |
| 64 | 65 | 0.637809106 | cat3 | from_4329 | to_4329 | art_long | art_long | M |
| 29 | 66 | 0.714967432 | cat3 | from_4330 | to_4330 | art_long | art_long | M |
| 41 | 66 | 0.649519247 | cat3 | from_4331 | to_4331 | art_long | art_long | M |
| 29 | 67 | 0.806516366 | cat3 | from_4332 | to_4332 | art_long | art_long | M |
| 41 | 67 | 0.672452995 | cat3 | from_4333 | to_4333 | art_long | art_long | M |
| 66 | 67 | 0.950783038 | cat3 | from_4334 | to_4334 | art_long | art_long | M |
| 12 | 68 | 0.623381723 | cat3 | from_4335 | to_4335 | art_long | art_long | M |
| 13 | 68 | 0.756516527 | cat3 | from_4336 | to_4336 | art_long | art_long | M |
| 15 | 68 | 0.605502025 | cat3 | from_4337 | to_4337 | art_long | art_long | M |
| 18 | 68 | 0.624396124 | cat3 | from_4338 | to_4338 | art_long | art_long | M |
| 19 | 68 | 0.664310644 | cat3 | from_4339 | to_4339 | art_long | art_long | M |
| 25 | 68 | 0.666117607 | cat3 | from_4340 | to_4340 | art_long | art_long | M |
| 31 | 68 | 0.673414337 | cat3 | from_4341 | to_4341 | art_long | art_long | M |
| 33 | 68 | 0.656857722 | cat3 | from_4342 | to_4342 | art_long | art_long | M |
| 48 | 68 | -0.628205749 | cat3 | from_4343 | to_4343 | art_long | art_long | M |
| 14 | 69 | 0.720737157 | cat3 | from_4344 | to_4344 | art_long | art_long | M |
| 30 | 69 | 0.895138672 | cat3 | from_4345 | to_4345 | art_long | art_long | M |
| 47 | 69 | 0.666716096 | cat3 | from_4346 | to_4346 | art_long | art_long | M |
| 54 | 69 | 0.622866333 | cat3 | from_4347 | to_4347 | art_long | art_long | M |
| 55 | 69 | 0.731722275 | cat3 | from_4348 | to_4348 | art_long | art_long | M |
| 58 | 69 | 0.995491551 | cat3 | from_4349 | to_4349 | art_long | art_long | M |
| 59 | 69 | 0.691769656 | cat3 | from_4350 | to_4350 | art_long | art_long | M |
| 49 | 70 | 0.758261845 | cat3 | from_4351 | to_4351 | art_long | art_long | M |
| 57 | 70 | 0.616714355 | cat3 | from_4352 | to_4352 | art_long | art_long | M |
| 3 | 71 | 0.69508696 | cat3 | from_4353 | to_4353 | art_long | art_long | M |
| 29 | 71 | 0.970826527 | cat3 | from_4354 | to_4354 | art_long | art_long | M |
| 41 | 71 | 0.775290285 | cat3 | from_4355 | to_4355 | art_long | art_long | M |
| 66 | 71 | 0.743423766 | cat3 | from_4356 | to_4356 | art_long | art_long | M |
| 67 | 71 | 0.83340511 | cat3 | from_4357 | to_4357 | art_long | art_long | M |
| 30 | 72 | 0.908667701 | cat3 | from_4358 | to_4358 | art_long | art_long | M |
| 33 | 72 | 0.608910913 | cat3 | from_4359 | to_4359 | art_long | art_long | M |
| 47 | 72 | 0.82789468 | cat3 | from_4360 | to_4360 | art_long | art_long | M |
| 55 | 72 | 0.75097252 | cat3 | from_4361 | to_4361 | art_long | art_long | M |
| 58 | 72 | 0.782709517 | cat3 | from_4362 | to_4362 | art_long | art_long | M |
| 69 | 72 | 0.794044298 | cat3 | from_4363 | to_4363 | art_long | art_long | M |
| 14 | 73 | 0.999494755 | cat3 | from_4364 | to_4364 | art_long | art_long | M |
| 24 | 73 | 0.678669247 | cat3 | from_4365 | to_4365 | art_long | art_long | M |
| 25 | 73 | 0.750499722 | cat3 | from_4366 | to_4366 | art_long | art_long | M |

| | | | | | | | | |
|---|---|---|---|---|---|---|---|---|
| 28 | 73 | 0.80487174 | cat3 | from_4367 | to_4367 | art_long | art_long | M |
| 31 | 73 | 0.622680202 | cat3 | from_4368 | to_4368 | art_long | art_long | M |
| 33 | 73 | 0.628704377 | cat3 | from_4369 | to_4369 | art_long | art_long | M |
| 58 | 73 | 0.704901983 | cat3 | from_4370 | to_4370 | art_long | art_long | M |
| 59 | 73 | 0.979729912 | cat3 | from_4371 | to_4371 | art_long | art_long | M |
| 69 | 73 | 0.71900739 | cat3 | from_4372 | to_4372 | art_long | art_long | M |
| 14 | 74 | 0.891511249 | cat3 | from_4373 | to_4373 | art_long | art_long | M |
| 24 | 74 | 0.821788914 | cat3 | from_4374 | to_4374 | art_long | art_long | M |
| 25 | 74 | 0.709769134 | cat3 | from_4375 | to_4375 | art_long | art_long | M |
| 28 | 74 | 0.799169305 | cat3 | from_4376 | to_4376 | art_long | art_long | M |
| 32 | 74 | 0.605647946 | cat3 | from_4377 | to_4377 | art_long | art_long | M |
| 58 | 74 | 0.645906531 | cat3 | from_4378 | to_4378 | art_long | art_long | M |
| 59 | 74 | 0.908803573 | cat3 | from_4379 | to_4379 | art_long | art_long | M |
| 69 | 74 | 0.629781644 | cat3 | from_4380 | to_4380 | art_long | art_long | M |
| 73 | 74 | 0.903974942 | cat3 | from_4381 | to_4381 | art_long | art_long | M |
| 49 | 75 | 0.789340573 | cat3 | from_4382 | to_4382 | art_long | art_long | M |
| 56 | 75 | 0.646811656 | cat3 | from_4383 | to_4383 | art_long | art_long | M |
| 57 | 75 | 0.621854189 | cat3 | from_4384 | to_4384 | art_long | art_long | M |
| 70 | 75 | 0.897242696 | cat3 | from_4385 | to_4385 | art_long | art_long | M |
| 14 | 76 | 0.703193676 | cat3 | from_4386 | to_4386 | art_long | art_long | M |
| 24 | 76 | 0.930195497 | cat3 | from_4387 | to_4387 | art_long | art_long | M |
| 25 | 76 | 0.835188201 | cat3 | from_4388 | to_4388 | art_long | art_long | M |
| 28 | 76 | 0.913795207 | cat3 | from_4389 | to_4389 | art_long | art_long | M |
| 32 | 76 | 0.672363771 | cat3 | from_4390 | to_4390 | art_long | art_long | M |
| 33 | 76 | 0.62531638 | cat3 | from_4391 | to_4391 | art_long | art_long | M |
| 55 | 76 | 0.603977828 | cat3 | from_4392 | to_4392 | art_long | art_long | M |
| 59 | 76 | 0.818622718 | cat3 | from_4393 | to_4393 | art_long | art_long | M |
| 73 | 76 | 0.721006873 | cat3 | from_4394 | to_4394 | art_long | art_long | M |
| 74 | 76 | 0.809939704 | cat3 | from_4395 | to_4395 | art_long | art_long | M |
| 25 | 77 | 0.605635592 | cat3 | from_4396 | to_4396 | art_long | art_long | M |
| 30 | 77 | 0.637905608 | cat3 | from_4397 | to_4397 | art_long | art_long | M |
| 54 | 77 | 0.943797678 | cat3 | from_4398 | to_4398 | art_long | art_long | M |
| 55 | 77 | 0.646756243 | cat3 | from_4399 | to_4399 | art_long | art_long | M |
| 58 | 77 | 0.792796854 | cat3 | from_4400 | to_4400 | art_long | art_long | M |
| 69 | 77 | 0.756617285 | cat3 | from_4401 | to_4401 | art_long | art_long | M |
| 74 | 77 | 0.645170639 | cat3 | from_4402 | to_4402 | art_long | art_long | M |
| 76 | 77 | 0.63120904 | cat3 | from_4403 | to_4403 | art_long | art_long | M |
| 14 | 78 | 0.778014894 | cat3 | from_4404 | to_4404 | art_long | art_long | M |
| 24 | 78 | 0.91305249 | cat3 | from_4405 | to_4405 | art_long | art_long | M |
| 25 | 78 | 0.658241779 | cat3 | from_4406 | to_4406 | art_long | art_long | M |
| 28 | 78 | 0.830971863 | cat3 | from_4407 | to_4407 | art_long | art_long | M |
| 32 | 78 | 0.665018321 | cat3 | from_4408 | to_4408 | art_long | art_long | M |
| 59 | 78 | 0.843663828 | cat3 | from_4409 | to_4409 | art_long | art_long | M |
| 73 | 78 | 0.797154936 | cat3 | from_4410 | to_4410 | art_long | art_long | M |
| 74 | 78 | 0.969341016 | cat3 | from_4411 | to_4411 | art_long | art_long | M |
| 76 | 78 | 0.867818309 | cat3 | from_4412 | to_4412 | art_long | art_long | M |
| 77 | 78 | 0.671188133 | cat3 | from_4413 | to_4413 | art_long | art_long | M |
| 12 | 79 | 0.968299136 | cat3 | from_4414 | to_4414 | art_long | art_long | M |
| 13 | 79 | 0.811122671 | cat3 | from_4415 | to_4415 | art_long | art_long | M |
| 15 | 79 | 0.605112793 | cat3 | from_4416 | to_4416 | art_long | art_long | M |
| 18 | 79 | 0.770935726 | cat3 | from_4417 | to_4417 | art_long | art_long | M |
| 19 | 79 | 0.977893739 | cat3 | from_4418 | to_4418 | art_long | art_long | M |
| 39 | 79 | 0.685042884 | cat3 | from_4419 | to_4419 | art_long | art_long | M |
| 40 | 79 | 0.807468565 | cat3 | from_4420 | to_4420 | art_long | art_long | M |
| 68 | 79 | 0.696309032 | cat3 | from_4421 | to_4421 | art_long | art_long | M |
| 24 | 80 | 0.60590222 | cat3 | from_4422 | to_4422 | art_long | art_long | M |
| 27 | 80 | 0.608576402 | cat3 | from_4423 | to_4423 | art_long | art_long | M |
| 30 | 80 | 0.618775832 | cat3 | from_4424 | to_4424 | art_long | art_long | M |
| 54 | 80 | 0.899129775 | cat3 | from_4425 | to_4425 | art_long | art_long | M |
| 55 | 80 | 0.705493088 | cat3 | from_4426 | to_4426 | art_long | art_long | M |
| 58 | 80 | 0.733209949 | cat3 | from_4427 | to_4427 | art_long | art_long | M |
| 69 | 80 | 0.683042592 | cat3 | from_4428 | to_4428 | art_long | art_long | M |
| 76 | 80 | 0.663117903 | cat3 | from_4429 | to_4429 | art_long | art_long | M |
| 77 | 80 | 0.983164452 | cat3 | from_4430 | to_4430 | art_long | art_long | M |
| 78 | 80 | 0.659081407 | cat3 | from_4431 | to_4431 | art_long | art_long | M |
| 25 | 81 | 0.679603483 | cat3 | from_4432 | to_4432 | art_long | art_long | M |
| 31 | 81 | 0.652012297 | cat3 | from_4433 | to_4433 | art_long | art_long | M |
| 33 | 81 | 0.640206376 | cat3 | from_4434 | to_4434 | art_long | art_long | M |
| 54 | 81 | 0.940512153 | cat3 | from_4435 | to_4435 | art_long | art_long | M |
| 58 | 81 | 0.675252632 | cat3 | from_4436 | to_4436 | art_long | art_long | M |
| 68 | 81 | 0.651837203 | cat3 | from_4437 | to_4437 | art_long | art_long | M |
| 69 | 81 | 0.686255491 | cat3 | from_4438 | to_4438 | art_long | art_long | M |
| 77 | 81 | 0.84434973 | cat3 | from_4439 | to_4439 | art_long | art_long | M |
| 80 | 81 | 0.758945046 | cat3 | from_4440 | to_4440 | art_long | art_long | M |
| 3 | 82 | 0.718967378 | cat3 | from_4441 | to_4441 | art_long | art_long | M |
| 29 | 82 | 0.71650428 | cat3 | from_4442 | to_4442 | art_long | art_long | M |
| 41 | 82 | 0.671041906 | cat3 | from_4443 | to_4443 | art_long | art_long | M |
| 66 | 82 | 0.673189148 | cat3 | from_4444 | to_4444 | art_long | art_long | M |
| 67 | 82 | 0.683353717 | cat3 | from_4445 | to_4445 | art_long | art_long | M |
| 71 | 82 | 0.782315254 | cat3 | from_4446 | to_4446 | art_long | art_long | M |
| 14 | 83 | 0.616581305 | cat3 | from_4447 | to_4447 | art_long | art_long | M |
| 25 | 83 | 0.680954563 | cat3 | from_4448 | to_4448 | art_long | art_long | M |
| 30 | 83 | 0.788568437 | cat3 | from_4449 | to_4449 | art_long | art_long | M |
| 31 | 83 | 0.64645979 | cat3 | from_4450 | to_4450 | art_long | art_long | M |
| 33 | 83 | 0.63810488 | cat3 | from_4451 | to_4451 | art_long | art_long | M |
| 54 | 83 | 0.839209206 | cat3 | from_4452 | to_4452 | art_long | art_long | M |
| 58 | 83 | 0.892461374 | cat3 | from_4453 | to_4453 | art_long | art_long | M |
| 68 | 83 | 0.618273956 | cat3 | from_4454 | to_4454 | art_long | art_long | M |
| 69 | 83 | 0.905469859 | cat3 | from_4455 | to_4455 | art_long | art_long | M |
| 72 | 83 | 0.745044767 | cat3 | from_4456 | to_4456 | art_long | art_long | M |
| 73 | 83 | 0.614163655 | cat3 | from_4457 | to_4457 | art_long | art_long | M |
| 77 | 83 | 0.846068414 | cat3 | from_4458 | to_4458 | art_long | art_long | M |
| 80 | 83 | 0.756737264 | cat3 | from_4459 | to_4459 | art_long | art_long | M |
| 81 | 83 | 0.925468126 | cat3 | from_4460 | to_4460 | art_long | art_long | M |
| 8 | 84 | 0.703400566 | cat3 | from_4461 | to_4461 | art_long | art_long | M |
| 9 | 84 | 0.635126289 | cat3 | from_4462 | to_4462 | art_long | art_long | M |
| 10 | 84 | 0.64047466 | cat3 | from_4463 | to_4463 | art_long | art_long | M |
| 49 | 84 | 0.744662401 | cat3 | from_4464 | to_4464 | art_long | art_long | M |
| 70 | 84 | 0.714351246 | cat3 | from_4465 | to_4465 | art_long | art_long | M |
| 75 | 84 | 0.888523339 | cat3 | from_4466 | to_4466 | art_long | art_long | M |
| 34 | 85 | 0.729606489 | cat3 | from_4467 | to_4467 | art_long | art_long | M |
| 49 | 85 | 0.855285894 | cat3 | from_4468 | to_4468 | art_long | art_long | M |
| 54 | 85 | -0.633585171 | cat3 | from_4469 | to_4469 | art_long | art_long | M |
| 56 | 85 | 0.708723711 | cat3 | from_4470 | to_4470 | art_long | art_long | M |

| | | | | | | | | |
|---|---|---|---|---|---|---|---|---|
| 65 | 85 | 0.627841123 | cat3 | from_4471 | to_4471 | art_long | art_long | M |
| 70 | 85 | 0.706545752 | cat3 | from_4472 | to_4472 | art_long | art_long | M |
| 75 | 85 | 0.695418288 | cat3 | from_4473 | to_4473 | art_long | art_long | M |
| 30 | 86 | 0.633372149 | cat3 | from_4474 | to_4474 | art_long | art_long | M |
| 44 | 86 | 0.636514385 | cat3 | from_4475 | to_4475 | art_long | art_long | M |
| 46 | 86 | 0.64063625 | cat3 | from_4476 | to_4476 | art_long | art_long | M |
| 47 | 86 | 0.646051697 | cat3 | from_4477 | to_4477 | art_long | art_long | M |
| 55 | 86 | 0.613305416 | cat3 | from_4478 | to_4478 | art_long | art_long | M |
| 72 | 86 | 0.671045291 | cat3 | from_4479 | to_4479 | art_long | art_long | M |
| 28 | 87 | 0.644855572 | cat3 | from_4480 | to_4480 | art_long | art_long | M |
| 30 | 87 | 0.917955341 | cat3 | from_4481 | to_4481 | art_long | art_long | M |
| 47 | 87 | 0.982032365 | cat3 | from_4482 | to_4482 | art_long | art_long | M |
| 55 | 87 | 0.968970641 | cat3 | from_4483 | to_4483 | art_long | art_long | M |
| 58 | 87 | 0.82136758 | cat3 | from_4484 | to_4484 | art_long | art_long | M |
| 69 | 87 | 0.794229346 | cat3 | from_4485 | to_4485 | art_long | art_long | M |
| 72 | 87 | 0.865991809 | cat3 | from_4486 | to_4486 | art_long | art_long | M |
| 86 | 87 | 0.664526625 | cat3 | from_4487 | to_4487 | art_long | art_long | M |
| 8 | 88 | 0.695078091 | cat3 | from_4488 | to_4488 | art_long | art_long | M |
| 9 | 88 | 0.940842006 | cat3 | from_4489 | to_4489 | art_long | art_long | M |
| 10 | 88 | 0.946167075 | cat3 | from_4490 | to_4490 | art_long | art_long | M |
| 22 | 88 | 0.897572076 | cat3 | from_4491 | to_4491 | art_long | art_long | M |
| 23 | 88 | 0.973411897 | cat3 | from_4492 | to_4492 | art_long | art_long | M |
| 57 | 88 | 0.620887109 | cat3 | from_4493 | to_4493 | art_long | art_long | M |
| 65 | 88 | 0.613977557 | cat3 | from_4494 | to_4494 | art_long | art_long | M |
| 3 | 89 | 0.607604019 | cat3 | from_4495 | to_4495 | art_long | art_long | M |
| 29 | 89 | 0.848509845 | cat3 | from_4496 | to_4496 | art_long | art_long | M |
| 41 | 89 | 0.660593121 | cat3 | from_4497 | to_4497 | art_long | art_long | M |
| 66 | 89 | 0.645471835 | cat3 | from_4498 | to_4498 | art_long | art_long | M |
| 67 | 89 | 0.725726132 | cat3 | from_4499 | to_4499 | art_long | art_long | M |
| 71 | 89 | 0.874542807 | cat3 | from_4500 | to_4500 | art_long | art_long | M |
| 82 | 89 | 0.915003929 | cat3 | from_4501 | to_4501 | art_long | art_long | M |
| 25 | 90 | 0.67758213 | cat3 | from_4502 | to_4502 | art_long | art_long | M |
| 28 | 90 | 0.710652629 | cat3 | from_4503 | to_4503 | art_long | art_long | M |
| 30 | 90 | 0.608178509 | cat3 | from_4504 | to_4504 | art_long | art_long | M |
| 31 | 90 | 0.713334957 | cat3 | from_4505 | to_4505 | art_long | art_long | M |
| 33 | 90 | 0.748449479 | cat3 | from_4506 | to_4506 | art_long | art_long | M |
| 37 | 90 | 0.629257345 | cat3 | from_4507 | to_4507 | art_long | art_long | M |
| 47 | 90 | 0.665527607 | cat3 | from_4508 | to_4508 | art_long | art_long | M |
| 57 | 90 | -0.660116536 | cat3 | from_4509 | to_4509 | art_long | art_long | M |
| 59 | 90 | 0.636439888 | cat3 | from_4510 | to_4510 | art_long | art_long | M |
| 72 | 90 | 0.704112304 | cat3 | from_4511 | to_4511 | art_long | art_long | M |
| 76 | 90 | 0.642051533 | cat3 | from_4512 | to_4512 | art_long | art_long | M |
| 87 | 90 | 0.632298657 | cat3 | from_4513 | to_4513 | art_long | art_long | M |
| 23 | 91 | 0.697144591 | cat3 | from_4514 | to_4514 | art_long | art_long | M |
| 27 | 91 | 0.687224848 | cat3 | from_4515 | to_4515 | art_long | art_long | M |
| 30 | 91 | 0.627547077 | cat3 | from_4516 | to_4516 | art_long | art_long | M |
| 58 | 91 | 0.625376497 | cat3 | from_4517 | to_4517 | art_long | art_long | M |
| 69 | 91 | 0.641147293 | cat3 | from_4518 | to_4518 | art_long | art_long | M |
| 72 | 91 | 0.663195948 | cat3 | from_4519 | to_4519 | art_long | art_long | M |
| 88 | 91 | 0.619896121 | cat3 | from_4520 | to_4520 | art_long | art_long | M |
| 21 | 92 | 0.692982708 | cat3 | from_4521 | to_4521 | art_long | art_long | M |
| 42 | 92 | 0.992639028 | cat3 | from_4522 | to_4522 | art_long | art_long | M |
| 43 | 92 | 0.60223735 | cat3 | from_4523 | to_4523 | art_long | art_long | M |
| 44 | 92 | 0.620822357 | cat3 | from_4524 | to_4524 | art_long | art_long | M |
| 45 | 92 | 0.932214561 | cat3 | from_4525 | to_4525 | art_long | art_long | M |
| 46 | 92 | 0.633677473 | cat3 | from_4526 | to_4526 | art_long | art_long | M |
| 54 | 92 | -0.64036952 | cat3 | from_4527 | to_4527 | art_long | art_long | M |
| 62 | 92 | 0.91734072 | cat3 | from_4528 | to_4528 | art_long | art_long | M |
| 8 | 93 | 0.715021048 | cat3 | from_4529 | to_4529 | art_long | art_long | M |
| 19 | 93 | 0.646227104 | cat3 | from_4530 | to_4530 | art_long | art_long | M |
| 54 | 93 | 0.75763991 | cat3 | from_4531 | to_4531 | art_long | art_long | M |
| 79 | 93 | 0.719347103 | cat3 | from_4532 | to_4532 | art_long | art_long | M |
| 81 | 93 | 0.686905858 | cat3 | from_4533 | to_4533 | art_long | art_long | M |
| 92 | 93 | -0.623613713 | cat3 | from_4534 | to_4534 | art_long | art_long | M |
| 30 | 94 | 0.613445915 | cat3 | from_4535 | to_4535 | art_long | art_long | M |
| 32 | 94 | 0.658331645 | cat3 | from_4536 | to_4536 | art_long | art_long | M |
| 44 | 94 | 0.634892879 | cat3 | from_4537 | to_4537 | art_long | art_long | M |
| 46 | 94 | 0.635490619 | cat3 | from_4538 | to_4538 | art_long | art_long | M |
| 47 | 94 | 0.611893757 | cat3 | from_4539 | to_4539 | art_long | art_long | M |
| 55 | 94 | 0.622340696 | cat3 | from_4540 | to_4540 | art_long | art_long | M |
| 72 | 94 | 0.617402062 | cat3 | from_4541 | to_4541 | art_long | art_long | M |
| 86 | 94 | 0.988889074 | cat3 | from_4542 | to_4542 | art_long | art_long | M |
| 87 | 94 | 0.642904427 | cat3 | from_4543 | to_4543 | art_long | art_long | M |
| 24 | 95 | 0.735839536 | cat3 | from_4544 | to_4544 | art_long | art_long | M |
| 27 | 95 | 0.709714633 | cat3 | from_4545 | to_4545 | art_long | art_long | M |
| 28 | 95 | 0.652346425 | cat3 | from_4546 | to_4546 | art_long | art_long | M |
| 30 | 95 | 0.694920376 | cat3 | from_4547 | to_4547 | art_long | art_long | M |
| 47 | 95 | 0.788803893 | cat3 | from_4548 | to_4548 | art_long | art_long | M |
| 55 | 95 | 0.931050063 | cat3 | from_4549 | to_4549 | art_long | art_long | M |
| 58 | 95 | 0.711333776 | cat3 | from_4550 | to_4550 | art_long | art_long | M |
| 69 | 95 | 0.64311436 | cat3 | from_4551 | to_4551 | art_long | art_long | M |
| 76 | 95 | 0.665340923 | cat3 | from_4552 | to_4552 | art_long | art_long | M |
| 77 | 95 | 0.776555403 | cat3 | from_4553 | to_4553 | art_long | art_long | M |
| 78 | 95 | 0.6087449 | cat3 | from_4554 | to_4554 | art_long | art_long | M |
| 80 | 95 | 0.850200303 | cat3 | from_4555 | to_4555 | art_long | art_long | M |
| 87 | 95 | 0.818280276 | cat3 | from_4556 | to_4556 | art_long | art_long | M |
| 41 | 96 | 0.811306276 | cat3 | from_4557 | to_4557 | art_long | art_long | M |
| 66 | 96 | 0.621064171 | cat3 | from_4558 | to_4558 | art_long | art_long | M |
| 82 | 96 | 0.6105959 | cat3 | from_4559 | to_4559 | art_long | art_long | M |
| 14 | 97 | 0.745057039 | cat3 | from_4560 | to_4560 | art_long | art_long | M |
| 24 | 97 | 0.637516427 | cat3 | from_4561 | to_4561 | art_long | art_long | M |
| 26 | 97 | 0.684563602 | cat3 | from_4562 | to_4562 | art_long | art_long | M |
| 28 | 97 | 0.858011176 | cat3 | from_4563 | to_4563 | art_long | art_long | M |
| 30 | 97 | 0.613846412 | cat3 | from_4564 | to_4564 | art_long | art_long | M |
| 37 | 97 | 0.683216797 | cat3 | from_4565 | to_4565 | art_long | art_long | M |
| 47 | 97 | 0.703495201 | cat3 | from_4566 | to_4566 | art_long | art_long | M |
| 55 | 97 | 0.608745182 | cat3 | from_4567 | to_4567 | art_long | art_long | M |
| 59 | 97 | 0.817367179 | cat3 | from_4568 | to_4568 | art_long | art_long | M |
| 64 | 97 | 0.603616248 | cat3 | from_4569 | to_4569 | art_long | art_long | M |
| 72 | 97 | 0.63428883 | cat3 | from_4570 | to_4570 | art_long | art_long | M |
| 73 | 97 | 0.74364639 | cat3 | from_4571 | to_4571 | art_long | art_long | M |
| 76 | 97 | 0.669017146 | cat3 | from_4572 | to_4572 | art_long | art_long | M |
| 87 | 97 | 0.698226822 | cat3 | from_4573 | to_4573 | art_long | art_long | M |
| 90 | 97 | 0.899565499 | cat3 | from_4574 | to_4574 | art_long | art_long | M |

| | | | | | | | | |
|---|---|---|---|---|---|---|---|---|
| 29 | 98 | 0.704098761 | cat3 | from_4575 | to_4575 | art_long | art_long | M |
| 41 | 98 | 0.72693675 | cat3 | from_4576 | to_4576 | art_long | art_long | M |
| 71 | 98 | 0.714821811 | cat3 | from_4577 | to_4577 | art_long | art_long | M |
| 82 | 98 | 0.676048855 | cat3 | from_4578 | to_4578 | art_long | art_long | M |
| 89 | 98 | 0.837604014 | cat3 | from_4579 | to_4579 | art_long | art_long | M |
| 44 | 99 | 0.890835929 | cat3 | from_4580 | to_4580 | art_long | art_long | M |
| 46 | 99 | 0.868212559 | cat3 | from_4581 | to_4581 | art_long | art_long | M |
| 50 | 99 | 0.643937489 | cat3 | from_4582 | to_4582 | art_long | art_long | M |
| 52 | 99 | 0.738240186 | cat3 | from_4583 | to_4583 | art_long | art_long | M |
| 60 | 99 | 0.902813235 | cat3 | from_4584 | to_4584 | art_long | art_long | M |
| 61 | 99 | 0.859877207 | cat3 | from_4585 | to_4585 | art_long | art_long | M |
| 86 | 99 | 0.646007839 | cat3 | from_4586 | to_4586 | art_long | art_long | M |
| 94 | 99 | 0.621321825 | cat3 | from_4587 | to_4587 | art_long | art_long | M |
| 42 | 100 | 0.909967489 | cat3 | from_4588 | to_4588 | art_long | art_long | M |
| 44 | 100 | 0.614594972 | cat3 | from_4589 | to_4589 | art_long | art_long | M |
| 45 | 100 | 0.967019097 | cat3 | from_4590 | to_4590 | art_long | art_long | M |
| 46 | 100 | 0.658701399 | cat3 | from_4591 | to_4591 | art_long | art_long | M |
| 62 | 100 | 0.928449513 | cat3 | from_4592 | to_4592 | art_long | art_long | M |
| 81 | 100 | -0.656984588 | cat3 | from_4593 | to_4593 | art_long | art_long | M |
| 92 | 100 | 0.873288612 | cat3 | from_4594 | to_4594 | art_long | art_long | M |
| 4 | 101 | 0.654905218 | cat3 | from_4595 | to_4595 | art_long | art_long | M |
| 17 | 101 | 0.621621107 | cat3 | from_4596 | to_4596 | art_long | art_long | M |
| 21 | 101 | 0.62409066 | cat3 | from_4597 | to_4597 | art_long | art_long | M |
| 42 | 101 | 0.977872022 | cat3 | from_4598 | to_4598 | art_long | art_long | M |
| 43 | 101 | 0.783089003 | cat3 | from_4599 | to_4599 | art_long | art_long | M |
| 44 | 101 | 0.658978443 | cat3 | from_4600 | to_4600 | art_long | art_long | M |
| 45 | 101 | 0.973535393 | cat3 | from_4601 | to_4601 | art_long | art_long | M |
| 46 | 101 | 0.652894641 | cat3 | from_4602 | to_4602 | art_long | art_long | M |
| 60 | 101 | 0.611596902 | cat3 | from_4603 | to_4603 | art_long | art_long | M |
| 62 | 101 | 0.98792391 | cat3 | from_4604 | to_4604 | art_long | art_long | M |
| 92 | 101 | 0.951359121 | cat3 | from_4605 | to_4605 | art_long | art_long | M |
| 100 | 101 | 0.884995793 | cat3 | from_4606 | to_4606 | art_long | art_long | M |
| 35 | 102 | 0.647931666 | cat3 | from_4607 | to_4607 | art_long | art_long | M |
| 42 | 102 | 0.612350017 | cat3 | from_4608 | to_4608 | art_long | art_long | M |
| 44 | 102 | 0.887672803 | cat3 | from_4609 | to_4609 | art_long | art_long | M |
| 45 | 102 | 0.659607915 | cat3 | from_4610 | to_4610 | art_long | art_long | M |
| 46 | 102 | 0.933716522 | cat3 | from_4611 | to_4611 | art_long | art_long | M |
| 48 | 102 | 0.664560427 | cat3 | from_4612 | to_4612 | art_long | art_long | M |
| 52 | 102 | 0.768441163 | cat3 | from_4613 | to_4613 | art_long | art_long | M |
| 60 | 102 | 0.620549238 | cat3 | from_4614 | to_4614 | art_long | art_long | M |
| 61 | 102 | 0.684869324 | cat3 | from_4615 | to_4615 | art_long | art_long | M |
| 62 | 102 | 0.60639321 | cat3 | from_4616 | to_4616 | art_long | art_long | M |
| 99 | 102 | 0.717112821 | cat3 | from_4617 | to_4617 | art_long | art_long | M |
| 100 | 102 | 0.733166638 | cat3 | from_4618 | to_4618 | art_long | art_long | M |
| 42 | 103 | 0.605822903 | cat3 | from_4619 | to_4619 | art_long | art_long | M |
| 54 | 103 | -0.688557156 | cat3 | from_4620 | to_4620 | art_long | art_long | M |
| 81 | 103 | -0.632707398 | cat3 | from_4621 | to_4621 | art_long | art_long | M |
| 90 | 103 | 0.619887217 | cat3 | from_4622 | to_4622 | art_long | art_long | M |
| 92 | 103 | 0.652042555 | cat3 | from_4623 | to_4623 | art_long | art_long | M |
| 93 | 103 | -0.79296352 | cat3 | from_4624 | to_4624 | art_long | art_long | M |
| 97 | 103 | 0.633403257 | cat3 | from_4625 | to_4625 | art_long | art_long | M |
| 17 | 104 | 0.706070091 | cat3 | from_4626 | to_4626 | art_long | art_long | M |
| 31 | 104 | -0.678846278 | cat3 | from_4627 | to_4627 | art_long | art_long | M |
| 33 | 104 | -0.670851959 | cat3 | from_4628 | to_4628 | art_long | art_long | M |
| 35 | 104 | 0.825113842 | cat3 | from_4629 | to_4629 | art_long | art_long | M |
| 36 | 104 | 0.700467116 | cat3 | from_4630 | to_4630 | art_long | art_long | M |
| 38 | 104 | 0.817300828 | cat3 | from_4631 | to_4631 | art_long | art_long | M |
| 48 | 104 | 0.965116525 | cat3 | from_4632 | to_4632 | art_long | art_long | M |
| 50 | 104 | 0.628158097 | cat3 | from_4633 | to_4633 | art_long | art_long | M |
| 20 | 105 | 0.61966305 | cat3 | from_4634 | to_4634 | art_long | art_long | M |
| 44 | 105 | 0.697722764 | cat3 | from_4635 | to_4635 | art_long | art_long | M |
| 46 | 105 | 0.67440154 | cat3 | from_4636 | to_4636 | art_long | art_long | M |
| 50 | 105 | 0.668571562 | cat3 | from_4637 | to_4637 | art_long | art_long | M |
| 60 | 105 | 0.751437407 | cat3 | from_4638 | to_4638 | art_long | art_long | M |
| 61 | 105 | 0.77887459 | cat3 | from_4639 | to_4639 | art_long | art_long | M |
| 99 | 105 | 0.944274253 | cat3 | from_4640 | to_4640 | art_long | art_long | M |
| 11 | 106 | 0.619955159 | cat3 | from_4641 | to_4641 | art_long | art_long | M |
| 35 | 106 | 0.612470054 | cat3 | from_4642 | to_4642 | art_long | art_long | M |
| 38 | 106 | 0.61543913 | cat3 | from_4643 | to_4643 | art_long | art_long | M |
| 44 | 106 | 0.973702385 | cat3 | from_4644 | to_4644 | art_long | art_long | M |
| 45 | 106 | 0.617458039 | cat3 | from_4645 | to_4645 | art_long | art_long | M |
| 46 | 106 | 0.964074461 | cat3 | from_4646 | to_4646 | art_long | art_long | M |
| 48 | 106 | 0.645586963 | cat3 | from_4647 | to_4647 | art_long | art_long | M |
| 52 | 106 | 0.818459742 | cat3 | from_4648 | to_4648 | art_long | art_long | M |
| 60 | 106 | 0.868681031 | cat3 | from_4649 | to_4649 | art_long | art_long | M |
| 61 | 106 | 0.898419937 | cat3 | from_4650 | to_4650 | art_long | art_long | M |
| 62 | 106 | 0.630777707 | cat3 | from_4651 | to_4651 | art_long | art_long | M |
| 94 | 106 | 0.610597266 | cat3 | from_4652 | to_4652 | art_long | art_long | M |
| 99 | 106 | 0.825461653 | cat3 | from_4653 | to_4653 | art_long | art_long | M |
| 102 | 106 | 0.862798664 | cat3 | from_4654 | to_4654 | art_long | art_long | M |
| 104 | 106 | 0.642293761 | cat3 | from_4655 | to_4655 | art_long | art_long | M |
| 105 | 106 | 0.621341449 | cat3 | from_4656 | to_4656 | art_long | art_long | M |
| 4 | 107 | 0.634222346 | cat3 | from_4657 | to_4657 | art_long | art_long | M |
| 7 | 107 | 0.61549189 | cat3 | from_4658 | to_4658 | art_long | art_long | M |
| 17 | 107 | 0.668659054 | cat3 | from_4659 | to_4659 | art_long | art_long | M |
| 36 | 107 | 0.704001852 | cat3 | from_4660 | to_4660 | art_long | art_long | M |
| 42 | 107 | 0.747277584 | cat3 | from_4661 | to_4661 | art_long | art_long | M |
| 43 | 107 | 0.932431206 | cat3 | from_4662 | to_4662 | art_long | art_long | M |
| 45 | 107 | 0.812048667 | cat3 | from_4663 | to_4663 | art_long | art_long | M |
| 60 | 107 | 0.630536832 | cat3 | from_4664 | to_4664 | art_long | art_long | M |
| 62 | 107 | 0.879612707 | cat3 | from_4665 | to_4665 | art_long | art_long | M |
| 92 | 107 | 0.67936226 | cat3 | from_4666 | to_4666 | art_long | art_long | M |
| 100 | 107 | 0.670388515 | cat3 | from_4667 | to_4667 | art_long | art_long | M |
| 101 | 107 | 0.869730065 | cat3 | from_4668 | to_4668 | art_long | art_long | M |
| 106 | 107 | 0.609463152 | cat3 | from_4669 | to_4669 | art_long | art_long | M |
| 36 | 108 | 0.674949822 | cat3 | from_4670 | to_4670 | art_long | art_long | M |
| 70 | 108 | 0.604295817 | cat3 | from_4671 | to_4671 | art_long | art_long | M |
| 75 | 108 | 0.618780895 | cat3 | from_4672 | to_4672 | art_long | art_long | M |
| 24 | 109 | 0.654516832 | cat3 | from_4673 | to_4673 | art_long | art_long | M |
| 27 | 109 | 0.647500404 | cat3 | from_4674 | to_4674 | art_long | art_long | M |
| 28 | 109 | 0.679700576 | cat3 | from_4675 | to_4675 | art_long | art_long | M |
| 30 | 109 | 0.869641204 | cat3 | from_4676 | to_4676 | art_long | art_long | M |
| 47 | 109 | 0.921776106 | cat3 | from_4677 | to_4677 | art_long | art_long | M |
| 55 | 109 | 0.992294041 | cat3 | from_4678 | to_4678 | art_long | art_long | M |

| | | | | | | | | |
|---|---|---|---|---|---|---|---|---|
| 58 | 109 | 0.837610936 | cat3 | from_4679 | to_4679 | art_long | art_long | M |
| 69 | 109 | 0.791409006 | cat3 | from_4680 | to_4680 | art_long | art_long | M |
| 72 | 109 | 0.747033345 | cat3 | from_4681 | to_4681 | art_long | art_long | M |
| 76 | 109 | 0.627788529 | cat3 | from_4682 | to_4682 | art_long | art_long | M |
| 77 | 109 | 0.725713685 | cat3 | from_4683 | to_4683 | art_long | art_long | M |
| 80 | 109 | 0.769838481 | cat3 | from_4684 | to_4684 | art_long | art_long | M |
| 83 | 109 | 0.621847296 | cat3 | from_4685 | to_4685 | art_long | art_long | M |
| 86 | 109 | 0.605975674 | cat3 | from_4686 | to_4686 | art_long | art_long | M |
| 87 | 109 | 0.95691702 | cat3 | from_4687 | to_4687 | art_long | art_long | M |
| 94 | 109 | 0.627058194 | cat3 | from_4688 | to_4688 | art_long | art_long | M |
| 95 | 109 | 0.946491139 | cat3 | from_4689 | to_4689 | art_long | art_long | M |
| 14 | 110 | 0.96733871 | cat3 | from_4690 | to_4690 | art_long | art_long | M |
| 25 | 110 | 0.76541431 | cat3 | from_4691 | to_4691 | art_long | art_long | M |
| 28 | 110 | 0.702446601 | cat3 | from_4692 | to_4692 | art_long | art_long | M |
| 31 | 110 | 0.724286964 | cat3 | from_4693 | to_4693 | art_long | art_long | M |
| 33 | 110 | 0.716099219 | cat3 | from_4694 | to_4694 | art_long | art_long | M |
| 58 | 110 | 0.670442245 | cat3 | from_4695 | to_4695 | art_long | art_long | M |
| 59 | 110 | 0.921312951 | cat3 | from_4696 | to_4696 | art_long | art_long | M |
| 69 | 110 | 0.695488624 | cat3 | from_4697 | to_4697 | art_long | art_long | M |
| 73 | 110 | 0.965849598 | cat3 | from_4698 | to_4698 | art_long | art_long | M |
| 74 | 110 | 0.874141826 | cat3 | from_4699 | to_4699 | art_long | art_long | M |
| 76 | 110 | 0.659304407 | cat3 | from_4700 | to_4700 | art_long | art_long | M |
| 78 | 110 | 0.751611256 | cat3 | from_4701 | to_4701 | art_long | art_long | M |
| 83 | 110 | 0.662941821 | cat3 | from_4702 | to_4702 | art_long | art_long | M |
| 97 | 110 | 0.640139719 | cat3 | from_4703 | to_4703 | art_long | art_long | M |
| 30 | 111 | 0.874105332 | cat3 | from_4704 | to_4704 | art_long | art_long | M |
| 47 | 111 | 0.947218718 | cat3 | from_4705 | to_4705 | art_long | art_long | M |
| 55 | 111 | 0.835845365 | cat3 | from_4706 | to_4706 | art_long | art_long | M |
| 58 | 111 | 0.70114846 | cat3 | from_4707 | to_4707 | art_long | art_long | M |
| 69 | 111 | 0.698751941 | cat3 | from_4708 | to_4708 | art_long | art_long | M |
| 72 | 111 | 0.908832576 | cat3 | from_4709 | to_4709 | art_long | art_long | M |
| 86 | 111 | 0.65825365 | cat3 | from_4710 | to_4710 | art_long | art_long | M |
| 87 | 111 | 0.939711566 | cat3 | from_4711 | to_4711 | art_long | art_long | M |
| 90 | 111 | 0.728065332 | cat3 | from_4712 | to_4712 | art_long | art_long | M |
| 97 | 111 | 0.757700719 | cat3 | from_4713 | to_4713 | art_long | art_long | M |
| 109 | 111 | 0.800290349 | cat3 | from_4714 | to_4714 | art_long | art_long | M |
| 14 | 112 | 0.79702599 | cat3 | from_4715 | to_4715 | art_long | art_long | M |
| 25 | 112 | 0.972659175 | cat3 | from_4716 | to_4716 | art_long | art_long | M |
| 31 | 112 | 0.916204286 | cat3 | from_4717 | to_4717 | art_long | art_long | M |
| 33 | 112 | 0.923036636 | cat3 | from_4718 | to_4718 | art_long | art_long | M |
| 48 | 112 | -0.610767041 | cat3 | from_4719 | to_4719 | art_long | art_long | M |
| 51 | 112 | 0.624175173 | cat3 | from_4720 | to_4720 | art_long | art_long | M |
| 54 | 112 | 0.620796176 | cat3 | from_4721 | to_4721 | art_long | art_long | M |
| 59 | 112 | 0.758365489 | cat3 | from_4722 | to_4722 | art_long | art_long | M |
| 68 | 112 | 0.742230598 | cat3 | from_4723 | to_4723 | art_long | art_long | M |
| 73 | 112 | 0.794788933 | cat3 | from_4724 | to_4724 | art_long | art_long | M |
| 74 | 112 | 0.701206514 | cat3 | from_4725 | to_4725 | art_long | art_long | M |
| 76 | 112 | 0.715612979 | cat3 | from_4726 | to_4726 | art_long | art_long | M |
| 81 | 112 | 0.721135005 | cat3 | from_4727 | to_4727 | art_long | art_long | M |
| 83 | 112 | 0.733615875 | cat3 | from_4728 | to_4728 | art_long | art_long | M |
| 90 | 112 | 0.619316206 | cat3 | from_4729 | to_4729 | art_long | art_long | M |
| 110 | 112 | 0.828573463 | cat3 | from_4730 | to_4730 | art_long | art_long | M |
| 22 | 113 | 0.644433212 | cat3 | from_4731 | to_4731 | art_long | art_long | M |
| 34 | 113 | 0.765501725 | cat3 | from_4732 | to_4732 | art_long | art_long | M |
| 64 | 113 | 0.739812487 | cat3 | from_4733 | to_4733 | art_long | art_long | M |
| 65 | 113 | 0.877686034 | cat3 | from_4734 | to_4734 | art_long | art_long | M |
| 12 | 114 | 0.736514708 | cat3 | from_4735 | to_4735 | art_long | art_long | M |
| 13 | 114 | 0.929112052 | cat3 | from_4736 | to_4736 | art_long | art_long | M |
| 15 | 114 | 0.633097344 | cat3 | from_4737 | to_4737 | art_long | art_long | M |
| 19 | 114 | 0.794398125 | cat3 | from_4738 | to_4738 | art_long | art_long | M |
| 20 | 114 | 0.712029771 | cat3 | from_4739 | to_4739 | art_long | art_long | M |
| 39 | 114 | 0.655514965 | cat3 | from_4740 | to_4740 | art_long | art_long | M |
| 40 | 114 | 0.762006706 | cat3 | from_4741 | to_4741 | art_long | art_long | M |
| 68 | 114 | 0.758868692 | cat3 | from_4742 | to_4742 | art_long | art_long | M |
| 79 | 114 | 0.783077485 | cat3 | from_4743 | to_4743 | art_long | art_long | M |
| 26 | 115 | 0.948076908 | cat3 | from_4744 | to_4744 | art_long | art_long | M |
| 37 | 115 | 0.928837925 | cat3 | from_4745 | to_4745 | art_long | art_long | M |
| 90 | 115 | 0.669247033 | cat3 | from_4746 | to_4746 | art_long | art_long | M |
| 93 | 115 | -0.614108933 | cat3 | from_4747 | to_4747 | art_long | art_long | M |
| 97 | 115 | 0.72888287 | cat3 | from_4748 | to_4748 | art_long | art_long | M |
| 111 | 115 | 0.62822465 | cat3 | from_4749 | to_4749 | art_long | art_long | M |
| 5 | 116 | 0.866977936 | cat3 | from_4750 | to_4750 | art_long | art_long | M |
| 16 | 116 | 0.620050787 | cat3 | from_4751 | to_4751 | art_long | art_long | M |
| 63 | 116 | 0.799268074 | cat3 | from_4752 | to_4752 | art_long | art_long | M |
| 65 | 116 | 0.657639653 | cat3 | from_4753 | to_4753 | art_long | art_long | M |
| 34 | 117 | 0.74600089 | cat3 | from_4754 | to_4754 | art_long | art_long | M |
| 64 | 117 | 0.855913314 | cat3 | from_4755 | to_4755 | art_long | art_long | M |
| 65 | 117 | 0.634006086 | cat3 | from_4756 | to_4756 | art_long | art_long | M |
| 91 | 117 | 0.722770838 | cat3 | from_4757 | to_4757 | art_long | art_long | M |
| 97 | 117 | 0.728741973 | cat3 | from_4758 | to_4758 | art_long | art_long | M |
| 26 | 118 | 0.732724384 | cat3 | from_4759 | to_4759 | art_long | art_long | M |
| 37 | 118 | 0.70847862 | cat3 | from_4760 | to_4760 | art_long | art_long | M |
| 55 | 118 | 0.623929717 | cat3 | from_4761 | to_4761 | art_long | art_long | M |
| 58 | 118 | 0.633808417 | cat3 | from_4762 | to_4762 | art_long | art_long | M |
| 69 | 118 | 0.612609147 | cat3 | from_4763 | to_4763 | art_long | art_long | M |
| 87 | 118 | 0.621695948 | cat3 | from_4764 | to_4764 | art_long | art_long | M |
| 109 | 118 | 0.643913926 | cat3 | from_4765 | to_4765 | art_long | art_long | M |
| 115 | 118 | 0.603979782 | cat3 | from_4766 | to_4766 | art_long | art_long | M |
| 24 | 119 | 0.647142794 | cat3 | from_4767 | to_4767 | art_long | art_long | M |
| 27 | 119 | 0.655398015 | cat3 | from_4768 | to_4768 | art_long | art_long | M |
| 28 | 119 | 0.643373218 | cat3 | from_4769 | to_4769 | art_long | art_long | M |
| 30 | 119 | 0.853397698 | cat3 | from_4770 | to_4770 | art_long | art_long | M |
| 47 | 119 | 0.749857494 | cat3 | from_4771 | to_4771 | art_long | art_long | M |
| 54 | 119 | 0.6854846 | cat3 | from_4772 | to_4772 | art_long | art_long | M |
| 55 | 119 | 0.882478389 | cat3 | from_4773 | to_4773 | art_long | art_long | M |
| 58 | 119 | 0.94725034 | cat3 | from_4774 | to_4774 | art_long | art_long | M |
| 59 | 119 | 0.64341149 | cat3 | from_4775 | to_4775 | art_long | art_long | M |
| 69 | 119 | 0.913052655 | cat3 | from_4776 | to_4776 | art_long | art_long | M |
| 72 | 119 | 0.689119496 | cat3 | from_4777 | to_4777 | art_long | art_long | M |
| 73 | 119 | 0.602784983 | cat3 | from_4778 | to_4778 | art_long | art_long | M |
| 74 | 119 | 0.656355918 | cat3 | from_4779 | to_4779 | art_long | art_long | M |
| 76 | 119 | 0.633940615 | cat3 | from_4780 | to_4780 | art_long | art_long | M |
| 77 | 119 | 0.875935392 | cat3 | from_4781 | to_4781 | art_long | art_long | M |
| 78 | 119 | 0.664354027 | cat3 | from_4782 | to_4782 | art_long | art_long | M |

| | | | | | | | | |
|---|---|---|---|---|---|---|---|---|
| 80 | 119 | 0.867435102 | cat3 | from_4783 | to_4783 | art_long | art_long | M |
| 81 | 119 | 0.617337964 | cat3 | from_4784 | to_4784 | art_long | art_long | M |
| 83 | 119 | 0.80229444 | cat3 | from_4785 | to_4785 | art_long | art_long | M |
| 87 | 119 | 0.84877046 | cat3 | from_4786 | to_4786 | art_long | art_long | M |
| 95 | 119 | 0.893293354 | cat3 | from_4787 | to_4787 | art_long | art_long | M |
| 109 | 119 | 0.933396064 | cat3 | from_4788 | to_4788 | art_long | art_long | M |
| 111 | 119 | 0.649147057 | cat3 | from_4789 | to_4789 | art_long | art_long | M |
| 118 | 119 | 0.654852897 | cat3 | from_4790 | to_4790 | art_long | art_long | M |
| 54 | 120 | 0.944583376 | cat3 | from_4791 | to_4791 | art_long | art_long | M |
| 58 | 120 | 0.697217193 | cat3 | from_4792 | to_4792 | art_long | art_long | M |
| 69 | 120 | 0.684377937 | cat3 | from_4793 | to_4793 | art_long | art_long | M |
| 74 | 120 | 0.695052246 | cat3 | from_4794 | to_4794 | art_long | art_long | M |
| 77 | 120 | 0.912581219 | cat3 | from_4795 | to_4795 | art_long | art_long | M |
| 78 | 120 | 0.662655402 | cat3 | from_4796 | to_4796 | art_long | art_long | M |
| 80 | 120 | 0.838180036 | cat3 | from_4797 | to_4797 | art_long | art_long | M |
| 81 | 120 | 0.886473964 | cat3 | from_4798 | to_4798 | art_long | art_long | M |
| 83 | 120 | 0.827208829 | cat3 | from_4799 | to_4799 | art_long | art_long | M |
| 93 | 120 | 0.757897245 | cat3 | from_4800 | to_4800 | art_long | art_long | M |
| 103 | 120 | -0.741803513 | cat3 | from_4801 | to_4801 | art_long | art_long | M |
| 119 | 120 | 0.710844072 | cat3 | from_4802 | to_4802 | art_long | art_long | M |
| 29 | 121 | 0.628941047 | cat3 | from_4803 | to_4803 | art_long | art_long | M |
| 41 | 121 | 0.875367281 | cat3 | from_4804 | to_4804 | art_long | art_long | M |
| 71 | 121 | 0.654714126 | cat3 | from_4805 | to_4805 | art_long | art_long | M |
| 82 | 121 | 0.79164097 | cat3 | from_4806 | to_4806 | art_long | art_long | M |
| 89 | 121 | 0.785868346 | cat3 | from_4807 | to_4807 | art_long | art_long | M |
| 96 | 121 | 0.752789638 | cat3 | from_4808 | to_4808 | art_long | art_long | M |
| 98 | 121 | 0.841438916 | cat3 | from_4809 | to_4809 | art_long | art_long | M |
| 14 | 122 | 0.716660391 | cat3 | from_4810 | to_4810 | art_long | art_long | M |
| 24 | 122 | 0.994603534 | cat3 | from_4811 | to_4811 | art_long | art_long | M |
| 25 | 122 | 0.631653888 | cat3 | from_4812 | to_4812 | art_long | art_long | M |
| 28 | 122 | 0.970025522 | cat3 | from_4813 | to_4813 | art_long | art_long | M |
| 32 | 122 | 0.680164014 | cat3 | from_4814 | to_4814 | art_long | art_long | M |
| 55 | 122 | 0.648668509 | cat3 | from_4815 | to_4815 | art_long | art_long | M |
| 59 | 122 | 0.848235027 | cat3 | from_4816 | to_4816 | art_long | art_long | M |
| 64 | 122 | 0.636193206 | cat3 | from_4817 | to_4817 | art_long | art_long | M |
| 73 | 122 | 0.736404149 | cat3 | from_4818 | to_4818 | art_long | art_long | M |
| 74 | 122 | 0.837103749 | cat3 | from_4819 | to_4819 | art_long | art_long | M |
| 76 | 122 | 0.937704945 | cat3 | from_4820 | to_4820 | art_long | art_long | M |
| 78 | 122 | 0.908153713 | cat3 | from_4821 | to_4821 | art_long | art_long | M |
| 95 | 122 | 0.716262489 | cat3 | from_4822 | to_4822 | art_long | art_long | M |
| 97 | 122 | 0.709021549 | cat3 | from_4823 | to_4823 | art_long | art_long | M |
| 109 | 122 | 0.667150443 | cat3 | from_4824 | to_4824 | art_long | art_long | M |
| 110 | 122 | 0.637369485 | cat3 | from_4825 | to_4825 | art_long | art_long | M |
| 119 | 122 | 0.658520507 | cat3 | from_4826 | to_4826 | art_long | art_long | M |
| 34 | 123 | 0.696513333 | cat3 | from_4827 | to_4827 | art_long | art_long | M |
| 49 | 123 | 0.741494126 | cat3 | from_4828 | to_4828 | art_long | art_long | M |
| 56 | 123 | 0.739833998 | cat3 | from_4829 | to_4829 | art_long | art_long | M |
| 57 | 123 | 0.638144165 | cat3 | from_4830 | to_4830 | art_long | art_long | M |
| 70 | 123 | 0.909879656 | cat3 | from_4831 | to_4831 | art_long | art_long | M |
| 75 | 123 | 0.837993486 | cat3 | from_4832 | to_4832 | art_long | art_long | M |
| 85 | 123 | 0.887235824 | cat3 | from_4833 | to_4833 | art_long | art_long | M |
| 14 | 124 | 0.61201098 | cat3 | from_4834 | to_4834 | art_long | art_long | M |
| 29 | 124 | 0.768587889 | cat3 | from_4835 | to_4835 | art_long | art_long | M |
| 41 | 124 | 0.687551585 | cat3 | from_4836 | to_4836 | art_long | art_long | M |
| 66 | 124 | 0.627476252 | cat3 | from_4837 | to_4837 | art_long | art_long | M |
| 67 | 124 | 0.685918896 | cat3 | from_4838 | to_4838 | art_long | art_long | M |
| 71 | 124 | 0.800005187 | cat3 | from_4839 | to_4839 | art_long | art_long | M |
| 73 | 124 | 0.609119565 | cat3 | from_4840 | to_4840 | art_long | art_long | M |
| 82 | 124 | 0.669250869 | cat3 | from_4841 | to_4841 | art_long | art_long | M |
| 89 | 124 | 0.679152288 | cat3 | from_4842 | to_4842 | art_long | art_long | M |
| 6 | 125 | 0.763700496 | cat3 | from_4843 | to_4843 | art_long | art_long | M |
| 8 | 125 | 0.685849143 | cat3 | from_4844 | to_4844 | art_long | art_long | M |
| 9 | 125 | 0.699049129 | cat3 | from_4845 | to_4845 | art_long | art_long | M |
| 10 | 125 | 0.719999576 | cat3 | from_4846 | to_4846 | art_long | art_long | M |
| 23 | 125 | 0.671309901 | cat3 | from_4847 | to_4847 | art_long | art_long | M |
| 49 | 125 | 0.645901221 | cat3 | from_4848 | to_4848 | art_long | art_long | M |
| 65 | 125 | 0.785623219 | cat3 | from_4849 | to_4849 | art_long | art_long | M |
| 75 | 125 | 0.646405459 | cat3 | from_4850 | to_4850 | art_long | art_long | M |
| 84 | 125 | 0.795283618 | cat3 | from_4851 | to_4851 | art_long | art_long | M |
| 88 | 125 | 0.60568863 | cat3 | from_4852 | to_4852 | art_long | art_long | M |
| 8 | 126 | 0.732983196 | cat3 | from_4853 | to_4853 | art_long | art_long | M |
| 9 | 126 | 0.916222222 | cat3 | from_4854 | to_4854 | art_long | art_long | M |
| 10 | 126 | 0.896875359 | cat3 | from_4855 | to_4855 | art_long | art_long | M |
| 22 | 126 | 0.820539681 | cat3 | from_4856 | to_4856 | art_long | art_long | M |
| 23 | 126 | 0.849201472 | cat3 | from_4857 | to_4857 | art_long | art_long | M |
| 57 | 126 | 0.730690896 | cat3 | from_4858 | to_4858 | art_long | art_long | M |
| 88 | 126 | 0.932420661 | cat3 | from_4859 | to_4859 | art_long | art_long | M |
| 93 | 126 | 0.617833554 | cat3 | from_4860 | to_4860 | art_long | art_long | M |
| 4 | 127 | 0.70188879 | cat3 | from_4861 | to_4861 | art_long | art_long | M |
| 5 | 127 | 0.707183493 | cat3 | from_4862 | to_4862 | art_long | art_long | M |
| 34 | 127 | 0.933111516 | cat3 | from_4863 | to_4863 | art_long | art_long | M |
| 49 | 127 | 0.715185071 | cat3 | from_4864 | to_4864 | art_long | art_long | M |
| 65 | 127 | 0.846056185 | cat3 | from_4865 | to_4865 | art_long | art_long | M |
| 70 | 127 | 0.624849398 | cat3 | from_4866 | to_4866 | art_long | art_long | M |
| 85 | 127 | 0.766476177 | cat3 | from_4867 | to_4867 | art_long | art_long | M |
| 113 | 127 | 0.650650995 | cat3 | from_4868 | to_4868 | art_long | art_long | M |
| 117 | 127 | 0.771741565 | cat3 | from_4869 | to_4869 | art_long | art_long | M |
| 123 | 127 | 0.698682008 | cat3 | from_4870 | to_4870 | art_long | art_long | M |
| 125 | 127 | 0.625007338 | cat3 | from_4871 | to_4871 | art_long | art_long | M |
| 11 | 128 | 0.633780638 | cat3 | from_4872 | to_4872 | art_long | art_long | M |
| 38 | 128 | 0.642244302 | cat3 | from_4873 | to_4873 | art_long | art_long | M |
| 43 | 128 | 0.615260013 | cat3 | from_4874 | to_4874 | art_long | art_long | M |
| 44 | 128 | 0.966648228 | cat3 | from_4875 | to_4875 | art_long | art_long | M |
| 46 | 128 | 0.931681958 | cat3 | from_4876 | to_4876 | art_long | art_long | M |
| 52 | 128 | 0.844768111 | cat3 | from_4877 | to_4877 | art_long | art_long | M |
| 60 | 128 | 0.974166072 | cat3 | from_4878 | to_4878 | art_long | art_long | M |
| 61 | 128 | 0.87593353 | cat3 | from_4879 | to_4879 | art_long | art_long | M |
| 62 | 128 | 0.620202761 | cat3 | from_4880 | to_4880 | art_long | art_long | M |
| 94 | 128 | 0.60212893 | cat3 | from_4881 | to_4881 | art_long | art_long | M |
| 99 | 128 | 0.893549899 | cat3 | from_4882 | to_4882 | art_long | art_long | M |
| 101 | 128 | 0.63181589 | cat3 | from_4883 | to_4883 | art_long | art_long | M |
| 102 | 128 | 0.744982901 | cat3 | from_4884 | to_4884 | art_long | art_long | M |
| 105 | 128 | 0.710103432 | cat3 | from_4885 | to_4885 | art_long | art_long | M |
| 106 | 128 | 0.956857638 | cat3 | from_4886 | to_4886 | art_long | art_long | M |

| | | | | | | | | |
|---|---|---|---|---|---|---|---|---|
| 107 | 128 | 0.650542451 | cat3 | from_4887 | to_4887 | art_long | art_long | M |
| 26 | 129 | 0.823089277 | cat3 | from_4888 | to_4888 | art_long | art_long | M |
| 37 | 129 | 0.921715993 | cat3 | from_4889 | to_4889 | art_long | art_long | M |
| 90 | 129 | 0.604604634 | cat3 | from_4890 | to_4890 | art_long | art_long | M |
| 115 | 129 | 0.756134312 | cat3 | from_4891 | to_4891 | art_long | art_long | M |
| 118 | 129 | 0.653217346 | cat3 | from_4892 | to_4892 | art_long | art_long | M |
| 26 | 130 | 0.94091948 | cat3 | from_4893 | to_4893 | art_long | art_long | M |
| 37 | 130 | 0.969844277 | cat3 | from_4894 | to_4894 | art_long | art_long | M |
| 97 | 130 | 0.606908684 | cat3 | from_4895 | to_4895 | art_long | art_long | M |
| 115 | 130 | 0.833570932 | cat3 | from_4896 | to_4896 | art_long | art_long | M |
| 118 | 130 | 0.761657724 | cat3 | from_4897 | to_4897 | art_long | art_long | M |
| 129 | 130 | 0.942415303 | cat3 | from_4898 | to_4898 | art_long | art_long | M |
| 6 | 131 | -0.673263194 | cat3 | from_4899 | to_4899 | art_long | art_long | M |
| 20 | 131 | -0.63251498 | cat3 | from_4900 | to_4900 | art_long | art_long | M |
| 55 | 131 | 0.664514219 | cat3 | from_4901 | to_4901 | art_long | art_long | M |
| 95 | 131 | 0.734141169 | cat3 | from_4902 | to_4902 | art_long | art_long | M |
| 109 | 131 | 0.674659693 | cat3 | from_4903 | to_4903 | art_long | art_long | M |
| 118 | 131 | 0.917964132 | cat3 | from_4904 | to_4904 | art_long | art_long | M |
| 119 | 131 | 0.636463086 | cat3 | from_4905 | to_4905 | art_long | art_long | M |
| 130 | 131 | 0.65446709 | cat3 | from_4906 | to_4906 | art_long | art_long | M |
| 17 | 132 | 0.780656021 | cat3 | from_4907 | to_4907 | art_long | art_long | M |
| 35 | 132 | 0.965340441 | cat3 | from_4908 | to_4908 | art_long | art_long | M |
| 36 | 132 | 0.716273837 | cat3 | from_4909 | to_4909 | art_long | art_long | M |
| 38 | 132 | 0.99455135 | cat3 | from_4910 | to_4910 | art_long | art_long | M |
| 44 | 132 | 0.656090208 | cat3 | from_4911 | to_4911 | art_long | art_long | M |
| 46 | 132 | 0.641328922 | cat3 | from_4912 | to_4912 | art_long | art_long | M |
| 48 | 132 | 0.90443394 | cat3 | from_4913 | to_4913 | art_long | art_long | M |
| 50 | 132 | 0.791817227 | cat3 | from_4914 | to_4914 | art_long | art_long | M |
| 60 | 132 | 0.626269182 | cat3 | from_4915 | to_4915 | art_long | art_long | M |
| 104 | 132 | 0.855536031 | cat3 | from_4916 | to_4916 | art_long | art_long | M |
| 106 | 132 | 0.633433145 | cat3 | from_4917 | to_4917 | art_long | art_long | M |
| 128 | 132 | 0.654601642 | cat3 | from_4918 | to_4918 | art_long | art_long | M |
| 17 | 133 | 0.7552673 | cat3 | from_4919 | to_4919 | art_long | art_long | M |
| 35 | 133 | 0.72376648 | cat3 | from_4920 | to_4920 | art_long | art_long | M |
| 38 | 133 | 0.730259051 | cat3 | from_4921 | to_4921 | art_long | art_long | M |
| 42 | 133 | 0.71775946 | cat3 | from_4922 | to_4922 | art_long | art_long | M |
| 54 | 133 | -0.639513815 | cat3 | from_4923 | to_4923 | art_long | art_long | M |
| 77 | 133 | -0.616692442 | cat3 | from_4924 | to_4924 | art_long | art_long | M |
| 80 | 133 | -0.655076002 | cat3 | from_4925 | to_4925 | art_long | art_long | M |
| 92 | 133 | 0.752740383 | cat3 | from_4926 | to_4926 | art_long | art_long | M |
| 101 | 133 | 0.667364922 | cat3 | from_4927 | to_4927 | art_long | art_long | M |
| 132 | 133 | 0.687519806 | cat3 | from_4928 | to_4928 | art_long | art_long | M |
| 8 | 134 | -0.618309539 | cat3 | from_4929 | to_4929 | art_long | art_long | M |
| 17 | 134 | 0.848008276 | cat3 | from_4930 | to_4930 | art_long | art_long | M |
| 35 | 134 | 0.747743836 | cat3 | from_4931 | to_4931 | art_long | art_long | M |
| 36 | 134 | 0.737392478 | cat3 | from_4932 | to_4932 | art_long | art_long | M |
| 38 | 134 | 0.682208494 | cat3 | from_4933 | to_4933 | art_long | art_long | M |
| 45 | 134 | 0.656332709 | cat3 | from_4934 | to_4934 | art_long | art_long | M |
| 48 | 134 | 0.813916864 | cat3 | from_4935 | to_4935 | art_long | art_long | M |
| 62 | 134 | 0.623196366 | cat3 | from_4936 | to_4936 | art_long | art_long | M |
| 68 | 134 | -0.607138114 | cat3 | from_4937 | to_4937 | art_long | art_long | M |
| 81 | 134 | -0.642262453 | cat3 | from_4938 | to_4938 | art_long | art_long | M |
| 83 | 134 | -0.605715436 | cat3 | from_4939 | to_4939 | art_long | art_long | M |
| 100 | 134 | 0.708286528 | cat3 | from_4940 | to_4940 | art_long | art_long | M |
| 104 | 134 | 0.75955833 | cat3 | from_4941 | to_4941 | art_long | art_long | M |
| 132 | 134 | 0.686019073 | cat3 | from_4942 | to_4942 | art_long | art_long | M |
| 133 | 134 | 0.646555876 | cat3 | from_4943 | to_4943 | art_long | art_long | M |
| 44 | 135 | 0.645343947 | cat3 | from_4944 | to_4944 | art_long | art_long | M |
| 46 | 135 | 0.626956311 | cat3 | from_4945 | to_4945 | art_long | art_long | M |
| 49 | 135 | -0.665103509 | cat3 | from_4946 | to_4946 | art_long | art_long | M |
| 61 | 135 | 0.808790378 | cat3 | from_4947 | to_4947 | art_long | art_long | M |
| 90 | 135 | -0.637108389 | cat3 | from_4948 | to_4948 | art_long | art_long | M |
| 104 | 135 | 0.689080228 | cat3 | from_4949 | to_4949 | art_long | art_long | M |
| 106 | 135 | 0.802297307 | cat3 | from_4950 | to_4950 | art_long | art_long | M |
| 115 | 135 | -0.648067351 | cat3 | from_4951 | to_4951 | art_long | art_long | M |
| 128 | 135 | 0.682835083 | cat3 | from_4952 | to_4952 | art_long | art_long | M |
| 26 | 136 | 0.793812404 | cat3 | from_4953 | to_4953 | art_long | art_long | M |
| 37 | 136 | 0.7956052 | cat3 | from_4954 | to_4954 | art_long | art_long | M |
| 74 | 136 | 0.640245247 | cat3 | from_4955 | to_4955 | art_long | art_long | M |
| 78 | 136 | 0.614609826 | cat3 | from_4956 | to_4956 | art_long | art_long | M |
| 108 | 136 | 0.694108392 | cat3 | from_4957 | to_4957 | art_long | art_long | M |
| 118 | 136 | 0.716977613 | cat3 | from_4958 | to_4958 | art_long | art_long | M |
| 129 | 136 | 0.82499974 | cat3 | from_4959 | to_4959 | art_long | art_long | M |
| 130 | 136 | 0.904550225 | cat3 | from_4960 | to_4960 | art_long | art_long | M |
| 131 | 136 | 0.642023473 | cat3 | from_4961 | to_4961 | art_long | art_long | M |
| 54 | 137 | 0.68904101 | cat3 | from_4962 | to_4962 | art_long | art_long | M |
| 58 | 137 | 0.636059775 | cat3 | from_4963 | to_4963 | art_long | art_long | M |
| 69 | 137 | 0.63361659 | cat3 | from_4964 | to_4964 | art_long | art_long | M |
| 77 | 137 | 0.691568573 | cat3 | from_4965 | to_4965 | art_long | art_long | M |
| 80 | 137 | 0.619703018 | cat3 | from_4966 | to_4966 | art_long | art_long | M |
| 81 | 137 | 0.675193451 | cat3 | from_4967 | to_4967 | art_long | art_long | M |
| 83 | 137 | 0.692963725 | cat3 | from_4968 | to_4968 | art_long | art_long | M |
| 100 | 137 | -0.604198554 | cat3 | from_4969 | to_4969 | art_long | art_long | M |
| 118 | 137 | 0.800578374 | cat3 | from_4970 | to_4970 | art_long | art_long | M |
| 119 | 137 | 0.608752073 | cat3 | from_4971 | to_4971 | art_long | art_long | M |
| 120 | 137 | 0.751816632 | cat3 | from_4972 | to_4972 | art_long | art_long | M |
| 131 | 137 | 0.687214845 | cat3 | from_4973 | to_4973 | art_long | art_long | M |
| 136 | 137 | 0.778371301 | cat3 | from_4974 | to_4974 | art_long | art_long | M |
| 17 | 138 | 0.979544578 | cat3 | from_4975 | to_4975 | art_long | art_long | M |
| 35 | 138 | 0.680207485 | cat3 | from_4976 | to_4976 | art_long | art_long | M |
| 36 | 138 | 0.96421495 | cat3 | from_4977 | to_4977 | art_long | art_long | M |
| 38 | 138 | 0.71579332 | cat3 | from_4978 | to_4978 | art_long | art_long | M |
| 48 | 138 | 0.72579358 | cat3 | from_4979 | to_4979 | art_long | art_long | M |
| 104 | 138 | 0.755673251 | cat3 | from_4980 | to_4980 | art_long | art_long | M |
| 107 | 138 | 0.645007962 | cat3 | from_4981 | to_4981 | art_long | art_long | M |
| 108 | 138 | 0.605830213 | cat3 | from_4982 | to_4982 | art_long | art_long | M |
| 132 | 138 | 0.746824972 | cat3 | from_4983 | to_4983 | art_long | art_long | M |
| 133 | 138 | 0.648594493 | cat3 | from_4984 | to_4984 | art_long | art_long | M |
| 134 | 138 | 0.865500141 | cat3 | from_4985 | to_4985 | art_long | art_long | M |
| 32 | 139 | 0.62695533 | cat3 | from_4986 | to_4986 | art_long | art_long | M |
| 42 | 139 | 0.662404786 | cat3 | from_4987 | to_4987 | art_long | art_long | M |
| 43 | 139 | 0.71763333 | cat3 | from_4988 | to_4988 | art_long | art_long | M |
| 45 | 139 | 0.810214004 | cat3 | from_4989 | to_4989 | art_long | art_long | M |
| 62 | 139 | 0.825995605 | cat3 | from_4990 | to_4990 | art_long | art_long | M |

| | | | | | | | | |
|---|---|---|---|---|---|---|---|---|
| 91 | 139 | -0.614929085 | cat3 | from_4991 | to_4991 | art_long | art_long | M |
| 100 | 139 | 0.804448902 | cat3 | from_4992 | to_4992 | art_long | art_long | M |
| 101 | 139 | 0.742817269 | cat3 | from_4993 | to_4993 | art_long | art_long | M |
| 107 | 139 | 0.804028018 | cat3 | from_4994 | to_4994 | art_long | art_long | M |
| 135 | 139 | 0.640642891 | cat3 | from_4995 | to_4995 | art_long | art_long | M |
| 25 | 140 | 0.68545839 | cat3 | from_4996 | to_4996 | art_long | art_long | M |
| 30 | 140 | 0.801349386 | cat3 | from_4997 | to_4997 | art_long | art_long | M |
| 54 | 140 | 0.880310261 | cat3 | from_4998 | to_4998 | art_long | art_long | M |
| 55 | 140 | 0.654430214 | cat3 | from_4999 | to_4999 | art_long | art_long | M |
| 58 | 140 | 0.911774399 | cat3 | from_5000 | to_5000 | art_long | art_long | M |
| 69 | 140 | 0.906201223 | cat3 | from_5001 | to_5001 | art_long | art_long | M |
| 72 | 140 | 0.74114007 | cat3 | from_5002 | to_5002 | art_long | art_long | M |
| 74 | 140 | 0.606330169 | cat3 | from_5003 | to_5003 | art_long | art_long | M |
| 77 | 140 | 0.923546843 | cat3 | from_5004 | to_5004 | art_long | art_long | M |
| 80 | 140 | 0.862791575 | cat3 | from_5005 | to_5005 | art_long | art_long | M |
| 81 | 140 | 0.904009685 | cat3 | from_5006 | to_5006 | art_long | art_long | M |
| 83 | 140 | 0.98018088 | cat3 | from_5007 | to_5007 | art_long | art_long | M |
| 87 | 140 | 0.665168445 | cat3 | from_5008 | to_5008 | art_long | art_long | M |
| 95 | 140 | 0.653802645 | cat3 | from_5009 | to_5009 | art_long | art_long | M |
| 109 | 140 | 0.728364109 | cat3 | from_5010 | to_5010 | art_long | art_long | M |
| 110 | 140 | 0.611407686 | cat3 | from_5011 | to_5011 | art_long | art_long | M |
| 112 | 140 | 0.697195417 | cat3 | from_5012 | to_5012 | art_long | art_long | M |
| 119 | 140 | 0.880501651 | cat3 | from_5013 | to_5013 | art_long | art_long | M |
| 120 | 140 | 0.850819827 | cat3 | from_5014 | to_5014 | art_long | art_long | M |
| 137 | 140 | 0.699705287 | cat3 | from_5015 | to_5015 | art_long | art_long | M |
| 37 | 141 | 0.618596381 | cat3 | from_5016 | to_5016 | art_long | art_long | M |
| 58 | 141 | 0.673353506 | cat3 | from_5017 | to_5017 | art_long | art_long | M |
| 69 | 141 | 0.683745886 | cat3 | from_5018 | to_5018 | art_long | art_long | M |
| 81 | 141 | 0.627361953 | cat3 | from_5019 | to_5019 | art_long | art_long | M |
| 83 | 141 | 0.707338264 | cat3 | from_5020 | to_5020 | art_long | art_long | M |
| 118 | 141 | 0.85551063 | cat3 | from_5021 | to_5021 | art_long | art_long | M |
| 120 | 141 | 0.624052267 | cat3 | from_5022 | to_5022 | art_long | art_long | M |
| 130 | 141 | 0.67699416 | cat3 | from_5023 | to_5023 | art_long | art_long | M |
| 131 | 141 | 0.671736825 | cat3 | from_5024 | to_5024 | art_long | art_long | M |
| 136 | 141 | 0.769781108 | cat3 | from_5025 | to_5025 | art_long | art_long | M |
| 137 | 141 | 0.966550811 | cat3 | from_5026 | to_5026 | art_long | art_long | M |
| 140 | 141 | 0.692385992 | cat3 | from_5027 | to_5027 | art_long | art_long | M |
| 14 | 142 | 0.606556779 | cat3 | from_5028 | to_5028 | art_long | art_long | M |
| 25 | 142 | 0.669084305 | cat3 | from_5029 | to_5029 | art_long | art_long | M |
| 26 | 142 | 0.730727992 | cat3 | from_5030 | to_5030 | art_long | art_long | M |
| 31 | 142 | 0.604744384 | cat3 | from_5031 | to_5031 | art_long | art_long | M |
| 33 | 142 | 0.604261008 | cat3 | from_5032 | to_5032 | art_long | art_long | M |
| 37 | 142 | 0.806757462 | cat3 | from_5033 | to_5033 | art_long | art_long | M |
| 73 | 142 | 0.604144586 | cat3 | from_5034 | to_5034 | art_long | art_long | M |
| 108 | 142 | 0.679853907 | cat3 | from_5035 | to_5035 | art_long | art_long | M |
| 110 | 142 | 0.616736683 | cat3 | from_5036 | to_5036 | art_long | art_long | M |
| 112 | 142 | 0.710306655 | cat3 | from_5037 | to_5037 | art_long | art_long | M |
| 118 | 142 | 0.608497065 | cat3 | from_5038 | to_5038 | art_long | art_long | M |
| 129 | 142 | 0.835663514 | cat3 | from_5039 | to_5039 | art_long | art_long | M |
| 130 | 142 | 0.858746329 | cat3 | from_5040 | to_5040 | art_long | art_long | M |
| 136 | 142 | 0.923006769 | cat3 | from_5041 | to_5041 | art_long | art_long | M |
| 137 | 142 | 0.763068702 | cat3 | from_5042 | to_5042 | art_long | art_long | M |
| 141 | 142 | 0.803882415 | cat3 | from_5043 | to_5043 | art_long | art_long | M |
| 17 | 143 | 0.666059505 | cat3 | from_5044 | to_5044 | art_long | art_long | M |
| 35 | 143 | 0.793661803 | cat3 | from_5045 | to_5045 | art_long | art_long | M |
| 36 | 143 | 0.676916973 | cat3 | from_5046 | to_5046 | art_long | art_long | M |
| 38 | 143 | 0.832950589 | cat3 | from_5047 | to_5047 | art_long | art_long | M |
| 48 | 143 | 0.873241471 | cat3 | from_5048 | to_5048 | art_long | art_long | M |
| 50 | 143 | 0.767225307 | cat3 | from_5049 | to_5049 | art_long | art_long | M |
| 90 | 143 | -0.606865034 | cat3 | from_5050 | to_5050 | art_long | art_long | M |
| 104 | 143 | 0.911046297 | cat3 | from_5051 | to_5051 | art_long | art_long | M |
| 117 | 143 | -0.606594921 | cat3 | from_5052 | to_5052 | art_long | art_long | M |
| 132 | 143 | 0.860546687 | cat3 | from_5053 | to_5053 | art_long | art_long | M |
| 134 | 143 | 0.608108276 | cat3 | from_5054 | to_5054 | art_long | art_long | M |
| 138 | 143 | 0.734745361 | cat3 | from_5055 | to_5055 | art_long | art_long | M |
| 20 | 144 | -0.602896463 | cat3 | from_5056 | to_5056 | art_long | art_long | M |
| 26 | 144 | 0.633211308 | cat3 | from_5057 | to_5057 | art_long | art_long | M |
| 58 | 144 | 0.639677282 | cat3 | from_5058 | to_5058 | art_long | art_long | M |
| 69 | 144 | 0.612743266 | cat3 | from_5059 | to_5059 | art_long | art_long | M |
| 95 | 144 | 0.605781188 | cat3 | from_5060 | to_5060 | art_long | art_long | M |
| 109 | 144 | 0.60330703 | cat3 | from_5061 | to_5061 | art_long | art_long | M |
| 118 | 144 | 0.968411666 | cat3 | from_5062 | to_5062 | art_long | art_long | M |
| 119 | 144 | 0.684408837 | cat3 | from_5063 | to_5063 | art_long | art_long | M |
| 130 | 144 | 0.702915352 | cat3 | from_5064 | to_5064 | art_long | art_long | M |
| 131 | 144 | 0.92221603 | cat3 | from_5065 | to_5065 | art_long | art_long | M |
| 136 | 144 | 0.761425442 | cat3 | from_5066 | to_5066 | art_long | art_long | M |
| 137 | 144 | 0.890471073 | cat3 | from_5067 | to_5067 | art_long | art_long | M |
| 141 | 144 | 0.885183798 | cat3 | from_5068 | to_5068 | art_long | art_long | M |
| 142 | 144 | 0.626870014 | cat3 | from_5069 | to_5069 | art_long | art_long | M |
| 8 | 145 | 0.830436462 | cat3 | from_5070 | to_5070 | art_long | art_long | M |
| 12 | 145 | 0.617151757 | cat3 | from_5071 | to_5071 | art_long | art_long | M |
| 19 | 145 | 0.612322398 | cat3 | from_5072 | to_5072 | art_long | art_long | M |
| 54 | 145 | 0.639917872 | cat3 | from_5073 | to_5073 | art_long | art_long | M |
| 64 | 145 | -0.625178307 | cat3 | from_5074 | to_5074 | art_long | art_long | M |
| 79 | 145 | 0.735025389 | cat3 | from_5075 | to_5075 | art_long | art_long | M |
| 81 | 145 | 0.739946315 | cat3 | from_5076 | to_5076 | art_long | art_long | M |
| 93 | 145 | 0.853229023 | cat3 | from_5077 | to_5077 | art_long | art_long | M |
| 100 | 145 | -0.687994129 | cat3 | from_5078 | to_5078 | art_long | art_long | M |
| 103 | 145 | -0.714077888 | cat3 | from_5079 | to_5079 | art_long | art_long | M |
| 120 | 145 | 0.602613112 | cat3 | from_5080 | to_5080 | art_long | art_long | M |
| 134 | 145 | -0.614745123 | cat3 | from_5081 | to_5081 | art_long | art_long | M |
| 49 | 146 | 0.715503427 | cat3 | from_5082 | to_5082 | art_long | art_long | M |
| 56 | 146 | 0.756213559 | cat3 | from_5083 | to_5083 | art_long | art_long | M |
| 57 | 146 | 0.719134516 | cat3 | from_5084 | to_5084 | art_long | art_long | M |
| 70 | 146 | 0.894514294 | cat3 | from_5085 | to_5085 | art_long | art_long | M |
| 75 | 146 | 0.952762417 | cat3 | from_5086 | to_5086 | art_long | art_long | M |
| 84 | 146 | 0.708255254 | cat3 | from_5087 | to_5087 | art_long | art_long | M |
| 85 | 146 | 0.784752213 | cat3 | from_5088 | to_5088 | art_long | art_long | M |
| 108 | 146 | 0.619994161 | cat3 | from_5089 | to_5089 | art_long | art_long | M |
| 123 | 146 | 0.939159706 | cat3 | from_5090 | to_5090 | art_long | art_long | M |
| 4 | 147 | 0.64390971 | cat3 | from_5091 | to_5091 | art_long | art_long | M |
| 21 | 147 | 0.700502615 | cat3 | from_5092 | to_5092 | art_long | art_long | M |
| 42 | 147 | 0.953983624 | cat3 | from_5093 | to_5093 | art_long | art_long | M |
| 43 | 147 | 0.714958405 | cat3 | from_5094 | to_5094 | art_long | art_long | M |

| | | | | | | | | |
|---|---|---|---|---|---|---|---|---|
| 44 | 147 | 0.60447872 | cat3 | from_5095 | to_5095 | art_long | art_long | M |
| 45 | 147 | 0.871902488 | cat3 | from_5096 | to_5096 | art_long | art_long | M |
| 60 | 147 | 0.63926291 | cat3 | from_5097 | to_5097 | art_long | art_long | M |
| 62 | 147 | 0.890956558 | cat3 | from_5098 | to_5098 | art_long | art_long | M |
| 92 | 147 | 0.961742022 | cat3 | from_5099 | to_5099 | art_long | art_long | M |
| 100 | 147 | 0.746796198 | cat3 | from_5100 | to_5100 | art_long | art_long | M |
| 101 | 147 | 0.950066694 | cat3 | from_5101 | to_5101 | art_long | art_long | M |
| 107 | 147 | 0.763486273 | cat3 | from_5102 | to_5102 | art_long | art_long | M |
| 133 | 147 | 0.76936275 | cat3 | from_5103 | to_5103 | art_long | art_long | M |
| 43 | 148 | 0.656216483 | cat3 | from_5104 | to_5104 | art_long | art_long | M |
| 44 | 148 | 0.833696174 | cat3 | from_5105 | to_5105 | art_long | art_long | M |
| 46 | 148 | 0.782850762 | cat3 | from_5106 | to_5106 | art_long | art_long | M |
| 50 | 148 | 0.690530005 | cat3 | from_5107 | to_5107 | art_long | art_long | M |
| 52 | 148 | 0.681873046 | cat3 | from_5108 | to_5108 | art_long | art_long | M |
| 60 | 148 | 0.938759223 | cat3 | from_5109 | to_5109 | art_long | art_long | M |
| 61 | 148 | 0.887555075 | cat3 | from_5110 | to_5110 | art_long | art_long | M |
| 99 | 148 | 0.961267383 | cat3 | from_5111 | to_5111 | art_long | art_long | M |
| 105 | 148 | 0.910440616 | cat3 | from_5112 | to_5112 | art_long | art_long | M |
| 106 | 148 | 0.800496261 | cat3 | from_5113 | to_5113 | art_long | art_long | M |
| 128 | 148 | 0.905747789 | cat3 | from_5114 | to_5114 | art_long | art_long | M |
| 4 | 149 | 0.627060457 | cat3 | from_5115 | to_5115 | art_long | art_long | M |
| 7 | 149 | 0.605515435 | cat3 | from_5116 | to_5116 | art_long | art_long | M |
| 17 | 149 | 0.663780569 | cat3 | from_5117 | to_5117 | art_long | art_long | M |
| 36 | 149 | 0.7105276 | cat3 | from_5118 | to_5118 | art_long | art_long | M |
| 42 | 149 | 0.717013471 | cat3 | from_5119 | to_5119 | art_long | art_long | M |
| 43 | 149 | 0.936428636 | cat3 | from_5120 | to_5120 | art_long | art_long | M |
| 45 | 149 | 0.78271149 | cat3 | from_5121 | to_5121 | art_long | art_long | M |
| 60 | 149 | 0.630266627 | cat3 | from_5122 | to_5122 | art_long | art_long | M |
| 62 | 149 | 0.855585674 | cat3 | from_5123 | to_5123 | art_long | art_long | M |
| 92 | 149 | 0.647838006 | cat3 | from_5124 | to_5124 | art_long | art_long | M |
| 100 | 149 | 0.633910413 | cat3 | from_5125 | to_5125 | art_long | art_long | M |
| 101 | 149 | 0.846969805 | cat3 | from_5126 | to_5126 | art_long | art_long | M |
| 107 | 149 | 0.998813122 | cat3 | from_5127 | to_5127 | art_long | art_long | M |
| 128 | 149 | 0.645557252 | cat3 | from_5128 | to_5128 | art_long | art_long | M |
| 138 | 149 | 0.641287409 | cat3 | from_5129 | to_5129 | art_long | art_long | M |
| 139 | 149 | 0.788696731 | cat3 | from_5130 | to_5130 | art_long | art_long | M |
| 147 | 149 | 0.742999401 | cat3 | from_5131 | to_5131 | art_long | art_long | M |
| 3 | 150 | 0.646846967 | cat3 | from_5132 | to_5132 | art_long | art_long | M |
| 29 | 150 | 0.841400704 | cat3 | from_5133 | to_5133 | art_long | art_long | M |
| 41 | 150 | 0.699810485 | cat3 | from_5134 | to_5134 | art_long | art_long | M |
| 66 | 150 | 0.688363335 | cat3 | from_5135 | to_5135 | art_long | art_long | M |
| 67 | 150 | 0.747531032 | cat3 | from_5136 | to_5136 | art_long | art_long | M |
| 71 | 150 | 0.872571666 | cat3 | from_5137 | to_5137 | art_long | art_long | M |
| 82 | 150 | 0.946600043 | cat3 | from_5138 | to_5138 | art_long | art_long | M |
| 89 | 150 | 0.993118887 | cat3 | from_5139 | to_5139 | art_long | art_long | M |
| 98 | 150 | 0.809392122 | cat3 | from_5140 | to_5140 | art_long | art_long | M |
| 121 | 150 | 0.815802506 | cat3 | from_5141 | to_5141 | art_long | art_long | M |
| 124 | 150 | 0.711847652 | cat3 | from_5142 | to_5142 | art_long | art_long | M |
| 1 | 151 | -0.708074125 | cat3 | from_5143 | to_5143 | art_long | art_long | M |
| 8 | 152 | 0.702896971 | cat3 | from_5144 | to_5144 | art_long | art_long | M |
| 50 | 152 | 0.648012643 | cat3 | from_5145 | to_5145 | art_long | art_long | M |
| 61 | 152 | 0.621724592 | cat3 | from_5146 | to_5146 | art_long | art_long | M |
| 93 | 152 | 0.663775884 | cat3 | from_5147 | to_5147 | art_long | art_long | M |
| 99 | 152 | 0.609328394 | cat3 | from_5148 | to_5148 | art_long | art_long | M |
| 105 | 152 | 0.753429452 | cat3 | from_5149 | to_5149 | art_long | art_long | M |
| 145 | 152 | 0.694339983 | cat3 | from_5150 | to_5150 | art_long | art_long | M |
| 148 | 152 | 0.694051883 | cat3 | from_5151 | to_5151 | art_long | art_long | M |
| 151 | 152 | 0.699886504 | cat3 | from_5152 | to_5152 | art_long | art_long | M |
| 8 | 153 | 0.612876235 | cat3 | from_5153 | to_5153 | art_long | art_long | M |
| 50 | 153 | 0.644669092 | cat3 | from_5154 | to_5154 | art_long | art_long | M |
| 61 | 153 | 0.658604326 | cat3 | from_5155 | to_5155 | art_long | art_long | M |
| 93 | 153 | 0.622524339 | cat3 | from_5156 | to_5156 | art_long | art_long | M |
| 99 | 153 | 0.643267807 | cat3 | from_5157 | to_5157 | art_long | art_long | M |
| 105 | 153 | 0.744689679 | cat3 | from_5158 | to_5158 | art_long | art_long | M |
| 148 | 153 | 0.713943878 | cat3 | from_5159 | to_5159 | art_long | art_long | M |
| 151 | 153 | 0.734796124 | cat3 | from_5160 | to_5160 | art_long | art_long | M |
| 152 | 153 | 0.98392353 | cat3 | from_5161 | to_5161 | art_long | art_long | M |
| 17 | 154 | 0.794596709 | cat3 | from_5162 | to_5162 | art_long | art_short | J |
| 36 | 154 | 0.850576017 | cat3 | from_5163 | to_5163 | art_long | art_short | J |
| 72 | 154 | -0.615105061 | cat3 | from_5164 | to_5164 | art_long | art_short | J |
| 91 | 154 | -0.628650417 | cat3 | from_5165 | to_5165 | art_long | art_short | J |
| 104 | 154 | 0.629648224 | cat3 | from_5166 | to_5166 | art_long | art_short | J |
| 108 | 154 | 0.61733862 | cat3 | from_5167 | to_5167 | art_long | art_short | J |
| 111 | 154 | -0.613846851 | cat3 | from_5168 | to_5168 | art_long | art_short | J |
| 134 | 154 | 0.611866943 | cat3 | from_5169 | to_5169 | art_long | art_short | J |
| 138 | 154 | 0.880713876 | cat3 | from_5170 | to_5170 | art_long | art_short | J |
| 143 | 154 | 0.767178724 | cat3 | from_5171 | to_5171 | art_long | art_short | J |
| 2 | 155 | 0.971154277 | cat3 | from_5172 | to_5172 | art_long | art_short | J |
| 6 | 155 | -0.633966514 | cat3 | from_5173 | to_5173 | art_long | art_short | J |
| 53 | 155 | 0.713722857 | cat3 | from_5174 | to_5174 | art_long | art_short | J |
| 112 | 155 | -0.611331039 | cat3 | from_5175 | to_5175 | art_long | art_short | J |
| 8 | 156 | 0.69248078 | cat3 | from_5176 | to_5176 | art_long | art_short | J |
| 10 | 156 | 0.611622925 | cat3 | from_5177 | to_5177 | art_long | art_short | J |
| 13 | 156 | 0.646216285 | cat3 | from_5178 | to_5178 | art_long | art_short | J |
| 19 | 156 | 0.617512539 | cat3 | from_5179 | to_5179 | art_long | art_short | J |
| 40 | 156 | 0.6239042 | cat3 | from_5180 | to_5180 | art_long | art_short | J |
| 79 | 156 | 0.623803202 | cat3 | from_5181 | to_5181 | art_long | art_short | J |
| 114 | 156 | 0.800122034 | cat3 | from_5182 | to_5182 | art_long | art_short | J |
| 125 | 156 | 0.712814721 | cat3 | from_5183 | to_5183 | art_long | art_short | J |
| 6 | 157 | 0.838904149 | cat3 | from_5184 | to_5184 | art_long | art_short | J |
| 8 | 157 | 0.831800577 | cat3 | from_5185 | to_5185 | art_long | art_short | J |
| 9 | 157 | 0.644482765 | cat3 | from_5186 | to_5186 | art_long | art_short | J |
| 10 | 157 | 0.637450823 | cat3 | from_5187 | to_5187 | art_long | art_short | J |
| 84 | 157 | 0.622387528 | cat3 | from_5188 | to_5188 | art_long | art_short | J |
| 125 | 157 | 0.703756945 | cat3 | from_5189 | to_5189 | art_long | art_short | J |
| 145 | 157 | 0.660263145 | cat3 | from_5190 | to_5190 | art_long | art_short | J |
| 152 | 157 | 0.612861885 | cat3 | from_5191 | to_5191 | art_long | art_short | J |
| 156 | 157 | 0.670122731 | cat3 | from_5192 | to_5192 | art_short | art_short | H |
| 8 | 158 | 0.695485449 | cat3 | from_5193 | to_5193 | art_long | art_short | J |
| 13 | 158 | 0.635124084 | cat3 | from_5194 | to_5194 | art_long | art_short | J |
| 48 | 158 | -0.61848685 | cat3 | from_5195 | to_5195 | art_long | art_short | J |
| 63 | 158 | 0.620360406 | cat3 | from_5196 | to_5196 | art_long | art_short | J |
| 79 | 158 | 0.609261407 | cat3 | from_5197 | to_5197 | art_long | art_short | J |
| 104 | 158 | -0.606331585 | cat3 | from_5198 | to_5198 | art_long | art_short | J |

| | | | | | | | | |
|---|---|---|---|---|---|---|---|---|
| 114 | 158 | 0.801836619 | cat3 | from_5199 | to_5199 | art_long | art_short | J |
| 125 | 158 | 0.741049706 | cat3 | from_5200 | to_5200 | art_long | art_short | J |
| 131 | 158 | -0.608401343 | cat3 | from_5201 | to_5201 | art_long | art_short | J |
| 134 | 158 | -0.60351616 | cat3 | from_5202 | to_5202 | art_long | art_short | J |
| 156 | 158 | 0.989626235 | cat3 | from_5203 | to_5203 | art_short | art_short | H |
| 157 | 158 | 0.671688315 | cat3 | from_5204 | to_5204 | art_short | art_short | H |
| 4 | 159 | 0.820719684 | cat3 | from_5205 | to_5205 | art_long | art_short | J |
| 6 | 159 | 0.764473435 | cat3 | from_5206 | to_5206 | art_long | art_short | J |
| 7 | 159 | 0.82112863 | cat3 | from_5207 | to_5207 | art_long | art_short | J |
| 11 | 159 | 0.712634066 | cat3 | from_5208 | to_5208 | art_long | art_short | J |
| 65 | 159 | 0.664171213 | cat3 | from_5209 | to_5209 | art_long | art_short | J |
| 113 | 159 | 0.611564969 | cat3 | from_5210 | to_5210 | art_long | art_short | J |
| 118 | 159 | -0.641090718 | cat3 | from_5211 | to_5211 | art_long | art_short | J |
| 137 | 159 | -0.650826675 | cat3 | from_5212 | to_5212 | art_long | art_short | J |
| 141 | 159 | -0.650303414 | cat3 | from_5213 | to_5213 | art_long | art_short | J |
| 144 | 159 | -0.66628993 | cat3 | from_5214 | to_5214 | art_long | art_short | J |
| 4 | 160 | 0.952199199 | cat3 | from_5215 | to_5215 | art_long | art_short | J |
| 6 | 160 | 0.71753791 | cat3 | from_5216 | to_5216 | art_long | art_short | J |
| 7 | 160 | 0.834865281 | cat3 | from_5217 | to_5217 | art_long | art_short | J |
| 11 | 160 | 0.736956303 | cat3 | from_5218 | to_5218 | art_long | art_short | J |
| 34 | 160 | 0.691685754 | cat3 | from_5219 | to_5219 | art_long | art_short | J |
| 127 | 160 | 0.676953767 | cat3 | from_5220 | to_5220 | art_long | art_short | J |
| 147 | 160 | 0.618103324 | cat3 | from_5221 | to_5221 | art_long | art_short | J |
| 159 | 160 | 0.853814977 | cat3 | from_5222 | to_5222 | art_short | art_short | H |
| 8 | 161 | 0.675993276 | cat3 | from_5223 | to_5223 | art_long | art_short | J |
| 9 | 161 | 0.828083022 | cat3 | from_5224 | to_5224 | art_long | art_short | J |
| 10 | 161 | 0.8062816 | cat3 | from_5225 | to_5225 | art_long | art_short | J |
| 22 | 161 | 0.749183336 | cat3 | from_5226 | to_5226 | art_long | art_short | J |
| 23 | 161 | 0.825442255 | cat3 | from_5227 | to_5227 | art_long | art_short | J |
| 57 | 161 | 0.692399384 | cat3 | from_5228 | to_5228 | art_long | art_short | J |
| 88 | 161 | 0.861165226 | cat3 | from_5229 | to_5229 | art_long | art_short | J |
| 126 | 161 | 0.913115051 | cat3 | from_5230 | to_5230 | art_long | art_short | J |
| 2 | 162 | 0.658353205 | cat3 | from_5231 | to_5231 | art_long | art_short | J |
| 16 | 162 | 0.603031164 | cat3 | from_5232 | to_5232 | art_long | art_short | J |
| 155 | 162 | 0.667251721 | cat3 | from_5233 | to_5233 | art_short | art_short | H |
| 13 | 163 | 0.75520846 | cat3 | from_5234 | to_5234 | art_long | art_short | J |
| 15 | 163 | 0.635131308 | cat3 | from_5235 | to_5235 | art_long | art_short | J |
| 16 | 163 | 0.603335346 | cat3 | from_5236 | to_5236 | art_long | art_short | J |
| 40 | 163 | 0.701207304 | cat3 | from_5237 | to_5237 | art_long | art_short | J |
| 114 | 163 | 0.71936841 | cat3 | from_5238 | to_5238 | art_long | art_short | J |
| 162 | 163 | 0.928637674 | cat3 | from_5239 | to_5239 | art_short | art_short | H |
| 12 | 164 | 0.675294678 | cat3 | from_5240 | to_5240 | art_long | art_short | J |
| 16 | 164 | 0.959191928 | cat3 | from_5241 | to_5241 | art_long | art_short | J |
| 18 | 164 | 0.690148688 | cat3 | from_5242 | to_5242 | art_long | art_short | J |
| 19 | 164 | 0.628064261 | cat3 | from_5243 | to_5243 | art_long | art_short | J |
| 39 | 164 | 0.903624617 | cat3 | from_5244 | to_5244 | art_long | art_short | J |
| 40 | 164 | 0.811169363 | cat3 | from_5245 | to_5245 | art_long | art_short | J |
| 53 | 164 | 0.72694898 | cat3 | from_5246 | to_5246 | art_long | art_short | J |
| 155 | 164 | 0.618991066 | cat3 | from_5247 | to_5247 | art_short | art_short | H |
| 162 | 164 | 0.619285289 | cat3 | from_5248 | to_5248 | art_short | art_short | H |
| 163 | 164 | 0.657296093 | cat3 | from_5249 | to_5249 | art_short | art_short | H |
| 12 | 165 | 0.756547202 | cat3 | from_5250 | to_5250 | art_long | art_short | J |
| 13 | 165 | 0.89665634 | cat3 | from_5251 | to_5251 | art_long | art_short | J |
| 15 | 165 | 0.692948414 | cat3 | from_5252 | to_5252 | art_long | art_short | J |
| 18 | 165 | 0.612519606 | cat3 | from_5253 | to_5253 | art_long | art_short | J |
| 19 | 165 | 0.737220469 | cat3 | from_5254 | to_5254 | art_long | art_short | J |
| 20 | 165 | 0.702839709 | cat3 | from_5255 | to_5255 | art_long | art_short | J |
| 39 | 165 | 0.68073338 | cat3 | from_5256 | to_5256 | art_long | art_short | J |
| 40 | 165 | 0.69131516 | cat3 | from_5257 | to_5257 | art_long | art_short | J |
| 68 | 165 | 0.752238086 | cat3 | from_5258 | to_5258 | art_long | art_short | J |
| 79 | 165 | 0.752174594 | cat3 | from_5259 | to_5259 | art_long | art_short | J |
| 114 | 165 | 0.961738113 | cat3 | from_5260 | to_5260 | art_long | art_short | J |
| 156 | 165 | 0.761508963 | cat3 | from_5261 | to_5261 | art_short | art_short | H |
| 158 | 165 | 0.781900417 | cat3 | from_5262 | to_5262 | art_short | art_short | H |
| 163 | 165 | 0.707101927 | cat3 | from_5263 | to_5263 | art_short | art_short | H |
| 25 | 166 | -0.607308445 | cat3 | from_5264 | to_5264 | art_long | art_short | J |
| 31 | 166 | -0.640818063 | cat3 | from_5265 | to_5265 | art_long | art_short | J |
| 33 | 166 | -0.630058368 | cat3 | from_5266 | to_5266 | art_long | art_short | J |
| 35 | 166 | 0.85848656 | cat3 | from_5267 | to_5267 | art_long | art_short | J |
| 38 | 166 | 0.80725212 | cat3 | from_5268 | to_5268 | art_long | art_short | J |
| 48 | 166 | 0.959387917 | cat3 | from_5269 | to_5269 | art_long | art_short | J |
| 51 | 166 | -0.633280132 | cat3 | from_5270 | to_5270 | art_long | art_short | J |
| 63 | 166 | -0.610485 | cat3 | from_5271 | to_5271 | art_long | art_short | J |
| 68 | 166 | -0.687474291 | cat3 | from_5272 | to_5272 | art_long | art_short | J |
| 102 | 166 | 0.658808058 | cat3 | from_5273 | to_5273 | art_long | art_short | J |
| 104 | 166 | 0.923196396 | cat3 | from_5274 | to_5274 | art_long | art_short | J |
| 110 | 166 | -0.615739454 | cat3 | from_5275 | to_5275 | art_long | art_short | J |
| 112 | 166 | -0.665667041 | cat3 | from_5276 | to_5276 | art_long | art_short | J |
| 132 | 166 | 0.811183279 | cat3 | from_5277 | to_5277 | art_long | art_short | J |
| 134 | 166 | 0.766247272 | cat3 | from_5278 | to_5278 | art_long | art_short | J |
| 138 | 166 | 0.656506387 | cat3 | from_5279 | to_5279 | art_long | art_short | J |
| 143 | 166 | 0.894346195 | cat3 | from_5280 | to_5280 | art_long | art_short | J |
| 12 | 167 | 0.995857313 | cat3 | from_5281 | to_5281 | art_long | art_short | J |
| 13 | 167 | 0.775148565 | cat3 | from_5282 | to_5282 | art_long | art_short | J |
| 15 | 167 | 0.639573789 | cat3 | from_5283 | to_5283 | art_long | art_short | J |
| 16 | 167 | 0.638850699 | cat3 | from_5284 | to_5284 | art_long | art_short | J |
| 18 | 167 | 0.787485241 | cat3 | from_5285 | to_5285 | art_long | art_short | J |
| 19 | 167 | 0.955181273 | cat3 | from_5286 | to_5286 | art_long | art_short | J |
| 39 | 167 | 0.726187675 | cat3 | from_5287 | to_5287 | art_long | art_short | J |
| 40 | 167 | 0.838863674 | cat3 | from_5288 | to_5288 | art_long | art_short | J |
| 79 | 167 | 0.961535017 | cat3 | from_5289 | to_5289 | art_long | art_short | J |
| 114 | 167 | 0.725181625 | cat3 | from_5290 | to_5290 | art_long | art_short | J |
| 163 | 167 | 0.626215964 | cat3 | from_5291 | to_5291 | art_short | art_short | H |
| 164 | 167 | 0.679554875 | cat3 | from_5292 | to_5292 | art_short | art_short | H |
| 165 | 167 | 0.732368682 | cat3 | from_5293 | to_5293 | art_short | art_short | H |
| 39 | 168 | 0.621244445 | cat3 | from_5294 | to_5294 | art_long | art_short | J |
| 53 | 168 | 0.781068242 | cat3 | from_5295 | to_5295 | art_long | art_short | J |
| 165 | 168 | 0.626035942 | cat3 | from_5296 | to_5296 | art_short | art_short | H |
| 13 | 169 | 0.728111019 | cat3 | from_5297 | to_5297 | art_long | art_short | J |
| 15 | 169 | 0.66700401 | cat3 | from_5298 | to_5298 | art_long | art_short | J |
| 16 | 169 | 0.697877878 | cat3 | from_5299 | to_5299 | art_long | art_short | J |
| 20 | 169 | 0.720633454 | cat3 | from_5300 | to_5300 | art_long | art_short | J |
| 21 | 169 | 0.706412044 | cat3 | from_5301 | to_5301 | art_long | art_short | J |
| 39 | 169 | 0.756999159 | cat3 | from_5302 | to_5302 | art_long | art_short | J |

| | | | | | | | | |
|---|---|---|---|---|---|---|---|---|
| 40 | 169 | 0.654001157 | cat3 | from_5303 | to_5303 | art_long | art_short | J |
| 114 | 169 | 0.782780957 | cat3 | from_5304 | to_5304 | art_long | art_short | J |
| 116 | 169 | 0.634823939 | cat3 | from_5305 | to_5305 | art_long | art_short | J |
| 158 | 169 | 0.603348868 | cat3 | from_5306 | to_5306 | art_short | art_short | H |
| 162 | 169 | 0.633522144 | cat3 | from_5307 | to_5307 | art_short | art_short | H |
| 163 | 169 | 0.783539154 | cat3 | from_5308 | to_5308 | art_short | art_short | H |
| 164 | 169 | 0.756971656 | cat3 | from_5309 | to_5309 | art_short | art_short | H |
| 165 | 169 | 0.855338662 | cat3 | from_5310 | to_5310 | art_short | art_short | H |
| 168 | 169 | 0.739327517 | cat3 | from_5311 | to_5311 | art_short | art_short | H |
| 42 | 170 | 0.669433492 | cat3 | from_5312 | to_5312 | art_long | art_short | J |
| 45 | 170 | 0.812609335 | cat3 | from_5313 | to_5313 | art_long | art_short | J |
| 62 | 170 | 0.795907399 | cat3 | from_5314 | to_5314 | art_long | art_short | J |
| 91 | 170 | -0.617976354 | cat3 | from_5315 | to_5315 | art_long | art_short | J |
| 100 | 170 | 0.862667261 | cat3 | from_5316 | to_5316 | art_long | art_short | J |
| 101 | 170 | 0.703164251 | cat3 | from_5317 | to_5317 | art_long | art_short | J |
| 102 | 170 | 0.625094794 | cat3 | from_5318 | to_5318 | art_long | art_short | J |
| 107 | 170 | 0.663092833 | cat3 | from_5319 | to_5319 | art_long | art_short | J |
| 134 | 170 | 0.630194682 | cat3 | from_5320 | to_5320 | art_long | art_short | J |
| 139 | 170 | 0.968605083 | cat3 | from_5321 | to_5321 | art_long | art_short | J |
| 141 | 170 | -0.623299187 | cat3 | from_5322 | to_5322 | art_long | art_short | J |
| 149 | 170 | 0.638055182 | cat3 | from_5323 | to_5323 | art_long | art_short | J |
| 42 | 171 | 0.725319375 | cat3 | from_5324 | to_5324 | art_long | art_short | J |
| 45 | 171 | 0.85138287 | cat3 | from_5325 | to_5325 | art_long | art_short | J |
| 62 | 171 | 0.836757616 | cat3 | from_5326 | to_5326 | art_long | art_short | J |
| 91 | 171 | -0.606731752 | cat3 | from_5327 | to_5327 | art_long | art_short | J |
| 92 | 171 | 0.659951937 | cat3 | from_5328 | to_5328 | art_long | art_short | J |
| 100 | 171 | 0.887798384 | cat3 | from_5329 | to_5329 | art_long | art_short | J |
| 101 | 171 | 0.75493833 | cat3 | from_5330 | to_5330 | art_long | art_short | J |
| 102 | 171 | 0.635553243 | cat3 | from_5331 | to_5331 | art_long | art_short | J |
| 107 | 171 | 0.694851353 | cat3 | from_5332 | to_5332 | art_long | art_short | J |
| 134 | 171 | 0.632270748 | cat3 | from_5333 | to_5333 | art_long | art_short | J |
| 139 | 171 | 0.968497076 | cat3 | from_5334 | to_5334 | art_long | art_short | J |
| 141 | 171 | -0.629953065 | cat3 | from_5335 | to_5335 | art_long | art_short | J |
| 149 | 171 | 0.668916564 | cat3 | from_5336 | to_5336 | art_long | art_short | J |
| 170 | 171 | 0.996356107 | cat3 | from_5337 | to_5337 | art_short | art_short | H |
| 22 | 172 | 0.667512688 | cat3 | from_5338 | to_5338 | art_long | art_short | J |
| 27 | 172 | 0.894721559 | cat3 | from_5339 | to_5339 | art_long | art_short | J |
| 55 | 172 | 0.624211111 | cat3 | from_5340 | to_5340 | art_long | art_short | J |
| 64 | 172 | 0.729569514 | cat3 | from_5341 | to_5341 | art_long | art_short | J |
| 87 | 172 | 0.614887014 | cat3 | from_5342 | to_5342 | art_long | art_short | J |
| 91 | 172 | 0.851242633 | cat3 | from_5343 | to_5343 | art_long | art_short | J |
| 109 | 172 | 0.627467089 | cat3 | from_5344 | to_5344 | art_long | art_short | J |
| 117 | 172 | 0.682572217 | cat3 | from_5345 | to_5345 | art_long | art_short | J |
| 24 | 173 | 0.845692253 | cat3 | from_5346 | to_5346 | art_long | art_short | J |
| 25 | 173 | 0.602852866 | cat3 | from_5347 | to_5347 | art_long | art_short | J |
| 28 | 173 | 0.919754095 | cat3 | from_5348 | to_5348 | art_long | art_short | J |
| 47 | 173 | 0.66329658 | cat3 | from_5349 | to_5349 | art_long | art_short | J |
| 50 | 173 | -0.642008361 | cat3 | from_5350 | to_5350 | art_long | art_short | J |
| 55 | 173 | 0.671061963 | cat3 | from_5351 | to_5351 | art_long | art_short | J |
| 59 | 173 | 0.743131901 | cat3 | from_5352 | to_5352 | art_long | art_short | J |
| 64 | 173 | 0.650700861 | cat3 | from_5353 | to_5353 | art_long | art_short | J |
| 73 | 173 | 0.609798014 | cat3 | from_5354 | to_5354 | art_long | art_short | J |
| 76 | 173 | 0.84707048 | cat3 | from_5355 | to_5355 | art_long | art_short | J |
| 78 | 173 | 0.64521934 | cat3 | from_5356 | to_5356 | art_long | art_short | J |
| 87 | 173 | 0.641801728 | cat3 | from_5357 | to_5357 | art_long | art_short | J |
| 90 | 173 | 0.822426573 | cat3 | from_5358 | to_5358 | art_long | art_short | J |
| 95 | 173 | 0.615437144 | cat3 | from_5359 | to_5359 | art_long | art_short | J |
| 97 | 173 | 0.838464302 | cat3 | from_5360 | to_5360 | art_long | art_short | J |
| 109 | 173 | 0.649646547 | cat3 | from_5361 | to_5361 | art_long | art_short | J |
| 122 | 173 | 0.87181002 | cat3 | from_5362 | to_5362 | art_long | art_short | J |
| 14 | 174 | 0.854751567 | cat3 | from_5363 | to_5363 | art_long | art_short | J |
| 24 | 174 | 0.821877625 | cat3 | from_5364 | to_5364 | art_long | art_short | J |
| 25 | 174 | 0.776827456 | cat3 | from_5365 | to_5365 | art_long | art_short | J |
| 28 | 174 | 0.896552583 | cat3 | from_5366 | to_5366 | art_long | art_short | J |
| 31 | 174 | 0.683059011 | cat3 | from_5367 | to_5367 | art_long | art_short | J |
| 33 | 174 | 0.710106503 | cat3 | from_5368 | to_5368 | art_long | art_short | J |
| 59 | 174 | 0.914745532 | cat3 | from_5369 | to_5369 | art_long | art_short | J |
| 73 | 174 | 0.863674898 | cat3 | from_5370 | to_5370 | art_long | art_short | J |
| 74 | 174 | 0.841101578 | cat3 | from_5371 | to_5371 | art_long | art_short | J |
| 76 | 174 | 0.865576613 | cat3 | from_5372 | to_5372 | art_long | art_short | J |
| 78 | 174 | 0.823773564 | cat3 | from_5373 | to_5373 | art_long | art_short | J |
| 90 | 174 | 0.70413156 | cat3 | from_5374 | to_5374 | art_long | art_short | J |
| 97 | 174 | 0.761992587 | cat3 | from_5375 | to_5375 | art_long | art_short | J |
| 110 | 174 | 0.875541391 | cat3 | from_5376 | to_5376 | art_long | art_short | J |
| 112 | 174 | 0.730863939 | cat3 | from_5377 | to_5377 | art_long | art_short | J |
| 119 | 174 | 0.605454936 | cat3 | from_5378 | to_5378 | art_long | art_short | J |
| 122 | 174 | 0.858147886 | cat3 | from_5379 | to_5379 | art_long | art_short | J |
| 173 | 174 | 0.87296228 | cat3 | from_5380 | to_5380 | art_short | art_short | H |
| 82 | 175 | 0.81909991 | cat3 | from_5381 | to_5381 | art_long | art_short | J |
| 89 | 175 | 0.75984126 | cat3 | from_5382 | to_5382 | art_long | art_short | J |
| 121 | 175 | 0.610747367 | cat3 | from_5383 | to_5383 | art_long | art_short | J |
| 150 | 175 | 0.767809349 | cat3 | from_5384 | to_5384 | art_long | art_short | J |
| 26 | 176 | -0.626324408 | cat3 | from_5385 | to_5385 | art_long | art_short | J |
| 46 | 176 | 0.608541553 | cat3 | from_5386 | to_5386 | art_long | art_short | J |
| 61 | 176 | 0.822968598 | cat3 | from_5387 | to_5387 | art_long | art_short | J |
| 102 | 176 | 0.654648984 | cat3 | from_5388 | to_5388 | art_long | art_short | J |
| 106 | 176 | 0.697560551 | cat3 | from_5389 | to_5389 | art_long | art_short | J |
| 115 | 176 | -0.654342104 | cat3 | from_5390 | to_5390 | art_long | art_short | J |
| 117 | 176 | -0.626553211 | cat3 | from_5391 | to_5391 | art_long | art_short | J |
| 127 | 176 | -0.608093796 | cat3 | from_5392 | to_5392 | art_long | art_short | J |
| 135 | 176 | 0.783038871 | cat3 | from_5393 | to_5393 | art_long | art_short | J |
| 170 | 176 | 0.608533641 | cat3 | from_5394 | to_5394 | art_short | art_short | H |
| 17 | 177 | 0.682597897 | cat3 | from_5395 | to_5395 | art_long | art_short | J |
| 27 | 177 | -0.631111206 | cat3 | from_5396 | to_5396 | art_long | art_short | J |
| 35 | 177 | 0.687987147 | cat3 | from_5397 | to_5397 | art_long | art_short | J |
| 38 | 177 | 0.681191348 | cat3 | from_5398 | to_5398 | art_long | art_short | J |
| 42 | 177 | 0.635059104 | cat3 | from_5399 | to_5399 | art_long | art_short | J |
| 54 | 177 | -0.678170367 | cat3 | from_5400 | to_5400 | art_long | art_short | J |
| 77 | 177 | -0.687100863 | cat3 | from_5401 | to_5401 | art_long | art_short | J |
| 80 | 177 | -0.694270364 | cat3 | from_5402 | to_5402 | art_long | art_short | J |
| 92 | 177 | 0.672395239 | cat3 | from_5403 | to_5403 | art_long | art_short | J |
| 93 | 177 | -0.609748357 | cat3 | from_5404 | to_5404 | art_long | art_short | J |
| 120 | 177 | -0.649193489 | cat3 | from_5405 | to_5405 | art_long | art_short | J |
| 126 | 177 | -0.623844665 | cat3 | from_5406 | to_5406 | art_long | art_short | J |

| | | | | | | | | |
|---|---|---|---|---|---|---|---|---|
| 132 | 177 | 0.627375882 | cat3 | from_5407 | to_5407 | art_long | art_short | J |
| 133 | 177 | 0.90760645 | cat3 | from_5408 | to_5408 | art_long | art_short | J |
| 134 | 177 | 0.647377477 | cat3 | from_5409 | to_5409 | art_long | art_short | J |
| 138 | 177 | 0.645153396 | cat3 | from_5410 | to_5410 | art_long | art_short | J |
| 147 | 177 | 0.649579523 | cat3 | from_5411 | to_5411 | art_long | art_short | J |
| 42 | 178 | 0.808426647 | cat3 | from_5412 | to_5412 | art_long | art_short | J |
| 45 | 178 | 0.733641822 | cat3 | from_5413 | to_5413 | art_long | art_short | J |
| 52 | 178 | 0.701410114 | cat3 | from_5414 | to_5414 | art_long | art_short | J |
| 62 | 178 | 0.757247985 | cat3 | from_5415 | to_5415 | art_long | art_short | J |
| 80 | 178 | -0.63776805 | cat3 | from_5416 | to_5416 | art_long | art_short | J |
| 92 | 178 | 0.821919482 | cat3 | from_5417 | to_5417 | art_long | art_short | J |
| 100 | 178 | 0.622831832 | cat3 | from_5418 | to_5418 | art_long | art_short | J |
| 101 | 178 | 0.805085907 | cat3 | from_5419 | to_5419 | art_long | art_short | J |
| 107 | 178 | 0.646590875 | cat3 | from_5420 | to_5420 | art_long | art_short | J |
| 133 | 178 | 0.637292831 | cat3 | from_5421 | to_5421 | art_long | art_short | J |
| 147 | 178 | 0.854393888 | cat3 | from_5422 | to_5422 | art_long | art_short | J |
| 149 | 178 | 0.629704939 | cat3 | from_5423 | to_5423 | art_long | art_short | J |
| 160 | 178 | 0.623185456 | cat3 | from_5424 | to_5424 | art_short | art_short | H |
| 177 | 178 | 0.710933905 | cat3 | from_5425 | to_5425 | art_short | art_short | H |
| 2 | 179 | 0.929042604 | cat3 | from_5426 | to_5426 | art_long | art_short | J |
| 6 | 179 | -0.624919216 | cat3 | from_5427 | to_5427 | art_long | art_short | J |
| 16 | 179 | 0.677424207 | cat3 | from_5428 | to_5428 | art_long | art_short | J |
| 53 | 179 | 0.860530904 | cat3 | from_5429 | to_5429 | art_long | art_short | J |
| 155 | 179 | 0.955556561 | cat3 | from_5430 | to_5430 | art_short | art_short | H |
| 164 | 179 | 0.715530266 | cat3 | from_5431 | to_5431 | art_short | art_short | H |
| 15 | 180 | 0.61659243 | cat3 | from_5432 | to_5432 | art_long | art_short | J |
| 53 | 180 | 0.820458461 | cat3 | from_5433 | to_5433 | art_long | art_short | J |
| 168 | 180 | 0.954978056 | cat3 | from_5434 | to_5434 | art_short | art_short | H |
| 169 | 180 | 0.670106132 | cat3 | from_5435 | to_5435 | art_short | art_short | H |
| 179 | 180 | 0.667874658 | cat3 | from_5436 | to_5436 | art_short | art_short | H |
| 3 | 181 | 0.691870289 | cat3 | from_5437 | to_5437 | art_long | art_short | J |
| 29 | 181 | 0.89175115 | cat3 | from_5438 | to_5438 | art_long | art_short | J |
| 41 | 181 | 0.642094154 | cat3 | from_5439 | to_5439 | art_long | art_short | J |
| 66 | 181 | 0.64154096 | cat3 | from_5440 | to_5440 | art_long | art_short | J |
| 67 | 181 | 0.701196548 | cat3 | from_5441 | to_5441 | art_long | art_short | J |
| 71 | 181 | 0.911819393 | cat3 | from_5442 | to_5442 | art_long | art_short | J |
| 82 | 181 | 0.815018034 | cat3 | from_5443 | to_5443 | art_long | art_short | J |
| 89 | 181 | 0.825255736 | cat3 | from_5444 | to_5444 | art_long | art_short | J |
| 98 | 181 | 0.609769537 | cat3 | from_5445 | to_5445 | art_long | art_short | J |
| 124 | 181 | 0.658865231 | cat3 | from_5446 | to_5446 | art_long | art_short | J |
| 150 | 181 | 0.821666188 | cat3 | from_5447 | to_5447 | art_long | art_short | J |
| 175 | 181 | 0.742189487 | cat3 | from_5448 | to_5448 | art_short | art_short | H |
| 23 | 182 | 0.6316727 | cat3 | from_5449 | to_5449 | art_long | art_short | J |
| 32 | 182 | -0.648547818 | cat3 | from_5450 | to_5450 | art_long | art_short | J |
| 49 | 182 | 0.627556484 | cat3 | from_5451 | to_5451 | art_long | art_short | J |
| 56 | 182 | 0.905624346 | cat3 | from_5452 | to_5452 | art_long | art_short | J |
| 57 | 182 | 0.773610649 | cat3 | from_5453 | to_5453 | art_long | art_short | J |
| 70 | 182 | 0.729113115 | cat3 | from_5454 | to_5454 | art_long | art_short | J |
| 75 | 182 | 0.71428884 | cat3 | from_5455 | to_5455 | art_long | art_short | J |
| 85 | 182 | 0.686731215 | cat3 | from_5456 | to_5456 | art_long | art_short | J |
| 123 | 182 | 0.770882902 | cat3 | from_5457 | to_5457 | art_long | art_short | J |
| 146 | 182 | 0.761279366 | cat3 | from_5458 | to_5458 | art_long | art_short | J |
| 44 | 183 | 0.702981017 | cat3 | from_5459 | to_5459 | art_long | art_short | J |
| 46 | 183 | 0.660053202 | cat3 | from_5460 | to_5460 | art_long | art_short | J |
| 50 | 183 | 0.655411486 | cat3 | from_5461 | to_5461 | art_long | art_short | J |
| 52 | 183 | 0.686524118 | cat3 | from_5462 | to_5462 | art_long | art_short | J |
| 60 | 183 | 0.813960211 | cat3 | from_5463 | to_5463 | art_long | art_short | J |
| 61 | 183 | 0.862310086 | cat3 | from_5464 | to_5464 | art_long | art_short | J |
| 64 | 183 | -0.635694137 | cat3 | from_5465 | to_5465 | art_long | art_short | J |
| 99 | 183 | 0.8694358 | cat3 | from_5466 | to_5466 | art_long | art_short | J |
| 105 | 183 | 0.873888119 | cat3 | from_5467 | to_5467 | art_long | art_short | J |
| 106 | 183 | 0.696346235 | cat3 | from_5468 | to_5468 | art_long | art_short | J |
| 128 | 183 | 0.773318841 | cat3 | from_5469 | to_5469 | art_long | art_short | J |
| 148 | 183 | 0.90965265 | cat3 | from_5470 | to_5470 | art_long | art_short | J |
| 152 | 183 | 0.621568052 | cat3 | from_5471 | to_5471 | art_long | art_short | J |
| 176 | 183 | 0.698752343 | cat3 | from_5472 | to_5472 | art_short | art_short | H |
| 42 | 184 | 0.787932152 | cat3 | from_5473 | to_5473 | art_long | art_short | J |
| 43 | 184 | 0.90367064 | cat3 | from_5474 | to_5474 | art_long | art_short | J |
| 45 | 184 | 0.821582313 | cat3 | from_5475 | to_5475 | art_long | art_short | J |
| 60 | 184 | 0.626867295 | cat3 | from_5476 | to_5476 | art_long | art_short | J |
| 62 | 184 | 0.878967967 | cat3 | from_5477 | to_5477 | art_long | art_short | J |
| 92 | 184 | 0.741291302 | cat3 | from_5478 | to_5478 | art_long | art_short | J |
| 100 | 184 | 0.691404216 | cat3 | from_5479 | to_5479 | art_long | art_short | J |
| 101 | 184 | 0.877590286 | cat3 | from_5480 | to_5480 | art_long | art_short | J |
| 107 | 184 | 0.928438519 | cat3 | from_5481 | to_5481 | art_long | art_short | J |
| 128 | 184 | 0.614260952 | cat3 | from_5482 | to_5482 | art_long | art_short | J |
| 139 | 184 | 0.840880394 | cat3 | from_5483 | to_5483 | art_long | art_short | J |
| 147 | 184 | 0.803324613 | cat3 | from_5484 | to_5484 | art_long | art_short | J |
| 149 | 184 | 0.921971899 | cat3 | from_5485 | to_5485 | art_long | art_short | J |
| 160 | 184 | 0.618704382 | cat3 | from_5486 | to_5486 | art_short | art_short | H |
| 170 | 184 | 0.736945086 | cat3 | from_5487 | to_5487 | art_short | art_short | H |
| 171 | 184 | 0.77755055 | cat3 | from_5488 | to_5488 | art_short | art_short | H |
| 178 | 184 | 0.833328667 | cat3 | from_5489 | to_5489 | art_short | art_short | H |
| 3 | 185 | 0.759332037 | cat3 | from_5490 | to_5490 | art_long | art_short | J |
| 82 | 185 | 0.788755316 | cat3 | from_5491 | to_5491 | art_long | art_short | J |
| 89 | 185 | 0.696318298 | cat3 | from_5492 | to_5492 | art_long | art_short | J |
| 150 | 185 | 0.727453972 | cat3 | from_5493 | to_5493 | art_long | art_short | J |
| 175 | 185 | 0.774115725 | cat3 | from_5494 | to_5494 | art_short | art_short | H |
| 5 | 186 | 0.610654908 | cat3 | from_5495 | to_5495 | art_long | art_short | J |
| 12 | 186 | 0.670647344 | cat3 | from_5496 | to_5496 | art_long | art_short | J |
| 13 | 186 | 0.730725849 | cat3 | from_5497 | to_5497 | art_long | art_short | J |
| 19 | 186 | 0.65821652 | cat3 | from_5498 | to_5498 | art_long | art_short | J |
| 35 | 186 | -0.634431809 | cat3 | from_5499 | to_5499 | art_long | art_short | J |
| 63 | 186 | 0.800243474 | cat3 | from_5500 | to_5500 | art_long | art_short | J |
| 79 | 186 | 0.672413723 | cat3 | from_5501 | to_5501 | art_long | art_short | J |
| 114 | 186 | 0.700093491 | cat3 | from_5502 | to_5502 | art_long | art_short | J |
| 116 | 186 | 0.723119757 | cat3 | from_5503 | to_5503 | art_long | art_short | J |
| 156 | 186 | 0.864741012 | cat3 | from_5504 | to_5504 | art_short | art_short | H |
| 158 | 186 | 0.830204557 | cat3 | from_5505 | to_5505 | art_short | art_short | H |
| 163 | 186 | 0.623482352 | cat3 | from_5506 | to_5506 | art_short | art_short | H |
| 165 | 186 | 0.691808888 | cat3 | from_5507 | to_5507 | art_short | art_short | H |
| 167 | 186 | 0.676782134 | cat3 | from_5508 | to_5508 | art_short | art_short | H |
| 2 | 187 | 0.664124469 | cat3 | from_5509 | to_5509 | art_long | art_short | J |
| 16 | 187 | 0.617737463 | cat3 | from_5510 | to_5510 | art_long | art_short | J |

| | | | | | | | | |
|---|---|---|---|---|---|---|---|---|
| 53 | 187 | 0.837561837 | cat3 | from_5511 | to_5511 | art_long | art_short | J |
| 155 | 187 | 0.660034666 | cat3 | from_5512 | to_5512 | art_short | art_short | H |
| 164 | 187 | 0.606730122 | cat3 | from_5513 | to_5513 | art_short | art_short | H |
| 168 | 187 | 0.901169269 | cat3 | from_5514 | to_5514 | art_short | art_short | H |
| 169 | 187 | 0.654804471 | cat3 | from_5515 | to_5515 | art_short | art_short | H |
| 179 | 187 | 0.769655268 | cat3 | from_5516 | to_5516 | art_short | art_short | H |
| 180 | 187 | 0.968947743 | cat3 | from_5517 | to_5517 | art_short | art_short | H |
| 4 | 188 | 0.832193707 | cat3 | from_5518 | to_5518 | art_long | art_short | J |
| 6 | 188 | 0.794998479 | cat3 | from_5519 | to_5519 | art_long | art_short | J |
| 7 | 188 | 0.821322906 | cat3 | from_5520 | to_5520 | art_long | art_short | J |
| 11 | 188 | 0.708715941 | cat3 | from_5521 | to_5521 | art_long | art_short | J |
| 65 | 188 | 0.645745412 | cat3 | from_5522 | to_5522 | art_long | art_short | J |
| 118 | 188 | -0.65183888 | cat3 | from_5523 | to_5523 | art_long | art_short | J |
| 131 | 188 | -0.618215163 | cat3 | from_5524 | to_5524 | art_long | art_short | J |
| 137 | 188 | -0.615420559 | cat3 | from_5525 | to_5525 | art_long | art_short | J |
| 141 | 188 | -0.622415929 | cat3 | from_5526 | to_5526 | art_long | art_short | J |
| 144 | 188 | -0.663286293 | cat3 | from_5527 | to_5527 | art_long | art_short | J |
| 159 | 188 | 0.996466062 | cat3 | from_5528 | to_5528 | art_short | art_short | H |
| 160 | 188 | 0.879402759 | cat3 | from_5529 | to_5529 | art_short | art_short | H |
| 29 | 189 | 0.699742325 | cat3 | from_5530 | to_5530 | art_long | art_short | J |
| 66 | 189 | 0.912963922 | cat3 | from_5531 | to_5531 | art_long | art_short | J |
| 67 | 189 | 0.874241332 | cat3 | from_5532 | to_5532 | art_long | art_short | J |
| 71 | 189 | 0.715728802 | cat3 | from_5533 | to_5533 | art_long | art_short | J |
| 82 | 189 | 0.721799926 | cat3 | from_5534 | to_5534 | art_long | art_short | J |
| 89 | 189 | 0.658750322 | cat3 | from_5535 | to_5535 | art_long | art_short | J |
| 150 | 189 | 0.686256808 | cat3 | from_5536 | to_5536 | art_long | art_short | J |
| 175 | 189 | 0.687967015 | cat3 | from_5537 | to_5537 | art_short | art_short | H |
| 181 | 189 | 0.777278099 | cat3 | from_5538 | to_5538 | art_short | art_short | H |
| 32 | 190 | 0.769560316 | cat3 | from_5539 | to_5539 | art_long | art_short | J |
| 42 | 190 | 0.646415548 | cat3 | from_5540 | to_5540 | art_long | art_short | J |
| 45 | 190 | 0.627343074 | cat3 | from_5541 | to_5541 | art_long | art_short | J |
| 62 | 190 | 0.63011543 | cat3 | from_5542 | to_5542 | art_long | art_short | J |
| 86 | 190 | 0.694303085 | cat3 | from_5543 | to_5543 | art_long | art_short | J |
| 92 | 190 | 0.640025218 | cat3 | from_5544 | to_5544 | art_long | art_short | J |
| 94 | 190 | 0.676615988 | cat3 | from_5545 | to_5545 | art_long | art_short | J |
| 97 | 190 | 0.657101355 | cat3 | from_5546 | to_5546 | art_long | art_short | J |
| 101 | 190 | 0.642968721 | cat3 | from_5547 | to_5547 | art_long | art_short | J |
| 147 | 190 | 0.627706299 | cat3 | from_5548 | to_5548 | art_long | art_short | J |
| 178 | 190 | 0.628405366 | cat3 | from_5549 | to_5549 | art_short | art_short | H |
| 28 | 191 | 0.637128737 | cat3 | from_5550 | to_5550 | art_long | art_short | J |
| 32 | 191 | 0.829189836 | cat3 | from_5551 | to_5551 | art_long | art_short | J |
| 42 | 191 | 0.605706271 | cat3 | from_5552 | to_5552 | art_long | art_short | J |
| 45 | 191 | 0.642741337 | cat3 | from_5553 | to_5553 | art_long | art_short | J |
| 62 | 191 | 0.618316781 | cat3 | from_5554 | to_5554 | art_long | art_short | J |
| 86 | 191 | 0.664342792 | cat3 | from_5555 | to_5555 | art_long | art_short | J |
| 94 | 191 | 0.664832655 | cat3 | from_5556 | to_5556 | art_long | art_short | J |
| 97 | 191 | 0.612008081 | cat3 | from_5557 | to_5557 | art_long | art_short | J |
| 100 | 191 | 0.665005615 | cat3 | from_5558 | to_5558 | art_long | art_short | J |
| 170 | 191 | 0.612478433 | cat3 | from_5559 | to_5559 | art_short | art_short | H |
| 171 | 191 | 0.629926585 | cat3 | from_5560 | to_5560 | art_short | art_short | H |
| 173 | 191 | 0.680800235 | cat3 | from_5561 | to_5561 | art_short | art_short | H |
| 190 | 191 | 0.952284007 | cat3 | from_5562 | to_5562 | art_short | art_short | H |
| 12 | 192 | 0.657914703 | cat3 | from_5563 | to_5563 | art_long | art_short | J |
| 13 | 192 | 0.670534963 | cat3 | from_5564 | to_5564 | art_long | art_short | J |
| 15 | 192 | 0.795274508 | cat3 | from_5565 | to_5565 | art_long | art_short | J |
| 16 | 192 | 0.768691723 | cat3 | from_5566 | to_5566 | art_long | art_short | J |
| 18 | 192 | 0.693379053 | cat3 | from_5567 | to_5567 | art_long | art_short | J |
| 19 | 192 | 0.605982272 | cat3 | from_5568 | to_5568 | art_long | art_short | J |
| 39 | 192 | 0.655040555 | cat3 | from_5569 | to_5569 | art_long | art_short | J |
| 40 | 192 | 0.618596252 | cat3 | from_5570 | to_5570 | art_long | art_short | J |
| 162 | 192 | 0.727151579 | cat3 | from_5571 | to_5571 | art_short | art_short | H |
| 163 | 192 | 0.73798828 | cat3 | from_5572 | to_5572 | art_short | art_short | H |
| 164 | 192 | 0.735677264 | cat3 | from_5573 | to_5573 | art_short | art_short | H |
| 167 | 192 | 0.673576949 | cat3 | from_5574 | to_5574 | art_short | art_short | H |
| 169 | 192 | 0.656322005 | cat3 | from_5575 | to_5575 | art_short | art_short | H |
| 75 | 193 | 0.655882683 | cat3 | from_5576 | to_5576 | art_long | art_short | J |
| 84 | 193 | 0.711692618 | cat3 | from_5577 | to_5577 | art_long | art_short | J |
| 14 | 194 | 0.751418054 | cat3 | from_5578 | to_5578 | art_long | art_short | J |
| 24 | 194 | 0.878619626 | cat3 | from_5579 | to_5579 | art_long | art_short | J |
| 25 | 194 | 0.711347403 | cat3 | from_5580 | to_5580 | art_long | art_short | J |
| 28 | 194 | 0.947545962 | cat3 | from_5581 | to_5581 | art_long | art_short | J |
| 33 | 194 | 0.629379832 | cat3 | from_5582 | to_5582 | art_long | art_short | J |
| 50 | 194 | -0.616597327 | cat3 | from_5583 | to_5583 | art_long | art_short | J |
| 55 | 194 | 0.623698344 | cat3 | from_5584 | to_5584 | art_long | art_short | J |
| 59 | 194 | 0.861134133 | cat3 | from_5585 | to_5585 | art_long | art_short | J |
| 73 | 194 | 0.763534337 | cat3 | from_5586 | to_5586 | art_long | art_short | J |
| 74 | 194 | 0.746710446 | cat3 | from_5587 | to_5587 | art_long | art_short | J |
| 76 | 194 | 0.89575673 | cat3 | from_5588 | to_5588 | art_long | art_short | J |
| 78 | 194 | 0.776075252 | cat3 | from_5589 | to_5589 | art_long | art_short | J |
| 87 | 194 | 0.602416766 | cat3 | from_5590 | to_5590 | art_long | art_short | J |
| 90 | 194 | 0.775934473 | cat3 | from_5591 | to_5591 | art_long | art_short | J |
| 97 | 194 | 0.822864923 | cat3 | from_5592 | to_5592 | art_long | art_short | J |
| 109 | 194 | 0.631698826 | cat3 | from_5593 | to_5593 | art_long | art_short | J |
| 110 | 194 | 0.736453935 | cat3 | from_5594 | to_5594 | art_long | art_short | J |
| 112 | 194 | 0.61805171 | cat3 | from_5595 | to_5595 | art_long | art_short | J |
| 122 | 194 | 0.908790639 | cat3 | from_5596 | to_5596 | art_long | art_short | J |
| 173 | 194 | 0.967969794 | cat3 | from_5597 | to_5597 | art_short | art_short | H |
| 174 | 194 | 0.966304916 | cat3 | from_5598 | to_5598 | art_short | art_short | H |
| 190 | 194 | 0.608814337 | cat3 | from_5599 | to_5599 | art_short | art_short | H |
| 191 | 194 | 0.643869699 | cat3 | from_5600 | to_5600 | art_short | art_short | H |
| 24 | 195 | 0.631858901 | cat3 | from_5601 | to_5601 | art_long | art_short | J |
| 27 | 195 | 0.60679997 | cat3 | from_5602 | to_5602 | art_long | art_short | J |
| 30 | 195 | 0.632907502 | cat3 | from_5603 | to_5603 | art_long | art_short | J |
| 47 | 195 | 0.75156567 | cat3 | from_5604 | to_5604 | art_long | art_short | J |
| 51 | 195 | -0.641456497 | cat3 | from_5605 | to_5605 | art_long | art_short | J |
| 55 | 195 | 0.860905123 | cat3 | from_5606 | to_5606 | art_long | art_short | J |
| 70 | 195 | -0.645730394 | cat3 | from_5607 | to_5607 | art_long | art_short | J |
| 77 | 195 | 0.636970859 | cat3 | from_5608 | to_5608 | art_long | art_short | J |
| 80 | 195 | 0.750177852 | cat3 | from_5609 | to_5609 | art_long | art_short | J |
| 87 | 195 | 0.733587031 | cat3 | from_5610 | to_5610 | art_long | art_short | J |
| 95 | 195 | 0.925492188 | cat3 | from_5611 | to_5611 | art_long | art_short | J |
| 109 | 195 | 0.853032455 | cat3 | from_5612 | to_5612 | art_long | art_short | J |
| 119 | 195 | 0.722558208 | cat3 | from_5613 | to_5613 | art_long | art_short | J |
| 131 | 195 | 0.655394218 | cat3 | from_5614 | to_5614 | art_long | art_short | J |

| | | | | | | | | |
|---|---|---|---|---|---|---|---|---|
| 173 | 195 | 0.65084754 | cat3 | from_5615 | to_5615 | art_short | art_short | H |
| 2 | 196 | 0.664950392 | cat3 | from_5616 | to_5616 | art_long | art_short | J |
| 152 | 196 | -0.63892406 | cat3 | from_5617 | to_5617 | art_long | art_short | J |
| 153 | 196 | -0.613849879 | cat3 | from_5618 | to_5618 | art_long | art_short | J |
| 155 | 196 | 0.717011981 | cat3 | from_5619 | to_5619 | art_short | art_short | H |
| 162 | 196 | 0.957612418 | cat3 | from_5620 | to_5620 | art_short | art_short | H |
| 163 | 196 | 0.822078446 | cat3 | from_5621 | to_5621 | art_short | art_short | H |
| 179 | 196 | 0.607278889 | cat3 | from_5622 | to_5622 | art_short | art_short | H |
| 192 | 196 | 0.620073632 | cat3 | from_5623 | to_5623 | art_short | art_short | H |
| 12 | 197 | 0.769037264 | cat3 | from_5624 | to_5624 | art_long | art_short | J |
| 16 | 197 | 0.889632804 | cat3 | from_5625 | to_5625 | art_long | art_short | J |
| 18 | 197 | 0.654267921 | cat3 | from_5626 | to_5626 | art_long | art_short | J |
| 19 | 197 | 0.694318199 | cat3 | from_5627 | to_5627 | art_long | art_short | J |
| 39 | 197 | 0.828990344 | cat3 | from_5628 | to_5628 | art_long | art_short | J |
| 40 | 197 | 0.833467686 | cat3 | from_5629 | to_5629 | art_long | art_short | J |
| 53 | 197 | 0.638895887 | cat3 | from_5630 | to_5630 | art_long | art_short | J |
| 79 | 197 | 0.618328109 | cat3 | from_5631 | to_5631 | art_long | art_short | J |
| 155 | 197 | 0.646473077 | cat3 | from_5632 | to_5632 | art_short | art_short | H |
| 162 | 197 | 0.677344976 | cat3 | from_5633 | to_5633 | art_short | art_short | H |
| 163 | 197 | 0.734370917 | cat3 | from_5634 | to_5634 | art_short | art_short | H |
| 164 | 197 | 0.96690448 | cat3 | from_5635 | to_5635 | art_short | art_short | H |
| 167 | 197 | 0.784380345 | cat3 | from_5636 | to_5636 | art_short | art_short | H |
| 169 | 197 | 0.709888801 | cat3 | from_5637 | to_5637 | art_short | art_short | H |
| 179 | 197 | 0.729290659 | cat3 | from_5638 | to_5638 | art_short | art_short | H |
| 192 | 197 | 0.754109975 | cat3 | from_5639 | to_5639 | art_short | art_short | H |
| 196 | 197 | 0.637522565 | cat3 | from_5640 | to_5640 | art_short | art_short | H |
| 6 | 198 | 0.607127246 | cat3 | from_5641 | to_5641 | art_long | art_short | J |
| 8 | 198 | 0.725394362 | cat3 | from_5642 | to_5642 | art_long | art_short | J |
| 9 | 198 | 0.706513465 | cat3 | from_5643 | to_5643 | art_long | art_short | J |
| 10 | 198 | 0.707872731 | cat3 | from_5644 | to_5644 | art_long | art_short | J |
| 35 | 198 | -0.614970989 | cat3 | from_5645 | to_5645 | art_long | art_short | J |
| 65 | 198 | 0.608134067 | cat3 | from_5646 | to_5646 | art_long | art_short | J |
| 84 | 198 | 0.638692574 | cat3 | from_5647 | to_5647 | art_long | art_short | J |
| 125 | 198 | 0.786956207 | cat3 | from_5648 | to_5648 | art_long | art_short | J |
| 126 | 198 | 0.607270823 | cat3 | from_5649 | to_5649 | art_long | art_short | J |
| 156 | 198 | 0.745583733 | cat3 | from_5650 | to_5650 | art_short | art_short | H |
| 157 | 198 | 0.769864835 | cat3 | from_5651 | to_5651 | art_short | art_short | H |
| 158 | 198 | 0.715153566 | cat3 | from_5652 | to_5652 | art_short | art_short | H |
| 161 | 198 | 0.625769052 | cat3 | from_5653 | to_5653 | art_short | art_short | H |
| 186 | 198 | 0.685916182 | cat3 | from_5654 | to_5654 | art_short | art_short | H |
| 193 | 198 | 0.629333016 | cat3 | from_5655 | to_5655 | art_short | art_short | H |
| 49 | 199 | 0.756172887 | cat3 | from_5656 | to_5656 | art_long | art_short | J |
| 56 | 199 | 0.734127077 | cat3 | from_5657 | to_5657 | art_long | art_short | J |
| 65 | 199 | 0.659119933 | cat3 | from_5658 | to_5658 | art_long | art_short | J |
| 70 | 199 | 0.650530868 | cat3 | from_5659 | to_5659 | art_long | art_short | J |
| 75 | 199 | 0.837635136 | cat3 | from_5660 | to_5660 | art_long | art_short | J |
| 84 | 199 | 0.664269531 | cat3 | from_5661 | to_5661 | art_long | art_short | J |
| 85 | 199 | 0.814714542 | cat3 | from_5662 | to_5662 | art_long | art_short | J |
| 94 | 199 | -0.603348893 | cat3 | from_5663 | to_5663 | art_long | art_short | J |
| 113 | 199 | 0.656715846 | cat3 | from_5664 | to_5664 | art_long | art_short | J |
| 123 | 199 | 0.777437118 | cat3 | from_5665 | to_5665 | art_long | art_short | J |
| 146 | 199 | 0.863437702 | cat3 | from_5666 | to_5666 | art_long | art_short | J |
| 182 | 199 | 0.714438545 | cat3 | from_5667 | to_5667 | art_short | art_short | H |
| 193 | 199 | 0.816000158 | cat3 | from_5668 | to_5668 | art_short | art_short | H |
| 24 | 200 | 0.621388985 | cat3 | from_5669 | to_5669 | art_long | art_short | J |
| 51 | 200 | -0.659126167 | cat3 | from_5670 | to_5670 | art_long | art_short | J |
| 55 | 200 | 0.672406611 | cat3 | from_5671 | to_5671 | art_long | art_short | J |
| 70 | 200 | -0.647706523 | cat3 | from_5672 | to_5672 | art_long | art_short | J |
| 80 | 200 | 0.677578664 | cat3 | from_5673 | to_5673 | art_long | art_short | J |
| 95 | 200 | 0.839570719 | cat3 | from_5674 | to_5674 | art_long | art_short | J |
| 109 | 200 | 0.66213115 | cat3 | from_5675 | to_5675 | art_long | art_short | J |
| 131 | 200 | 0.623057817 | cat3 | from_5676 | to_5676 | art_long | art_short | J |
| 195 | 200 | 0.933232388 | cat3 | from_5677 | to_5677 | art_short | art_short | H |
| 8 | 201 | 0.646994758 | cat3 | from_5678 | to_5678 | art_long | art_short | J |
| 9 | 201 | 0.738774015 | cat3 | from_5679 | to_5679 | art_long | art_short | J |
| 10 | 201 | 0.782120732 | cat3 | from_5680 | to_5680 | art_long | art_short | J |
| 23 | 201 | 0.761994948 | cat3 | from_5681 | to_5681 | art_long | art_short | J |
| 36 | 201 | -0.632886315 | cat3 | from_5682 | to_5682 | art_long | art_short | J |
| 88 | 201 | 0.701893951 | cat3 | from_5683 | to_5683 | art_long | art_short | J |
| 125 | 201 | 0.737646223 | cat3 | from_5684 | to_5684 | art_long | art_short | J |
| 156 | 201 | 0.643002142 | cat3 | from_5685 | to_5685 | art_short | art_short | H |
| 158 | 201 | 0.654576886 | cat3 | from_5686 | to_5686 | art_short | art_short | H |
| 193 | 201 | 0.602562649 | cat3 | from_5687 | to_5687 | art_short | art_short | H |
| 198 | 201 | 0.661620331 | cat3 | from_5688 | to_5688 | art_short | art_short | H |
| 51 | 202 | 0.633161223 | cat3 | from_5689 | to_5689 | art_long | art_short | J |
| 61 | 203 | 0.727753734 | cat3 | from_5690 | to_5690 | art_long | art_short | J |
| 70 | 203 | -0.627872432 | cat3 | from_5691 | to_5691 | art_long | art_short | J |
| 102 | 203 | 0.656369646 | cat3 | from_5692 | to_5692 | art_long | art_short | J |
| 106 | 203 | 0.632268665 | cat3 | from_5693 | to_5693 | art_long | art_short | J |
| 115 | 203 | -0.633860262 | cat3 | from_5694 | to_5694 | art_long | art_short | J |
| 127 | 203 | -0.608752575 | cat3 | from_5695 | to_5695 | art_long | art_short | J |
| 135 | 203 | 0.76231608 | cat3 | from_5696 | to_5696 | art_long | art_short | J |
| 170 | 203 | 0.642437405 | cat3 | from_5697 | to_5697 | art_short | art_short | H |
| 176 | 203 | 0.981275209 | cat3 | from_5698 | to_5698 | art_short | art_short | H |
| 44 | 204 | 0.624840977 | cat3 | from_5699 | to_5699 | art_long | art_short | J |
| 46 | 204 | 0.666544671 | cat3 | from_5700 | to_5700 | art_long | art_short | J |
| 51 | 204 | -0.648820056 | cat3 | from_5701 | to_5701 | art_long | art_short | J |
| 61 | 204 | 0.691303991 | cat3 | from_5702 | to_5702 | art_long | art_short | J |
| 70 | 204 | -0.639935214 | cat3 | from_5703 | to_5703 | art_long | art_short | J |
| 102 | 204 | 0.779777458 | cat3 | from_5704 | to_5704 | art_long | art_short | J |
| 104 | 204 | 0.630899559 | cat3 | from_5705 | to_5705 | art_long | art_short | J |
| 106 | 204 | 0.733333235 | cat3 | from_5706 | to_5706 | art_long | art_short | J |
| 135 | 204 | 0.808963935 | cat3 | from_5707 | to_5707 | art_long | art_short | J |
| 139 | 204 | 0.66717084 | cat3 | from_5708 | to_5708 | art_long | art_short | J |
| 166 | 204 | 0.688563559 | cat3 | from_5709 | to_5709 | art_short | art_short | H |
| 170 | 204 | 0.725445777 | cat3 | from_5710 | to_5710 | art_short | art_short | H |
| 171 | 204 | 0.689692663 | cat3 | from_5711 | to_5711 | art_short | art_short | H |
| 176 | 204 | 0.916417657 | cat3 | from_5712 | to_5712 | art_short | art_short | H |
| 203 | 204 | 0.951312739 | cat3 | from_5713 | to_5713 | art_short | art_short | H |
| 4 | 205 | 0.619347218 | cat3 | from_5714 | to_5714 | art_long | art_short | J |
| 7 | 205 | 0.717130049 | cat3 | from_5715 | to_5715 | art_long | art_short | J |
| 11 | 205 | 0.814113032 | cat3 | from_5716 | to_5716 | art_long | art_short | J |
| 22 | 205 | 0.675772134 | cat3 | from_5717 | to_5717 | art_long | art_short | J |
| 26 | 205 | -0.613995866 | cat3 | from_5718 | to_5718 | art_long | art_short | J |

| | | | | | | | | |
|---|---|---|---|---|---|---|---|---|
| 37 | 205 | -0.680323043 | cat3 | from_5719 | to_5719 | art_long | art_short | J |
| 88 | 205 | 0.642359937 | cat3 | from_5720 | to_5720 | art_long | art_short | J |
| 90 | 205 | -0.64784145 | cat3 | from_5721 | to_5721 | art_long | art_short | J |
| 115 | 205 | -0.646775561 | cat3 | from_5722 | to_5722 | art_long | art_short | J |
| 126 | 205 | 0.691162061 | cat3 | from_5723 | to_5723 | art_long | art_short | J |
| 129 | 205 | -0.690044502 | cat3 | from_5724 | to_5724 | art_long | art_short | J |
| 130 | 205 | -0.633344549 | cat3 | from_5725 | to_5725 | art_long | art_short | J |
| 135 | 205 | 0.654944378 | cat3 | from_5726 | to_5726 | art_long | art_short | J |
| 159 | 205 | 0.615363541 | cat3 | from_5727 | to_5727 | art_short | art_short | H |
| 160 | 205 | 0.626946783 | cat3 | from_5728 | to_5728 | art_short | art_short | H |
| 161 | 205 | 0.707120744 | cat3 | from_5729 | to_5729 | art_short | art_short | H |
| 188 | 205 | 0.638418335 | cat3 | from_5730 | to_5730 | art_short | art_short | H |
| 21 | 206 | -0.627710643 | cat3 | from_5731 | to_5731 | art_long | art_short | J |
| 42 | 206 | -0.631182228 | cat3 | from_5732 | to_5732 | art_long | art_short | J |
| 49 | 206 | -0.616913093 | cat3 | from_5733 | to_5733 | art_long | art_short | J |
| 54 | 206 | 0.839618125 | cat3 | from_5734 | to_5734 | art_long | art_short | J |
| 77 | 206 | 0.891936508 | cat3 | from_5735 | to_5735 | art_long | art_short | J |
| 80 | 206 | 0.937371722 | cat3 | from_5736 | to_5736 | art_long | art_short | J |
| 81 | 206 | 0.695081343 | cat3 | from_5737 | to_5737 | art_long | art_short | J |
| 83 | 206 | 0.613072848 | cat3 | from_5738 | to_5738 | art_long | art_short | J |
| 85 | 206 | -0.653013825 | cat3 | from_5739 | to_5739 | art_long | art_short | J |
| 92 | 206 | -0.670992325 | cat3 | from_5740 | to_5740 | art_long | art_short | J |
| 95 | 206 | 0.768823937 | cat3 | from_5741 | to_5741 | art_long | art_short | J |
| 109 | 206 | 0.64059032 | cat3 | from_5742 | to_5742 | art_long | art_short | J |
| 119 | 206 | 0.708401803 | cat3 | from_5743 | to_5743 | art_long | art_short | J |
| 120 | 206 | 0.742366195 | cat3 | from_5744 | to_5744 | art_long | art_short | J |
| 127 | 206 | -0.637523071 | cat3 | from_5745 | to_5745 | art_long | art_short | J |
| 133 | 206 | -0.762022127 | cat3 | from_5746 | to_5746 | art_long | art_short | J |
| 140 | 206 | 0.717626503 | cat3 | from_5747 | to_5747 | art_long | art_short | J |
| 147 | 206 | -0.700986874 | cat3 | from_5748 | to_5748 | art_long | art_short | J |
| 177 | 206 | -0.654316342 | cat3 | from_5749 | to_5749 | art_short | art_short | H |
| 195 | 206 | 0.794026735 | cat3 | from_5750 | to_5750 | art_short | art_short | H |
| 200 | 206 | 0.765933114 | cat3 | from_5751 | to_5751 | art_short | art_short | H |
| 4 | 207 | 0.606079155 | cat3 | from_5752 | to_5752 | art_long | art_short | J |
| 7 | 207 | 0.610276112 | cat3 | from_5753 | to_5753 | art_long | art_short | J |
| 22 | 207 | 0.629556213 | cat3 | from_5754 | to_5754 | art_long | art_short | J |
| 34 | 207 | 0.920450437 | cat3 | from_5755 | to_5755 | art_long | art_short | J |
| 64 | 207 | 0.635895923 | cat3 | from_5756 | to_5756 | art_long | art_short | J |
| 65 | 207 | 0.934624708 | cat3 | from_5757 | to_5757 | art_long | art_short | J |
| 85 | 207 | 0.626193857 | cat3 | from_5758 | to_5758 | art_long | art_short | J |
| 113 | 207 | 0.904799025 | cat3 | from_5759 | to_5759 | art_long | art_short | J |
| 116 | 207 | 0.618956963 | cat3 | from_5760 | to_5760 | art_long | art_short | J |
| 117 | 207 | 0.639988745 | cat3 | from_5761 | to_5761 | art_long | art_short | J |
| 125 | 207 | 0.636948334 | cat3 | from_5762 | to_5762 | art_long | art_short | J |
| 127 | 207 | 0.848229517 | cat3 | from_5763 | to_5763 | art_long | art_short | J |
| 159 | 207 | 0.679595112 | cat3 | from_5764 | to_5764 | art_short | art_short | H |
| 160 | 207 | 0.610030057 | cat3 | from_5765 | to_5765 | art_short | art_short | H |
| 188 | 207 | 0.669583032 | cat3 | from_5766 | to_5766 | art_short | art_short | H |
| 198 | 207 | 0.620623964 | cat3 | from_5767 | to_5767 | art_short | art_short | H |
| 199 | 207 | 0.667262524 | cat3 | from_5768 | to_5768 | art_short | art_short | H |
| 29 | 208 | 0.68004568 | cat3 | from_5769 | to_5769 | art_long | art_short | J |
| 66 | 208 | 0.689100423 | cat3 | from_5770 | to_5770 | art_long | art_short | J |
| 71 | 208 | 0.672170342 | cat3 | from_5771 | to_5771 | art_long | art_short | J |
| 82 | 208 | 0.643120034 | cat3 | from_5772 | to_5772 | art_long | art_short | J |
| 89 | 208 | 0.622970337 | cat3 | from_5773 | to_5773 | art_long | art_short | J |
| 96 | 208 | 0.695538581 | cat3 | from_5774 | to_5774 | art_long | art_short | J |
| 150 | 208 | 0.641106758 | cat3 | from_5775 | to_5775 | art_long | art_short | J |
| 175 | 208 | 0.841038422 | cat3 | from_5776 | to_5776 | art_short | art_short | H |
| 181 | 208 | 0.744472367 | cat3 | from_5777 | to_5777 | art_short | art_short | H |
| 189 | 208 | 0.740445868 | cat3 | from_5778 | to_5778 | art_short | art_short | H |
| 29 | 209 | 0.689783 | cat3 | from_5779 | to_5779 | art_long | art_short | J |
| 66 | 209 | 0.654796255 | cat3 | from_5780 | to_5780 | art_long | art_short | J |
| 71 | 209 | 0.685827295 | cat3 | from_5781 | to_5781 | art_long | art_short | J |
| 82 | 209 | 0.875552489 | cat3 | from_5782 | to_5782 | art_long | art_short | J |
| 89 | 209 | 0.847660921 | cat3 | from_5783 | to_5783 | art_long | art_short | J |
| 96 | 209 | 0.624318418 | cat3 | from_5784 | to_5784 | art_long | art_short | J |
| 121 | 209 | 0.730207561 | cat3 | from_5785 | to_5785 | art_long | art_short | J |
| 150 | 209 | 0.866233166 | cat3 | from_5786 | to_5786 | art_long | art_short | J |
| 175 | 209 | 0.96390411 | cat3 | from_5787 | to_5787 | art_short | art_short | H |
| 181 | 209 | 0.76234705 | cat3 | from_5788 | to_5788 | art_short | art_short | H |
| 185 | 209 | 0.800750876 | cat3 | from_5789 | to_5789 | art_short | art_short | H |
| 189 | 209 | 0.7310813 | cat3 | from_5790 | to_5790 | art_short | art_short | H |
| 208 | 209 | 0.866321968 | cat3 | from_5791 | to_5791 | art_short | art_short | H |
| 96 | 210 | 0.634828158 | cat3 | from_5792 | to_5792 | art_long | art_short | J |
| 175 | 210 | 0.870351336 | cat3 | from_5793 | to_5793 | art_short | art_short | H |
| 185 | 210 | 0.638677616 | cat3 | from_5794 | to_5794 | art_short | art_short | H |
| 200 | 210 | 0.661026833 | cat3 | from_5795 | to_5795 | art_short | art_short | H |
| 208 | 210 | 0.88063123 | cat3 | from_5796 | to_5796 | art_short | art_short | H |
| 209 | 210 | 0.858809469 | cat3 | from_5797 | to_5797 | art_short | art_short | H |
| 30 | 211 | 0.975083179 | cat3 | from_5798 | to_5798 | art_long | art_short | J |
| 47 | 211 | 0.902512086 | cat3 | from_5799 | to_5799 | art_long | art_short | J |
| 55 | 211 | 0.855644875 | cat3 | from_5800 | to_5800 | art_long | art_short | J |
| 58 | 211 | 0.783188301 | cat3 | from_5801 | to_5801 | art_long | art_short | J |
| 69 | 211 | 0.777853652 | cat3 | from_5802 | to_5802 | art_long | art_short | J |
| 72 | 211 | 0.909667159 | cat3 | from_5803 | to_5803 | art_long | art_short | J |
| 83 | 211 | 0.656919525 | cat3 | from_5804 | to_5804 | art_long | art_short | J |
| 86 | 211 | 0.635097807 | cat3 | from_5805 | to_5805 | art_long | art_short | J |
| 87 | 211 | 0.926084521 | cat3 | from_5806 | to_5806 | art_long | art_short | J |
| 90 | 211 | 0.6834167 | cat3 | from_5807 | to_5807 | art_long | art_short | J |
| 95 | 211 | 0.666195728 | cat3 | from_5808 | to_5808 | art_long | art_short | J |
| 97 | 211 | 0.640239402 | cat3 | from_5809 | to_5809 | art_long | art_short | J |
| 109 | 211 | 0.849014824 | cat3 | from_5810 | to_5810 | art_long | art_short | J |
| 111 | 211 | 0.919989283 | cat3 | from_5811 | to_5811 | art_long | art_short | J |
| 119 | 211 | 0.75693949 | cat3 | from_5812 | to_5812 | art_long | art_short | J |
| 140 | 211 | 0.676005528 | cat3 | from_5813 | to_5813 | art_long | art_short | J |
| 172 | 211 | 0.602130252 | cat3 | from_5814 | to_5814 | art_short | art_short | H |
| 173 | 211 | 0.607911038 | cat3 | from_5815 | to_5815 | art_short | art_short | H |
| 195 | 211 | 0.67125874 | cat3 | from_5816 | to_5816 | art_short | art_short | H |
| 20 | 212 | -0.606961633 | cat3 | from_5817 | to_5817 | art_long | art_short | J |
| 56 | 212 | 0.705187469 | cat3 | from_5818 | to_5818 | art_long | art_short | J |
| 57 | 212 | 0.7488004 | cat3 | from_5819 | to_5819 | art_long | art_short | J |
| 75 | 212 | 0.64168451 | cat3 | from_5820 | to_5820 | art_long | art_short | J |
| 86 | 212 | -0.631697527 | cat3 | from_5821 | to_5821 | art_long | art_short | J |
| 105 | 212 | -0.613304259 | cat3 | from_5822 | to_5822 | art_long | art_short | J |

| | | | | | | | | |
|---|---|---|---|---|---|---|---|---|
| 123 | 212 | 0.657593627 | cat3 | from_5823 | to_5823 | art_long | art_short | J |
| 146 | 212 | 0.782202738 | cat3 | from_5824 | to_5824 | art_long | art_short | J |
| 193 | 212 | 0.606391517 | cat3 | from_5825 | to_5825 | art_short | art_short | H |
| 199 | 212 | 0.760466526 | cat3 | from_5826 | to_5826 | art_short | art_short | H |
| 17 | 213 | 0.652828179 | cat3 | from_5827 | to_5827 | art_long | art_short | J |
| 27 | 213 | -0.695023437 | cat3 | from_5828 | to_5828 | art_long | art_short | J |
| 36 | 213 | 0.625739835 | cat3 | from_5829 | to_5829 | art_long | art_short | J |
| 50 | 213 | 0.767120379 | cat3 | from_5830 | to_5830 | art_long | art_short | J |
| 64 | 213 | -0.615096367 | cat3 | from_5831 | to_5831 | art_long | art_short | J |
| 138 | 213 | 0.68860003 | cat3 | from_5832 | to_5832 | art_long | art_short | J |
| 143 | 213 | 0.607824276 | cat3 | from_5833 | to_5833 | art_long | art_short | J |
| 154 | 213 | 0.764095758 | cat3 | from_5834 | to_5834 | art_short | art_short | H |
| 172 | 213 | -0.676528366 | cat3 | from_5835 | to_5835 | art_short | art_short | H |
| 177 | 213 | 0.692573169 | cat3 | from_5836 | to_5836 | art_short | art_short | H |
| 35 | 214 | 0.849576426 | cat3 | from_5837 | to_5837 | art_long | art_short | J |
| 38 | 214 | 0.862015809 | cat3 | from_5838 | to_5838 | art_long | art_short | J |
| 48 | 214 | 0.820838868 | cat3 | from_5839 | to_5839 | art_long | art_short | J |
| 50 | 214 | 0.88153794 | cat3 | from_5840 | to_5840 | art_long | art_short | J |
| 104 | 214 | 0.768040697 | cat3 | from_5841 | to_5841 | art_long | art_short | J |
| 132 | 214 | 0.851782342 | cat3 | from_5842 | to_5842 | art_long | art_short | J |
| 143 | 214 | 0.895129639 | cat3 | from_5843 | to_5843 | art_long | art_short | J |
| 166 | 214 | 0.860517598 | cat3 | from_5844 | to_5844 | art_short | art_short | H |
| 177 | 214 | 0.623263979 | cat3 | from_5845 | to_5845 | art_short | art_short | H |
| 213 | 214 | 0.7294406 | cat3 | from_5846 | to_5846 | art_short | art_short | H |
| 29 | 215 | 0.609949079 | cat3 | from_5847 | to_5847 | art_long | art_short | J |
| 39 | 215 | 0.64791389 | cat3 | from_5848 | to_5848 | art_long | art_short | J |
| 71 | 215 | 0.620230214 | cat3 | from_5849 | to_5849 | art_long | art_short | J |
| 82 | 215 | 0.744091528 | cat3 | from_5850 | to_5850 | art_long | art_short | J |
| 89 | 215 | 0.773872846 | cat3 | from_5851 | to_5851 | art_long | art_short | J |
| 98 | 215 | 0.625495175 | cat3 | from_5852 | to_5852 | art_long | art_short | J |
| 150 | 215 | 0.768222221 | cat3 | from_5853 | to_5853 | art_long | art_short | J |
| 164 | 215 | 0.620926783 | cat3 | from_5854 | to_5854 | art_short | art_short | H |
| 181 | 215 | 0.647636445 | cat3 | from_5855 | to_5855 | art_short | art_short | H |
| 209 | 215 | 0.609895559 | cat3 | from_5856 | to_5856 | art_short | art_short | H |
| 12 | 216 | 0.861797364 | cat3 | from_5857 | to_5857 | art_long | art_short | J |
| 15 | 216 | 0.741565446 | cat3 | from_5858 | to_5858 | art_long | art_short | J |
| 16 | 216 | 0.741660276 | cat3 | from_5859 | to_5859 | art_long | art_short | J |
| 18 | 216 | 0.866374425 | cat3 | from_5860 | to_5860 | art_long | art_short | J |
| 19 | 216 | 0.721716042 | cat3 | from_5861 | to_5861 | art_long | art_short | J |
| 39 | 216 | 0.694321493 | cat3 | from_5862 | to_5862 | art_long | art_short | J |
| 40 | 216 | 0.612912809 | cat3 | from_5863 | to_5863 | art_long | art_short | J |
| 79 | 216 | 0.730687039 | cat3 | from_5864 | to_5864 | art_long | art_short | J |
| 164 | 216 | 0.755439514 | cat3 | from_5865 | to_5865 | art_short | art_short | H |
| 167 | 216 | 0.857975593 | cat3 | from_5866 | to_5866 | art_short | art_short | H |
| 192 | 216 | 0.788311498 | cat3 | from_5867 | to_5867 | art_short | art_short | H |
| 197 | 216 | 0.816503382 | cat3 | from_5868 | to_5868 | art_short | art_short | H |
| 1 | 217 | 0.687417585 | cat3 | from_5869 | to_5869 | art_long | art_short | J |
| 14 | 217 | -0.630854537 | cat3 | from_5870 | to_5870 | art_long | art_short | J |
| 73 | 217 | -0.626323926 | cat3 | from_5871 | to_5871 | art_long | art_short | J |
| 149 | 217 | -0.606471574 | cat3 | from_5872 | to_5872 | art_long | art_short | J |
| 30 | 218 | 0.96995483 | cat3 | from_5873 | to_5873 | art_long | art_short | J |
| 47 | 218 | 0.840189794 | cat3 | from_5874 | to_5874 | art_long | art_short | J |
| 55 | 218 | 0.888638242 | cat3 | from_5875 | to_5875 | art_long | art_short | J |
| 58 | 218 | 0.865630046 | cat3 | from_5876 | to_5876 | art_long | art_short | J |
| 69 | 218 | 0.844912723 | cat3 | from_5877 | to_5877 | art_long | art_short | J |
| 72 | 218 | 0.822013796 | cat3 | from_5878 | to_5878 | art_long | art_short | J |
| 77 | 218 | 0.732870056 | cat3 | from_5879 | to_5879 | art_long | art_short | J |
| 80 | 218 | 0.742130655 | cat3 | from_5880 | to_5880 | art_long | art_short | J |
| 83 | 218 | 0.764343931 | cat3 | from_5881 | to_5881 | art_long | art_short | J |
| 87 | 218 | 0.898789594 | cat3 | from_5882 | to_5882 | art_long | art_short | J |
| 95 | 218 | 0.80684306 | cat3 | from_5883 | to_5883 | art_long | art_short | J |
| 109 | 218 | 0.912507991 | cat3 | from_5884 | to_5884 | art_long | art_short | J |
| 111 | 218 | 0.787766719 | cat3 | from_5885 | to_5885 | art_long | art_short | J |
| 119 | 218 | 0.892405273 | cat3 | from_5886 | to_5886 | art_long | art_short | J |
| 140 | 218 | 0.81019245 | cat3 | from_5887 | to_5887 | art_long | art_short | J |
| 172 | 218 | 0.628791514 | cat3 | from_5888 | to_5888 | art_short | art_short | H |
| 173 | 218 | 0.609060409 | cat3 | from_5889 | to_5889 | art_short | art_short | H |
| 174 | 218 | 0.6208774 | cat3 | from_5890 | to_5890 | art_short | art_short | H |
| 194 | 218 | 0.631275341 | cat3 | from_5891 | to_5891 | art_short | art_short | H |
| 195 | 218 | 0.782566673 | cat3 | from_5892 | to_5892 | art_short | art_short | H |
| 206 | 218 | 0.68274633 | cat3 | from_5893 | to_5893 | art_short | art_short | H |
| 211 | 218 | 0.947656607 | cat3 | from_5894 | to_5894 | art_short | art_short | H |
| 1 | 219 | 0.648004684 | cat3 | from_5895 | to_5895 | art_long | art_untreat | E |
| 30 | 219 | 0.741114913 | cat3 | from_5896 | to_5896 | art_long | art_untreat | E |
| 72 | 219 | 0.632127831 | cat3 | from_5897 | to_5897 | art_long | art_untreat | E |
| 211 | 219 | 0.785311052 | cat3 | from_5898 | to_5898 | art_short | art_untreat | C |
| 218 | 219 | 0.752857321 | cat3 | from_5899 | to_5899 | art_short | art_untreat | C |
| 13 | 220 | 0.602630504 | cat3 | from_5900 | to_5900 | art_long | art_untreat | E |
| 15 | 220 | 0.621636039 | cat3 | from_5901 | to_5901 | art_long | art_untreat | E |
| 16 | 220 | 0.667569763 | cat3 | from_5902 | to_5902 | art_long | art_untreat | E |
| 39 | 220 | 0.623786029 | cat3 | from_5903 | to_5903 | art_long | art_untreat | E |
| 114 | 220 | 0.624810147 | cat3 | from_5904 | to_5904 | art_long | art_untreat | E |
| 162 | 220 | 0.764705738 | cat3 | from_5905 | to_5905 | art_short | art_untreat | C |
| 163 | 220 | 0.839767503 | cat3 | from_5906 | to_5906 | art_short | art_untreat | C |
| 164 | 220 | 0.769072563 | cat3 | from_5907 | to_5907 | art_short | art_untreat | C |
| 165 | 220 | 0.742224447 | cat3 | from_5908 | to_5908 | art_short | art_untreat | C |
| 168 | 220 | 0.663670395 | cat3 | from_5909 | to_5909 | art_short | art_untreat | C |
| 169 | 220 | 0.936509501 | cat3 | from_5910 | to_5910 | art_short | art_untreat | C |
| 180 | 220 | 0.666255511 | cat3 | from_5911 | to_5911 | art_short | art_untreat | C |
| 187 | 220 | 0.682894229 | cat3 | from_5912 | to_5912 | art_short | art_untreat | C |
| 192 | 220 | 0.703481521 | cat3 | from_5913 | to_5913 | art_short | art_untreat | C |
| 196 | 220 | 0.739907692 | cat3 | from_5914 | to_5914 | art_short | art_untreat | C |
| 197 | 220 | 0.771867942 | cat3 | from_5915 | to_5915 | art_short | art_untreat | C |
| 31 | 221 | 0.718613207 | cat3 | from_5916 | to_5916 | art_long | art_untreat | E |
| 33 | 221 | 0.748231826 | cat3 | from_5917 | to_5917 | art_long | art_untreat | E |
| 57 | 221 | -0.614346617 | cat3 | from_5918 | to_5918 | art_long | art_untreat | E |
| 90 | 221 | 0.785684513 | cat3 | from_5919 | to_5919 | art_long | art_untreat | E |
| 173 | 221 | 0.767175972 | cat3 | from_5920 | to_5920 | art_short | art_untreat | C |
| 174 | 221 | 0.680113329 | cat3 | from_5921 | to_5921 | art_short | art_untreat | C |
| 194 | 221 | 0.731039521 | cat3 | from_5922 | to_5922 | art_short | art_untreat | C |
| 219 | 221 | 0.734091344 | cat3 | from_5923 | to_5923 | art_untreat | art_untreat | A |
| 32 | 222 | -0.674348934 | cat3 | from_5924 | to_5924 | art_long | art_untreat | E |
| 43 | 222 | -0.610525334 | cat3 | from_5925 | to_5925 | art_long | art_untreat | E |
| 49 | 222 | 0.633239631 | cat3 | from_5926 | to_5926 | art_long | art_untreat | E |

| | | | | | | | | |
|---|---|---|---|---|---|---|---|---|
| 56 | 222 | 0.772864459 | cat3 | from_5927 | to_5927 | art_long | art_untreat | E |
| 74 | 222 | -0.620586537 | cat3 | from_5928 | to_5928 | art_long | art_untreat | E |
| 85 | 222 | 0.67103589 | cat3 | from_5929 | to_5929 | art_long | art_untreat | E |
| 182 | 222 | 0.830668606 | cat3 | from_5930 | to_5930 | art_short | art_untreat | C |
| 199 | 222 | 0.769766403 | cat3 | from_5931 | to_5931 | art_short | art_untreat | C |
| 201 | 222 | 0.612918123 | cat3 | from_5932 | to_5932 | art_short | art_untreat | C |
| 44 | 223 | 0.673512513 | cat3 | from_5933 | to_5933 | art_long | art_untreat | E |
| 46 | 223 | 0.672751558 | cat3 | from_5934 | to_5934 | art_long | art_untreat | E |
| 52 | 223 | 0.809099191 | cat3 | from_5935 | to_5935 | art_long | art_untreat | E |
| 60 | 223 | 0.636351489 | cat3 | from_5936 | to_5936 | art_long | art_untreat | E |
| 61 | 223 | 0.734460111 | cat3 | from_5937 | to_5937 | art_long | art_untreat | E |
| 99 | 223 | 0.667151238 | cat3 | from_5938 | to_5938 | art_long | art_untreat | E |
| 102 | 223 | 0.645174247 | cat3 | from_5939 | to_5939 | art_long | art_untreat | E |
| 106 | 223 | 0.694541722 | cat3 | from_5940 | to_5940 | art_long | art_untreat | E |
| 128 | 223 | 0.660049343 | cat3 | from_5941 | to_5941 | art_long | art_untreat | E |
| 148 | 223 | 0.635615039 | cat3 | from_5942 | to_5942 | art_long | art_untreat | E |
| 176 | 223 | 0.833065582 | cat3 | from_5943 | to_5943 | art_short | art_untreat | C |
| 183 | 223 | 0.844444374 | cat3 | from_5944 | to_5944 | art_short | art_untreat | C |
| 203 | 223 | 0.755871206 | cat3 | from_5945 | to_5945 | art_short | art_untreat | C |
| 204 | 223 | 0.743494716 | cat3 | from_5946 | to_5946 | art_short | art_untreat | C |
| 29 | 224 | 0.773277899 | cat3 | from_5947 | to_5947 | art_long | art_untreat | E |
| 71 | 224 | 0.723692339 | cat3 | from_5948 | to_5948 | art_long | art_untreat | E |
| 82 | 224 | 0.632817287 | cat3 | from_5949 | to_5949 | art_long | art_untreat | E |
| 89 | 224 | 0.68261699 | cat3 | from_5950 | to_5950 | art_long | art_untreat | E |
| 124 | 224 | 0.768274144 | cat3 | from_5951 | to_5951 | art_long | art_untreat | E |
| 150 | 224 | 0.681704799 | cat3 | from_5952 | to_5952 | art_long | art_untreat | E |
| 175 | 224 | 0.682930481 | cat3 | from_5953 | to_5953 | art_short | art_untreat | C |
| 181 | 224 | 0.783913096 | cat3 | from_5954 | to_5954 | art_short | art_untreat | C |
| 208 | 224 | 0.736206974 | cat3 | from_5955 | to_5955 | art_short | art_untreat | C |
| 209 | 224 | 0.730699297 | cat3 | from_5956 | to_5956 | art_short | art_untreat | C |
| 30 | 225 | 0.838920237 | cat3 | from_5957 | to_5957 | art_long | art_untreat | E |
| 47 | 225 | 0.700522665 | cat3 | from_5958 | to_5958 | art_long | art_untreat | E |
| 55 | 225 | 0.717530752 | cat3 | from_5959 | to_5959 | art_long | art_untreat | E |
| 72 | 225 | 0.680634681 | cat3 | from_5960 | to_5960 | art_long | art_untreat | E |
| 87 | 225 | 0.715573921 | cat3 | from_5961 | to_5961 | art_long | art_untreat | E |
| 95 | 225 | 0.643043188 | cat3 | from_5962 | to_5962 | art_long | art_untreat | E |
| 109 | 225 | 0.722905791 | cat3 | from_5963 | to_5963 | art_long | art_untreat | E |
| 111 | 225 | 0.643511803 | cat3 | from_5964 | to_5964 | art_long | art_untreat | E |
| 119 | 225 | 0.646096161 | cat3 | from_5965 | to_5965 | art_long | art_untreat | E |
| 140 | 225 | 0.603877139 | cat3 | from_5966 | to_5966 | art_long | art_untreat | E |
| 195 | 225 | 0.780001063 | cat3 | from_5967 | to_5967 | art_short | art_untreat | C |
| 206 | 225 | 0.678368925 | cat3 | from_5968 | to_5968 | art_short | art_untreat | C |
| 211 | 225 | 0.879523193 | cat3 | from_5969 | to_5969 | art_short | art_untreat | C |
| 218 | 225 | 0.906950282 | cat3 | from_5970 | to_5970 | art_short | art_untreat | C |
| 219 | 225 | 0.916531785 | cat3 | from_5971 | to_5971 | art_untreat | art_untreat | A |
| 221 | 225 | 0.735866623 | cat3 | from_5972 | to_5972 | art_untreat | art_untreat | A |
| 52 | 226 | 0.633728595 | cat3 | from_5973 | to_5973 | art_long | art_untreat | E |
| 74 | 226 | -0.646393957 | cat3 | from_5974 | to_5974 | art_long | art_untreat | E |
| 78 | 226 | -0.609352653 | cat3 | from_5975 | to_5975 | art_long | art_untreat | E |
| 99 | 226 | 0.654377338 | cat3 | from_5976 | to_5976 | art_long | art_untreat | E |
| 105 | 226 | 0.702190823 | cat3 | from_5977 | to_5977 | art_long | art_untreat | E |
| 176 | 226 | 0.712018229 | cat3 | from_5978 | to_5978 | art_short | art_untreat | C |
| 183 | 226 | 0.766975035 | cat3 | from_5979 | to_5979 | art_short | art_untreat | C |
| 203 | 226 | 0.64973853 | cat3 | from_5980 | to_5980 | art_short | art_untreat | C |
| 217 | 226 | 0.651919955 | cat3 | from_5981 | to_5981 | art_short | art_untreat | C |
| 223 | 226 | 0.880974376 | cat3 | from_5982 | to_5982 | art_untreat | art_untreat | A |
| 198 | 227 | 0.711557799 | cat3 | from_5983 | to_5983 | art_short | ec | K |
| 113 | 228 | 0.789166191 | cat3 | from_5984 | to_5984 | art_long | ec | N |
| 152 | 228 | -0.615966921 | cat3 | from_5985 | to_5985 | art_long | ec | N |
| 153 | 228 | -0.649203263 | cat3 | from_5986 | to_5986 | art_long | ec | N |
| 207 | 228 | 0.671753424 | cat3 | from_5987 | to_5987 | art_short | ec | K |
| 212 | 228 | 0.650608121 | cat3 | from_5988 | to_5988 | art_short | ec | K |
| 227 | 228 | 0.899560102 | cat3 | from_5989 | to_5989 | ec | ec | P |
| 12 | 229 | 0.680477941 | cat3 | from_5990 | to_5990 | art_long | ec | N |
| 40 | 229 | 0.643564485 | cat3 | from_5991 | to_5991 | art_long | ec | N |
| 156 | 229 | 0.612753743 | cat3 | from_5992 | to_5992 | art_short | ec | K |
| 163 | 229 | 0.612680281 | cat3 | from_5993 | to_5993 | art_short | ec | K |
| 164 | 229 | 0.620343222 | cat3 | from_5994 | to_5994 | art_short | ec | K |
| 165 | 229 | 0.608139925 | cat3 | from_5995 | to_5995 | art_short | ec | K |
| 167 | 229 | 0.693137623 | cat3 | from_5996 | to_5996 | art_short | ec | K |
| 197 | 229 | 0.760087557 | cat3 | from_5997 | to_5997 | art_short | ec | K |
| 217 | 229 | 0.765832513 | cat3 | from_5998 | to_5998 | art_short | ec | K |
| 220 | 229 | 0.623611785 | cat3 | from_5999 | to_5999 | art_untreat | ec | F |
| 53 | 230 | 0.684504099 | cat3 | from_6000 | to_6000 | art_long | ec | N |
| 168 | 230 | 0.823033653 | cat3 | from_6001 | to_6001 | art_short | ec | K |
| 169 | 230 | 0.609092002 | cat3 | from_6002 | to_6002 | art_short | ec | K |
| 179 | 230 | 0.711816912 | cat3 | from_6003 | to_6003 | art_short | ec | K |
| 180 | 230 | 0.89372958 | cat3 | from_6004 | to_6004 | art_short | ec | K |
| 187 | 230 | 0.886067567 | cat3 | from_6005 | to_6005 | art_short | ec | K |
| 220 | 230 | 0.7439303 | cat3 | from_6006 | to_6006 | art_untreat | ec | F |
| 1 | 231 | 0.652502128 | cat3 | from_6007 | to_6007 | art_long | ec | N |
| 34 | 231 | 0.783109348 | cat3 | from_6008 | to_6008 | art_long | ec | N |
| 65 | 231 | 0.647425264 | cat3 | from_6009 | to_6009 | art_long | ec | N |
| 113 | 231 | 0.72517669 | cat3 | from_6010 | to_6010 | art_long | ec | N |
| 127 | 231 | 0.667830674 | cat3 | from_6011 | to_6011 | art_long | ec | N |
| 153 | 231 | -0.631463818 | cat3 | from_6012 | to_6012 | art_long | ec | N |
| 207 | 231 | 0.834111941 | cat3 | from_6013 | to_6013 | art_short | ec | K |
| 227 | 231 | 0.630273076 | cat3 | from_6014 | to_6014 | ec | ec | P |
| 228 | 231 | 0.666684764 | cat3 | from_6015 | to_6015 | ec | ec | P |
| 139 | 232 | 0.652262923 | cat3 | from_6016 | to_6016 | art_long | ec | N |
| 170 | 232 | 0.692587865 | cat3 | from_6017 | to_6017 | art_short | ec | K |
| 171 | 232 | 0.704977237 | cat3 | from_6018 | to_6018 | art_short | ec | K |
| 178 | 232 | 0.662502057 | cat3 | from_6019 | to_6019 | art_short | ec | K |
| 184 | 232 | 0.623376836 | cat3 | from_6020 | to_6020 | art_short | ec | K |
| 223 | 232 | 0.730947784 | cat3 | from_6021 | to_6021 | art_untreat | ec | F |
| 226 | 232 | 0.731323999 | cat3 | from_6022 | to_6022 | art_untreat | ec | F |
| 6 | 233 | -0.649632568 | cat3 | from_6023 | to_6023 | art_long | ec | N |
| 20 | 233 | -0.607754814 | cat3 | from_6024 | to_6024 | art_long | ec | N |
| 99 | 233 | -0.605386919 | cat3 | from_6025 | to_6025 | art_long | ec | N |
| 105 | 233 | -0.645249511 | cat3 | from_6026 | to_6026 | art_long | ec | N |
| 131 | 233 | 0.648426191 | cat3 | from_6027 | to_6027 | art_long | ec | N |
| 148 | 233 | -0.646166722 | cat3 | from_6028 | to_6028 | art_long | ec | N |
| 152 | 233 | -0.642843062 | cat3 | from_6029 | to_6029 | art_long | ec | N |
| 153 | 233 | -0.661750321 | cat3 | from_6030 | to_6030 | art_long | ec | N |

| | | | | | | | | |
|---|---|---|---|---|---|---|---|---|
| 195 | 233 | 0.62936167 | cat3 | from_6031 | to_6031 | art_short | ec | K |
| 200 | 233 | 0.691430065 | cat3 | from_6032 | to_6032 | art_short | ec | K |
| 29 | 234 | 0.609818818 | cat3 | from_6033 | to_6033 | art_long | ec | N |
| 89 | 234 | 0.695489947 | cat3 | from_6034 | to_6034 | art_long | ec | N |
| 98 | 234 | 0.75220121 | cat3 | from_6035 | to_6035 | art_long | ec | N |
| 121 | 234 | 0.721788727 | cat3 | from_6036 | to_6036 | art_long | ec | N |
| 150 | 234 | 0.667790709 | cat3 | from_6037 | to_6037 | art_long | ec | N |
| 175 | 234 | 0.727478585 | cat3 | from_6038 | to_6038 | art_short | ec | K |
| 209 | 234 | 0.749063628 | cat3 | from_6039 | to_6039 | art_short | ec | K |
| 29 | 235 | 0.743990196 | cat3 | from_6040 | to_6040 | art_long | ec | N |
| 71 | 235 | 0.646474718 | cat3 | from_6041 | to_6041 | art_long | ec | N |
| 89 | 235 | 0.637245676 | cat3 | from_6042 | to_6042 | art_long | ec | N |
| 98 | 235 | 0.701573464 | cat3 | from_6043 | to_6043 | art_long | ec | N |
| 121 | 235 | 0.623459945 | cat3 | from_6044 | to_6044 | art_long | ec | N |
| 175 | 235 | 0.621329229 | cat3 | from_6045 | to_6045 | art_short | ec | K |
| 181 | 235 | 0.702376879 | cat3 | from_6046 | to_6046 | art_short | ec | K |
| 208 | 235 | 0.659884458 | cat3 | from_6047 | to_6047 | art_short | ec | K |
| 209 | 235 | 0.65415856 | cat3 | from_6048 | to_6048 | art_short | ec | K |
| 224 | 235 | 0.650858484 | cat3 | from_6049 | to_6049 | art_untreat | ec | F |
| 234 | 235 | 0.921764297 | cat3 | from_6050 | to_6050 | ec | ec | P |
| 152 | 236 | -0.621042509 | cat3 | from_6051 | to_6051 | art_long | ec | N |
| 153 | 236 | -0.733061809 | cat3 | from_6052 | to_6052 | art_long | ec | N |
| 227 | 236 | 0.76303312 | cat3 | from_6053 | to_6053 | ec | ec | P |
| 228 | 236 | 0.642592726 | cat3 | from_6054 | to_6054 | ec | ec | P |
| 232 | 236 | 0.702671773 | cat3 | from_6055 | to_6055 | ec | ec | P |
| 1 | 237 | 0.716541325 | cat3 | from_6056 | to_6056 | art_long | ec | N |
| 178 | 237 | 0.778916992 | cat3 | from_6057 | to_6057 | art_short | ec | K |
| 223 | 237 | 0.669645683 | cat3 | from_6058 | to_6058 | art_untreat | ec | F |
| 226 | 237 | 0.674116003 | cat3 | from_6059 | to_6059 | art_untreat | ec | F |
| 232 | 237 | 0.946090316 | cat3 | from_6060 | to_6060 | ec | ec | P |
| 236 | 237 | 0.711408275 | cat3 | from_6061 | to_6061 | ec | ec | P |
| 1 | 238 | 0.614314379 | cat3 | from_6062 | to_6062 | art_long | ec | N |
| 32 | 238 | -0.658388147 | cat3 | from_6063 | to_6063 | art_long | ec | N |
| 53 | 238 | 0.813819842 | cat3 | from_6064 | to_6064 | art_long | ec | N |
| 164 | 238 | 0.683343358 | cat3 | from_6065 | to_6065 | art_short | ec | K |
| 168 | 238 | 0.761078381 | cat3 | from_6066 | to_6066 | art_short | ec | K |
| 179 | 238 | 0.764361658 | cat3 | from_6067 | to_6067 | art_short | ec | K |
| 180 | 238 | 0.775274756 | cat3 | from_6068 | to_6068 | art_short | ec | K |
| 187 | 238 | 0.729686242 | cat3 | from_6069 | to_6069 | art_short | ec | K |
| 197 | 238 | 0.675797515 | cat3 | from_6070 | to_6070 | art_short | ec | K |
| 216 | 238 | 0.614553933 | cat3 | from_6071 | to_6071 | art_short | ec | K |
| 230 | 238 | 0.860707908 | cat3 | from_6072 | to_6072 | ec | ec | P |
| 32 | 239 | -0.603426486 | cat3 | from_6073 | to_6073 | art_long | ec | N |
| 43 | 239 | -0.602810538 | cat3 | from_6074 | to_6074 | art_long | ec | N |
| 56 | 239 | 0.621706671 | cat3 | from_6075 | to_6075 | art_long | ec | N |
| 107 | 239 | -0.605153369 | cat3 | from_6076 | to_6076 | art_long | ec | N |
| 149 | 239 | -0.609276791 | cat3 | from_6077 | to_6077 | art_long | ec | N |
| 182 | 239 | 0.70097646 | cat3 | from_6078 | to_6078 | art_short | ec | K |
| 193 | 239 | 0.680285218 | cat3 | from_6079 | to_6079 | art_short | ec | K |
| 199 | 239 | 0.695857865 | cat3 | from_6080 | to_6080 | art_short | ec | K |
| 201 | 239 | 0.615455184 | cat3 | from_6081 | to_6081 | art_short | ec | K |
| 217 | 239 | 0.656590627 | cat3 | from_6082 | to_6082 | art_short | ec | K |
| 222 | 239 | 0.918972009 | cat3 | from_6083 | to_6083 | art_untreat | ec | F |
| 54 | 240 | -0.612873002 | cat3 | from_6084 | to_6084 | art_long | ec | N |
| 68 | 240 | -0.670138916 | cat3 | from_6085 | to_6085 | art_long | ec | N |
| 81 | 240 | -0.645142085 | cat3 | from_6086 | to_6086 | art_long | ec | N |
| 83 | 240 | -0.6485928 | cat3 | from_6087 | to_6087 | art_long | ec | N |
| 120 | 240 | -0.60589989 | cat3 | from_6088 | to_6088 | art_long | ec | N |
| 134 | 240 | 0.744697548 | cat3 | from_6089 | to_6089 | art_long | ec | N |
| 138 | 240 | 0.61884821 | cat3 | from_6090 | to_6090 | art_long | ec | N |
| 140 | 240 | -0.635113778 | cat3 | from_6091 | to_6091 | art_long | ec | N |
| 154 | 240 | 0.612699197 | cat3 | from_6092 | to_6092 | art_short | ec | K |
| 166 | 240 | 0.713471483 | cat3 | from_6093 | to_6093 | art_short | ec | K |
| 170 | 240 | 0.615901265 | cat3 | from_6094 | to_6094 | art_short | ec | K |
| 171 | 240 | 0.620129956 | cat3 | from_6095 | to_6095 | art_short | ec | K |
| 177 | 240 | 0.754973581 | cat3 | from_6096 | to_6096 | art_short | ec | K |
| 232 | 240 | 0.652717498 | cat3 | from_6097 | to_6097 | ec | ec | P |
| 237 | 240 | 0.648805784 | cat3 | from_6098 | to_6098 | ec | ec | P |
| 48 | 241 | 0.60835453 | cat3 | from_6099 | to_6099 | art_long | ec | N |
| 68 | 241 | -0.655333666 | cat3 | from_6100 | to_6100 | art_long | ec | N |
| 91 | 241 | -0.679281464 | cat3 | from_6101 | to_6101 | art_long | ec | N |
| 104 | 241 | 0.62315695 | cat3 | from_6102 | to_6102 | art_long | ec | N |
| 134 | 241 | 0.731836336 | cat3 | from_6103 | to_6103 | art_long | ec | N |
| 138 | 241 | 0.65750387 | cat3 | from_6104 | to_6104 | art_long | ec | N |
| 139 | 241 | 0.628933162 | cat3 | from_6105 | to_6105 | art_long | ec | N |
| 143 | 241 | 0.642109044 | cat3 | from_6106 | to_6106 | art_long | ec | N |
| 154 | 241 | 0.71960511 | cat3 | from_6107 | to_6107 | art_short | ec | K |
| 166 | 241 | 0.743868481 | cat3 | from_6108 | to_6108 | art_short | ec | K |
| 170 | 241 | 0.693406417 | cat3 | from_6109 | to_6109 | art_short | ec | K |
| 171 | 241 | 0.666091171 | cat3 | from_6110 | to_6110 | art_short | ec | K |
| 204 | 241 | 0.632454261 | cat3 | from_6111 | to_6111 | art_short | ec | K |
| 233 | 241 | 0.69830108 | cat3 | from_6112 | to_6112 | ec | ec | P |
| 240 | 241 | 0.902028789 | cat3 | from_6113 | to_6113 | ec | ec | P |
| 156 | 242 | 0.660047904 | cat3 | from_6114 | to_6114 | art_short | ec | K |
| 157 | 242 | 0.672068613 | cat3 | from_6115 | to_6115 | art_short | ec | K |
| 158 | 242 | 0.651212585 | cat3 | from_6116 | to_6116 | art_short | ec | K |
| 198 | 242 | 0.818832002 | cat3 | from_6117 | to_6117 | art_short | ec | K |
| 201 | 242 | 0.679453966 | cat3 | from_6118 | to_6118 | art_short | ec | K |
| 217 | 242 | 0.631370874 | cat3 | from_6119 | to_6119 | art_short | ec | K |
| 227 | 242 | 0.688517216 | cat3 | from_6120 | to_6120 | ec | ec | P |
| 229 | 242 | 0.612548957 | cat3 | from_6121 | to_6121 | ec | ec | P |
| 231 | 242 | 0.648746519 | cat3 | from_6122 | to_6122 | ec | ec | P |
| 236 | 242 | 0.617116387 | cat3 | from_6123 | to_6123 | ec | ec | P |
| 113 | 243 | 0.852146825 | cat3 | from_6124 | to_6124 | art_long | ec | N |
| 152 | 243 | -0.66249897 | cat3 | from_6125 | to_6125 | art_long | ec | N |
| 153 | 243 | -0.697102717 | cat3 | from_6126 | to_6126 | art_long | ec | N |
| 207 | 243 | 0.752307378 | cat3 | from_6127 | to_6127 | art_short | ec | K |
| 227 | 243 | 0.845772984 | cat3 | from_6128 | to_6128 | ec | ec | P |
| 228 | 243 | 0.976199236 | cat3 | from_6129 | to_6129 | ec | ec | P |
| 231 | 243 | 0.793825872 | cat3 | from_6130 | to_6130 | ec | ec | P |
| 236 | 243 | 0.633782217 | cat3 | from_6131 | to_6131 | ec | ec | P |
| 242 | 244 | 0.690274184 | cat3 | from_6132 | to_6132 | ec | ec | P |
| 81 | 245 | 0.697186669 | cat3 | from_6133 | to_6133 | art_long | ec | N |
| 134 | 245 | -0.608721969 | cat3 | from_6134 | to_6134 | art_long | ec | N |

| | | | | | | | | |
|---|---|---|---|---|---|---|---|---|
| 145 | 245 | 0.670540922 | cat3 | from_6135 | to_6135 | art_long | ec | N |
| 206 | 245 | 0.629587523 | cat3 | from_6136 | to_6136 | art_short | ec | K |
| 219 | 245 | 0.775212251 | cat3 | from_6137 | to_6137 | art_untreat | ec | F |
| 225 | 245 | 0.661150163 | cat3 | from_6138 | to_6138 | art_untreat | ec | F |
| 229 | 245 | 0.63160043 | cat3 | from_6139 | to_6139 | ec | ec | P |
| 242 | 245 | 0.690755393 | cat3 | from_6140 | to_6140 | ec | ec | P |
| 49 | 246 | -0.647821482 | cat3 | from_6141 | to_6141 | art_long | ec | N |
| 51 | 246 | -0.714565394 | cat3 | from_6142 | to_6142 | art_long | ec | N |
| 63 | 246 | -0.629187269 | cat3 | from_6143 | to_6143 | art_long | ec | N |
| 70 | 246 | -0.720369015 | cat3 | from_6144 | to_6144 | art_long | ec | N |
| 75 | 246 | -0.683388118 | cat3 | from_6145 | to_6145 | art_long | ec | N |
| 84 | 246 | -0.656132416 | cat3 | from_6146 | to_6146 | art_long | ec | N |
| 146 | 246 | -0.604905161 | cat3 | from_6147 | to_6147 | art_long | ec | N |
| 191 | 246 | 0.634698126 | cat3 | from_6148 | to_6148 | art_short | ec | K |
| 195 | 246 | 0.710164856 | cat3 | from_6149 | to_6149 | art_short | ec | K |
| 200 | 246 | 0.675505413 | cat3 | from_6150 | to_6150 | art_short | ec | K |
| 203 | 246 | 0.630498731 | cat3 | from_6151 | to_6151 | art_short | ec | K |
| 204 | 246 | 0.70241703 | cat3 | from_6152 | to_6152 | art_short | ec | K |
| 223 | 246 | 0.65751448 | cat3 | from_6153 | to_6153 | art_untreat | ec | F |
| 225 | 246 | 0.659599964 | cat3 | from_6154 | to_6154 | art_untreat | ec | F |
| 12 | 247 | 0.854223767 | cat3 | from_6155 | to_6155 | art_long | ec | N |
| 13 | 247 | 0.662915381 | cat3 | from_6156 | to_6156 | art_long | ec | N |
| 19 | 247 | 0.845606658 | cat3 | from_6157 | to_6157 | art_long | ec | N |
| 40 | 247 | 0.718720071 | cat3 | from_6158 | to_6158 | art_long | ec | N |
| 79 | 247 | 0.862546225 | cat3 | from_6159 | to_6159 | art_long | ec | N |
| 85 | 247 | -0.655127365 | cat3 | from_6160 | to_6160 | art_long | ec | N |
| 92 | 247 | -0.646905796 | cat3 | from_6161 | to_6161 | art_long | ec | N |
| 93 | 247 | 0.72118062 | cat3 | from_6162 | to_6162 | art_long | ec | N |
| 97 | 247 | -0.614355304 | cat3 | from_6163 | to_6163 | art_long | ec | N |
| 103 | 247 | -0.638677138 | cat3 | from_6164 | to_6164 | art_long | ec | N |
| 114 | 247 | 0.631619244 | cat3 | from_6165 | to_6165 | art_long | ec | N |
| 115 | 247 | -0.6809424 | cat3 | from_6166 | to_6166 | art_long | ec | N |
| 117 | 247 | -0.630015219 | cat3 | from_6167 | to_6167 | art_long | ec | N |
| 133 | 247 | -0.71661914 | cat3 | from_6168 | to_6168 | art_long | ec | N |
| 145 | 247 | 0.622315507 | cat3 | from_6169 | to_6169 | art_long | ec | N |
| 147 | 247 | -0.614303962 | cat3 | from_6170 | to_6170 | art_long | ec | N |
| 163 | 247 | 0.622912588 | cat3 | from_6171 | to_6171 | art_short | ec | K |
| 167 | 247 | 0.887803493 | cat3 | from_6172 | to_6172 | art_short | ec | K |
| 186 | 247 | 0.655134788 | cat3 | from_6173 | to_6173 | art_short | ec | K |
| 216 | 247 | 0.622949016 | cat3 | from_6174 | to_6174 | art_short | ec | K |
| 229 | 247 | 0.698071016 | cat3 | from_6175 | to_6175 | ec | ec | P |
| 8 | 248 | 0.604705106 | cat3 | from_6176 | to_6176 | art_long | ec | N |
| 50 | 248 | 0.644805771 | cat3 | from_6177 | to_6177 | art_long | ec | N |
| 61 | 248 | 0.83320993 | cat3 | from_6178 | to_6178 | art_long | ec | N |
| 93 | 248 | 0.715000826 | cat3 | from_6179 | to_6179 | art_long | ec | N |
| 99 | 248 | 0.6633405 | cat3 | from_6180 | to_6180 | art_long | ec | N |
| 105 | 248 | 0.741885565 | cat3 | from_6181 | to_6181 | art_long | ec | N |
| 145 | 248 | 0.666595336 | cat3 | from_6182 | to_6182 | art_long | ec | N |
| 148 | 248 | 0.774836712 | cat3 | from_6183 | to_6183 | art_long | ec | N |
| 151 | 248 | 0.660698547 | cat3 | from_6184 | to_6184 | art_long | ec | N |
| 152 | 248 | 0.890802688 | cat3 | from_6185 | to_6185 | art_long | ec | N |
| 153 | 248 | 0.870353617 | cat3 | from_6186 | to_6186 | art_long | ec | N |
| 157 | 248 | 0.675659336 | cat3 | from_6187 | to_6187 | art_short | ec | K |
| 176 | 248 | 0.632951482 | cat3 | from_6188 | to_6188 | art_short | ec | K |
| 183 | 248 | 0.777872382 | cat3 | from_6189 | to_6189 | art_short | ec | K |
| 2 | 6 | -0.607424959 | cat4 | from_6190 | to_6190 | art_long | art_long | M |
| 4 | 6 | 0.650799702 | cat4 | from_6191 | to_6191 | art_long | art_long | M |
| 4 | 7 | 0.927566449 | cat4 | from_6192 | to_6192 | art_long | art_long | M |
| 6 | 8 | 0.66345499 | cat4 | from_6193 | to_6193 | art_long | art_long | M |
| 6 | 9 | 0.621412115 | cat4 | from_6194 | to_6194 | art_long | art_long | M |
| 8 | 9 | 0.894662737 | cat4 | from_6195 | to_6195 | art_long | art_long | M |
| 6 | 10 | 0.616361137 | cat4 | from_6196 | to_6196 | art_long | art_long | M |
| 8 | 10 | 0.876816306 | cat4 | from_6197 | to_6197 | art_long | art_long | M |
| 9 | 10 | 0.99681237 | cat4 | from_6198 | to_6198 | art_long | art_long | M |
| 4 | 11 | 0.786108061 | cat4 | from_6199 | to_6199 | art_long | art_long | M |
| 7 | 11 | 0.751757573 | cat4 | from_6200 | to_6200 | art_long | art_long | M |
| 12 | 13 | 0.779897947 | cat4 | from_6201 | to_6201 | art_long | art_long | M |
| 5 | 15 | 0.691862987 | cat4 | from_6202 | to_6202 | art_long | art_long | M |
| 12 | 15 | 0.651007882 | cat4 | from_6203 | to_6203 | art_long | art_long | M |
| 13 | 15 | 0.845217582 | cat4 | from_6204 | to_6204 | art_long | art_long | M |
| 2 | 16 | 0.611189345 | cat4 | from_6205 | to_6205 | art_long | art_long | M |
| 12 | 16 | 0.620196432 | cat4 | from_6206 | to_6206 | art_long | art_long | M |
| 15 | 16 | 0.603310551 | cat4 | from_6207 | to_6207 | art_long | art_long | M |
| 5 | 18 | 0.629641413 | cat4 | from_6208 | to_6208 | art_long | art_long | M |
| 12 | 18 | 0.809992501 | cat4 | from_6209 | to_6209 | art_long | art_long | M |
| 13 | 18 | 0.667480701 | cat4 | from_6210 | to_6210 | art_long | art_long | M |
| 15 | 18 | 0.796161832 | cat4 | from_6211 | to_6211 | art_long | art_long | M |
| 16 | 18 | 0.763142167 | cat4 | from_6212 | to_6212 | art_long | art_long | M |
| 12 | 19 | 0.952752224 | cat4 | from_6213 | to_6213 | art_long | art_long | M |
| 13 | 19 | 0.826208252 | cat4 | from_6214 | to_6214 | art_long | art_long | M |
| 15 | 19 | 0.618490822 | cat4 | from_6215 | to_6215 | art_long | art_long | M |
| 16 | 19 | 0.624661267 | cat4 | from_6216 | to_6216 | art_long | art_long | M |
| 18 | 19 | 0.77197214 | cat4 | from_6217 | to_6217 | art_long | art_long | M |
| 13 | 20 | 0.603490318 | cat4 | from_6218 | to_6218 | art_long | art_long | M |
| 16 | 21 | 0.593827193 | cat4 | from_6219 | to_6219 | art_long | art_long | M |
| 20 | 21 | 0.863215177 | cat4 | from_6220 | to_6220 | art_long | art_long | M |
| 9 | 22 | 0.698984814 | cat4 | from_6221 | to_6221 | art_long | art_long | M |
| 10 | 22 | 0.715661295 | cat4 | from_6222 | to_6222 | art_long | art_long | M |
| 11 | 22 | 0.644139366 | cat4 | from_6223 | to_6223 | art_long | art_long | M |
| 8 | 23 | 0.689325892 | cat4 | from_6224 | to_6224 | art_long | art_long | M |
| 9 | 23 | 0.916270666 | cat4 | from_6225 | to_6225 | art_long | art_long | M |
| 10 | 23 | 0.9283552 | cat4 | from_6226 | to_6226 | art_long | art_long | M |
| 11 | 23 | 0.611725113 | cat4 | from_6227 | to_6227 | art_long | art_long | M |
| 22 | 23 | 0.830731465 | cat4 | from_6228 | to_6228 | art_long | art_long | M |
| 14 | 24 | 0.653381001 | cat4 | from_6229 | to_6229 | art_long | art_long | M |
| 14 | 25 | 0.742957444 | cat4 | from_6230 | to_6230 | art_long | art_long | M |
| 24 | 25 | 0.598070921 | cat4 | from_6231 | to_6231 | art_long | art_long | M |
| 20 | 27 | -0.618066175 | cat4 | from_6232 | to_6232 | art_long | art_long | M |
| 22 | 27 | 0.742188931 | cat4 | from_6233 | to_6233 | art_long | art_long | M |
| 14 | 28 | 0.790060333 | cat4 | from_6234 | to_6234 | art_long | art_long | M |
| 24 | 28 | 0.940161714 | cat4 | from_6235 | to_6235 | art_long | art_long | M |
| 25 | 28 | 0.671911847 | cat4 | from_6236 | to_6236 | art_long | art_long | M |
| 26 | 28 | 0.577305316 | cat4 | from_6237 | to_6237 | art_long | art_long | M |
| 3 | 29 | 0.606335785 | cat4 | from_6238 | to_6238 | art_long | art_long | M |

| | | | | | | | | |
|---|---|---|---|---|---|---|---|---|
| 2 | 31 | -0.617112395 | cat4 | from_6239 | to_6239 | art_long | art_long | M |
| 14 | 31 | 0.622823134 | cat4 | from_6240 | to_6240 | art_long | art_long | M |
| 25 | 31 | 0.888587576 | cat4 | from_6241 | to_6241 | art_long | art_long | M |
| 24 | 32 | 0.690186053 | cat4 | from_6242 | to_6242 | art_long | art_long | M |
| 28 | 32 | 0.622959606 | cat4 | from_6243 | to_6243 | art_long | art_long | M |
| 2 | 33 | -0.582563194 | cat4 | from_6244 | to_6244 | art_long | art_long | M |
| 14 | 33 | 0.626155453 | cat4 | from_6245 | to_6245 | art_long | art_long | M |
| 25 | 33 | 0.914009254 | cat4 | from_6246 | to_6246 | art_long | art_long | M |
| 31 | 33 | 0.996048471 | cat4 | from_6247 | to_6247 | art_long | art_long | M |
| 4 | 34 | 0.705596366 | cat4 | from_6248 | to_6248 | art_long | art_long | M |
| 5 | 34 | 0.675433886 | cat4 | from_6249 | to_6249 | art_long | art_long | M |
| 7 | 34 | 0.621241452 | cat4 | from_6250 | to_6250 | art_long | art_long | M |
| 17 | 35 | 0.710597612 | cat4 | from_6251 | to_6251 | art_long | art_long | M |
| 17 | 36 | 0.966072086 | cat4 | from_6252 | to_6252 | art_long | art_long | M |
| 35 | 36 | 0.587111838 | cat4 | from_6253 | to_6253 | art_long | art_long | M |
| 26 | 37 | 0.960565026 | cat4 | from_6254 | to_6254 | art_long | art_long | M |
| 17 | 38 | 0.753199655 | cat4 | from_6255 | to_6255 | art_long | art_long | M |
| 35 | 38 | 0.980058532 | cat4 | from_6256 | to_6256 | art_long | art_long | M |
| 36 | 38 | 0.66441904 | cat4 | from_6257 | to_6257 | art_long | art_long | M |
| 12 | 39 | 0.7410295 | cat4 | from_6258 | to_6258 | art_long | art_long | M |
| 13 | 39 | 0.632829509 | cat4 | from_6259 | to_6259 | art_long | art_long | M |
| 15 | 39 | 0.593552814 | cat4 | from_6260 | to_6260 | art_long | art_long | M |
| 16 | 39 | 0.914776947 | cat4 | from_6261 | to_6261 | art_long | art_long | M |
| 18 | 39 | 0.832706241 | cat4 | from_6262 | to_6262 | art_long | art_long | M |
| 19 | 39 | 0.767938847 | cat4 | from_6263 | to_6263 | art_long | art_long | M |
| 20 | 39 | 0.640664338 | cat4 | from_6264 | to_6264 | art_long | art_long | M |
| 21 | 39 | 0.632690042 | cat4 | from_6265 | to_6265 | art_long | art_long | M |
| 12 | 40 | 0.798870575 | cat4 | from_6266 | to_6266 | art_long | art_long | M |
| 13 | 40 | 0.687011701 | cat4 | from_6267 | to_6267 | art_long | art_long | M |
| 16 | 40 | 0.793852907 | cat4 | from_6268 | to_6268 | art_long | art_long | M |
| 18 | 40 | 0.64349749 | cat4 | from_6269 | to_6269 | art_long | art_long | M |
| 19 | 40 | 0.880204717 | cat4 | from_6270 | to_6270 | art_long | art_long | M |
| 39 | 40 | 0.861907174 | cat4 | from_6271 | to_6271 | art_long | art_long | M |
| 29 | 41 | 0.739385858 | cat4 | from_6272 | to_6272 | art_long | art_long | M |
| 4 | 42 | 0.578873537 | cat4 | from_6273 | to_6273 | art_long | art_long | M |
| 21 | 42 | 0.632662979 | cat4 | from_6274 | to_6274 | art_long | art_long | M |
| 6 | 43 | 0.597283289 | cat4 | from_6275 | to_6275 | art_long | art_long | M |
| 36 | 43 | 0.606226945 | cat4 | from_6276 | to_6276 | art_long | art_long | M |
| 42 | 43 | 0.625171034 | cat4 | from_6277 | to_6277 | art_long | art_long | M |
| 11 | 44 | 0.643869338 | cat4 | from_6278 | to_6278 | art_long | art_long | M |
| 35 | 44 | 0.651830422 | cat4 | from_6279 | to_6279 | art_long | art_long | M |
| 38 | 44 | 0.644332774 | cat4 | from_6280 | to_6280 | art_long | art_long | M |
| 42 | 44 | 0.663770992 | cat4 | from_6281 | to_6281 | art_long | art_long | M |
| 4 | 45 | 0.597170361 | cat4 | from_6282 | to_6282 | art_long | art_long | M |
| 7 | 45 | 0.601659532 | cat4 | from_6283 | to_6283 | art_long | art_long | M |
| 17 | 45 | 0.589857503 | cat4 | from_6284 | to_6284 | art_long | art_long | M |
| 42 | 45 | 0.943530506 | cat4 | from_6285 | to_6285 | art_long | art_long | M |
| 43 | 45 | 0.697403085 | cat4 | from_6286 | to_6286 | art_long | art_long | M |
| 44 | 45 | 0.641863599 | cat4 | from_6287 | to_6287 | art_long | art_long | M |
| 11 | 46 | 0.647702236 | cat4 | from_6288 | to_6288 | art_long | art_long | M |
| 35 | 46 | 0.674224454 | cat4 | from_6289 | to_6289 | art_long | art_long | M |
| 38 | 46 | 0.64669456 | cat4 | from_6290 | to_6290 | art_long | art_long | M |
| 42 | 46 | 0.662136034 | cat4 | from_6291 | to_6291 | art_long | art_long | M |
| 44 | 46 | 0.987892294 | cat4 | from_6292 | to_6292 | art_long | art_long | M |
| 45 | 46 | 0.670549025 | cat4 | from_6293 | to_6293 | art_long | art_long | M |
| 28 | 47 | 0.599424394 | cat4 | from_6294 | to_6294 | art_long | art_long | M |
| 30 | 47 | 0.851052788 | cat4 | from_6295 | to_6295 | art_long | art_long | M |
| 17 | 48 | 0.701346096 | cat4 | from_6296 | to_6296 | art_long | art_long | M |
| 31 | 48 | -0.669532732 | cat4 | from_6297 | to_6297 | art_long | art_long | M |
| 33 | 48 | -0.659051325 | cat4 | from_6298 | to_6298 | art_long | art_long | M |
| 35 | 48 | 0.927081171 | cat4 | from_6299 | to_6299 | art_long | art_long | M |
| 36 | 48 | 0.628842665 | cat4 | from_6300 | to_6300 | art_long | art_long | M |
| 38 | 48 | 0.891550108 | cat4 | from_6301 | to_6301 | art_long | art_long | M |
| 44 | 48 | 0.594295095 | cat4 | from_6302 | to_6302 | art_long | art_long | M |
| 46 | 48 | 0.631013835 | cat4 | from_6303 | to_6303 | art_long | art_long | M |
| 28 | 50 | -0.577860117 | cat4 | from_6304 | to_6304 | art_long | art_long | M |
| 35 | 50 | 0.694069195 | cat4 | from_6305 | to_6305 | art_long | art_long | M |
| 38 | 50 | 0.775696356 | cat4 | from_6306 | to_6306 | art_long | art_long | M |
| 48 | 50 | 0.627861399 | cat4 | from_6307 | to_6307 | art_long | art_long | M |
| 11 | 52 | 0.584495127 | cat4 | from_6308 | to_6308 | art_long | art_long | M |
| 35 | 52 | 0.586666931 | cat4 | from_6309 | to_6309 | art_long | art_long | M |
| 38 | 52 | 0.594283871 | cat4 | from_6310 | to_6310 | art_long | art_long | M |
| 42 | 52 | 0.59163696 | cat4 | from_6311 | to_6311 | art_long | art_long | M |
| 44 | 52 | 0.865510076 | cat4 | from_6312 | to_6312 | art_long | art_long | M |
| 46 | 52 | 0.863200838 | cat4 | from_6313 | to_6313 | art_long | art_long | M |
| 2 | 53 | 0.738803943 | cat4 | from_6314 | to_6314 | art_long | art_long | M |
| 16 | 53 | 0.758902943 | cat4 | from_6315 | to_6315 | art_long | art_long | M |
| 18 | 53 | 0.580450597 | cat4 | from_6316 | to_6316 | art_long | art_long | M |
| 39 | 53 | 0.695003287 | cat4 | from_6317 | to_6317 | art_long | art_long | M |
| 25 | 54 | 0.642124357 | cat4 | from_6318 | to_6318 | art_long | art_long | M |
| 42 | 54 | -0.614195801 | cat4 | from_6319 | to_6319 | art_long | art_long | M |
| 24 | 55 | 0.613813229 | cat4 | from_6320 | to_6320 | art_long | art_long | M |
| 27 | 55 | 0.608077034 | cat4 | from_6321 | to_6321 | art_long | art_long | M |
| 28 | 55 | 0.659922647 | cat4 | from_6322 | to_6322 | art_long | art_long | M |
| 30 | 55 | 0.849653088 | cat4 | from_6323 | to_6323 | art_long | art_long | M |
| 47 | 55 | 0.956728527 | cat4 | from_6324 | to_6324 | art_long | art_long | M |
| 22 | 56 | 0.702827373 | cat4 | from_6325 | to_6325 | art_long | art_long | M |
| 22 | 57 | 0.632111176 | cat4 | from_6326 | to_6326 | art_long | art_long | M |
| 56 | 57 | 0.793045634 | cat4 | from_6327 | to_6327 | art_long | art_long | M |
| 14 | 58 | 0.703606748 | cat4 | from_6328 | to_6328 | art_long | art_long | M |
| 30 | 58 | 0.89744481 | cat4 | from_6329 | to_6329 | art_long | art_long | M |
| 47 | 58 | 0.694580115 | cat4 | from_6330 | to_6330 | art_long | art_long | M |
| 54 | 58 | 0.631819504 | cat4 | from_6331 | to_6331 | art_long | art_long | M |
| 55 | 58 | 0.77671212 | cat4 | from_6332 | to_6332 | art_long | art_long | M |
| 14 | 59 | 0.973523969 | cat4 | from_6333 | to_6333 | art_long | art_long | M |
| 24 | 59 | 0.797302554 | cat4 | from_6334 | to_6334 | art_long | art_long | M |
| 25 | 59 | 0.758449355 | cat4 | from_6335 | to_6335 | art_long | art_long | M |
| 26 | 59 | 0.600303281 | cat4 | from_6336 | to_6336 | art_long | art_long | M |
| 28 | 59 | 0.907600852 | cat4 | from_6337 | to_6337 | art_long | art_long | M |
| 31 | 59 | 0.588560142 | cat4 | from_6338 | to_6338 | art_long | art_long | M |
| 33 | 59 | 0.61046551 | cat4 | from_6339 | to_6339 | art_long | art_long | M |
| 58 | 59 | 0.690490783 | cat4 | from_6340 | to_6340 | art_long | art_long | M |
| 11 | 60 | 0.603438329 | cat4 | from_6341 | to_6341 | art_long | art_long | M |
| 38 | 60 | 0.620859369 | cat4 | from_6342 | to_6342 | art_long | art_long | M |

| | | | | | | | | |
|---|---|---|---|---|---|---|---|---|
| 43 | 60 | 0.624054552 | cat4 | from_6343 | to_6343 | art_long | art_long | M |
| 44 | 60 | 0.896483765 | cat4 | from_6344 | to_6344 | art_long | art_long | M |
| 46 | 60 | 0.856052013 | cat4 | from_6345 | to_6345 | art_long | art_long | M |
| 52 | 60 | 0.837643189 | cat4 | from_6346 | to_6346 | art_long | art_long | M |
| 43 | 61 | 0.615880866 | cat4 | from_6347 | to_6347 | art_long | art_long | M |
| 44 | 61 | 0.808422162 | cat4 | from_6348 | to_6348 | art_long | art_long | M |
| 46 | 61 | 0.803101221 | cat4 | from_6349 | to_6349 | art_long | art_long | M |
| 50 | 61 | 0.61929135 | cat4 | from_6350 | to_6350 | art_long | art_long | M |
| 52 | 61 | 0.623560936 | cat4 | from_6351 | to_6351 | art_long | art_long | M |
| 60 | 61 | 0.808622843 | cat4 | from_6352 | to_6352 | art_long | art_long | M |
| 4 | 62 | 0.61042263 | cat4 | from_6353 | to_6353 | art_long | art_long | M |
| 7 | 62 | 0.595707088 | cat4 | from_6354 | to_6354 | art_long | art_long | M |
| 17 | 62 | 0.604402361 | cat4 | from_6355 | to_6355 | art_long | art_long | M |
| 42 | 62 | 0.957262617 | cat4 | from_6356 | to_6356 | art_long | art_long | M |
| 43 | 62 | 0.756331928 | cat4 | from_6357 | to_6357 | art_long | art_long | M |
| 44 | 62 | 0.672386083 | cat4 | from_6358 | to_6358 | art_long | art_long | M |
| 45 | 62 | 0.979816664 | cat4 | from_6359 | to_6359 | art_long | art_long | M |
| 46 | 62 | 0.670434325 | cat4 | from_6360 | to_6360 | art_long | art_long | M |
| 52 | 62 | 0.578332511 | cat4 | from_6361 | to_6361 | art_long | art_long | M |
| 5 | 63 | 0.927676847 | cat4 | from_6362 | to_6362 | art_long | art_long | M |
| 13 | 63 | 0.661562839 | cat4 | from_6363 | to_6363 | art_long | art_long | M |
| 15 | 63 | 0.751057476 | cat4 | from_6364 | to_6364 | art_long | art_long | M |
| 18 | 63 | 0.696957279 | cat4 | from_6365 | to_6365 | art_long | art_long | M |
| 24 | 64 | 0.626540229 | cat4 | from_6366 | to_6366 | art_long | art_long | M |
| 27 | 64 | 0.733053443 | cat4 | from_6367 | to_6367 | art_long | art_long | M |
| 28 | 64 | 0.660831713 | cat4 | from_6368 | to_6368 | art_long | art_long | M |
| 34 | 64 | 0.614748221 | cat4 | from_6369 | to_6369 | art_long | art_long | M |
| 50 | 64 | -0.672795254 | cat4 | from_6370 | to_6370 | art_long | art_long | M |
| 22 | 65 | 0.634568631 | cat4 | from_6371 | to_6371 | art_long | art_long | M |
| 23 | 65 | 0.637759262 | cat4 | from_6372 | to_6372 | art_long | art_long | M |
| 34 | 65 | 0.813957818 | cat4 | from_6373 | to_6373 | art_long | art_long | M |
| 49 | 65 | 0.597694257 | cat4 | from_6374 | to_6374 | art_long | art_long | M |
| 64 | 65 | 0.629072705 | cat4 | from_6375 | to_6375 | art_long | art_long | M |
| 29 | 66 | 0.711767029 | cat4 | from_6376 | to_6376 | art_long | art_long | M |
| 41 | 66 | 0.652842836 | cat4 | from_6377 | to_6377 | art_long | art_long | M |
| 29 | 67 | 0.804263855 | cat4 | from_6378 | to_6378 | art_long | art_long | M |
| 41 | 67 | 0.674748904 | cat4 | from_6379 | to_6379 | art_long | art_long | M |
| 66 | 67 | 0.950952848 | cat4 | from_6380 | to_6380 | art_long | art_long | M |
| 12 | 68 | 0.626804943 | cat4 | from_6381 | to_6381 | art_long | art_long | M |
| 13 | 68 | 0.7597457 | cat4 | from_6382 | to_6382 | art_long | art_long | M |
| 15 | 68 | 0.606376723 | cat4 | from_6383 | to_6383 | art_long | art_long | M |
| 18 | 68 | 0.614570238 | cat4 | from_6384 | to_6384 | art_long | art_long | M |
| 19 | 68 | 0.666251881 | cat4 | from_6385 | to_6385 | art_long | art_long | M |
| 25 | 68 | 0.669171928 | cat4 | from_6386 | to_6386 | art_long | art_long | M |
| 31 | 68 | 0.677906701 | cat4 | from_6387 | to_6387 | art_long | art_long | M |
| 33 | 68 | 0.661683274 | cat4 | from_6388 | to_6388 | art_long | art_long | M |
| 48 | 68 | -0.622964371 | cat4 | from_6389 | to_6389 | art_long | art_long | M |
| 14 | 69 | 0.719875114 | cat4 | from_6390 | to_6390 | art_long | art_long | M |
| 30 | 69 | 0.895561569 | cat4 | from_6391 | to_6391 | art_long | art_long | M |
| 47 | 69 | 0.666035197 | cat4 | from_6392 | to_6392 | art_long | art_long | M |
| 54 | 69 | 0.608967082 | cat4 | from_6393 | to_6393 | art_long | art_long | M |
| 55 | 69 | 0.731716065 | cat4 | from_6394 | to_6394 | art_long | art_long | M |
| 58 | 69 | 0.995245727 | cat4 | from_6395 | to_6395 | art_long | art_long | M |
| 59 | 69 | 0.687143108 | cat4 | from_6396 | to_6396 | art_long | art_long | M |
| 49 | 70 | 0.751467416 | cat4 | from_6397 | to_6397 | art_long | art_long | M |
| 57 | 70 | 0.618892778 | cat4 | from_6398 | to_6398 | art_long | art_long | M |
| 3 | 71 | 0.685973706 | cat4 | from_6399 | to_6399 | art_long | art_long | M |
| 29 | 71 | 0.97081858 | cat4 | from_6400 | to_6400 | art_long | art_long | M |
| 41 | 71 | 0.771257463 | cat4 | from_6401 | to_6401 | art_long | art_long | M |
| 66 | 71 | 0.73970147 | cat4 | from_6402 | to_6402 | art_long | art_long | M |
| 67 | 71 | 0.830764997 | cat4 | from_6403 | to_6403 | art_long | art_long | M |
| 30 | 72 | 0.907846023 | cat4 | from_6404 | to_6404 | art_long | art_long | M |
| 33 | 72 | 0.580084375 | cat4 | from_6405 | to_6405 | art_long | art_long | M |
| 47 | 72 | 0.830653335 | cat4 | from_6406 | to_6406 | art_long | art_long | M |
| 55 | 72 | 0.755307275 | cat4 | from_6407 | to_6407 | art_long | art_long | M |
| 58 | 72 | 0.78084284 | cat4 | from_6408 | to_6408 | art_long | art_long | M |
| 69 | 72 | 0.794022727 | cat4 | from_6409 | to_6409 | art_long | art_long | M |
| 14 | 73 | 0.999019551 | cat4 | from_6410 | to_6410 | art_long | art_long | M |
| 24 | 73 | 0.677957114 | cat4 | from_6411 | to_6411 | art_long | art_long | M |
| 25 | 73 | 0.749314877 | cat4 | from_6412 | to_6412 | art_long | art_long | M |
| 28 | 73 | 0.804851868 | cat4 | from_6413 | to_6413 | art_long | art_long | M |
| 31 | 73 | 0.619681791 | cat4 | from_6414 | to_6414 | art_long | art_long | M |
| 33 | 73 | 0.625284784 | cat4 | from_6415 | to_6415 | art_long | art_long | M |
| 58 | 73 | 0.70332761 | cat4 | from_6416 | to_6416 | art_long | art_long | M |
| 59 | 73 | 0.979532458 | cat4 | from_6417 | to_6417 | art_long | art_long | M |
| 69 | 73 | 0.716094742 | cat4 | from_6418 | to_6418 | art_long | art_long | M |
| 14 | 74 | 0.887504722 | cat4 | from_6419 | to_6419 | art_long | art_long | M |
| 24 | 74 | 0.823057324 | cat4 | from_6420 | to_6420 | art_long | art_long | M |
| 25 | 74 | 0.711946762 | cat4 | from_6421 | to_6421 | art_long | art_long | M |
| 28 | 74 | 0.799294033 | cat4 | from_6422 | to_6422 | art_long | art_long | M |
| 32 | 74 | 0.608924761 | cat4 | from_6423 | to_6423 | art_long | art_long | M |
| 58 | 74 | 0.639748197 | cat4 | from_6424 | to_6424 | art_long | art_long | M |
| 59 | 74 | 0.908334613 | cat4 | from_6425 | to_6425 | art_long | art_long | M |
| 69 | 74 | 0.621047215 | cat4 | from_6426 | to_6426 | art_long | art_long | M |
| 73 | 74 | 0.902358685 | cat4 | from_6427 | to_6427 | art_long | art_long | M |
| 49 | 75 | 0.781566665 | cat4 | from_6428 | to_6428 | art_long | art_long | M |
| 56 | 75 | 0.639061771 | cat4 | from_6429 | to_6429 | art_long | art_long | M |
| 57 | 75 | 0.623849471 | cat4 | from_6430 | to_6430 | art_long | art_long | M |
| 70 | 75 | 0.899282837 | cat4 | from_6431 | to_6431 | art_long | art_long | M |
| 14 | 76 | 0.702903941 | cat4 | from_6432 | to_6432 | art_long | art_long | M |
| 24 | 76 | 0.922494931 | cat4 | from_6433 | to_6433 | art_long | art_long | M |
| 25 | 76 | 0.827367484 | cat4 | from_6434 | to_6434 | art_long | art_long | M |
| 28 | 76 | 0.910123913 | cat4 | from_6435 | to_6435 | art_long | art_long | M |
| 32 | 76 | 0.653351677 | cat4 | from_6436 | to_6436 | art_long | art_long | M |
| 33 | 76 | 0.612322405 | cat4 | from_6437 | to_6437 | art_long | art_long | M |
| 55 | 76 | 0.603378882 | cat4 | from_6438 | to_6438 | art_long | art_long | M |
| 59 | 76 | 0.815287047 | cat4 | from_6439 | to_6439 | art_long | art_long | M |
| 73 | 76 | 0.71923898 | cat4 | from_6440 | to_6440 | art_long | art_long | M |
| 74 | 76 | 0.802657387 | cat4 | from_6441 | to_6441 | art_long | art_long | M |
| 25 | 77 | 0.596907644 | cat4 | from_6442 | to_6442 | art_long | art_long | M |
| 30 | 77 | 0.639328702 | cat4 | from_6443 | to_6443 | art_long | art_long | M |
| 54 | 77 | 0.927174267 | cat4 | from_6444 | to_6444 | art_long | art_long | M |
| 55 | 77 | 0.647528761 | cat4 | from_6445 | to_6445 | art_long | art_long | M |
| 58 | 77 | 0.793116843 | cat4 | from_6446 | to_6446 | art_long | art_long | M |

| | | | | | | | | |
|---|---|---|---|---|---|---|---|---|
| 69 | 77 | 0.75751287 | cat4 | from_6447 | to_6447 | art_long | art_long | M |
| 74 | 77 | 0.636642852 | cat4 | from_6448 | to_6448 | art_long | art_long | M |
| 76 | 77 | 0.632056261 | cat4 | from_6449 | to_6449 | art_long | art_long | M |
| 14 | 78 | 0.777022108 | cat4 | from_6450 | to_6450 | art_long | art_long | M |
| 24 | 78 | 0.902831124 | cat4 | from_6451 | to_6451 | art_long | art_long | M |
| 25 | 78 | 0.648851128 | cat4 | from_6452 | to_6452 | art_long | art_long | M |
| 28 | 78 | 0.825713136 | cat4 | from_6453 | to_6453 | art_long | art_long | M |
| 32 | 78 | 0.641934217 | cat4 | from_6454 | to_6454 | art_long | art_long | M |
| 59 | 78 | 0.838560903 | cat4 | from_6455 | to_6455 | art_long | art_long | M |
| 73 | 78 | 0.794039899 | cat4 | from_6456 | to_6456 | art_long | art_long | M |
| 74 | 78 | 0.958663382 | cat4 | from_6457 | to_6457 | art_long | art_long | M |
| 76 | 78 | 0.868001164 | cat4 | from_6458 | to_6458 | art_long | art_long | M |
| 77 | 78 | 0.67240743 | cat4 | from_6459 | to_6459 | art_long | art_long | M |
| 12 | 79 | 0.968465794 | cat4 | from_6460 | to_6460 | art_long | art_long | M |
| 13 | 79 | 0.813351689 | cat4 | from_6461 | to_6461 | art_long | art_long | M |
| 15 | 79 | 0.606054801 | cat4 | from_6462 | to_6462 | art_long | art_long | M |
| 18 | 79 | 0.760576393 | cat4 | from_6463 | to_6463 | art_long | art_long | M |
| 19 | 79 | 0.977434834 | cat4 | from_6464 | to_6464 | art_long | art_long | M |
| 39 | 79 | 0.67959024 | cat4 | from_6465 | to_6465 | art_long | art_long | M |
| 40 | 79 | 0.779848792 | cat4 | from_6466 | to_6466 | art_long | art_long | M |
| 68 | 79 | 0.699673661 | cat4 | from_6467 | to_6467 | art_long | art_long | M |
| 24 | 80 | 0.594983855 | cat4 | from_6468 | to_6468 | art_long | art_long | M |
| 27 | 80 | 0.601603847 | cat4 | from_6469 | to_6469 | art_long | art_long | M |
| 30 | 80 | 0.620774398 | cat4 | from_6470 | to_6470 | art_long | art_long | M |
| 54 | 80 | 0.878889305 | cat4 | from_6471 | to_6471 | art_long | art_long | M |
| 55 | 80 | 0.707558436 | cat4 | from_6472 | to_6472 | art_long | art_long | M |
| 58 | 80 | 0.733415001 | cat4 | from_6473 | to_6473 | art_long | art_long | M |
| 69 | 80 | 0.684507622 | cat4 | from_6474 | to_6474 | art_long | art_long | M |
| 74 | 80 | 0.57989243 | cat4 | from_6475 | to_6475 | art_long | art_long | M |
| 76 | 80 | 0.663793017 | cat4 | from_6476 | to_6476 | art_long | art_long | M |
| 77 | 80 | 0.983017731 | cat4 | from_6477 | to_6477 | art_long | art_long | M |
| 78 | 80 | 0.660689948 | cat4 | from_6478 | to_6478 | art_long | art_long | M |
| 8 | 81 | 0.592891305 | cat4 | from_6479 | to_6479 | art_long | art_long | M |
| 25 | 81 | 0.681886039 | cat4 | from_6480 | to_6480 | art_long | art_long | M |
| 31 | 81 | 0.654849729 | cat4 | from_6481 | to_6481 | art_long | art_long | M |
| 33 | 81 | 0.643145308 | cat4 | from_6482 | to_6482 | art_long | art_long | M |
| 54 | 81 | 0.940175922 | cat4 | from_6483 | to_6483 | art_long | art_long | M |
| 58 | 81 | 0.669327802 | cat4 | from_6484 | to_6484 | art_long | art_long | M |
| 68 | 81 | 0.654640703 | cat4 | from_6485 | to_6485 | art_long | art_long | M |
| 69 | 81 | 0.677660434 | cat4 | from_6486 | to_6486 | art_long | art_long | M |
| 77 | 81 | 0.835242646 | cat4 | from_6487 | to_6487 | art_long | art_long | M |
| 80 | 81 | 0.747382643 | cat4 | from_6488 | to_6488 | art_long | art_long | M |
| 3 | 82 | 0.698451208 | cat4 | from_6489 | to_6489 | art_long | art_long | M |
| 29 | 82 | 0.714952246 | cat4 | from_6490 | to_6490 | art_long | art_long | M |
| 41 | 82 | 0.660121892 | cat4 | from_6491 | to_6491 | art_long | art_long | M |
| 66 | 82 | 0.662521876 | cat4 | from_6492 | to_6492 | art_long | art_long | M |
| 67 | 82 | 0.675084064 | cat4 | from_6493 | to_6493 | art_long | art_long | M |
| 71 | 82 | 0.78088338 | cat4 | from_6494 | to_6494 | art_long | art_long | M |
| 14 | 83 | 0.607510879 | cat4 | from_6495 | to_6495 | art_long | art_long | M |
| 25 | 83 | 0.65009382 | cat4 | from_6496 | to_6496 | art_long | art_long | M |
| 30 | 83 | 0.785542247 | cat4 | from_6497 | to_6497 | art_long | art_long | M |
| 31 | 83 | 0.60391324 | cat4 | from_6498 | to_6498 | art_long | art_long | M |
| 33 | 83 | 0.594287891 | cat4 | from_6499 | to_6499 | art_long | art_long | M |
| 54 | 83 | 0.794830961 | cat4 | from_6500 | to_6500 | art_long | art_long | M |
| 58 | 83 | 0.883586001 | cat4 | from_6501 | to_6501 | art_long | art_long | M |
| 68 | 83 | 0.584166507 | cat4 | from_6502 | to_6502 | art_long | art_long | M |
| 69 | 83 | 0.8995428 | cat4 | from_6503 | to_6503 | art_long | art_long | M |
| 72 | 83 | 0.749613839 | cat4 | from_6504 | to_6504 | art_long | art_long | M |
| 73 | 83 | 0.599296612 | cat4 | from_6505 | to_6505 | art_long | art_long | M |
| 77 | 83 | 0.840966331 | cat4 | from_6506 | to_6506 | art_long | art_long | M |
| 80 | 83 | 0.756197224 | cat4 | from_6507 | to_6507 | art_long | art_long | M |
| 81 | 83 | 0.891387316 | cat4 | from_6508 | to_6508 | art_long | art_long | M |
| 6 | 84 | 0.584266641 | cat4 | from_6509 | to_6509 | art_long | art_long | M |
| 8 | 84 | 0.709538346 | cat4 | from_6510 | to_6510 | art_long | art_long | M |
| 9 | 84 | 0.635309279 | cat4 | from_6511 | to_6511 | art_long | art_long | M |
| 10 | 84 | 0.637179583 | cat4 | from_6512 | to_6512 | art_long | art_long | M |
| 49 | 84 | 0.729869013 | cat4 | from_6513 | to_6513 | art_long | art_long | M |
| 70 | 84 | 0.720843184 | cat4 | from_6514 | to_6514 | art_long | art_long | M |
| 75 | 84 | 0.890229616 | cat4 | from_6515 | to_6515 | art_long | art_long | M |
| 34 | 85 | 0.729739966 | cat4 | from_6516 | to_6516 | art_long | art_long | M |
| 49 | 85 | 0.855025339 | cat4 | from_6517 | to_6517 | art_long | art_long | M |
| 54 | 85 | -0.631004546 | cat4 | from_6518 | to_6518 | art_long | art_long | M |
| 56 | 85 | 0.708692508 | cat4 | from_6519 | to_6519 | art_long | art_long | M |
| 65 | 85 | 0.627465535 | cat4 | from_6520 | to_6520 | art_long | art_long | M |
| 70 | 85 | 0.696763457 | cat4 | from_6521 | to_6521 | art_long | art_long | M |
| 75 | 85 | 0.684998715 | cat4 | from_6522 | to_6522 | art_long | art_long | M |
| 30 | 86 | 0.626696695 | cat4 | from_6523 | to_6523 | art_long | art_long | M |
| 32 | 86 | 0.590250168 | cat4 | from_6524 | to_6524 | art_long | art_long | M |
| 44 | 86 | 0.603746925 | cat4 | from_6525 | to_6525 | art_long | art_long | M |
| 46 | 86 | 0.626689735 | cat4 | from_6526 | to_6526 | art_long | art_long | M |
| 47 | 86 | 0.624290019 | cat4 | from_6527 | to_6527 | art_long | art_long | M |
| 55 | 86 | 0.597293069 | cat4 | from_6528 | to_6528 | art_long | art_long | M |
| 72 | 86 | 0.656810154 | cat4 | from_6529 | to_6529 | art_long | art_long | M |
| 28 | 87 | 0.637608265 | cat4 | from_6530 | to_6530 | art_long | art_long | M |
| 30 | 87 | 0.918207722 | cat4 | from_6531 | to_6531 | art_long | art_long | M |
| 47 | 87 | 0.978699007 | cat4 | from_6532 | to_6532 | art_long | art_long | M |
| 55 | 87 | 0.96801302 | cat4 | from_6533 | to_6533 | art_long | art_long | M |
| 58 | 87 | 0.820956139 | cat4 | from_6534 | to_6534 | art_long | art_long | M |
| 69 | 87 | 0.795066711 | cat4 | from_6535 | to_6535 | art_long | art_long | M |
| 72 | 87 | 0.866923341 | cat4 | from_6536 | to_6536 | art_long | art_long | M |
| 80 | 87 | 0.594429538 | cat4 | from_6537 | to_6537 | art_long | art_long | M |
| 83 | 87 | 0.597374246 | cat4 | from_6538 | to_6538 | art_long | art_long | M |
| 86 | 87 | 0.655160491 | cat4 | from_6539 | to_6539 | art_long | art_long | M |
| 8 | 88 | 0.60759503 | cat4 | from_6540 | to_6540 | art_long | art_long | M |
| 9 | 88 | 0.861905995 | cat4 | from_6541 | to_6541 | art_long | art_long | M |
| 10 | 88 | 0.874859849 | cat4 | from_6542 | to_6542 | art_long | art_long | M |
| 22 | 88 | 0.867604211 | cat4 | from_6543 | to_6543 | art_long | art_long | M |
| 23 | 88 | 0.923637104 | cat4 | from_6544 | to_6544 | art_long | art_long | M |
| 65 | 88 | 0.603120418 | cat4 | from_6545 | to_6545 | art_long | art_long | M |
| 25 | 90 | 0.676464002 | cat4 | from_6546 | to_6546 | art_long | art_long | M |
| 28 | 90 | 0.710653946 | cat4 | from_6547 | to_6547 | art_long | art_long | M |
| 30 | 90 | 0.605461205 | cat4 | from_6548 | to_6548 | art_long | art_long | M |
| 31 | 90 | 0.709074212 | cat4 | from_6549 | to_6549 | art_long | art_long | M |
| 33 | 90 | 0.743380059 | cat4 | from_6550 | to_6550 | art_long | art_long | M |

| | | | | | | | | |
|---|---|---|---|---|---|---|---|---|
| 37 | 90 | 0.629325465 | cat4 | from_6551 | to_6551 | art_long | art_long | M |
| 47 | 90 | 0.651929932 | cat4 | from_6552 | to_6552 | art_long | art_long | M |
| 57 | 90 | -0.657117005 | cat4 | from_6553 | to_6553 | art_long | art_long | M |
| 59 | 90 | 0.636547109 | cat4 | from_6554 | to_6554 | art_long | art_long | M |
| 72 | 90 | 0.696056019 | cat4 | from_6555 | to_6555 | art_long | art_long | M |
| 76 | 90 | 0.640551493 | cat4 | from_6556 | to_6556 | art_long | art_long | M |
| 87 | 90 | 0.62797534 | cat4 | from_6557 | to_6557 | art_long | art_long | M |
| 23 | 91 | 0.696978478 | cat4 | from_6558 | to_6558 | art_long | art_long | M |
| 27 | 91 | 0.68667993 | cat4 | from_6559 | to_6559 | art_long | art_long | M |
| 30 | 91 | 0.625632805 | cat4 | from_6560 | to_6560 | art_long | art_long | M |
| 58 | 91 | 0.624585091 | cat4 | from_6561 | to_6561 | art_long | art_long | M |
| 69 | 91 | 0.639437646 | cat4 | from_6562 | to_6562 | art_long | art_long | M |
| 72 | 91 | 0.656991878 | cat4 | from_6563 | to_6563 | art_long | art_long | M |
| 83 | 91 | 0.579105071 | cat4 | from_6564 | to_6564 | art_long | art_long | M |
| 88 | 91 | 0.583185883 | cat4 | from_6565 | to_6565 | art_long | art_long | M |
| 4 | 92 | 0.587676471 | cat4 | from_6566 | to_6566 | art_long | art_long | M |
| 21 | 92 | 0.689368815 | cat4 | from_6567 | to_6567 | art_long | art_long | M |
| 42 | 92 | 0.972652744 | cat4 | from_6568 | to_6568 | art_long | art_long | M |
| 43 | 92 | 0.598778048 | cat4 | from_6569 | to_6569 | art_long | art_long | M |
| 44 | 92 | 0.611164371 | cat4 | from_6570 | to_6570 | art_long | art_long | M |
| 45 | 92 | 0.931962149 | cat4 | from_6571 | to_6571 | art_long | art_long | M |
| 46 | 92 | 0.632452556 | cat4 | from_6572 | to_6572 | art_long | art_long | M |
| 52 | 92 | 0.582905616 | cat4 | from_6573 | to_6573 | art_long | art_long | M |
| 54 | 92 | -0.639088205 | cat4 | from_6574 | to_6574 | art_long | art_long | M |
| 62 | 92 | 0.910295074 | cat4 | from_6575 | to_6575 | art_long | art_long | M |
| 77 | 92 | -0.593265858 | cat4 | from_6576 | to_6576 | art_long | art_long | M |
| 80 | 92 | -0.591623682 | cat4 | from_6577 | to_6577 | art_long | art_long | M |
| 8 | 93 | 0.700208484 | cat4 | from_6578 | to_6578 | art_long | art_long | M |
| 19 | 93 | 0.639880304 | cat4 | from_6579 | to_6579 | art_long | art_long | M |
| 54 | 93 | 0.743394881 | cat4 | from_6580 | to_6580 | art_long | art_long | M |
| 79 | 93 | 0.708545508 | cat4 | from_6581 | to_6581 | art_long | art_long | M |
| 81 | 93 | 0.679006997 | cat4 | from_6582 | to_6582 | art_long | art_long | M |
| 92 | 93 | -0.620442312 | cat4 | from_6583 | to_6583 | art_long | art_long | M |
| 30 | 94 | 0.60894414 | cat4 | from_6584 | to_6584 | art_long | art_long | M |
| 32 | 94 | 0.65434056 | cat4 | from_6585 | to_6585 | art_long | art_long | M |
| 44 | 94 | 0.607816846 | cat4 | from_6586 | to_6586 | art_long | art_long | M |
| 46 | 94 | 0.624968006 | cat4 | from_6587 | to_6587 | art_long | art_long | M |
| 47 | 94 | 0.595107761 | cat4 | from_6588 | to_6588 | art_long | art_long | M |
| 55 | 94 | 0.609925913 | cat4 | from_6589 | to_6589 | art_long | art_long | M |
| 58 | 94 | 0.584323197 | cat4 | from_6590 | to_6590 | art_long | art_long | M |
| 61 | 94 | 0.576410631 | cat4 | from_6591 | to_6591 | art_long | art_long | M |
| 72 | 94 | 0.607044001 | cat4 | from_6592 | to_6592 | art_long | art_long | M |
| 86 | 94 | 0.988688541 | cat4 | from_6593 | to_6593 | art_long | art_long | M |
| 87 | 94 | 0.636273334 | cat4 | from_6594 | to_6594 | art_long | art_long | M |
| 24 | 95 | 0.728463735 | cat4 | from_6595 | to_6595 | art_long | art_long | M |
| 27 | 95 | 0.705553012 | cat4 | from_6596 | to_6596 | art_long | art_long | M |
| 28 | 95 | 0.648836214 | cat4 | from_6597 | to_6597 | art_long | art_long | M |
| 30 | 95 | 0.695750544 | cat4 | from_6598 | to_6598 | art_long | art_long | M |
| 47 | 95 | 0.784086476 | cat4 | from_6599 | to_6599 | art_long | art_long | M |
| 55 | 95 | 0.927308941 | cat4 | from_6600 | to_6600 | art_long | art_long | M |
| 58 | 95 | 0.71184612 | cat4 | from_6601 | to_6601 | art_long | art_long | M |
| 69 | 95 | 0.644089579 | cat4 | from_6602 | to_6602 | art_long | art_long | M |
| 76 | 95 | 0.666008943 | cat4 | from_6603 | to_6603 | art_long | art_long | M |
| 77 | 95 | 0.777125577 | cat4 | from_6604 | to_6604 | art_long | art_long | M |
| 78 | 95 | 0.609828821 | cat4 | from_6605 | to_6605 | art_long | art_long | M |
| 80 | 95 | 0.850286938 | cat4 | from_6606 | to_6606 | art_long | art_long | M |
| 87 | 95 | 0.8183182 | cat4 | from_6607 | to_6607 | art_long | art_long | M |
| 41 | 96 | 0.810958259 | cat4 | from_6608 | to_6608 | art_long | art_long | M |
| 66 | 96 | 0.625054424 | cat4 | from_6609 | to_6609 | art_long | art_long | M |
| 82 | 96 | 0.584871999 | cat4 | from_6610 | to_6610 | art_long | art_long | M |
| 14 | 97 | 0.744816581 | cat4 | from_6611 | to_6611 | art_long | art_long | M |
| 24 | 97 | 0.631569119 | cat4 | from_6612 | to_6612 | art_long | art_long | M |
| 25 | 97 | 0.576638629 | cat4 | from_6613 | to_6613 | art_long | art_long | M |
| 26 | 97 | 0.680029046 | cat4 | from_6614 | to_6614 | art_long | art_long | M |
| 28 | 97 | 0.854778868 | cat4 | from_6615 | to_6615 | art_long | art_long | M |
| 30 | 97 | 0.614593782 | cat4 | from_6616 | to_6616 | art_long | art_long | M |
| 37 | 97 | 0.682202566 | cat4 | from_6617 | to_6617 | art_long | art_long | M |
| 47 | 97 | 0.698913318 | cat4 | from_6618 | to_6618 | art_long | art_long | M |
| 55 | 97 | 0.6076052 | cat4 | from_6619 | to_6619 | art_long | art_long | M |
| 59 | 97 | 0.814350765 | cat4 | from_6620 | to_6620 | art_long | art_long | M |
| 64 | 97 | 0.598937497 | cat4 | from_6621 | to_6621 | art_long | art_long | M |
| 72 | 97 | 0.633637488 | cat4 | from_6622 | to_6622 | art_long | art_long | M |
| 73 | 97 | 0.742069537 | cat4 | from_6623 | to_6623 | art_long | art_long | M |
| 76 | 97 | 0.669555356 | cat4 | from_6624 | to_6624 | art_long | art_long | M |
| 87 | 97 | 0.698358016 | cat4 | from_6625 | to_6625 | art_long | art_long | M |
| 90 | 97 | 0.898015717 | cat4 | from_6626 | to_6626 | art_long | art_long | M |
| 93 | 97 | -0.580683912 | cat4 | from_6627 | to_6627 | art_long | art_long | M |
| 29 | 98 | 0.701364122 | cat4 | from_6628 | to_6628 | art_long | art_long | M |
| 41 | 98 | 0.712232137 | cat4 | from_6629 | to_6629 | art_long | art_long | M |
| 71 | 98 | 0.712391723 | cat4 | from_6630 | to_6630 | art_long | art_long | M |
| 82 | 98 | 0.677665878 | cat4 | from_6631 | to_6631 | art_long | art_long | M |
| 44 | 99 | 0.895628537 | cat4 | from_6632 | to_6632 | art_long | art_long | M |
| 46 | 99 | 0.868856001 | cat4 | from_6633 | to_6633 | art_long | art_long | M |
| 50 | 99 | 0.604491628 | cat4 | from_6634 | to_6634 | art_long | art_long | M |
| 52 | 99 | 0.740713361 | cat4 | from_6635 | to_6635 | art_long | art_long | M |
| 60 | 99 | 0.896176881 | cat4 | from_6636 | to_6636 | art_long | art_long | M |
| 61 | 99 | 0.831868398 | cat4 | from_6637 | to_6637 | art_long | art_long | M |
| 86 | 99 | 0.618789877 | cat4 | from_6638 | to_6638 | art_long | art_long | M |
| 94 | 99 | 0.599693528 | cat4 | from_6639 | to_6639 | art_long | art_long | M |
| 8 | 100 | -0.602989149 | cat4 | from_6640 | to_6640 | art_long | art_long | M |
| 42 | 100 | 0.907223329 | cat4 | from_6641 | to_6641 | art_long | art_long | M |
| 44 | 100 | 0.622176279 | cat4 | from_6642 | to_6642 | art_long | art_long | M |
| 45 | 100 | 0.960462422 | cat4 | from_6643 | to_6643 | art_long | art_long | M |
| 46 | 100 | 0.66407332 | cat4 | from_6644 | to_6644 | art_long | art_long | M |
| 62 | 100 | 0.929311629 | cat4 | from_6645 | to_6645 | art_long | art_long | M |
| 81 | 100 | -0.659821862 | cat4 | from_6646 | to_6646 | art_long | art_long | M |
| 92 | 100 | 0.87006338 | cat4 | from_6647 | to_6647 | art_long | art_long | M |
| 4 | 101 | 0.629834616 | cat4 | from_6648 | to_6648 | art_long | art_long | M |
| 17 | 101 | 0.594846204 | cat4 | from_6649 | to_6649 | art_long | art_long | M |
| 21 | 101 | 0.600947436 | cat4 | from_6650 | to_6650 | art_long | art_long | M |
| 42 | 101 | 0.977724713 | cat4 | from_6651 | to_6651 | art_long | art_long | M |
| 43 | 101 | 0.756699686 | cat4 | from_6652 | to_6652 | art_long | art_long | M |
| 44 | 101 | 0.672938738 | cat4 | from_6653 | to_6653 | art_long | art_long | M |
| 45 | 101 | 0.958197444 | cat4 | from_6654 | to_6654 | art_long | art_long | M |

| | | | | | | | | |
|---|---|---|---|---|---|---|---|---|
| 46 | 101 | 0.659785487 | cat4 | from_6655 | to_6655 | art_long | art_long | M |
| 52 | 101 | 0.589690313 | cat4 | from_6656 | to_6656 | art_long | art_long | M |
| 60 | 101 | 0.612710581 | cat4 | from_6657 | to_6657 | art_long | art_long | M |
| 62 | 101 | 0.987947948 | cat4 | from_6658 | to_6658 | art_long | art_long | M |
| 92 | 101 | 0.940421392 | cat4 | from_6659 | to_6659 | art_long | art_long | M |
| 100 | 101 | 0.886016715 | cat4 | from_6660 | to_6660 | art_long | art_long | M |
| 35 | 102 | 0.633031923 | cat4 | from_6661 | to_6661 | art_long | art_long | M |
| 42 | 102 | 0.635896804 | cat4 | from_6662 | to_6662 | art_long | art_long | M |
| 44 | 102 | 0.894075227 | cat4 | from_6663 | to_6663 | art_long | art_long | M |
| 45 | 102 | 0.638179249 | cat4 | from_6664 | to_6664 | art_long | art_long | M |
| 46 | 102 | 0.926383347 | cat4 | from_6665 | to_6665 | art_long | art_long | M |
| 48 | 102 | 0.636727101 | cat4 | from_6666 | to_6666 | art_long | art_long | M |
| 52 | 102 | 0.765444532 | cat4 | from_6667 | to_6667 | art_long | art_long | M |
| 60 | 102 | 0.615713021 | cat4 | from_6668 | to_6668 | art_long | art_long | M |
| 61 | 102 | 0.647688278 | cat4 | from_6669 | to_6669 | art_long | art_long | M |
| 62 | 102 | 0.619583973 | cat4 | from_6670 | to_6670 | art_long | art_long | M |
| 92 | 102 | 0.578801684 | cat4 | from_6671 | to_6671 | art_long | art_long | M |
| 99 | 102 | 0.73098846 | cat4 | from_6672 | to_6672 | art_long | art_long | M |
| 100 | 102 | 0.734520731 | cat4 | from_6673 | to_6673 | art_long | art_long | M |
| 101 | 102 | 0.581925902 | cat4 | from_6674 | to_6674 | art_long | art_long | M |
| 42 | 103 | 0.623752496 | cat4 | from_6675 | to_6675 | art_long | art_long | M |
| 54 | 103 | -0.694431111 | cat4 | from_6676 | to_6676 | art_long | art_long | M |
| 81 | 103 | -0.634119625 | cat4 | from_6677 | to_6677 | art_long | art_long | M |
| 90 | 103 | 0.603549898 | cat4 | from_6678 | to_6678 | art_long | art_long | M |
| 92 | 103 | 0.645143496 | cat4 | from_6679 | to_6679 | art_long | art_long | M |
| 93 | 103 | -0.764560879 | cat4 | from_6680 | to_6680 | art_long | art_long | M |
| 97 | 103 | 0.62788077 | cat4 | from_6681 | to_6681 | art_long | art_long | M |
| 17 | 104 | 0.707866697 | cat4 | from_6682 | to_6682 | art_long | art_long | M |
| 31 | 104 | -0.659311275 | cat4 | from_6683 | to_6683 | art_long | art_long | M |
| 33 | 104 | -0.650466347 | cat4 | from_6684 | to_6684 | art_long | art_long | M |
| 35 | 104 | 0.81936042 | cat4 | from_6685 | to_6685 | art_long | art_long | M |
| 36 | 104 | 0.702659037 | cat4 | from_6686 | to_6686 | art_long | art_long | M |
| 38 | 104 | 0.813506117 | cat4 | from_6687 | to_6687 | art_long | art_long | M |
| 48 | 104 | 0.963063377 | cat4 | from_6688 | to_6688 | art_long | art_long | M |
| 50 | 104 | 0.631147954 | cat4 | from_6689 | to_6689 | art_long | art_long | M |
| 20 | 105 | 0.577553235 | cat4 | from_6690 | to_6690 | art_long | art_long | M |
| 24 | 105 | -0.577894669 | cat4 | from_6691 | to_6691 | art_long | art_long | M |
| 44 | 105 | 0.707713669 | cat4 | from_6692 | to_6692 | art_long | art_long | M |
| 46 | 105 | 0.680622875 | cat4 | from_6693 | to_6693 | art_long | art_long | M |
| 50 | 105 | 0.637452426 | cat4 | from_6694 | to_6694 | art_long | art_long | M |
| 60 | 105 | 0.751024773 | cat4 | from_6695 | to_6695 | art_long | art_long | M |
| 61 | 105 | 0.759988009 | cat4 | from_6696 | to_6696 | art_long | art_long | M |
| 99 | 105 | 0.945430619 | cat4 | from_6697 | to_6697 | art_long | art_long | M |
| 11 | 106 | 0.584032239 | cat4 | from_6698 | to_6698 | art_long | art_long | M |
| 35 | 106 | 0.579512307 | cat4 | from_6699 | to_6699 | art_long | art_long | M |
| 42 | 106 | 0.583941883 | cat4 | from_6700 | to_6700 | art_long | art_long | M |
| 44 | 106 | 0.9648459 | cat4 | from_6701 | to_6701 | art_long | art_long | M |
| 46 | 106 | 0.933380147 | cat4 | from_6702 | to_6702 | art_long | art_long | M |
| 48 | 106 | 0.592588711 | cat4 | from_6703 | to_6703 | art_long | art_long | M |
| 52 | 106 | 0.794920733 | cat4 | from_6704 | to_6704 | art_long | art_long | M |
| 60 | 106 | 0.829668705 | cat4 | from_6705 | to_6705 | art_long | art_long | M |
| 61 | 106 | 0.814695013 | cat4 | from_6706 | to_6706 | art_long | art_long | M |
| 62 | 106 | 0.637094007 | cat4 | from_6707 | to_6707 | art_long | art_long | M |
| 99 | 106 | 0.825430147 | cat4 | from_6708 | to_6708 | art_long | art_long | M |
| 100 | 106 | 0.583538421 | cat4 | from_6709 | to_6709 | art_long | art_long | M |
| 101 | 106 | 0.616247034 | cat4 | from_6710 | to_6710 | art_long | art_long | M |
| 102 | 106 | 0.87064637 | cat4 | from_6711 | to_6711 | art_long | art_long | M |
| 105 | 106 | 0.629978362 | cat4 | from_6712 | to_6712 | art_long | art_long | M |
| 4 | 107 | 0.63531835 | cat4 | from_6713 | to_6713 | art_long | art_long | M |
| 7 | 107 | 0.616699413 | cat4 | from_6714 | to_6714 | art_long | art_long | M |
| 17 | 107 | 0.669652919 | cat4 | from_6715 | to_6715 | art_long | art_long | M |
| 36 | 107 | 0.704658491 | cat4 | from_6716 | to_6716 | art_long | art_long | M |
| 42 | 107 | 0.714593955 | cat4 | from_6717 | to_6717 | art_long | art_long | M |
| 43 | 107 | 0.932567348 | cat4 | from_6718 | to_6718 | art_long | art_long | M |
| 45 | 107 | 0.810605982 | cat4 | from_6719 | to_6719 | art_long | art_long | M |
| 60 | 107 | 0.624643873 | cat4 | from_6720 | to_6720 | art_long | art_long | M |
| 62 | 107 | 0.859364573 | cat4 | from_6721 | to_6721 | art_long | art_long | M |
| 92 | 107 | 0.676556647 | cat4 | from_6722 | to_6722 | art_long | art_long | M |
| 100 | 107 | 0.658149191 | cat4 | from_6723 | to_6723 | art_long | art_long | M |
| 101 | 107 | 0.844451455 | cat4 | from_6724 | to_6724 | art_long | art_long | M |
| 17 | 108 | 0.592911649 | cat4 | from_6725 | to_6725 | art_long | art_long | M |
| 36 | 108 | 0.678079285 | cat4 | from_6726 | to_6726 | art_long | art_long | M |
| 70 | 108 | 0.60960203 | cat4 | from_6727 | to_6727 | art_long | art_long | M |
| 75 | 108 | 0.62396779 | cat4 | from_6728 | to_6728 | art_long | art_long | M |
| 96 | 108 | 0.594399855 | cat4 | from_6729 | to_6729 | art_long | art_long | M |
| 24 | 109 | 0.636484431 | cat4 | from_6730 | to_6730 | art_long | art_long | M |
| 27 | 109 | 0.635041777 | cat4 | from_6731 | to_6731 | art_long | art_long | M |
| 28 | 109 | 0.667972848 | cat4 | from_6732 | to_6732 | art_long | art_long | M |
| 30 | 109 | 0.869206331 | cat4 | from_6733 | to_6733 | art_long | art_long | M |
| 47 | 109 | 0.922375208 | cat4 | from_6734 | to_6734 | art_long | art_long | M |
| 55 | 109 | 0.992300687 | cat4 | from_6735 | to_6735 | art_long | art_long | M |
| 58 | 109 | 0.835254673 | cat4 | from_6736 | to_6736 | art_long | art_long | M |
| 69 | 109 | 0.79140585 | cat4 | from_6737 | to_6737 | art_long | art_long | M |
| 72 | 109 | 0.751028482 | cat4 | from_6738 | to_6738 | art_long | art_long | M |
| 76 | 109 | 0.627603538 | cat4 | from_6739 | to_6739 | art_long | art_long | M |
| 77 | 109 | 0.726164519 | cat4 | from_6740 | to_6740 | art_long | art_long | M |
| 80 | 109 | 0.77140661 | cat4 | from_6741 | to_6741 | art_long | art_long | M |
| 83 | 109 | 0.629531961 | cat4 | from_6742 | to_6742 | art_long | art_long | M |
| 86 | 109 | 0.592349431 | cat4 | from_6743 | to_6743 | art_long | art_long | M |
| 87 | 109 | 0.956761438 | cat4 | from_6744 | to_6744 | art_long | art_long | M |
| 94 | 109 | 0.61659284 | cat4 | from_6745 | to_6745 | art_long | art_long | M |
| 95 | 109 | 0.943870684 | cat4 | from_6746 | to_6746 | art_long | art_long | M |
| 97 | 109 | 0.578766808 | cat4 | from_6747 | to_6747 | art_long | art_long | M |
| 14 | 110 | 0.966400356 | cat4 | from_6748 | to_6748 | art_long | art_long | M |
| 24 | 110 | 0.581702661 | cat4 | from_6749 | to_6749 | art_long | art_long | M |
| 25 | 110 | 0.765034774 | cat4 | from_6750 | to_6750 | art_long | art_long | M |
| 28 | 110 | 0.702878469 | cat4 | from_6751 | to_6751 | art_long | art_long | M |
| 31 | 110 | 0.721818691 | cat4 | from_6752 | to_6752 | art_long | art_long | M |
| 33 | 110 | 0.713353571 | cat4 | from_6753 | to_6753 | art_long | art_long | M |
| 58 | 110 | 0.668146437 | cat4 | from_6754 | to_6754 | art_long | art_long | M |
| 59 | 110 | 0.92140124 | cat4 | from_6755 | to_6755 | art_long | art_long | M |
| 68 | 110 | 0.584307663 | cat4 | from_6756 | to_6756 | art_long | art_long | M |
| 69 | 110 | 0.691567298 | cat4 | from_6757 | to_6757 | art_long | art_long | M |
| 73 | 110 | 0.965794474 | cat4 | from_6758 | to_6758 | art_long | art_long | M |

| | | | | | | | | |
|---|---|---|---|---|---|---|---|---|
| 74 | 110 | 0.873420144 | cat4 | from_6759 | to_6759 | art_long | art_long | M |
| 76 | 110 | 0.656810832 | cat4 | from_6760 | to_6760 | art_long | art_long | M |
| 78 | 110 | 0.747509672 | cat4 | from_6761 | to_6761 | art_long | art_long | M |
| 83 | 110 | 0.644571778 | cat4 | from_6762 | to_6762 | art_long | art_long | M |
| 97 | 110 | 0.637928859 | cat4 | from_6763 | to_6763 | art_long | art_long | M |
| 30 | 111 | 0.872739918 | cat4 | from_6764 | to_6764 | art_long | art_long | M |
| 47 | 111 | 0.948053121 | cat4 | from_6765 | to_6765 | art_long | art_long | M |
| 55 | 111 | 0.839173825 | cat4 | from_6766 | to_6766 | art_long | art_long | M |
| 58 | 111 | 0.699021819 | cat4 | from_6767 | to_6767 | art_long | art_long | M |
| 69 | 111 | 0.69909994 | cat4 | from_6768 | to_6768 | art_long | art_long | M |
| 72 | 111 | 0.910329354 | cat4 | from_6769 | to_6769 | art_long | art_long | M |
| 86 | 111 | 0.641533844 | cat4 | from_6770 | to_6770 | art_long | art_long | M |
| 87 | 111 | 0.939116542 | cat4 | from_6771 | to_6771 | art_long | art_long | M |
| 90 | 111 | 0.717774632 | cat4 | from_6772 | to_6772 | art_long | art_long | M |
| 91 | 111 | 0.578746872 | cat4 | from_6773 | to_6773 | art_long | art_long | M |
| 94 | 111 | 0.576161347 | cat4 | from_6774 | to_6774 | art_long | art_long | M |
| 95 | 111 | 0.578741445 | cat4 | from_6775 | to_6775 | art_long | art_long | M |
| 97 | 111 | 0.754868134 | cat4 | from_6776 | to_6776 | art_long | art_long | M |
| 109 | 111 | 0.803760589 | cat4 | from_6777 | to_6777 | art_long | art_long | M |
| 14 | 112 | 0.790772209 | cat4 | from_6778 | to_6778 | art_long | art_long | M |
| 25 | 112 | 0.972614149 | cat4 | from_6779 | to_6779 | art_long | art_long | M |
| 28 | 112 | 0.578958387 | cat4 | from_6780 | to_6780 | art_long | art_long | M |
| 31 | 112 | 0.917274144 | cat4 | from_6781 | to_6781 | art_long | art_long | M |
| 33 | 112 | 0.923915927 | cat4 | from_6782 | to_6782 | art_long | art_long | M |
| 48 | 112 | -0.605057434 | cat4 | from_6783 | to_6783 | art_long | art_long | M |
| 51 | 112 | 0.610860304 | cat4 | from_6784 | to_6784 | art_long | art_long | M |
| 54 | 112 | 0.62619286 | cat4 | from_6785 | to_6785 | art_long | art_long | M |
| 59 | 112 | 0.757618137 | cat4 | from_6786 | to_6786 | art_long | art_long | M |
| 68 | 112 | 0.745368132 | cat4 | from_6787 | to_6787 | art_long | art_long | M |
| 69 | 112 | 0.579627748 | cat4 | from_6788 | to_6788 | art_long | art_long | M |
| 73 | 112 | 0.791910129 | cat4 | from_6789 | to_6789 | art_long | art_long | M |
| 74 | 112 | 0.703799077 | cat4 | from_6790 | to_6790 | art_long | art_long | M |
| 76 | 112 | 0.70541577 | cat4 | from_6791 | to_6791 | art_long | art_long | M |
| 78 | 112 | 0.583977866 | cat4 | from_6792 | to_6792 | art_long | art_long | M |
| 81 | 112 | 0.723319165 | cat4 | from_6793 | to_6793 | art_long | art_long | M |
| 83 | 112 | 0.695017199 | cat4 | from_6794 | to_6794 | art_long | art_long | M |
| 90 | 112 | 0.617304074 | cat4 | from_6795 | to_6795 | art_long | art_long | M |
| 110 | 112 | 0.826642596 | cat4 | from_6796 | to_6796 | art_long | art_long | M |
| 7 | 113 | 0.578795495 | cat4 | from_6797 | to_6797 | art_long | art_long | M |
| 22 | 113 | 0.639022658 | cat4 | from_6798 | to_6798 | art_long | art_long | M |
| 34 | 113 | 0.762969508 | cat4 | from_6799 | to_6799 | art_long | art_long | M |
| 50 | 113 | -0.576913279 | cat4 | from_6800 | to_6800 | art_long | art_long | M |
| 64 | 113 | 0.740557881 | cat4 | from_6801 | to_6801 | art_long | art_long | M |
| 65 | 113 | 0.870376141 | cat4 | from_6802 | to_6802 | art_long | art_long | M |
| 12 | 114 | 0.737403906 | cat4 | from_6803 | to_6803 | art_long | art_long | M |
| 13 | 114 | 0.929814442 | cat4 | from_6804 | to_6804 | art_long | art_long | M |
| 15 | 114 | 0.630396812 | cat4 | from_6805 | to_6805 | art_long | art_long | M |
| 19 | 114 | 0.791742754 | cat4 | from_6806 | to_6806 | art_long | art_long | M |
| 20 | 114 | 0.72002622 | cat4 | from_6807 | to_6807 | art_long | art_long | M |
| 39 | 114 | 0.641562755 | cat4 | from_6808 | to_6808 | art_long | art_long | M |
| 40 | 114 | 0.71533678 | cat4 | from_6809 | to_6809 | art_long | art_long | M |
| 68 | 114 | 0.761419577 | cat4 | from_6810 | to_6810 | art_long | art_long | M |
| 79 | 114 | 0.784710929 | cat4 | from_6811 | to_6811 | art_long | art_long | M |
| 26 | 115 | 0.932998986 | cat4 | from_6812 | to_6812 | art_long | art_long | M |
| 37 | 115 | 0.921809169 | cat4 | from_6813 | to_6813 | art_long | art_long | M |
| 90 | 115 | 0.662988455 | cat4 | from_6814 | to_6814 | art_long | art_long | M |
| 93 | 115 | -0.603223117 | cat4 | from_6815 | to_6815 | art_long | art_long | M |
| 97 | 115 | 0.728152843 | cat4 | from_6816 | to_6816 | art_long | art_long | M |
| 111 | 115 | 0.633734482 | cat4 | from_6817 | to_6817 | art_long | art_long | M |
| 5 | 116 | 0.866441049 | cat4 | from_6818 | to_6818 | art_long | art_long | M |
| 16 | 116 | 0.602699728 | cat4 | from_6819 | to_6819 | art_long | art_long | M |
| 34 | 116 | 0.576978784 | cat4 | from_6820 | to_6820 | art_long | art_long | M |
| 63 | 116 | 0.797717878 | cat4 | from_6821 | to_6821 | art_long | art_long | M |
| 65 | 116 | 0.6428464 | cat4 | from_6822 | to_6822 | art_long | art_long | M |
| 113 | 116 | 0.596888801 | cat4 | from_6823 | to_6823 | art_long | art_long | M |
| 28 | 117 | 0.579533151 | cat4 | from_6824 | to_6824 | art_long | art_long | M |
| 34 | 117 | 0.744262446 | cat4 | from_6825 | to_6825 | art_long | art_long | M |
| 59 | 117 | 0.578587595 | cat4 | from_6826 | to_6826 | art_long | art_long | M |
| 64 | 117 | 0.855786375 | cat4 | from_6827 | to_6827 | art_long | art_long | M |
| 65 | 117 | 0.629020475 | cat4 | from_6828 | to_6828 | art_long | art_long | M |
| 90 | 117 | 0.585649518 | cat4 | from_6829 | to_6829 | art_long | art_long | M |
| 91 | 117 | 0.722539906 | cat4 | from_6830 | to_6830 | art_long | art_long | M |
| 97 | 117 | 0.726305632 | cat4 | from_6831 | to_6831 | art_long | art_long | M |
| 26 | 118 | 0.733695648 | cat4 | from_6832 | to_6832 | art_long | art_long | M |
| 37 | 118 | 0.707960058 | cat4 | from_6833 | to_6833 | art_long | art_long | M |
| 55 | 118 | 0.608556585 | cat4 | from_6834 | to_6834 | art_long | art_long | M |
| 58 | 118 | 0.63006594 | cat4 | from_6835 | to_6835 | art_long | art_long | M |
| 69 | 118 | 0.606925118 | cat4 | from_6836 | to_6836 | art_long | art_long | M |
| 87 | 118 | 0.61312252 | cat4 | from_6837 | to_6837 | art_long | art_long | M |
| 95 | 118 | 0.583611559 | cat4 | from_6838 | to_6838 | art_long | art_long | M |
| 109 | 118 | 0.630618593 | cat4 | from_6839 | to_6839 | art_long | art_long | M |
| 115 | 118 | 0.593073486 | cat4 | from_6840 | to_6840 | art_long | art_long | M |
| 14 | 119 | 0.589991439 | cat4 | from_6841 | to_6841 | art_long | art_long | M |
| 24 | 119 | 0.633079828 | cat4 | from_6842 | to_6842 | art_long | art_long | M |
| 27 | 119 | 0.645978342 | cat4 | from_6843 | to_6843 | art_long | art_long | M |
| 28 | 119 | 0.634888765 | cat4 | from_6844 | to_6844 | art_long | art_long | M |
| 30 | 119 | 0.85387965 | cat4 | from_6845 | to_6845 | art_long | art_long | M |
| 47 | 119 | 0.752213278 | cat4 | from_6846 | to_6846 | art_long | art_long | M |
| 54 | 119 | 0.663868096 | cat4 | from_6847 | to_6847 | art_long | art_long | M |
| 55 | 119 | 0.883419839 | cat4 | from_6848 | to_6848 | art_long | art_long | M |
| 58 | 119 | 0.94563786 | cat4 | from_6849 | to_6849 | art_long | art_long | M |
| 59 | 119 | 0.635178113 | cat4 | from_6850 | to_6850 | art_long | art_long | M |
| 69 | 119 | 0.912891392 | cat4 | from_6851 | to_6851 | art_long | art_long | M |
| 72 | 119 | 0.692857264 | cat4 | from_6852 | to_6852 | art_long | art_long | M |
| 73 | 119 | 0.597331323 | cat4 | from_6853 | to_6853 | art_long | art_long | M |
| 74 | 119 | 0.642052305 | cat4 | from_6854 | to_6854 | art_long | art_long | M |
| 76 | 119 | 0.634432413 | cat4 | from_6855 | to_6855 | art_long | art_long | M |
| 77 | 119 | 0.875922137 | cat4 | from_6856 | to_6856 | art_long | art_long | M |
| 78 | 119 | 0.665969969 | cat4 | from_6857 | to_6857 | art_long | art_long | M |
| 80 | 119 | 0.868345558 | cat4 | from_6858 | to_6858 | art_long | art_long | M |
| 81 | 119 | 0.6039519 | cat4 | from_6859 | to_6859 | art_long | art_long | M |
| 83 | 119 | 0.803089047 | cat4 | from_6860 | to_6860 | art_long | art_long | M |
| 87 | 119 | 0.85007887 | cat4 | from_6861 | to_6861 | art_long | art_long | M |
| 94 | 119 | 0.586059771 | cat4 | from_6862 | to_6862 | art_long | art_long | M |

| | | | | | | | | |
|---|---|---|---|---|---|---|---|---|
| 95 | 119 | 0.892501591 | cat4 | from_6863 | to_6863 | art_long | art_long | M |
| 109 | 119 | 0.933955268 | cat4 | from_6864 | to_6864 | art_long | art_long | M |
| 111 | 119 | 0.653640699 | cat4 | from_6865 | to_6865 | art_long | art_long | M |
| 118 | 119 | 0.644696889 | cat4 | from_6866 | to_6866 | art_long | art_long | M |
| 54 | 120 | 0.943395493 | cat4 | from_6867 | to_6867 | art_long | art_long | M |
| 58 | 120 | 0.692507612 | cat4 | from_6868 | to_6868 | art_long | art_long | M |
| 69 | 120 | 0.677408227 | cat4 | from_6869 | to_6869 | art_long | art_long | M |
| 74 | 120 | 0.696665233 | cat4 | from_6870 | to_6870 | art_long | art_long | M |
| 77 | 120 | 0.904952908 | cat4 | from_6871 | to_6871 | art_long | art_long | M |
| 78 | 120 | 0.65572504 | cat4 | from_6872 | to_6872 | art_long | art_long | M |
| 80 | 120 | 0.828149074 | cat4 | from_6873 | to_6873 | art_long | art_long | M |
| 81 | 120 | 0.886996735 | cat4 | from_6874 | to_6874 | art_long | art_long | M |
| 83 | 120 | 0.798676145 | cat4 | from_6875 | to_6875 | art_long | art_long | M |
| 92 | 120 | -0.601345813 | cat4 | from_6876 | to_6876 | art_long | art_long | M |
| 93 | 120 | 0.751264166 | cat4 | from_6877 | to_6877 | art_long | art_long | M |
| 103 | 120 | -0.738575239 | cat4 | from_6878 | to_6878 | art_long | art_long | M |
| 119 | 120 | 0.698876092 | cat4 | from_6879 | to_6879 | art_long | art_long | M |
| 29 | 121 | 0.627351143 | cat4 | from_6880 | to_6880 | art_long | art_long | M |
| 41 | 121 | 0.875984138 | cat4 | from_6881 | to_6881 | art_long | art_long | M |
| 71 | 121 | 0.652759102 | cat4 | from_6882 | to_6882 | art_long | art_long | M |
| 82 | 121 | 0.783222206 | cat4 | from_6883 | to_6883 | art_long | art_long | M |
| 96 | 121 | 0.750502948 | cat4 | from_6884 | to_6884 | art_long | art_long | M |
| 98 | 121 | 0.829753238 | cat4 | from_6885 | to_6885 | art_long | art_long | M |
| 14 | 122 | 0.714875407 | cat4 | from_6886 | to_6886 | art_long | art_long | M |
| 24 | 122 | 0.99431374 | cat4 | from_6887 | to_6887 | art_long | art_long | M |
| 25 | 122 | 0.633286046 | cat4 | from_6888 | to_6888 | art_long | art_long | M |
| 28 | 122 | 0.970012158 | cat4 | from_6889 | to_6889 | art_long | art_long | M |
| 32 | 122 | 0.678568139 | cat4 | from_6890 | to_6890 | art_long | art_long | M |
| 55 | 122 | 0.632672197 | cat4 | from_6891 | to_6891 | art_long | art_long | M |
| 59 | 122 | 0.848512753 | cat4 | from_6892 | to_6892 | art_long | art_long | M |
| 64 | 122 | 0.637657069 | cat4 | from_6893 | to_6893 | art_long | art_long | M |
| 73 | 122 | 0.736246058 | cat4 | from_6894 | to_6894 | art_long | art_long | M |
| 74 | 122 | 0.837627095 | cat4 | from_6895 | to_6895 | art_long | art_long | M |
| 76 | 122 | 0.932729357 | cat4 | from_6896 | to_6896 | art_long | art_long | M |
| 78 | 122 | 0.901158516 | cat4 | from_6897 | to_6897 | art_long | art_long | M |
| 95 | 122 | 0.71145308 | cat4 | from_6898 | to_6898 | art_long | art_long | M |
| 97 | 122 | 0.705012393 | cat4 | from_6899 | to_6899 | art_long | art_long | M |
| 109 | 122 | 0.653327785 | cat4 | from_6900 | to_6900 | art_long | art_long | M |
| 110 | 122 | 0.637932701 | cat4 | from_6901 | to_6901 | art_long | art_long | M |
| 119 | 122 | 0.648080925 | cat4 | from_6902 | to_6902 | art_long | art_long | M |
| 34 | 123 | 0.692959916 | cat4 | from_6903 | to_6903 | art_long | art_long | M |
| 49 | 123 | 0.740098166 | cat4 | from_6904 | to_6904 | art_long | art_long | M |
| 56 | 123 | 0.737757945 | cat4 | from_6905 | to_6905 | art_long | art_long | M |
| 57 | 123 | 0.639804506 | cat4 | from_6906 | to_6906 | art_long | art_long | M |
| 70 | 123 | 0.908322114 | cat4 | from_6907 | to_6907 | art_long | art_long | M |
| 75 | 123 | 0.836954073 | cat4 | from_6908 | to_6908 | art_long | art_long | M |
| 85 | 123 | 0.883733831 | cat4 | from_6909 | to_6909 | art_long | art_long | M |
| 3 | 124 | 0.595243827 | cat4 | from_6910 | to_6910 | art_long | art_long | M |
| 14 | 124 | 0.607430938 | cat4 | from_6911 | to_6911 | art_long | art_long | M |
| 29 | 124 | 0.764215366 | cat4 | from_6912 | to_6912 | art_long | art_long | M |
| 41 | 124 | 0.690847408 | cat4 | from_6913 | to_6913 | art_long | art_long | M |
| 59 | 124 | 0.584692983 | cat4 | from_6914 | to_6914 | art_long | art_long | M |
| 66 | 124 | 0.631317762 | cat4 | from_6915 | to_6915 | art_long | art_long | M |
| 67 | 124 | 0.688162083 | cat4 | from_6916 | to_6916 | art_long | art_long | M |
| 71 | 124 | 0.795010771 | cat4 | from_6917 | to_6917 | art_long | art_long | M |
| 73 | 124 | 0.607771841 | cat4 | from_6918 | to_6918 | art_long | art_long | M |
| 82 | 124 | 0.657035654 | cat4 | from_6919 | to_6919 | art_long | art_long | M |
| 6 | 125 | 0.767943138 | cat4 | from_6920 | to_6920 | art_long | art_long | M |
| 8 | 125 | 0.691148305 | cat4 | from_6921 | to_6921 | art_long | art_long | M |
| 9 | 125 | 0.700923244 | cat4 | from_6922 | to_6922 | art_long | art_long | M |
| 10 | 125 | 0.719960715 | cat4 | from_6923 | to_6923 | art_long | art_long | M |
| 23 | 125 | 0.664163301 | cat4 | from_6924 | to_6924 | art_long | art_long | M |
| 49 | 125 | 0.641072814 | cat4 | from_6925 | to_6925 | art_long | art_long | M |
| 65 | 125 | 0.767325941 | cat4 | from_6926 | to_6926 | art_long | art_long | M |
| 75 | 125 | 0.652849127 | cat4 | from_6927 | to_6927 | art_long | art_long | M |
| 84 | 125 | 0.798274586 | cat4 | from_6928 | to_6928 | art_long | art_long | M |
| 8 | 126 | 0.715994509 | cat4 | from_6929 | to_6929 | art_long | art_long | M |
| 9 | 126 | 0.906555263 | cat4 | from_6930 | to_6930 | art_long | art_long | M |
| 10 | 126 | 0.889882668 | cat4 | from_6931 | to_6931 | art_long | art_long | M |
| 22 | 126 | 0.821329215 | cat4 | from_6932 | to_6932 | art_long | art_long | M |
| 23 | 126 | 0.848043362 | cat4 | from_6933 | to_6933 | art_long | art_long | M |
| 27 | 126 | 0.592360837 | cat4 | from_6934 | to_6934 | art_long | art_long | M |
| 57 | 126 | 0.722616139 | cat4 | from_6935 | to_6935 | art_long | art_long | M |
| 88 | 126 | 0.900994166 | cat4 | from_6936 | to_6936 | art_long | art_long | M |
| 93 | 126 | 0.619241181 | cat4 | from_6937 | to_6937 | art_long | art_long | M |
| 115 | 126 | -0.586421449 | cat4 | from_6938 | to_6938 | art_long | art_long | M |
| 4 | 127 | 0.698352444 | cat4 | from_6939 | to_6939 | art_long | art_long | M |
| 5 | 127 | 0.704239435 | cat4 | from_6940 | to_6940 | art_long | art_long | M |
| 34 | 127 | 0.933158826 | cat4 | from_6941 | to_6941 | art_long | art_long | M |
| 49 | 127 | 0.714830077 | cat4 | from_6942 | to_6942 | art_long | art_long | M |
| 65 | 127 | 0.845351726 | cat4 | from_6943 | to_6943 | art_long | art_long | M |
| 70 | 127 | 0.614940313 | cat4 | from_6944 | to_6944 | art_long | art_long | M |
| 85 | 127 | 0.766588833 | cat4 | from_6945 | to_6945 | art_long | art_long | M |
| 113 | 127 | 0.648307674 | cat4 | from_6946 | to_6946 | art_long | art_long | M |
| 116 | 127 | 0.584674462 | cat4 | from_6947 | to_6947 | art_long | art_long | M |
| 117 | 127 | 0.769982821 | cat4 | from_6948 | to_6948 | art_long | art_long | M |
| 123 | 127 | 0.695132047 | cat4 | from_6949 | to_6949 | art_long | art_long | M |
| 125 | 127 | 0.616271532 | cat4 | from_6950 | to_6950 | art_long | art_long | M |
| 11 | 128 | 0.633646692 | cat4 | from_6951 | to_6951 | art_long | art_long | M |
| 35 | 128 | 0.595845883 | cat4 | from_6952 | to_6952 | art_long | art_long | M |
| 38 | 128 | 0.641232083 | cat4 | from_6953 | to_6953 | art_long | art_long | M |
| 43 | 128 | 0.607334802 | cat4 | from_6954 | to_6954 | art_long | art_long | M |
| 44 | 128 | 0.95567377 | cat4 | from_6955 | to_6955 | art_long | art_long | M |
| 45 | 128 | 0.582054567 | cat4 | from_6956 | to_6956 | art_long | art_long | M |
| 46 | 128 | 0.931353194 | cat4 | from_6957 | to_6957 | art_long | art_long | M |
| 52 | 128 | 0.845728348 | cat4 | from_6958 | to_6958 | art_long | art_long | M |
| 60 | 128 | 0.974283379 | cat4 | from_6959 | to_6959 | art_long | art_long | M |
| 61 | 128 | 0.868758306 | cat4 | from_6960 | to_6960 | art_long | art_long | M |
| 62 | 128 | 0.622974547 | cat4 | from_6961 | to_6961 | art_long | art_long | M |
| 86 | 128 | 0.582785895 | cat4 | from_6962 | to_6962 | art_long | art_long | M |
| 94 | 128 | 0.596607237 | cat4 | from_6963 | to_6963 | art_long | art_long | M |
| 99 | 128 | 0.888197667 | cat4 | from_6964 | to_6964 | art_long | art_long | M |
| 101 | 128 | 0.633601179 | cat4 | from_6965 | to_6965 | art_long | art_long | M |
| 102 | 128 | 0.736999008 | cat4 | from_6966 | to_6966 | art_long | art_long | M |

| | | | | | | | | |
|---|---|---|---|---|---|---|---|---|
| 105 | 128 | 0.711235494 | cat4 | from_6967 | to_6967 | art_long | art_long | M |
| 106 | 128 | 0.91354593 | cat4 | from_6968 | to_6968 | art_long | art_long | M |
| 107 | 128 | 0.643812904 | cat4 | from_6969 | to_6969 | art_long | art_long | M |
| 25 | 129 | 0.579171229 | cat4 | from_6970 | to_6970 | art_long | art_long | M |
| 26 | 129 | 0.823186542 | cat4 | from_6971 | to_6971 | art_long | art_long | M |
| 37 | 129 | 0.916163006 | cat4 | from_6972 | to_6972 | art_long | art_long | M |
| 90 | 129 | 0.602243749 | cat4 | from_6973 | to_6973 | art_long | art_long | M |
| 115 | 129 | 0.732370413 | cat4 | from_6974 | to_6974 | art_long | art_long | M |
| 118 | 129 | 0.654211624 | cat4 | from_6975 | to_6975 | art_long | art_long | M |
| 26 | 130 | 0.93982474 | cat4 | from_6976 | to_6976 | art_long | art_long | M |
| 28 | 130 | 0.601203705 | cat4 | from_6977 | to_6977 | art_long | art_long | M |
| 37 | 130 | 0.963819084 | cat4 | from_6978 | to_6978 | art_long | art_long | M |
| 59 | 130 | 0.601113056 | cat4 | from_6979 | to_6979 | art_long | art_long | M |
| 97 | 130 | 0.597055306 | cat4 | from_6980 | to_6980 | art_long | art_long | M |
| 115 | 130 | 0.808547413 | cat4 | from_6981 | to_6981 | art_long | art_long | M |
| 118 | 130 | 0.761623876 | cat4 | from_6982 | to_6982 | art_long | art_long | M |
| 129 | 130 | 0.943286665 | cat4 | from_6983 | to_6983 | art_long | art_long | M |
| 6 | 131 | -0.640297239 | cat4 | from_6984 | to_6984 | art_long | art_long | M |
| 20 | 131 | -0.606989214 | cat4 | from_6985 | to_6985 | art_long | art_long | M |
| 55 | 131 | 0.641413024 | cat4 | from_6986 | to_6986 | art_long | art_long | M |
| 95 | 131 | 0.725554698 | cat4 | from_6987 | to_6987 | art_long | art_long | M |
| 109 | 131 | 0.654393613 | cat4 | from_6988 | to_6988 | art_long | art_long | M |
| 118 | 131 | 0.917745741 | cat4 | from_6989 | to_6989 | art_long | art_long | M |
| 119 | 131 | 0.620789873 | cat4 | from_6990 | to_6990 | art_long | art_long | M |
| 129 | 131 | 0.588770754 | cat4 | from_6991 | to_6991 | art_long | art_long | M |
| 130 | 131 | 0.658231009 | cat4 | from_6992 | to_6992 | art_long | art_long | M |
| 17 | 132 | 0.739238477 | cat4 | from_6993 | to_6993 | art_long | art_long | M |
| 35 | 132 | 0.94599588 | cat4 | from_6994 | to_6994 | art_long | art_long | M |
| 36 | 132 | 0.671695056 | cat4 | from_6995 | to_6995 | art_long | art_long | M |
| 38 | 132 | 0.970300521 | cat4 | from_6996 | to_6996 | art_long | art_long | M |
| 44 | 132 | 0.674677891 | cat4 | from_6997 | to_6997 | art_long | art_long | M |
| 46 | 132 | 0.648016538 | cat4 | from_6998 | to_6998 | art_long | art_long | M |
| 48 | 132 | 0.876143881 | cat4 | from_6999 | to_6999 | art_long | art_long | M |
| 50 | 132 | 0.740197266 | cat4 | from_7000 | to_7000 | art_long | art_long | M |
| 52 | 132 | 0.580553141 | cat4 | from_7001 | to_7001 | art_long | art_long | M |
| 60 | 132 | 0.624171755 | cat4 | from_7002 | to_7002 | art_long | art_long | M |
| 104 | 132 | 0.81068463 | cat4 | from_7003 | to_7003 | art_long | art_long | M |
| 106 | 132 | 0.658326108 | cat4 | from_7004 | to_7004 | art_long | art_long | M |
| 128 | 132 | 0.65308479 | cat4 | from_7005 | to_7005 | art_long | art_long | M |
| 17 | 133 | 0.747278766 | cat4 | from_7006 | to_7006 | art_long | art_long | M |
| 35 | 133 | 0.72405969 | cat4 | from_7007 | to_7007 | art_long | art_long | M |
| 36 | 133 | 0.582724628 | cat4 | from_7008 | to_7008 | art_long | art_long | M |
| 38 | 133 | 0.729724832 | cat4 | from_7009 | to_7009 | art_long | art_long | M |
| 42 | 133 | 0.710582702 | cat4 | from_7010 | to_7010 | art_long | art_long | M |
| 45 | 133 | 0.599588345 | cat4 | from_7011 | to_7011 | art_long | art_long | M |
| 54 | 133 | -0.640728676 | cat4 | from_7012 | to_7012 | art_long | art_long | M |
| 62 | 133 | 0.595827847 | cat4 | from_7013 | to_7013 | art_long | art_long | M |
| 77 | 133 | -0.611038055 | cat4 | from_7014 | to_7014 | art_long | art_long | M |
| 80 | 133 | -0.647035774 | cat4 | from_7015 | to_7015 | art_long | art_long | M |
| 92 | 133 | 0.752967626 | cat4 | from_7016 | to_7016 | art_long | art_long | M |
| 101 | 133 | 0.665529739 | cat4 | from_7017 | to_7017 | art_long | art_long | M |
| 115 | 133 | 0.598756138 | cat4 | from_7018 | to_7018 | art_long | art_long | M |
| 132 | 133 | 0.681155666 | cat4 | from_7019 | to_7019 | art_long | art_long | M |
| 8 | 134 | -0.599721315 | cat4 | from_7020 | to_7020 | art_long | art_long | M |
| 17 | 134 | 0.848955492 | cat4 | from_7021 | to_7021 | art_long | art_long | M |
| 35 | 134 | 0.741983339 | cat4 | from_7022 | to_7022 | art_long | art_long | M |
| 36 | 134 | 0.739408988 | cat4 | from_7023 | to_7023 | art_long | art_long | M |
| 38 | 134 | 0.678641952 | cat4 | from_7024 | to_7024 | art_long | art_long | M |
| 45 | 134 | 0.653286171 | cat4 | from_7025 | to_7025 | art_long | art_long | M |
| 48 | 134 | 0.812201751 | cat4 | from_7026 | to_7026 | art_long | art_long | M |
| 62 | 134 | 0.599663139 | cat4 | from_7027 | to_7027 | art_long | art_long | M |
| 68 | 134 | -0.592583192 | cat4 | from_7028 | to_7028 | art_long | art_long | M |
| 81 | 134 | -0.631146282 | cat4 | from_7029 | to_7029 | art_long | art_long | M |
| 83 | 134 | -0.608608719 | cat4 | from_7030 | to_7030 | art_long | art_long | M |
| 100 | 134 | 0.689800746 | cat4 | from_7031 | to_7031 | art_long | art_long | M |
| 104 | 134 | 0.76116521 | cat4 | from_7032 | to_7032 | art_long | art_long | M |
| 132 | 134 | 0.645442879 | cat4 | from_7033 | to_7033 | art_long | art_long | M |
| 133 | 134 | 0.638290131 | cat4 | from_7034 | to_7034 | art_long | art_long | M |
| 44 | 135 | 0.625374069 | cat4 | from_7035 | to_7035 | art_long | art_long | M |
| 46 | 135 | 0.620719786 | cat4 | from_7036 | to_7036 | art_long | art_long | M |
| 49 | 135 | -0.665030186 | cat4 | from_7037 | to_7037 | art_long | art_long | M |
| 61 | 135 | 0.808384215 | cat4 | from_7038 | to_7038 | art_long | art_long | M |
| 90 | 135 | -0.636779681 | cat4 | from_7039 | to_7039 | art_long | art_long | M |
| 104 | 135 | 0.687614327 | cat4 | from_7040 | to_7040 | art_long | art_long | M |
| 106 | 135 | 0.739540099 | cat4 | from_7041 | to_7041 | art_long | art_long | M |
| 115 | 135 | -0.645179233 | cat4 | from_7042 | to_7042 | art_long | art_long | M |
| 128 | 135 | 0.679868129 | cat4 | from_7043 | to_7043 | art_long | art_long | M |
| 11 | 136 | -0.590777453 | cat4 | from_7044 | to_7044 | art_long | art_long | M |
| 14 | 136 | 0.57760093 | cat4 | from_7045 | to_7045 | art_long | art_long | M |
| 26 | 136 | 0.789020106 | cat4 | from_7046 | to_7046 | art_long | art_long | M |
| 37 | 136 | 0.781472986 | cat4 | from_7047 | to_7047 | art_long | art_long | M |
| 59 | 136 | 0.597241664 | cat4 | from_7048 | to_7048 | art_long | art_long | M |
| 73 | 136 | 0.590542334 | cat4 | from_7049 | to_7049 | art_long | art_long | M |
| 74 | 136 | 0.641906826 | cat4 | from_7050 | to_7050 | art_long | art_long | M |
| 78 | 136 | 0.588062355 | cat4 | from_7051 | to_7051 | art_long | art_long | M |
| 108 | 136 | 0.697053273 | cat4 | from_7052 | to_7052 | art_long | art_long | M |
| 110 | 136 | 0.591081271 | cat4 | from_7053 | to_7053 | art_long | art_long | M |
| 118 | 136 | 0.712495547 | cat4 | from_7054 | to_7054 | art_long | art_long | M |
| 129 | 136 | 0.826632862 | cat4 | from_7055 | to_7055 | art_long | art_long | M |
| 130 | 136 | 0.904038031 | cat4 | from_7056 | to_7056 | art_long | art_long | M |
| 131 | 136 | 0.645064835 | cat4 | from_7057 | to_7057 | art_long | art_long | M |
| 54 | 137 | 0.692859302 | cat4 | from_7058 | to_7058 | art_long | art_long | M |
| 58 | 137 | 0.628199307 | cat4 | from_7059 | to_7059 | art_long | art_long | M |
| 69 | 137 | 0.622721348 | cat4 | from_7060 | to_7060 | art_long | art_long | M |
| 77 | 137 | 0.680596897 | cat4 | from_7061 | to_7061 | art_long | art_long | M |
| 80 | 137 | 0.606101114 | cat4 | from_7062 | to_7062 | art_long | art_long | M |
| 81 | 137 | 0.677737908 | cat4 | from_7063 | to_7063 | art_long | art_long | M |
| 83 | 137 | 0.658352782 | cat4 | from_7064 | to_7064 | art_long | art_long | M |
| 100 | 137 | -0.609123329 | cat4 | from_7065 | to_7065 | art_long | art_long | M |
| 103 | 137 | -0.60443868 | cat4 | from_7066 | to_7066 | art_long | art_long | M |
| 113 | 137 | -0.582714272 | cat4 | from_7067 | to_7067 | art_long | art_long | M |
| 118 | 137 | 0.800900303 | cat4 | from_7068 | to_7068 | art_long | art_long | M |
| 119 | 137 | 0.59207778 | cat4 | from_7069 | to_7069 | art_long | art_long | M |
| 120 | 137 | 0.753030612 | cat4 | from_7070 | to_7070 | art_long | art_long | M |

| | | | | | | | | |
|---|---|---|---|---|---|---|---|---|
| 130 | 137 | 0.598786334 | cat4 | from_7071 | to_7071 | art_long | art_long | M |
| 131 | 137 | 0.690228389 | cat4 | from_7072 | to_7072 | art_long | art_long | M |
| 136 | 137 | 0.778986143 | cat4 | from_7073 | to_7073 | art_long | art_long | M |
| 17 | 138 | 0.978687328 | cat4 | from_7074 | to_7074 | art_long | art_long | M |
| 35 | 138 | 0.670694419 | cat4 | from_7075 | to_7075 | art_long | art_long | M |
| 36 | 138 | 0.964228581 | cat4 | from_7076 | to_7076 | art_long | art_long | M |
| 38 | 138 | 0.708695033 | cat4 | from_7077 | to_7077 | art_long | art_long | M |
| 48 | 138 | 0.722032969 | cat4 | from_7078 | to_7078 | art_long | art_long | M |
| 91 | 138 | -0.579364378 | cat4 | from_7079 | to_7079 | art_long | art_long | M |
| 104 | 138 | 0.75739667 | cat4 | from_7080 | to_7080 | art_long | art_long | M |
| 107 | 138 | 0.645397671 | cat4 | from_7081 | to_7081 | art_long | art_long | M |
| 108 | 138 | 0.610722792 | cat4 | from_7082 | to_7082 | art_long | art_long | M |
| 132 | 138 | 0.692378223 | cat4 | from_7083 | to_7083 | art_long | art_long | M |
| 133 | 138 | 0.635419699 | cat4 | from_7084 | to_7084 | art_long | art_long | M |
| 134 | 138 | 0.86621001 | cat4 | from_7085 | to_7085 | art_long | art_long | M |
| 32 | 139 | 0.613339053 | cat4 | from_7086 | to_7086 | art_long | art_long | M |
| 42 | 139 | 0.64866811 | cat4 | from_7087 | to_7087 | art_long | art_long | M |
| 43 | 139 | 0.714986135 | cat4 | from_7088 | to_7088 | art_long | art_long | M |
| 45 | 139 | 0.810201533 | cat4 | from_7089 | to_7089 | art_long | art_long | M |
| 62 | 139 | 0.81828001 | cat4 | from_7090 | to_7090 | art_long | art_long | M |
| 91 | 139 | -0.614941516 | cat4 | from_7091 | to_7091 | art_long | art_long | M |
| 92 | 139 | 0.579459033 | cat4 | from_7092 | to_7092 | art_long | art_long | M |
| 100 | 139 | 0.80037569 | cat4 | from_7093 | to_7093 | art_long | art_long | M |
| 101 | 139 | 0.733340183 | cat4 | from_7094 | to_7094 | art_long | art_long | M |
| 107 | 139 | 0.801962172 | cat4 | from_7095 | to_7095 | art_long | art_long | M |
| 134 | 139 | 0.582513933 | cat4 | from_7096 | to_7096 | art_long | art_long | M |
| 135 | 139 | 0.640302985 | cat4 | from_7097 | to_7097 | art_long | art_long | M |
| 138 | 139 | 0.581203091 | cat4 | from_7098 | to_7098 | art_long | art_long | M |
| 14 | 140 | 0.589130238 | cat4 | from_7099 | to_7099 | art_long | art_long | M |
| 25 | 140 | 0.679944123 | cat4 | from_7100 | to_7100 | art_long | art_long | M |
| 30 | 140 | 0.801147348 | cat4 | from_7101 | to_7101 | art_long | art_long | M |
| 54 | 140 | 0.869375242 | cat4 | from_7102 | to_7102 | art_long | art_long | M |
| 55 | 140 | 0.651612361 | cat4 | from_7103 | to_7103 | art_long | art_long | M |
| 58 | 140 | 0.911822683 | cat4 | from_7104 | to_7104 | art_long | art_long | M |
| 69 | 140 | 0.905865771 | cat4 | from_7105 | to_7105 | art_long | art_long | M |
| 72 | 140 | 0.738571309 | cat4 | from_7106 | to_7106 | art_long | art_long | M |
| 73 | 140 | 0.591237068 | cat4 | from_7107 | to_7107 | art_long | art_long | M |
| 74 | 140 | 0.601458958 | cat4 | from_7108 | to_7108 | art_long | art_long | M |
| 77 | 140 | 0.923235683 | cat4 | from_7109 | to_7109 | art_long | art_long | M |
| 80 | 140 | 0.861923149 | cat4 | from_7110 | to_7110 | art_long | art_long | M |
| 81 | 140 | 0.898336069 | cat4 | from_7111 | to_7111 | art_long | art_long | M |
| 83 | 140 | 0.968104759 | cat4 | from_7112 | to_7112 | art_long | art_long | M |
| 87 | 140 | 0.66479547 | cat4 | from_7113 | to_7113 | art_long | art_long | M |
| 95 | 140 | 0.654158548 | cat4 | from_7114 | to_7114 | art_long | art_long | M |
| 109 | 140 | 0.725889253 | cat4 | from_7115 | to_7115 | art_long | art_long | M |
| 110 | 140 | 0.609781792 | cat4 | from_7116 | to_7116 | art_long | art_long | M |
| 112 | 140 | 0.689080531 | cat4 | from_7117 | to_7117 | art_long | art_long | M |
| 119 | 140 | 0.878528511 | cat4 | from_7118 | to_7118 | art_long | art_long | M |
| 120 | 140 | 0.846659671 | cat4 | from_7119 | to_7119 | art_long | art_long | M |
| 137 | 140 | 0.692793772 | cat4 | from_7120 | to_7120 | art_long | art_long | M |
| 26 | 141 | 0.594126178 | cat4 | from_7121 | to_7121 | art_long | art_long | M |
| 37 | 141 | 0.617578364 | cat4 | from_7122 | to_7122 | art_long | art_long | M |
| 54 | 141 | 0.582614133 | cat4 | from_7123 | to_7123 | art_long | art_long | M |
| 58 | 141 | 0.667677291 | cat4 | from_7124 | to_7124 | art_long | art_long | M |
| 69 | 141 | 0.675465478 | cat4 | from_7125 | to_7125 | art_long | art_long | M |
| 77 | 141 | 0.590323346 | cat4 | from_7126 | to_7126 | art_long | art_long | M |
| 81 | 141 | 0.629732772 | cat4 | from_7127 | to_7127 | art_long | art_long | M |
| 83 | 141 | 0.678369577 | cat4 | from_7128 | to_7128 | art_long | art_long | M |
| 100 | 141 | -0.590326566 | cat4 | from_7129 | to_7129 | art_long | art_long | M |
| 113 | 141 | -0.594325101 | cat4 | from_7130 | to_7130 | art_long | art_long | M |
| 118 | 141 | 0.855953731 | cat4 | from_7131 | to_7131 | art_long | art_long | M |
| 119 | 141 | 0.582895702 | cat4 | from_7132 | to_7132 | art_long | art_long | M |
| 120 | 141 | 0.625979162 | cat4 | from_7133 | to_7133 | art_long | art_long | M |
| 129 | 141 | 0.603233219 | cat4 | from_7134 | to_7134 | art_long | art_long | M |
| 130 | 141 | 0.679434045 | cat4 | from_7135 | to_7135 | art_long | art_long | M |
| 131 | 141 | 0.674088258 | cat4 | from_7136 | to_7136 | art_long | art_long | M |
| 136 | 141 | 0.767401934 | cat4 | from_7137 | to_7137 | art_long | art_long | M |
| 137 | 141 | 0.966539408 | cat4 | from_7138 | to_7138 | art_long | art_long | M |
| 140 | 141 | 0.687713785 | cat4 | from_7139 | to_7139 | art_long | art_long | M |
| 14 | 142 | 0.592302277 | cat4 | from_7140 | to_7140 | art_long | art_long | M |
| 25 | 142 | 0.670631598 | cat4 | from_7141 | to_7141 | art_long | art_long | M |
| 26 | 142 | 0.727464066 | cat4 | from_7142 | to_7142 | art_long | art_long | M |
| 31 | 142 | 0.614337036 | cat4 | from_7143 | to_7143 | art_long | art_long | M |
| 33 | 142 | 0.614511985 | cat4 | from_7144 | to_7144 | art_long | art_long | M |
| 37 | 142 | 0.792551484 | cat4 | from_7145 | to_7145 | art_long | art_long | M |
| 73 | 142 | 0.596033545 | cat4 | from_7146 | to_7146 | art_long | art_long | M |
| 102 | 142 | -0.602866873 | cat4 | from_7147 | to_7147 | art_long | art_long | M |
| 108 | 142 | 0.683206101 | cat4 | from_7148 | to_7148 | art_long | art_long | M |
| 110 | 142 | 0.610755363 | cat4 | from_7149 | to_7149 | art_long | art_long | M |
| 112 | 142 | 0.714190657 | cat4 | from_7150 | to_7150 | art_long | art_long | M |
| 118 | 142 | 0.606505289 | cat4 | from_7151 | to_7151 | art_long | art_long | M |
| 129 | 142 | 0.837046572 | cat4 | from_7152 | to_7152 | art_long | art_long | M |
| 130 | 142 | 0.85958055 | cat4 | from_7153 | to_7153 | art_long | art_long | M |
| 136 | 142 | 0.926148812 | cat4 | from_7154 | to_7154 | art_long | art_long | M |
| 137 | 142 | 0.76413501 | cat4 | from_7155 | to_7155 | art_long | art_long | M |
| 141 | 142 | 0.800782206 | cat4 | from_7156 | to_7156 | art_long | art_long | M |
| 17 | 143 | 0.668447988 | cat4 | from_7157 | to_7157 | art_long | art_long | M |
| 35 | 143 | 0.785622611 | cat4 | from_7158 | to_7158 | art_long | art_long | M |
| 36 | 143 | 0.680011336 | cat4 | from_7159 | to_7159 | art_long | art_long | M |
| 38 | 143 | 0.827058295 | cat4 | from_7160 | to_7160 | art_long | art_long | M |
| 48 | 143 | 0.86999955 | cat4 | from_7161 | to_7161 | art_long | art_long | M |
| 50 | 143 | 0.769819823 | cat4 | from_7162 | to_7162 | art_long | art_long | M |
| 90 | 143 | -0.60159736 | cat4 | from_7163 | to_7163 | art_long | art_long | M |
| 104 | 143 | 0.911545017 | cat4 | from_7164 | to_7164 | art_long | art_long | M |
| 117 | 143 | -0.599200499 | cat4 | from_7165 | to_7165 | art_long | art_long | M |
| 132 | 143 | 0.808512476 | cat4 | from_7166 | to_7166 | art_long | art_long | M |
| 134 | 143 | 0.611358781 | cat4 | from_7167 | to_7167 | art_long | art_long | M |
| 135 | 143 | 0.590262935 | cat4 | from_7168 | to_7168 | art_long | art_long | M |
| 138 | 143 | 0.737948433 | cat4 | from_7169 | to_7169 | art_long | art_long | M |
| 20 | 144 | -0.589566844 | cat4 | from_7170 | to_7170 | art_long | art_long | M |
| 26 | 144 | 0.633749582 | cat4 | from_7171 | to_7171 | art_long | art_long | M |
| 37 | 144 | 0.598306997 | cat4 | from_7172 | to_7172 | art_long | art_long | M |
| 58 | 144 | 0.637426243 | cat4 | from_7173 | to_7173 | art_long | art_long | M |
| 69 | 144 | 0.609034255 | cat4 | from_7174 | to_7174 | art_long | art_long | M |

| | | | | | | | | |
|---|---|---|---|---|---|---|---|---|
| 77 | 144 | 0.589670343 | cat4 | from_7175 | to_7175 | art_long | art_long | M |
| 95 | 144 | 0.603086181 | cat4 | from_7176 | to_7176 | art_long | art_long | M |
| 109 | 144 | 0.59376305 | cat4 | from_7177 | to_7177 | art_long | art_long | M |
| 118 | 144 | 0.968240567 | cat4 | from_7178 | to_7178 | art_long | art_long | M |
| 119 | 144 | 0.676959224 | cat4 | from_7179 | to_7179 | art_long | art_long | M |
| 129 | 144 | 0.577001028 | cat4 | from_7180 | to_7180 | art_long | art_long | M |
| 130 | 144 | 0.701417337 | cat4 | from_7181 | to_7181 | art_long | art_long | M |
| 131 | 144 | 0.920819582 | cat4 | from_7182 | to_7182 | art_long | art_long | M |
| 136 | 144 | 0.752257726 | cat4 | from_7183 | to_7183 | art_long | art_long | M |
| 137 | 144 | 0.888817529 | cat4 | from_7184 | to_7184 | art_long | art_long | M |
| 141 | 144 | 0.884653575 | cat4 | from_7185 | to_7185 | art_long | art_long | M |
| 142 | 144 | 0.620667978 | cat4 | from_7186 | to_7186 | art_long | art_long | M |
| 8 | 145 | 0.823578181 | cat4 | from_7187 | to_7187 | art_long | art_long | M |
| 12 | 145 | 0.615054381 | cat4 | from_7188 | to_7188 | art_long | art_long | M |
| 19 | 145 | 0.611117521 | cat4 | from_7189 | to_7189 | art_long | art_long | M |
| 54 | 145 | 0.635253076 | cat4 | from_7190 | to_7190 | art_long | art_long | M |
| 64 | 145 | -0.623381564 | cat4 | from_7191 | to_7191 | art_long | art_long | M |
| 79 | 145 | 0.731412892 | cat4 | from_7192 | to_7192 | art_long | art_long | M |
| 81 | 145 | 0.73776846 | cat4 | from_7193 | to_7193 | art_long | art_long | M |
| 93 | 145 | 0.851695075 | cat4 | from_7194 | to_7194 | art_long | art_long | M |
| 100 | 145 | -0.682977083 | cat4 | from_7195 | to_7195 | art_long | art_long | M |
| 103 | 145 | -0.700348135 | cat4 | from_7196 | to_7196 | art_long | art_long | M |
| 120 | 145 | 0.601482943 | cat4 | from_7197 | to_7197 | art_long | art_long | M |
| 134 | 145 | -0.612326586 | cat4 | from_7198 | to_7198 | art_long | art_long | M |
| 49 | 146 | 0.710809047 | cat4 | from_7199 | to_7199 | art_long | art_long | M |
| 56 | 146 | 0.750003889 | cat4 | from_7200 | to_7200 | art_long | art_long | M |
| 57 | 146 | 0.720539 | cat4 | from_7201 | to_7201 | art_long | art_long | M |
| 65 | 146 | 0.586187563 | cat4 | from_7202 | to_7202 | art_long | art_long | M |
| 70 | 146 | 0.896097847 | cat4 | from_7203 | to_7203 | art_long | art_long | M |
| 75 | 146 | 0.953273545 | cat4 | from_7204 | to_7204 | art_long | art_long | M |
| 84 | 146 | 0.71279745 | cat4 | from_7205 | to_7205 | art_long | art_long | M |
| 85 | 146 | 0.776514749 | cat4 | from_7206 | to_7206 | art_long | art_long | M |
| 94 | 146 | -0.580198604 | cat4 | from_7207 | to_7207 | art_long | art_long | M |
| 108 | 146 | 0.62462311 | cat4 | from_7208 | to_7208 | art_long | art_long | M |
| 123 | 146 | 0.938304492 | cat4 | from_7209 | to_7209 | art_long | art_long | M |
| 4 | 147 | 0.614997268 | cat4 | from_7210 | to_7210 | art_long | art_long | M |
| 21 | 147 | 0.671724211 | cat4 | from_7211 | to_7211 | art_long | art_long | M |
| 42 | 147 | 0.955862038 | cat4 | from_7212 | to_7212 | art_long | art_long | M |
| 43 | 147 | 0.685630913 | cat4 | from_7213 | to_7213 | art_long | art_long | M |
| 44 | 147 | 0.623241932 | cat4 | from_7214 | to_7214 | art_long | art_long | M |
| 45 | 147 | 0.854883737 | cat4 | from_7215 | to_7215 | art_long | art_long | M |
| 46 | 147 | 0.595088568 | cat4 | from_7216 | to_7216 | art_long | art_long | M |
| 52 | 147 | 0.580335418 | cat4 | from_7217 | to_7217 | art_long | art_long | M |
| 60 | 147 | 0.63865945 | cat4 | from_7218 | to_7218 | art_long | art_long | M |
| 62 | 147 | 0.893564569 | cat4 | from_7219 | to_7219 | art_long | art_long | M |
| 92 | 147 | 0.947296998 | cat4 | from_7220 | to_7220 | art_long | art_long | M |
| 99 | 147 | 0.605660933 | cat4 | from_7221 | to_7221 | art_long | art_long | M |
| 100 | 147 | 0.750813057 | cat4 | from_7222 | to_7222 | art_long | art_long | M |
| 101 | 147 | 0.951766954 | cat4 | from_7223 | to_7223 | art_long | art_long | M |
| 107 | 147 | 0.735978002 | cat4 | from_7224 | to_7224 | art_long | art_long | M |
| 128 | 147 | 0.59880225 | cat4 | from_7225 | to_7225 | art_long | art_long | M |
| 133 | 147 | 0.76360261 | cat4 | from_7226 | to_7226 | art_long | art_long | M |
| 43 | 148 | 0.610599677 | cat4 | from_7227 | to_7227 | art_long | art_long | M |
| 44 | 148 | 0.843468382 | cat4 | from_7228 | to_7228 | art_long | art_long | M |
| 46 | 148 | 0.78031371 | cat4 | from_7229 | to_7229 | art_long | art_long | M |
| 50 | 148 | 0.627167891 | cat4 | from_7230 | to_7230 | art_long | art_long | M |
| 52 | 148 | 0.680932979 | cat4 | from_7231 | to_7231 | art_long | art_long | M |
| 60 | 148 | 0.918055807 | cat4 | from_7232 | to_7232 | art_long | art_long | M |
| 61 | 148 | 0.83737029 | cat4 | from_7233 | to_7233 | art_long | art_long | M |
| 99 | 148 | 0.960204098 | cat4 | from_7234 | to_7234 | art_long | art_long | M |
| 102 | 148 | 0.590116495 | cat4 | from_7235 | to_7235 | art_long | art_long | M |
| 105 | 148 | 0.907871619 | cat4 | from_7236 | to_7236 | art_long | art_long | M |
| 106 | 148 | 0.818038269 | cat4 | from_7237 | to_7237 | art_long | art_long | M |
| 128 | 148 | 0.888181228 | cat4 | from_7238 | to_7238 | art_long | art_long | M |
| 4 | 149 | 0.626794052 | cat4 | from_7239 | to_7239 | art_long | art_long | M |
| 7 | 149 | 0.60477783 | cat4 | from_7240 | to_7240 | art_long | art_long | M |
| 17 | 149 | 0.663128956 | cat4 | from_7241 | to_7241 | art_long | art_long | M |
| 36 | 149 | 0.708990212 | cat4 | from_7242 | to_7242 | art_long | art_long | M |
| 42 | 149 | 0.693070229 | cat4 | from_7243 | to_7243 | art_long | art_long | M |
| 43 | 149 | 0.935642831 | cat4 | from_7244 | to_7244 | art_long | art_long | M |
| 45 | 149 | 0.782389639 | cat4 | from_7245 | to_7245 | art_long | art_long | M |
| 60 | 149 | 0.627234521 | cat4 | from_7246 | to_7246 | art_long | art_long | M |
| 62 | 149 | 0.841486708 | cat4 | from_7247 | to_7247 | art_long | art_long | M |
| 92 | 149 | 0.64677562 | cat4 | from_7248 | to_7248 | art_long | art_long | M |
| 100 | 149 | 0.626517501 | cat4 | from_7249 | to_7249 | art_long | art_long | M |
| 101 | 149 | 0.828702404 | cat4 | from_7250 | to_7250 | art_long | art_long | M |
| 107 | 149 | 0.998339243 | cat4 | from_7251 | to_7251 | art_long | art_long | M |
| 128 | 149 | 0.641963399 | cat4 | from_7252 | to_7252 | art_long | art_long | M |
| 138 | 149 | 0.638598605 | cat4 | from_7253 | to_7253 | art_long | art_long | M |
| 139 | 149 | 0.788065808 | cat4 | from_7254 | to_7254 | art_long | art_long | M |
| 147 | 149 | 0.723078465 | cat4 | from_7255 | to_7255 | art_long | art_long | M |
| 3 | 150 | 0.595513963 | cat4 | from_7256 | to_7256 | art_long | art_long | M |
| 29 | 150 | 0.824837781 | cat4 | from_7257 | to_7257 | art_long | art_long | M |
| 41 | 150 | 0.663636367 | cat4 | from_7258 | to_7258 | art_long | art_long | M |
| 66 | 150 | 0.6530059 | cat4 | from_7259 | to_7259 | art_long | art_long | M |
| 67 | 150 | 0.71595533 | cat4 | from_7260 | to_7260 | art_long | art_long | M |
| 71 | 150 | 0.856189874 | cat4 | from_7261 | to_7261 | art_long | art_long | M |
| 82 | 150 | 0.938886601 | cat4 | from_7262 | to_7262 | art_long | art_long | M |
| 98 | 150 | 0.807639183 | cat4 | from_7263 | to_7263 | art_long | art_long | M |
| 121 | 150 | 0.783514385 | cat4 | from_7264 | to_7264 | art_long | art_long | M |
| 124 | 150 | 0.672750759 | cat4 | from_7265 | to_7265 | art_long | art_long | M |
| 1 | 151 | -0.708933894 | cat4 | from_7266 | to_7266 | art_long | art_long | M |
| 6 | 151 | 0.583253878 | cat4 | from_7267 | to_7267 | art_long | art_long | M |
| 93 | 152 | 0.581192527 | cat4 | from_7268 | to_7268 | art_long | art_long | M |
| 99 | 152 | 0.608736352 | cat4 | from_7269 | to_7269 | art_long | art_long | M |
| 105 | 152 | 0.708825908 | cat4 | from_7270 | to_7270 | art_long | art_long | M |
| 148 | 152 | 0.710794736 | cat4 | from_7271 | to_7271 | art_long | art_long | M |
| 89 | 153 | 0.604557467 | cat4 | from_7272 | to_7272 | art_long | art_long | M |
| 99 | 153 | 0.61399937 | cat4 | from_7273 | to_7273 | art_long | art_long | M |
| 105 | 153 | 0.669667306 | cat4 | from_7274 | to_7274 | art_long | art_long | M |
| 148 | 153 | 0.707575875 | cat4 | from_7275 | to_7275 | art_long | art_long | M |
| 152 | 153 | 0.984391642 | cat4 | from_7276 | to_7276 | art_long | art_long | M |
| 17 | 154 | 0.789203065 | cat4 | from_7277 | to_7277 | art_long | art_short | J |
| 36 | 154 | 0.846282327 | cat4 | from_7278 | to_7278 | art_long | art_short | J |

| | | | | | | | | |
|---|---|---|---|---|---|---|---|---|
| 47 | 154 | -0.61135778 | cat4 | from_7279 | to_7279 | art_long | art_short | J |
| 72 | 154 | -0.622851257 | cat4 | from_7280 | to_7280 | art_long | art_short | J |
| 91 | 154 | -0.61092095 | cat4 | from_7281 | to_7281 | art_long | art_short | J |
| 104 | 154 | 0.62962167 | cat4 | from_7282 | to_7282 | art_long | art_short | J |
| 108 | 154 | 0.621610339 | cat4 | from_7283 | to_7283 | art_long | art_short | J |
| 111 | 154 | -0.624126878 | cat4 | from_7284 | to_7284 | art_long | art_short | J |
| 134 | 154 | 0.612905553 | cat4 | from_7285 | to_7285 | art_long | art_short | J |
| 138 | 154 | 0.87926357 | cat4 | from_7286 | to_7286 | art_long | art_short | J |
| 143 | 154 | 0.766743519 | cat4 | from_7287 | to_7287 | art_long | art_short | J |
| 2 | 155 | 0.971116898 | cat4 | from_7288 | to_7288 | art_long | art_short | J |
| 6 | 155 | -0.617176908 | cat4 | from_7289 | to_7289 | art_long | art_short | J |
| 16 | 155 | 0.585208709 | cat4 | from_7290 | to_7290 | art_long | art_short | J |
| 51 | 155 | -0.590359806 | cat4 | from_7291 | to_7291 | art_long | art_short | J |
| 53 | 155 | 0.713904057 | cat4 | from_7292 | to_7292 | art_long | art_short | J |
| 110 | 155 | -0.600155667 | cat4 | from_7293 | to_7293 | art_long | art_short | J |
| 112 | 155 | -0.603999108 | cat4 | from_7294 | to_7294 | art_long | art_short | J |
| 8 | 156 | 0.698815801 | cat4 | from_7295 | to_7295 | art_long | art_short | J |
| 9 | 156 | 0.602332601 | cat4 | from_7296 | to_7296 | art_long | art_short | J |
| 10 | 156 | 0.608281768 | cat4 | from_7297 | to_7297 | art_long | art_short | J |
| 12 | 156 | 0.584692228 | cat4 | from_7298 | to_7298 | art_long | art_short | J |
| 13 | 156 | 0.654368272 | cat4 | from_7299 | to_7299 | art_long | art_short | J |
| 19 | 156 | 0.616497114 | cat4 | from_7300 | to_7300 | art_long | art_short | J |
| 35 | 156 | -0.580398744 | cat4 | from_7301 | to_7301 | art_long | art_short | J |
| 63 | 156 | 0.580368607 | cat4 | from_7302 | to_7302 | art_long | art_short | J |
| 79 | 156 | 0.628276541 | cat4 | from_7303 | to_7303 | art_long | art_short | J |
| 114 | 156 | 0.807440539 | cat4 | from_7304 | to_7304 | art_long | art_short | J |
| 116 | 156 | 0.580187508 | cat4 | from_7305 | to_7305 | art_long | art_short | J |
| 125 | 156 | 0.71798491 | cat4 | from_7306 | to_7306 | art_long | art_short | J |
| 132 | 156 | -0.601800778 | cat4 | from_7307 | to_7307 | art_long | art_short | J |
| 6 | 157 | 0.845165063 | cat4 | from_7308 | to_7308 | art_long | art_short | J |
| 8 | 157 | 0.833501576 | cat4 | from_7309 | to_7309 | art_long | art_short | J |
| 9 | 157 | 0.643525235 | cat4 | from_7310 | to_7310 | art_long | art_short | J |
| 10 | 157 | 0.633025312 | cat4 | from_7311 | to_7311 | art_long | art_short | J |
| 84 | 157 | 0.639023312 | cat4 | from_7312 | to_7312 | art_long | art_short | J |
| 125 | 157 | 0.709068497 | cat4 | from_7313 | to_7313 | art_long | art_short | J |
| 145 | 157 | 0.64513662 | cat4 | from_7314 | to_7314 | art_long | art_short | J |
| 156 | 157 | 0.685359153 | cat4 | from_7315 | to_7315 | art_short | art_short | H |
| 8 | 158 | 0.70193704 | cat4 | from_7316 | to_7316 | art_long | art_short | J |
| 9 | 158 | 0.590794138 | cat4 | from_7317 | to_7317 | art_long | art_short | J |
| 10 | 158 | 0.597723035 | cat4 | from_7318 | to_7318 | art_long | art_short | J |
| 13 | 158 | 0.64349786 | cat4 | from_7319 | to_7319 | art_long | art_short | J |
| 19 | 158 | 0.597789473 | cat4 | from_7320 | to_7320 | art_long | art_short | J |
| 48 | 158 | -0.604418452 | cat4 | from_7321 | to_7321 | art_long | art_short | J |
| 63 | 158 | 0.615689618 | cat4 | from_7322 | to_7322 | art_long | art_short | J |
| 79 | 158 | 0.614612922 | cat4 | from_7323 | to_7323 | art_long | art_short | J |
| 84 | 158 | 0.591104777 | cat4 | from_7324 | to_7324 | art_long | art_short | J |
| 104 | 158 | -0.578132247 | cat4 | from_7325 | to_7325 | art_long | art_short | J |
| 114 | 158 | 0.808644649 | cat4 | from_7326 | to_7326 | art_long | art_short | J |
| 116 | 158 | 0.577503642 | cat4 | from_7327 | to_7327 | art_long | art_short | J |
| 125 | 158 | 0.745868346 | cat4 | from_7328 | to_7328 | art_long | art_short | J |
| 131 | 158 | -0.576674867 | cat4 | from_7329 | to_7329 | art_long | art_short | J |
| 156 | 158 | 0.989784886 | cat4 | from_7330 | to_7330 | art_short | art_short | H |
| 157 | 158 | 0.684988676 | cat4 | from_7331 | to_7331 | art_short | art_short | H |
| 4 | 159 | 0.819768959 | cat4 | from_7332 | to_7332 | art_long | art_short | J |
| 6 | 159 | 0.770289673 | cat4 | from_7333 | to_7333 | art_long | art_short | J |
| 7 | 159 | 0.821781523 | cat4 | from_7334 | to_7334 | art_long | art_short | J |
| 11 | 159 | 0.700851141 | cat4 | from_7335 | to_7335 | art_long | art_short | J |
| 65 | 159 | 0.644075878 | cat4 | from_7336 | to_7336 | art_long | art_short | J |
| 113 | 159 | 0.611479469 | cat4 | from_7337 | to_7337 | art_long | art_short | J |
| 118 | 159 | -0.624809878 | cat4 | from_7338 | to_7338 | art_long | art_short | J |
| 137 | 159 | -0.625380596 | cat4 | from_7339 | to_7339 | art_long | art_short | J |
| 141 | 159 | -0.629786432 | cat4 | from_7340 | to_7340 | art_long | art_short | J |
| 144 | 159 | -0.653578028 | cat4 | from_7341 | to_7341 | art_long | art_short | J |
| 4 | 160 | 0.952298306 | cat4 | from_7342 | to_7342 | art_long | art_short | J |
| 6 | 160 | 0.713777384 | cat4 | from_7343 | to_7343 | art_long | art_short | J |
| 7 | 160 | 0.835215495 | cat4 | from_7344 | to_7344 | art_long | art_short | J |
| 11 | 160 | 0.734218506 | cat4 | from_7345 | to_7345 | art_long | art_short | J |
| 34 | 160 | 0.68899888 | cat4 | from_7346 | to_7346 | art_long | art_short | J |
| 107 | 160 | 0.586661602 | cat4 | from_7347 | to_7347 | art_long | art_short | J |
| 127 | 160 | 0.674301152 | cat4 | from_7348 | to_7348 | art_long | art_short | J |
| 131 | 160 | -0.588770158 | cat4 | from_7349 | to_7349 | art_long | art_short | J |
| 147 | 160 | 0.592766972 | cat4 | from_7350 | to_7350 | art_long | art_short | J |
| 149 | 160 | 0.583867666 | cat4 | from_7351 | to_7351 | art_long | art_short | J |
| 159 | 160 | 0.85126592 | cat4 | from_7352 | to_7352 | art_short | art_short | H |
| 8 | 161 | 0.672914243 | cat4 | from_7353 | to_7353 | art_long | art_short | J |
| 9 | 161 | 0.827264822 | cat4 | from_7354 | to_7354 | art_long | art_short | J |
| 10 | 161 | 0.806129082 | cat4 | from_7355 | to_7355 | art_long | art_short | J |
| 22 | 161 | 0.745885695 | cat4 | from_7356 | to_7356 | art_long | art_short | J |
| 23 | 161 | 0.824877127 | cat4 | from_7357 | to_7357 | art_long | art_short | J |
| 57 | 161 | 0.692142567 | cat4 | from_7358 | to_7358 | art_long | art_short | J |
| 88 | 161 | 0.805866638 | cat4 | from_7359 | to_7359 | art_long | art_short | J |
| 126 | 161 | 0.909256528 | cat4 | from_7360 | to_7360 | art_long | art_short | J |
| 2 | 162 | 0.658947889 | cat4 | from_7361 | to_7361 | art_long | art_short | J |
| 16 | 162 | 0.592037576 | cat4 | from_7362 | to_7362 | art_long | art_short | J |
| 116 | 162 | 0.585245787 | cat4 | from_7363 | to_7363 | art_long | art_short | J |
| 155 | 162 | 0.667213071 | cat4 | from_7364 | to_7364 | art_short | art_short | H |
| 12 | 163 | 0.591079316 | cat4 | from_7365 | to_7365 | art_long | art_short | J |
| 13 | 163 | 0.75935449 | cat4 | from_7366 | to_7366 | art_long | art_short | J |
| 15 | 163 | 0.635274091 | cat4 | from_7367 | to_7367 | art_long | art_short | J |
| 16 | 163 | 0.584179299 | cat4 | from_7368 | to_7368 | art_long | art_short | J |
| 19 | 163 | 0.586596573 | cat4 | from_7369 | to_7369 | art_long | art_short | J |
| 40 | 163 | 0.670134731 | cat4 | from_7370 | to_7370 | art_long | art_short | J |
| 114 | 163 | 0.724679482 | cat4 | from_7371 | to_7371 | art_long | art_short | J |
| 116 | 163 | 0.590018044 | cat4 | from_7372 | to_7372 | art_long | art_short | J |
| 162 | 163 | 0.927963382 | cat4 | from_7373 | to_7373 | art_short | art_short | H |
| 2 | 164 | 0.583128261 | cat4 | from_7374 | to_7374 | art_long | art_short | J |
| 12 | 164 | 0.671931916 | cat4 | from_7375 | to_7375 | art_long | art_short | J |
| 16 | 164 | 0.955789178 | cat4 | from_7376 | to_7376 | art_long | art_short | J |
| 18 | 164 | 0.689663963 | cat4 | from_7377 | to_7377 | art_long | art_short | J |
| 19 | 164 | 0.626011083 | cat4 | from_7378 | to_7378 | art_long | art_short | J |
| 39 | 164 | 0.903609548 | cat4 | from_7379 | to_7379 | art_long | art_short | J |
| 40 | 164 | 0.803886384 | cat4 | from_7380 | to_7380 | art_long | art_short | J |
| 53 | 164 | 0.726518427 | cat4 | from_7381 | to_7381 | art_long | art_short | J |
| 155 | 164 | 0.618511035 | cat4 | from_7382 | to_7382 | art_short | art_short | H |

| | | | | | | | | |
|---|---|---|---|---|---|---|---|---|
| 162 | 164 | 0.616998387 | cat4 | from_7383 | to_7383 | art_short | art_short | H |
| 163 | 164 | 0.650948973 | cat4 | from_7384 | to_7384 | art_short | art_short | H |
| 12 | 165 | 0.758281676 | cat4 | from_7385 | to_7385 | art_long | art_short | J |
| 13 | 165 | 0.898646443 | cat4 | from_7386 | to_7386 | art_long | art_short | J |
| 15 | 165 | 0.691430237 | cat4 | from_7387 | to_7387 | art_long | art_short | J |
| 18 | 165 | 0.597321952 | cat4 | from_7388 | to_7388 | art_long | art_short | J |
| 19 | 165 | 0.737437507 | cat4 | from_7389 | to_7389 | art_long | art_short | J |
| 20 | 165 | 0.70978619 | cat4 | from_7390 | to_7390 | art_long | art_short | J |
| 39 | 165 | 0.670584604 | cat4 | from_7391 | to_7391 | art_long | art_short | J |
| 40 | 165 | 0.654929998 | cat4 | from_7392 | to_7392 | art_long | art_short | J |
| 63 | 165 | 0.587219993 | cat4 | from_7393 | to_7393 | art_long | art_short | J |
| 68 | 165 | 0.755528431 | cat4 | from_7394 | to_7394 | art_long | art_short | J |
| 79 | 165 | 0.75520026 | cat4 | from_7395 | to_7395 | art_long | art_short | J |
| 114 | 165 | 0.962257236 | cat4 | from_7396 | to_7396 | art_long | art_short | J |
| 156 | 165 | 0.76750341 | cat4 | from_7397 | to_7397 | art_short | art_short | H |
| 158 | 165 | 0.787419003 | cat4 | from_7398 | to_7398 | art_short | art_short | H |
| 163 | 165 | 0.712434924 | cat4 | from_7399 | to_7399 | art_short | art_short | H |
| 14 | 166 | -0.580403986 | cat4 | from_7400 | to_7400 | art_long | art_short | J |
| 17 | 166 | 0.587149095 | cat4 | from_7401 | to_7401 | art_long | art_short | J |
| 25 | 166 | -0.603928636 | cat4 | from_7402 | to_7402 | art_long | art_short | J |
| 31 | 166 | -0.633277636 | cat4 | from_7403 | to_7403 | art_long | art_short | J |
| 33 | 166 | -0.622076828 | cat4 | from_7404 | to_7404 | art_long | art_short | J |
| 35 | 166 | 0.85761468 | cat4 | from_7405 | to_7405 | art_long | art_short | J |
| 38 | 166 | 0.806985774 | cat4 | from_7406 | to_7406 | art_long | art_short | J |
| 48 | 166 | 0.959387794 | cat4 | from_7407 | to_7407 | art_long | art_short | J |
| 50 | 166 | 0.587089656 | cat4 | from_7408 | to_7408 | art_long | art_short | J |
| 51 | 166 | -0.632504738 | cat4 | from_7409 | to_7409 | art_long | art_short | J |
| 63 | 166 | -0.609629759 | cat4 | from_7410 | to_7410 | art_long | art_short | J |
| 68 | 166 | -0.68231086 | cat4 | from_7411 | to_7411 | art_long | art_short | J |
| 102 | 166 | 0.632188946 | cat4 | from_7412 | to_7412 | art_long | art_short | J |
| 104 | 166 | 0.921030656 | cat4 | from_7413 | to_7413 | art_long | art_short | J |
| 110 | 166 | -0.614950063 | cat4 | from_7414 | to_7414 | art_long | art_short | J |
| 112 | 166 | -0.660043129 | cat4 | from_7415 | to_7415 | art_long | art_short | J |
| 132 | 166 | 0.786354438 | cat4 | from_7416 | to_7416 | art_long | art_short | J |
| 134 | 166 | 0.764432202 | cat4 | from_7417 | to_7417 | art_long | art_short | J |
| 138 | 166 | 0.652854436 | cat4 | from_7418 | to_7418 | art_long | art_short | J |
| 143 | 166 | 0.890652753 | cat4 | from_7419 | to_7419 | art_long | art_short | J |
| 12 | 167 | 0.995883556 | cat4 | from_7420 | to_7420 | art_long | art_short | J |
| 13 | 167 | 0.776727608 | cat4 | from_7421 | to_7421 | art_long | art_short | J |
| 15 | 167 | 0.640546104 | cat4 | from_7422 | to_7422 | art_long | art_short | J |
| 16 | 167 | 0.625931003 | cat4 | from_7423 | to_7423 | art_long | art_short | J |
| 18 | 167 | 0.779359194 | cat4 | from_7424 | to_7424 | art_long | art_short | J |
| 19 | 167 | 0.95530799 | cat4 | from_7425 | to_7425 | art_long | art_short | J |
| 39 | 167 | 0.722101093 | cat4 | from_7426 | to_7426 | art_long | art_short | J |
| 40 | 167 | 0.814922428 | cat4 | from_7427 | to_7427 | art_long | art_short | J |
| 68 | 167 | 0.590135012 | cat4 | from_7428 | to_7428 | art_long | art_short | J |
| 79 | 167 | 0.961701617 | cat4 | from_7429 | to_7429 | art_long | art_short | J |
| 114 | 167 | 0.725941355 | cat4 | from_7430 | to_7430 | art_long | art_short | J |
| 145 | 167 | 0.586152307 | cat4 | from_7431 | to_7431 | art_long | art_short | J |
| 156 | 167 | 0.577441302 | cat4 | from_7432 | to_7432 | art_short | art_short | H |
| 163 | 167 | 0.629667854 | cat4 | from_7433 | to_7433 | art_short | art_short | H |
| 164 | 167 | 0.676377773 | cat4 | from_7434 | to_7434 | art_short | art_short | H |
| 165 | 167 | 0.734205584 | cat4 | from_7435 | to_7435 | art_short | art_short | H |
| 27 | 168 | -0.592116949 | cat4 | from_7436 | to_7436 | art_long | art_short | J |
| 39 | 168 | 0.616300488 | cat4 | from_7437 | to_7437 | art_long | art_short | J |
| 53 | 168 | 0.779228374 | cat4 | from_7438 | to_7438 | art_long | art_short | J |
| 165 | 168 | 0.630976307 | cat4 | from_7439 | to_7439 | art_short | art_short | H |
| 13 | 169 | 0.72762393 | cat4 | from_7440 | to_7440 | art_long | art_short | J |
| 15 | 169 | 0.667823674 | cat4 | from_7441 | to_7441 | art_long | art_short | J |
| 16 | 169 | 0.688907556 | cat4 | from_7442 | to_7442 | art_long | art_short | J |
| 20 | 169 | 0.719059476 | cat4 | from_7443 | to_7443 | art_long | art_short | J |
| 21 | 169 | 0.707339388 | cat4 | from_7444 | to_7444 | art_long | art_short | J |
| 39 | 169 | 0.75494894 | cat4 | from_7445 | to_7445 | art_long | art_short | J |
| 40 | 169 | 0.638561285 | cat4 | from_7446 | to_7446 | art_long | art_short | J |
| 114 | 169 | 0.778358406 | cat4 | from_7447 | to_7447 | art_long | art_short | J |
| 116 | 169 | 0.635866602 | cat4 | from_7448 | to_7448 | art_long | art_short | J |
| 158 | 169 | 0.601665365 | cat4 | from_7449 | to_7449 | art_short | art_short | H |
| 162 | 169 | 0.634828624 | cat4 | from_7450 | to_7450 | art_short | art_short | H |
| 163 | 169 | 0.782899205 | cat4 | from_7451 | to_7451 | art_short | art_short | H |
| 164 | 169 | 0.755417294 | cat4 | from_7452 | to_7452 | art_short | art_short | H |
| 165 | 169 | 0.852519259 | cat4 | from_7453 | to_7453 | art_short | art_short | H |
| 168 | 169 | 0.739879865 | cat4 | from_7454 | to_7454 | art_short | art_short | H |
| 32 | 170 | 0.587578282 | cat4 | from_7455 | to_7455 | art_long | art_short | J |
| 42 | 170 | 0.627513836 | cat4 | from_7456 | to_7456 | art_long | art_short | J |
| 45 | 170 | 0.808459616 | cat4 | from_7457 | to_7457 | art_long | art_short | J |
| 62 | 170 | 0.767969116 | cat4 | from_7458 | to_7458 | art_long | art_short | J |
| 83 | 170 | -0.579928965 | cat4 | from_7459 | to_7459 | art_long | art_short | J |
| 91 | 170 | -0.614756528 | cat4 | from_7460 | to_7460 | art_long | art_short | J |
| 92 | 170 | 0.592532503 | cat4 | from_7461 | to_7461 | art_long | art_short | J |
| 100 | 170 | 0.841045124 | cat4 | from_7462 | to_7462 | art_long | art_short | J |
| 101 | 170 | 0.67132648 | cat4 | from_7463 | to_7463 | art_long | art_short | J |
| 102 | 170 | 0.576106621 | cat4 | from_7464 | to_7464 | art_long | art_short | J |
| 107 | 170 | 0.664021574 | cat4 | from_7465 | to_7465 | art_long | art_short | J |
| 134 | 170 | 0.632968324 | cat4 | from_7466 | to_7466 | art_long | art_short | J |
| 139 | 170 | 0.962721525 | cat4 | from_7467 | to_7467 | art_long | art_short | J |
| 141 | 170 | -0.611736103 | cat4 | from_7468 | to_7468 | art_long | art_short | J |
| 149 | 170 | 0.636991713 | cat4 | from_7469 | to_7469 | art_long | art_short | J |
| 32 | 171 | 0.57849456 | cat4 | from_7470 | to_7470 | art_long | art_short | J |
| 42 | 171 | 0.684907838 | cat4 | from_7471 | to_7471 | art_long | art_short | J |
| 43 | 171 | 0.601667103 | cat4 | from_7472 | to_7472 | art_long | art_short | J |
| 45 | 171 | 0.847978548 | cat4 | from_7473 | to_7473 | art_long | art_short | J |
| 62 | 171 | 0.810695482 | cat4 | from_7474 | to_7474 | art_long | art_short | J |
| 83 | 171 | -0.580016474 | cat4 | from_7475 | to_7475 | art_long | art_short | J |
| 91 | 171 | -0.60421699 | cat4 | from_7476 | to_7476 | art_long | art_short | J |
| 92 | 171 | 0.654911826 | cat4 | from_7477 | to_7477 | art_long | art_short | J |
| 100 | 171 | 0.868108473 | cat4 | from_7478 | to_7478 | art_long | art_short | J |
| 101 | 171 | 0.724765702 | cat4 | from_7479 | to_7479 | art_long | art_short | J |
| 102 | 171 | 0.589692337 | cat4 | from_7480 | to_7480 | art_long | art_short | J |
| 107 | 171 | 0.695722799 | cat4 | from_7481 | to_7481 | art_long | art_short | J |
| 134 | 171 | 0.634696261 | cat4 | from_7482 | to_7482 | art_long | art_short | J |
| 139 | 171 | 0.9637474 | cat4 | from_7483 | to_7483 | art_long | art_short | J |
| 141 | 171 | -0.619813381 | cat4 | from_7484 | to_7484 | art_long | art_short | J |
| 149 | 171 | 0.668152169 | cat4 | from_7485 | to_7485 | art_long | art_short | J |
| 170 | 171 | 0.996319844 | cat4 | from_7486 | to_7486 | art_short | art_short | H |

| | | | | | | | | |
|---|---|---|---|---|---|---|---|---|
| 22 | 172 | 0.666292457 | cat4 | from_7487 | to_7487 | art_long | art_short | J |
| 27 | 172 | 0.893365921 | cat4 | from_7488 | to_7488 | art_long | art_short | J |
| 30 | 172 | 0.600261895 | cat4 | from_7489 | to_7489 | art_long | art_short | J |
| 47 | 172 | 0.587117025 | cat4 | from_7490 | to_7490 | art_long | art_short | J |
| 55 | 172 | 0.618356066 | cat4 | from_7491 | to_7491 | art_long | art_short | J |
| 64 | 172 | 0.727511095 | cat4 | from_7492 | to_7492 | art_long | art_short | J |
| 87 | 172 | 0.612711964 | cat4 | from_7493 | to_7493 | art_long | art_short | J |
| 91 | 172 | 0.85119024 | cat4 | from_7494 | to_7494 | art_long | art_short | J |
| 109 | 172 | 0.622890349 | cat4 | from_7495 | to_7495 | art_long | art_short | J |
| 117 | 172 | 0.681993096 | cat4 | from_7496 | to_7496 | art_long | art_short | J |
| 14 | 173 | 0.596069753 | cat4 | from_7497 | to_7497 | art_long | art_short | J |
| 24 | 173 | 0.835338641 | cat4 | from_7498 | to_7498 | art_long | art_short | J |
| 25 | 173 | 0.593326486 | cat4 | from_7499 | to_7499 | art_long | art_short | J |
| 28 | 173 | 0.913856007 | cat4 | from_7500 | to_7500 | art_long | art_short | J |
| 47 | 173 | 0.66321722 | cat4 | from_7501 | to_7501 | art_long | art_short | J |
| 50 | 173 | -0.643987336 | cat4 | from_7502 | to_7502 | art_long | art_short | J |
| 55 | 173 | 0.672190838 | cat4 | from_7503 | to_7503 | art_long | art_short | J |
| 59 | 173 | 0.737988738 | cat4 | from_7504 | to_7504 | art_long | art_short | J |
| 64 | 173 | 0.642991702 | cat4 | from_7505 | to_7505 | art_long | art_short | J |
| 73 | 173 | 0.606871376 | cat4 | from_7506 | to_7506 | art_long | art_short | J |
| 76 | 173 | 0.847269974 | cat4 | from_7507 | to_7507 | art_long | art_short | J |
| 78 | 173 | 0.646676218 | cat4 | from_7508 | to_7508 | art_long | art_short | J |
| 87 | 173 | 0.643739085 | cat4 | from_7509 | to_7509 | art_long | art_short | J |
| 90 | 173 | 0.819247507 | cat4 | from_7510 | to_7510 | art_long | art_short | J |
| 95 | 173 | 0.616528523 | cat4 | from_7511 | to_7511 | art_long | art_short | J |
| 97 | 173 | 0.838557125 | cat4 | from_7512 | to_7512 | art_long | art_short | J |
| 109 | 173 | 0.651384595 | cat4 | from_7513 | to_7513 | art_long | art_short | J |
| 122 | 173 | 0.864532596 | cat4 | from_7514 | to_7514 | art_long | art_short | J |
| 14 | 174 | 0.853815764 | cat4 | from_7515 | to_7515 | art_long | art_short | J |
| 24 | 174 | 0.812811126 | cat4 | from_7516 | to_7516 | art_long | art_short | J |
| 25 | 174 | 0.767407116 | cat4 | from_7517 | to_7517 | art_long | art_short | J |
| 28 | 174 | 0.891587175 | cat4 | from_7518 | to_7518 | art_long | art_short | J |
| 30 | 174 | 0.600106196 | cat4 | from_7519 | to_7519 | art_long | art_short | J |
| 31 | 174 | 0.667660005 | cat4 | from_7520 | to_7520 | art_long | art_short | J |
| 33 | 174 | 0.693551156 | cat4 | from_7521 | to_7521 | art_long | art_short | J |
| 58 | 174 | 0.599896471 | cat4 | from_7522 | to_7522 | art_long | art_short | J |
| 59 | 174 | 0.909921547 | cat4 | from_7523 | to_7523 | art_long | art_short | J |
| 69 | 174 | 0.58995568 | cat4 | from_7524 | to_7524 | art_long | art_short | J |
| 73 | 174 | 0.860778539 | cat4 | from_7525 | to_7525 | art_long | art_short | J |
| 74 | 174 | 0.831803246 | cat4 | from_7526 | to_7526 | art_long | art_short | J |
| 76 | 174 | 0.865811464 | cat4 | from_7527 | to_7527 | art_long | art_short | J |
| 78 | 174 | 0.824395154 | cat4 | from_7528 | to_7528 | art_long | art_short | J |
| 90 | 174 | 0.701764904 | cat4 | from_7529 | to_7529 | art_long | art_short | J |
| 97 | 174 | 0.762382271 | cat4 | from_7530 | to_7530 | art_long | art_short | J |
| 110 | 174 | 0.871554614 | cat4 | from_7531 | to_7531 | art_long | art_short | J |
| 112 | 174 | 0.718328633 | cat4 | from_7532 | to_7532 | art_long | art_short | J |
| 119 | 174 | 0.607166241 | cat4 | from_7533 | to_7533 | art_long | art_short | J |
| 122 | 174 | 0.851879836 | cat4 | from_7534 | to_7534 | art_long | art_short | J |
| 173 | 174 | 0.873411974 | cat4 | from_7535 | to_7535 | art_short | art_short | H |
| 82 | 175 | 0.815259917 | cat4 | from_7536 | to_7536 | art_long | art_short | J |
| 121 | 175 | 0.610953839 | cat4 | from_7537 | to_7537 | art_long | art_short | J |
| 150 | 175 | 0.74702169 | cat4 | from_7538 | to_7538 | art_long | art_short | J |
| 14 | 176 | -0.57703672 | cat4 | from_7539 | to_7539 | art_long | art_short | J |
| 26 | 176 | -0.618826561 | cat4 | from_7540 | to_7540 | art_long | art_short | J |
| 37 | 176 | -0.59750836 | cat4 | from_7541 | to_7541 | art_long | art_short | J |
| 46 | 176 | 0.59346272 | cat4 | from_7542 | to_7542 | art_long | art_short | J |
| 61 | 176 | 0.82325867 | cat4 | from_7543 | to_7543 | art_long | art_short | J |
| 85 | 176 | -0.592069516 | cat4 | from_7544 | to_7544 | art_long | art_short | J |
| 97 | 176 | -0.579795314 | cat4 | from_7545 | to_7545 | art_long | art_short | J |
| 102 | 176 | 0.610965667 | cat4 | from_7546 | to_7546 | art_long | art_short | J |
| 106 | 176 | 0.61836516 | cat4 | from_7547 | to_7547 | art_long | art_short | J |
| 115 | 176 | -0.656468667 | cat4 | from_7548 | to_7548 | art_long | art_short | J |
| 117 | 176 | -0.622046802 | cat4 | from_7549 | to_7549 | art_long | art_short | J |
| 123 | 176 | -0.588573128 | cat4 | from_7550 | to_7550 | art_long | art_short | J |
| 127 | 176 | -0.60824819 | cat4 | from_7551 | to_7551 | art_long | art_short | J |
| 135 | 176 | 0.781775148 | cat4 | from_7552 | to_7552 | art_long | art_short | J |
| 139 | 176 | 0.587397743 | cat4 | from_7553 | to_7553 | art_long | art_short | J |
| 170 | 176 | 0.610877901 | cat4 | from_7554 | to_7554 | art_short | art_short | H |
| 171 | 176 | 0.580818311 | cat4 | from_7555 | to_7555 | art_short | art_short | H |
| 17 | 177 | 0.680583335 | cat4 | from_7556 | to_7556 | art_long | art_short | J |
| 27 | 177 | -0.630290427 | cat4 | from_7557 | to_7557 | art_long | art_short | J |
| 35 | 177 | 0.687622969 | cat4 | from_7558 | to_7558 | art_long | art_short | J |
| 38 | 177 | 0.681133097 | cat4 | from_7559 | to_7559 | art_long | art_short | J |
| 42 | 177 | 0.617782421 | cat4 | from_7560 | to_7560 | art_long | art_short | J |
| 54 | 177 | -0.672832158 | cat4 | from_7561 | to_7561 | art_long | art_short | J |
| 77 | 177 | -0.685864271 | cat4 | from_7562 | to_7562 | art_long | art_short | J |
| 80 | 177 | -0.691963778 | cat4 | from_7563 | to_7563 | art_long | art_short | J |
| 92 | 177 | 0.672039709 | cat4 | from_7564 | to_7564 | art_long | art_short | J |
| 93 | 177 | -0.608770297 | cat4 | from_7565 | to_7565 | art_long | art_short | J |
| 120 | 177 | -0.647764039 | cat4 | from_7566 | to_7566 | art_long | art_short | J |
| 126 | 177 | -0.622407279 | cat4 | from_7567 | to_7567 | art_long | art_short | J |
| 132 | 177 | 0.61021826 | cat4 | from_7568 | to_7568 | art_long | art_short | J |
| 133 | 177 | 0.906056959 | cat4 | from_7569 | to_7569 | art_long | art_short | J |
| 134 | 177 | 0.645096549 | cat4 | from_7570 | to_7570 | art_long | art_short | J |
| 138 | 177 | 0.640267936 | cat4 | from_7571 | to_7571 | art_long | art_short | J |
| 147 | 177 | 0.635631562 | cat4 | from_7572 | to_7572 | art_long | art_short | J |
| 42 | 178 | 0.81763715 | cat4 | from_7573 | to_7573 | art_long | art_short | J |
| 45 | 178 | 0.71890532 | cat4 | from_7574 | to_7574 | art_long | art_short | J |
| 52 | 178 | 0.70458886 | cat4 | from_7575 | to_7575 | art_long | art_short | J |
| 54 | 178 | -0.599213839 | cat4 | from_7576 | to_7576 | art_long | art_short | J |
| 62 | 178 | 0.764426808 | cat4 | from_7577 | to_7577 | art_long | art_short | J |
| 80 | 178 | -0.604193425 | cat4 | from_7578 | to_7578 | art_long | art_short | J |
| 92 | 178 | 0.809884963 | cat4 | from_7579 | to_7579 | art_long | art_short | J |
| 100 | 178 | 0.630445063 | cat4 | from_7580 | to_7580 | art_long | art_short | J |
| 101 | 178 | 0.81245836 | cat4 | from_7581 | to_7581 | art_long | art_short | J |
| 107 | 178 | 0.620850859 | cat4 | from_7582 | to_7582 | art_long | art_short | J |
| 133 | 178 | 0.63423578 | cat4 | from_7583 | to_7583 | art_long | art_short | J |
| 147 | 178 | 0.860737743 | cat4 | from_7584 | to_7584 | art_long | art_short | J |
| 149 | 178 | 0.611518223 | cat4 | from_7585 | to_7585 | art_long | art_short | J |
| 160 | 178 | 0.596843414 | cat4 | from_7586 | to_7586 | art_short | art_short | H |
| 177 | 178 | 0.69494187 | cat4 | from_7587 | to_7587 | art_short | art_short | H |
| 2 | 179 | 0.92889382 | cat4 | from_7588 | to_7588 | art_long | art_short | J |
| 6 | 179 | -0.610099305 | cat4 | from_7589 | to_7589 | art_long | art_short | J |
| 16 | 179 | 0.67267555 | cat4 | from_7590 | to_7590 | art_long | art_short | J |

| | | | | | | | | |
|---|---|---|---|---|---|---|---|---|
| 53 | 179 | 0.860579123 | cat4 | from_7591 | to_7591 | art_long | art_short | J |
| 155 | 179 | 0.955541013 | cat4 | from_7592 | to_7592 | art_short | art_short | H |
| 162 | 179 | 0.577966815 | cat4 | from_7593 | to_7593 | art_short | art_short | H |
| 164 | 179 | 0.715230615 | cat4 | from_7594 | to_7594 | art_short | art_short | H |
| 168 | 179 | 0.598450466 | cat4 | from_7595 | to_7595 | art_short | art_short | H |
| 15 | 180 | 0.616591197 | cat4 | from_7596 | to_7596 | art_long | art_short | J |
| 16 | 180 | 0.577697735 | cat4 | from_7597 | to_7597 | art_long | art_short | J |
| 53 | 180 | 0.820503538 | cat4 | from_7598 | to_7598 | art_long | art_short | J |
| 164 | 180 | 0.577084119 | cat4 | from_7599 | to_7599 | art_short | art_short | H |
| 168 | 180 | 0.951561478 | cat4 | from_7600 | to_7600 | art_short | art_short | H |
| 169 | 180 | 0.669925264 | cat4 | from_7601 | to_7601 | art_short | art_short | H |
| 179 | 180 | 0.667971507 | cat4 | from_7602 | to_7602 | art_short | art_short | H |
| 3 | 181 | 0.672197624 | cat4 | from_7603 | to_7603 | art_long | art_short | J |
| 29 | 181 | 0.889969257 | cat4 | from_7604 | to_7604 | art_long | art_short | J |
| 41 | 181 | 0.631684776 | cat4 | from_7605 | to_7605 | art_long | art_short | J |
| 66 | 181 | 0.631391876 | cat4 | from_7606 | to_7606 | art_long | art_short | J |
| 67 | 181 | 0.693108431 | cat4 | from_7607 | to_7607 | art_long | art_short | J |
| 71 | 181 | 0.910244772 | cat4 | from_7608 | to_7608 | art_long | art_short | J |
| 82 | 181 | 0.81574576 | cat4 | from_7609 | to_7609 | art_long | art_short | J |
| 98 | 181 | 0.611660702 | cat4 | from_7610 | to_7610 | art_long | art_short | J |
| 124 | 181 | 0.647067523 | cat4 | from_7611 | to_7611 | art_long | art_short | J |
| 150 | 181 | 0.81629823 | cat4 | from_7612 | to_7612 | art_long | art_short | J |
| 175 | 181 | 0.738707252 | cat4 | from_7613 | to_7613 | art_short | art_short | H |
| 22 | 182 | 0.576890364 | cat4 | from_7614 | to_7614 | art_long | art_short | J |
| 23 | 182 | 0.631726725 | cat4 | from_7615 | to_7615 | art_long | art_short | J |
| 32 | 182 | -0.642815536 | cat4 | from_7616 | to_7616 | art_long | art_short | J |
| 49 | 182 | 0.62721158 | cat4 | from_7617 | to_7617 | art_long | art_short | J |
| 56 | 182 | 0.905438874 | cat4 | from_7618 | to_7618 | art_long | art_short | J |
| 57 | 182 | 0.769553495 | cat4 | from_7619 | to_7619 | art_long | art_short | J |
| 70 | 182 | 0.718000398 | cat4 | from_7620 | to_7620 | art_long | art_short | J |
| 75 | 182 | 0.7025167 | cat4 | from_7621 | to_7621 | art_long | art_short | J |
| 85 | 182 | 0.6868906 | cat4 | from_7622 | to_7622 | art_long | art_short | J |
| 123 | 182 | 0.767058049 | cat4 | from_7623 | to_7623 | art_long | art_short | J |
| 146 | 182 | 0.752235287 | cat4 | from_7624 | to_7624 | art_long | art_short | J |
| 161 | 182 | 0.583558131 | cat4 | from_7625 | to_7625 | art_short | art_short | H |
| 44 | 183 | 0.691294232 | cat4 | from_7626 | to_7626 | art_long | art_short | J |
| 46 | 183 | 0.65873458 | cat4 | from_7627 | to_7627 | art_long | art_short | J |
| 50 | 183 | 0.647423388 | cat4 | from_7628 | to_7628 | art_long | art_short | J |
| 52 | 183 | 0.685739438 | cat4 | from_7629 | to_7629 | art_long | art_short | J |
| 60 | 183 | 0.813844345 | cat4 | from_7630 | to_7630 | art_long | art_short | J |
| 61 | 183 | 0.859419429 | cat4 | from_7631 | to_7631 | art_long | art_short | J |
| 64 | 183 | -0.636149409 | cat4 | from_7632 | to_7632 | art_long | art_short | J |
| 99 | 183 | 0.857644545 | cat4 | from_7633 | to_7633 | art_long | art_short | J |
| 105 | 183 | 0.86699864 | cat4 | from_7634 | to_7634 | art_long | art_short | J |
| 106 | 183 | 0.657617229 | cat4 | from_7635 | to_7635 | art_long | art_short | J |
| 128 | 183 | 0.773142679 | cat4 | from_7636 | to_7636 | art_long | art_short | J |
| 148 | 183 | 0.881314543 | cat4 | from_7637 | to_7637 | art_long | art_short | J |
| 176 | 183 | 0.694247696 | cat4 | from_7638 | to_7638 | art_short | art_short | H |
| 17 | 184 | 0.579158637 | cat4 | from_7639 | to_7639 | art_long | art_short | J |
| 36 | 184 | 0.577639207 | cat4 | from_7640 | to_7640 | art_long | art_short | J |
| 42 | 184 | 0.765412668 | cat4 | from_7641 | to_7641 | art_long | art_short | J |
| 43 | 184 | 0.902239182 | cat4 | from_7642 | to_7642 | art_long | art_short | J |
| 45 | 184 | 0.821505385 | cat4 | from_7643 | to_7643 | art_long | art_short | J |
| 52 | 184 | 0.587319731 | cat4 | from_7644 | to_7644 | art_long | art_short | J |
| 60 | 184 | 0.624945939 | cat4 | from_7645 | to_7645 | art_long | art_short | J |
| 62 | 184 | 0.866942506 | cat4 | from_7646 | to_7646 | art_long | art_short | J |
| 92 | 184 | 0.740741013 | cat4 | from_7647 | to_7647 | art_long | art_short | J |
| 100 | 184 | 0.685411958 | cat4 | from_7648 | to_7648 | art_long | art_short | J |
| 101 | 184 | 0.861522401 | cat4 | from_7649 | to_7649 | art_long | art_short | J |
| 107 | 184 | 0.927520153 | cat4 | from_7650 | to_7650 | art_long | art_short | J |
| 128 | 184 | 0.611978821 | cat4 | from_7651 | to_7651 | art_long | art_short | J |
| 138 | 184 | 0.57938277 | cat4 | from_7652 | to_7652 | art_long | art_short | J |
| 139 | 184 | 0.840633872 | cat4 | from_7653 | to_7653 | art_long | art_short | J |
| 147 | 184 | 0.785110119 | cat4 | from_7654 | to_7654 | art_long | art_short | J |
| 149 | 184 | 0.921887602 | cat4 | from_7655 | to_7655 | art_long | art_short | J |
| 160 | 184 | 0.618031048 | cat4 | from_7656 | to_7656 | art_short | art_short | H |
| 170 | 184 | 0.73431084 | cat4 | from_7657 | to_7657 | art_short | art_short | H |
| 171 | 184 | 0.775439567 | cat4 | from_7658 | to_7658 | art_short | art_short | H |
| 178 | 184 | 0.813599595 | cat4 | from_7659 | to_7659 | art_short | art_short | H |
| 3 | 185 | 0.760683037 | cat4 | from_7660 | to_7660 | art_long | art_short | J |
| 82 | 185 | 0.778633127 | cat4 | from_7661 | to_7661 | art_long | art_short | J |
| 150 | 185 | 0.69368989 | cat4 | from_7662 | to_7662 | art_long | art_short | J |
| 175 | 185 | 0.773365319 | cat4 | from_7663 | to_7663 | art_short | art_short | H |
| 181 | 185 | 0.588893336 | cat4 | from_7664 | to_7664 | art_short | art_short | H |
| 5 | 186 | 0.608705358 | cat4 | from_7665 | to_7665 | art_long | art_short | J |
| 9 | 186 | 0.583412701 | cat4 | from_7666 | to_7666 | art_long | art_short | J |
| 12 | 186 | 0.672022147 | cat4 | from_7667 | to_7667 | art_long | art_short | J |
| 13 | 186 | 0.736386884 | cat4 | from_7668 | to_7668 | art_long | art_short | J |
| 19 | 186 | 0.656929175 | cat4 | from_7669 | to_7669 | art_long | art_short | J |
| 35 | 186 | -0.627691865 | cat4 | from_7670 | to_7670 | art_long | art_short | J |
| 38 | 186 | -0.587625751 | cat4 | from_7671 | to_7671 | art_long | art_short | J |
| 63 | 186 | 0.79072231 | cat4 | from_7672 | to_7672 | art_long | art_short | J |
| 79 | 186 | 0.675926077 | cat4 | from_7673 | to_7673 | art_long | art_short | J |
| 114 | 186 | 0.710939413 | cat4 | from_7674 | to_7674 | art_long | art_short | J |
| 116 | 186 | 0.726671537 | cat4 | from_7675 | to_7675 | art_long | art_short | J |
| 132 | 186 | -0.590258853 | cat4 | from_7676 | to_7676 | art_long | art_short | J |
| 156 | 186 | 0.870485467 | cat4 | from_7677 | to_7677 | art_short | art_short | H |
| 158 | 186 | 0.836649551 | cat4 | from_7678 | to_7678 | art_short | art_short | H |
| 163 | 186 | 0.631254638 | cat4 | from_7679 | to_7679 | art_short | art_short | H |
| 165 | 186 | 0.700108326 | cat4 | from_7680 | to_7680 | art_short | art_short | H |
| 167 | 186 | 0.677572993 | cat4 | from_7681 | to_7681 | art_short | art_short | H |
| 2 | 187 | 0.664434534 | cat4 | from_7682 | to_7682 | art_long | art_short | J |
| 16 | 187 | 0.604160653 | cat4 | from_7683 | to_7683 | art_long | art_short | J |
| 53 | 187 | 0.836050351 | cat4 | from_7684 | to_7684 | art_long | art_short | J |
| 155 | 187 | 0.659553694 | cat4 | from_7685 | to_7685 | art_short | art_short | H |
| 162 | 187 | 0.577955391 | cat4 | from_7686 | to_7686 | art_short | art_short | H |
| 164 | 187 | 0.60347143 | cat4 | from_7687 | to_7687 | art_short | art_short | H |
| 168 | 187 | 0.902018524 | cat4 | from_7688 | to_7688 | art_short | art_short | H |
| 169 | 187 | 0.656020699 | cat4 | from_7689 | to_7689 | art_short | art_short | H |
| 179 | 187 | 0.768038163 | cat4 | from_7690 | to_7690 | art_short | art_short | H |
| 180 | 187 | 0.966320152 | cat4 | from_7691 | to_7691 | art_short | art_short | H |
| 4 | 188 | 0.832628095 | cat4 | from_7692 | to_7692 | art_long | art_short | J |
| 6 | 188 | 0.796636899 | cat4 | from_7693 | to_7693 | art_long | art_short | J |
| 7 | 188 | 0.822740007 | cat4 | from_7694 | to_7694 | art_long | art_short | J |

| | | | | | | | | |
|---|---|---|---|---|---|---|---|---|
| 11 | 188 | 0.701379775 | cat4 | from_7695 | to_7695 | art_long | art_short | J |
| 65 | 188 | 0.631834079 | cat4 | from_7696 | to_7696 | art_long | art_short | J |
| 118 | 188 | -0.640696025 | cat4 | from_7697 | to_7697 | art_long | art_short | J |
| 131 | 188 | -0.601436691 | cat4 | from_7698 | to_7698 | art_long | art_short | J |
| 137 | 188 | -0.597336606 | cat4 | from_7699 | to_7699 | art_long | art_short | J |
| 141 | 188 | -0.608221873 | cat4 | from_7700 | to_7700 | art_long | art_short | J |
| 144 | 188 | -0.655116275 | cat4 | from_7701 | to_7701 | art_long | art_short | J |
| 159 | 188 | 0.995812128 | cat4 | from_7702 | to_7702 | art_short | art_short | H |
| 160 | 188 | 0.878762104 | cat4 | from_7703 | to_7703 | art_short | art_short | H |
| 29 | 189 | 0.691219065 | cat4 | from_7704 | to_7704 | art_long | art_short | J |
| 66 | 189 | 0.912552042 | cat4 | from_7705 | to_7705 | art_long | art_short | J |
| 67 | 189 | 0.872834634 | cat4 | from_7706 | to_7706 | art_long | art_short | J |
| 71 | 189 | 0.706371054 | cat4 | from_7707 | to_7707 | art_long | art_short | J |
| 82 | 189 | 0.701263381 | cat4 | from_7708 | to_7708 | art_long | art_short | J |
| 150 | 189 | 0.633712457 | cat4 | from_7709 | to_7709 | art_long | art_short | J |
| 175 | 189 | 0.683608939 | cat4 | from_7710 | to_7710 | art_short | art_short | H |
| 181 | 189 | 0.756398849 | cat4 | from_7711 | to_7711 | art_short | art_short | H |
| 28 | 190 | 0.597613743 | cat4 | from_7712 | to_7712 | art_long | art_short | J |
| 32 | 190 | 0.766994378 | cat4 | from_7713 | to_7713 | art_long | art_short | J |
| 42 | 190 | 0.612792025 | cat4 | from_7714 | to_7714 | art_long | art_short | J |
| 45 | 190 | 0.625501734 | cat4 | from_7715 | to_7715 | art_long | art_short | J |
| 59 | 190 | 0.590144498 | cat4 | from_7716 | to_7716 | art_long | art_short | J |
| 62 | 190 | 0.610600374 | cat4 | from_7717 | to_7717 | art_long | art_short | J |
| 86 | 190 | 0.695485886 | cat4 | from_7718 | to_7718 | art_long | art_short | J |
| 92 | 190 | 0.636305533 | cat4 | from_7719 | to_7719 | art_long | art_short | J |
| 94 | 190 | 0.677240172 | cat4 | from_7720 | to_7720 | art_long | art_short | J |
| 97 | 190 | 0.652853074 | cat4 | from_7721 | to_7721 | art_long | art_short | J |
| 101 | 190 | 0.618749134 | cat4 | from_7722 | to_7722 | art_long | art_short | J |
| 147 | 190 | 0.599849357 | cat4 | from_7723 | to_7723 | art_long | art_short | J |
| 173 | 190 | 0.589292402 | cat4 | from_7724 | to_7724 | art_short | art_short | H |
| 174 | 190 | 0.584770336 | cat4 | from_7725 | to_7725 | art_short | art_short | H |
| 178 | 190 | 0.599593038 | cat4 | from_7726 | to_7726 | art_short | art_short | H |
| 184 | 190 | 0.576160139 | cat4 | from_7727 | to_7727 | art_short | art_short | H |
| 8 | 191 | -0.580146691 | cat4 | from_7728 | to_7728 | art_long | art_short | J |
| 9 | 191 | -0.593294045 | cat4 | from_7729 | to_7729 | art_long | art_short | J |
| 10 | 191 | -0.58538088 | cat4 | from_7730 | to_7730 | art_long | art_short | J |
| 28 | 191 | 0.637961108 | cat4 | from_7731 | to_7731 | art_long | art_short | J |
| 32 | 191 | 0.824063183 | cat4 | from_7732 | to_7732 | art_long | art_short | J |
| 42 | 191 | 0.576285449 | cat4 | from_7733 | to_7733 | art_long | art_short | J |
| 45 | 191 | 0.64138819 | cat4 | from_7734 | to_7734 | art_long | art_short | J |
| 62 | 191 | 0.601187354 | cat4 | from_7735 | to_7735 | art_long | art_short | J |
| 86 | 191 | 0.665383718 | cat4 | from_7736 | to_7736 | art_long | art_short | J |
| 92 | 191 | 0.580400876 | cat4 | from_7737 | to_7737 | art_long | art_short | J |
| 94 | 191 | 0.66547496 | cat4 | from_7738 | to_7738 | art_long | art_short | J |
| 97 | 191 | 0.60871968 | cat4 | from_7739 | to_7739 | art_long | art_short | J |
| 100 | 191 | 0.65234912 | cat4 | from_7740 | to_7740 | art_long | art_short | J |
| 122 | 191 | 0.593290058 | cat4 | from_7741 | to_7741 | art_long | art_short | J |
| 170 | 191 | 0.613849812 | cat4 | from_7742 | to_7742 | art_short | art_short | H |
| 171 | 191 | 0.631219506 | cat4 | from_7743 | to_7743 | art_short | art_short | H |
| 173 | 191 | 0.674957542 | cat4 | from_7744 | to_7744 | art_short | art_short | H |
| 182 | 191 | -0.576964728 | cat4 | from_7745 | to_7745 | art_short | art_short | H |
| 190 | 191 | 0.952380272 | cat4 | from_7746 | to_7746 | art_short | art_short | H |
| 12 | 192 | 0.661164293 | cat4 | from_7747 | to_7747 | art_long | art_short | J |
| 13 | 192 | 0.675763782 | cat4 | from_7748 | to_7748 | art_long | art_short | J |
| 15 | 192 | 0.794315343 | cat4 | from_7749 | to_7749 | art_long | art_short | J |
| 16 | 192 | 0.748798481 | cat4 | from_7750 | to_7750 | art_long | art_short | J |
| 18 | 192 | 0.681490206 | cat4 | from_7751 | to_7751 | art_long | art_short | J |
| 19 | 192 | 0.608384703 | cat4 | from_7752 | to_7752 | art_long | art_short | J |
| 39 | 192 | 0.648322593 | cat4 | from_7753 | to_7753 | art_long | art_short | J |
| 40 | 192 | 0.590559588 | cat4 | from_7754 | to_7754 | art_long | art_short | J |
| 162 | 192 | 0.728740527 | cat4 | from_7755 | to_7755 | art_short | art_short | H |
| 163 | 192 | 0.741922312 | cat4 | from_7756 | to_7756 | art_short | art_short | H |
| 164 | 192 | 0.729386846 | cat4 | from_7757 | to_7757 | art_short | art_short | H |
| 167 | 192 | 0.676469345 | cat4 | from_7758 | to_7758 | art_short | art_short | H |
| 169 | 192 | 0.657072788 | cat4 | from_7759 | to_7759 | art_short | art_short | H |
| 75 | 193 | 0.664828694 | cat4 | from_7760 | to_7760 | art_long | art_short | J |
| 84 | 193 | 0.723536183 | cat4 | from_7761 | to_7761 | art_long | art_short | J |
| 14 | 194 | 0.750634589 | cat4 | from_7762 | to_7762 | art_long | art_short | J |
| 24 | 194 | 0.877897361 | cat4 | from_7763 | to_7763 | art_long | art_short | J |
| 25 | 194 | 0.711186548 | cat4 | from_7764 | to_7764 | art_long | art_short | J |
| 28 | 194 | 0.947575914 | cat4 | from_7765 | to_7765 | art_long | art_short | J |
| 30 | 194 | 0.588250464 | cat4 | from_7766 | to_7766 | art_long | art_short | J |
| 31 | 194 | 0.583488223 | cat4 | from_7767 | to_7767 | art_long | art_short | J |
| 33 | 194 | 0.62751813 | cat4 | from_7768 | to_7768 | art_long | art_short | J |
| 50 | 194 | -0.608817897 | cat4 | from_7769 | to_7769 | art_long | art_short | J |
| 55 | 194 | 0.612126001 | cat4 | from_7770 | to_7770 | art_long | art_short | J |
| 59 | 194 | 0.861309263 | cat4 | from_7771 | to_7771 | art_long | art_short | J |
| 64 | 194 | 0.583103634 | cat4 | from_7772 | to_7772 | art_long | art_short | J |
| 73 | 194 | 0.763653379 | cat4 | from_7773 | to_7773 | art_long | art_short | J |
| 74 | 194 | 0.746501667 | cat4 | from_7774 | to_7774 | art_long | art_short | J |
| 76 | 194 | 0.892934344 | cat4 | from_7775 | to_7775 | art_long | art_short | J |
| 78 | 194 | 0.771948395 | cat4 | from_7776 | to_7776 | art_long | art_short | J |
| 87 | 194 | 0.596560646 | cat4 | from_7777 | to_7777 | art_long | art_short | J |
| 90 | 194 | 0.775983762 | cat4 | from_7778 | to_7778 | art_long | art_short | J |
| 95 | 194 | 0.592138387 | cat4 | from_7779 | to_7779 | art_long | art_short | J |
| 97 | 194 | 0.820435052 | cat4 | from_7780 | to_7780 | art_long | art_short | J |
| 109 | 194 | 0.621968302 | cat4 | from_7781 | to_7781 | art_long | art_short | J |
| 110 | 194 | 0.736771661 | cat4 | from_7782 | to_7782 | art_long | art_short | J |
| 112 | 194 | 0.617585908 | cat4 | from_7783 | to_7783 | art_long | art_short | J |
| 119 | 194 | 0.584955652 | cat4 | from_7784 | to_7784 | art_long | art_short | J |
| 122 | 194 | 0.90870897 | cat4 | from_7785 | to_7785 | art_long | art_short | J |
| 173 | 194 | 0.963004421 | cat4 | from_7786 | to_7786 | art_short | art_short | H |
| 174 | 194 | 0.962150896 | cat4 | from_7787 | to_7787 | art_short | art_short | H |
| 190 | 194 | 0.60941594 | cat4 | from_7788 | to_7788 | art_short | art_short | H |
| 191 | 194 | 0.644410379 | cat4 | from_7789 | to_7789 | art_short | art_short | H |
| 24 | 195 | 0.629179987 | cat4 | from_7790 | to_7790 | art_long | art_short | J |
| 27 | 195 | 0.605716387 | cat4 | from_7791 | to_7791 | art_long | art_short | J |
| 30 | 195 | 0.631840805 | cat4 | from_7792 | to_7792 | art_long | art_short | J |
| 47 | 195 | 0.740955496 | cat4 | from_7793 | to_7793 | art_long | art_short | J |
| 51 | 195 | -0.639591116 | cat4 | from_7794 | to_7794 | art_long | art_short | J |
| 55 | 195 | 0.852937509 | cat4 | from_7795 | to_7795 | art_long | art_short | J |
| 70 | 195 | -0.640236959 | cat4 | from_7796 | to_7796 | art_long | art_short | J |
| 76 | 195 | 0.580849481 | cat4 | from_7797 | to_7797 | art_long | art_short | J |
| 77 | 195 | 0.636146272 | cat4 | from_7798 | to_7798 | art_long | art_short | J |

| | | | | | | | | |
|---|---|---|---|---|---|---|---|---|
| 80 | 195 | 0.74812844 | cat4 | from_7799 | to_7799 | art_long | art_short | J |
| 87 | 195 | 0.731112333 | cat4 | from_7800 | to_7800 | art_long | art_short | J |
| 95 | 195 | 0.924695137 | cat4 | from_7801 | to_7801 | art_long | art_short | J |
| 109 | 195 | 0.846913101 | cat4 | from_7802 | to_7802 | art_long | art_short | J |
| 119 | 195 | 0.719381197 | cat4 | from_7803 | to_7803 | art_long | art_short | J |
| 122 | 195 | 0.600589429 | cat4 | from_7804 | to_7804 | art_long | art_short | J |
| 131 | 195 | 0.6519415 | cat4 | from_7805 | to_7805 | art_long | art_short | J |
| 173 | 195 | 0.64976836 | cat4 | from_7806 | to_7806 | art_short | art_short | H |
| 194 | 195 | 0.576419519 | cat4 | from_7807 | to_7807 | art_short | art_short | H |
| 2 | 196 | 0.66490171 | cat4 | from_7808 | to_7808 | art_long | art_short | J |
| 155 | 196 | 0.717043782 | cat4 | from_7809 | to_7809 | art_short | art_short | H |
| 162 | 196 | 0.955939725 | cat4 | from_7810 | to_7810 | art_short | art_short | H |
| 163 | 196 | 0.817095026 | cat4 | from_7811 | to_7811 | art_short | art_short | H |
| 164 | 196 | 0.583943743 | cat4 | from_7812 | to_7812 | art_short | art_short | H |
| 179 | 196 | 0.607366428 | cat4 | from_7813 | to_7813 | art_short | art_short | H |
| 192 | 196 | 0.617164733 | cat4 | from_7814 | to_7814 | art_short | art_short | H |
| 2 | 197 | 0.597263184 | cat4 | from_7815 | to_7815 | art_long | art_short | J |
| 12 | 197 | 0.758776486 | cat4 | from_7816 | to_7816 | art_long | art_short | J |
| 16 | 197 | 0.889944061 | cat4 | from_7817 | to_7817 | art_long | art_short | J |
| 18 | 197 | 0.655336289 | cat4 | from_7818 | to_7818 | art_long | art_short | J |
| 19 | 197 | 0.686832143 | cat4 | from_7819 | to_7819 | art_long | art_short | J |
| 39 | 197 | 0.828193533 | cat4 | from_7820 | to_7820 | art_long | art_short | J |
| 40 | 197 | 0.832375437 | cat4 | from_7821 | to_7821 | art_long | art_short | J |
| 53 | 197 | 0.635831231 | cat4 | from_7822 | to_7822 | art_long | art_short | J |
| 79 | 197 | 0.607010914 | cat4 | from_7823 | to_7823 | art_long | art_short | J |
| 155 | 197 | 0.643108711 | cat4 | from_7824 | to_7824 | art_short | art_short | H |
| 162 | 197 | 0.669384975 | cat4 | from_7825 | to_7825 | art_short | art_short | H |
| 163 | 197 | 0.718770682 | cat4 | from_7826 | to_7826 | art_short | art_short | H |
| 164 | 197 | 0.965381297 | cat4 | from_7827 | to_7827 | art_short | art_short | H |
| 167 | 197 | 0.774427213 | cat4 | from_7828 | to_7828 | art_short | art_short | H |
| 169 | 197 | 0.703924992 | cat4 | from_7829 | to_7829 | art_short | art_short | H |
| 179 | 197 | 0.726445178 | cat4 | from_7830 | to_7830 | art_short | art_short | H |
| 192 | 197 | 0.739402055 | cat4 | from_7831 | to_7831 | art_short | art_short | H |
| 196 | 197 | 0.635268164 | cat4 | from_7832 | to_7832 | art_short | art_short | H |
| 6 | 198 | 0.618650498 | cat4 | from_7833 | to_7833 | art_long | art_short | J |
| 8 | 198 | 0.730898079 | cat4 | from_7834 | to_7834 | art_long | art_short | J |
| 9 | 198 | 0.70744797 | cat4 | from_7835 | to_7835 | art_long | art_short | J |
| 10 | 198 | 0.706439487 | cat4 | from_7836 | to_7836 | art_long | art_short | J |
| 35 | 198 | -0.612052429 | cat4 | from_7837 | to_7837 | art_long | art_short | J |
| 38 | 198 | -0.580727687 | cat4 | from_7838 | to_7838 | art_long | art_short | J |
| 65 | 198 | 0.586692042 | cat4 | from_7839 | to_7839 | art_long | art_short | J |
| 84 | 198 | 0.649725763 | cat4 | from_7840 | to_7840 | art_long | art_short | J |
| 125 | 198 | 0.790894325 | cat4 | from_7841 | to_7841 | art_long | art_short | J |
| 126 | 198 | 0.587458269 | cat4 | from_7842 | to_7842 | art_long | art_short | J |
| 132 | 198 | -0.591814425 | cat4 | from_7843 | to_7843 | art_long | art_short | J |
| 156 | 198 | 0.753078623 | cat4 | from_7844 | to_7844 | art_short | art_short | H |
| 157 | 198 | 0.776459874 | cat4 | from_7845 | to_7845 | art_short | art_short | H |
| 158 | 198 | 0.723405168 | cat4 | from_7846 | to_7846 | art_short | art_short | H |
| 161 | 198 | 0.621283508 | cat4 | from_7847 | to_7847 | art_short | art_short | H |
| 186 | 198 | 0.695347705 | cat4 | from_7848 | to_7848 | art_short | art_short | H |
| 193 | 198 | 0.640645305 | cat4 | from_7849 | to_7849 | art_short | art_short | H |
| 49 | 199 | 0.755303964 | cat4 | from_7850 | to_7850 | art_long | art_short | J |
| 56 | 199 | 0.732742261 | cat4 | from_7851 | to_7851 | art_long | art_short | J |
| 65 | 199 | 0.65233264 | cat4 | from_7852 | to_7852 | art_long | art_short | J |
| 70 | 199 | 0.650689824 | cat4 | from_7853 | to_7853 | art_long | art_short | J |
| 75 | 199 | 0.835317514 | cat4 | from_7854 | to_7854 | art_long | art_short | J |
| 84 | 199 | 0.660249908 | cat4 | from_7855 | to_7855 | art_long | art_short | J |
| 85 | 199 | 0.812320076 | cat4 | from_7856 | to_7856 | art_long | art_short | J |
| 94 | 199 | -0.599926514 | cat4 | from_7857 | to_7857 | art_long | art_short | J |
| 113 | 199 | 0.657612136 | cat4 | from_7858 | to_7858 | art_long | art_short | J |
| 123 | 199 | 0.778148148 | cat4 | from_7859 | to_7859 | art_long | art_short | J |
| 146 | 199 | 0.862370258 | cat4 | from_7860 | to_7860 | art_long | art_short | J |
| 182 | 199 | 0.711707061 | cat4 | from_7861 | to_7861 | art_short | art_short | H |
| 193 | 199 | 0.808731036 | cat4 | from_7862 | to_7862 | art_short | art_short | H |
| 24 | 200 | 0.620623532 | cat4 | from_7863 | to_7863 | art_long | art_short | J |
| 51 | 200 | -0.658722762 | cat4 | from_7864 | to_7864 | art_long | art_short | J |
| 55 | 200 | 0.662914301 | cat4 | from_7865 | to_7865 | art_long | art_short | J |
| 70 | 200 | -0.638844478 | cat4 | from_7866 | to_7866 | art_long | art_short | J |
| 80 | 200 | 0.673762601 | cat4 | from_7867 | to_7867 | art_long | art_short | J |
| 95 | 200 | 0.837633357 | cat4 | from_7868 | to_7868 | art_long | art_short | J |
| 109 | 200 | 0.654475992 | cat4 | from_7869 | to_7869 | art_long | art_short | J |
| 131 | 200 | 0.621880371 | cat4 | from_7870 | to_7870 | art_long | art_short | J |
| 195 | 200 | 0.932968402 | cat4 | from_7871 | to_7871 | art_short | art_short | H |
| 8 | 201 | 0.646302129 | cat4 | from_7872 | to_7872 | art_long | art_short | J |
| 9 | 201 | 0.739196754 | cat4 | from_7873 | to_7873 | art_long | art_short | J |
| 10 | 201 | 0.782557904 | cat4 | from_7874 | to_7874 | art_long | art_short | J |
| 17 | 201 | -0.585217357 | cat4 | from_7875 | to_7875 | art_long | art_short | J |
| 23 | 201 | 0.760742088 | cat4 | from_7876 | to_7876 | art_long | art_short | J |
| 36 | 201 | -0.625597075 | cat4 | from_7877 | to_7877 | art_long | art_short | J |
| 88 | 201 | 0.648769334 | cat4 | from_7878 | to_7878 | art_long | art_short | J |
| 125 | 201 | 0.736317169 | cat4 | from_7879 | to_7879 | art_long | art_short | J |
| 138 | 201 | -0.576430225 | cat4 | from_7880 | to_7880 | art_long | art_short | J |
| 156 | 201 | 0.636698581 | cat4 | from_7881 | to_7881 | art_short | art_short | H |
| 158 | 201 | 0.650070715 | cat4 | from_7882 | to_7882 | art_short | art_short | H |
| 193 | 201 | 0.598088098 | cat4 | from_7883 | to_7883 | art_short | art_short | H |
| 198 | 201 | 0.65929941 | cat4 | from_7884 | to_7884 | art_short | art_short | H |
| 51 | 202 | 0.635360246 | cat4 | from_7885 | to_7885 | art_long | art_short | J |
| 144 | 202 | -0.581408195 | cat4 | from_7886 | to_7886 | art_long | art_short | J |
| 5 | 203 | -0.58275148 | cat4 | from_7887 | to_7887 | art_long | art_short | J |
| 14 | 203 | -0.599313644 | cat4 | from_7888 | to_7888 | art_long | art_short | J |
| 26 | 203 | -0.5865748 | cat4 | from_7889 | to_7889 | art_long | art_short | J |
| 61 | 203 | 0.7279122 | cat4 | from_7890 | to_7890 | art_long | art_short | J |
| 70 | 203 | -0.618198343 | cat4 | from_7891 | to_7891 | art_long | art_short | J |
| 73 | 203 | -0.582089395 | cat4 | from_7892 | to_7892 | art_long | art_short | J |
| 102 | 203 | 0.625953717 | cat4 | from_7893 | to_7893 | art_long | art_short | J |
| 106 | 203 | 0.576239146 | cat4 | from_7894 | to_7894 | art_long | art_short | J |
| 115 | 203 | -0.632544105 | cat4 | from_7895 | to_7895 | art_long | art_short | J |
| 117 | 203 | -0.583569922 | cat4 | from_7896 | to_7896 | art_long | art_short | J |
| 123 | 203 | -0.579676085 | cat4 | from_7897 | to_7897 | art_long | art_short | J |
| 127 | 203 | -0.609009503 | cat4 | from_7898 | to_7898 | art_long | art_short | J |
| 135 | 203 | 0.762288862 | cat4 | from_7899 | to_7899 | art_long | art_short | J |
| 139 | 203 | 0.592168766 | cat4 | from_7900 | to_7900 | art_long | art_short | J |
| 166 | 203 | 0.599589702 | cat4 | from_7901 | to_7901 | art_short | art_short | H |
| 170 | 203 | 0.641854871 | cat4 | from_7902 | to_7902 | art_short | art_short | H |

| | | | | | | | | |
|---|---|---|---|---|---|---|---|---|
| 171 | 203 | 0.601929795 | cat4 | from_7903 | to_7903 | art_short | art_short | H |
| 176 | 203 | 0.980308998 | cat4 | from_7904 | to_7904 | art_short | art_short | H |
| 44 | 204 | 0.606504591 | cat4 | from_7905 | to_7905 | art_long | art_short | J |
| 46 | 204 | 0.66058373 | cat4 | from_7906 | to_7906 | art_long | art_short | J |
| 48 | 204 | 0.594626146 | cat4 | from_7907 | to_7907 | art_long | art_short | J |
| 49 | 204 | -0.597273763 | cat4 | from_7908 | to_7908 | art_long | art_short | J |
| 51 | 204 | -0.647662665 | cat4 | from_7909 | to_7909 | art_long | art_short | J |
| 61 | 204 | 0.690842442 | cat4 | from_7910 | to_7910 | art_long | art_short | J |
| 70 | 204 | -0.633025852 | cat4 | from_7911 | to_7911 | art_long | art_short | J |
| 102 | 204 | 0.75028653 | cat4 | from_7912 | to_7912 | art_long | art_short | J |
| 104 | 204 | 0.629282654 | cat4 | from_7913 | to_7913 | art_long | art_short | J |
| 106 | 204 | 0.677377958 | cat4 | from_7914 | to_7914 | art_long | art_short | J |
| 135 | 204 | 0.808965705 | cat4 | from_7915 | to_7915 | art_long | art_short | J |
| 139 | 204 | 0.666934803 | cat4 | from_7916 | to_7916 | art_long | art_short | J |
| 166 | 204 | 0.688571206 | cat4 | from_7917 | to_7917 | art_short | art_short | H |
| 170 | 204 | 0.722948955 | cat4 | from_7918 | to_7918 | art_short | art_short | H |
| 171 | 204 | 0.68793753 | cat4 | from_7919 | to_7919 | art_short | art_short | H |
| 176 | 204 | 0.914536489 | cat4 | from_7920 | to_7920 | art_short | art_short | H |
| 203 | 204 | 0.951138449 | cat4 | from_7921 | to_7921 | art_short | art_short | H |
| 4 | 205 | 0.621478874 | cat4 | from_7922 | to_7922 | art_long | art_short | J |
| 7 | 205 | 0.719382913 | cat4 | from_7923 | to_7923 | art_long | art_short | J |
| 11 | 205 | 0.807635914 | cat4 | from_7924 | to_7924 | art_long | art_short | J |
| 22 | 205 | 0.665030882 | cat4 | from_7925 | to_7925 | art_long | art_short | J |
| 26 | 205 | -0.603597739 | cat4 | from_7926 | to_7926 | art_long | art_short | J |
| 37 | 205 | -0.676047995 | cat4 | from_7927 | to_7927 | art_long | art_short | J |
| 90 | 205 | -0.642993472 | cat4 | from_7928 | to_7928 | art_long | art_short | J |
| 115 | 205 | -0.650408082 | cat4 | from_7929 | to_7929 | art_long | art_short | J |
| 126 | 205 | 0.680036541 | cat4 | from_7930 | to_7930 | art_long | art_short | J |
| 129 | 205 | -0.670058056 | cat4 | from_7931 | to_7931 | art_long | art_short | J |
| 130 | 205 | -0.61387161 | cat4 | from_7932 | to_7932 | art_long | art_short | J |
| 135 | 205 | 0.652874779 | cat4 | from_7933 | to_7933 | art_long | art_short | J |
| 159 | 205 | 0.619909819 | cat4 | from_7934 | to_7934 | art_short | art_short | H |
| 160 | 205 | 0.628322805 | cat4 | from_7935 | to_7935 | art_short | art_short | H |
| 161 | 205 | 0.705902507 | cat4 | from_7936 | to_7936 | art_short | art_short | H |
| 188 | 205 | 0.642113745 | cat4 | from_7937 | to_7937 | art_short | art_short | H |
| 21 | 206 | -0.622662202 | cat4 | from_7938 | to_7938 | art_long | art_short | J |
| 42 | 206 | -0.625038636 | cat4 | from_7939 | to_7939 | art_long | art_short | J |
| 49 | 206 | -0.615908108 | cat4 | from_7940 | to_7940 | art_long | art_short | J |
| 54 | 206 | 0.838311408 | cat4 | from_7941 | to_7941 | art_long | art_short | J |
| 76 | 206 | 0.57862217 | cat4 | from_7942 | to_7942 | art_long | art_short | J |
| 77 | 206 | 0.886127933 | cat4 | from_7943 | to_7943 | art_long | art_short | J |
| 80 | 206 | 0.928903855 | cat4 | from_7944 | to_7944 | art_long | art_short | J |
| 81 | 206 | 0.696038453 | cat4 | from_7945 | to_7945 | art_long | art_short | J |
| 83 | 206 | 0.592144493 | cat4 | from_7946 | to_7946 | art_long | art_short | J |
| 85 | 206 | -0.65306759 | cat4 | from_7947 | to_7947 | art_long | art_short | J |
| 92 | 206 | -0.671434671 | cat4 | from_7948 | to_7948 | art_long | art_short | J |
| 95 | 206 | 0.764531578 | cat4 | from_7949 | to_7949 | art_long | art_short | J |
| 101 | 206 | -0.581137958 | cat4 | from_7950 | to_7950 | art_long | art_short | J |
| 103 | 206 | -0.589157922 | cat4 | from_7951 | to_7951 | art_long | art_short | J |
| 109 | 206 | 0.628220179 | cat4 | from_7952 | to_7952 | art_long | art_short | J |
| 119 | 206 | 0.698662059 | cat4 | from_7953 | to_7953 | art_long | art_short | J |
| 120 | 206 | 0.743158561 | cat4 | from_7954 | to_7954 | art_long | art_short | J |
| 127 | 206 | -0.637821197 | cat4 | from_7955 | to_7955 | art_long | art_short | J |
| 133 | 206 | -0.76271727 | cat4 | from_7956 | to_7956 | art_long | art_short | J |
| 140 | 206 | 0.714918409 | cat4 | from_7957 | to_7957 | art_long | art_short | J |
| 147 | 206 | -0.695559942 | cat4 | from_7958 | to_7958 | art_long | art_short | J |
| 177 | 206 | -0.653470029 | cat4 | from_7959 | to_7959 | art_short | art_short | H |
| 178 | 206 | -0.58901379 | cat4 | from_7960 | to_7960 | art_short | art_short | H |
| 195 | 206 | 0.792699972 | cat4 | from_7961 | to_7961 | art_short | art_short | H |
| 200 | 206 | 0.765732184 | cat4 | from_7962 | to_7962 | art_short | art_short | H |
| 4 | 207 | 0.6049457 | cat4 | from_7963 | to_7963 | art_long | art_short | J |
| 7 | 207 | 0.608313929 | cat4 | from_7964 | to_7964 | art_long | art_short | J |
| 22 | 207 | 0.628012775 | cat4 | from_7965 | to_7965 | art_long | art_short | J |
| 23 | 207 | 0.576915978 | cat4 | from_7966 | to_7966 | art_long | art_short | J |
| 34 | 207 | 0.920041048 | cat4 | from_7967 | to_7967 | art_long | art_short | J |
| 64 | 207 | 0.634510861 | cat4 | from_7968 | to_7968 | art_long | art_short | J |
| 65 | 207 | 0.93173771 | cat4 | from_7969 | to_7969 | art_long | art_short | J |
| 85 | 207 | 0.62599977 | cat4 | from_7970 | to_7970 | art_long | art_short | J |
| 113 | 207 | 0.903829705 | cat4 | from_7971 | to_7971 | art_long | art_short | J |
| 116 | 207 | 0.615232434 | cat4 | from_7972 | to_7972 | art_long | art_short | J |
| 117 | 207 | 0.639665549 | cat4 | from_7973 | to_7973 | art_long | art_short | J |
| 123 | 207 | 0.585252687 | cat4 | from_7974 | to_7974 | art_long | art_short | J |
| 125 | 207 | 0.632098951 | cat4 | from_7975 | to_7975 | art_long | art_short | J |
| 127 | 207 | 0.847848797 | cat4 | from_7976 | to_7976 | art_long | art_short | J |
| 159 | 207 | 0.672684488 | cat4 | from_7977 | to_7977 | art_short | art_short | H |
| 160 | 207 | 0.609299531 | cat4 | from_7978 | to_7978 | art_short | art_short | H |
| 188 | 207 | 0.665903315 | cat4 | from_7979 | to_7979 | art_short | art_short | H |
| 198 | 207 | 0.613190448 | cat4 | from_7980 | to_7980 | art_short | art_short | H |
| 199 | 207 | 0.666480669 | cat4 | from_7981 | to_7981 | art_short | art_short | H |
| 29 | 208 | 0.676357923 | cat4 | from_7982 | to_7982 | art_long | art_short | J |
| 66 | 208 | 0.692243262 | cat4 | from_7983 | to_7983 | art_long | art_short | J |
| 71 | 208 | 0.668099194 | cat4 | from_7984 | to_7984 | art_long | art_short | J |
| 82 | 208 | 0.631415779 | cat4 | from_7985 | to_7985 | art_long | art_short | J |
| 96 | 208 | 0.698954623 | cat4 | from_7986 | to_7986 | art_long | art_short | J |
| 150 | 208 | 0.604390711 | cat4 | from_7987 | to_7987 | art_long | art_short | J |
| 175 | 208 | 0.838992795 | cat4 | from_7988 | to_7988 | art_short | art_short | H |
| 181 | 208 | 0.732333591 | cat4 | from_7989 | to_7989 | art_short | art_short | H |
| 185 | 208 | 0.604592836 | cat4 | from_7990 | to_7990 | art_short | art_short | H |
| 189 | 208 | 0.743776614 | cat4 | from_7991 | to_7991 | art_short | art_short | H |
| 29 | 209 | 0.68977878 | cat4 | from_7992 | to_7992 | art_long | art_short | J |
| 66 | 209 | 0.651570169 | cat4 | from_7993 | to_7993 | art_long | art_short | J |
| 71 | 209 | 0.685828597 | cat4 | from_7994 | to_7994 | art_long | art_short | J |
| 82 | 209 | 0.8738767 | cat4 | from_7995 | to_7995 | art_long | art_short | J |
| 96 | 209 | 0.611764401 | cat4 | from_7996 | to_7996 | art_long | art_short | J |
| 121 | 209 | 0.728113534 | cat4 | from_7997 | to_7997 | art_long | art_short | J |
| 150 | 209 | 0.849808994 | cat4 | from_7998 | to_7998 | art_long | art_short | J |
| 175 | 209 | 0.963455448 | cat4 | from_7999 | to_7999 | art_short | art_short | H |
| 181 | 209 | 0.761006657 | cat4 | from_8000 | to_8000 | art_short | art_short | H |
| 185 | 209 | 0.797574921 | cat4 | from_8001 | to_8001 | art_short | art_short | H |
| 189 | 209 | 0.721667435 | cat4 | from_8002 | to_8002 | art_short | art_short | H |
| 208 | 209 | 0.861249074 | cat4 | from_8003 | to_8003 | art_short | art_short | H |
| 96 | 210 | 0.629785974 | cat4 | from_8004 | to_8004 | art_long | art_short | J |
| 175 | 210 | 0.870443625 | cat4 | from_8005 | to_8005 | art_short | art_short | H |
| 185 | 210 | 0.639226636 | cat4 | from_8006 | to_8006 | art_short | art_short | H |

| | | | | | | | | |
|---|---|---|---|---|---|---|---|---|
| 200 | 210 | 0.661116991 | cat4 | from_8007 | to_8007 | art_short | art_short | H |
| 208 | 210 | 0.879205796 | cat4 | from_8008 | to_8008 | art_short | art_short | H |
| 209 | 210 | 0.858050058 | cat4 | from_8009 | to_8009 | art_short | art_short | H |
| 30 | 211 | 0.97290195 | cat4 | from_8010 | to_8010 | art_long | art_short | J |
| 47 | 211 | 0.904176171 | cat4 | from_8011 | to_8011 | art_long | art_short | J |
| 55 | 211 | 0.858431758 | cat4 | from_8012 | to_8012 | art_long | art_short | J |
| 58 | 211 | 0.780604663 | cat4 | from_8013 | to_8013 | art_long | art_short | J |
| 69 | 211 | 0.777538963 | cat4 | from_8014 | to_8014 | art_long | art_short | J |
| 72 | 211 | 0.911160465 | cat4 | from_8015 | to_8015 | art_long | art_short | J |
| 83 | 211 | 0.664811559 | cat4 | from_8016 | to_8016 | art_long | art_short | J |
| 86 | 211 | 0.619605981 | cat4 | from_8017 | to_8017 | art_long | art_short | J |
| 87 | 211 | 0.925940197 | cat4 | from_8018 | to_8018 | art_long | art_short | J |
| 90 | 211 | 0.674317172 | cat4 | from_8019 | to_8019 | art_long | art_short | J |
| 91 | 211 | 0.576911342 | cat4 | from_8020 | to_8020 | art_long | art_short | J |
| 94 | 211 | 0.584699105 | cat4 | from_8021 | to_8021 | art_long | art_short | J |
| 95 | 211 | 0.665667037 | cat4 | from_8022 | to_8022 | art_long | art_short | J |
| 97 | 211 | 0.639025274 | cat4 | from_8023 | to_8023 | art_long | art_short | J |
| 109 | 211 | 0.851558691 | cat4 | from_8024 | to_8024 | art_long | art_short | J |
| 111 | 211 | 0.921533848 | cat4 | from_8025 | to_8025 | art_long | art_short | J |
| 119 | 211 | 0.7598123 | cat4 | from_8026 | to_8026 | art_long | art_short | J |
| 140 | 211 | 0.673305666 | cat4 | from_8027 | to_8027 | art_long | art_short | J |
| 172 | 211 | 0.596921937 | cat4 | from_8028 | to_8028 | art_short | art_short | H |
| 173 | 211 | 0.609915245 | cat4 | from_8029 | to_8029 | art_short | art_short | H |
| 195 | 211 | 0.665674969 | cat4 | from_8030 | to_8030 | art_short | art_short | H |
| 20 | 212 | -0.577277475 | cat4 | from_8031 | to_8031 | art_long | art_short | J |
| 56 | 212 | 0.699711479 | cat4 | from_8032 | to_8032 | art_long | art_short | J |
| 57 | 212 | 0.749999757 | cat4 | from_8033 | to_8033 | art_long | art_short | J |
| 72 | 212 | -0.600952732 | cat4 | from_8034 | to_8034 | art_long | art_short | J |
| 75 | 212 | 0.647391834 | cat4 | from_8035 | to_8035 | art_long | art_short | J |
| 86 | 212 | -0.619237956 | cat4 | from_8036 | to_8036 | art_long | art_short | J |
| 105 | 212 | -0.619910101 | cat4 | from_8037 | to_8037 | art_long | art_short | J |
| 123 | 212 | 0.659420514 | cat4 | from_8038 | to_8038 | art_long | art_short | J |
| 146 | 212 | 0.785202712 | cat4 | from_8039 | to_8039 | art_long | art_short | J |
| 193 | 212 | 0.613242232 | cat4 | from_8040 | to_8040 | art_short | art_short | H |
| 199 | 212 | 0.760431501 | cat4 | from_8041 | to_8041 | art_short | art_short | H |
| 17 | 213 | 0.654261429 | cat4 | from_8042 | to_8042 | art_long | art_short | J |
| 27 | 213 | -0.674482845 | cat4 | from_8043 | to_8043 | art_long | art_short | J |
| 36 | 213 | 0.629798085 | cat4 | from_8044 | to_8044 | art_long | art_short | J |
| 50 | 213 | 0.769851008 | cat4 | from_8045 | to_8045 | art_long | art_short | J |
| 55 | 213 | -0.588106612 | cat4 | from_8046 | to_8046 | art_long | art_short | J |
| 64 | 213 | -0.591811788 | cat4 | from_8047 | to_8047 | art_long | art_short | J |
| 109 | 213 | -0.5885961 | cat4 | from_8048 | to_8048 | art_long | art_short | J |
| 138 | 213 | 0.694296186 | cat4 | from_8049 | to_8049 | art_long | art_short | J |
| 143 | 213 | 0.613286233 | cat4 | from_8050 | to_8050 | art_long | art_short | J |
| 154 | 213 | 0.772341677 | cat4 | from_8051 | to_8051 | art_short | art_short | H |
| 172 | 213 | -0.666713452 | cat4 | from_8052 | to_8052 | art_short | art_short | H |
| 177 | 213 | 0.681900067 | cat4 | from_8053 | to_8053 | art_short | art_short | H |
| 35 | 214 | 0.849691504 | cat4 | from_8054 | to_8054 | art_long | art_short | J |
| 38 | 214 | 0.861990832 | cat4 | from_8055 | to_8055 | art_long | art_short | J |
| 48 | 214 | 0.820075793 | cat4 | from_8056 | to_8056 | art_long | art_short | J |
| 50 | 214 | 0.873097952 | cat4 | from_8057 | to_8057 | art_long | art_short | J |
| 59 | 214 | -0.592252706 | cat4 | from_8058 | to_8058 | art_long | art_short | J |
| 104 | 214 | 0.763151921 | cat4 | from_8059 | to_8059 | art_long | art_short | J |
| 132 | 214 | 0.834304291 | cat4 | from_8060 | to_8060 | art_long | art_short | J |
| 138 | 214 | 0.58296261 | cat4 | from_8061 | to_8061 | art_long | art_short | J |
| 143 | 214 | 0.887312283 | cat4 | from_8062 | to_8062 | art_long | art_short | J |
| 166 | 214 | 0.859874163 | cat4 | from_8063 | to_8063 | art_short | art_short | H |
| 177 | 214 | 0.623050199 | cat4 | from_8064 | to_8064 | art_short | art_short | H |
| 183 | 214 | 0.580193449 | cat4 | from_8065 | to_8065 | art_short | art_short | H |
| 213 | 214 | 0.713347804 | cat4 | from_8066 | to_8066 | art_short | art_short | H |
| 16 | 215 | 0.577049716 | cat4 | from_8067 | to_8067 | art_long | art_short | J |
| 29 | 215 | 0.607882181 | cat4 | from_8068 | to_8068 | art_long | art_short | J |
| 39 | 215 | 0.644138137 | cat4 | from_8069 | to_8069 | art_long | art_short | J |
| 71 | 215 | 0.61778545 | cat4 | from_8070 | to_8070 | art_long | art_short | J |
| 82 | 215 | 0.734516477 | cat4 | from_8071 | to_8071 | art_long | art_short | J |
| 98 | 215 | 0.613664937 | cat4 | from_8072 | to_8072 | art_long | art_short | J |
| 150 | 215 | 0.734105138 | cat4 | from_8073 | to_8073 | art_long | art_short | J |
| 164 | 215 | 0.617978861 | cat4 | from_8074 | to_8074 | art_short | art_short | H |
| 181 | 215 | 0.638883225 | cat4 | from_8075 | to_8075 | art_short | art_short | H |
| 209 | 215 | 0.607560845 | cat4 | from_8076 | to_8076 | art_short | art_short | H |
| 12 | 216 | 0.849452849 | cat4 | from_8077 | to_8077 | art_long | art_short | J |
| 15 | 216 | 0.735129801 | cat4 | from_8078 | to_8078 | art_long | art_short | J |
| 16 | 216 | 0.743346663 | cat4 | from_8079 | to_8079 | art_long | art_short | J |
| 18 | 216 | 0.866621104 | cat4 | from_8080 | to_8080 | art_long | art_short | J |
| 19 | 216 | 0.712880326 | cat4 | from_8081 | to_8081 | art_long | art_short | J |
| 39 | 216 | 0.693490008 | cat4 | from_8082 | to_8082 | art_long | art_short | J |
| 40 | 216 | 0.61563154 | cat4 | from_8083 | to_8083 | art_long | art_short | J |
| 79 | 216 | 0.71693268 | cat4 | from_8084 | to_8084 | art_long | art_short | J |
| 164 | 216 | 0.75390641 | cat4 | from_8085 | to_8085 | art_short | art_short | H |
| 167 | 216 | 0.846105638 | cat4 | from_8086 | to_8086 | art_short | art_short | H |
| 192 | 216 | 0.771482687 | cat4 | from_8087 | to_8087 | art_short | art_short | H |
| 197 | 216 | 0.81735325 | cat4 | from_8088 | to_8088 | art_short | art_short | H |
| 1 | 217 | 0.688482574 | cat4 | from_8089 | to_8089 | art_long | art_short | J |
| 14 | 217 | -0.627733942 | cat4 | from_8090 | to_8090 | art_long | art_short | J |
| 73 | 217 | -0.625743869 | cat4 | from_8091 | to_8091 | art_long | art_short | J |
| 107 | 217 | -0.592950782 | cat4 | from_8092 | to_8092 | art_long | art_short | J |
| 149 | 217 | -0.605699716 | cat4 | from_8093 | to_8093 | art_long | art_short | J |
| 30 | 218 | 0.96917979 | cat4 | from_8094 | to_8094 | art_long | art_short | J |
| 47 | 218 | 0.841636509 | cat4 | from_8095 | to_8095 | art_long | art_short | J |
| 55 | 218 | 0.890003187 | cat4 | from_8096 | to_8096 | art_long | art_short | J |
| 58 | 218 | 0.863918028 | cat4 | from_8097 | to_8097 | art_long | art_short | J |
| 69 | 218 | 0.844921935 | cat4 | from_8098 | to_8098 | art_long | art_short | J |
| 72 | 218 | 0.824438867 | cat4 | from_8099 | to_8099 | art_long | art_short | J |
| 77 | 218 | 0.733638292 | cat4 | from_8100 | to_8100 | art_long | art_short | J |
| 80 | 218 | 0.744112826 | cat4 | from_8101 | to_8101 | art_long | art_short | J |
| 83 | 218 | 0.767127183 | cat4 | from_8102 | to_8102 | art_long | art_short | J |
| 86 | 218 | 0.580182347 | cat4 | from_8103 | to_8103 | art_long | art_short | J |
| 87 | 218 | 0.899571784 | cat4 | from_8104 | to_8104 | art_long | art_short | J |
| 94 | 218 | 0.591230746 | cat4 | from_8105 | to_8105 | art_long | art_short | J |
| 95 | 218 | 0.806165972 | cat4 | from_8106 | to_8106 | art_long | art_short | J |
| 109 | 218 | 0.913651719 | cat4 | from_8107 | to_8107 | art_long | art_short | J |
| 111 | 218 | 0.7907789 | cat4 | from_8108 | to_8108 | art_long | art_short | J |
| 119 | 218 | 0.893569947 | cat4 | from_8109 | to_8109 | art_long | art_short | J |
| 140 | 218 | 0.807977418 | cat4 | from_8110 | to_8110 | art_long | art_short | J |

| | | | | | | | | |
|---|---|---|---|---|---|---|---|---|
| 172 | 218 | 0.62519885 | cat4 | from_8111 | to_8111 | art_short | art_short | H |
| 173 | 218 | 0.611301668 | cat4 | from_8112 | to_8112 | art_short | art_short | H |
| 174 | 218 | 0.62238705 | cat4 | from_8113 | to_8113 | art_short | art_short | H |
| 194 | 218 | 0.623068063 | cat4 | from_8114 | to_8114 | art_short | art_short | H |
| 195 | 218 | 0.778228408 | cat4 | from_8115 | to_8115 | art_short | art_short | H |
| 206 | 218 | 0.671821631 | cat4 | from_8116 | to_8116 | art_short | art_short | H |
| 211 | 218 | 0.948163793 | cat4 | from_8117 | to_8117 | art_short | art_short | H |
| 1 | 219 | 0.636640814 | cat4 | from_8118 | to_8118 | art_long | art_untreat | E |
| 30 | 219 | 0.739823996 | cat4 | from_8119 | to_8119 | art_long | art_untreat | E |
| 31 | 219 | 0.588294695 | cat4 | from_8120 | to_8120 | art_long | art_untreat | E |
| 72 | 219 | 0.62781643 | cat4 | from_8121 | to_8121 | art_long | art_untreat | E |
| 211 | 219 | 0.778694817 | cat4 | from_8122 | to_8122 | art_short | art_untreat | C |
| 218 | 219 | 0.748738208 | cat4 | from_8123 | to_8123 | art_short | art_untreat | C |
| 13 | 220 | 0.605577149 | cat4 | from_8124 | to_8124 | art_long | art_untreat | E |
| 15 | 220 | 0.622720266 | cat4 | from_8125 | to_8125 | art_long | art_untreat | E |
| 16 | 220 | 0.655810831 | cat4 | from_8126 | to_8126 | art_long | art_untreat | E |
| 21 | 220 | 0.590900838 | cat4 | from_8127 | to_8127 | art_long | art_untreat | E |
| 39 | 220 | 0.620710592 | cat4 | from_8128 | to_8128 | art_long | art_untreat | E |
| 114 | 220 | 0.626338561 | cat4 | from_8129 | to_8129 | art_long | art_untreat | E |
| 116 | 220 | 0.599130949 | cat4 | from_8130 | to_8130 | art_long | art_untreat | E |
| 162 | 220 | 0.765996209 | cat4 | from_8131 | to_8131 | art_short | art_untreat | C |
| 163 | 220 | 0.840182794 | cat4 | from_8132 | to_8132 | art_short | art_untreat | C |
| 164 | 220 | 0.766235657 | cat4 | from_8133 | to_8133 | art_short | art_untreat | C |
| 165 | 220 | 0.743057951 | cat4 | from_8134 | to_8134 | art_short | art_untreat | C |
| 168 | 220 | 0.666007522 | cat4 | from_8135 | to_8135 | art_short | art_untreat | C |
| 169 | 220 | 0.936540536 | cat4 | from_8136 | to_8136 | art_short | art_untreat | C |
| 180 | 220 | 0.66549607 | cat4 | from_8137 | to_8137 | art_short | art_untreat | C |
| 187 | 220 | 0.684970473 | cat4 | from_8138 | to_8138 | art_short | art_untreat | C |
| 192 | 220 | 0.705451816 | cat4 | from_8139 | to_8139 | art_short | art_untreat | C |
| 196 | 220 | 0.738742789 | cat4 | from_8140 | to_8140 | art_short | art_untreat | C |
| 197 | 220 | 0.763187525 | cat4 | from_8141 | to_8141 | art_short | art_untreat | C |
| 89 | 221 | 0.865842732 | cat4 | from_8142 | to_8142 | art_long | art_untreat | E |
| 32 | 222 | -0.654957919 | cat4 | from_8143 | to_8143 | art_long | art_untreat | E |
| 43 | 222 | -0.606080375 | cat4 | from_8144 | to_8144 | art_long | art_untreat | E |
| 49 | 222 | 0.632776388 | cat4 | from_8145 | to_8145 | art_long | art_untreat | E |
| 56 | 222 | 0.77180242 | cat4 | from_8146 | to_8146 | art_long | art_untreat | E |
| 74 | 222 | -0.613973507 | cat4 | from_8147 | to_8147 | art_long | art_untreat | E |
| 78 | 222 | -0.602546528 | cat4 | from_8148 | to_8148 | art_long | art_untreat | E |
| 85 | 222 | 0.669328207 | cat4 | from_8149 | to_8149 | art_long | art_untreat | E |
| 149 | 222 | -0.587064845 | cat4 | from_8150 | to_8150 | art_long | art_untreat | E |
| 182 | 222 | 0.828313159 | cat4 | from_8151 | to_8151 | art_short | art_untreat | C |
| 193 | 222 | 0.594912768 | cat4 | from_8152 | to_8152 | art_short | art_untreat | C |
| 199 | 222 | 0.77027824 | cat4 | from_8153 | to_8153 | art_short | art_untreat | C |
| 201 | 222 | 0.613636861 | cat4 | from_8154 | to_8154 | art_short | art_untreat | C |
| 44 | 223 | 0.665891999 | cat4 | from_8155 | to_8155 | art_long | art_untreat | E |
| 46 | 223 | 0.672870326 | cat4 | from_8156 | to_8156 | art_long | art_untreat | E |
| 52 | 223 | 0.808704579 | cat4 | from_8157 | to_8157 | art_long | art_untreat | E |
| 60 | 223 | 0.637505247 | cat4 | from_8158 | to_8158 | art_long | art_untreat | E |
| 61 | 223 | 0.730573724 | cat4 | from_8159 | to_8159 | art_long | art_untreat | E |
| 99 | 223 | 0.662519526 | cat4 | from_8160 | to_8160 | art_long | art_untreat | E |
| 102 | 223 | 0.63447706 | cat4 | from_8161 | to_8161 | art_long | art_untreat | E |
| 105 | 223 | 0.597771875 | cat4 | from_8162 | to_8162 | art_long | art_untreat | E |
| 106 | 223 | 0.661482355 | cat4 | from_8163 | to_8163 | art_long | art_untreat | E |
| 128 | 223 | 0.661113821 | cat4 | from_8164 | to_8164 | art_long | art_untreat | E |
| 148 | 223 | 0.62284071 | cat4 | from_8165 | to_8165 | art_long | art_untreat | E |
| 176 | 223 | 0.826491521 | cat4 | from_8166 | to_8166 | art_short | art_untreat | C |
| 183 | 223 | 0.844604081 | cat4 | from_8167 | to_8167 | art_short | art_untreat | C |
| 203 | 223 | 0.753450216 | cat4 | from_8168 | to_8168 | art_short | art_untreat | C |
| 204 | 223 | 0.742286362 | cat4 | from_8169 | to_8169 | art_short | art_untreat | C |
| 214 | 223 | 0.588472025 | cat4 | from_8170 | to_8170 | art_short | art_untreat | C |
| 29 | 224 | 0.771359585 | cat4 | from_8171 | to_8171 | art_long | art_untreat | E |
| 71 | 224 | 0.72161133 | cat4 | from_8172 | to_8172 | art_long | art_untreat | E |
| 82 | 224 | 0.625266223 | cat4 | from_8173 | to_8173 | art_long | art_untreat | E |
| 124 | 224 | 0.769577745 | cat4 | from_8174 | to_8174 | art_long | art_untreat | E |
| 150 | 224 | 0.652628974 | cat4 | from_8175 | to_8175 | art_long | art_untreat | E |
| 175 | 224 | 0.682936488 | cat4 | from_8176 | to_8176 | art_short | art_untreat | C |
| 181 | 224 | 0.775932399 | cat4 | from_8177 | to_8177 | art_short | art_untreat | C |
| 189 | 224 | 0.588961799 | cat4 | from_8178 | to_8178 | art_short | art_untreat | C |
| 208 | 224 | 0.737797961 | cat4 | from_8179 | to_8179 | art_short | art_untreat | C |
| 209 | 224 | 0.728662846 | cat4 | from_8180 | to_8180 | art_short | art_untreat | C |
| 17 | 225 | -0.577697461 | cat4 | from_8181 | to_8181 | art_long | art_untreat | E |
| 30 | 225 | 0.83707262 | cat4 | from_8182 | to_8182 | art_long | art_untreat | E |
| 36 | 225 | -0.591220806 | cat4 | from_8183 | to_8183 | art_long | art_untreat | E |
| 47 | 225 | 0.689842003 | cat4 | from_8184 | to_8184 | art_long | art_untreat | E |
| 55 | 225 | 0.71032289 | cat4 | from_8185 | to_8185 | art_long | art_untreat | E |
| 58 | 225 | 0.592990108 | cat4 | from_8186 | to_8186 | art_long | art_untreat | E |
| 72 | 225 | 0.67529034 | cat4 | from_8187 | to_8187 | art_long | art_untreat | E |
| 80 | 225 | 0.590220722 | cat4 | from_8188 | to_8188 | art_long | art_untreat | E |
| 87 | 225 | 0.712739023 | cat4 | from_8189 | to_8189 | art_long | art_untreat | E |
| 95 | 225 | 0.642340783 | cat4 | from_8190 | to_8190 | art_long | art_untreat | E |
| 109 | 225 | 0.717208997 | cat4 | from_8191 | to_8191 | art_long | art_untreat | E |
| 111 | 225 | 0.636952033 | cat4 | from_8192 | to_8192 | art_long | art_untreat | E |
| 119 | 225 | 0.642820544 | cat4 | from_8193 | to_8193 | art_long | art_untreat | E |
| 140 | 225 | 0.603638783 | cat4 | from_8194 | to_8194 | art_long | art_untreat | E |
| 173 | 225 | 0.577278985 | cat4 | from_8195 | to_8195 | art_short | art_untreat | C |
| 195 | 225 | 0.77999191 | cat4 | from_8196 | to_8196 | art_short | art_untreat | C |
| 206 | 225 | 0.677459095 | cat4 | from_8197 | to_8197 | art_short | art_untreat | C |
| 211 | 225 | 0.8713182 | cat4 | from_8198 | to_8198 | art_short | art_untreat | C |
| 218 | 225 | 0.901286663 | cat4 | from_8199 | to_8199 | art_short | art_untreat | C |
| 219 | 225 | 0.916519335 | cat4 | from_8200 | to_8200 | art_untreat | art_untreat | A |
| 14 | 226 | -0.592599976 | cat4 | from_8201 | to_8201 | art_long | art_untreat | E |
| 52 | 226 | 0.632942369 | cat4 | from_8202 | to_8202 | art_long | art_untreat | E |
| 59 | 226 | -0.591721918 | cat4 | from_8203 | to_8203 | art_long | art_untreat | E |
| 73 | 226 | -0.600027741 | cat4 | from_8204 | to_8204 | art_long | art_untreat | E |
| 74 | 226 | -0.646334092 | cat4 | from_8205 | to_8205 | art_long | art_untreat | E |
| 78 | 226 | -0.605999536 | cat4 | from_8206 | to_8206 | art_long | art_untreat | E |
| 99 | 226 | 0.646507902 | cat4 | from_8207 | to_8207 | art_long | art_untreat | E |
| 102 | 226 | 0.582199161 | cat4 | from_8208 | to_8208 | art_long | art_untreat | E |
| 105 | 226 | 0.697204722 | cat4 | from_8209 | to_8209 | art_long | art_untreat | E |
| 176 | 226 | 0.707837166 | cat4 | from_8210 | to_8210 | art_short | art_untreat | C |
| 183 | 226 | 0.76721009 | cat4 | from_8211 | to_8211 | art_short | art_untreat | C |
| 203 | 226 | 0.648489749 | cat4 | from_8212 | to_8212 | art_short | art_untreat | C |
| 217 | 226 | 0.651811052 | cat4 | from_8213 | to_8213 | art_short | art_untreat | C |
| 223 | 226 | 0.880982104 | cat4 | from_8214 | to_8214 | art_untreat | art_untreat | A |

| | | | | | | | | |
|---|---|---|---|---|---|---|---|---|
| 198 | 227 | 0.719353514 | cat4 | from_8215 | to_8215 | art_short | ec | K |
| 212 | 227 | 0.580226542 | cat4 | from_8216 | to_8216 | art_short | ec | K |
| 113 | 228 | 0.788992179 | cat4 | from_8217 | to_8217 | art_long | ec | N |
| 207 | 228 | 0.668245969 | cat4 | from_8218 | to_8218 | art_short | ec | K |
| 212 | 228 | 0.654876787 | cat4 | from_8219 | to_8219 | art_short | ec | K |
| 227 | 228 | 0.899474078 | cat4 | from_8220 | to_8220 | ec | ec | P |
| 12 | 229 | 0.683494734 | cat4 | from_8221 | to_8221 | art_long | ec | N |
| 40 | 229 | 0.611709073 | cat4 | from_8222 | to_8222 | art_long | ec | N |
| 79 | 229 | 0.582358007 | cat4 | from_8223 | to_8223 | art_long | ec | N |
| 156 | 229 | 0.621886059 | cat4 | from_8224 | to_8224 | art_short | ec | K |
| 158 | 229 | 0.585294913 | cat4 | from_8225 | to_8225 | art_short | ec | K |
| 163 | 229 | 0.619235349 | cat4 | from_8226 | to_8226 | art_short | ec | K |
| 164 | 229 | 0.613566444 | cat4 | from_8227 | to_8227 | art_short | ec | K |
| 165 | 229 | 0.615933097 | cat4 | from_8228 | to_8228 | art_short | ec | K |
| 167 | 229 | 0.6957656 | cat4 | from_8229 | to_8229 | art_short | ec | K |
| 197 | 229 | 0.74289793 | cat4 | from_8230 | to_8230 | art_short | ec | K |
| 217 | 229 | 0.744527782 | cat4 | from_8231 | to_8231 | art_short | ec | K |
| 220 | 229 | 0.626313228 | cat4 | from_8232 | to_8232 | art_untreat | ec | F |
| 53 | 230 | 0.680611512 | cat4 | from_8233 | to_8233 | art_long | ec | N |
| 168 | 230 | 0.824867604 | cat4 | from_8234 | to_8234 | art_short | ec | K |
| 169 | 230 | 0.609577341 | cat4 | from_8235 | to_8235 | art_short | ec | K |
| 179 | 230 | 0.70693253 | cat4 | from_8236 | to_8236 | art_short | ec | K |
| 180 | 230 | 0.886638351 | cat4 | from_8237 | to_8237 | art_short | ec | K |
| 187 | 230 | 0.886259555 | cat4 | from_8238 | to_8238 | art_short | ec | K |
| 220 | 230 | 0.744927105 | cat4 | from_8239 | to_8239 | art_untreat | ec | F |
| 1 | 231 | 0.636429467 | cat4 | from_8240 | to_8240 | art_long | ec | N |
| 34 | 231 | 0.782238946 | cat4 | from_8241 | to_8241 | art_long | ec | N |
| 65 | 231 | 0.644150672 | cat4 | from_8242 | to_8242 | art_long | ec | N |
| 113 | 231 | 0.725047645 | cat4 | from_8243 | to_8243 | art_long | ec | N |
| 127 | 231 | 0.667022919 | cat4 | from_8244 | to_8244 | art_long | ec | N |
| 159 | 231 | 0.579362498 | cat4 | from_8245 | to_8245 | art_short | ec | K |
| 207 | 231 | 0.834030385 | cat4 | from_8246 | to_8246 | art_short | ec | K |
| 227 | 231 | 0.623902153 | cat4 | from_8247 | to_8247 | ec | ec | P |
| 228 | 231 | 0.664671245 | cat4 | from_8248 | to_8248 | ec | ec | P |
| 139 | 232 | 0.644768196 | cat4 | from_8249 | to_8249 | art_long | ec | N |
| 170 | 232 | 0.695634813 | cat4 | from_8250 | to_8250 | art_short | ec | K |
| 171 | 232 | 0.707165731 | cat4 | from_8251 | to_8251 | art_short | ec | K |
| 178 | 232 | 0.616486213 | cat4 | from_8252 | to_8252 | art_short | ec | K |
| 184 | 232 | 0.619127735 | cat4 | from_8253 | to_8253 | art_short | ec | K |
| 223 | 232 | 0.718262312 | cat4 | from_8254 | to_8254 | art_untreat | ec | F |
| 226 | 232 | 0.721653495 | cat4 | from_8255 | to_8255 | art_untreat | ec | F |
| 6 | 233 | -0.5913683 | cat4 | from_8256 | to_8256 | art_long | ec | N |
| 99 | 233 | -0.620340731 | cat4 | from_8257 | to_8257 | art_long | ec | N |
| 105 | 233 | -0.656067497 | cat4 | from_8258 | to_8258 | art_long | ec | N |
| 131 | 233 | 0.651828345 | cat4 | from_8259 | to_8259 | art_long | ec | N |
| 148 | 233 | -0.661736266 | cat4 | from_8260 | to_8260 | art_long | ec | N |
| 152 | 233 | -0.628320911 | cat4 | from_8261 | to_8261 | art_long | ec | N |
| 153 | 233 | -0.618761576 | cat4 | from_8262 | to_8262 | art_long | ec | N |
| 195 | 233 | 0.617262788 | cat4 | from_8263 | to_8263 | art_short | ec | K |
| 200 | 233 | 0.682525542 | cat4 | from_8264 | to_8264 | art_short | ec | K |
| 29 | 234 | 0.608244502 | cat4 | from_8265 | to_8265 | art_long | ec | N |
| 68 | 234 | -0.577914396 | cat4 | from_8266 | to_8266 | art_long | ec | N |
| 98 | 234 | 0.740970246 | cat4 | from_8267 | to_8267 | art_long | ec | N |
| 121 | 234 | 0.723295442 | cat4 | from_8268 | to_8268 | art_long | ec | N |
| 150 | 234 | 0.638559743 | cat4 | from_8269 | to_8269 | art_long | ec | N |
| 175 | 234 | 0.727303849 | cat4 | from_8270 | to_8270 | art_short | ec | K |
| 181 | 234 | 0.589627983 | cat4 | from_8271 | to_8271 | art_short | ec | K |
| 208 | 234 | 0.587187569 | cat4 | from_8272 | to_8272 | art_short | ec | K |
| 209 | 234 | 0.746872291 | cat4 | from_8273 | to_8273 | art_short | ec | K |
| 224 | 234 | 0.583736176 | cat4 | from_8274 | to_8274 | art_untreat | ec | F |
| 29 | 235 | 0.739546957 | cat4 | from_8275 | to_8275 | art_long | ec | N |
| 41 | 235 | 0.596031695 | cat4 | from_8276 | to_8276 | art_long | ec | N |
| 71 | 235 | 0.64218564 | cat4 | from_8277 | to_8277 | art_long | ec | N |
| 98 | 235 | 0.685093199 | cat4 | from_8278 | to_8278 | art_long | ec | N |
| 121 | 235 | 0.626069184 | cat4 | from_8279 | to_8279 | art_long | ec | N |
| 175 | 235 | 0.620415944 | cat4 | from_8280 | to_8280 | art_short | ec | K |
| 181 | 235 | 0.689870309 | cat4 | from_8281 | to_8281 | art_short | ec | K |
| 208 | 235 | 0.66387331 | cat4 | from_8282 | to_8282 | art_short | ec | K |
| 209 | 235 | 0.649917765 | cat4 | from_8283 | to_8283 | art_short | ec | K |
| 224 | 235 | 0.653157423 | cat4 | from_8284 | to_8284 | art_untreat | ec | F |
| 234 | 235 | 0.921773368 | cat4 | from_8285 | to_8285 | ec | ec | P |
| 153 | 236 | -0.599645006 | cat4 | from_8286 | to_8286 | art_long | ec | N |
| 202 | 236 | 0.58501229 | cat4 | from_8287 | to_8287 | art_short | ec | K |
| 227 | 236 | 0.760616778 | cat4 | from_8288 | to_8288 | ec | ec | P |
| 228 | 236 | 0.644136032 | cat4 | from_8289 | to_8289 | ec | ec | P |
| 232 | 236 | 0.70339614 | cat4 | from_8290 | to_8290 | ec | ec | P |
| 1 | 237 | 0.68190534 | cat4 | from_8291 | to_8291 | art_long | ec | N |
| 178 | 237 | 0.738716926 | cat4 | from_8292 | to_8292 | art_short | ec | K |
| 184 | 237 | 0.585683591 | cat4 | from_8293 | to_8293 | art_short | ec | K |
| 223 | 237 | 0.661389477 | cat4 | from_8294 | to_8294 | art_untreat | ec | F |
| 226 | 237 | 0.668183502 | cat4 | from_8295 | to_8295 | art_untreat | ec | F |
| 232 | 237 | 0.946220418 | cat4 | from_8296 | to_8296 | ec | ec | P |
| 236 | 237 | 0.712606106 | cat4 | from_8297 | to_8297 | ec | ec | P |
| 1 | 238 | 0.589148534 | cat4 | from_8298 | to_8298 | art_long | ec | N |
| 32 | 238 | -0.635398214 | cat4 | from_8299 | to_8299 | art_long | ec | N |
| 53 | 238 | 0.813485318 | cat4 | from_8300 | to_8300 | art_long | ec | N |
| 164 | 238 | 0.681459172 | cat4 | from_8301 | to_8301 | art_short | ec | K |
| 168 | 238 | 0.762011448 | cat4 | from_8302 | to_8302 | art_short | ec | K |
| 179 | 238 | 0.76388597 | cat4 | from_8303 | to_8303 | art_short | ec | K |
| 180 | 238 | 0.774667528 | cat4 | from_8304 | to_8304 | art_short | ec | K |
| 187 | 238 | 0.730928243 | cat4 | from_8305 | to_8305 | art_short | ec | K |
| 197 | 238 | 0.669007511 | cat4 | from_8306 | to_8306 | art_short | ec | K |
| 216 | 238 | 0.606919514 | cat4 | from_8307 | to_8307 | art_short | ec | K |
| 230 | 238 | 0.859079551 | cat4 | from_8308 | to_8308 | ec | ec | P |
| 14 | 239 | -0.592614328 | cat4 | from_8309 | to_8309 | art_long | ec | N |
| 43 | 239 | -0.591911013 | cat4 | from_8310 | to_8310 | art_long | ec | N |
| 56 | 239 | 0.617557543 | cat4 | from_8311 | to_8311 | art_long | ec | N |
| 73 | 239 | -0.591199246 | cat4 | from_8312 | to_8312 | art_long | ec | N |
| 107 | 239 | -0.595888094 | cat4 | from_8313 | to_8313 | art_long | ec | N |
| 149 | 239 | -0.603859551 | cat4 | from_8314 | to_8314 | art_long | ec | N |
| 182 | 239 | 0.693908805 | cat4 | from_8315 | to_8315 | art_short | ec | K |
| 193 | 239 | 0.683825 | cat4 | from_8316 | to_8316 | art_short | ec | K |
| 199 | 239 | 0.696591375 | cat4 | from_8317 | to_8317 | art_short | ec | K |
| 201 | 239 | 0.615913825 | cat4 | from_8318 | to_8318 | art_short | ec | K |

| | | | | | | | | |
|---|---|---|---|---|---|---|---|---|
| 217 | 239 | 0.641473795 | cat4 | from_8319 | to_8319 | art_short | ec | K |
| 222 | 239 | 0.917632477 | cat4 | from_8320 | to_8320 | art_untreat | ec | F |
| 48 | 240 | 0.584105558 | cat4 | from_8321 | to_8321 | art_long | ec | N |
| 54 | 240 | -0.590947502 | cat4 | from_8322 | to_8322 | art_long | ec | N |
| 68 | 240 | -0.650846618 | cat4 | from_8323 | to_8323 | art_long | ec | N |
| 81 | 240 | -0.630539305 | cat4 | from_8324 | to_8324 | art_long | ec | N |
| 83 | 240 | -0.653610364 | cat4 | from_8325 | to_8325 | art_long | ec | N |
| 91 | 240 | -0.587701903 | cat4 | from_8326 | to_8326 | art_long | ec | N |
| 93 | 240 | -0.587800223 | cat4 | from_8327 | to_8327 | art_long | ec | N |
| 120 | 240 | -0.593841027 | cat4 | from_8328 | to_8328 | art_long | ec | N |
| 134 | 240 | 0.746770965 | cat4 | from_8329 | to_8329 | art_long | ec | N |
| 138 | 240 | 0.623806137 | cat4 | from_8330 | to_8330 | art_long | ec | N |
| 140 | 240 | -0.634247057 | cat4 | from_8331 | to_8331 | art_long | ec | N |
| 154 | 240 | 0.617769681 | cat4 | from_8332 | to_8332 | art_short | ec | K |
| 166 | 240 | 0.710443118 | cat4 | from_8333 | to_8333 | art_short | ec | K |
| 170 | 240 | 0.619546964 | cat4 | from_8334 | to_8334 | art_short | ec | K |
| 171 | 240 | 0.623118017 | cat4 | from_8335 | to_8335 | art_short | ec | K |
| 177 | 240 | 0.750551082 | cat4 | from_8336 | to_8336 | art_short | ec | K |
| 232 | 240 | 0.657255668 | cat4 | from_8337 | to_8337 | ec | ec | P |
| 237 | 240 | 0.652196149 | cat4 | from_8338 | to_8338 | ec | ec | P |
| 48 | 241 | 0.603548822 | cat4 | from_8339 | to_8339 | art_long | ec | N |
| 68 | 241 | -0.628011755 | cat4 | from_8340 | to_8340 | art_long | ec | N |
| 72 | 241 | -0.597038795 | cat4 | from_8341 | to_8341 | art_long | ec | N |
| 83 | 241 | -0.602849884 | cat4 | from_8342 | to_8342 | art_long | ec | N |
| 91 | 241 | -0.670413245 | cat4 | from_8343 | to_8343 | art_long | ec | N |
| 104 | 241 | 0.626154082 | cat4 | from_8344 | to_8344 | art_long | ec | N |
| 134 | 241 | 0.733538316 | cat4 | from_8345 | to_8345 | art_long | ec | N |
| 138 | 241 | 0.663547109 | cat4 | from_8346 | to_8346 | art_long | ec | N |
| 139 | 241 | 0.618765253 | cat4 | from_8347 | to_8347 | art_long | ec | N |
| 143 | 241 | 0.646918197 | cat4 | from_8348 | to_8348 | art_long | ec | N |
| 154 | 241 | 0.72702738 | cat4 | from_8349 | to_8349 | art_short | ec | K |
| 166 | 241 | 0.736988336 | cat4 | from_8350 | to_8350 | art_short | ec | K |
| 170 | 241 | 0.696306105 | cat4 | from_8351 | to_8351 | art_short | ec | K |
| 171 | 241 | 0.668360169 | cat4 | from_8352 | to_8352 | art_short | ec | K |
| 204 | 241 | 0.626052481 | cat4 | from_8353 | to_8353 | art_short | ec | K |
| 232 | 241 | 0.607312983 | cat4 | from_8354 | to_8354 | ec | ec | P |
| 233 | 241 | 0.706305809 | cat4 | from_8355 | to_8355 | ec | ec | P |
| 240 | 241 | 0.902779935 | cat4 | from_8356 | to_8356 | ec | ec | P |
| 156 | 242 | 0.672417931 | cat4 | from_8357 | to_8357 | art_short | ec | K |
| 157 | 242 | 0.684146975 | cat4 | from_8358 | to_8358 | art_short | ec | K |
| 158 | 242 | 0.662896861 | cat4 | from_8359 | to_8359 | art_short | ec | K |
| 198 | 242 | 0.823799065 | cat4 | from_8360 | to_8360 | art_short | ec | K |
| 201 | 242 | 0.675548187 | cat4 | from_8361 | to_8361 | art_short | ec | K |
| 217 | 242 | 0.603848192 | cat4 | from_8362 | to_8362 | art_short | ec | K |
| 227 | 242 | 0.697948886 | cat4 | from_8363 | to_8363 | ec | ec | P |
| 229 | 242 | 0.621201847 | cat4 | from_8364 | to_8364 | ec | ec | P |
| 231 | 242 | 0.641505424 | cat4 | from_8365 | to_8365 | ec | ec | P |
| 232 | 242 | 0.58092804 | cat4 | from_8366 | to_8366 | ec | ec | P |
| 236 | 242 | 0.616801795 | cat4 | from_8367 | to_8367 | ec | ec | P |
| 239 | 242 | 0.586263604 | cat4 | from_8368 | to_8368 | ec | ec | P |
| 113 | 243 | 0.852318673 | cat4 | from_8369 | to_8369 | art_long | ec | N |
| 141 | 243 | -0.579202569 | cat4 | from_8370 | to_8370 | art_long | ec | N |
| 152 | 243 | -0.593014829 | cat4 | from_8371 | to_8371 | art_long | ec | N |
| 153 | 243 | -0.585083561 | cat4 | from_8372 | to_8372 | art_long | ec | N |
| 207 | 243 | 0.750075859 | cat4 | from_8373 | to_8373 | art_short | ec | K |
| 212 | 243 | 0.581939112 | cat4 | from_8374 | to_8374 | art_short | ec | K |
| 227 | 243 | 0.84441287 | cat4 | from_8375 | to_8375 | ec | ec | P |
| 228 | 243 | 0.976082645 | cat4 | from_8376 | to_8376 | ec | ec | P |
| 231 | 243 | 0.792527943 | cat4 | from_8377 | to_8377 | ec | ec | P |
| 236 | 243 | 0.635394762 | cat4 | from_8378 | to_8378 | ec | ec | P |
| 239 | 243 | 0.594042756 | cat4 | from_8379 | to_8379 | ec | ec | P |
| 213 | 244 | 0.602523839 | cat4 | from_8380 | to_8380 | art_short | ec | K |
| 242 | 244 | 0.697242305 | cat4 | from_8381 | to_8381 | ec | ec | P |
| 31 | 245 | 0.60715091 | cat4 | from_8382 | to_8382 | art_long | ec | N |
| 33 | 245 | 0.587132514 | cat4 | from_8383 | to_8383 | art_long | ec | N |
| 42 | 245 | -0.58331171 | cat4 | from_8384 | to_8384 | art_long | ec | N |
| 54 | 245 | 0.599384966 | cat4 | from_8385 | to_8385 | art_long | ec | N |
| 81 | 245 | 0.69892364 | cat4 | from_8386 | to_8386 | art_long | ec | N |
| 100 | 245 | -0.594250705 | cat4 | from_8387 | to_8387 | art_long | ec | N |
| 134 | 245 | -0.585487307 | cat4 | from_8388 | to_8388 | art_long | ec | N |
| 145 | 245 | 0.662450578 | cat4 | from_8389 | to_8389 | art_long | ec | N |
| 206 | 245 | 0.628877255 | cat4 | from_8390 | to_8390 | art_short | ec | K |
| 219 | 245 | 0.764357973 | cat4 | from_8391 | to_8391 | art_untreat | ec | F |
| 225 | 245 | 0.652609494 | cat4 | from_8392 | to_8392 | art_untreat | ec | F |
| 229 | 245 | 0.639300918 | cat4 | from_8393 | to_8393 | ec | ec | P |
| 242 | 245 | 0.699398747 | cat4 | from_8394 | to_8394 | ec | ec | P |
| 49 | 246 | -0.642453202 | cat4 | from_8395 | to_8395 | art_long | ec | N |
| 51 | 246 | -0.715881995 | cat4 | from_8396 | to_8396 | art_long | ec | N |
| 63 | 246 | -0.619437408 | cat4 | from_8397 | to_8397 | art_long | ec | N |
| 70 | 246 | -0.691014624 | cat4 | from_8398 | to_8398 | art_long | ec | N |
| 75 | 246 | -0.653297874 | cat4 | from_8399 | to_8399 | art_long | ec | N |
| 84 | 246 | -0.612598604 | cat4 | from_8400 | to_8400 | art_long | ec | N |
| 94 | 246 | 0.586023425 | cat4 | from_8401 | to_8401 | art_long | ec | N |
| 146 | 246 | -0.58132486 | cat4 | from_8402 | to_8402 | art_long | ec | N |
| 176 | 246 | 0.603080871 | cat4 | from_8403 | to_8403 | art_short | ec | K |
| 191 | 246 | 0.635430994 | cat4 | from_8404 | to_8404 | art_short | ec | K |
| 195 | 246 | 0.704122163 | cat4 | from_8405 | to_8405 | art_short | ec | K |
| 200 | 246 | 0.672511165 | cat4 | from_8406 | to_8406 | art_short | ec | K |
| 203 | 246 | 0.628578222 | cat4 | from_8407 | to_8407 | art_short | ec | K |
| 204 | 246 | 0.697739334 | cat4 | from_8408 | to_8408 | art_short | ec | K |
| 223 | 246 | 0.64556723 | cat4 | from_8409 | to_8409 | art_untreat | ec | F |
| 225 | 246 | 0.654558564 | cat4 | from_8410 | to_8410 | art_untreat | ec | F |
| 12 | 247 | 0.85245446 | cat4 | from_8411 | to_8411 | art_long | ec | N |
| 13 | 247 | 0.670488795 | cat4 | from_8412 | to_8412 | art_long | ec | N |
| 19 | 247 | 0.841663477 | cat4 | from_8413 | to_8413 | art_long | ec | N |
| 40 | 247 | 0.672226625 | cat4 | from_8414 | to_8414 | art_long | ec | N |
| 42 | 247 | -0.616852989 | cat4 | from_8415 | to_8415 | art_long | ec | N |
| 79 | 247 | 0.862235816 | cat4 | from_8416 | to_8416 | art_long | ec | N |
| 85 | 247 | -0.647445657 | cat4 | from_8417 | to_8417 | art_long | ec | N |
| 92 | 247 | -0.641465348 | cat4 | from_8418 | to_8418 | art_long | ec | N |
| 93 | 247 | 0.697177866 | cat4 | from_8419 | to_8419 | art_long | ec | N |
| 97 | 247 | -0.610409822 | cat4 | from_8420 | to_8420 | art_long | ec | N |
| 103 | 247 | -0.651970064 | cat4 | from_8421 | to_8421 | art_long | ec | N |
| 114 | 247 | 0.644011152 | cat4 | from_8422 | to_8422 | art_long | ec | N |

| | | | | | | | | |
|---|---|---|---|---|---|---|---|---|
| 115 | 247 | -0.685488928 | cat4 | from_8423 | to_8423 | art_long | ec | N |
| 117 | 247 | -0.61192418 | cat4 | from_8424 | to_8424 | art_long | ec | N |
| 133 | 247 | -0.713878459 | cat4 | from_8425 | to_8425 | art_long | ec | N |
| 145 | 247 | 0.612145051 | cat4 | from_8426 | to_8426 | art_long | ec | N |
| 147 | 247 | -0.628914447 | cat4 | from_8427 | to_8427 | art_long | ec | N |
| 156 | 247 | 0.58285466 | cat4 | from_8428 | to_8428 | art_short | ec | K |
| 163 | 247 | 0.630627644 | cat4 | from_8429 | to_8429 | art_short | ec | K |
| 165 | 247 | 0.577776439 | cat4 | from_8430 | to_8430 | art_short | ec | K |
| 167 | 247 | 0.884995606 | cat4 | from_8431 | to_8431 | art_short | ec | K |
| 177 | 247 | -0.585747833 | cat4 | from_8432 | to_8432 | art_short | ec | K |
| 186 | 247 | 0.667930806 | cat4 | from_8433 | to_8433 | art_short | ec | K |
| 216 | 247 | 0.596514475 | cat4 | from_8434 | to_8434 | art_short | ec | K |
| 229 | 247 | 0.704661926 | cat4 | from_8435 | to_8435 | ec | ec | P |
| 6 | 248 | 0.585802923 | cat4 | from_8436 | to_8436 | art_long | ec | N |
| 8 | 248 | 0.610874848 | cat4 | from_8437 | to_8437 | art_long | ec | N |
| 43 | 248 | 0.596657198 | cat4 | from_8438 | to_8438 | art_long | ec | N |
| 50 | 248 | 0.649373183 | cat4 | from_8439 | to_8439 | art_long | ec | N |
| 61 | 248 | 0.83177345 | cat4 | from_8440 | to_8440 | art_long | ec | N |
| 93 | 248 | 0.702100851 | cat4 | from_8441 | to_8441 | art_long | ec | N |
| 99 | 248 | 0.620501323 | cat4 | from_8442 | to_8442 | art_long | ec | N |
| 105 | 248 | 0.706753834 | cat4 | from_8443 | to_8443 | art_long | ec | N |
| 115 | 248 | -0.585893842 | cat4 | from_8444 | to_8444 | art_long | ec | N |
| 135 | 248 | 0.582203235 | cat4 | from_8445 | to_8445 | art_long | ec | N |
| 145 | 248 | 0.662233924 | cat4 | from_8446 | to_8446 | art_long | ec | N |
| 148 | 248 | 0.703701086 | cat4 | from_8447 | to_8447 | art_long | ec | N |
| 151 | 248 | 0.663926605 | cat4 | from_8448 | to_8448 | art_long | ec | N |
| 152 | 248 | 0.66730554 | cat4 | from_8449 | to_8449 | art_long | ec | N |
| 153 | 248 | 0.586793791 | cat4 | from_8450 | to_8450 | art_long | ec | N |
| 157 | 248 | 0.680690272 | cat4 | from_8451 | to_8451 | art_short | ec | K |
| 176 | 248 | 0.635233623 | cat4 | from_8452 | to_8452 | art_short | ec | K |
| 183 | 248 | 0.767813382 | cat4 | from_8453 | to_8453 | art_short | ec | K |
