## Appendix Table S8 for "Distinguishable topological properties of functional genome networks in HIV-1 reservoirs"

**Supplementary Table 8. List of primers used in this study**

| Primer ID | Sequence | FW or RV | Description |
|---|---|---|---|
| HCP#63 | 5'-TGACGGGGTCACCCACACTGTGCCCATCTA-3' | FW | Actin beta [ACTB] |
| HCP#64 | 5'-CTAGAATTTGCGGTGGACGATGGAGGG-3' | RV | Actin beta [ACTB] |
| HCP#228 | 5'-ACACAGACCTTCAGATCACTCC-3' | FW | CHUK |
| HCP#229 | 5'-GATATTACTGAGGGCCACTTCC-3' | RV | CHUK |
| HCP#230 | 5'-GAACGTCGAAAAGAAAAGTCTCG-3' | FW | HIF1A |
| HCP#231 | 5'-CCTTATCAAGATGCGAACTCACA-3' | RV | HIF1A |
| HCP#232 | 5'-ACGCTCTGGGAAATCTGCTA-3' | FW | JAK1 |
| HCP#233 | 5'-ATGATGGCTCGGAAGAAAGG-3' | RV | JAK1 |
| HCP#234 | 5'-ACGTGACATCCTCGATAAAACTG-3' | FW | OAS2 |
| HCP#235 | 5'-GAACCCATCAAGGGACTTCTG-3' | RV | OAS2 |
